# Neural microstates in real-world behaviour captured on the smartphone

**DOI:** 10.1101/2024.07.22.604605

**Authors:** Ruchella Kock, Arko Ghosh

**Author notes:** Corresponding address Ruchella Kock & Arko Ghosh Leiden University Wassenaarseweg 52 Leiden, 2333 AK [Arko Ghosh] [Ruchella Kock].

## Abstract

Microstates are periods of quasi-stability in large-scale neural networks, and they are ubiquitous spanning sleep, rest and behaviour. One possibility is that the temporal and spatial features of these states vary from person to person contributing to inter-individual behavioural differences. Another possibility is that the microstates support momentary behavioural needs and as a result their properties may vary with behavioural fluctuations in the same person. Here, we leverage the history of smartphone touchscreen interaction dynamics and combined smartphone and EEG recordings (N = 61), to address if and how microstates contribute to real-world behaviour. We find that microstates measured when at rest correlated with the history (∼30 days) of smartphone interaction dynamics captured in the real world, indicating their role in the behaviour. During smartphone behaviour, the configuration of microstates varied over time. However, this variance was largely unrelated to the rich behavioural dynamics, as in similar temporal and spatial features were observed during rapid and slow smartphone interactions. We show that in the real world microstates do not respond to the fluctuations of the ongoing behaviour. Still, as microstates measured at rest are correlated to the inter-individual behavioural differences captured on the smartphone, we propose that microstates exert a top-down influence to broadly orchestrate real-world behaviour.

## Introduction

Human brain activity is spatially rich and highly dynamic. In large-scale neural networks a small number of functional states dominate this activity (Smitha et al., 2017). In EEG measurements, neural states can be captured as microstates, i.e., global patterns of scalp potential topographies lasting ∼60- 120 milliseconds (Lehmann et al., 1987). Four microstates — denoted A, B, C, D — have been repeatedly observed across the population (Khanna et al., 2014; Michel & Koenig, 2018). These four microstates have been linked to the following functional networks: A with auditory, B with visual, C with default mode, self-reflections, or saliency network, and D with the attention network (Britz et al., 2010; Tarailis et al., 2023). The temporal and spatial features of microstates are well captured using the parameters of duration, occurrence, *coverage (fraction of time that the microstate is active)*, and transitions probabilities (Lehmann et al., 1987). Variations of microstates parameters have been linked to brain functions, ranging from sleep to wakefulness (Bréchet et al., 2020; Brodbeck et al., 2012). However, as these insights are mostly based on artificial laboratory experiments, the role of microstates in spontaneous real-world behaviour remains unclear.

It is possible that inter-individual differences in microstate parameters contribute to diverse real-world behaviours. The relationship between inter-individual differences in microstates and behaviour is unclear. Still, microstates measured at rest are correlated to some inter-individual factors. Microstates A, B, C and D correlated to chronological age (Koenig et al 2002). The occurrence of microstate D was higher in males compared to females and the duration of microstate C was lower in males (Tomescu et al., 2018). Moreover, microstates are different in people who are healthy compared to people with neurological disorders (Lehmann et al., 2005; Murphy et al., 2020; Nishida et al., 2013). However, these interesting inter-individual differences do not easily inform how microstates contribute to real-world behaviours.

In the real world there is a rich range of behaviours, and it is also possible that microstates alter from one behaviour to the next to support the diverse computational needs. This idea is supported by laboratory-based observations, where within individuals, the microstate parameters vary depending on the behavioural context. For instance, the parameters of microstate D (attention network) and C (default mode) are altered during the performance of cognitive tasks compared to when at rest (Bréchet et al., 2019; Kim et al., 2021; Seitzman et al., 2017). Across verbal tasks compared to visual tasks, the parameters of microstates A and B vary, although there are inconsistent reports in terms of their direction (Antonova et al., 2022; Milz et al., 2016). Furthermore, the level of arousal during different behavioural contexts is associated with changes in the parameters of microstates A and C (Krylova et al., 2021; Li et al., 2023). These findings suggest that intra-individual variations in microstate parameters exhibit properties that alter to the behavioural context, albeit the evidence does not extend to the real world.

Smartphones offer a fresh approach to quantify the inter and intra-individual differences in real-world behaviour enabling us to address the putative microstates-behavioural links (Ceolini et al., 2022; Ceolini & Ghosh, 2023; Duckrow et al., 2021). A range of behaviours are expressed on the smartphone, from texting a friend to browsing the internet. Quantifying these rich behaviours and linking them to microstates is not straightforward. One intuitive possibility is to compare microstates parameters across different smartphone contexts such as between apps. But this approach is hindered by the large number of different apps in use (> 1000) and within each app there may be a range of behaviours: for instance, a gaming app may also support messaging. An emerging alternative leverages the time series of smartphone touchscreen interactions to separate the behaviours according to their next interval patterns. To capture these behavioral dynamics, an interval between two smartphone interactions (*k*), is considered in conjunction to the next interval (*k* + 1). These intervals can be pooled together in two-dimensional bins using a joint-interval distribution (JID). Using the JID framework, fast consecutive intervals can be separately considered from the slow consecutive intervals. Notably, using these behavioural dynamics, fast consecutive intervals has been correlated to sensorimotor cognitive tasks whereas the slower intervals were correlated to executive functions (Ceolini et al., 2022).

Addressing the intra-individual links between microstates and the JID behavioural representation raises a new challenge. Essentially, the different combinations of microstate parameters may reflect the ongoing behavioural dynamics. For instance, during slower-intervals, duration and coverage of a microstate may increase while occurrence may decrease. This creates the possibility of complex inter-relations between the many microstate parameters (say 4 microstates x 3 parameters) and the high dimensional behaviour (say 2500 two-dimensional bins of the JID). Non-negative matrix factorization (NNMF) is a powerful dimensionality-reduction technique that reveals interpretable low-rank approximations of a matrix (Lee et al., 1998). NNMF can be used to untangle interpretable configurations of microstates parameters that are expressed during different smartphone behaviours. As in to yield a few prototypical microstate configurations and their behavioural distribution.

In this study we quantified inter and intra-individual differences in microstates and behaviours. Mass univariate regression models captured the inter-individual differences in JIDs collected ∼30 days prior to the EEG measurement and corresponding microstates quantified at rest. To quantify intra-individual differences, we used NNMF, at the individual-level, to extract the prototypical microstate configurations across the two-dimensional bins.

## Results

Microstates showed both inter and intra-individual differences, and here we address the behavioural correlates of these differences. These microstates parameters were linked to diverse smartphone behaviours captured using a joint-interval distribution (JID). We reveal inter-individual differences between rest microstate parameters and historical JIDs. Within individuals, we leveraged NNMF and identified broad configurations of microstates parameters during smartphone behaviours. However, this configuration only marginally changed across different behavioural dynamics, be it fast or slow. Suggesting that diverse smartphone behaviours are not driven by on-demand re-configurations of microstates parameters.

### Microstates at rest

Microstates were identified during eyes closed resting-state, creating four global microstates. Microstate C exhibits anterior-to-posterior activity and microstate D shows substantial front-central activity. Microstate A and B were not identified. We named the other two microstates E and F (Tarailis et al., 2023). Microstate E displays substantial centro-parietal activity whereas microstate F displayed left-right activity. For each individual, we calculated the microstate parameters and found that they only marginally varied across the four microstates. Across most individuals, each microstate typically covered ∼25% of the whole measurement (coverage), occurred approximately 3 times per second (occurrence) and lasted ∼75 milliseconds (duration) (Figure 1). Together these four states explained 52% of the global variance (GEV) (Supplementary Figure 1). The GEV is a measure showing the percentage of variance of the original EEG data that is explained by the microstate templates.

**Figure 1.**
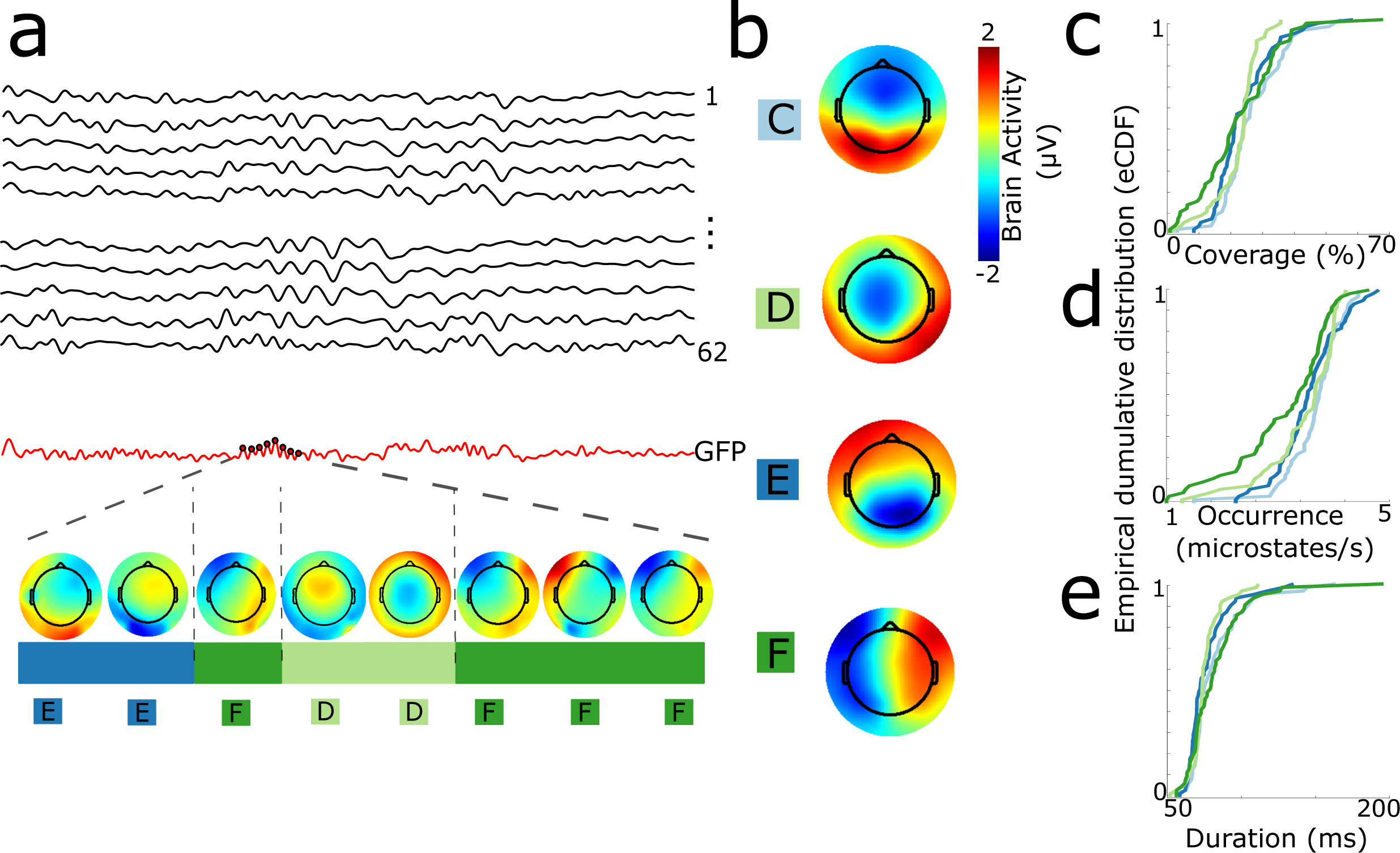
Microstates measured at rest. (a) Sketch on how to identify EEG microstates. **(b)** Four microstates were identified during rest denoted C, D, E and F. **(c to e)** Empirical cumulative distribution function (ECDF) of the rest microstate parameters across the population. The line colours represent the corresponding microstates. Overall, the distributions of microstate parameters marginally varied across the four states. **(c)** ECDF for the total percentage of time each microstate covered the rest recording. **(d)** ECDF for the median occurrence of the microstates **(e)** ECDF for the median duration of the microstates. Minimum duration was set to 50 ms.

### Inter-individual differences in microstates at rest

First, we addressed the stability of the microstate parameters captured at rest, given that inter-individual differences in unstable microstate parameters may simply arise by chance. A subset of individuals participated in more than one EEG session (N=25 out of 61) conducted approximately one week apart. We correlated the microstate parameters across these two sessions and found high correlations for most of the individuals (Median R^2^ = 0.7177 for coverage, Median R^2^ = 0.8478 for occurrence, Median R2 = 0.7352 for duration)(Supplementary Figure 2). These findings suggest individual’s microstate parameters remain largely stable over a period of a week.

Next, we correlated the inter-individual differences in the microstate parameters captured at rest and the historical behaviour captured in the real world (Figure 2). Smartphone behaviour was captured by using the JID which is composed of multiple two-dimensional bins each reflecting different behavioural intervals. The behavioral probability density in each bin may vary across individuals.

**Figure 2.**
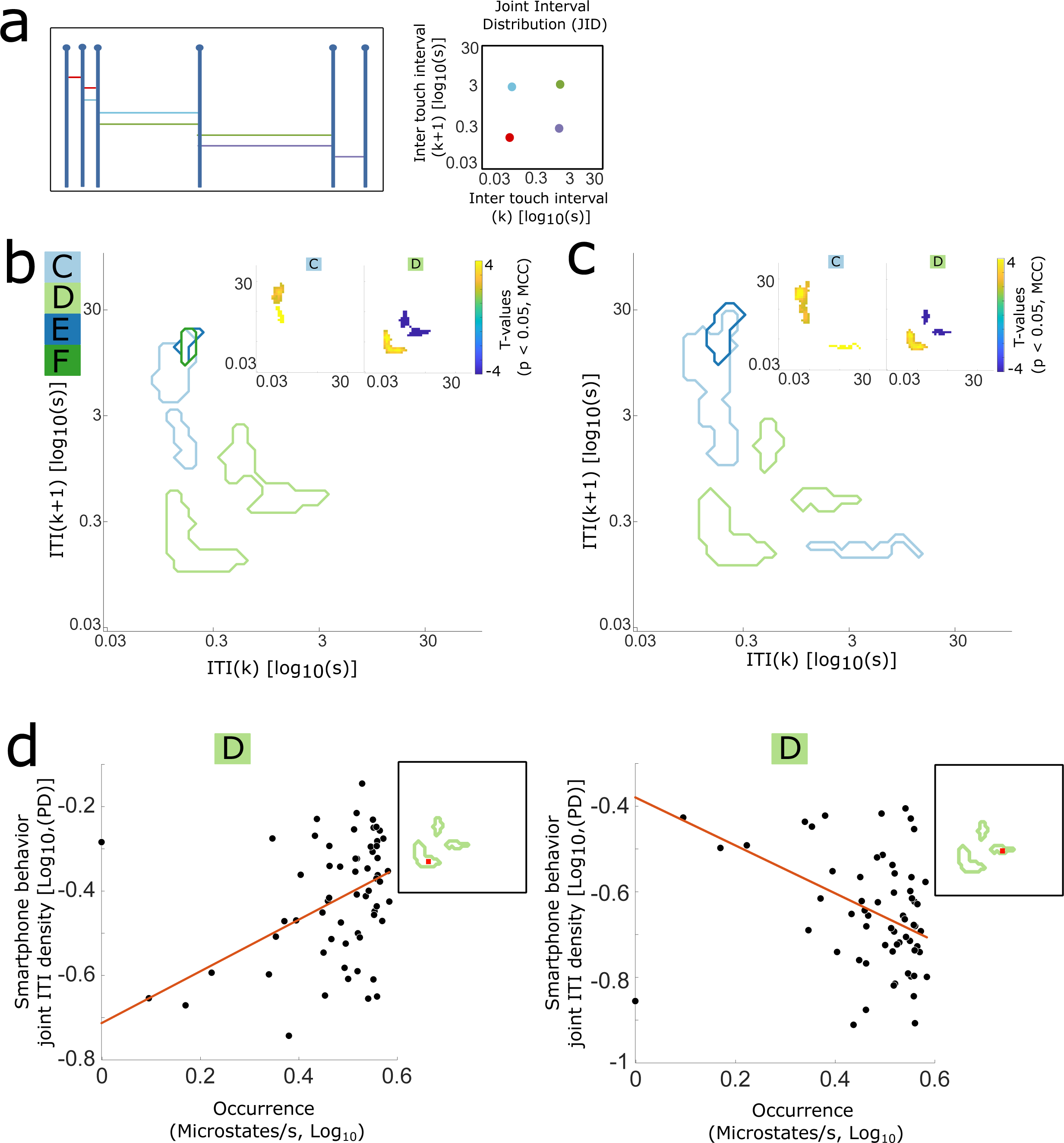
Inter-individual differences in historical smartphone behaviors and microstates captured at rest. **(a)** Sketch showing how smartphone behavior can be captured using a joint-interval distribution (JID). Multiple consecutive intervals (k and k+1) between smartphone interactions can be pooled into two-dimensional bins. Adapted from Ceolini & Ghosh (2023) with permission. **(b)** Univariate linear regressions reveal a relationship between JIDs and microstate parameters across the population. Contours of the JID show significant relationships between the behavioral bins and the four microstate coverage parameters. T-values show microstate C coverage increases during fast followed by slow intervals. Microstate D coverage increases during fast intervals and decreases during slow intervals. The *t*-values were corrected for multiple comparisons using spatiotemporal clustering (1000 bootstraps, α = 0.05). **(c)** Same as b for the occurrence parameter. **(d)** Scatterplot for two example two-dimensional bins (red dot) show microstate D occurrence increases during fast and decreases during slow behaviors. The red line shows the iterative least squares prediction. The behaviors and microstate parameters are shown in log10 space.

Interestingly, these inter-individual differences in behaviour were weakly correlated with different microstate parameters captured at rest. The coverage and occurrence parameter of microstate D increased during fast consecutive intervals and decreased during the slower intervals. The coverage and occurrence of microstate C increased during fast followed by slow intervals. The coverage of the other microstates (E and F) was weakly correlated to fast followed by slow intervals. The duration of microstates E and F were weakly correlated to slow intervals and fast followed by slow intervals (Supplementary Figure 3). Taken together, an individual microstate parameter is stable over time and there were statistically significant inter-individual differences. For results on inter-individual differences between smartphone behavior and global microstates that were not constrained to four states see Supplementary Notes Figure 2.

### Microstates during smartphone behaviour

To address whether microstates alter depending on an individual’s behavioural demands, we captured microstates during smartphone behaviour (Figure 3). At the individual-level approximately ten microstates were identified. These microstates were clustered across the population resulting in twelve population-level microstates. Out of these twelve, seven microstates displayed large amplitudes across the scalp, the rest were likely artifacts and not considered further (Supplementary Notes Figure 2). The four microstates commonly reported in the literature — microstate A, B, C and D — were found (Michel & Koenig, 2018). Additionally, three other microstates which we labelled as microstate E, F and G were identified. Microstates E has large amplitudes over the frontal electrodes. Microstates F and G both displayed front-central activity, with the activation stronger in the left hemisphere for microstate F and stronger in the right hemisphere for microstate G.

**Figure 3.**
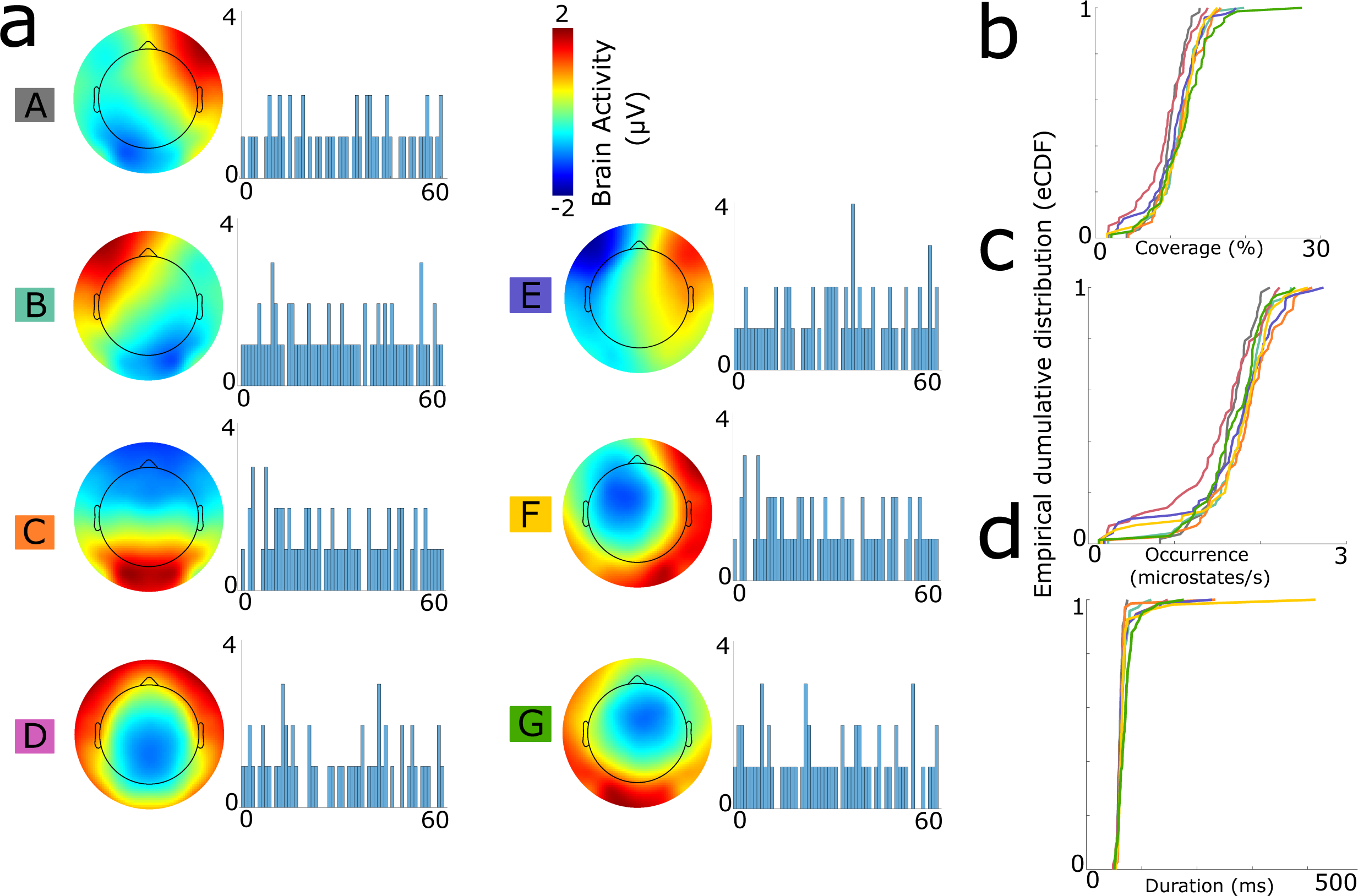
Microstates identified during smartphone behaviour. **(a)** Seven consistent microstates were present across the population. Microstate A to D match commonly reported microstates captured at rest. Microstates E to G have been reported more sparsely. The microstates displayed were clustered from individual-level microstates, and the individuals in each cluster are shown with a bar plot. Some individuals had more than one microstate that resembled the population-level microstate. **(b to d)** Empirical cumulative distribution function (ECDF) of the microstate parameters across the population show large similarities across the seven microstates. The line colours represent the corresponding microstates. The ECDF of **(b)** displays the coverage, **(c)** median occurrence and **(d)** median duration.

The population distributions of the microstate parameters (i.e. coverage, occurrence, duration) were relatively similar across the seven microstates (Figure 3). In most individuals, the seven microstates had a coverage ranging between 0 to 30% (Population Median = 11%). The occurrence of all the microstates ranged between 0 to 3 (Population Median = 1.7 states per second). The median duration ranged up to 250 milliseconds (Population Median = 62 milliseconds). Except for microstate F where the duration ranged up to 450 milliseconds in a subset of the individuals. The GEV for each microstate was ∼10%, except for microstate B and F where a subset of individuals had GEV values that ranged up to 60%. Together these seven microstate parameters had a GEV of ∼20% across the population (Supplementary Figure 1).

### Intra-individual differences in microstates during smartphone behaviour

Non-negative matrix factorization (NNMF) was utilized to extract the pattern of microstate parameters (coverage, occurrence, duration) underlying the diverse smartphone behaviours (Figure 4). NNMF identified typical configurations of microstate parameters (meta-parameters) and helped isolate where they are expressed across the behaviours (meta-behaviour). Except for two individuals, the NNMF resulted in rank 1 decompositions. There were substantial inter-individual differences in the meta-parameters. Despite these differences, the meta-parameters were expressed in two common ways across the behaviours. The first pattern was displayed in most individuals and showed that the same meta-parameters were present across the whole behavioural space. Indicating that although there is a specific configuration of microstate parameters during behaviour, it does not vary across faster or slower behaviours. For a subset of individuals (N = 22) the meta-parameters were present in only a few behaviours, this may be linked to the microstates driven by artifacts (see Supplementary Notes Figure 3). For an alternative approached based on block-bootstrapping which confirms the null findings see Supplementary Notes Figure 4 and 5.

**Figure 4.**
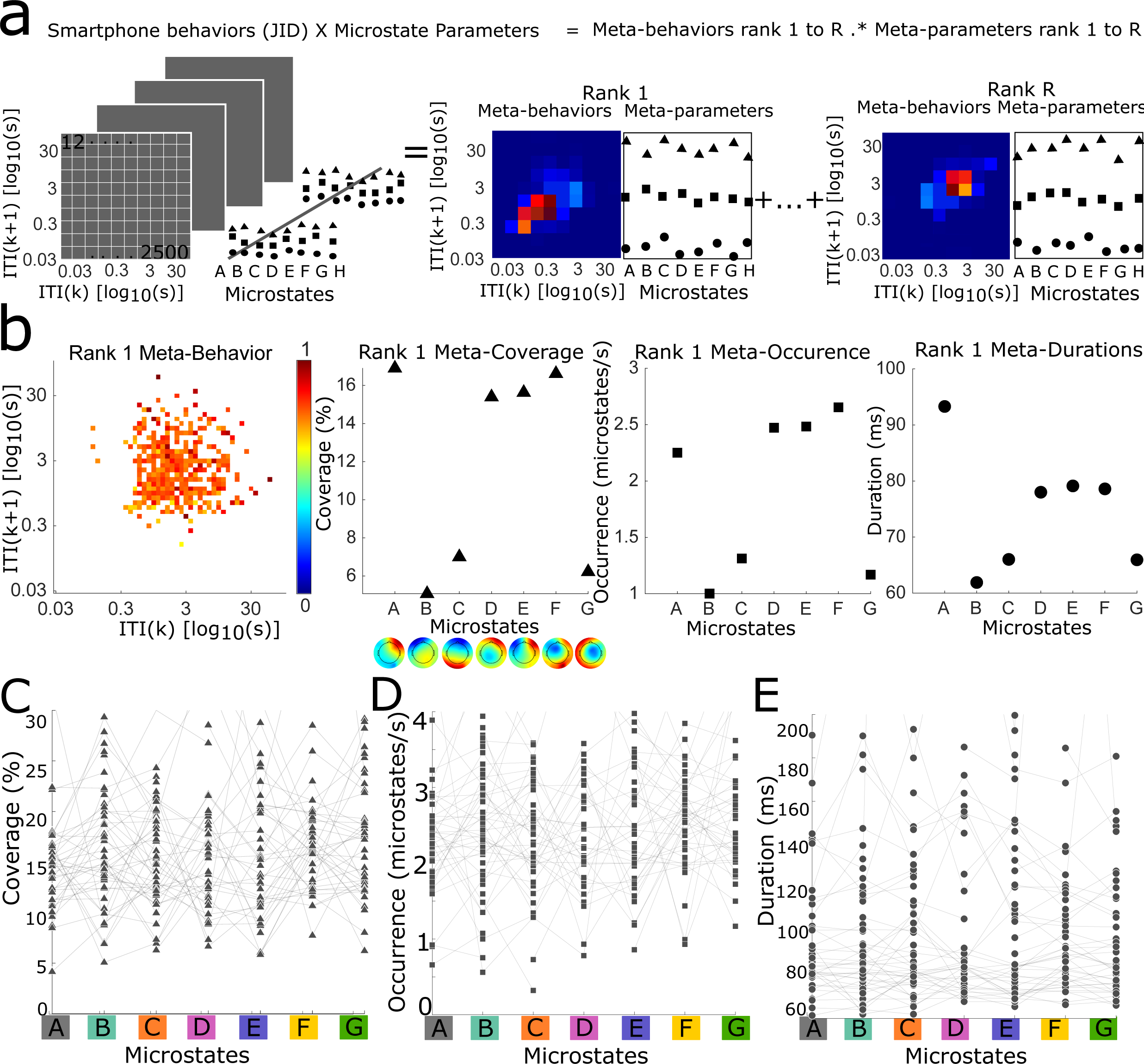
Configuration of microstates parameters during smartphone behaviour. **(a)** Sketch showing how non-negative matrix factorization (NNMF) decomposes the microstate parameters – coverage (triangle), occurrence (square), and duration (circle) – across each two-dimensional bin of smartphone behaviours. This results into rank-one or more approximations of the configuration of microstate parameters (meta-parameters) expressed across different smartphone behaviours (meta-behaviours). The meta behaviours are shown in log10 scale. **(b)** NNMF decompositions of one example individual. The same configuration of microstate parameters was present across all the behaviours, be it multiple fast or slow smartphone interactions. The microstates were identified at the individual-level and clustered across the population. Topography plots display the individual-level microstate maps. The microstate labels correspond to the population-level clusters displayed in Figure 3. When multiple microstates were clustered to the same population-level microstates these are denoted with a number such as C1, C2. **(c to e)** The meta-parameters across the population display substantial variance between individuals. The median parameter was calculated for individuals with multiple microstates clustered in the same population-level microstates. The line connects the values of the individual. **(e)** Displays the meta-coverage across the population, coverage values above 30% are not shown for readability. The values ranged up to ∼70% **(f)** Displays the meta-occurrence any occurrence above 4 are not displayed, the occurrence ranged up to ∼9 times per second. **(g)** Displays the meta-duration, durations above 200 are not displayed, the duration ranged up to ∼1 second.

Another microstate parameter is the probability of transitioning from one state to the next. We repeated the NNMF to identify typical transition parameters (meta-transitions) across the smartphone behaviours (meta-behaviours). The NNMF resulted in 88% rank 1, 8% rank 2 and 3% more than rank 3 decompositions (Supplementary Figure 4, Supplementary Notes Figure 6). Once again, we discovered substantial inter-individual differences in the transition probabilities. Still, for the majority of individuals, the meta-transition pattern was displayed across the whole behavioural space, suggesting a common transition pattern regardless of fast or slow behavioural dynamics.

To comprehensively address the relationship between multiple microstate parameters and smartphone behavior, we calculated the duration of a microstate before and after each transition (Supplementary Figure 4). Microstate transition probabilities are closely related to the occurrence parameter. Microstates that occur more frequently have a higher probability there will be a transition to them. While the transition probability is linked to the occurrence, duration is not reflected in the transition probability. We included the duration by using NNMF to extract meta-transition durations across the behaviors (meta-behaviors). The NNMF resulted in 60 % rank 1 decompositions, 30 % rank 2 decompositions, and 10 % larger than 2 (Supplementary Notes Figure 6). Similar to the transition probability, the NNMF with rank 2 decompositions separated transitions to and from microstates that appear dominated by artifacts another decomposition separated the remaining microstates. The durations before or after transitioning only marginally varied between the microstates. This pattern was consistent across the participants as well as across the whole behavioral space.

Overall, the NNMF identified configurations of microstate parameters that varied across individuals but not across different types of behaviours.

## Discussion

In this study, we addressed the role of microstates in real-world behaviour. Inter-individual differences in a subset of microstate parameters captured at rest, were weakly correlated with historical smartphone behaviours. Specifically, microstate D increased during fast intervals and decreased during slow intervals while microstate C decreased during fast followed by slow intervals. Within individuals, there may be a characteristic configuration of microstate parameters, but these did not alter based on smartphone behavioural dynamics. In other words, microstates parameters were not tuned to facilitate diverse behavioural demands, be it fast or slow intervals of smartphone behaviours. Our findings point towards microstates exerting a top-down control over behaviour.

The correlations between inter-individual differences in microstates captured at rest and real- world behaviour captured on the smartphone suggest that microstates contribute to behaviour in a trait-like manner. First, we found that microstate parameters were largely stable by using measurement sessions separated by a week. This is in line with prior studies which show good test-retest reliability of microstate parameters (Khanna et al., 2014; Kleinert et al., 2024). Second, the microstates captured during smartphone behaviour were largely invariant to the ongoing behavioural fluctuations. Finally, the inter-individual variations in behaviour explained by microstate are in line with previous work linking microstates to personality traits or expertise (Cui et al., 2021; Schiller et al., 2020; Tomescu et al., 2022). For instance, Cui et al. (2021) compared gaming experts to non-experts and found that gamers of games rich in visual stimuli had higher occurrence and coverage for microstate B that has been linked to visual functions. These and our findings are in line with emerging evidence from FMRI where resting-state neural networks are proposed to be stable and trait-like, reproducible across sessions within individuals, varying across individuals, and correlated to behaviours (Seitzman et al., 2019; Zanesco et al., 2020).

One possible explanation for the inter-individual patterns is that there may be common factors that dictate both behaviour and microstates. However, we can rule out two important candidate factors – age and gender. Considering age, the smartphone behavioural dynamics are known to alter through the adult lifespan, with faster behaviours becoming less common in the elderly (Ceolini et al., 2022).

Furthermore, microstate parameters also differ across the lifespan (Koenig et al., 2002). Still, given the narrow distribution of age in our study (16 to 27 years old), it is unlikely to drive the reported patterns. Considering gender, the smartphone behavioural dynamics shows weak differences between males and females. In any case, in our study we found no correlation between gender and microstate parameters.

The specificity with which certain microstates were correlated to the inter-individual differences in smartphone behaviour offer some clues on the underlying processes. Of all microstates, D was most correlated to the behaviour and the correlations spanned slow and fast consecutive intervals.

Microstate D is commonly linked to executive functions such as attention (Bréchet et al., 2019; Kim et al., 2021; Seitzman et al., 2017; Tarailis et al., 2023). In a previous study we have shown that inter-individual differences across executive functioning tasks were correlated with both fast and slow smartphone behavioural dynamics (Ceolini et al., 2022). We speculate that executive functions are shaped by microstate D resulting in the behavioural differences. However, emerging evidence does not strongly support the link between microstate D captured at rest and subsequent executive functioning in laboratory tasks (Chenot et al., 2024). Still, these findings were not linked to behaviors in the real-world. Microstates C, E and F showed correlations to behaviours where fast intervals were followed by slow intervals; although the latter two were only trends below alpha. All these three microstates have been linked to introspective processes (Tarailis et al., 2023). We speculate that microstates also shape introspective processes supporting the transition from fast to slow – perhaps thoughtful – smartphone behavioural dynamics.

Within individuals, microstates are known to vary between different behavioural contexts – such as at rest vs. when engaged in a task (Zanesco et al., 2020). Consistent with these findings we found that individuals may have a unique configuration of microstate parameters during smartphone behaviour.

However, the microstate parameters did not systematically fluctuate in corresponding to the diverse smartphone behaviours. Even when considering the microstate parameters separately we found no statistically significant relationship to behavioural dynamics within individuals. There may be characteristic configuration of microstates during behaviour, but they may not alter to the ongoing demands imposed by behaviour.

Considering all our findings, we speculate that microstates influence real-world behaviour through a top-down process. We found that microstates collected at rest behave as a trait-like property correlated to historical behaviours. During ongoing behaviour there is a characteristic configuration of microstates. Both these results indicate that microstates have a computational template that is related to behaviour. Still, during ongoing behaviours, the microstates do not dynamically alter, suggesting that bottom-up behavioural demands do not influence the computational template.

This study has some notable limitations. First, a general limitation of microstate analysis is that global states are identified and then fitted onto individual individuals, and thus overlooking the individual-level patterns. Here, we too identified global microstates when capturing inter-individual differences to allow for more robust comparisons across individuals. This analytical convenience comes at the cost of potentially missing the individualized patterns. Still, in the intra-individual analysis we did identify microstates at the individual-level. Second, while NNMF identified interpretable linear configurations of microstate parameters across the behaviours, we cannot rule out the possibility of non-linear relationships. NNMF decomposes a matrix, assuming that there is an underlying pattern in the matrix and that it can be constructed through linear combinations of the vectors (Wang & Zhang, 2013). Our results primarily indicate that there is a rank-1 linear decomposition to the describe the relationship between microstates and behavior. While a rank-1 decomposition might point to a single optimal decomposition for the matrix, it is important to recognize that NNMF inherently cannot achieve less than rank-1. Therefore, it remains possible that there is no underlying pattern that can be explained through linear relationship. In other words, microstates may still alter on-demand through highly complex non-linear patterns. Finally, it remains challenging to identify universal neuro-behavioural patterns in real-world behaviour; although all individuals were using smartphones, their goals and motivations underlying their behaviour may have been different.

Notwithstanding these limitations, our study provides a fresh new approach to study microstates during real-world behaviour. Real-world behaviour has many complexities, but these must be addressed to better understand the functional relevance of neural states.

## Methods

### Participant

Healthy individuals, ranging from 16 to 27 years old (Median = 22 years old), were recruited as part of a larger study on smartphone behaviour (Supplementary Notes Figure 7). Individuals were all right-handed as indicated through self-report. The recruitment took place on Leiden University campus through flyer advertisements. Sixty-one individuals (26 females) were included in the analysis after the pre-processing. The pre-processing ensured high-precision synchronization between the smartphone and EEG datasets based on the methods of Kock et al. (2023). The study was approved by the Leiden University Ethics committee. All individuals signed a written informed consent form and received debriefing after the study. The data was anonymized prior to the data analysis. As part of the larger study, a subset of the individuals (N=25), participated in two separate sessions approximately one week apart. The procedure across the two sessions were identical. The procedure across the two sessions were identical.

### EEG data collection and processing

EEG data was collected during while individuals were resting as well as using their smartphone.

The EEG measurement started with ∼10 minutes of rest. Through a randomized design individuals starting with either eyes-open or eyes-closed, then alternated halfway between conditions. During the eyes-open condition, individuals fixated on a screen in front of them. The conditions were confirmed by identifying the number of blinks with Independent Component Analysis. The eyes-closed condition contained a small number of blinks as opposed to the eyes-open condition. Next, EEG was recorded while individuals were using their smartphone (see next section for more details).

The individuals were seated in a faraday cage during the measurement. The EEG cap consisted of 64 equidistant electrodes and the 64-channel DC amplifier BrainAmp (Brain Products GmbH, Gilching) was used for the collection. The data was collected at a resolution of 1000 Hz. The EEG data was processed by replacing channels with impedance larger than 10 kΩ with interpolated data, performing Independent Component Analysis (Infomax) for blink artifacts removal, and subsequently removing the two electrodes placed under the eye from further analysis. The data was bandpass filtered between 1 to 40 Hz, selected based on common filtering ranges from previous studies (Michel & Koenig, 2018). The processing was performed offline using EEGLAB and Matlab 2019b (MathWorks, Natick)(Delorme & Makeig, 2004).

### Smartphone data collection and processing

Smartphone behaviour was recorded with the TapCounter app (QuantActions AG, Zurich), operating passively in the background. TapCounter collects the timestamps of smartphone interactions and the name of the corresponding app in use (such as Instagram). The data is collected at millisecond precision, with a standard deviation of 15 milliseconds (Balerna & Ghosh, 2018). Individuals were instructed to install the app on their Android phones for at least two weeks prior to coming to the laboratory. Based on their usage history their top two most frequently used social and top two non- social app were selected. They alternated every 10 minutes between the apps and were encouraged to take short breaks in between. Individuals could freely use the selected apps, without any explicit instructions on their interactions. They were using their right-thumb during the interactions, which we validated the through online video streaming.

We captured the smartphone behavioural dynamics with a joint-interval distribution (JID) following the methods introduced by Duckrow et al. (2021). This involved accumulating the time distance — inter-touch intervals (ITI) — between multiple consecutive smartphone interactions into two-dimensional (50 x 50) bins. Using a kernel density estimate over the log transformed inter-touch intervals the distribution of behavioural dynamics was captured.

### Microstates analysis

The microstates were identified during rest and smartphone behaviour separately, using the Microstate EEGLAB toolbox (Poulsen et al., 2018). At rest we followed the conventional method identifying global microstates and restricted the number of microstates to four (Michel & Koenig, 2018). A global selection facilitated inter-individual analysis. Four microstates are well documented at rest, allowing a comparison with the literature. For an additional analysis without restricting the number of states see Supplementary Notes Figure 1 During behaviour the number of states is not well documented therefore a preselection of the number of states was not applied. To capture potential intra-individual differences in the microstates during behaviour, we identified microstates at the individual-level.

First, the Global Field Power (GFP) was calculated and GFP peaks with a minimum distance of 10 milliseconds to each other were identified. Next, 1000 GFP peaks were randomly selected to perform the analysis. We clustered the GFP peaks with the modified k-means algorithm (Pascual-Marqui et al., 1995). The optimal number of clusters (between 2 to 10) was selected based on the global explained variance (GEV). This resulted in between 2 to 10 microstates. Commonly four to five microstates are selected to align with existing literature. However, more than four states may exist, and we allowed a wider range to capture all the potential microstates (Michel & Koenig, 2018). The final step included backfitting the microstates to the EEG data. During the back-fitting, segments smaller than 30 milliseconds were smoothed by replacing the microstate labels in these segments to the most likely class. This allows for better temporal continuity in the microstates (Pascual-Marqui et al., 1995; Poulsen et al., 2018). The minimum duration for a microstate was set to 50 milliseconds. After backfitting the microstates parameters coverage, occurrence, duration, and transition probabilities were calculated.

Next, we identified population-level microstates by repeating the modified k-means algorithm with the microstates of all the individuals. The optimal number of microstates (between 2 to 15) was selected based on the optimal GEV. The clustering was repeated 1000 times to receive the most stable result, accounting for the non-convex nature of the algorithm. The repetition with the best sum of squared distance was selected (Lisboa et al., 2013). For some individuals, multiple individual-level microstates were clustered as the same population-level microstate. This is possible as the population-level microstates are derivates of the individual-level ones. Therefore, a microstate that may not fit any population wide patterns would still be clustered into one population-level microstate. We reordered and hand labelled the population-level microstates in accordance with naming conventions in the literature (Tarailis et al., 2023).

### Inter-individual differences in microstates and smartphone behaviours

We identified microstates for the inter-individual analysis on the EEG data collected when individuals were resting with their eyes closed. As eyes open and eyes closed microstate parameters are known to vary, we selected eyes closed which is more broadly used (Seitzman et al., 2017). To understand inter-individual differences in microstates parameters we first calculated the stability of the rest microstate parameters (coverage, occurrence, and duration) across the two sessions with a correlation coefficient (Pearson R). Next, we used iterative least squares regression to reveal the inter-individual relationships between microstate parameters and smartphone behaviours captured through the JID. We trained a regression model for each microstate parameter separately and selected only the parameters of the first session, given the high correlation between the parameters across the two sessions. As individuals may differ in microstate parameters and these may be correlated to different behavioural dynamics, the regression model predicted each two-dimensional bin of the JID. The JID was calculated over ∼30 days of smartphone interactions prior to the measurement (19 individuals had less than 30 days of data, median = 30 days, mean = 27 days). Both the microstate parameters and JIDs violated the assumption of normal distribution and were log10 transformed before the analysis. Finally, to correct for multiple comparisons we used spatiotemporal clustering (α = 0.05, 1000 bootstraps). The regression analysis was performed using the LInear MOdeling (LIMO EEG) toolbox (Pernet et al., 2011).

To address the effect of gender on the inter-individual differences we calculated the Pearson correlation between the microstate parameters across the genders. The microstate parameters were highly correlated. Still, we performed the Wilcoxon Rank-sum test to compare the distributions of each microstate parameter across genders and found no significant differences.

### Intra-individual differences between microstates and smartphone behaviours

Within individuals, non-negative matrix factorization (NNMF) was used to identify the configuration of microstate parameters expressed across smartphone behaviours. This resulted in meta-parameters that show the typical configuration of microstates parameters over the behavioural dynamics (meta-behaviours). Prior to the NNMF, any bin with fewer than 25 percentile of interactions — captured across all the two-dimensional bins — was excluded from further analysis. NNMF can decompose a matrix into one or more low rank approximations. We approximated the optimal number of ranks (between 1 and 15) based on cross correlation. The cross-correlation was repeated 100 times with random initializations to ensure stable responses (Wu et al., 2016). We masked the data by randomly removing values in the matrix before performing the NNMF. Next, we reconstructed the masked matrix from the decompositions and compared it to the original matrix to calculate a test-error. The optimal rank was chosen based on the smallest test error averaged across the repetitions. Once the rank was chosen, we performed the stable and reproducible NNMF (STAR-NNMF) method. To perform STAR-NNMF, we first repeated the NNMF, without masking, for 1000 repetitions (Ceolini & Ghosh, 2023). Next, the method selects the most stable and reproducible meta-behaviours and meta-parameters based on the highest median pairwise cross-correlation across all repetitions. The procedure was performed for each individual to capture the intra-individual differences. NNMF was applied using the spams toolbox for MATLAB (mexTrainDL, spams, Inria).

We capture the relationship between smartphone behavior and the transition between microstates using NNMF. Specifically, transition probability, transition duration before, and transition duration after. Transition probability is the probability that a microstate is followed by another microstate (Michel & Koenig, 2018). Transition duration was defined as the average duration of the active microstate prior to and following the transition. The transition parameters — transition probability, transition duration before, and transition duration after — were calculated over each JID two-dimensional bin. Since the distribution of transition durations was skewed, the transition durations before and after were log10 transformed prior to the NNMF analysis.

### Data and code availability

The publication will contain links to the code shared on GitHub and data shared on Dataverse.nl.

The data is published 1 month after publication.

## Author contributions

A.G. conceived the study. A.G. and R.K. designed the study. R.K. analyzed the data aided by A.G. R.K. drafted the report aided by A.G. Both authors helped edit the manuscript.

## Acknowledgment

The authors would like to thank the student assistants who contributed to the data collection at Leiden University.

## Related figures

**Supplementary** Figure 1. Empirical cumulative distribution (ECDF) function for the global explained variance (GEV) across the population. The line colours represent the corresponding microstates. **(a)** GEV of microstates captured during rest. **(b)** GEV for microstates captured during smartphone behaviour.

**Supplementary** Figure 2. Microstate parameters at rest recorded across two different sessions - approximately one week apart - were highly correlated as shown with the Pearson R^2^.

**Supplementary** Figure 3. Marginal relationship between the microstate duration captured at rest and historical smartphone behaviours. **(a)** Contours of the joint-interval distribution (JID) show significant relationships to microstate E and F duration parameters. **b)** T-values show that microstate E duration decreases during multiple slow intervals and microstate F duration decreases during fast followed by slow intervals. These relationships were marginal. T-values were corrected for multiple comparisons using spatiotemporal clustering (1000 bootstraps, α = 0.05).

**Supplementary** Figure 4. Individual-level microstate and non-negative matrix factorization (NNMF) decompositions of all the individuals. (1) Microstates with the corresponding GEV values. (2) Meta- parameters show the microstate parameters – coverage (triangle), occurrence (square), and duration (circle) – expressed across the smartphone behaviours (meta-behaviours). The meta-behaviours are displayed in log10 space. The microstate labels correspond to the population-level clusters. When multiple microstates were clustered to the same population-level microstates these are denoted with a number such as C1,C2. (3) NNMF decompositions displaying the meta-transitions probabilities expressed across the smartphone behaviours (meta-behaviours). The meta-transition probabilities vary across the different microstates with transitions to some microstates happening more often than others. (4) NNMF decompositions displaying the durations occurring before or after a transition across the smartphone behavioural dynamics (meta-behaviours). The difference between the meta-durations before or after is minimal. The durations are log10 transformed.

## References

1. Balerna, M., & Ghosh, A. (2018). The details of past actions on a smartphone touchscreen are reflected by intrinsic sensorimotor dynamics. Npj Digital Medicine 2018 *1*:1, *1*(1), 1–5. 10.1038/s41746-017-0011-3

2. Bréchet, L., Brunet, D., Perogamvros, L., Tononi, G., & Michel, C. M. (2020). EEG microstates of dreams. Scientific Reports 2020 *10*:1, *10*(1), 1–9. 10.1038/s41598-020-74075-z

3. Britz, J., Van De Ville, D., & Michel, C. M. (2010). BOLD correlates of EEG topography reveal rapid resting-state network dynamics. NeuroImage, 52(4), 1162–1170. 10.1016/J.NEUROIMAGE.2010.02.052

4. Brodbeck, V., Kuhn, A., von Wegner, F., Morzelewski, A., Tagliazucchi, E., Borisov, S., Michel, C. M., & Laufs, H. (2012). EEG microstates of wakefulness and NREM sleep. NeuroImage, 62(3), 2129–2139. 10.1016/J.NEUROIMAGE.2012.05.060

5. Ceolini, E., & Ghosh, A. (2023). Common multi-day rhythms in smartphone behavior. Npj Digital Medicine 2023 *6*:1, *6*(1), 1–9. 10.1038/s41746-023-00799-7

6. Ceolini, E., Kock, R., Band, G. P. H., Stoet, G., & Ghosh, A. (2022). Temporal clusters of age-related behavioral alterations captured in smartphone touchscreen interactions. ISCIENCE, 25, 104791. 10.1016/j.isci.2022.104791

7. Chenot, Q., Hamery, C., Truninger, M., Langer, N., De boissezon, X., & Scannella, S. (2024). Investigating the relationship between resting-state EEG microstates and executive functions: A null finding. Cortex, 178, 1–17. 10.1016/J.CORTEX.2024.05.019

8. Delorme, A., & Makeig, S. (2004). EEGLAB: an open source toolbox for analysis of single-trial EEG dynamics including independent component analysis. Journal of Neuroscience Methods, 134(1), 9–21. 10.1016/J.JNEUMETH.2003.10.009

9. Duckrow, R. B., Ceolini, E., Zaveri, H. P., Brooks, C., & Ghosh, A. (2021). Artificial neural network trained on smartphone behavior can trace epileptiform activity in epilepsy Smartphone Artificial Neural Network Reconstructed epileptiform discharges Behavioural features Real Artificial Neural Network Output. ISCIENCE, 24, 102538. 10.1016/j.isci.2021.102538

10. Khanna, A., Pascual-Leone, A., & Farzan, F. (2014). Reliability of Resting-State Microstate Features in Electroencephalography. PLOS ONE, 9(12), e114163. 10.1371/JOURNAL.PONE.0114163

11. Lee, D., Port, N. L., Kruse, W., & Georgopoulos, A. P. (1998). Variability and Correlated Noise in the Discharge of Neurons in Motor and Parietal Areas of the Primate Cortex.

12. Lehmann, D., Ozaki, H., & Pal, I. (1987). EEG alpha map series: brain micro-states by space-oriented adaptive segmentation. Electroencephalography and Clinical Neurophysiology, 67(3), 271–288. 10.1016/0013-4694(87)90025-3

13. Lisboa, P. J. G., Etchells, T. A., Jarman, I. H., & Chambers, S. J. (2013). Finding reproducible cluster partitions for the k-means algorithm. BMC Bioinformatics, 14(SUPPL.1), 1–19. 10.1186/1471-2105-14-S1-S8/TABLES/3

14. Michel, C. M., & Koenig, T. (2018). EEG microstates as a tool for studying the temporal dynamics of whole-brain neuronal networks: A review. NeuroImage, 180, 577–593. 10.1016/J.NEUROIMAGE.2017.11.062

15. Pascual-Marqui, R. D., Michel, C. M., & Lehmann, D. (1995). Segmentation of Brain Electrical Activity into Microstates; Model Estimation and Validation. IEEE Transactions on Biomedical Engineering, 42(7), 658–665. 10.1109/10.391164

16. Pernet, C. R., Chauveau, N., Gaspar, C., & Rousselet, G. A. (2011). LIMO EEG: a toolbox for hierarchical LInear MOdeling of ElectroEncephaloGraphic data. Computational Intelligence and Neuroscience, 2011. 10.1155/2011/831409

17. Poulsen, A. T., Pedroni, A., Langer, N., & Hansen, L. K. (2018). Microstate EEGlab toolbox: An introductory guide. 10.1101/289850

18. Seitzman, B. A., Abell, M., Bartley, S. C., Erickson, M. A., Bolbecker, A. R., & Hetrick, W. P. (2017). Cognitive manipulation of brain electric microstates. NeuroImage, 146, 533–543. 10.1016/J.NEUROIMAGE.2016.10.002

19. Seitzman, B. A., Gratton, C., Laumann, T. O., Gordon, E. M., Adeyemo, B., Dworetsky, A., Kraus, B. T., Gilmore, A. W., Berg, J. J., Ortega, M., Nguyen, A., Greene, D. J., McDermott, K. B., Nelson, S. M., Lessov-Schlaggar, C. N., Schlaggar, B. L., Dosenbach, N. U. F., & Petersen, S. E. (2019). Trait-like variants in human functional brain networks. Proceedings of the National Academy of Sciences of the United States of America, 116(45), 22851–22861. 10.1073/PNAS.1902932116/SUPPL_FILE/PNAS.1902932116.SAPP.PDF

20. Smitha, K. A., Akhil Raja, K., Arun, K. M., Rajesh, P. G., Thomas, B., Kapilamoorthy, T. R., & Kesavadas, C. (2017). Resting state fMRI: A review on methods in resting state connectivity analysis and resting state networks. Neuroradiology Journal, 30(4), 305–317. 10.1177/1971400917697342/ASSET/IMAGES/LARGE/10.1177_1971400917697342-FIG7.JPEG

21. Tarailis, P., Koenig, T., Michel, C. M., & Griškova-Bulanova, I. (2023). The Functional Aspects of Resting EEG Microstates: A Systematic Review. Brain Topography 2023 *37*:2, *37*(2), 181–217. 10.1007/S10548-023-00958-9

22. Tomescu, M. I., Rihs, T. A., Rochas, V., Hardmeier, M., Britz, J., Allali, G., Fuhr, P., Eliez, S., & Michel, C. M. (2018). From swing to cane: Sex differences of EEG resting-state temporal patterns during maturation and aging. Developmental Cognitive Neuroscience, 31, 58–66. 10.1016/J.DCN.2018.04.011

23. Wang, Y. X., & Zhang, Y. J. (2013). Nonnegative matrix factorization: A comprehensive review. IEEE Transactions on Knowledge and Data Engineering, 25(6), 1336–1353. 10.1109/TKDE.2012.51

24. Wu, S., Joseph, A., Hammonds, A. S., Celniker, S. E., Yu, B., & Frise, E. (2016). Stability-driven nonnegative matrix factorization to interpret Spatial gene expression and build local gene networks. Proceedings of the National Academy of Sciences of the United States of America, 113(16), 4290–4295. 10.1073/PNAS.1521171113/SUPPL_FILE/PNAS.1521171113.SD06.XLSX

25. Zanesco, A. P., King, B. G., Skwara, A. C., & Saron, C. D. (2020). Within and between-person correlates of the temporal dynamics of resting EEG microstates. NeuroImage, 211, 116631. 10.1016/J.NEUROIMAGE.2020.116631

