## Supplementary figures and images for "Neural microstates in real-world behaviour captured on the smartphone"

### Supplementary Figure 1

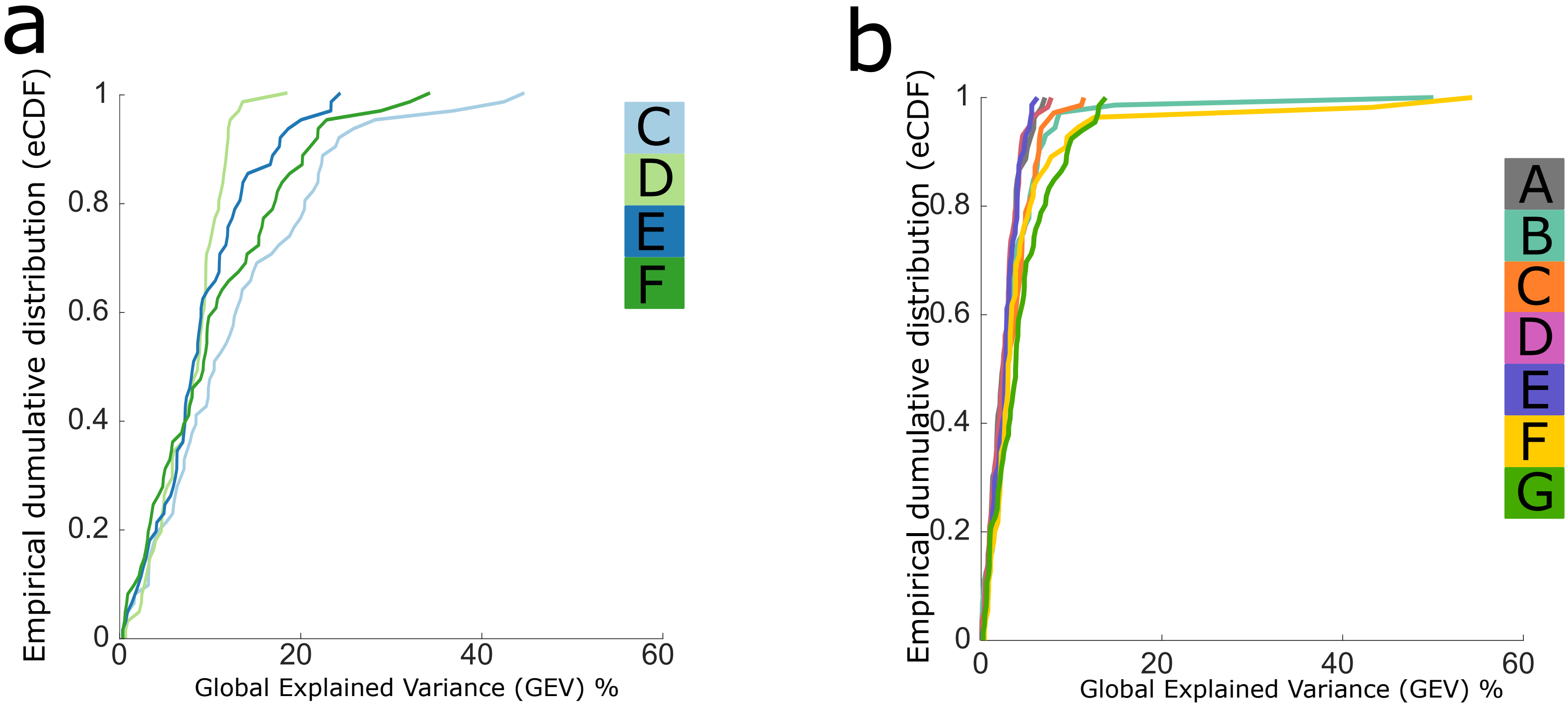

### Supplementary Figure 2

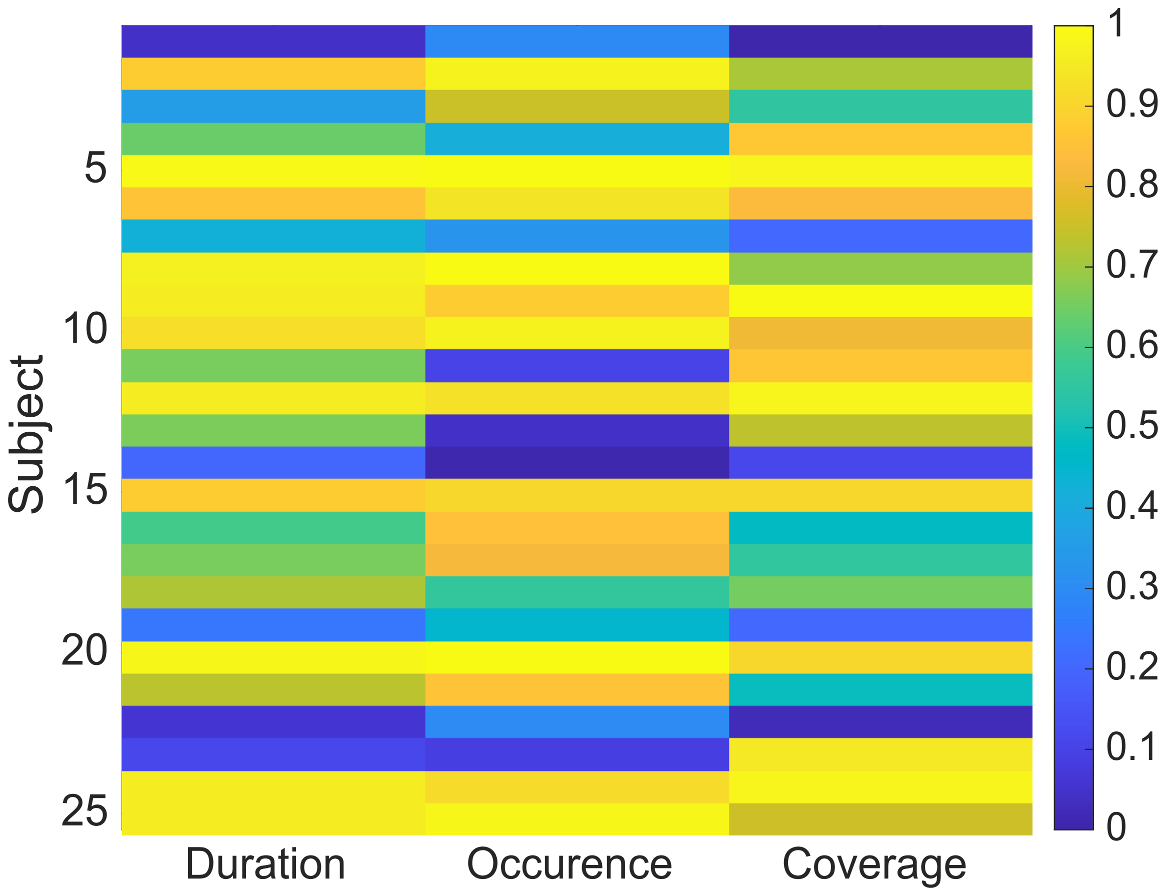

### Supplementary Figure 3

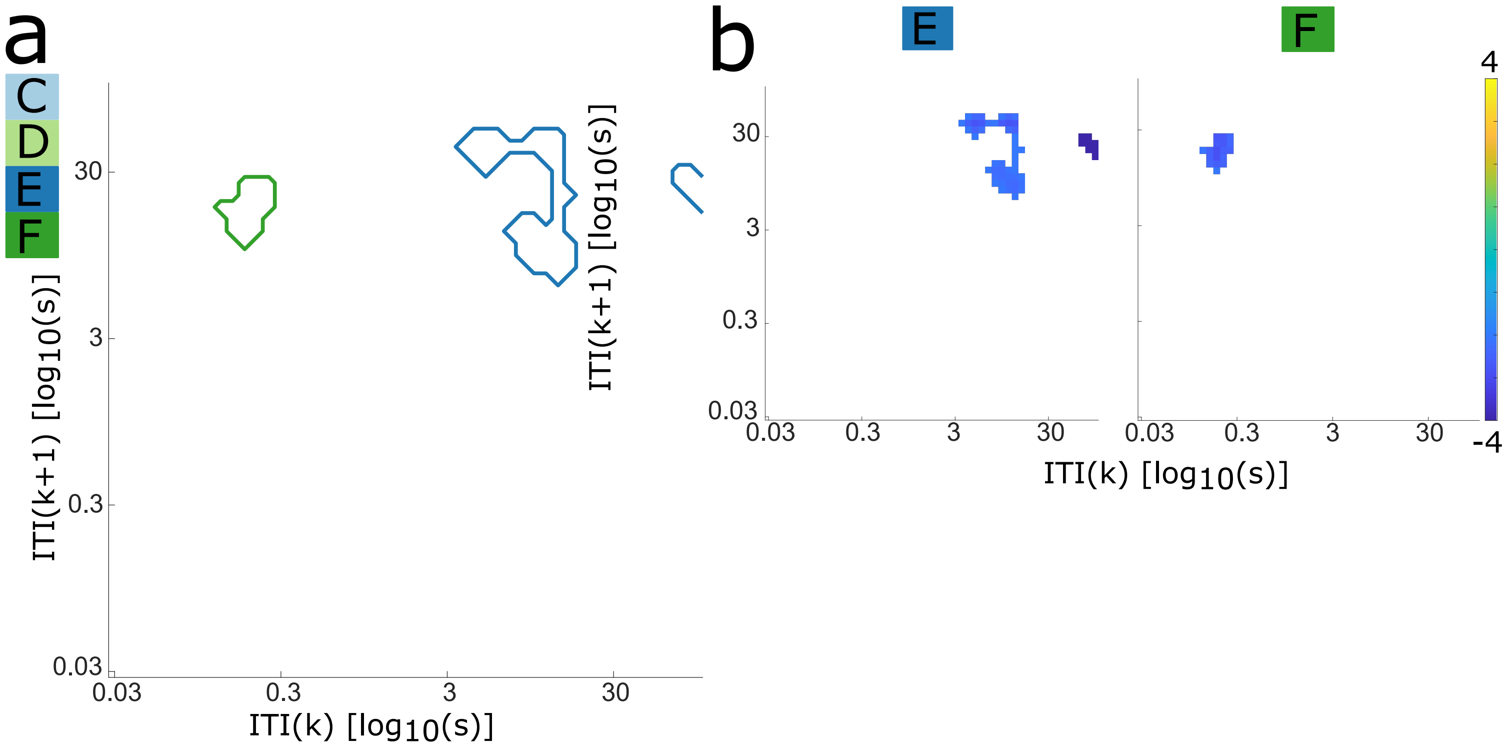
