## Supplementary Figure 4 for "Neural microstates in real-world behaviour captured on the smartphone"

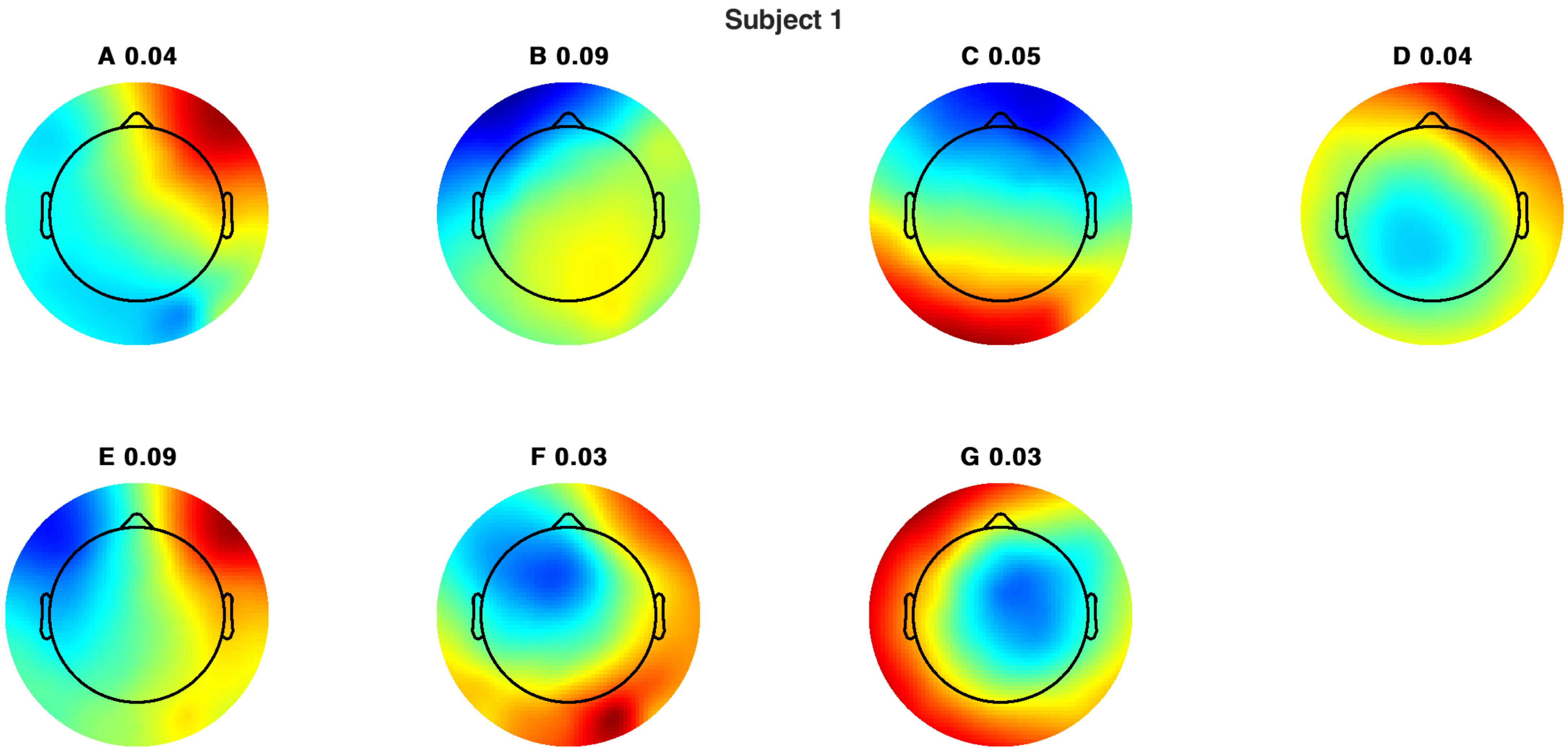
Subject 1
A 0.04
B 0.09
C 0.05
D 0.04
E 0.09
F 0.03
G 0.03

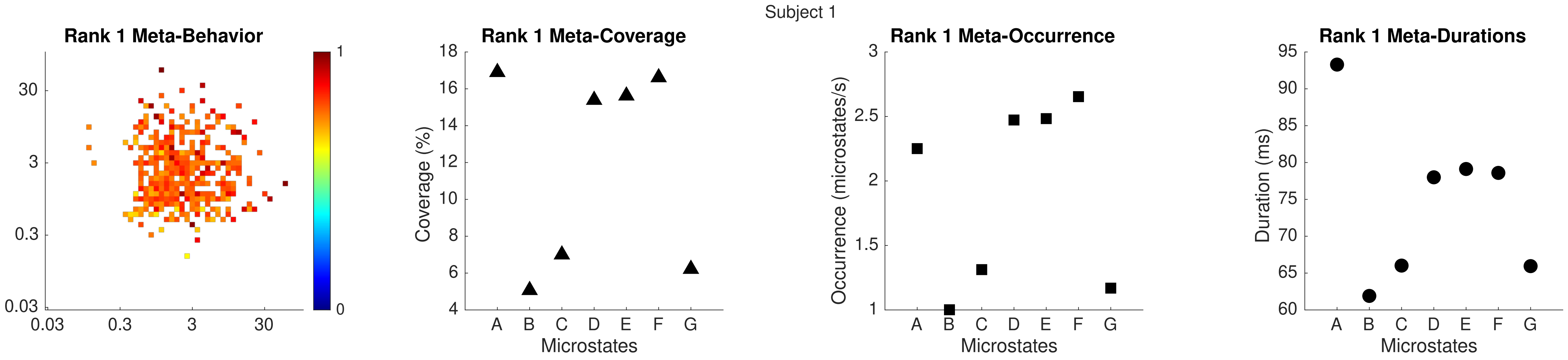
Subject 1
Rank 1 Meta-Behavior
Rank 1 Meta-Coverage
Rank 1 Meta-Occurrence
Rank 1 Meta-Durations
Coverage (%)
Occurrence (microstates/s)
Duration (ms)
Microstates
Microstates
Microstates
A B C D E F G

Subject 1

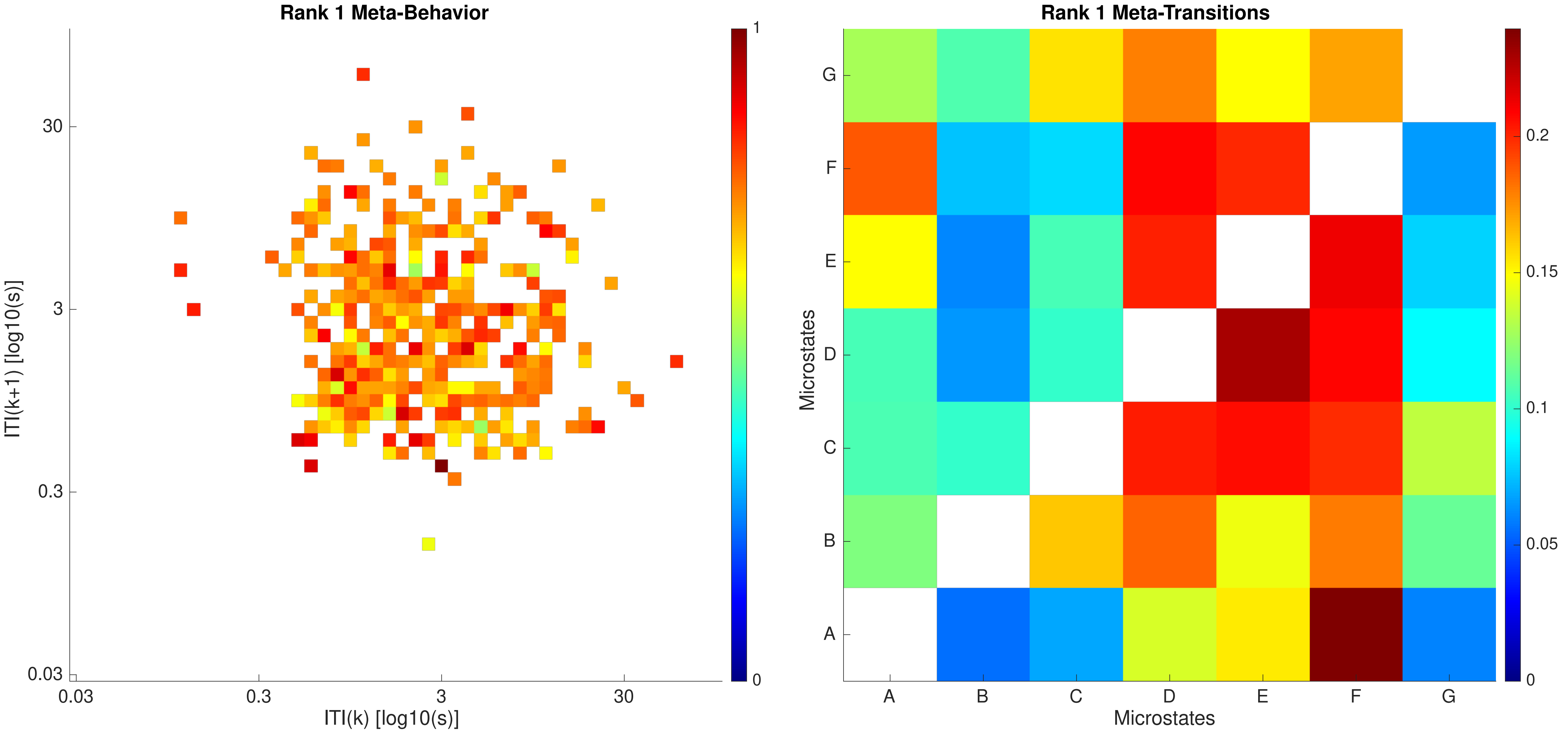

Rank 1 Meta-Behavior
ITI(k+1) [log10(s)]
30
3
0.3
0.03
ITI(k) [log10(s)]
0.03
0.3
3
30
1
0
Rank 1 Meta-Transitions
Microstates
G
F
E
D
C
B
A
Microstates
A
B
C
D
E
F
G
0.2
0.15
0.1
0.05
0

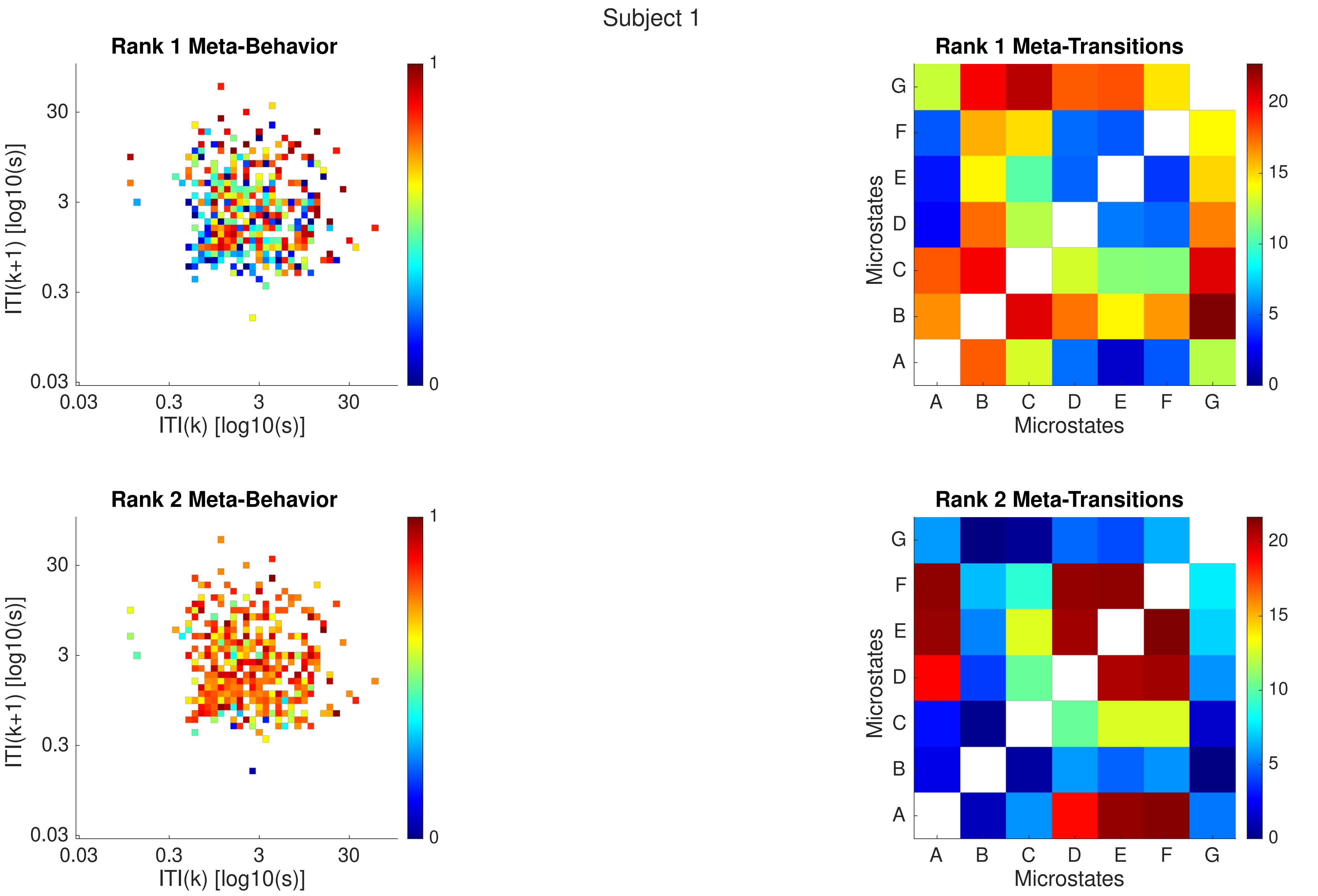
Subject 1
Rank 1 Meta-Behavior
ITI(k+1) [log10(s)]
ITI(k) [log10(s)]
0.03
0.3
3
30
0
1
Rank 1 Meta-Transitions
Microstates
A
B
C
D
E
F
G
0
5
10
15
20
Rank 2 Meta-Behavior
ITI(k+1) [log10(s)]
ITI(k) [log10(s)]
Rank 2 Meta-Transitions
Microstates

Subject 1

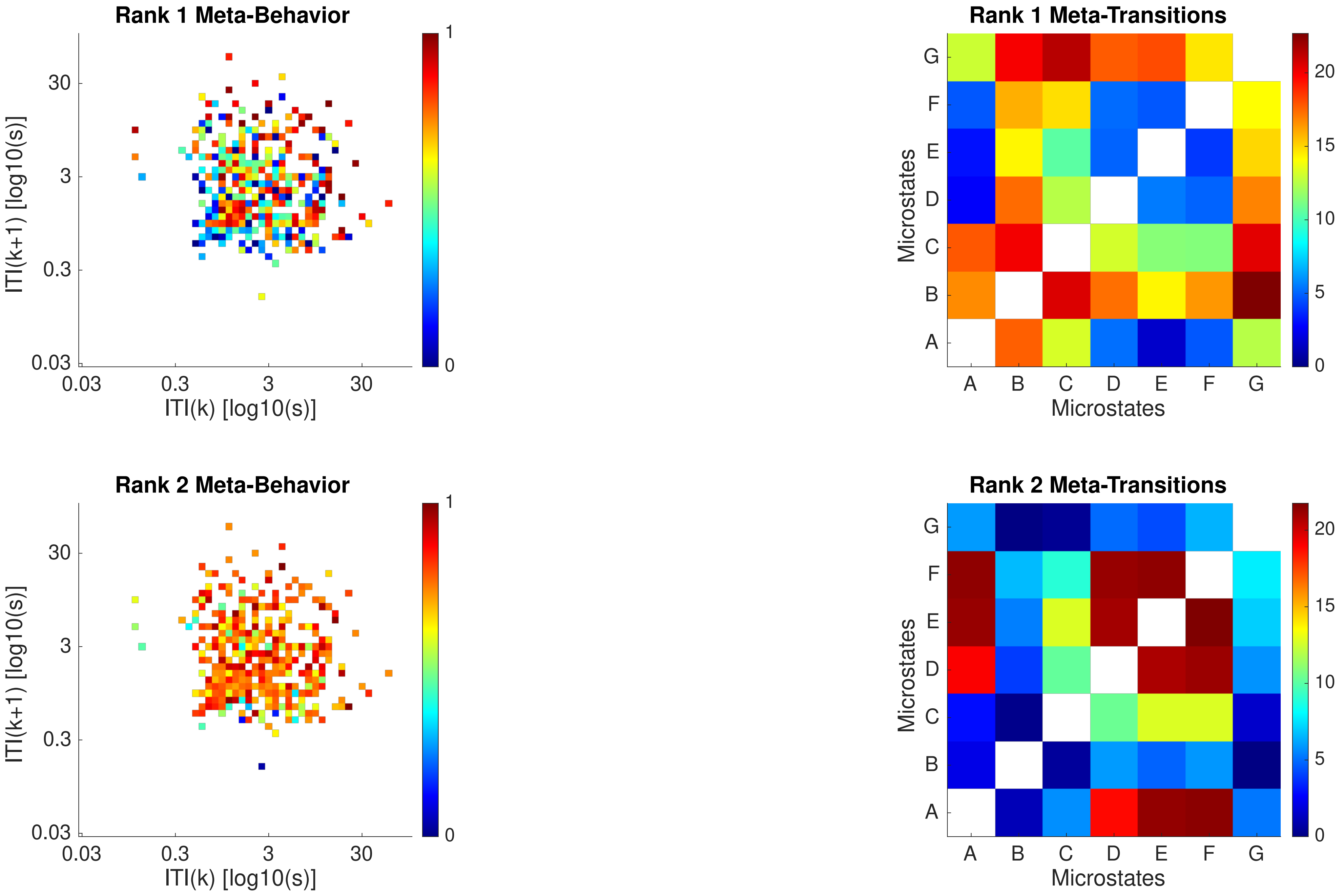

Rank 1 Meta-Behavior
ITI(k+1) [log10(s)]
ITI(k) [log10(s)]
0.03
0.3
3
30
0
1
Rank 1 Meta-Transitions
Microstates
A
B
C
D
E
F
G
0
5
10
15
20
Rank 2 Meta-Behavior
ITI(k+1) [log10(s)]
ITI(k) [log10(s)]
Rank 2 Meta-Transitions
Microstates

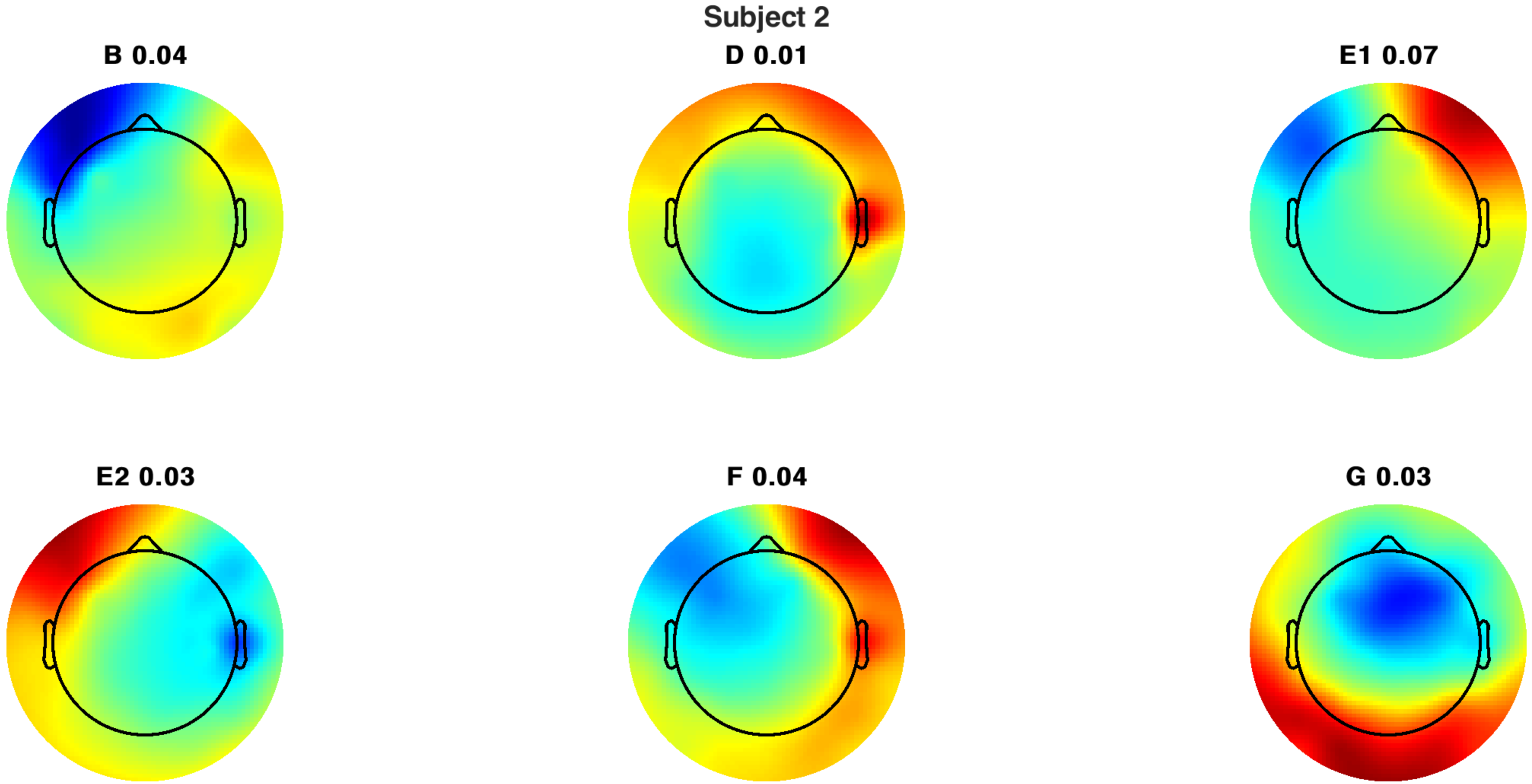
Subject 2
B 0.04
D 0.01
E1 0.07
E2 0.03
F 0.04
G 0.03

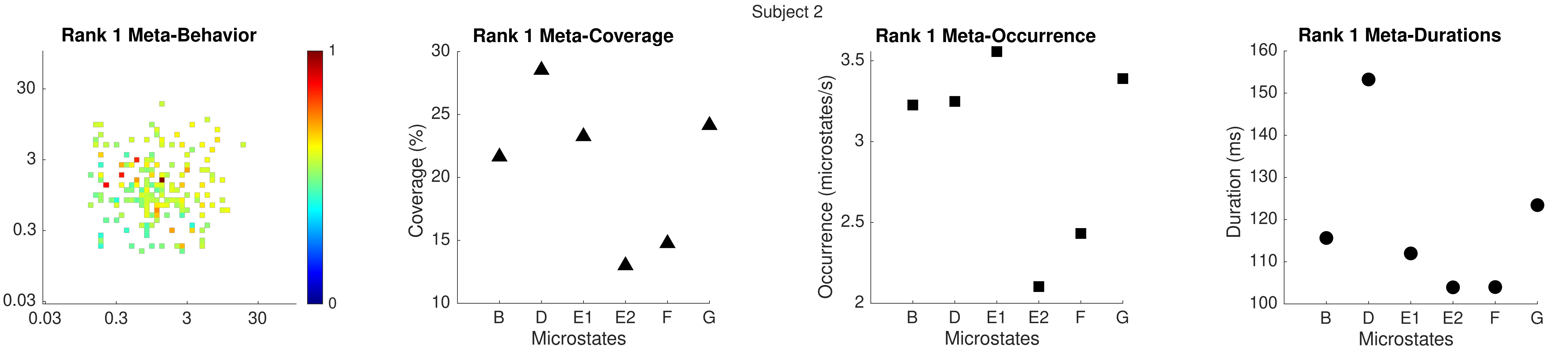

Subject 2
Rank 1 Meta-Behavior
30
3
0.3
0.03
0.03
0.3
3
30
1
0
Rank 1 Meta-Coverage
Coverage (%)
30
25
20
15
10
B
D
E1
E2
F
G
Microstates
Rank 1 Meta-Occurrence
Occurrence (microstates/s)
3.5
3
2.5
2
B
D
E1
E2
F
G
Microstates
Rank 1 Meta-Durations
Duration (ms)
160
150
140
130
120
110
100
B
D
E1
E2
F
G
Microstates

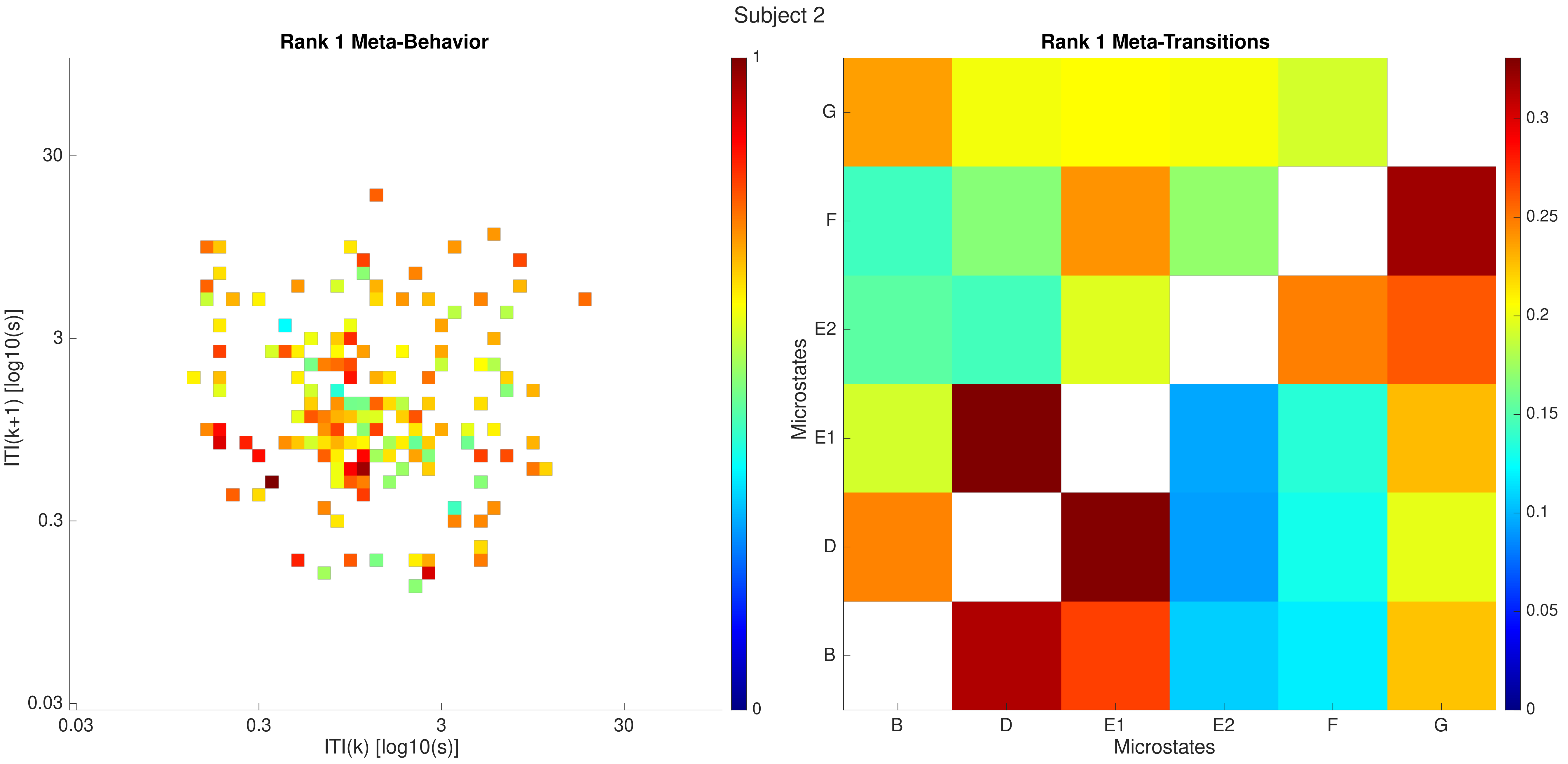
Subject 2
Rank 1 Meta-Behavior
ITI(k+1) [log10(s)]
ITI(k) [log10(s)]
0.03
0.3
3
30
0
1
Rank 1 Meta-Transitions
Microstates
B
D
E1
E2
F
G
0
0.05
0.1
0.15
0.2
0.25
0.3

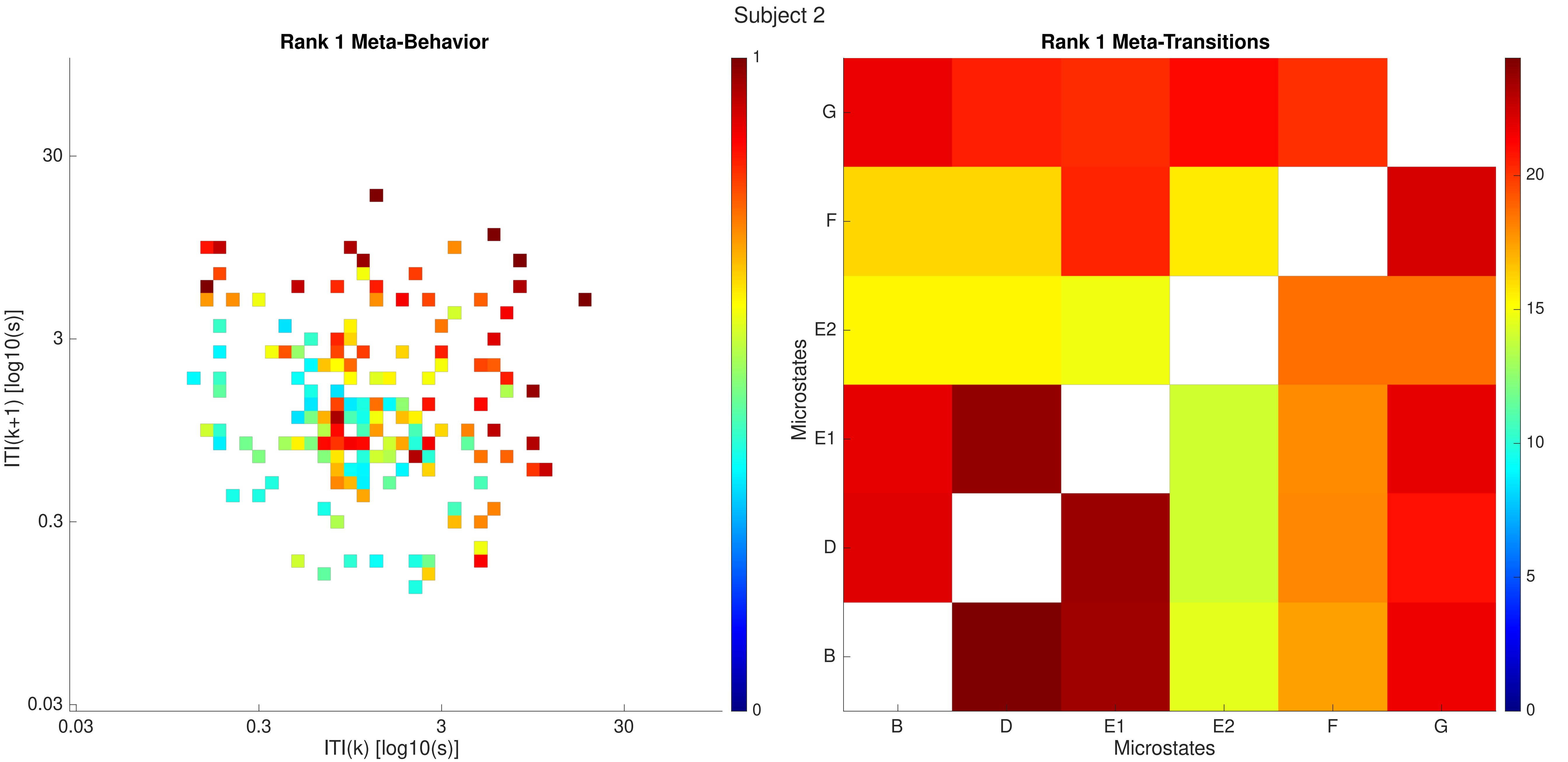
Subject 2
Rank 1 Meta-Behavior
ITI(k+1) [log10(s)]
30
3
0.3
0.03
ITI(k) [log10(s)]
0.03
0.3
3
30
1
0
Rank 1 Meta-Transitions
Microstates
G
F
E2
E1
D
B
Microstates
B
D
E1
E2
F
G
20
15
10
5
0

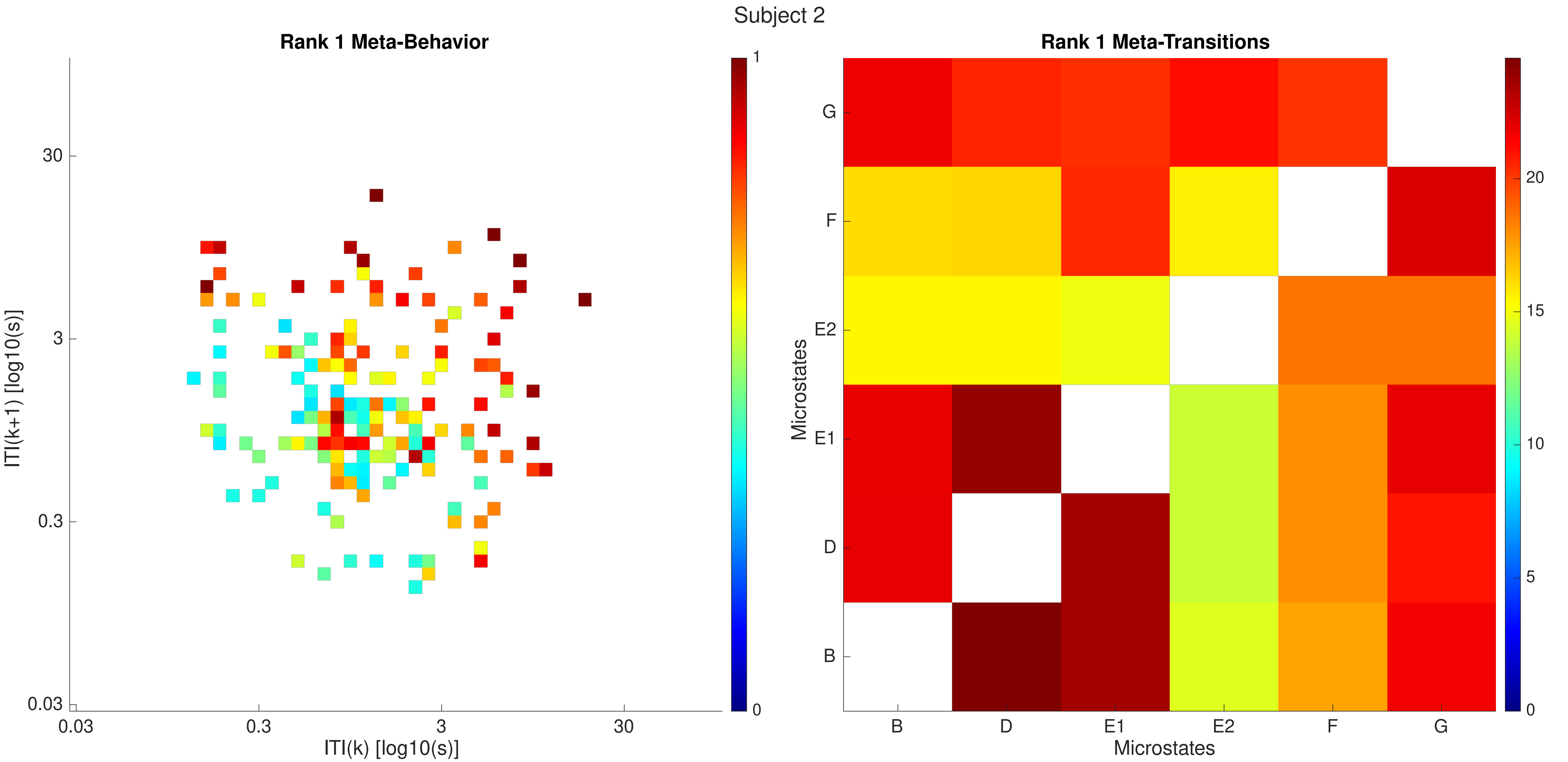

Subject 2
Rank 1 Meta-Behavior
ITI(k+1) [log10(s)]
30
3
0.3
0.03
0.03
0.3
3
30
ITI(k) [log10(s)]
1
0
Rank 1 Meta-Transitions
Microstates
G
F
E2
E1
D
B
B
D
E1
E2
F
G
Microstates
20
15
10
5
0

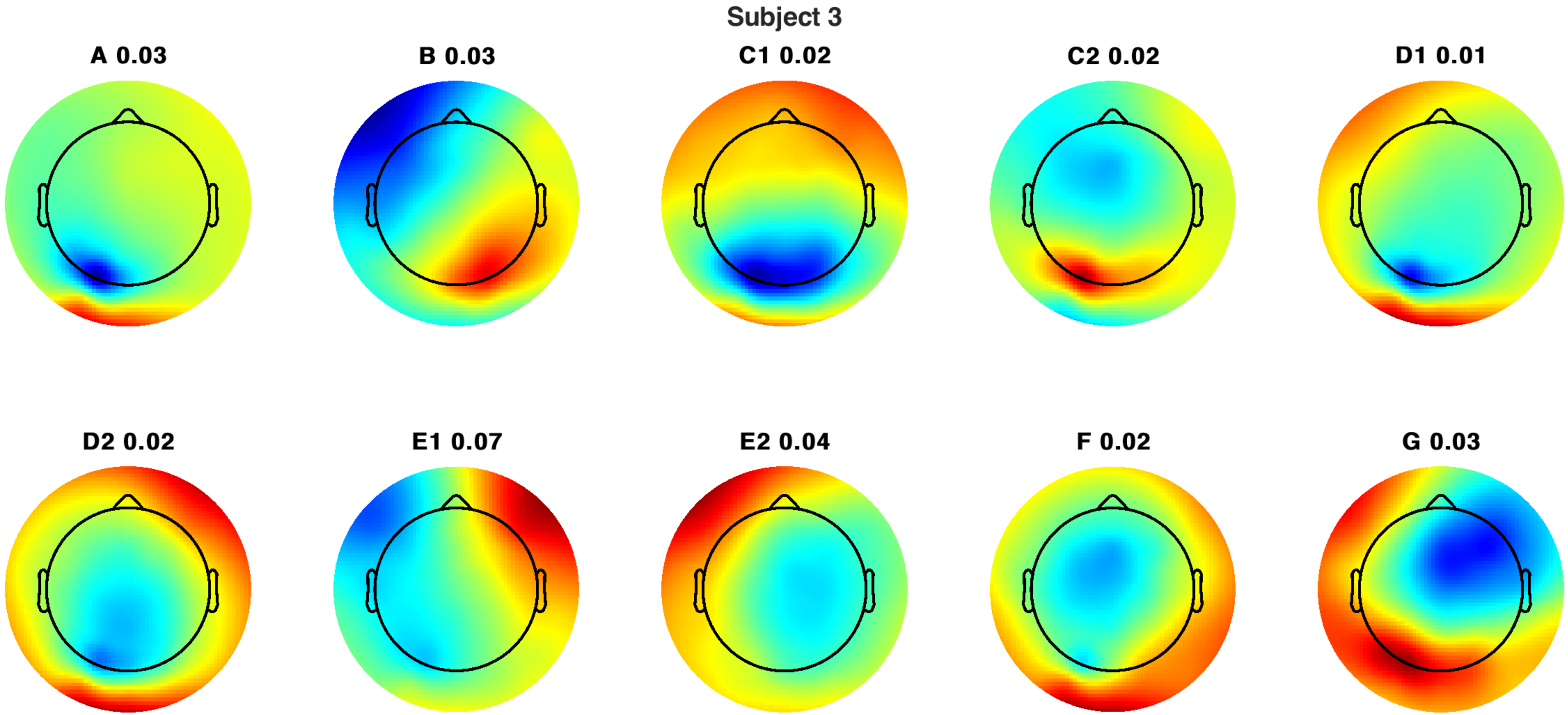
Subject 3
A 0.03
B 0.03
C1 0.02
C2 0.02
D1 0.01
D2 0.02
E1 0.07
E2 0.04
F 0.02
G 0.03

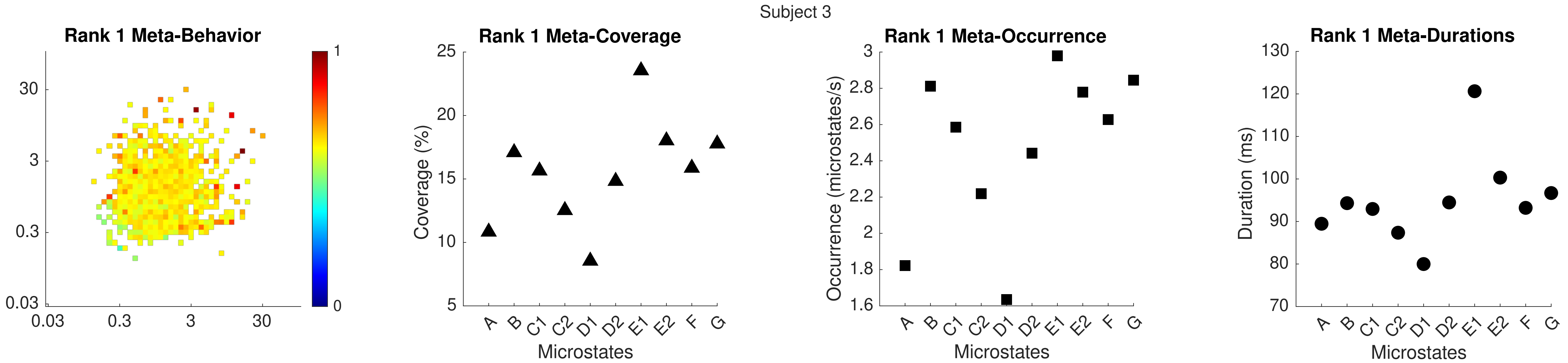

Subject 3
Rank 1 Meta-Behavior
30
3
0.3
0.03
0.03
0.3
3
30
1
0
Rank 1 Meta-Coverage
Coverage (%)
25
20
15
10
5
A B C1 C2 D1 D2 E1 E2 F G
Microstates
Rank 1 Meta-Occurrence
Occurrence (microstates/s)
3
2.8
2.6
2.4
2.2
2
1.8
1.6
A B C1 C2 D1 D2 E1 E2 F G
Microstates
Rank 1 Meta-Durations
Duration (ms)
130
120
110
100
90
80
70
A B C1 C2 D1 D2 E1 E2 F G
Microstates

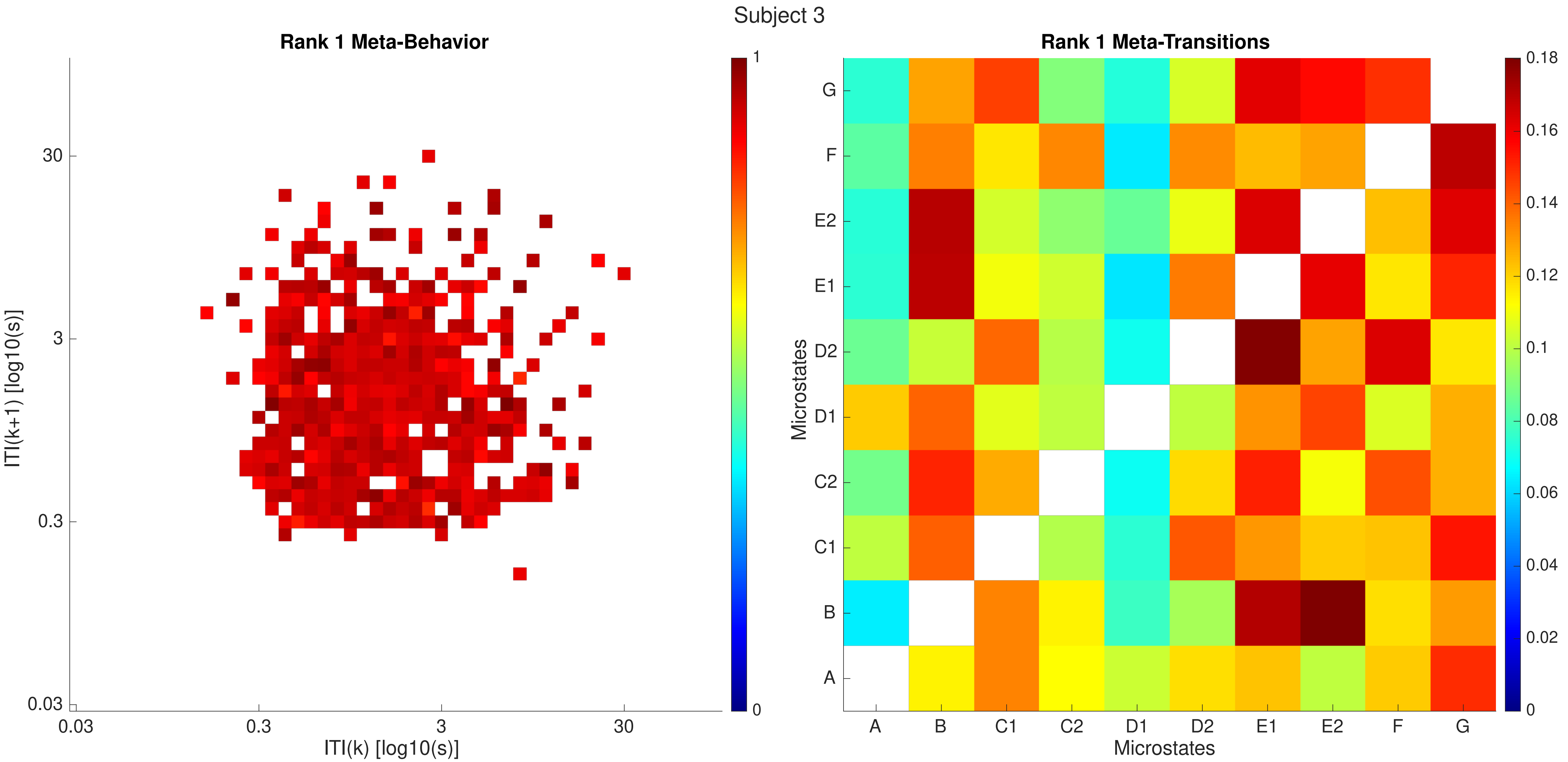

Subject 3
Rank 1 Meta-Behavior
ITI(k+1) [log10(s)]
ITI(k) [log10(s)]
0.03
0.3
3
30
0
1
Rank 1 Meta-Transitions
Microstates
A
B
C1
C2
D1
D2
E1
E2
F
G
0
0.02
0.04
0.06
0.08
0.1
0.12
0.14
0.16
0.18

Subject 3

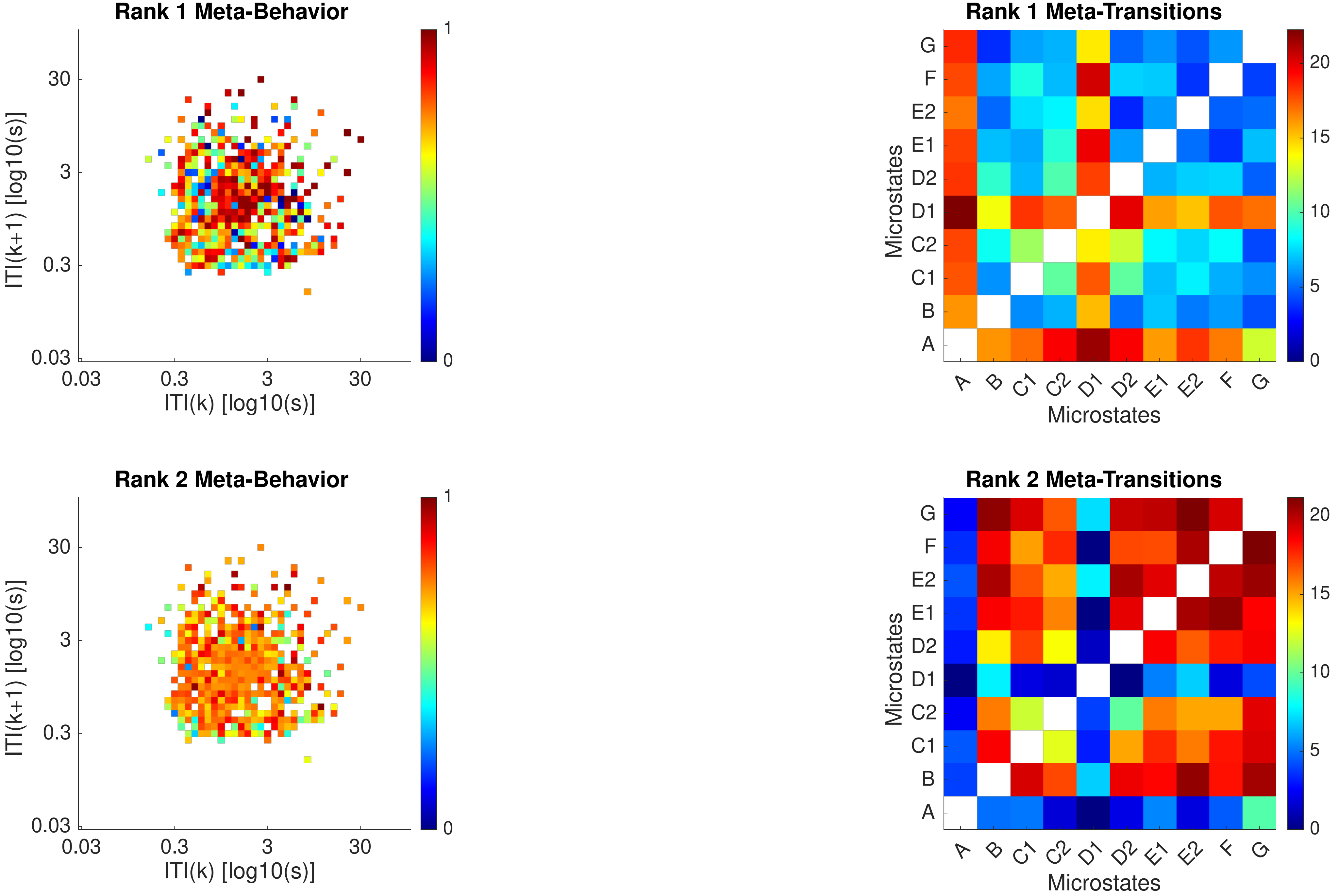

Rank 1 Meta-Behavior
ITI(k+1) [log10(s)]
ITI(k) [log10(s)]
0.03
0.3
3
30
0
1
Rank 1 Meta-Transitions
Microstates
A
B
C1
C2
D1
D2
E1
E2
F
G
0
5
10
15
20
Rank 2 Meta-Behavior
Rank 2 Meta-Transitions

Subject 3

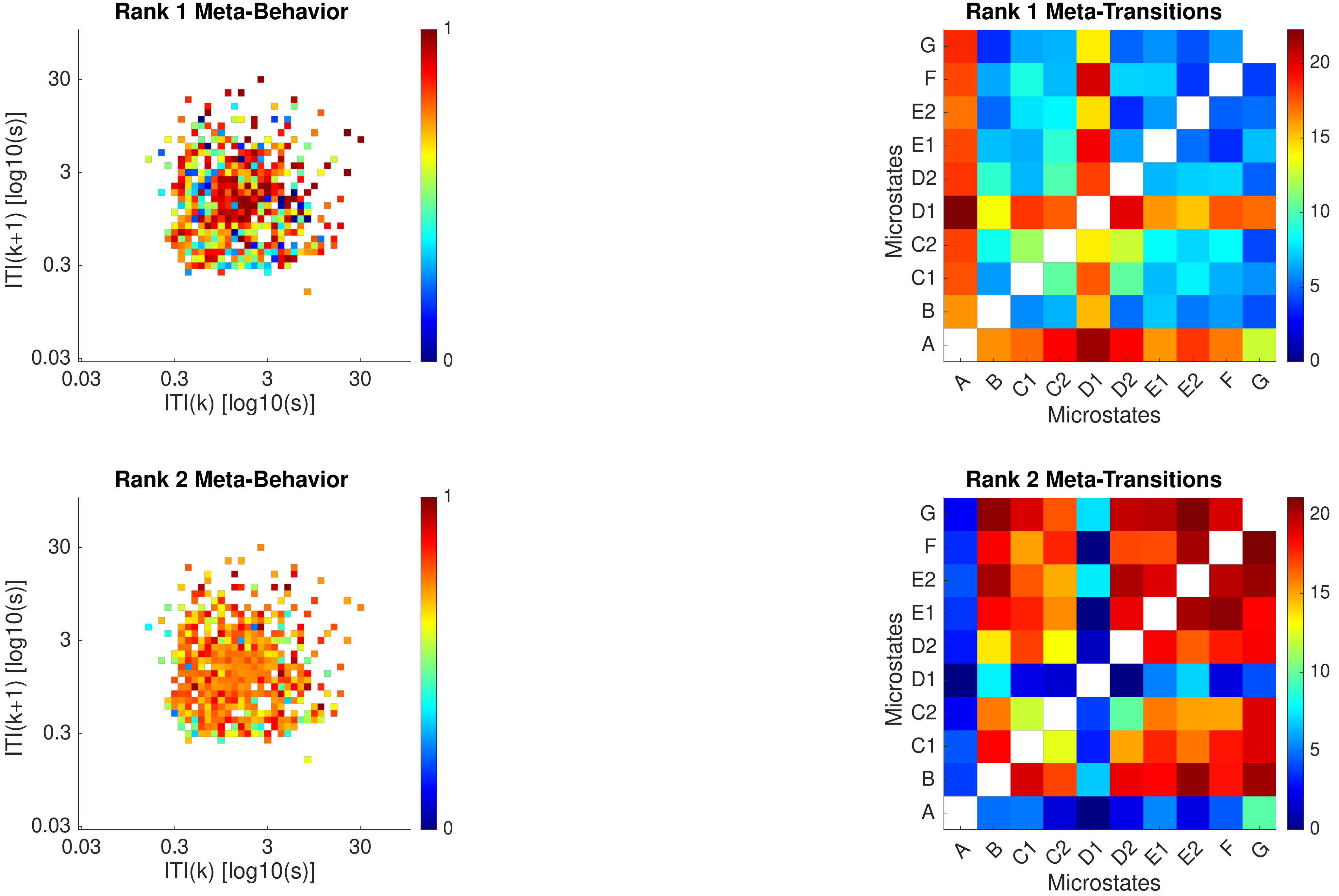
Rank 1 Meta-Behavior
ITI(k+1) [log10(s)]
ITI(k) [log10(s)]
0.03
0.3
3
30
0
1
Rank 1 Meta-Transitions
Microstates
A
B
C1
C2
D1
D2
E1
E2
F
G
0
5
10
15
20
Rank 2 Meta-Behavior
ITI(k+1) [log10(s)]
ITI(k) [log10(s)]
Rank 2 Meta-Transitions
Microstates

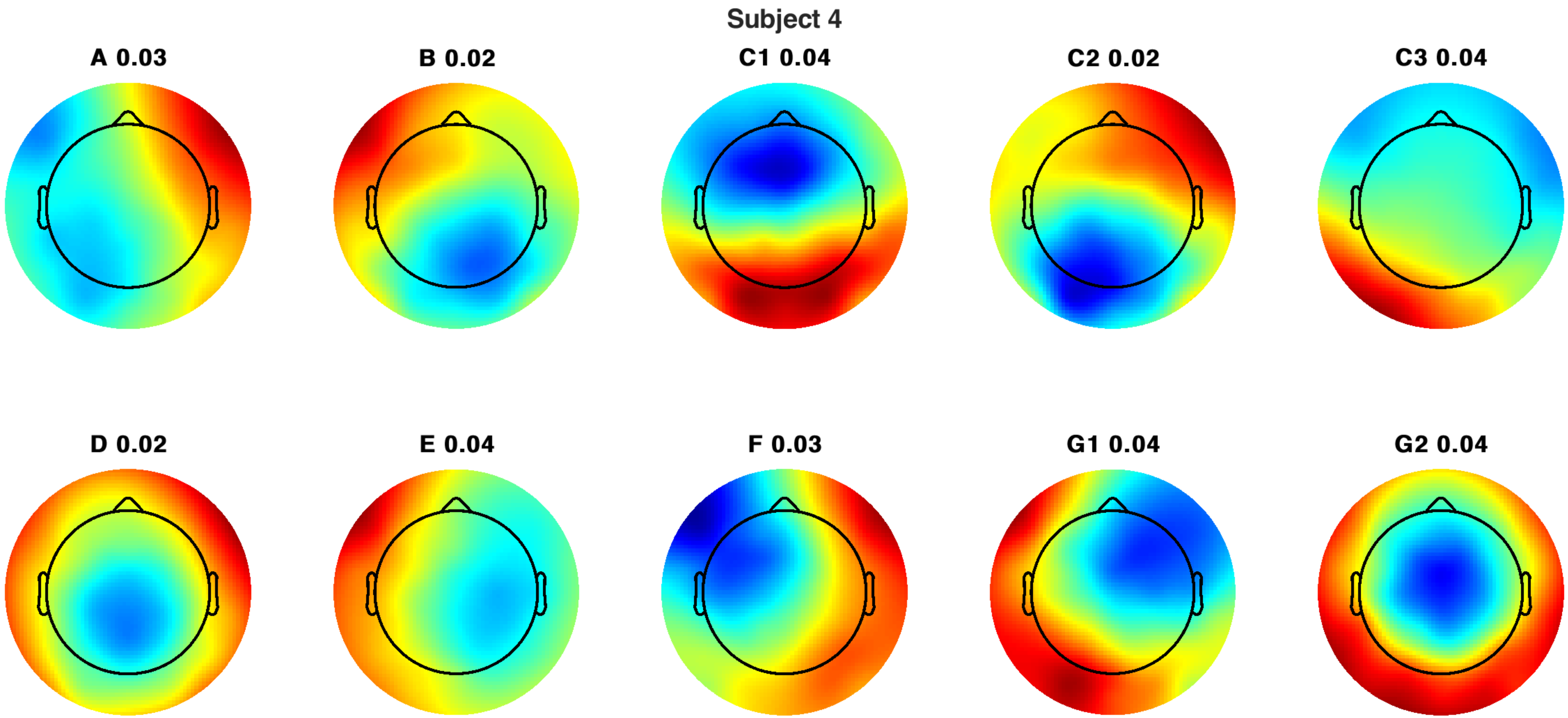
Subject 4
A 0.03
B 0.02
C1 0.04
C2 0.02
C3 0.04
D 0.02
E 0.04
F 0.03
G1 0.04
G2 0.04

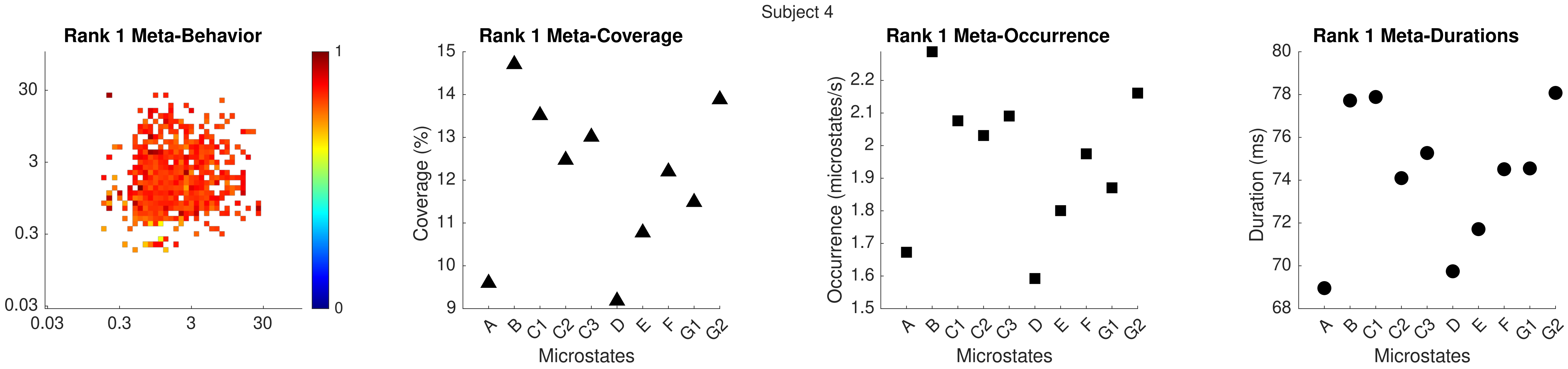

Subject 4
Rank 1 Meta-Behavior
Rank 1 Meta-Coverage
Rank 1 Meta-Occurrence
Rank 1 Meta-Durations
Coverage (%)
Occurrence (microstates/s)
Duration (ms)
Microstates
A B C1 C2 C3 D E F G1 G2

Subject 4

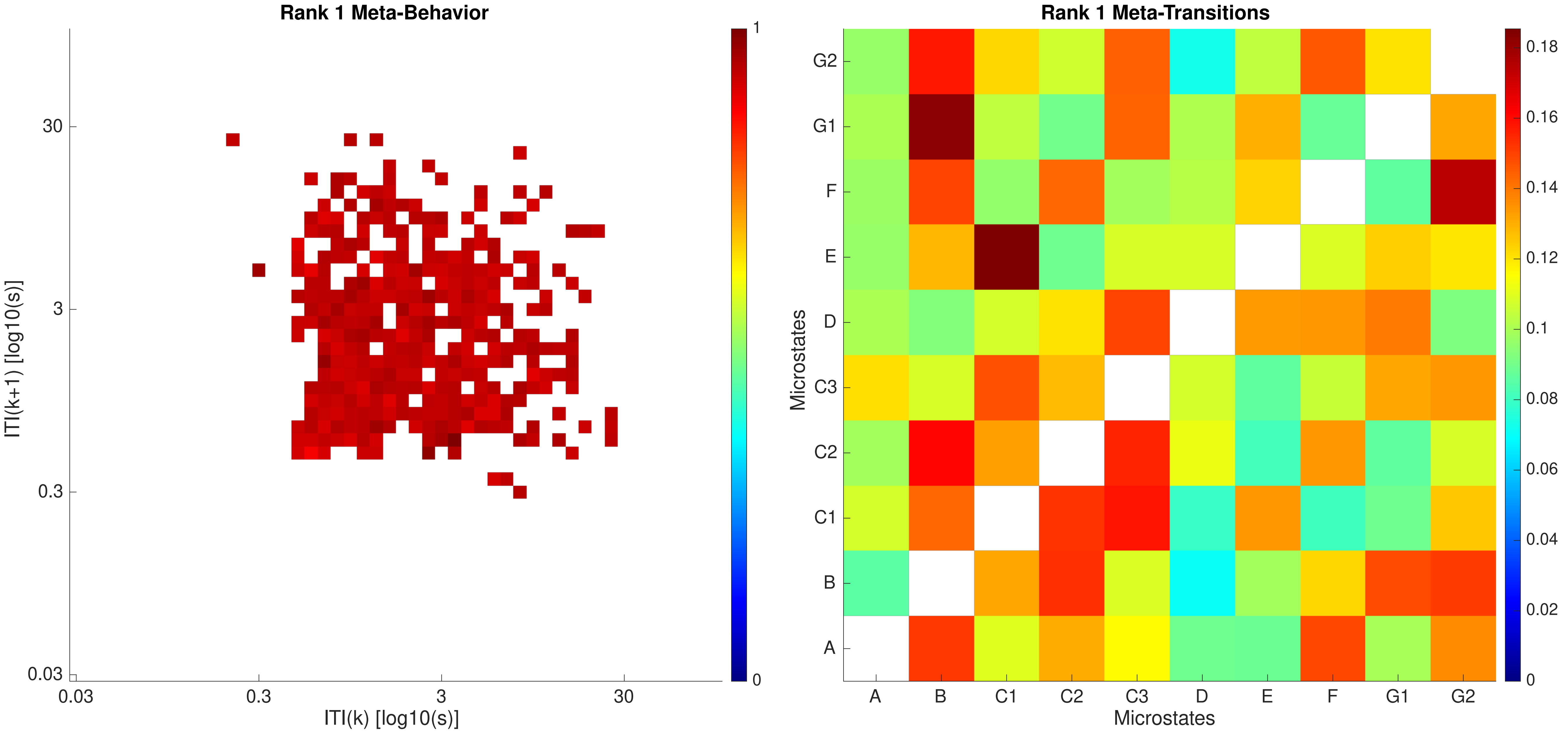

Rank 1 Meta-Behavior
ITI(k+1) [log10(s)]
ITI(k) [log10(s)]
0.03
0.3
3
30
1
0
Rank 1 Meta-Transitions
Microstates
A
B
C1
C2
C3
D
E
F
G1
G2
0.18
0.16
0.14
0.12
0.1
0.08
0.06
0.04
0.02
0

Subject 4

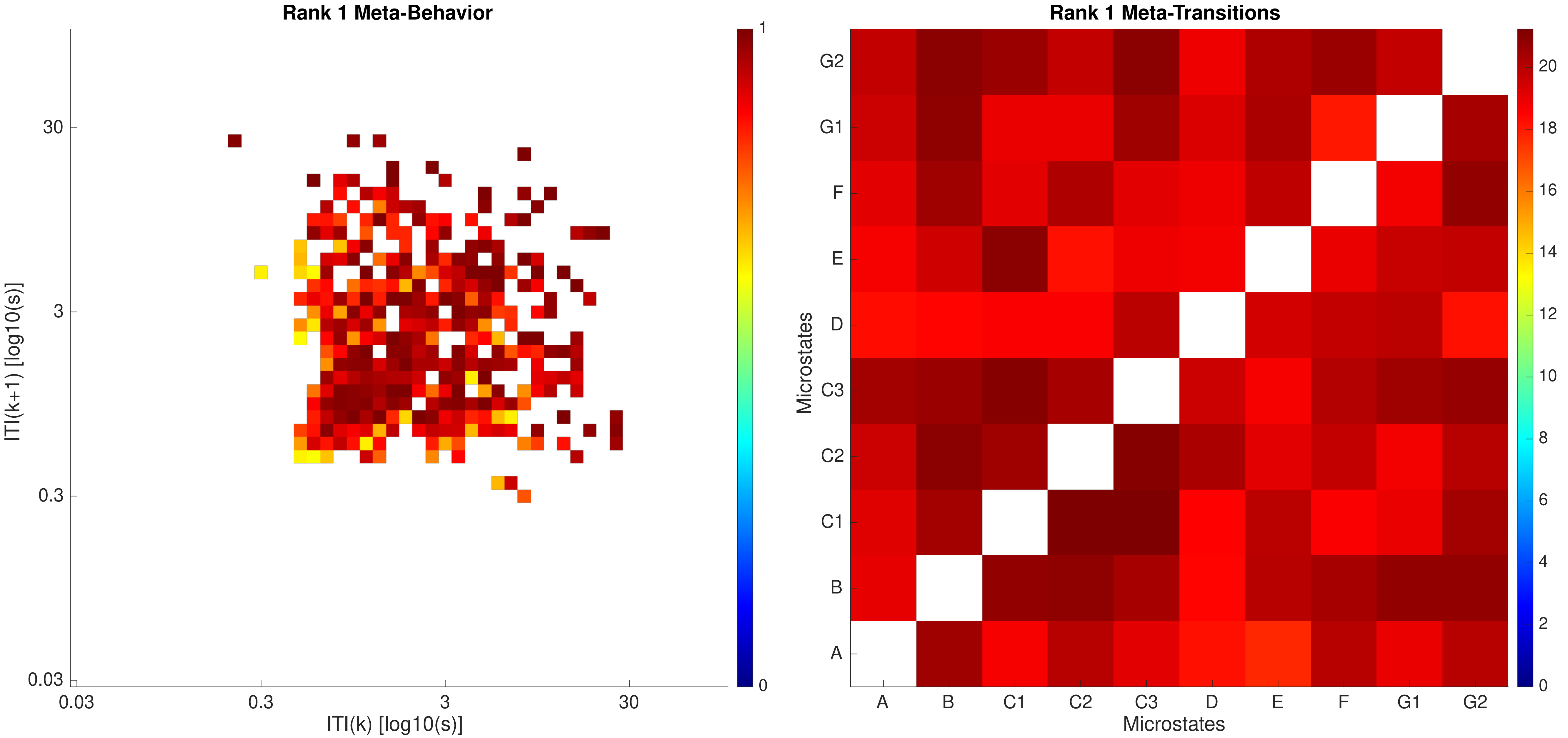

Rank 1 Meta-Behavior
ITI(k+1) [log10(s)]
ITI(k) [log10(s)]
0.03
0.3
3
30
0
1
Rank 1 Meta-Transitions
Microstates
A
B
C1
C2
C3
D
E
F
G1
G2
0
2
4
6
8
10
12
14
16
18
20

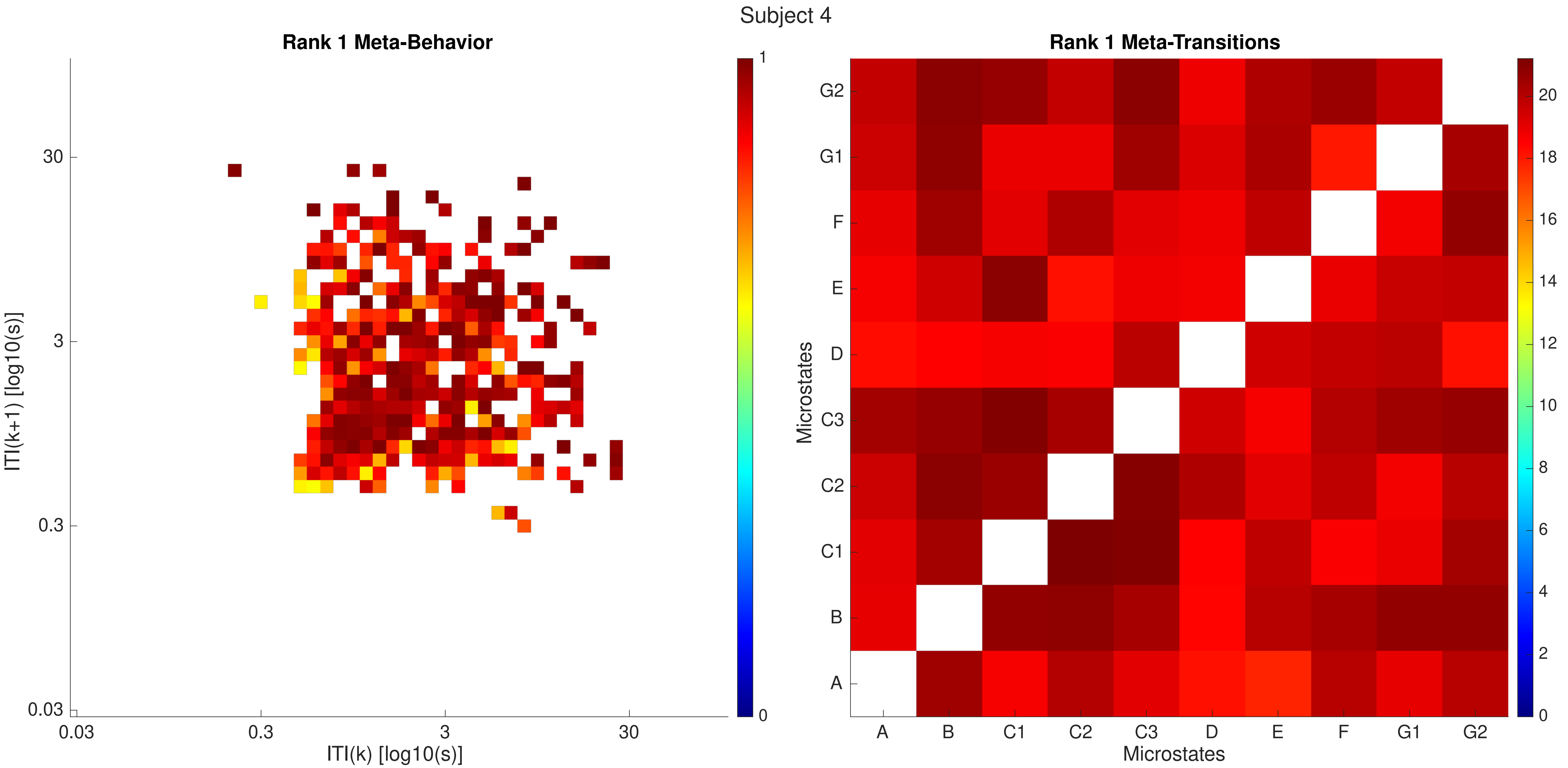
Subject 4
Rank 1 Meta-Behavior
ITI(k+1) [log10(s)]
ITI(k) [log10(s)]
0.03
0.3
3
30
0
1
Rank 1 Meta-Transitions
Microstates
A
B
C1
C2
C3
D
E
F
G1
G2
0
2
4
6
8
10
12
14
16
18
20

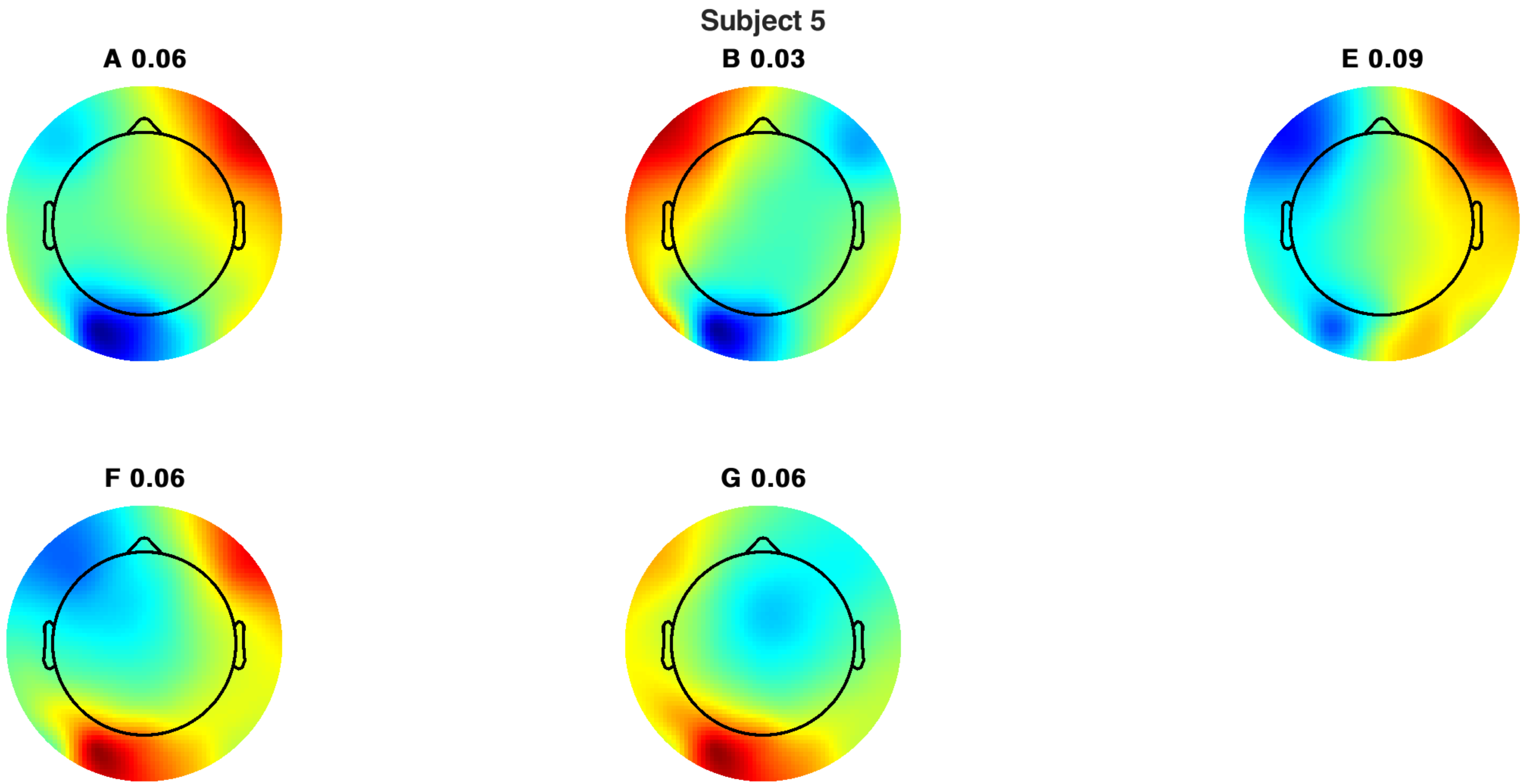
Subject 5
A 0.06
B 0.03
E 0.09
F 0.06
G 0.06

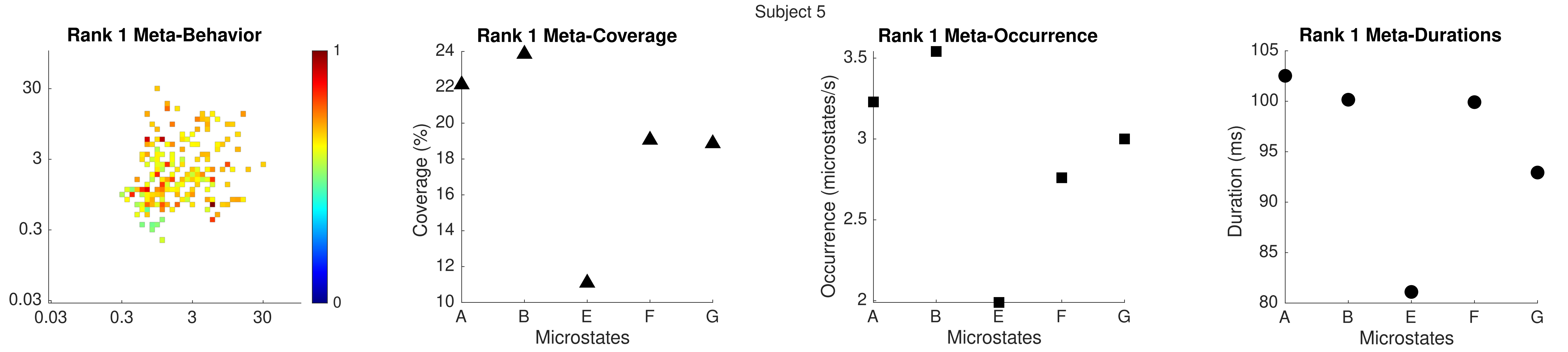

Subject 5
Rank 1 Meta-Behavior
Rank 1 Meta-Coverage
Coverage (%)
Microstates
Rank 1 Meta-Occurrence
Occurrence (microstates/s)
Microstates
Rank 1 Meta-Durations
Duration (ms)
Microstates

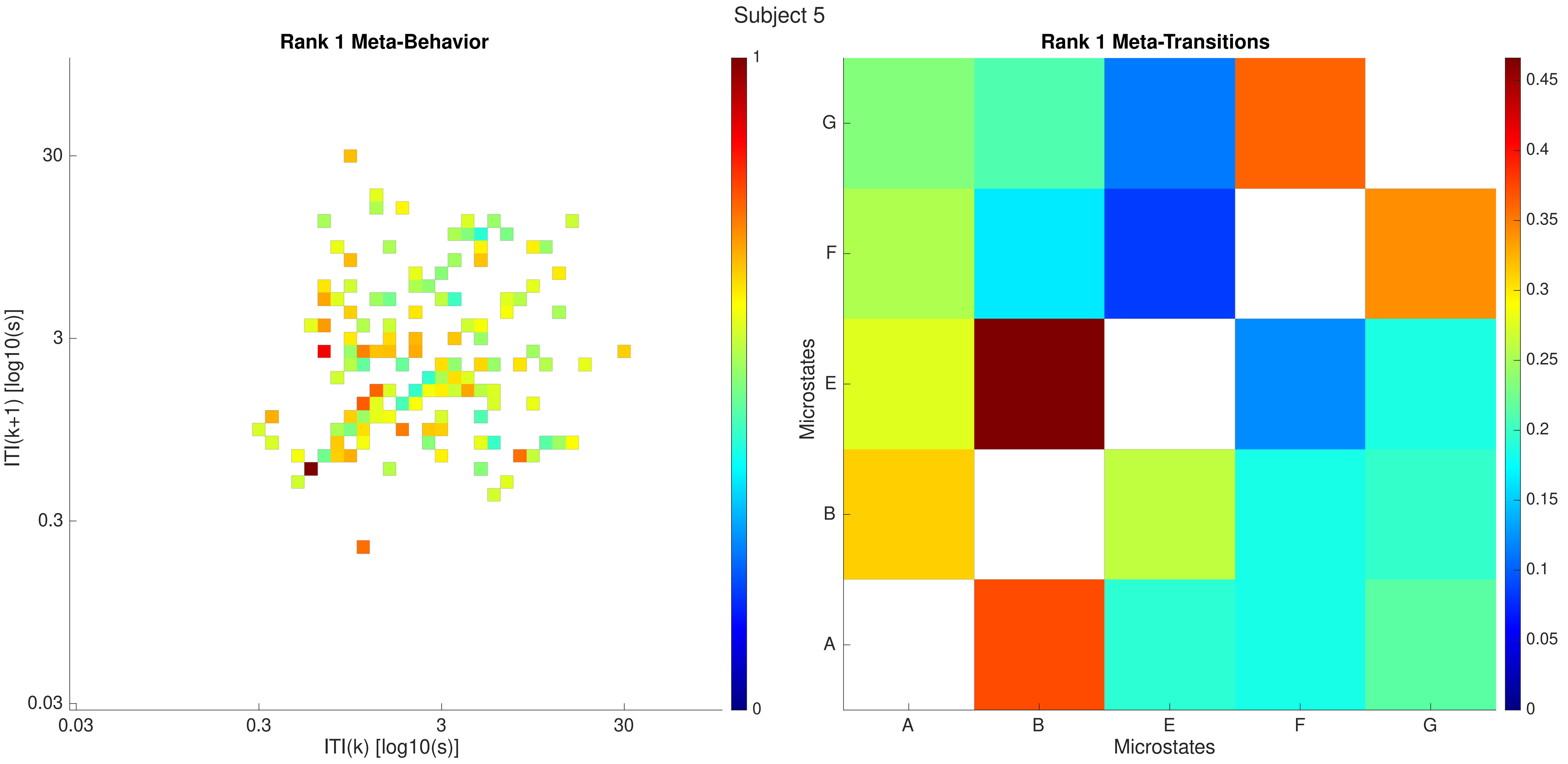

Subject 5
Rank 1 Meta-Behavior
ITI(k+1) [log10(s)]
ITI(k) [log10(s)]
0.03
0.3
3
30
0
1
Rank 1 Meta-Transitions
Microstates
A
B
E
F
G
0
0.05
0.1
0.15
0.2
0.25
0.3
0.35
0.4
0.45

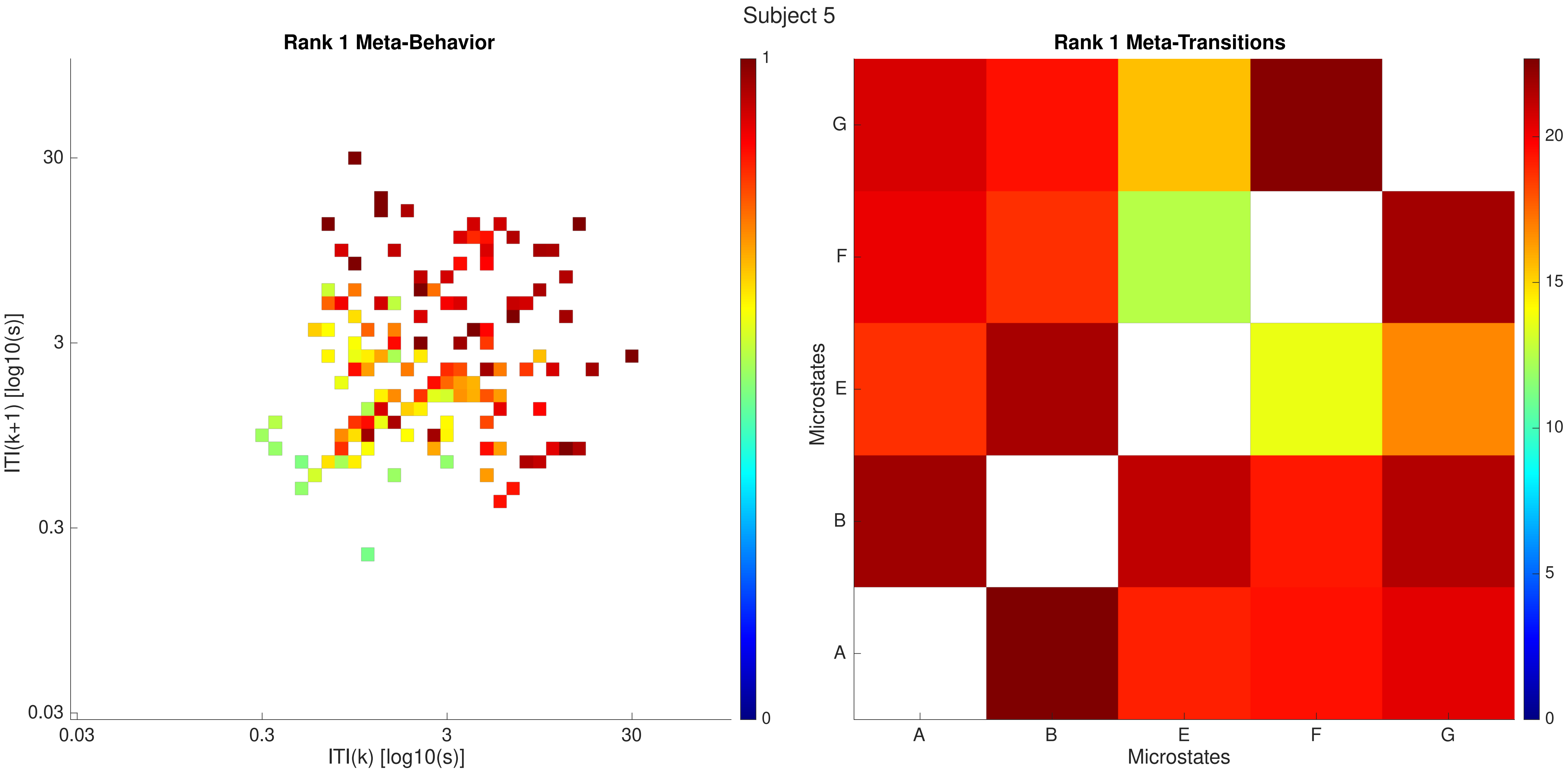

Subject 5
Rank 1 Meta-Behavior
ITI(k+1) [log10(s)]
ITI(k) [log10(s)]
0.03
0.3
3
30
0
1
Rank 1 Meta-Transitions
Microstates
A
B
E
F
G
0
5
10
15
20

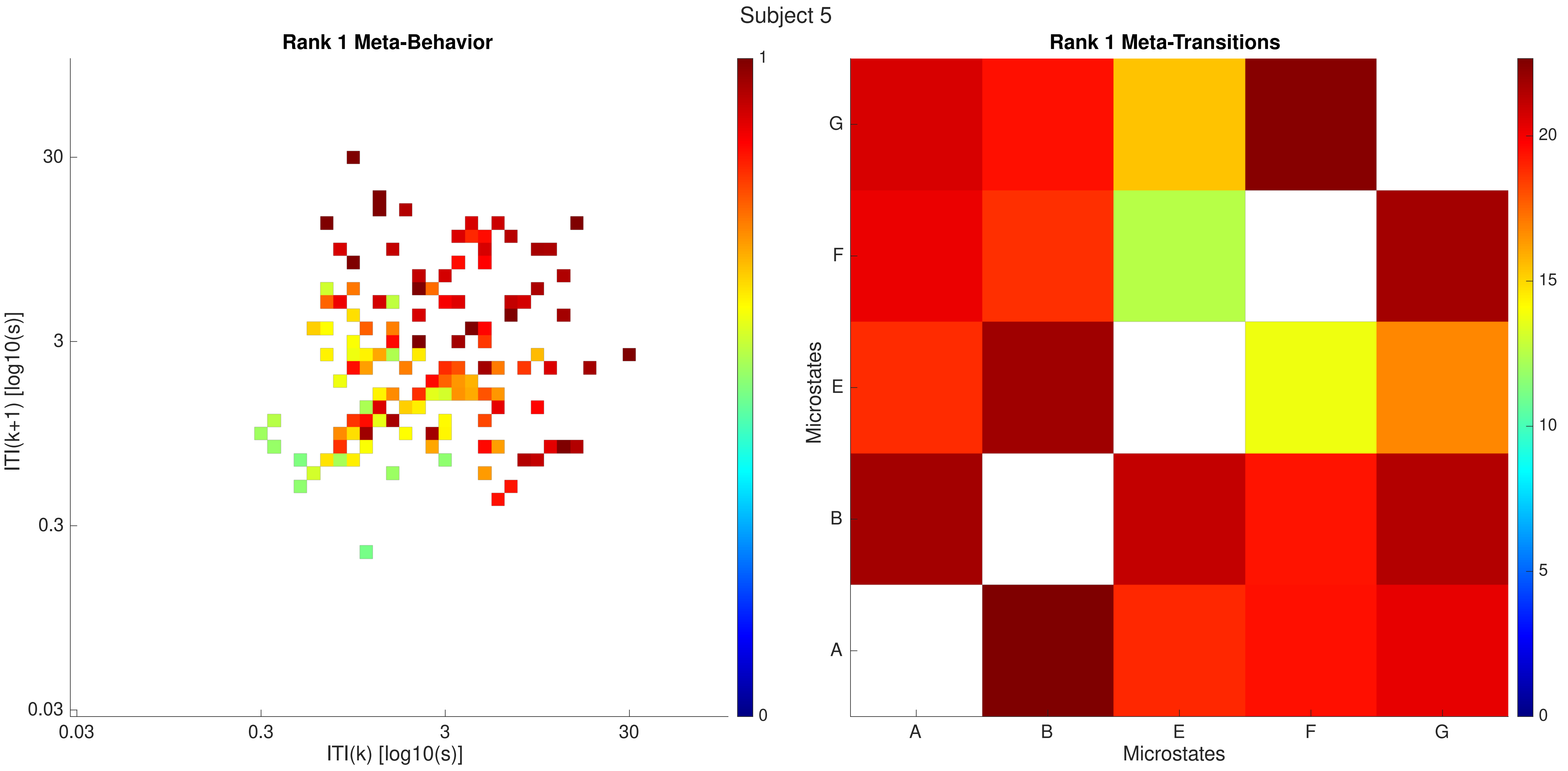

Subject 5
Rank 1 Meta-Behavior
ITI(k+1) [log10(s)]
ITI(k) [log10(s)]
0.03
0.3
3
30
1
0
Rank 1 Meta-Transitions
Microstates
A
B
E
F
G
0
5
10
15
20

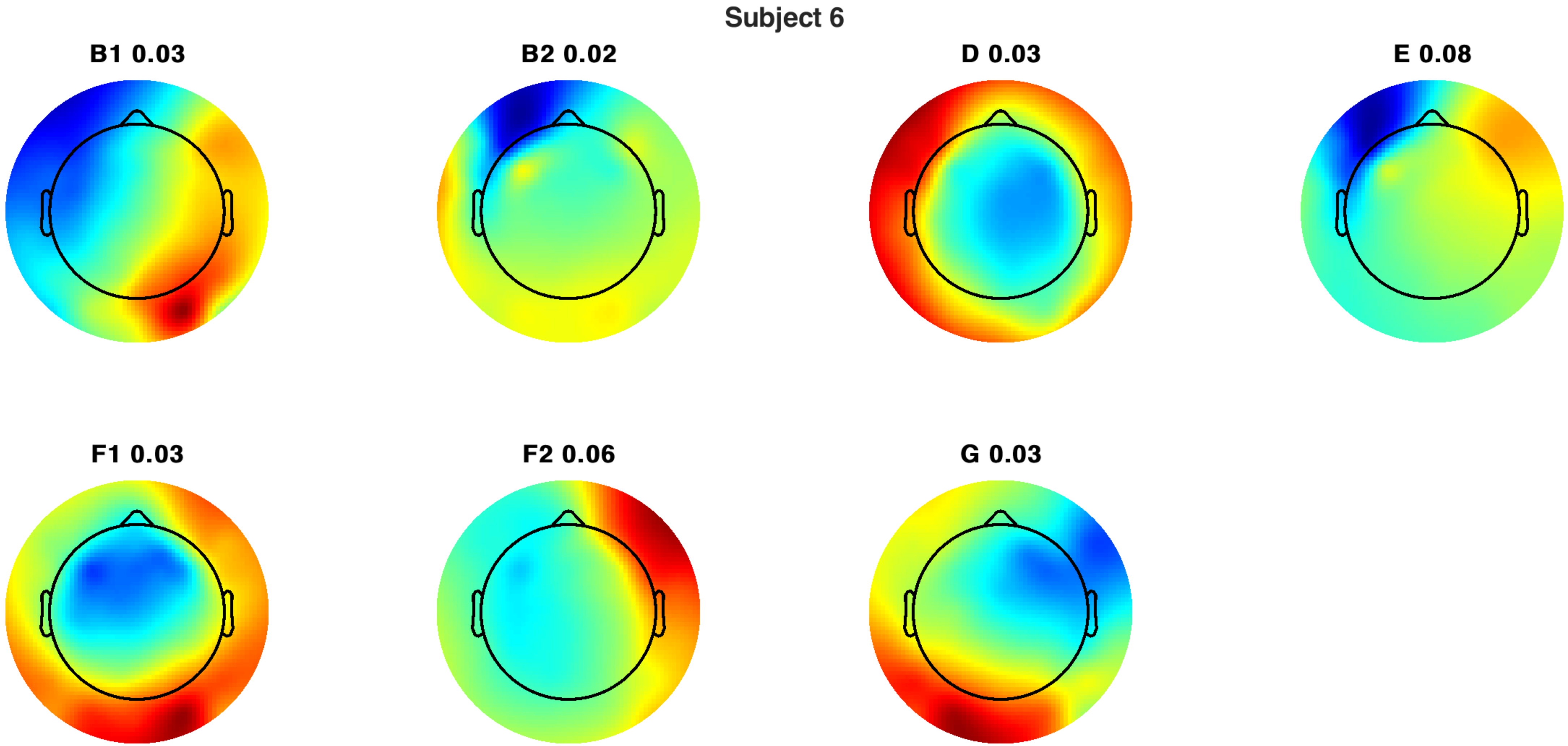
Subject 6
B1 0.03
B2 0.02
D 0.03
E 0.08
F1 0.03
F2 0.06
G 0.03

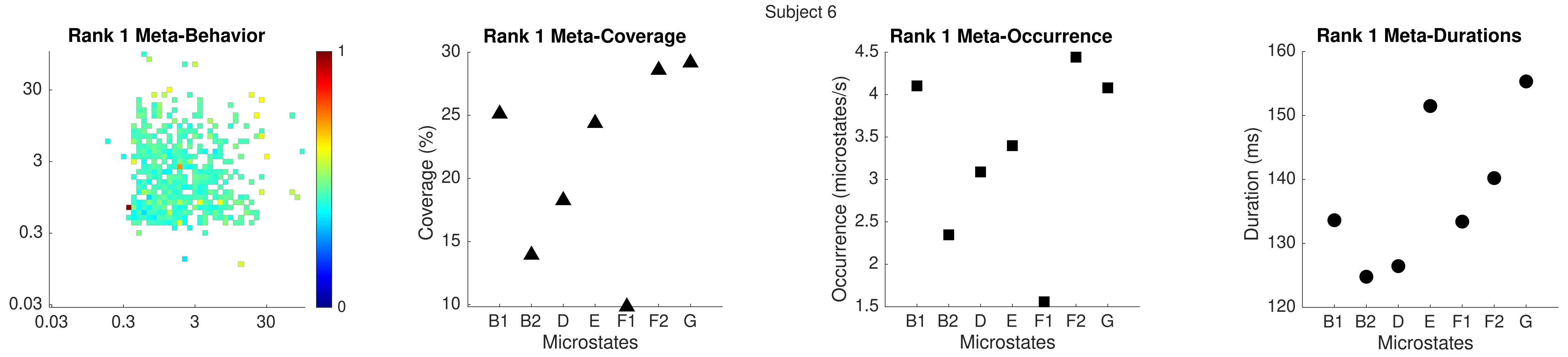
Subject 6
Rank 1 Meta-Behavior
Rank 1 Meta-Coverage
Rank 1 Meta-Occurrence
Rank 1 Meta-Durations
Coverage (%)
Occurrence (microstates/s)
Duration (ms)
Microstates
B1 B2 D E F1 F2 G

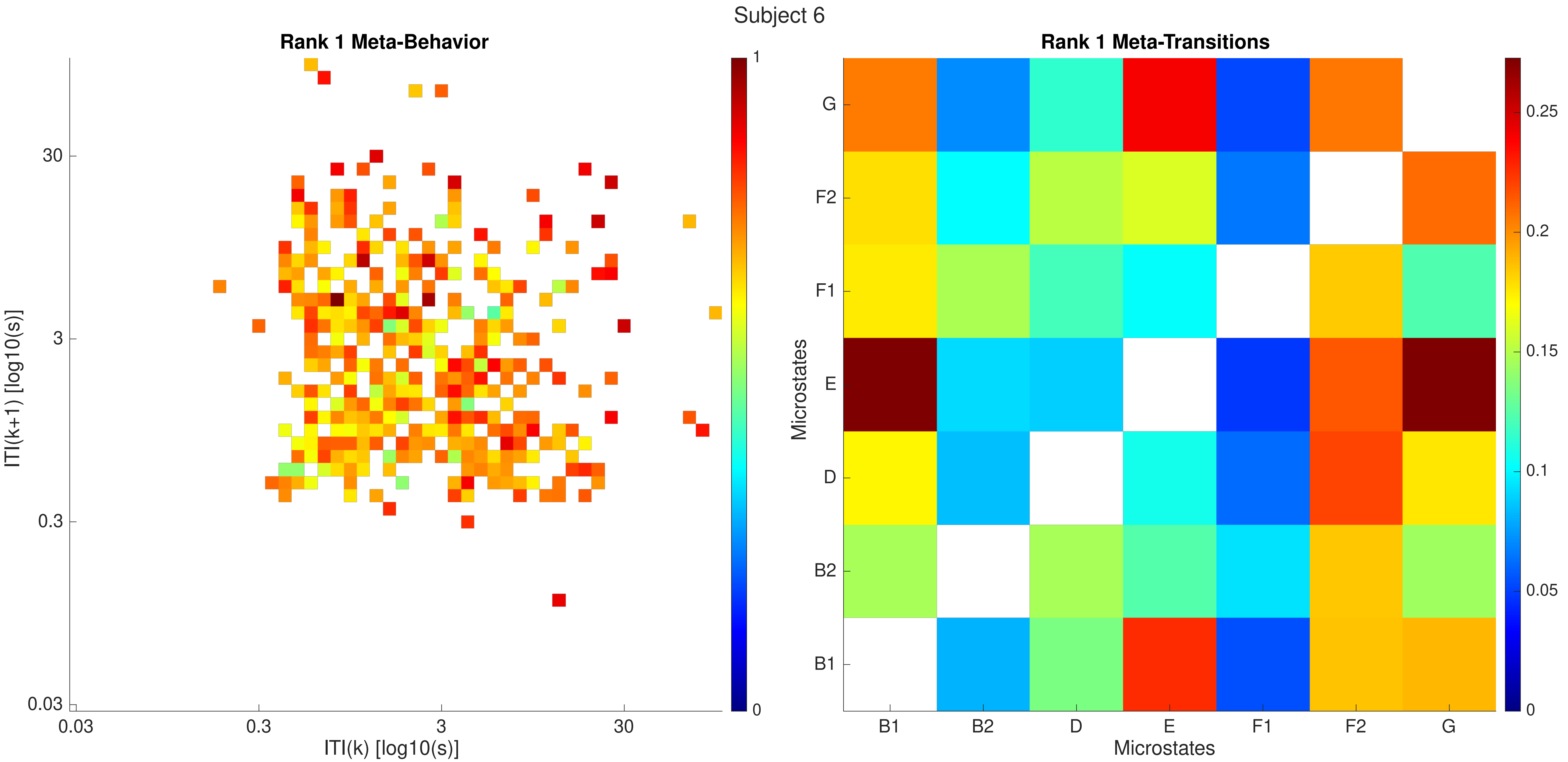
Subject 6
Rank 1 Meta-Behavior
ITI(k+1) [log10(s)]
30
3
0.3
0.03
0.03
0.3
3
30
ITI(k) [log10(s)]
1
0
Rank 1 Meta-Transitions
Microstates
G
F2
F1
E
D
B2
B1
B1
B2
D
E
F1
F2
G
Microstates
0.25
0.2
0.15
0.1
0.05
0

Subject 6

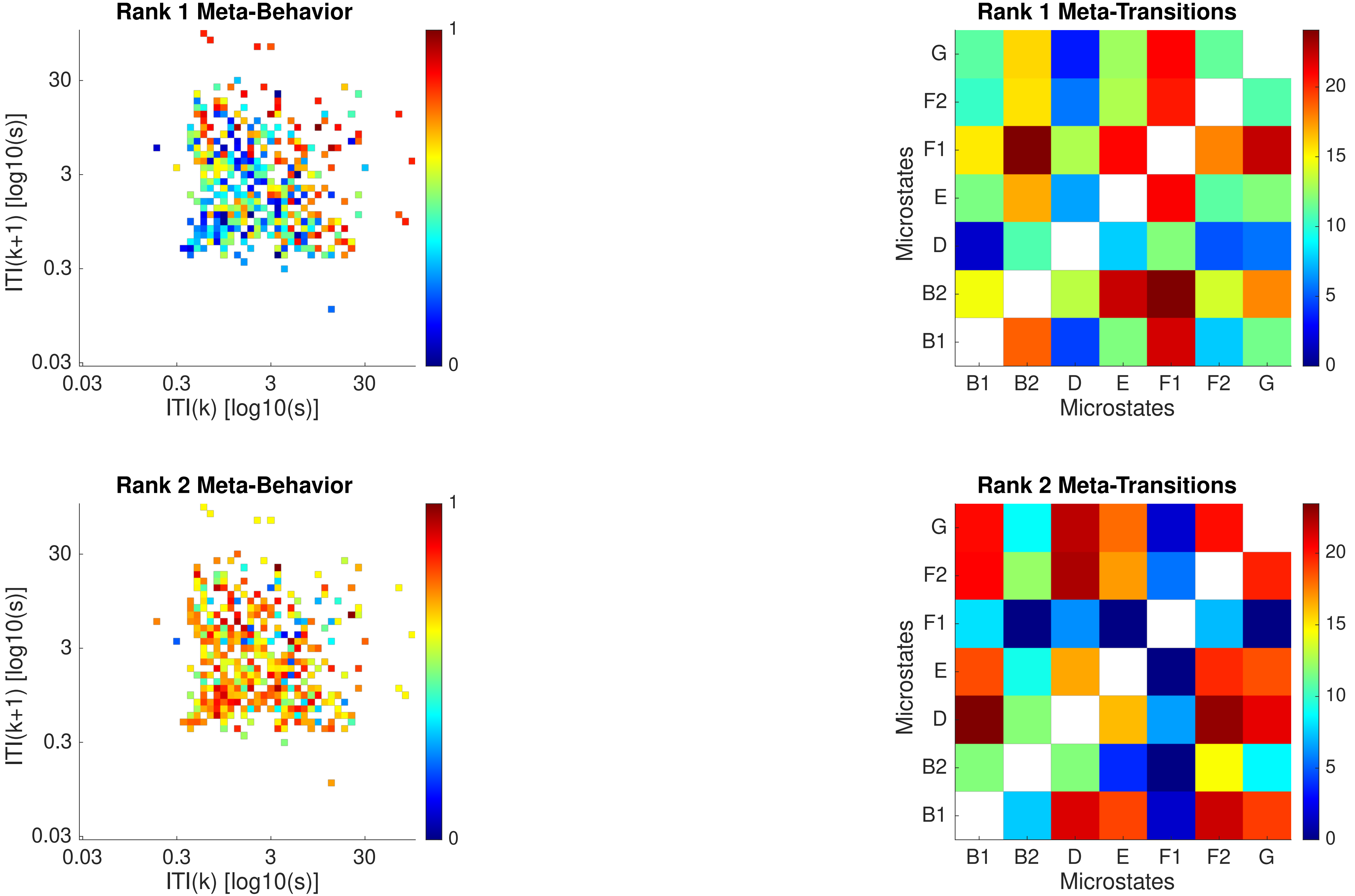
Rank 1 Meta-Behavior
ITI(k+1) [log10(s)]
ITI(k) [log10(s)]
0.03
0.3
3
30
0
1
Rank 1 Meta-Transitions
Microstates
G
F2
F1
E
D
B2
B1
0
5
10
15
20
Rank 2 Meta-Behavior
Rank 2 Meta-Transitions

Subject 6

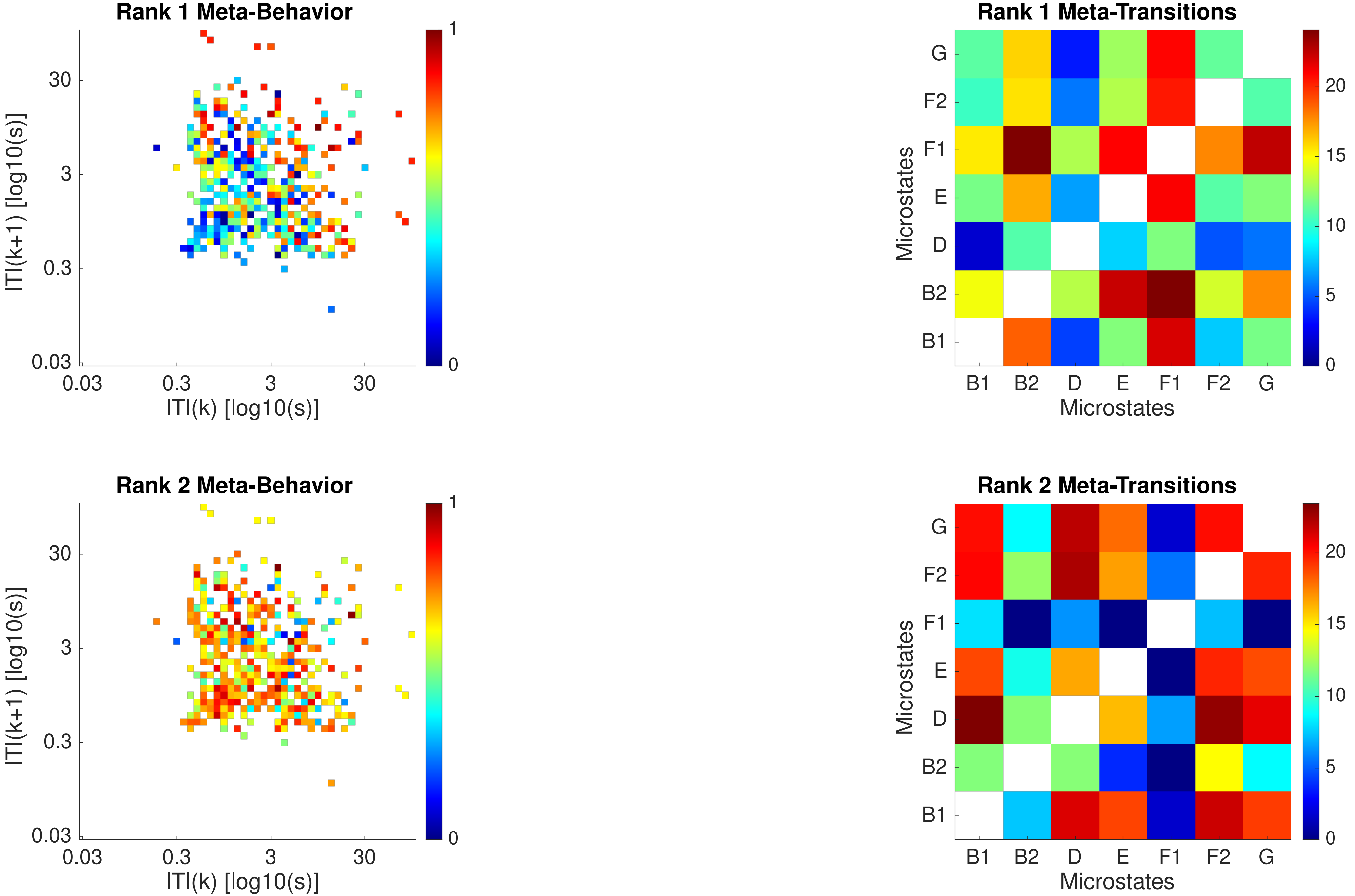

Rank 1 Meta-Behavior
ITI(k+1) [log10(s)]
ITI(k) [log10(s)]
0.03
0.3
3
30
0
1
Rank 1 Meta-Transitions
Microstates
B1
B2
D
E
F1
F2
G
0
5
10
15
20
Rank 2 Meta-Behavior
Rank 2 Meta-Transitions

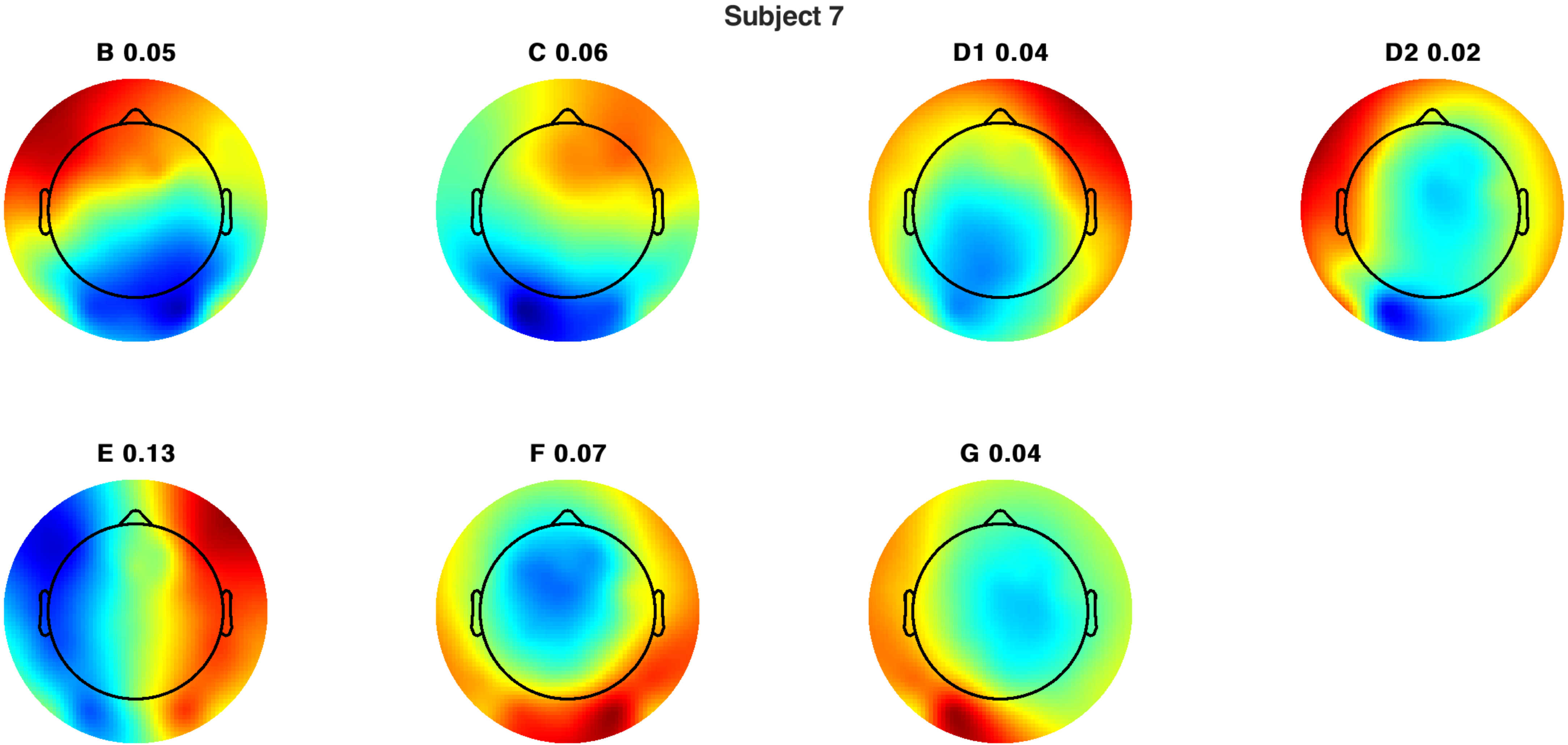
Subject 7
B 0.05
C 0.06
D1 0.04
D2 0.02
E 0.13
F 0.07
G 0.04

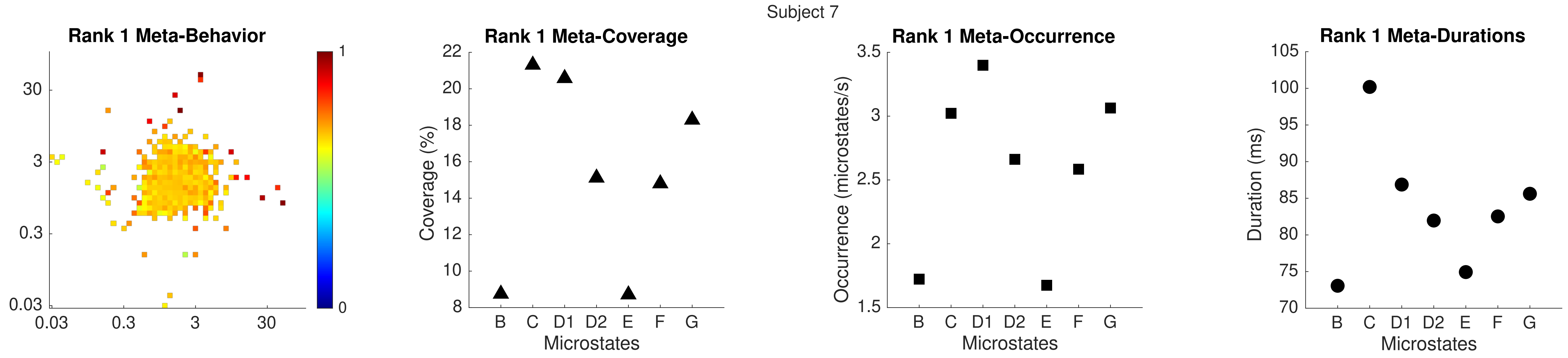
Subject 7
Rank 1 Meta-Behavior
Rank 1 Meta-Coverage
Rank 1 Meta-Occurrence
Rank 1 Meta-Durations
Coverage (%)
Occurrence (microstates/s)
Duration (ms)
Microstates
B C D1 D2 E F G

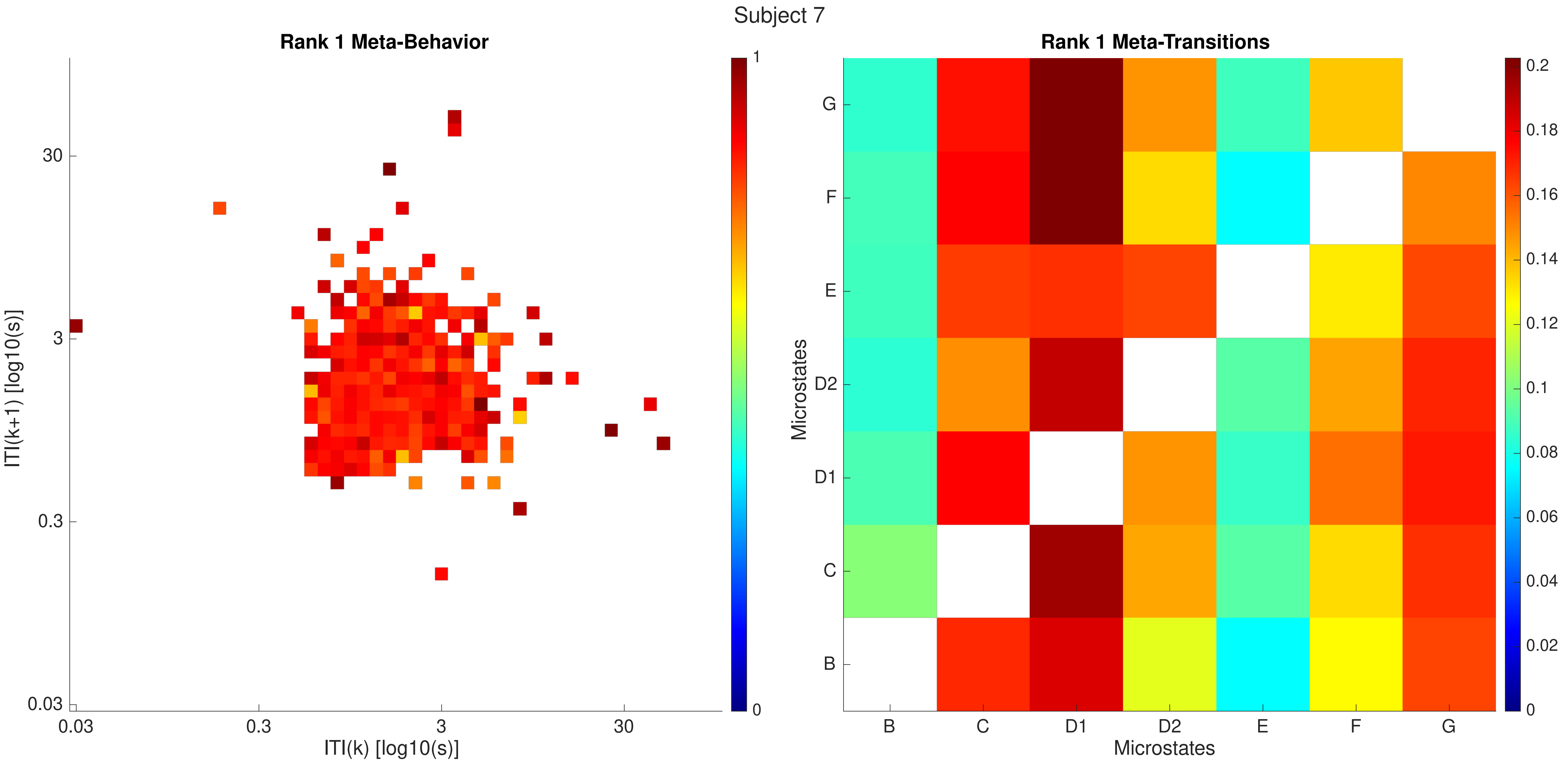
Subject 7
Rank 1 Meta-Behavior
ITI(k+1) [log10(s)]
ITI(k) [log10(s)]
30
3
0.3
0.03
1
0
Rank 1 Meta-Transitions
Microstates
G
F
E
D2
D1
C
B
0.2
0.18
0.16
0.14
0.12
0.1
0.08
0.06
0.04
0.02

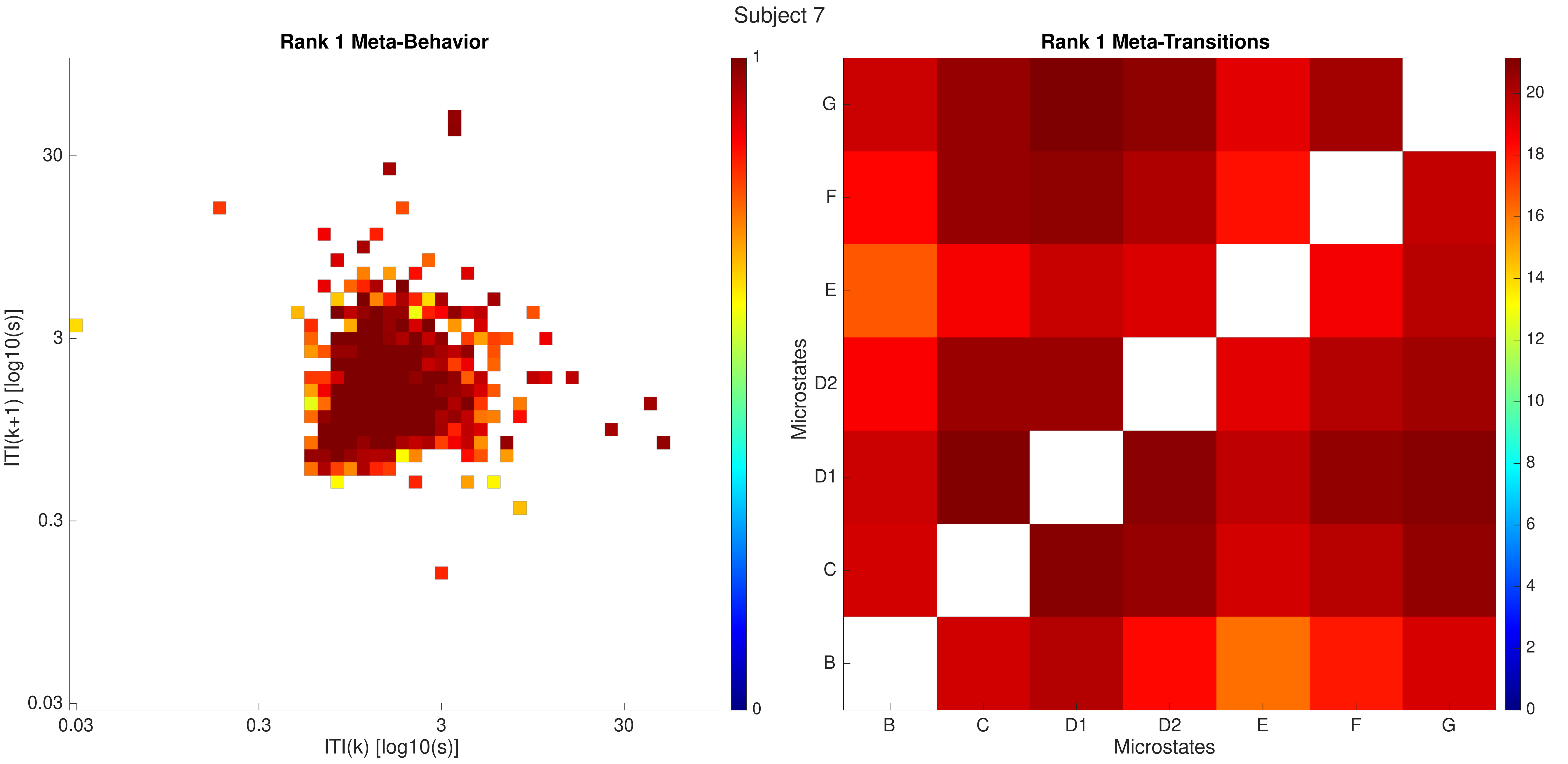

Subject 7
Rank 1 Meta-Behavior
ITI(k+1) [log10(s)]
ITI(k) [log10(s)]
30
3
0.3
0.03
0.03
0.3
3
30
1
0
Rank 1 Meta-Transitions
Microstates
G
F
E
D2
D1
C
B
B
C
D1
D2
E
F
G
Microstates
20
18
16
14
12
10
8
6
4
2
0

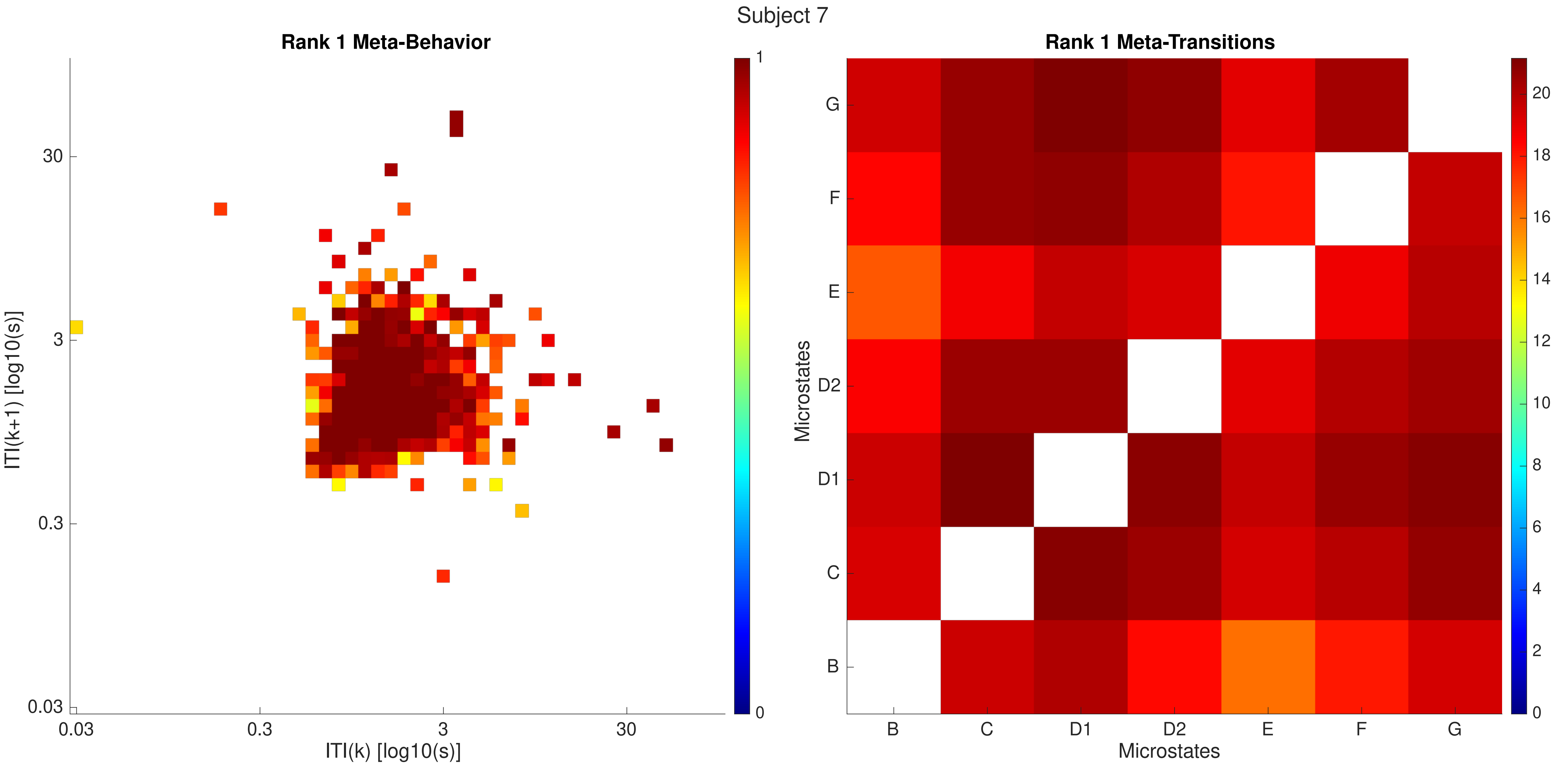
Subject 7
Rank 1 Meta-Behavior
ITI(k+1) [log10(s)]
ITI(k) [log10(s)]
0.03
0.3
3
30
0
1
Rank 1 Meta-Transitions
Microstates
Microstates
B
C
D1
D2
E
F
G
0
2
4
6
8
10
12
14
16
18
20

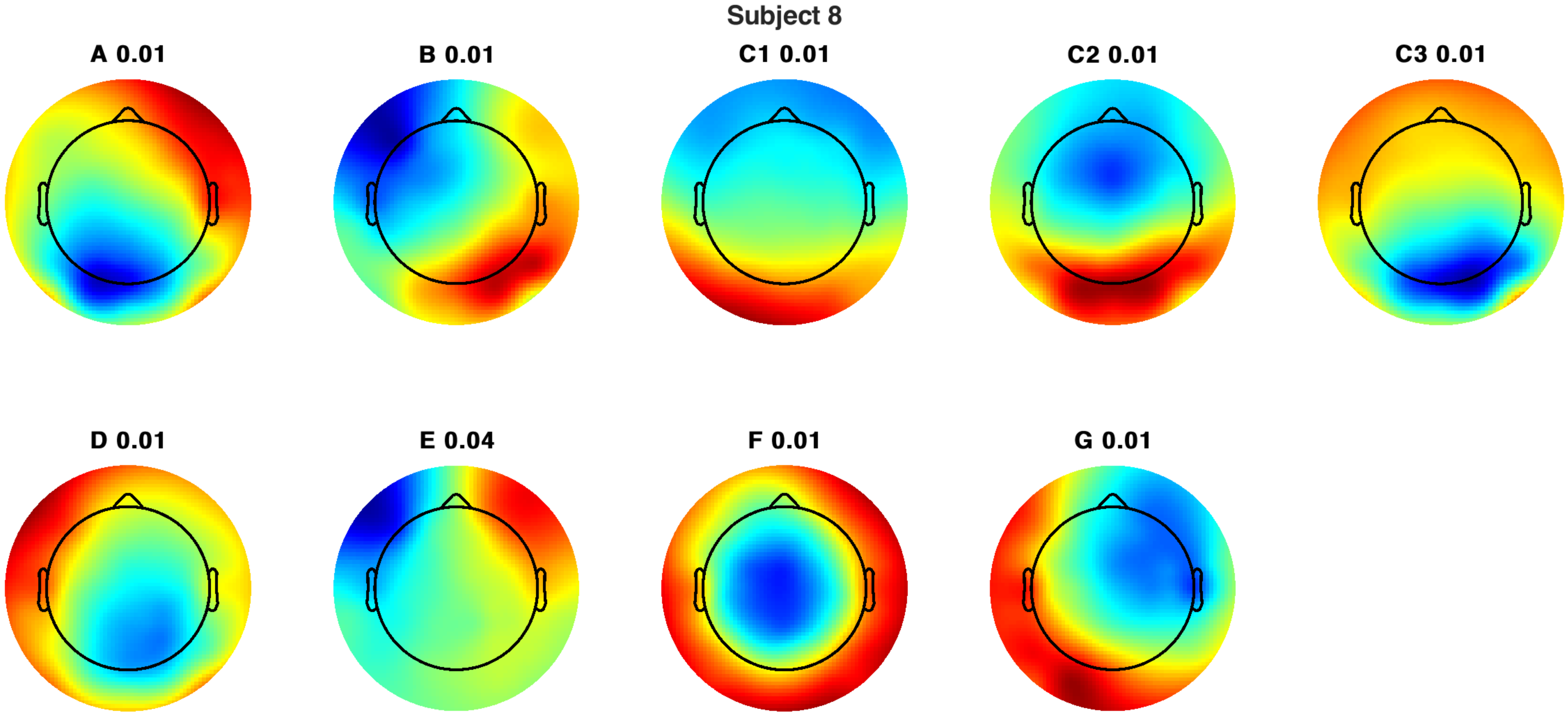
Subject 8
A 0.01
B 0.01
C1 0.01
C2 0.01
C3 0.01
D 0.01
E 0.04
F 0.01
G 0.01

Subject 8

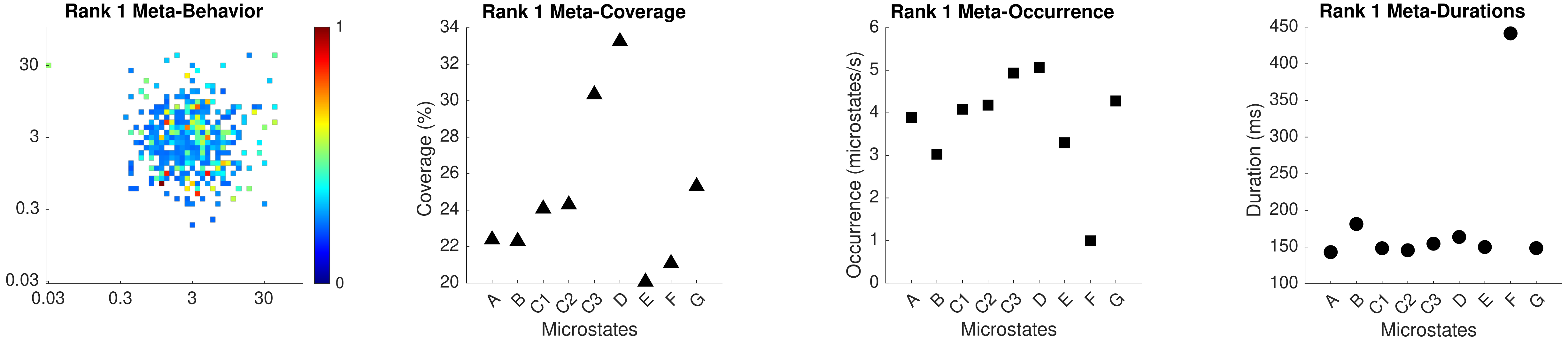

Rank 1 Meta-Behavior
Rank 1 Meta-Coverage
Coverage (%)
Microstates
Rank 1 Meta-Occurrence
Occurrence (microstates/s)
Microstates
Rank 1 Meta-Durations
Duration (ms)
Microstates
A B C1 C2 C3 D E F G

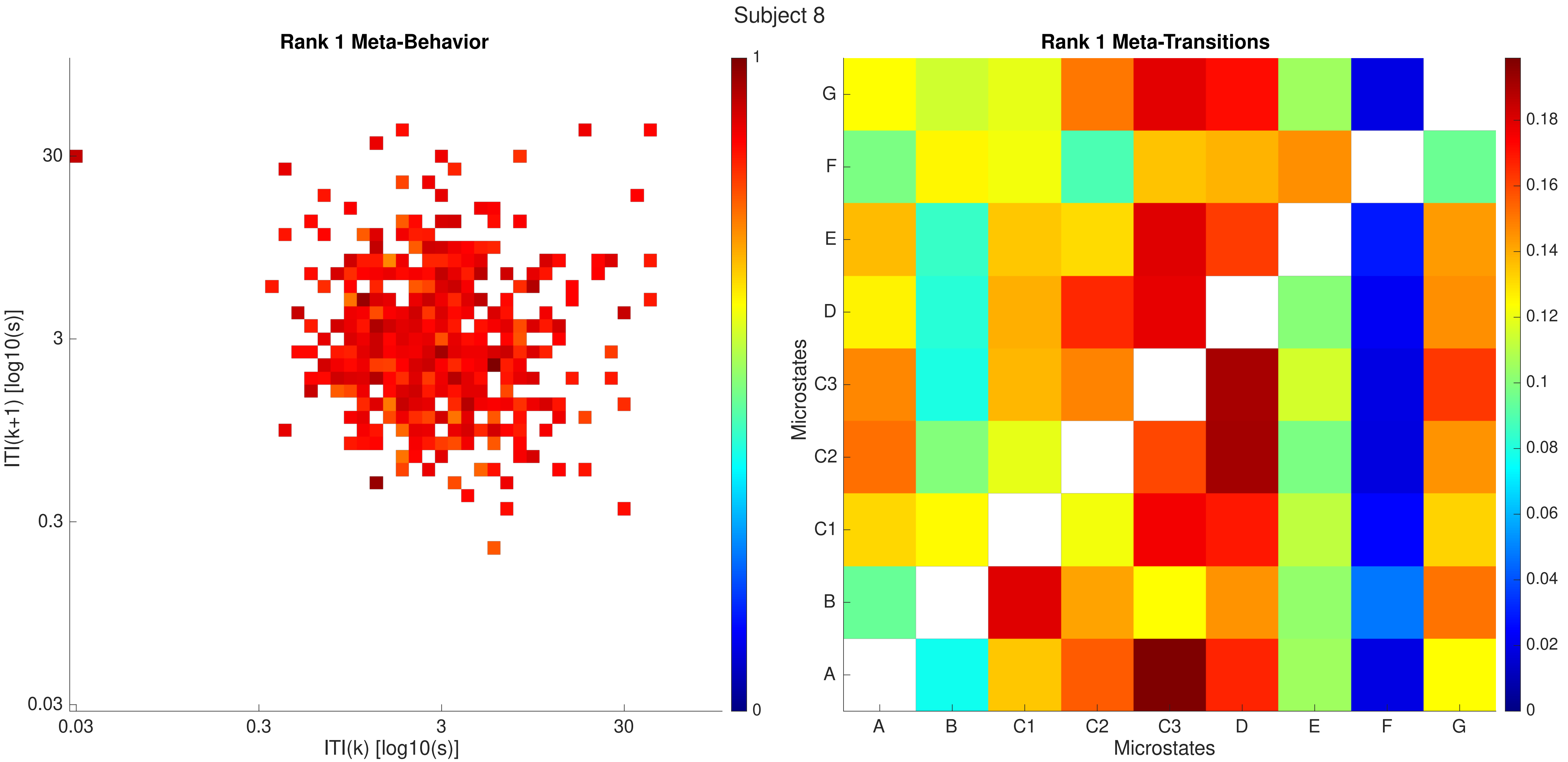

Subject 8
Rank 1 Meta-Behavior
ITI(k+1) [log10(s)]
30
3
0.3
0.03
0.03
0.3
3
30
ITI(k) [log10(s)]
1
0
Rank 1 Meta-Transitions
Microstates
G
F
E
D
C3
C2
C1
B
A
A
B
C1
C2
C3
D
E
F
G
Microstates
0.18
0.16
0.14
0.12
0.1
0.08
0.06
0.04
0.02
0

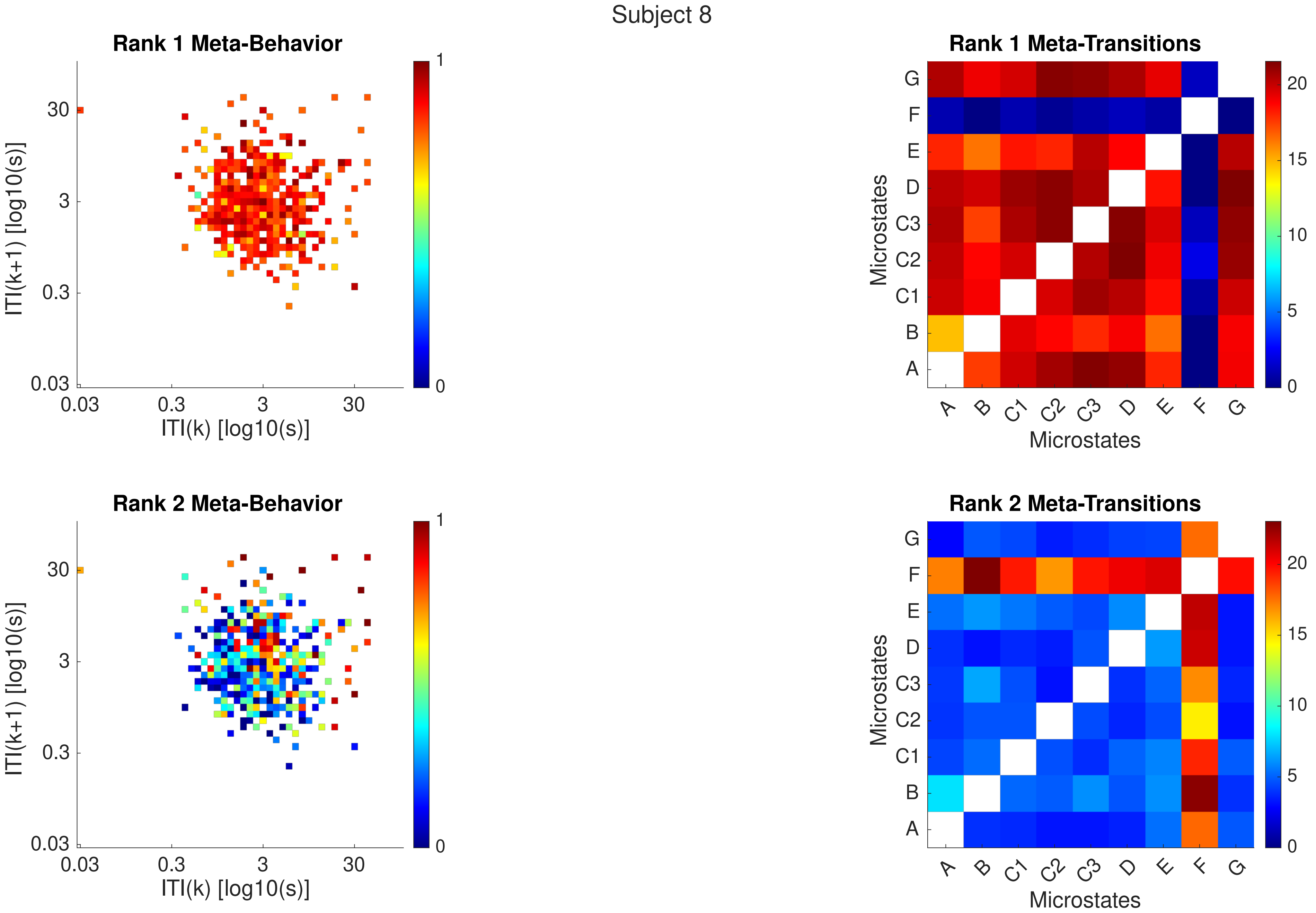

Subject 8
Rank 1 Meta-Behavior
ITI(k+1) [log10(s)]
ITI(k) [log10(s)]
0.03
0.3
3
30
0
1
Rank 1 Meta-Transitions
Microstates
A
B
C1
C2
C3
D
E
F
G
0
5
10
15
20
Rank 2 Meta-Behavior
ITI(k+1) [log10(s)]
ITI(k) [log10(s)]
Rank 2 Meta-Transitions
Microstates

Subject 8

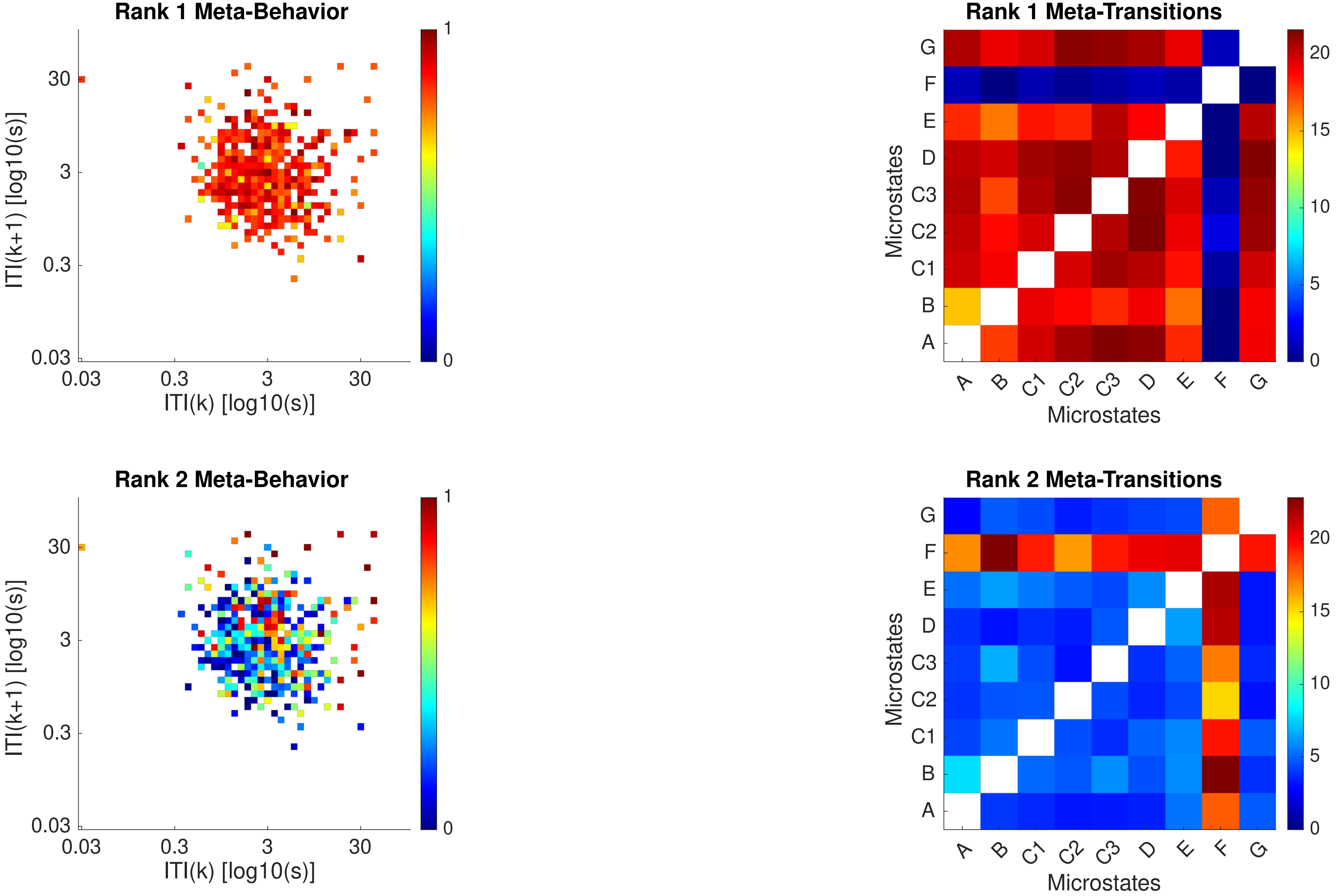

Rank 1 Meta-Behavior
Rank 1 Meta-Transitions
Rank 2 Meta-Behavior
Rank 2 Meta-Transitions
ITI(k+1) [log10(s)]
ITI(k) [log10(s)]
0.03
0.3
3
30
0
1
Microstates
A
B
C1
C2
C3
D
E
F
G
0
5
10
15
20

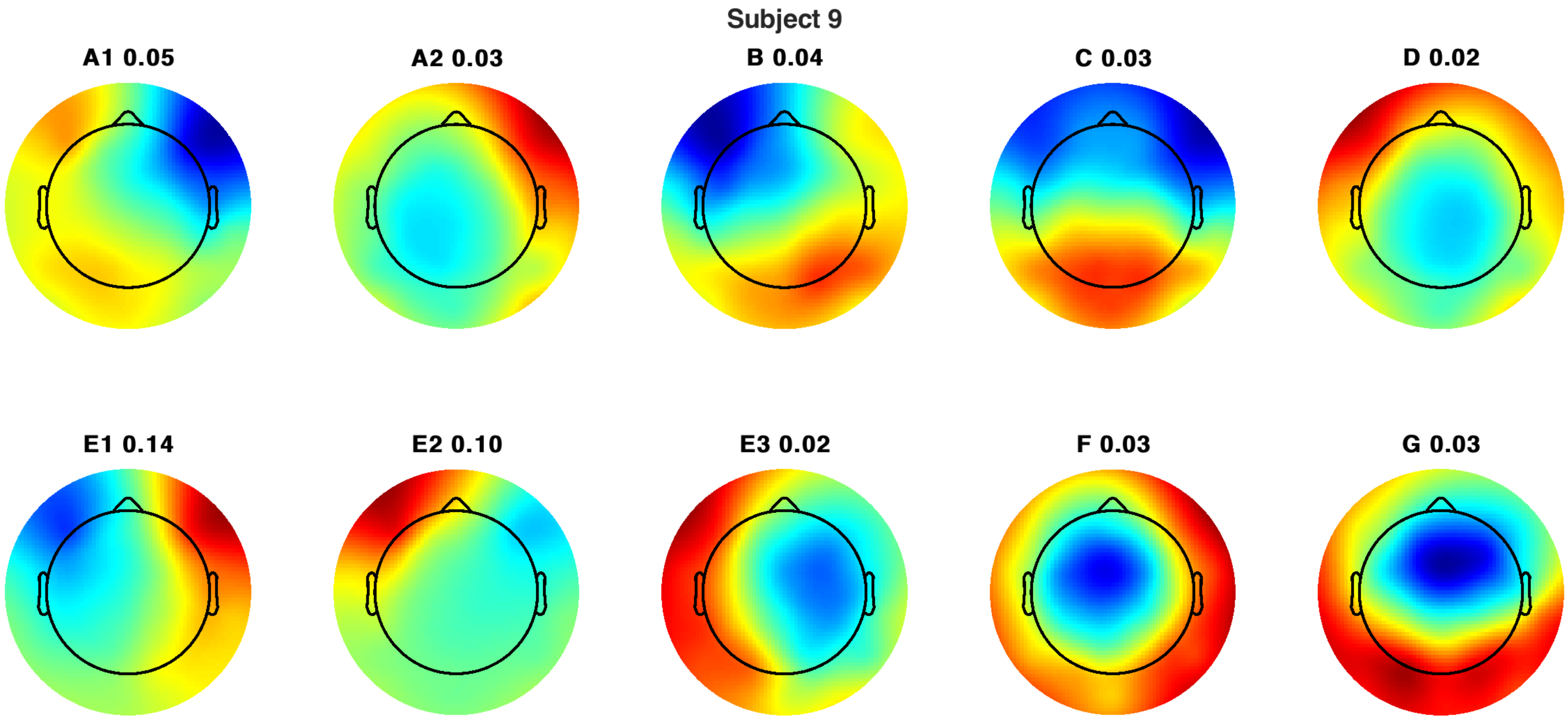
Subject 9
A1 0.05
A2 0.03
B 0.04
C 0.03
D 0.02
E1 0.14
E2 0.10
E3 0.02
F 0.03
G 0.03

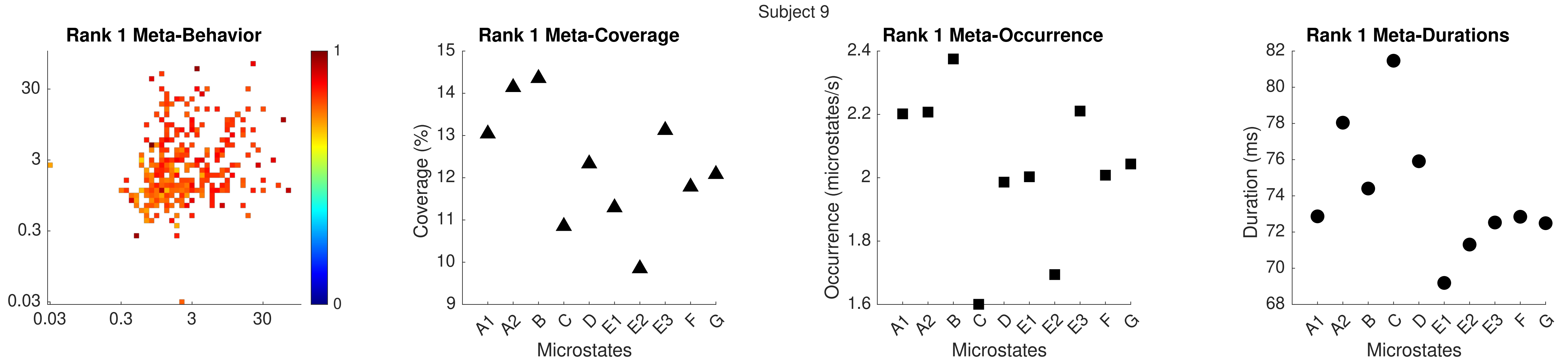

Subject 9
Rank 1 Meta-Behavior
Rank 1 Meta-Coverage
Rank 1 Meta-Occurrence
Rank 1 Meta-Durations
Coverage (%)
Occurrence (microstates/s)
Duration (ms)
Microstates
A1 A2 B C D E1 E2 E3 F G

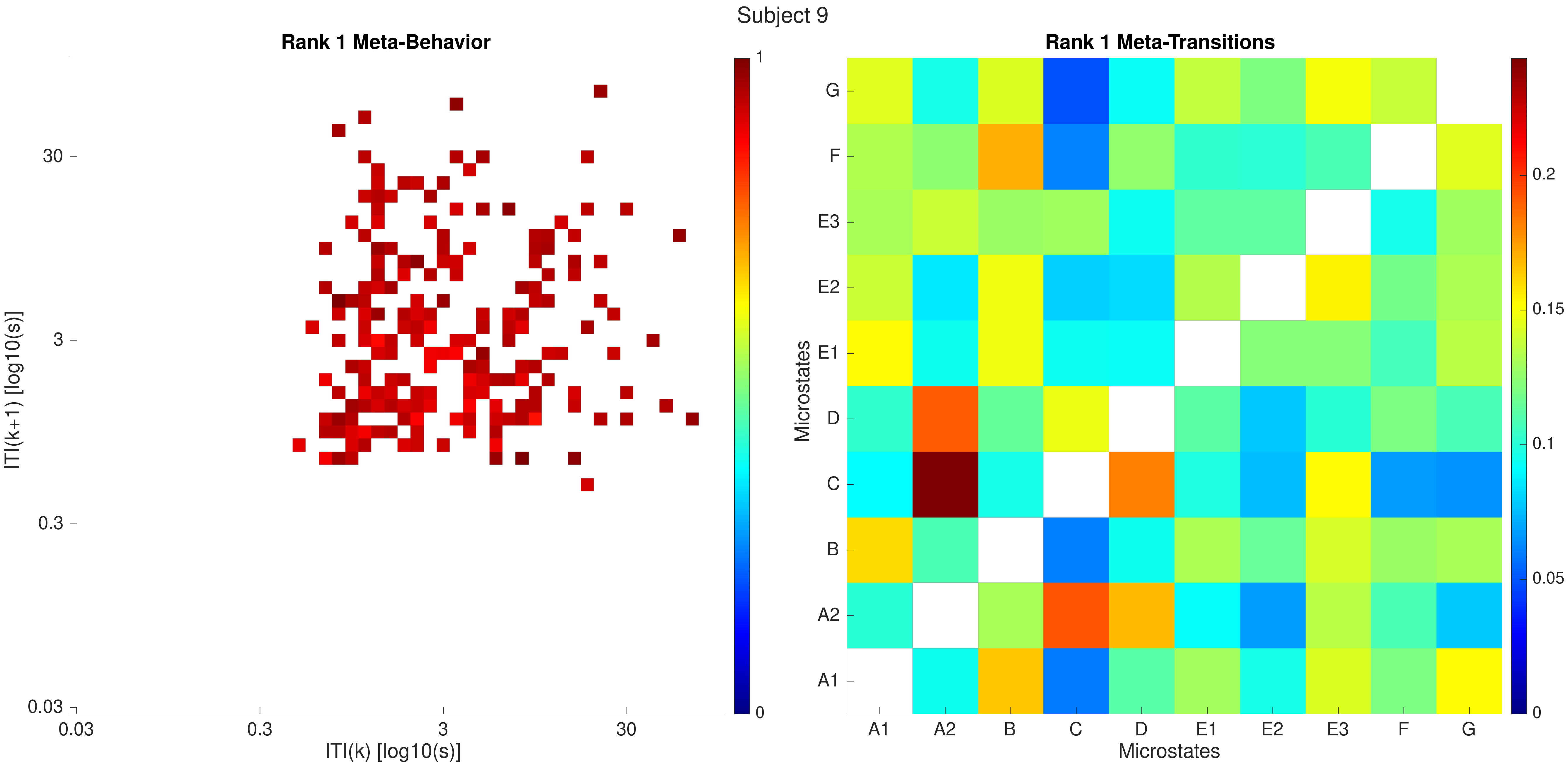

Subject 9
Rank 1 Meta-Behavior
ITI(k+1) [log10(s)]
ITI(k) [log10(s)]
0.03
0.3
3
30
0
1
Rank 1 Meta-Transitions
Microstates
A1
A2
B
C
D
E1
E2
E3
F
G
0
0.05
0.1
0.15
0.2

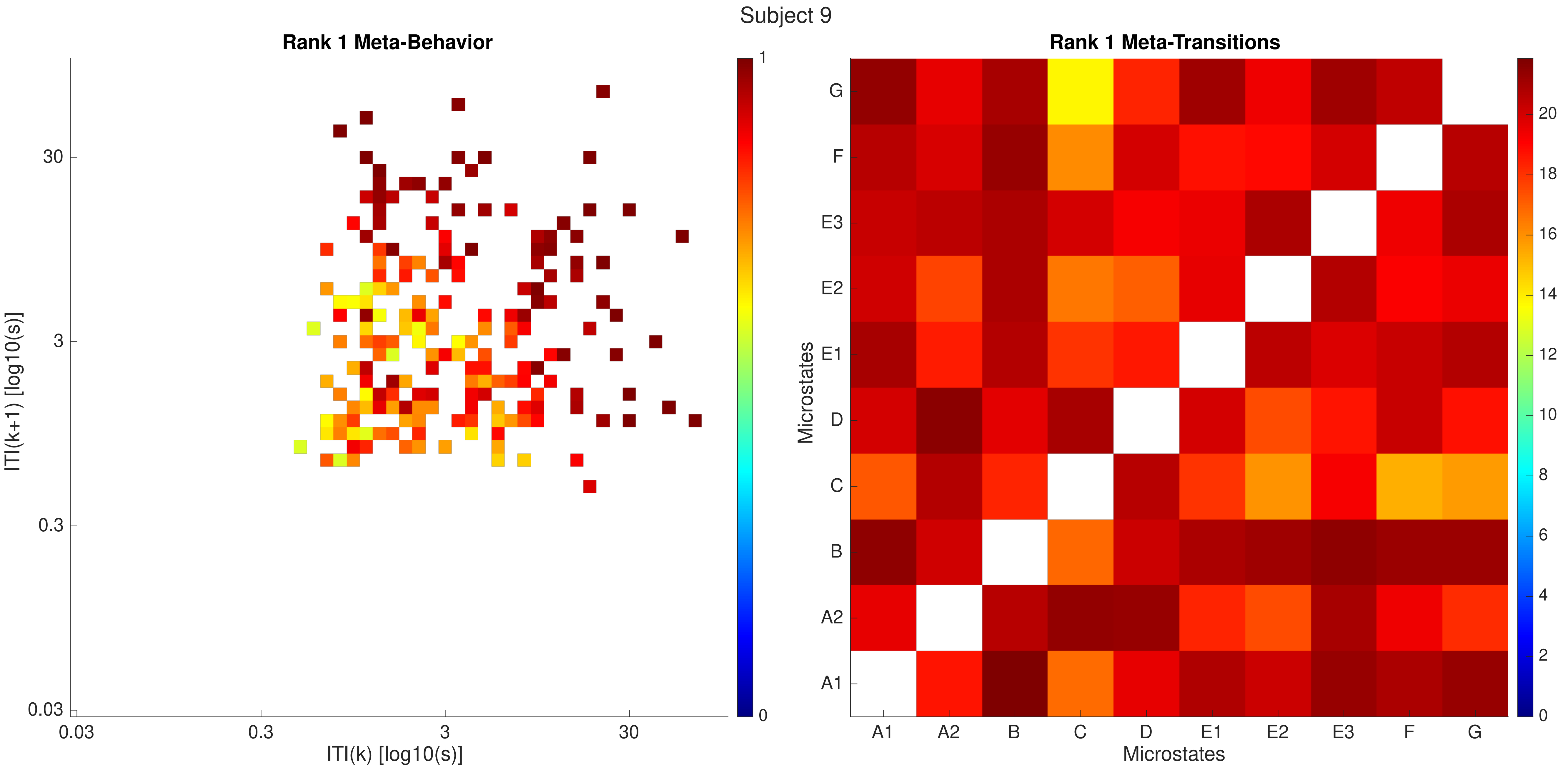
Subject 9
Rank 1 Meta-Behavior
ITI(k+1) [log10(s)]
ITI(k) [log10(s)]
0.03
0.3
3
30
0
1
Rank 1 Meta-Transitions
Microstates
A1
A2
B
C
D
E1
E2
E3
F
G
0
2
4
6
8
10
12
14
16
18
20

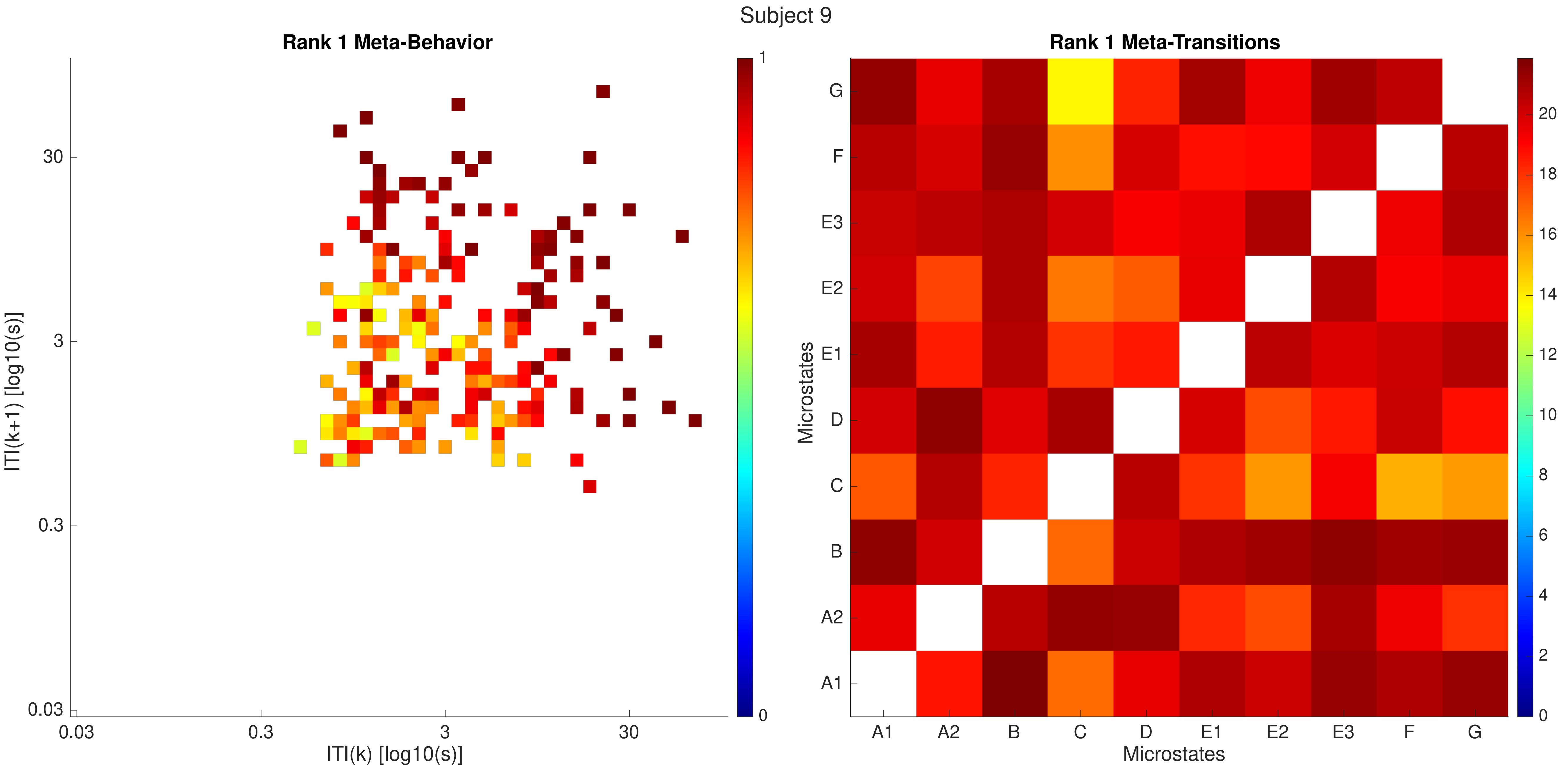

Subject 9
Rank 1 Meta-Behavior
ITI(k+1) [log10(s)]
ITI(k) [log10(s)]
0.03
0.3
3
30
1
0
Rank 1 Meta-Transitions
Microstates
A1
A2
B
C
D
E1
E2
E3
F
G
0
2
4
6
8
10
12
14
16
18
20

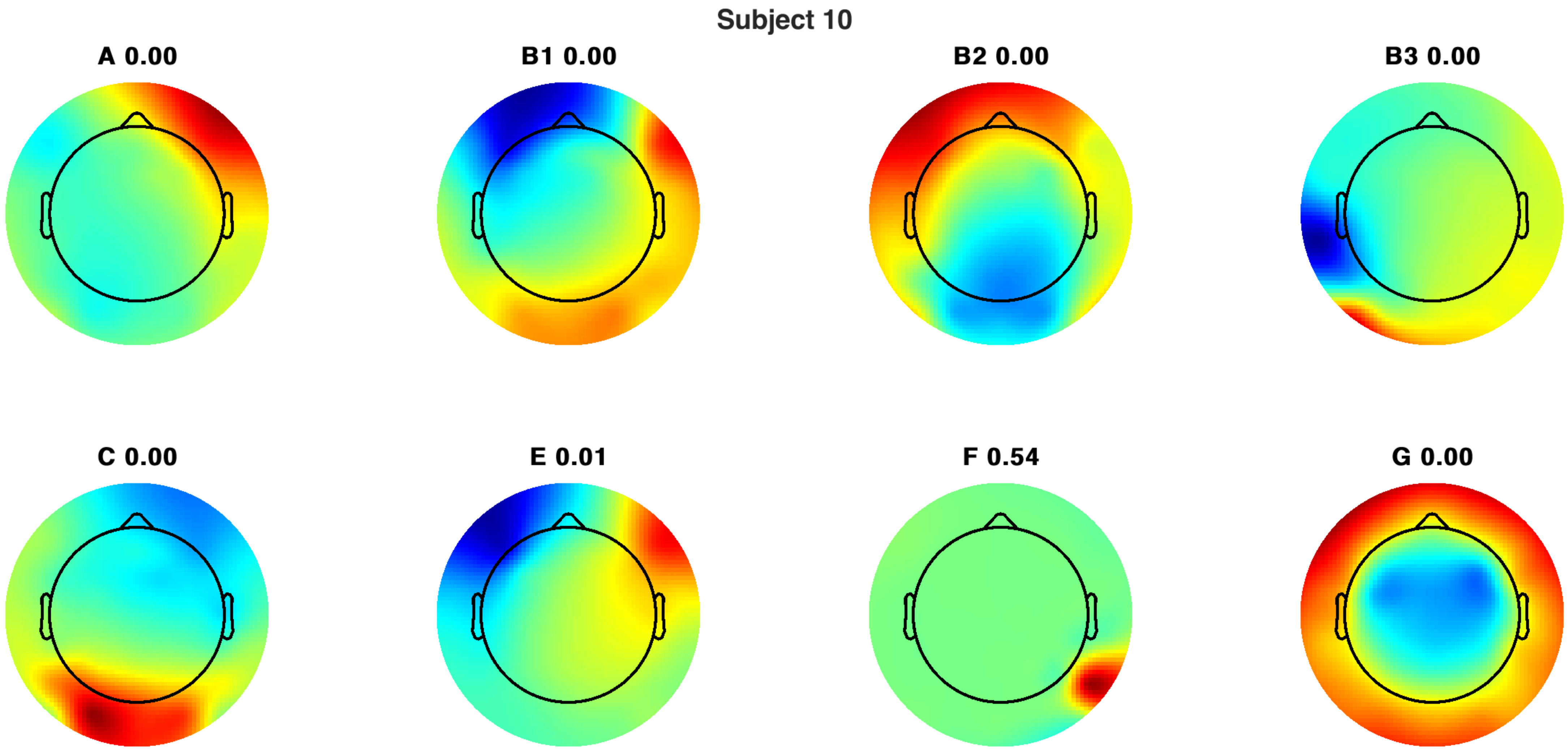
Subject 10
A 0.00
B1 0.00
B2 0.00
B3 0.00
C 0.00
E 0.01
F 0.54
G 0.00

### Subject 10

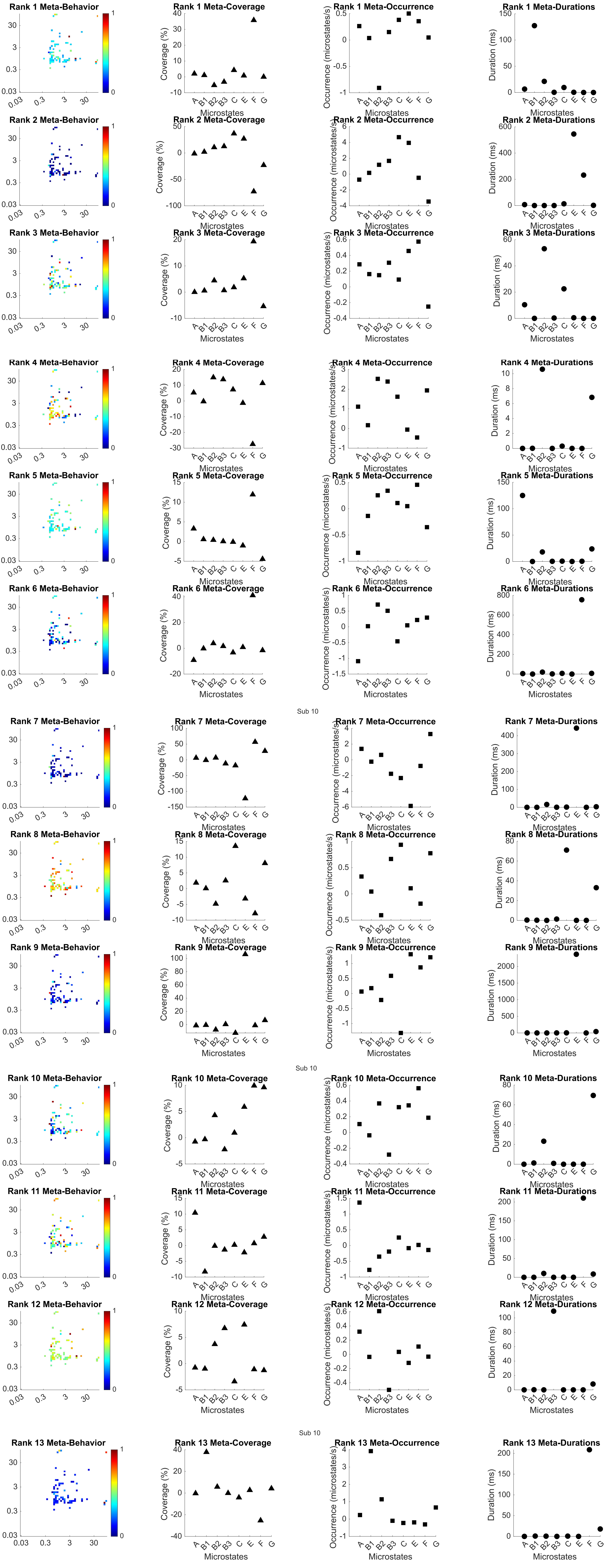
Rank 1 Meta-Behavior
Rank 1 Meta-Coverage
Rank 1 Meta-Occurrence
Rank 1 Meta-Durations
Rank 2 Meta-Behavior
Rank 2 Meta-Coverage
Rank 2 Meta-Occurrence
Rank 2 Meta-Durations
Rank 3 Meta-Behavior
Rank 3 Meta-Coverage
Rank 3 Meta-Occurrence
Rank 3 Meta-Durations
Rank 4 Meta-Behavior
Rank 4 Meta-Coverage
Rank 4 Meta-Occurrence
Rank 4 Meta-Durations
Rank 5 Meta-Behavior
Rank 5 Meta-Coverage
Rank 5 Meta-Occurrence
Rank 5 Meta-Durations
Rank 6 Meta-Behavior
Rank 6 Meta-Coverage
Rank 6 Meta-Occurrence
Rank 6 Meta-Durations
Sub 10
Rank 7 Meta-Behavior
Rank 7 Meta-Coverage
Rank 7 Meta-Occurrence
Rank 7 Meta-Durations
Rank 8 Meta-Behavior
Rank 8 Meta-Coverage
Rank 8 Meta-Occurrence
Rank 8 Meta-Durations
Rank 9 Meta-Behavior
Rank 9 Meta-Coverage
Rank 9 Meta-Occurrence
Rank 9 Meta-Durations
Sub 10
Rank 10 Meta-Behavior
Rank 10 Meta-Coverage
Rank 10 Meta-Occurrence
Rank 10 Meta-Durations
Rank 11 Meta-Behavior
Rank 11 Meta-Coverage
Rank 11 Meta-Occurrence
Rank 11 Meta-Durations
Rank 12 Meta-Behavior
Rank 12 Meta-Coverage
Rank 12 Meta-Occurrence
Rank 12 Meta-Durations
Sub 10
Rank 13 Meta-Behavior
Rank 13 Meta-Coverage
Rank 13 Meta-Occurrence
Rank 13 Meta-Durations
Coverage (%)
Occurrence (microstates/s)
Duration (ms)
Microstates
A B1 B2 B3 C E F G

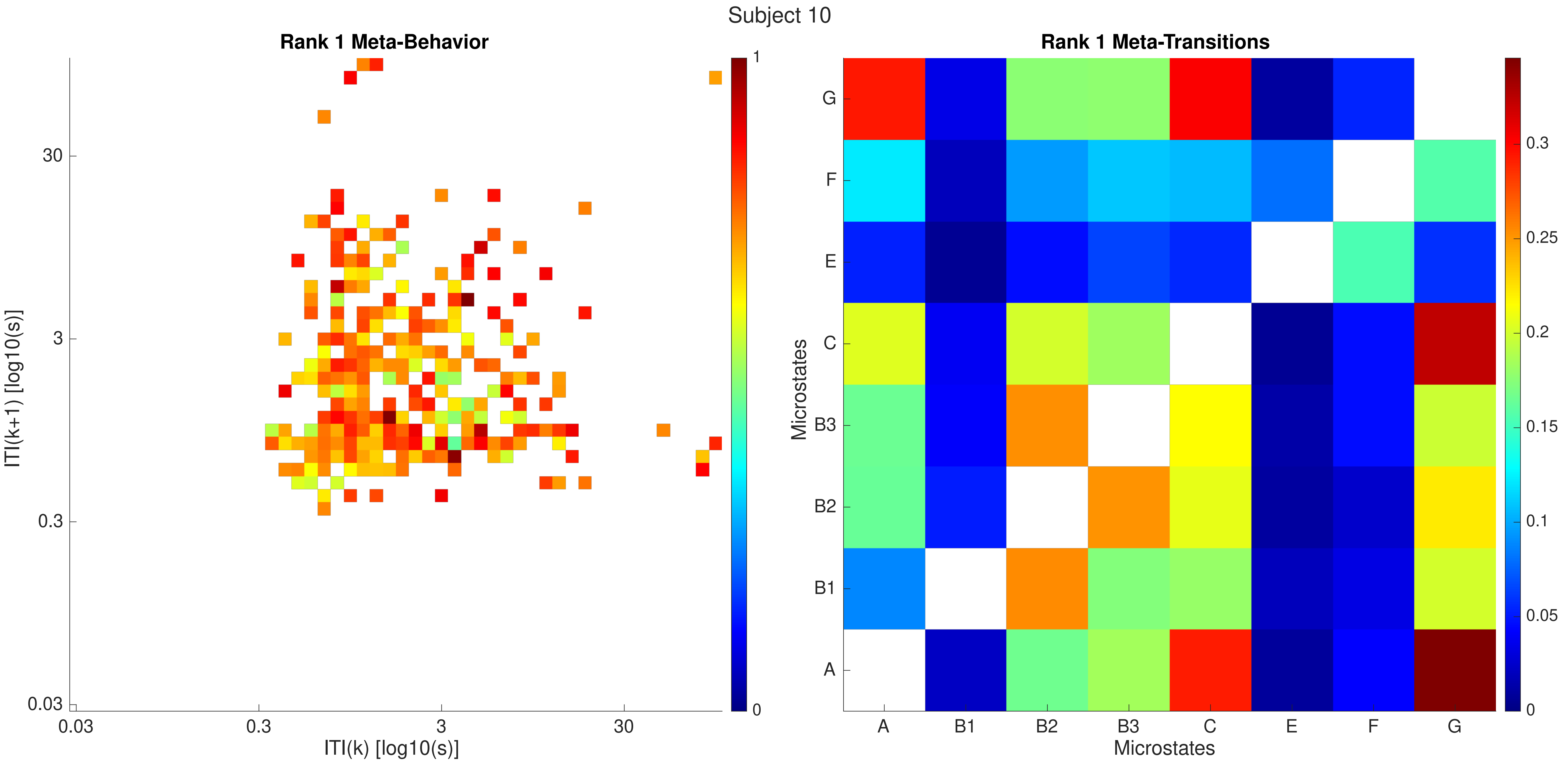
Subject 10
Rank 1 Meta-Behavior
ITI(k+1) [log10(s)]
ITI(k) [log10(s)]
0.03
0.3
3
30
0
1
Rank 1 Meta-Transitions
Microstates
Microstates
A
B1
B2
B3
C
E
F
G
0
0.05
0.1
0.15
0.2
0.25
0.3

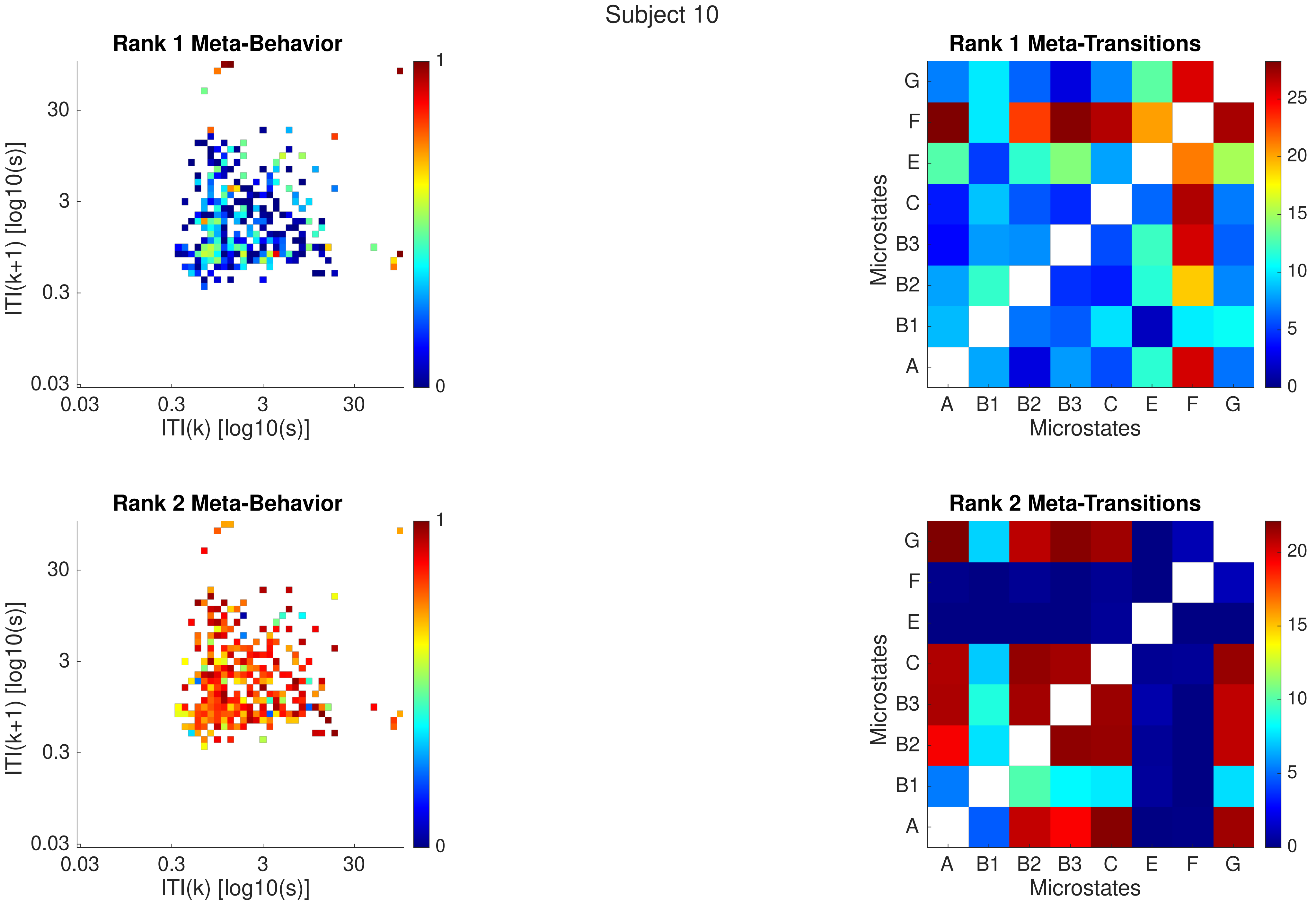
Subject 10
Rank 1 Meta-Behavior
ITI(k+1) [log10(s)]
ITI(k) [log10(s)]
0.03
0.3
3
30
0
1
Rank 1 Meta-Transitions
Microstates
A
B1
B2
B3
C
E
F
G
0
5
10
15
20
25
Rank 2 Meta-Behavior
Rank 2 Meta-Transitions

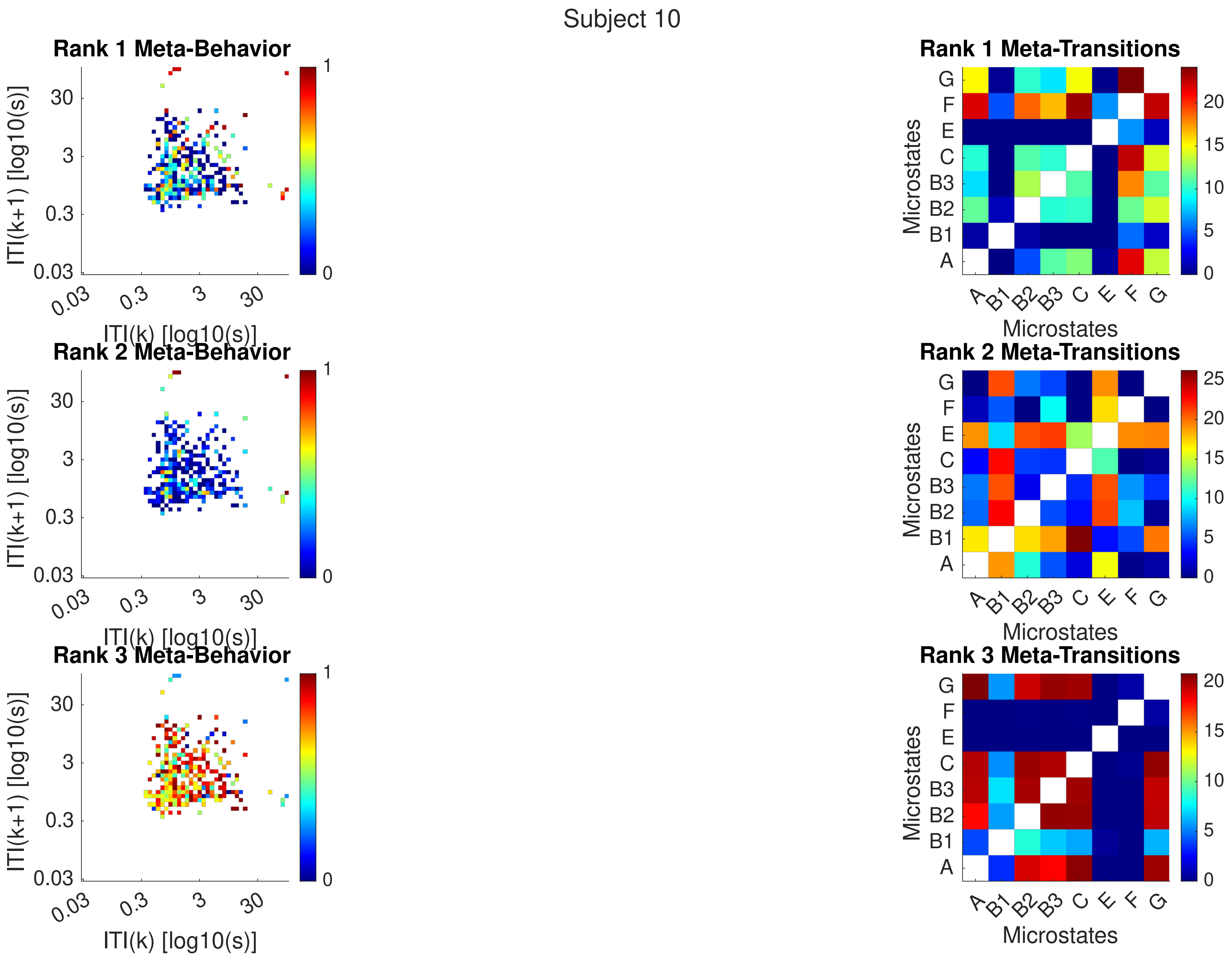
Subject 10
Rank 1 Meta-Behavior
Rank 2 Meta-Behavior
Rank 3 Meta-Behavior
ITI(k+1) [log10(s)]
ITI(k) [log10(s)]
0.03
0.3
3
30
Rank 1 Meta-Transitions
Rank 2 Meta-Transitions
Rank 3 Meta-Transitions
Microstates
A
B1
B2
B3
C
E
F
G

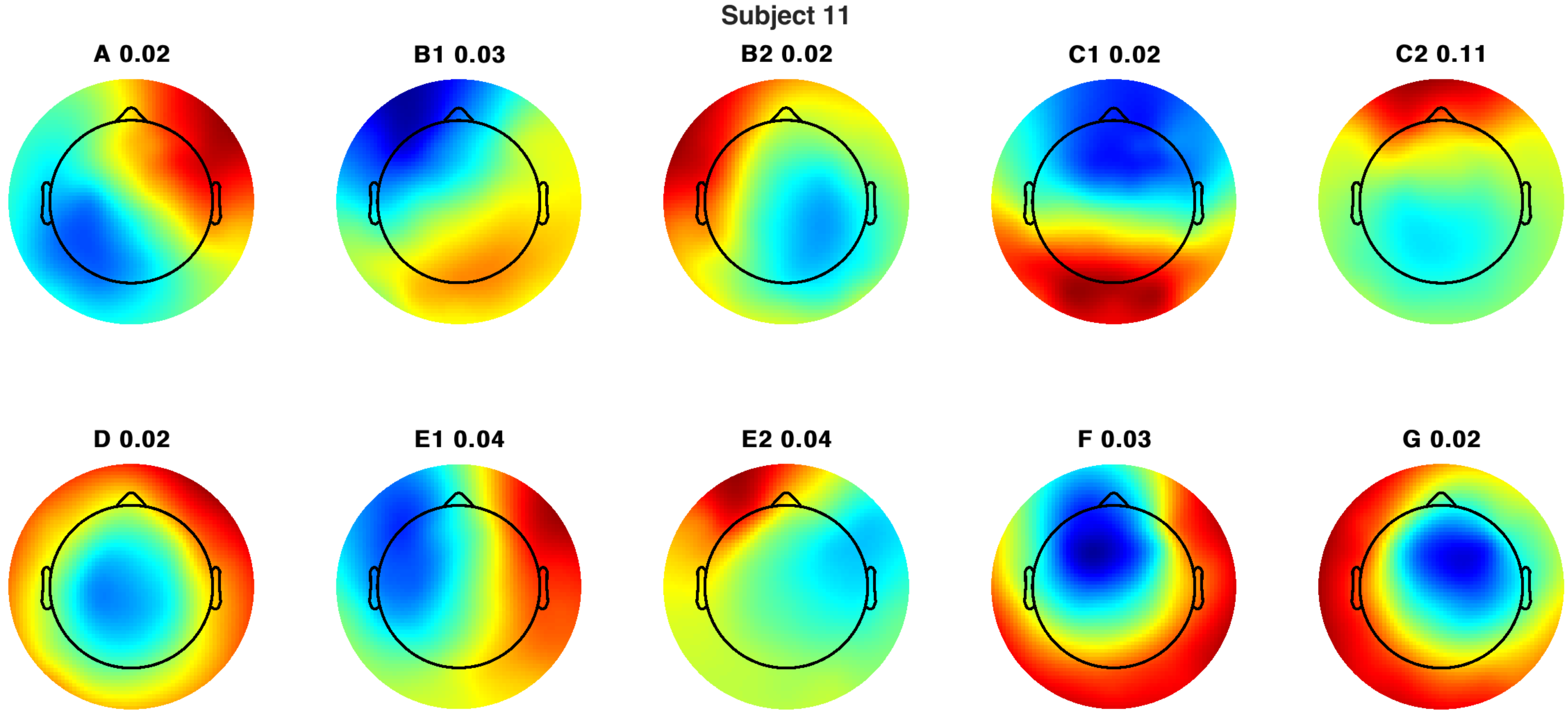
Subject 11
A 0.02
B1 0.03
B2 0.02
C1 0.02
C2 0.11
D 0.02
E1 0.04
E2 0.04
F 0.03
G 0.02

Subject 11

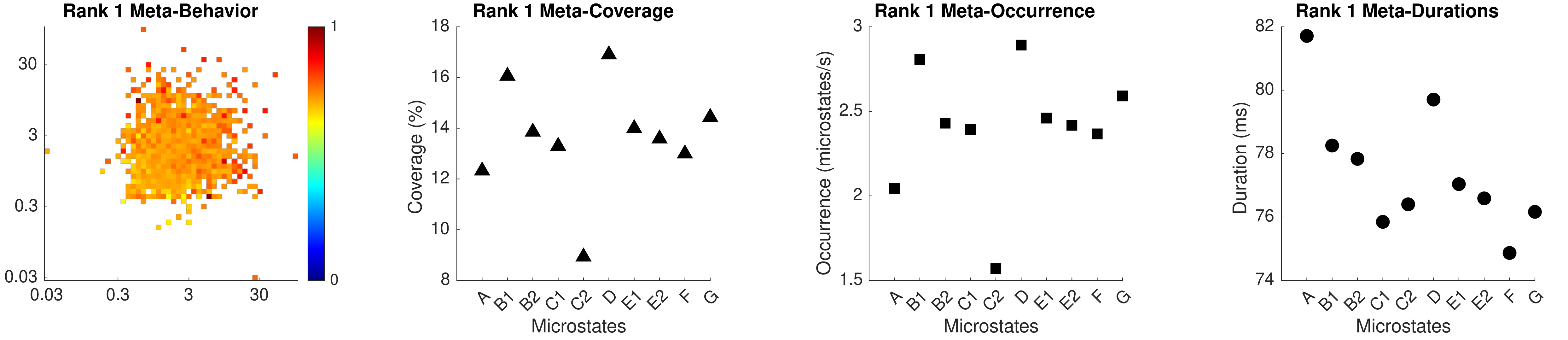

Rank 1 Meta-Behavior
30
3
0.3
0.03
0.03
0.3
3
30
1
0
Rank 1 Meta-Coverage
Coverage (%)
18
16
14
12
10
8
A
B1
B2
C1
C2
D
E1
E2
F
G
Microstates
Rank 1 Meta-Occurrence
Occurrence (microstates/s)
3
2.5
2
1.5
A
B1
B2
C1
C2
D
E1
E2
F
G
Microstates
Rank 1 Meta-Durations
Duration (ms)
82
80
78
76
74
A
B1
B2
C1
C2
D
E1
E2
F
G
Microstates

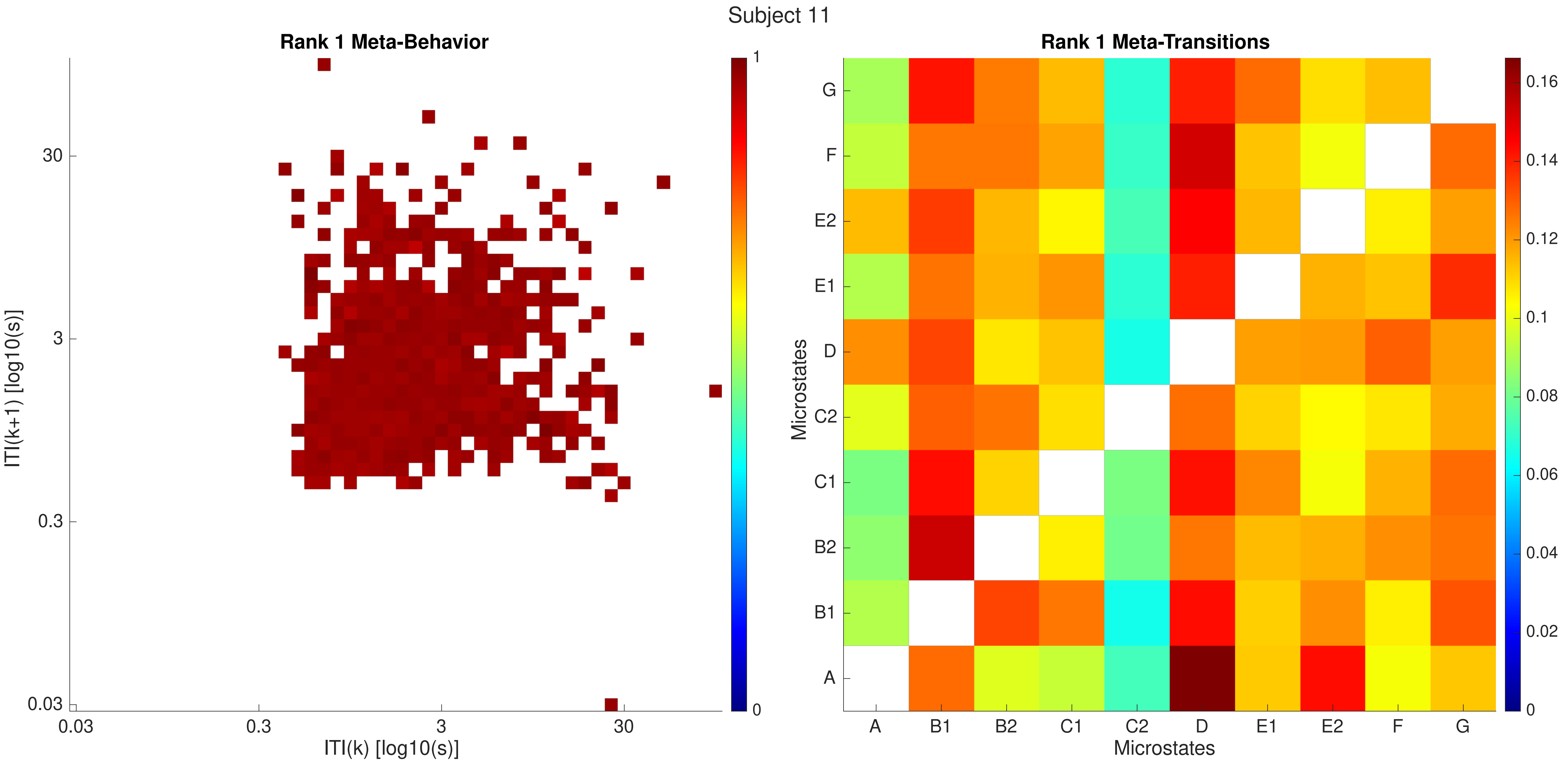
Subject 11
Rank 1 Meta-Behavior
ITI(k+1) [log10(s)]
ITI(k) [log10(s)]
0.03
0.3
3
30
1
0
Rank 1 Meta-Transitions
Microstates
A
B1
B2
C1
C2
D
E1
E2
F
G
0.16
0.14
0.12
0.1
0.08
0.06
0.04
0.02
0

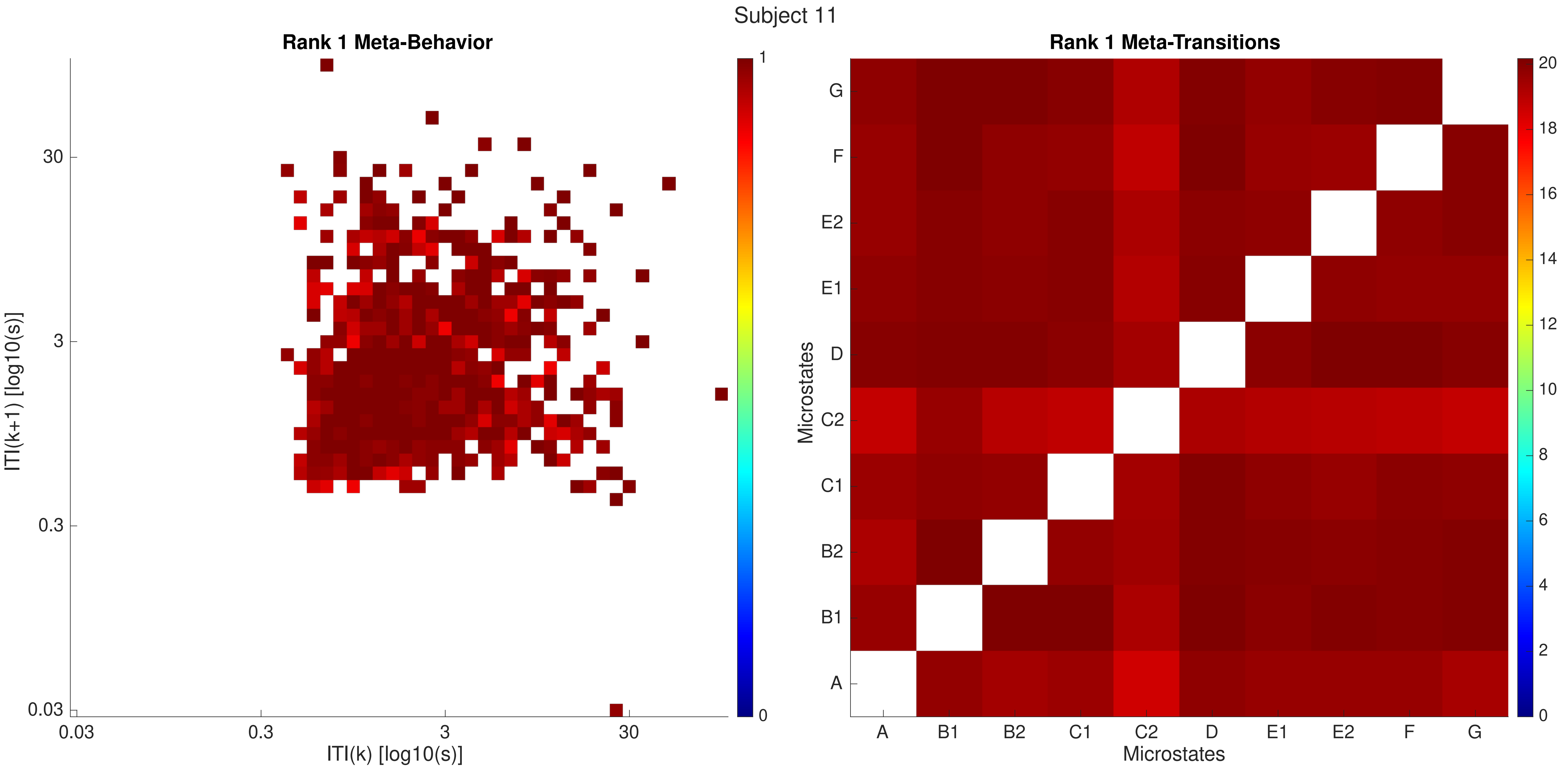

Subject 11
Rank 1 Meta-Behavior
ITI(k+1) [log10(s)]
ITI(k) [log10(s)]
0.03
0.3
3
30
0
1
Rank 1 Meta-Transitions
Microstates
A
B1
B2
C1
C2
D
E1
E2
F
G
0
2
4
6
8
10
12
14
16
18
20

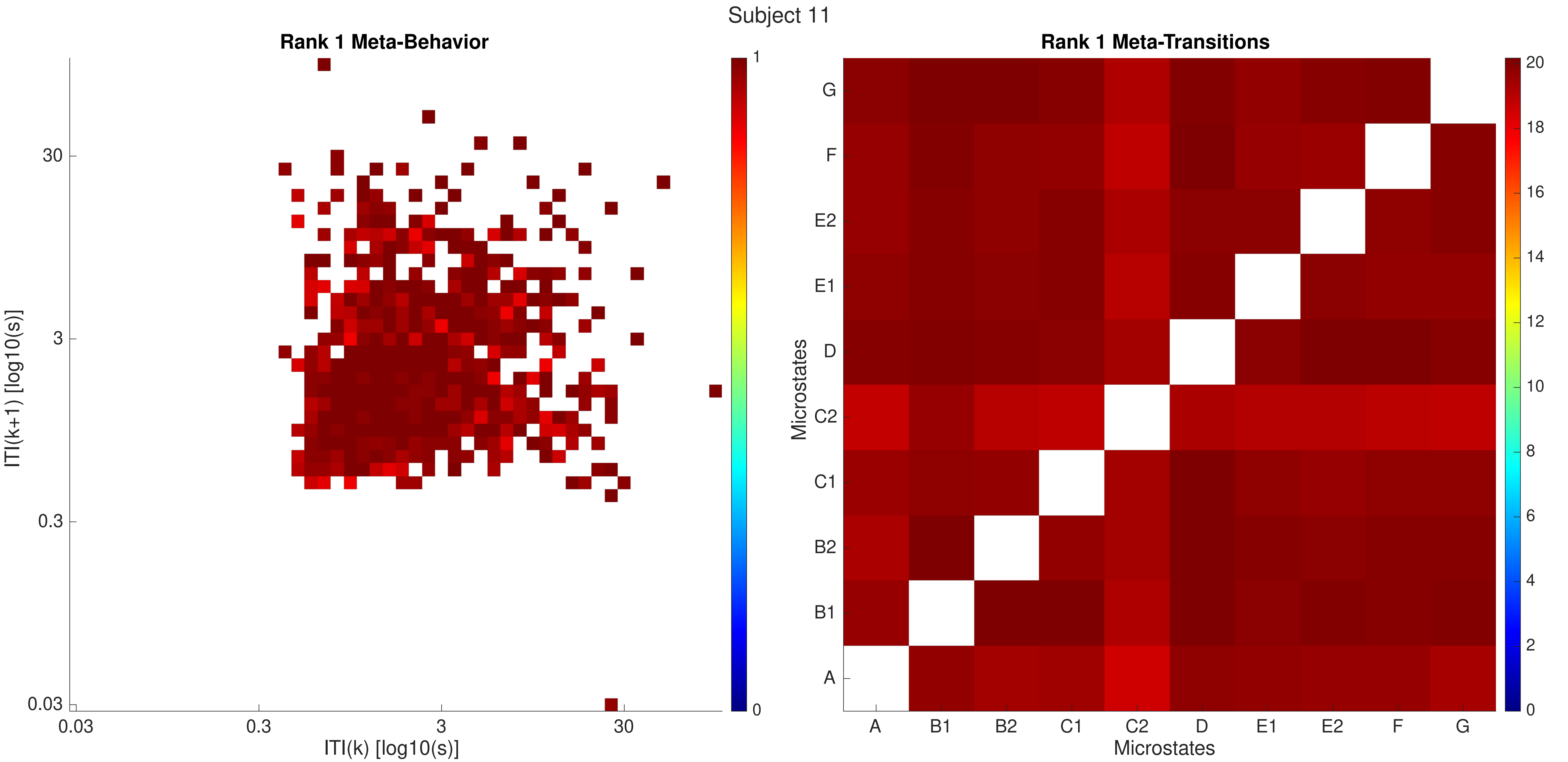
Subject 11
Rank 1 Meta-Behavior
ITI(k+1) [log10(s)]
30
3
0.3
0.03
ITI(k) [log10(s)]
1
0
Rank 1 Meta-Transitions
Microstates
A
B1
B2
C1
C2
D
E1
E2
F
G
20
18
16
14
12
10
8
6
4
2
0

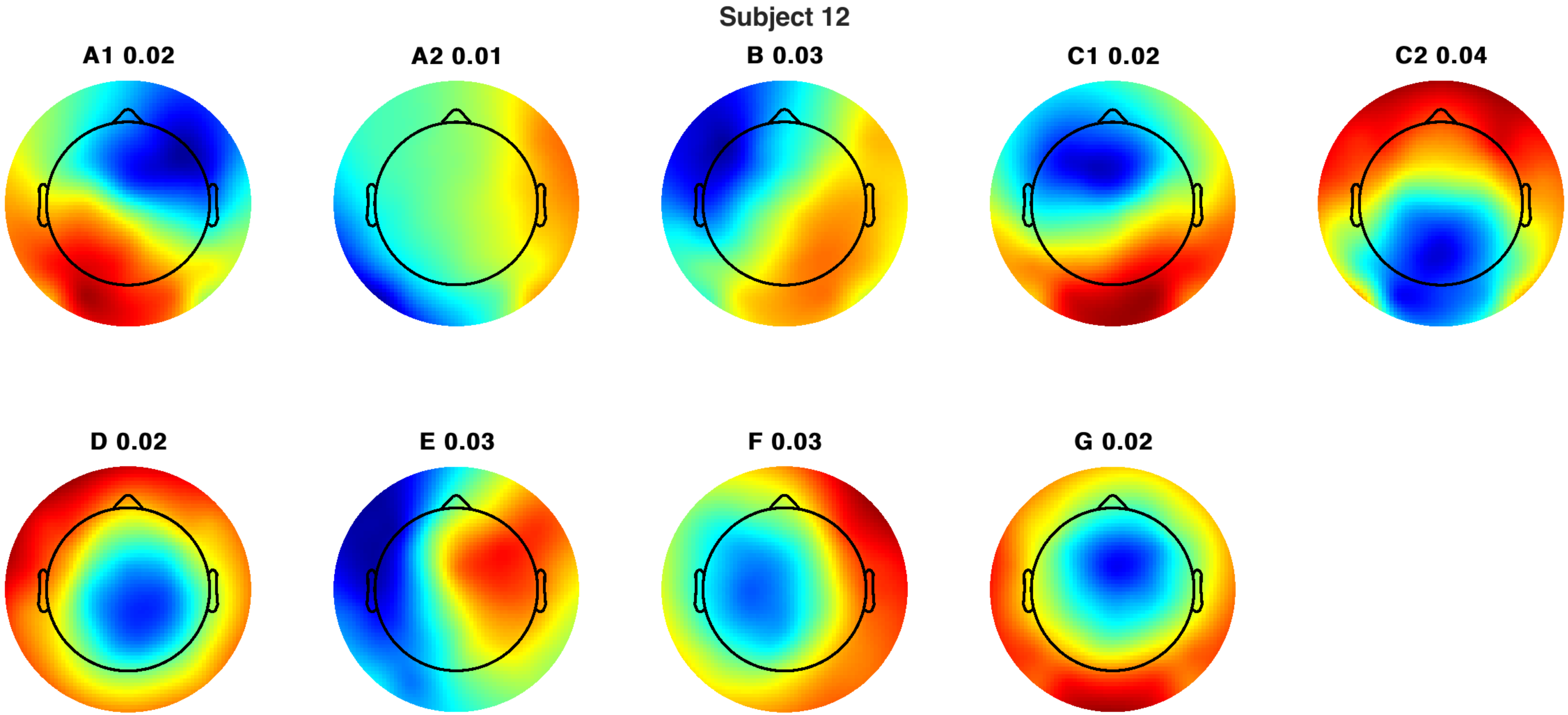
Subject 12
A1 0.02
A2 0.01
B 0.03
C1 0.02
C2 0.04
D 0.02
E 0.03
F 0.03
G 0.02

Subject 12

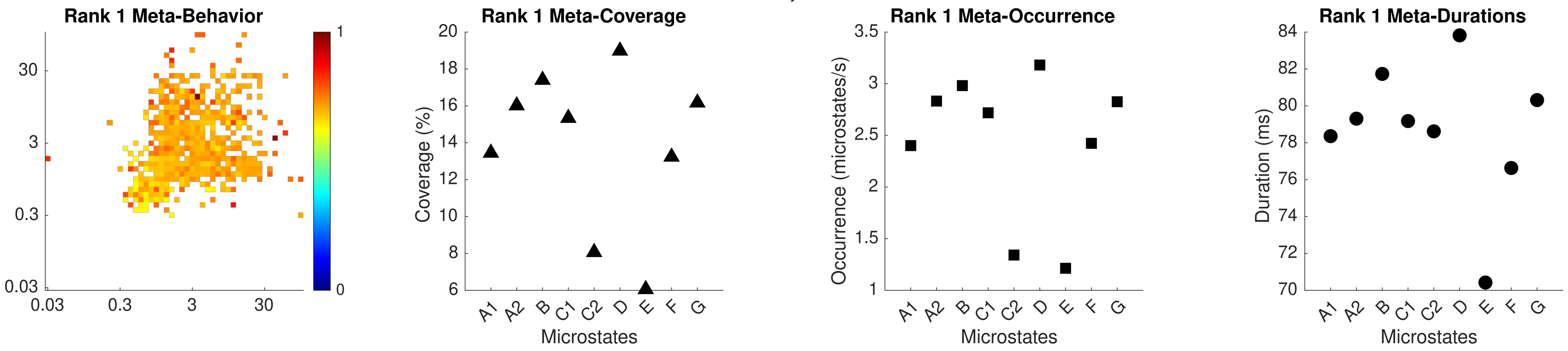

Rank 1 Meta-Behavior
Rank 1 Meta-Coverage
Rank 1 Meta-Occurrence
Rank 1 Meta-Durations
Coverage (%)
Occurrence (microstates/s)
Duration (ms)
Microstates
A1 A2 B C1 C2 D E F G

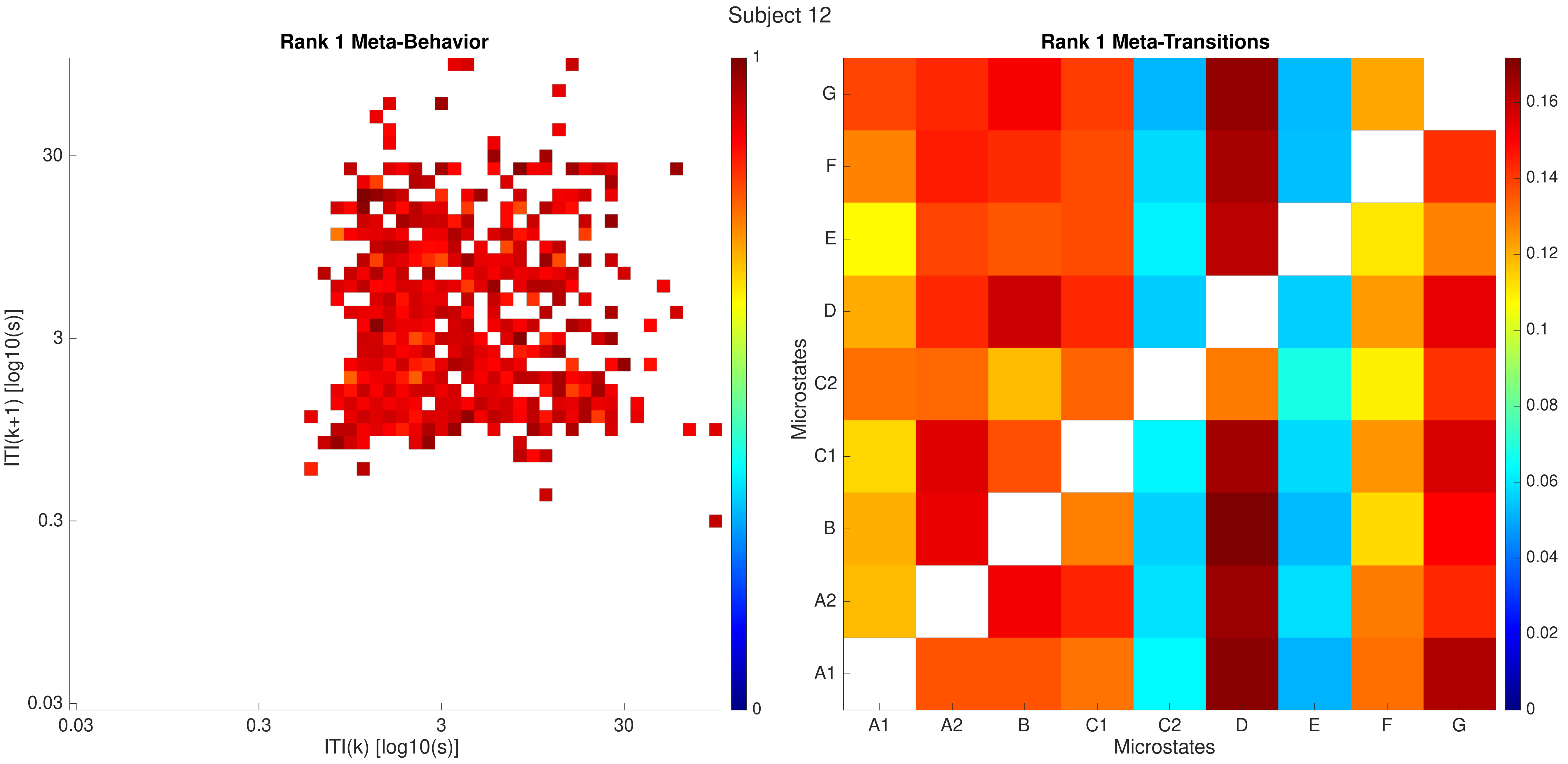
Subject 12
Rank 1 Meta-Behavior
ITI(k+1) [log10(s)]
ITI(k) [log10(s)]
0.03
0.3
3
30
1
0
Rank 1 Meta-Transitions
Microstates
A1
A2
B
C1
C2
D
E
F
G
0.16
0.14
0.12
0.1
0.08
0.06
0.04
0.02

Subject 12

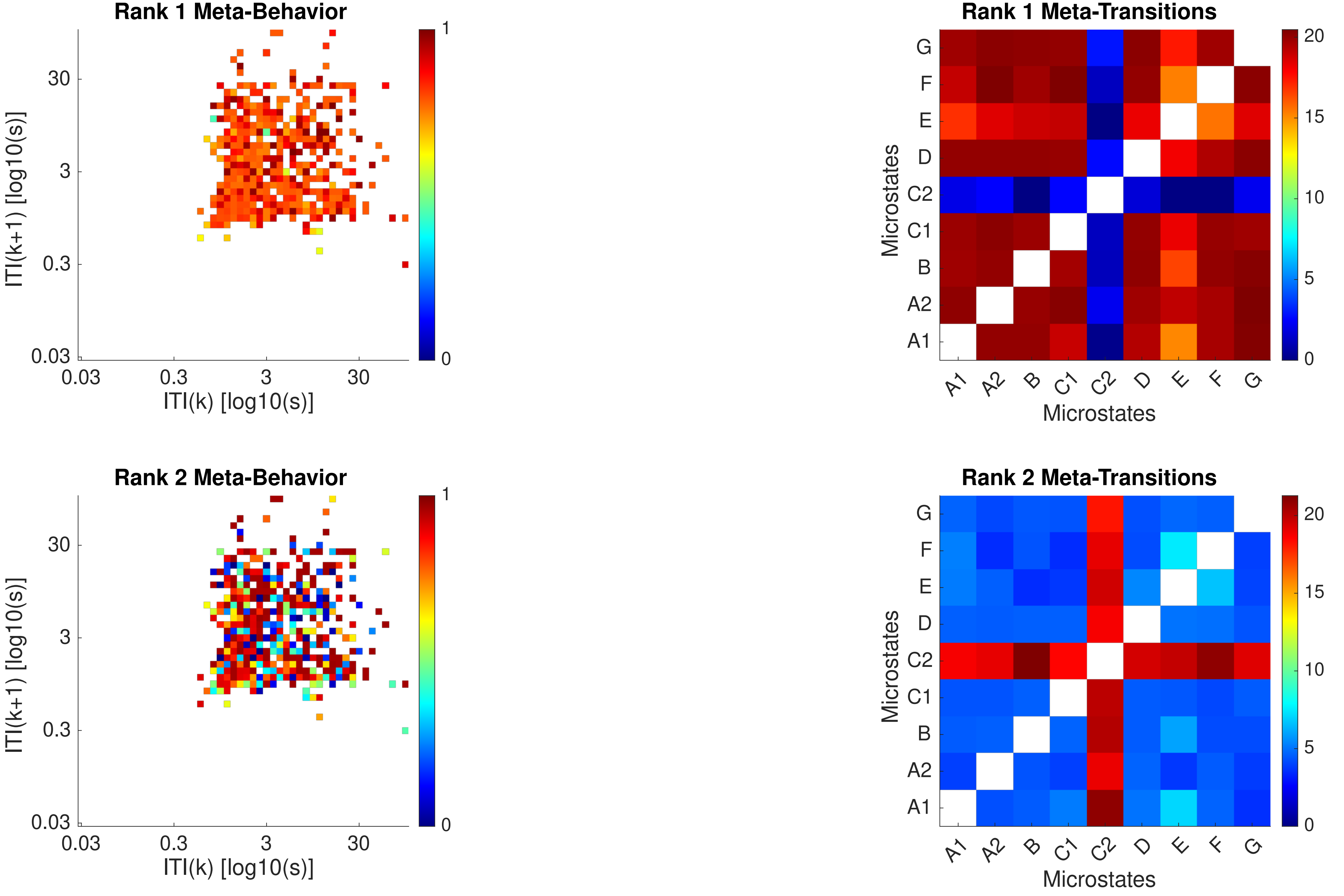

Rank 1 Meta-Behavior
ITI(k+1) [log10(s)]
ITI(k) [log10(s)]
0.03
0.3
3
30
0
1
Rank 1 Meta-Transitions
Microstates
G
F
E
D
C2
C1
B
A2
A1
0
5
10
15
20
Rank 2 Meta-Behavior
ITI(k+1) [log10(s)]
ITI(k) [log10(s)]
Rank 2 Meta-Transitions
Microstates

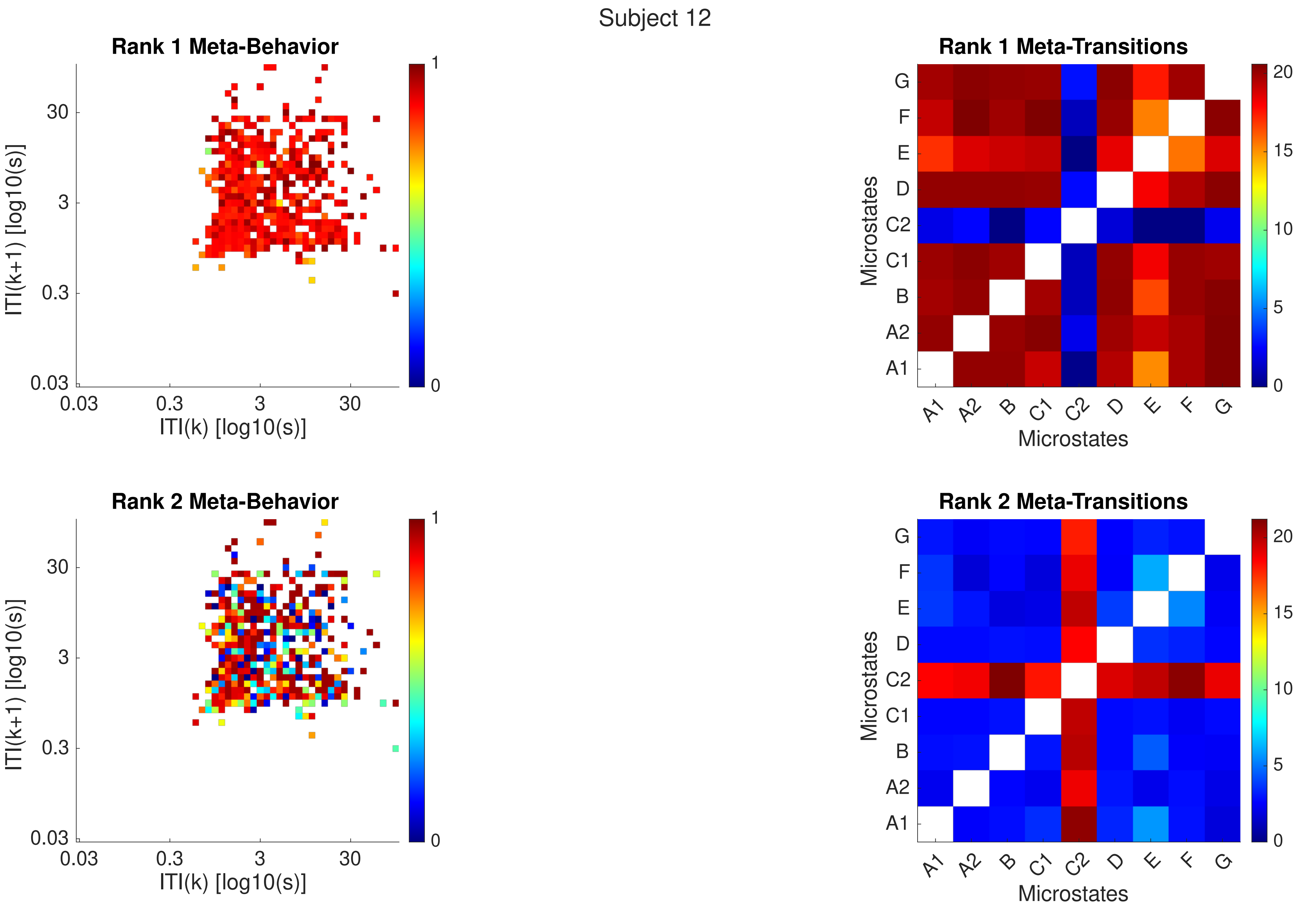
Subject 12
Rank 1 Meta-Behavior
ITI(k+1) [log10(s)]
ITI(k) [log10(s)]
0.03
0.3
3
30
0
1
Rank 1 Meta-Transitions
Microstates
A1
A2
B
C1
C2
D
E
F
G
0
5
10
15
20
Rank 2 Meta-Behavior
Rank 2 Meta-Transitions

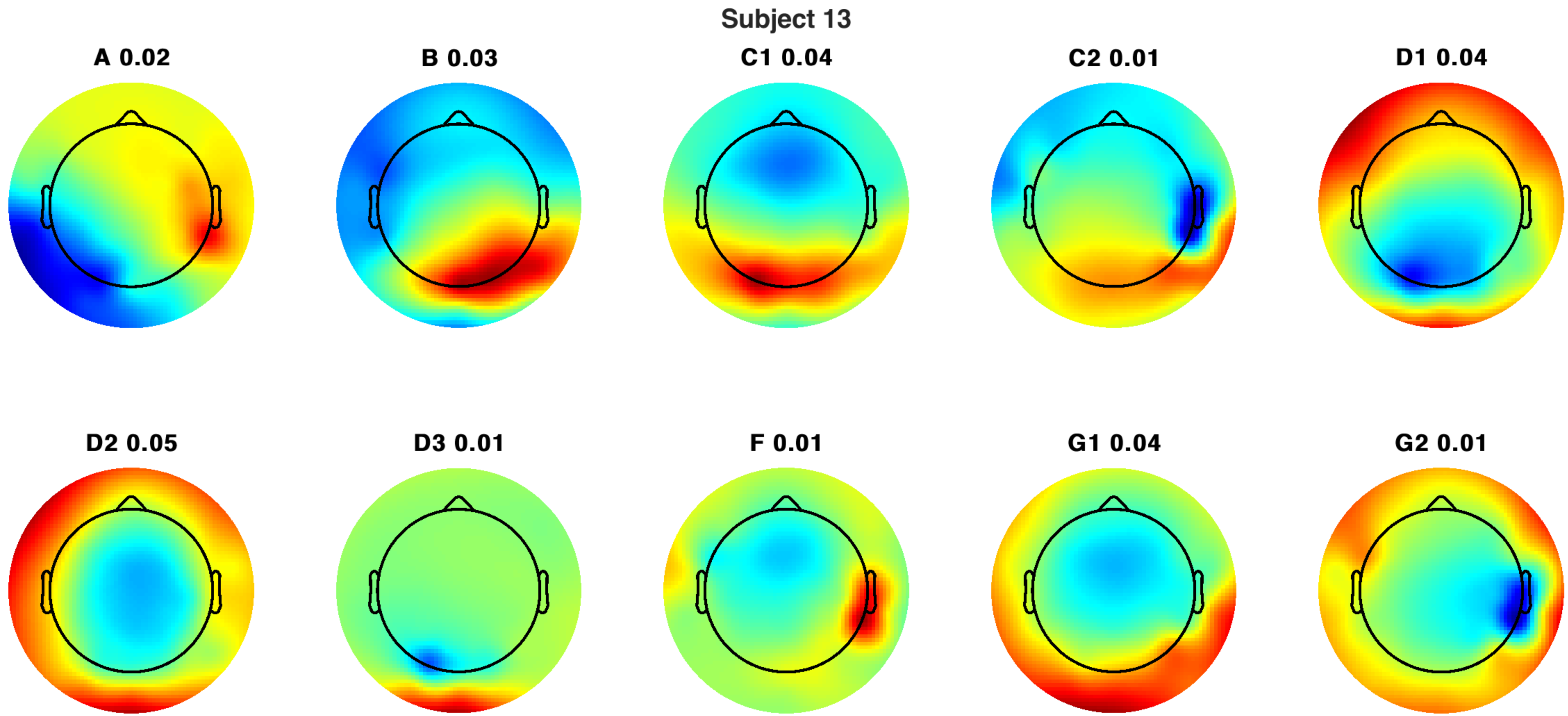
Subject 13
A 0.02
B 0.03
C1 0.04
C2 0.01
D1 0.04
D2 0.05
D3 0.01
F 0.01
G1 0.04
G2 0.01

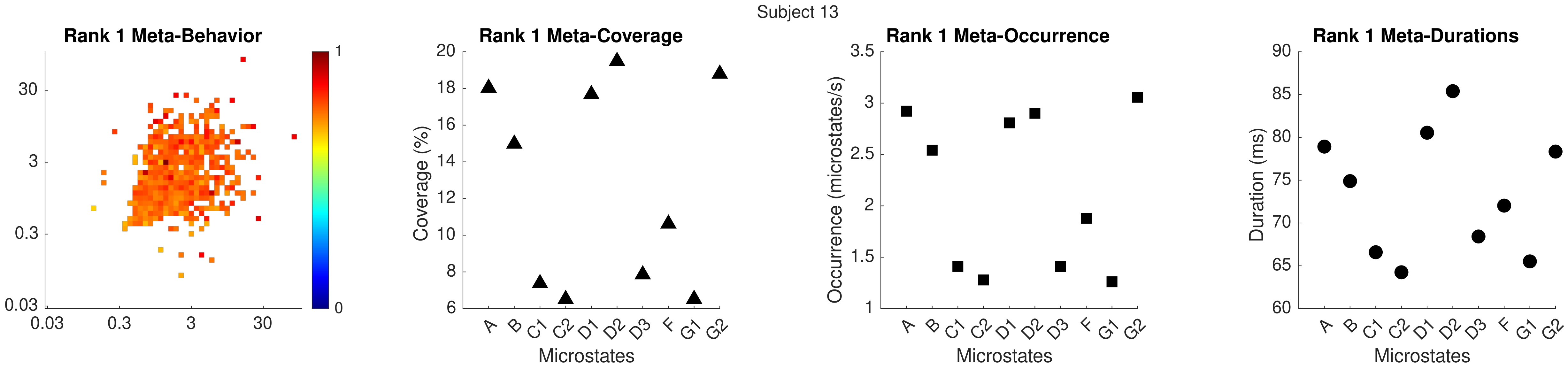
Subject 13
Rank 1 Meta-Behavior
Rank 1 Meta-Coverage
Rank 1 Meta-Occurrence
Rank 1 Meta-Durations
Coverage (%)
Occurrence (microstates/s)
Duration (ms)
Microstates
A B C1 C2 D1 D2 D3 F G1 G2

Subject 13

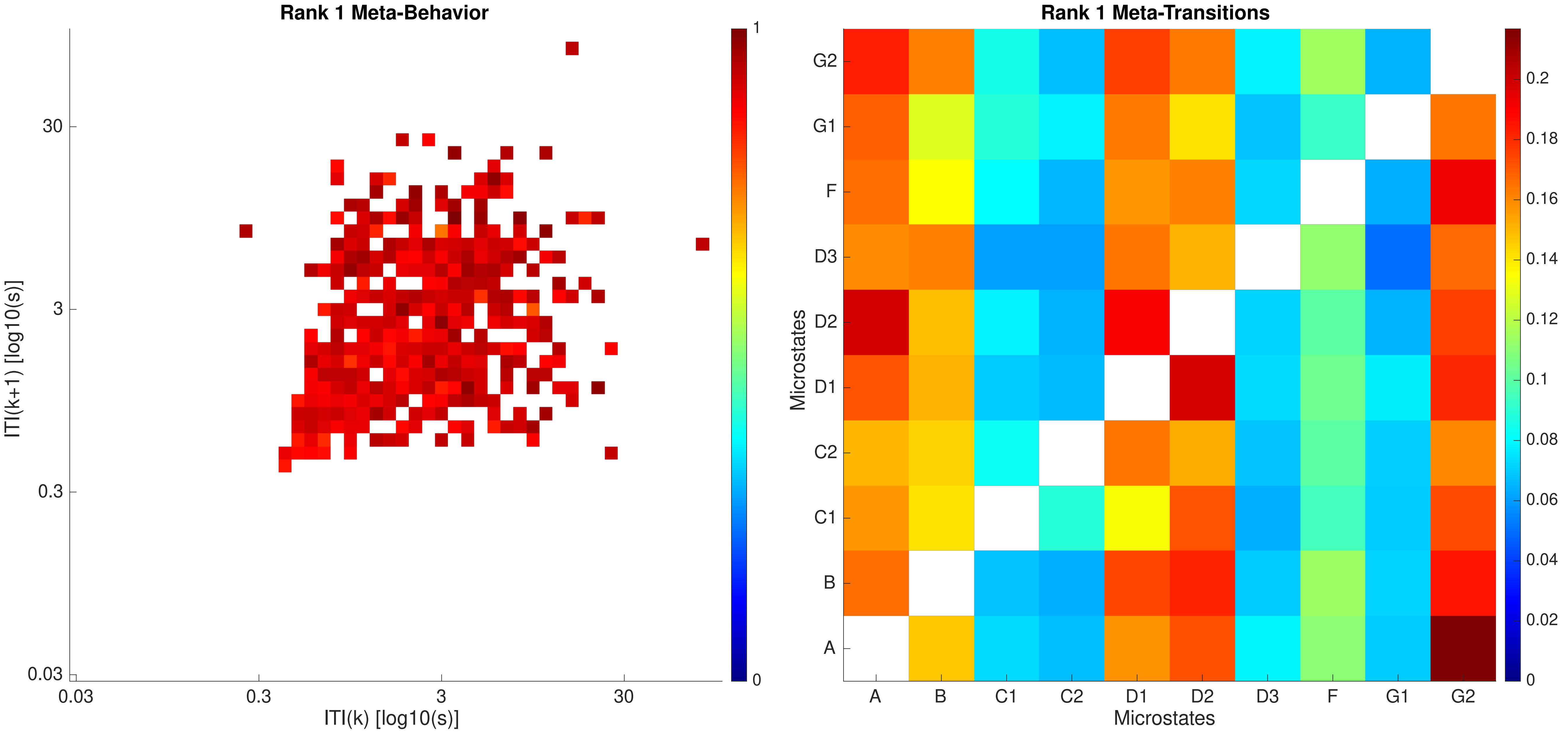

Rank 1 Meta-Behavior
ITI(k+1) [log10(s)]
ITI(k) [log10(s)]
0.03
0.3
3
30
0
1
Rank 1 Meta-Transitions
Microstates
A
B
C1
C2
D1
D2
D3
F
G1
G2
0
0.02
0.04
0.06
0.08
0.1
0.12
0.14
0.16
0.18
0.2

Subject 13

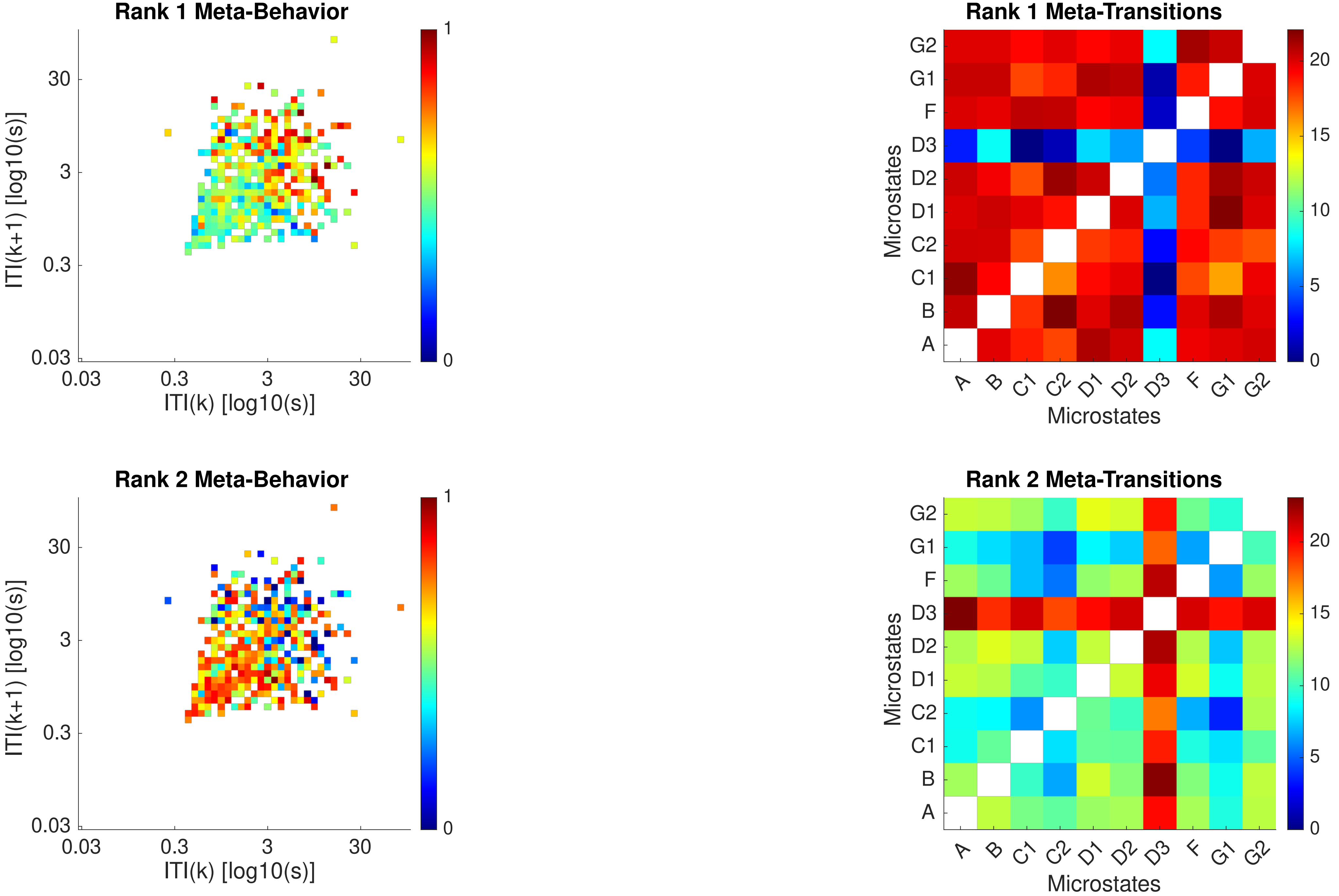

Rank 1 Meta-Behavior
ITI(k+1) [log10(s)]
ITI(k) [log10(s)]
0.03
0.3
3
30
0
1
Rank 1 Meta-Transitions
Microstates
A
B
C1
C2
D1
D2
D3
F
G1
G2
0
5
10
15
20
Rank 2 Meta-Behavior
ITI(k+1) [log10(s)]
ITI(k) [log10(s)]
Rank 2 Meta-Transitions
Microstates

Subject 13

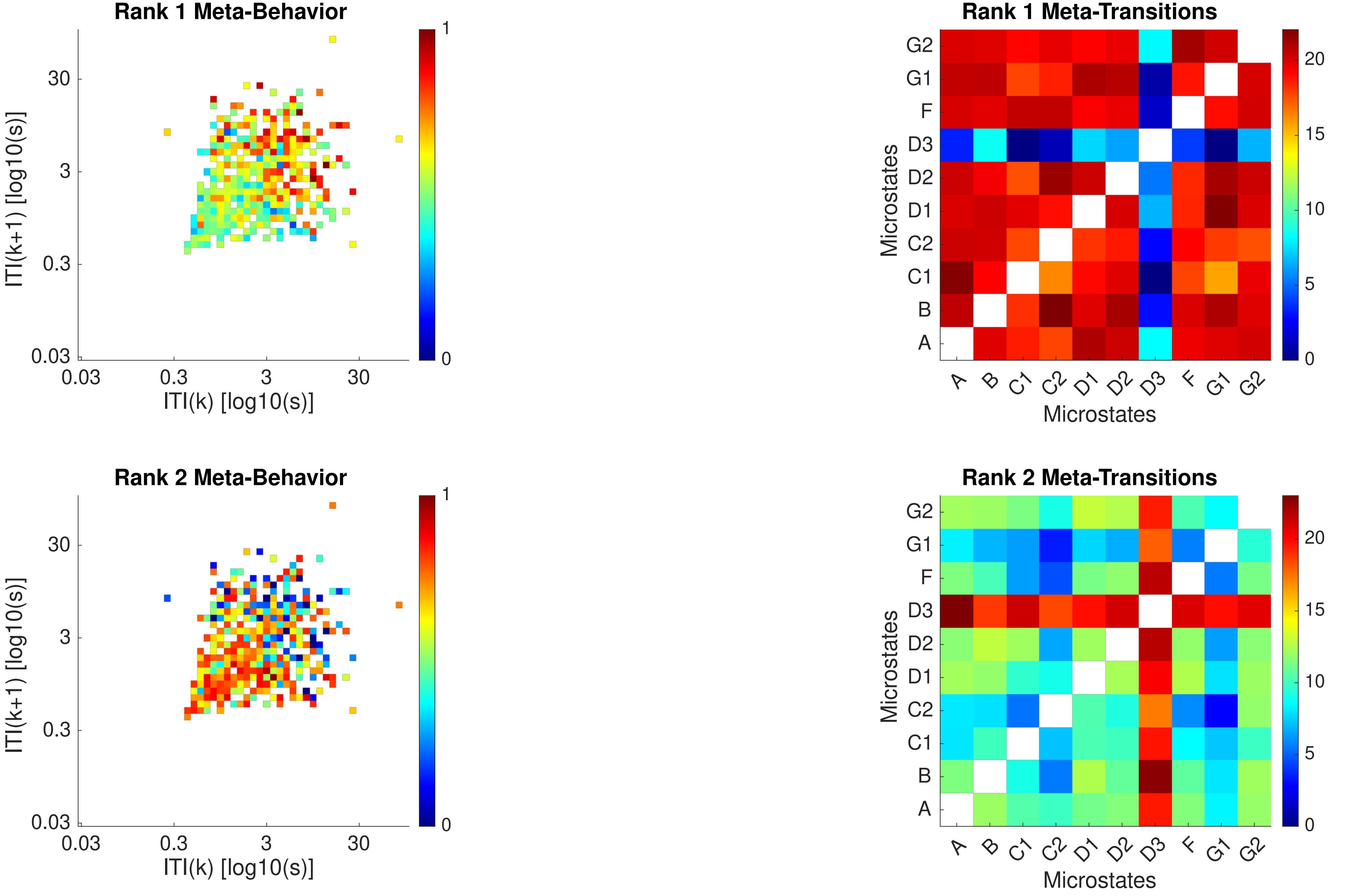
Rank 1 Meta-Behavior
ITI(k+1) [log10(s)]
ITI(k) [log10(s)]
0.03
0.3
3
30
0
1
Rank 1 Meta-Transitions
Microstates
A
B
C1
C2
D1
D2
D3
F
G1
G2
0
5
10
15
20
Rank 2 Meta-Behavior
Rank 2 Meta-Transitions

Subject 14

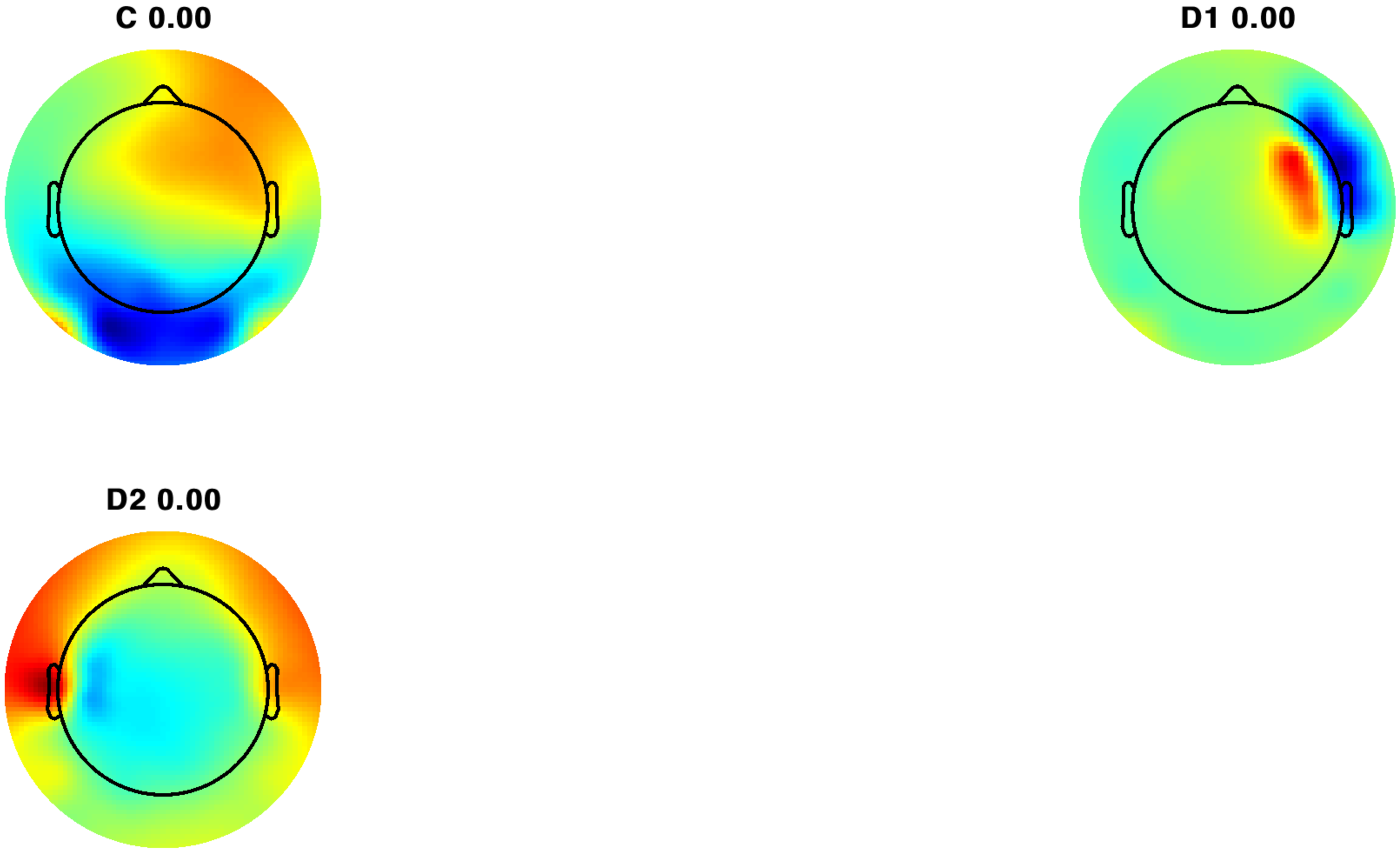

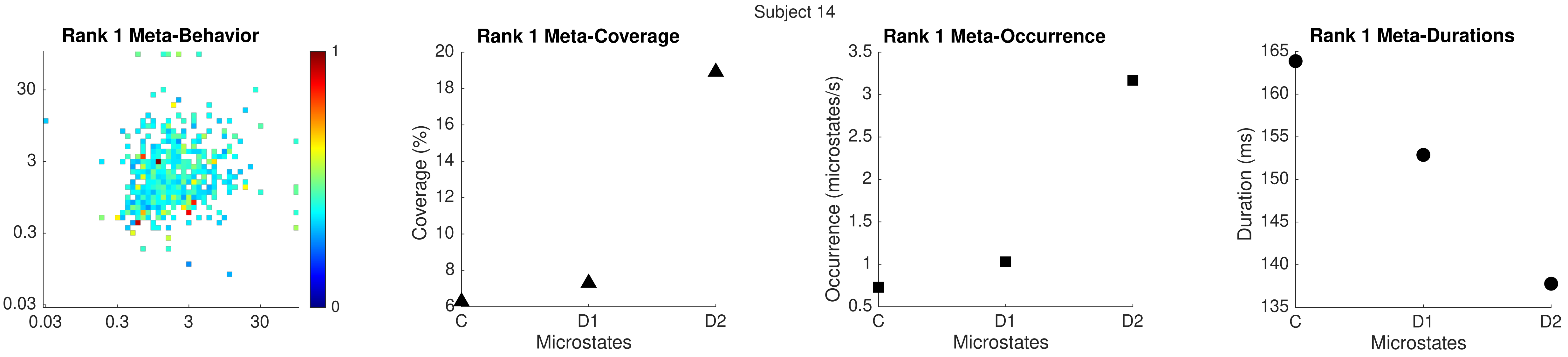
Subject 14
Rank 1 Meta-Behavior
30
3
0.3
0.03
0.03
0.3
3
30
1
0
Rank 1 Meta-Coverage
Coverage (%)
20
18
16
14
12
10
8
6
C
D1
D2
Microstates
Rank 1 Meta-Occurrence
Occurrence (microstates/s)
3.5
3
2.5
2
1.5
1
0.5
C
D1
D2
Microstates
Rank 1 Meta-Durations
Duration (ms)
165
160
155
150
145
140
135
C
D1
D2
Microstates

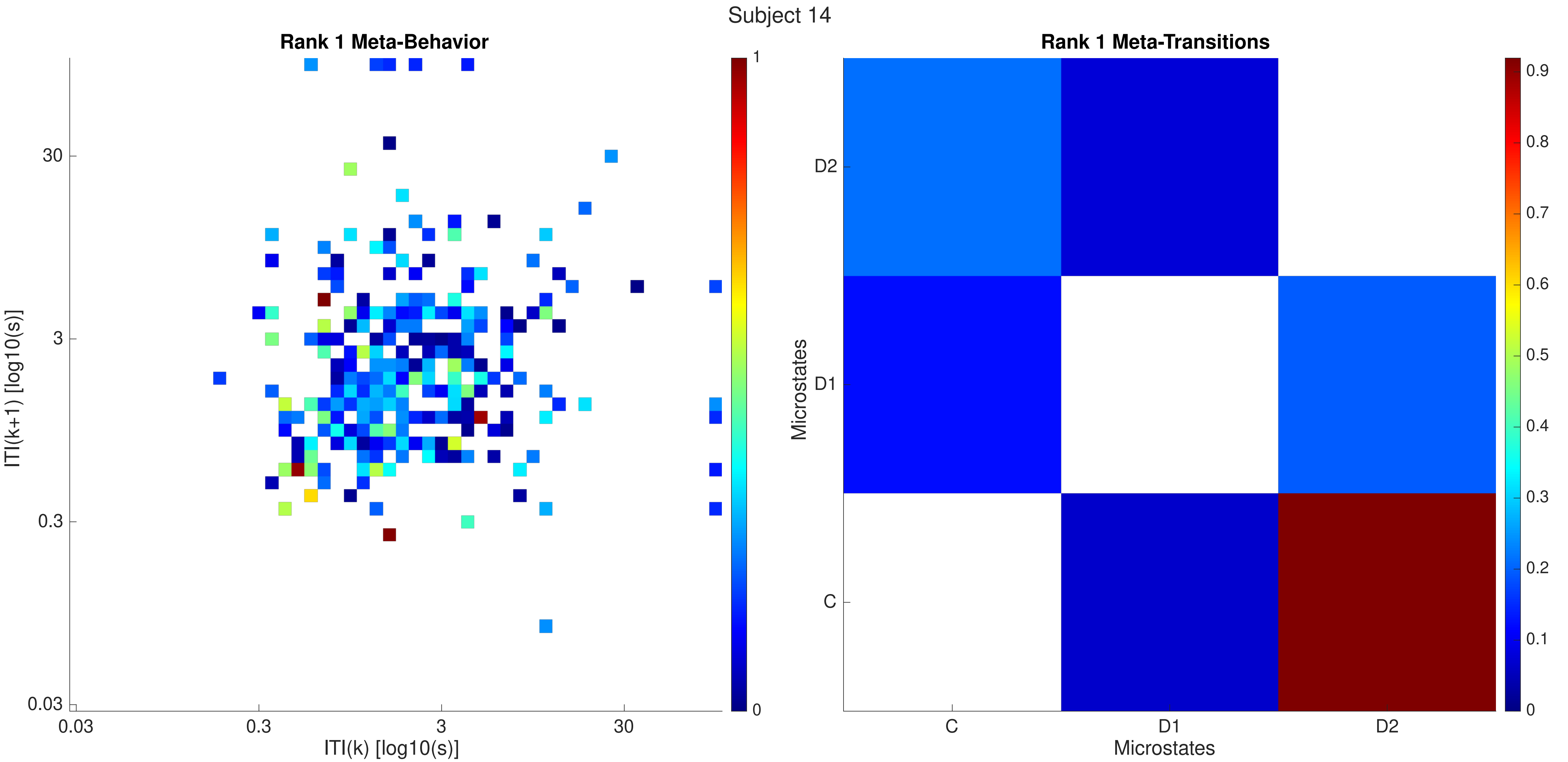

Subject 14
Rank 1 Meta-Behavior
ITI(k+1) [log10(s)]
ITI(k) [log10(s)]
30
3
0.3
0.03
1
0
Rank 1 Meta-Transitions
Microstates
D2
D1
C
Microstates
0.9
0.8
0.7
0.6
0.5
0.4
0.3
0.2
0.1
0

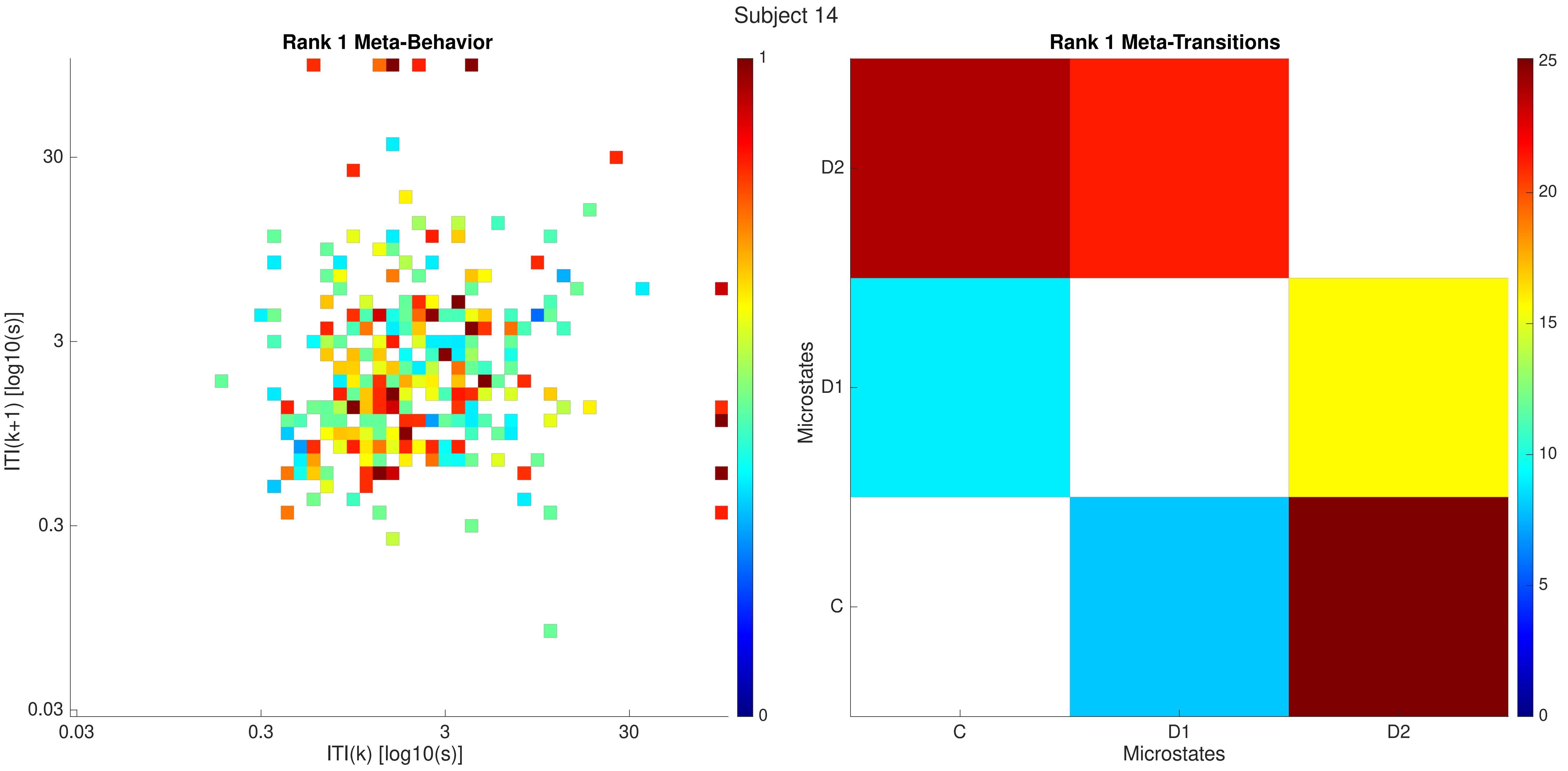
Subject 14
Rank 1 Meta-Behavior
ITI(k+1) [log10(s)]
ITI(k) [log10(s)]
0.03
0.3
3
30
1
0
Rank 1 Meta-Transitions
Microstates
Microstates
C
D1
D2
25
20
15
10
5
0

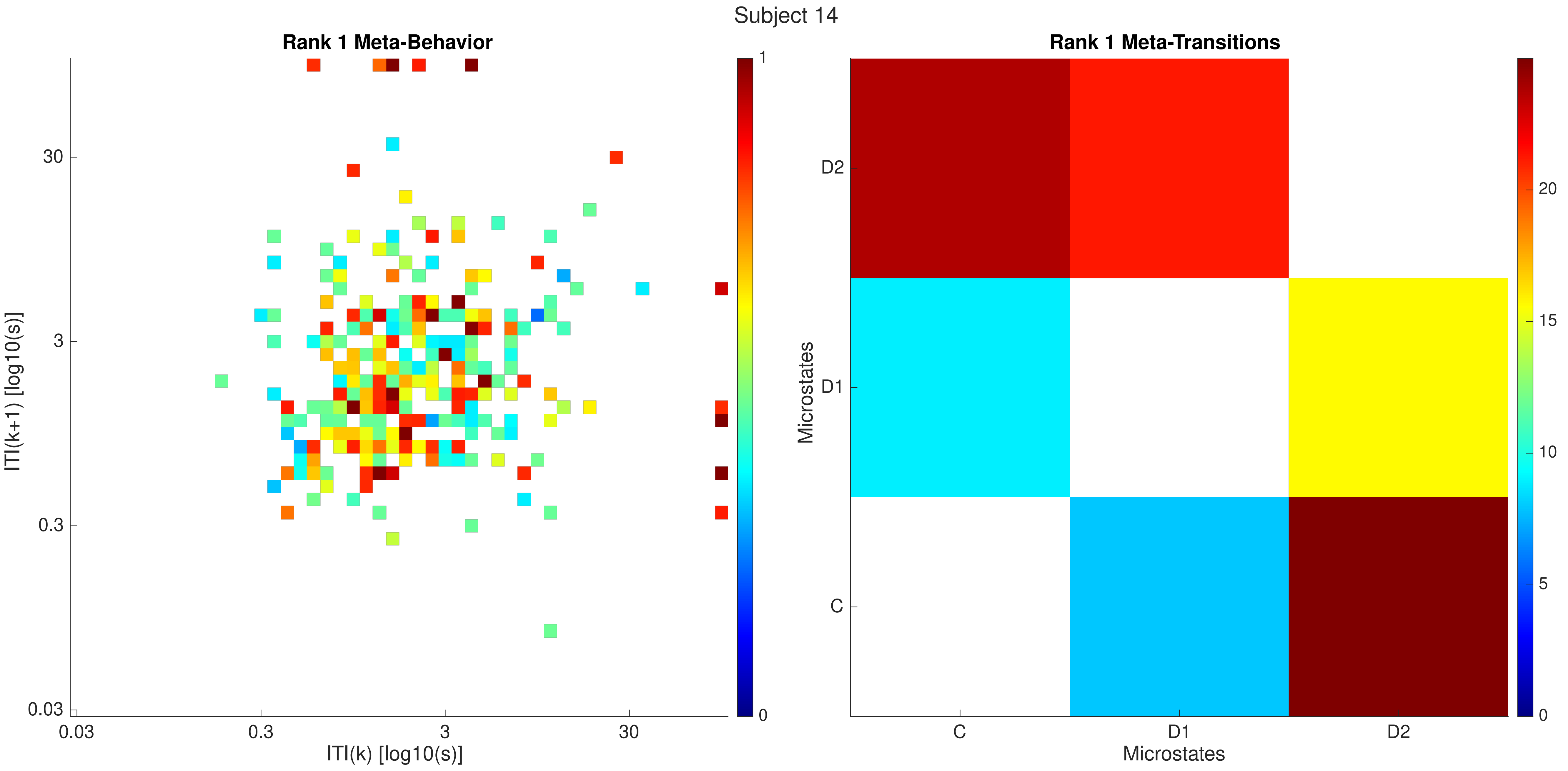

Subject 14
Rank 1 Meta-Behavior
ITI(k+1) [log10(s)]
ITI(k) [log10(s)]
0.03
0.3
3
30
0
1
Rank 1 Meta-Transitions
Microstates
C
D1
D2
0
5
10
15
20

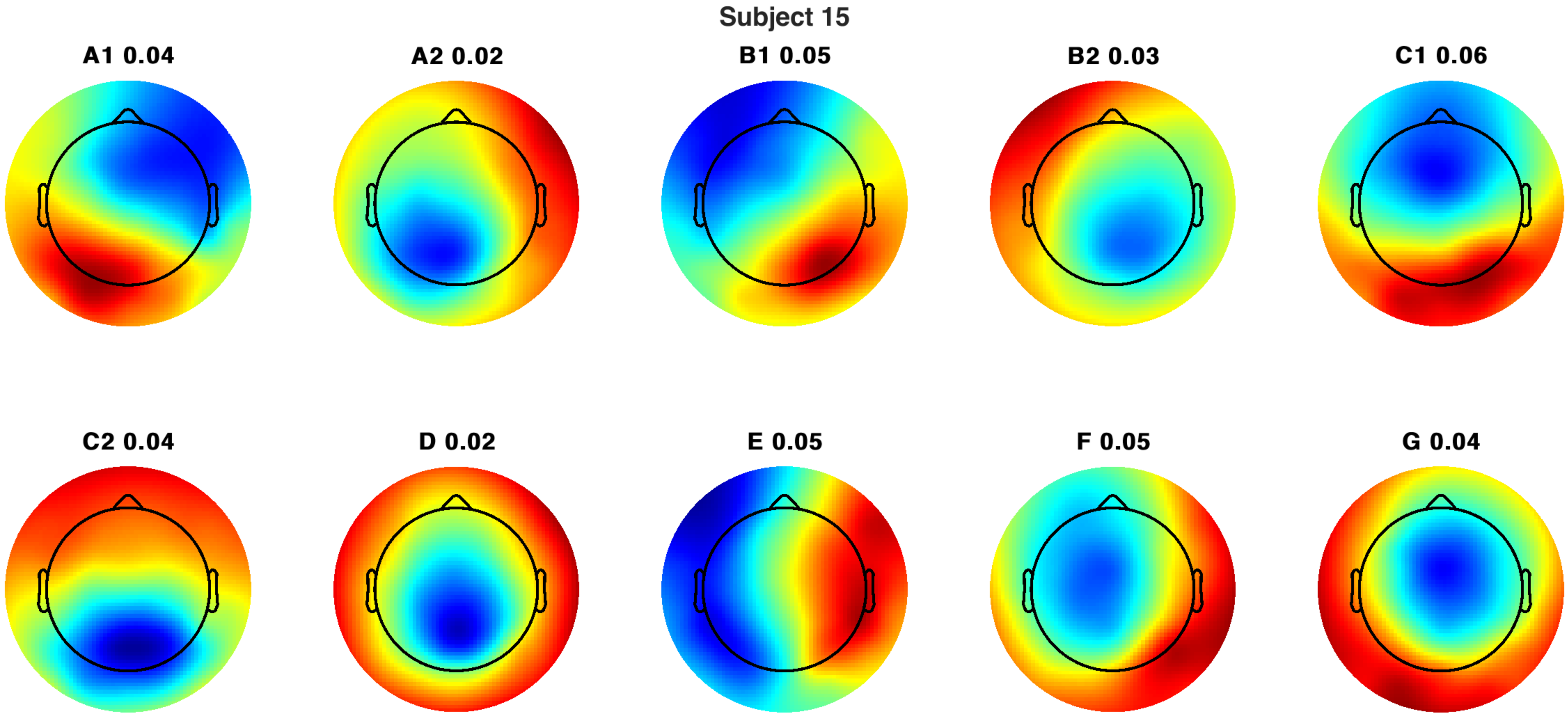
Subject 15
A1 0.04
A2 0.02
B1 0.05
B2 0.03
C1 0.06
C2 0.04
D 0.02
E 0.05
F 0.05
G 0.04

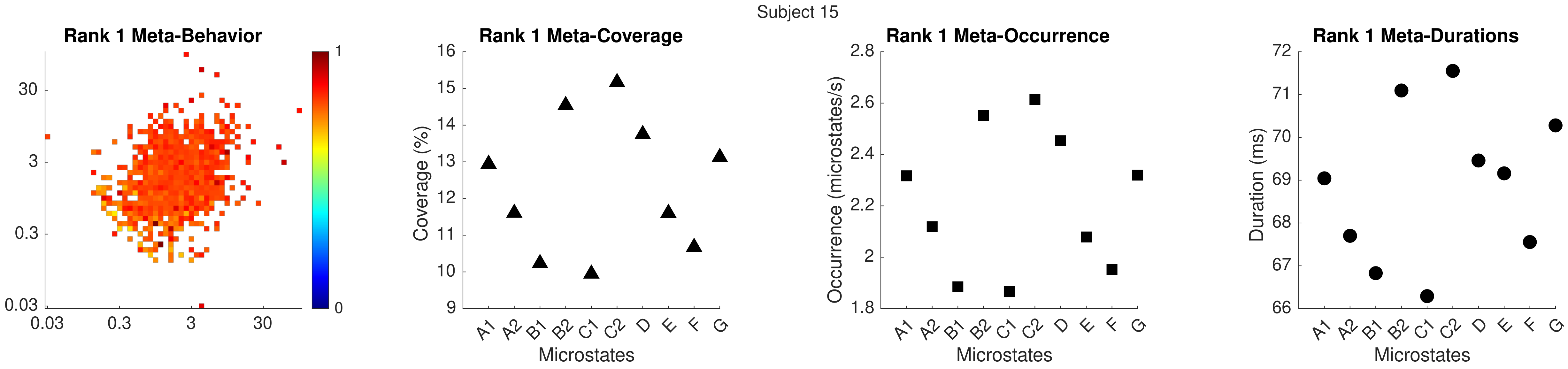

Subject 15
Rank 1 Meta-Behavior
30
3
0.3
0.03
0.03
0.3
3
30
1
0
Rank 1 Meta-Coverage
Coverage (%)
16
15
14
13
12
11
10
9
A1 A2 B1 B2 C1 C2 D E F G
Microstates
Rank 1 Meta-Occurrence
Occurrence (microstates/s)
2.8
2.6
2.4
2.2
2
1.8
A1 A2 B1 B2 C1 C2 D E F G
Microstates
Rank 1 Meta-Durations
Duration (ms)
72
71
70
69
68
67
66
A1 A2 B1 B2 C1 C2 D E F G
Microstates

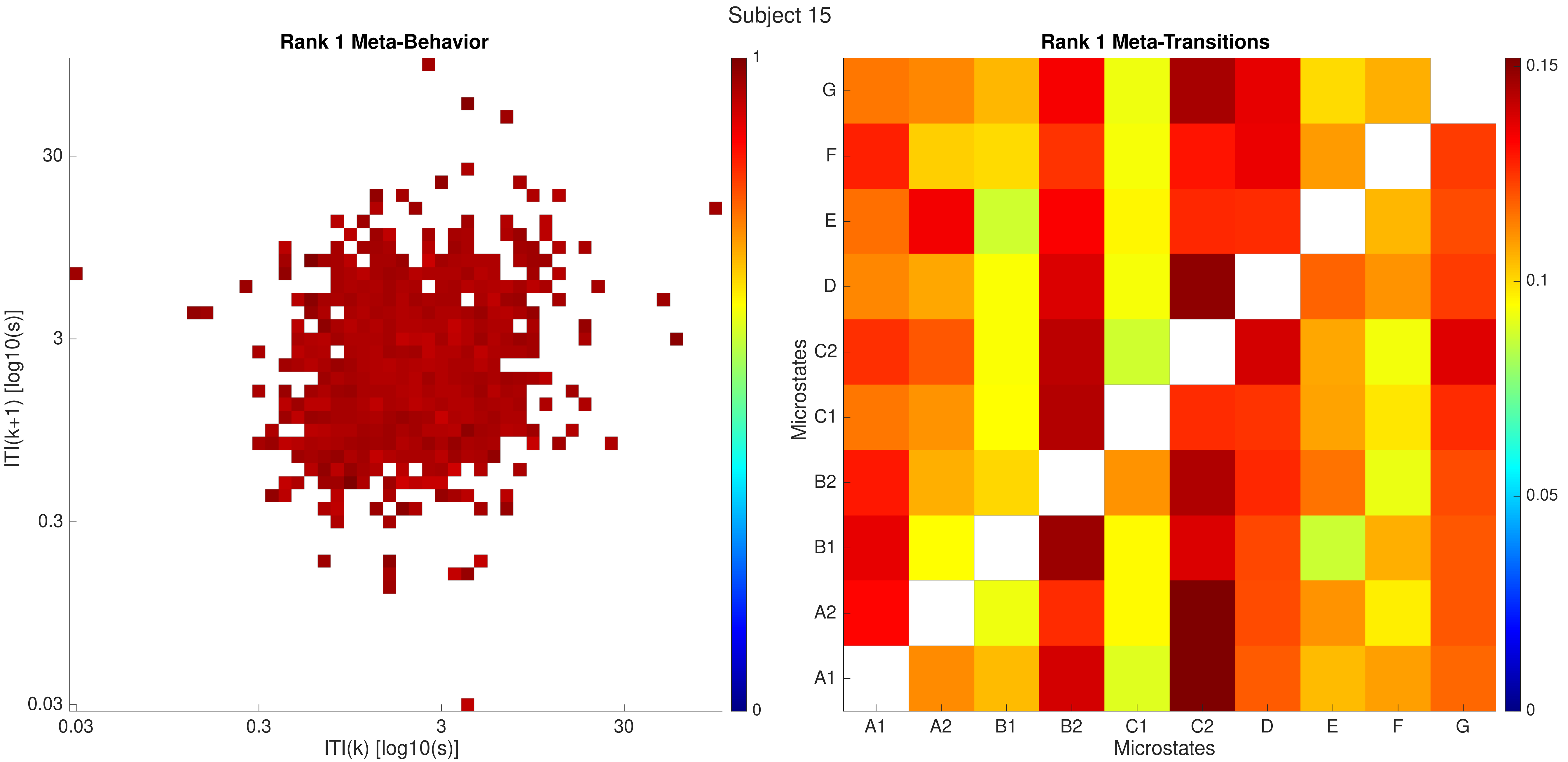
Subject 15
Rank 1 Meta-Behavior
ITI(k+1) [log10(s)]
ITI(k) [log10(s)]
0.03
0.3
3
30
1
0
Rank 1 Meta-Transitions
Microstates
A1
A2
B1
B2
C1
C2
D
E
F
G
0.15
0.1
0.05
0

Subject 15

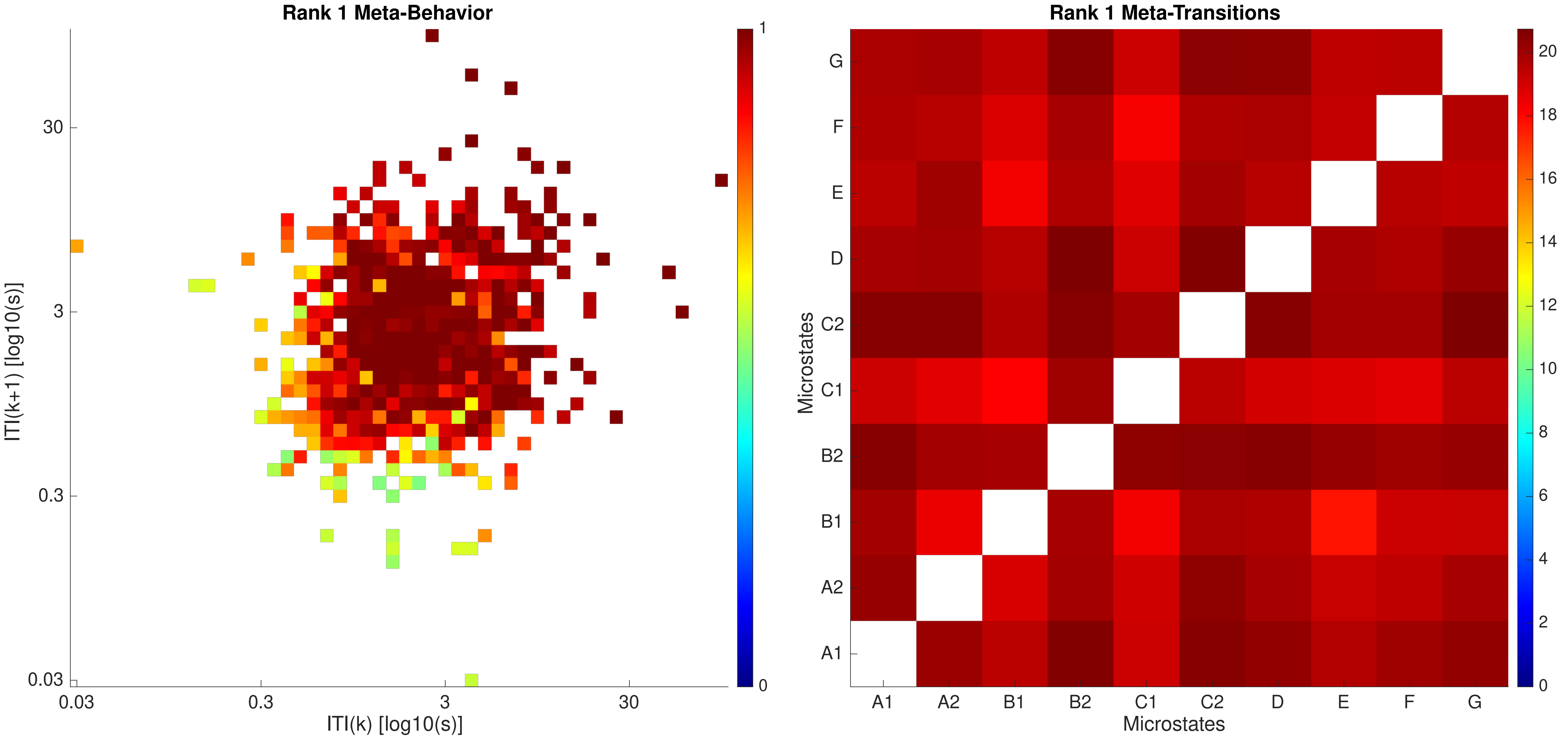
Rank 1 Meta-Behavior
ITI(k+1) [log10(s)]
ITI(k) [log10(s)]
30
3
0.3
0.03
0.03
0.3
3
30
1
0
Rank 1 Meta-Transitions
Microstates
G
F
E
D
C2
C1
B2
B1
A2
A1
A1
A2
B1
B2
C1
C2
D
E
F
G
Microstates
20
18
16
14
12
10
8
6
4
2
0

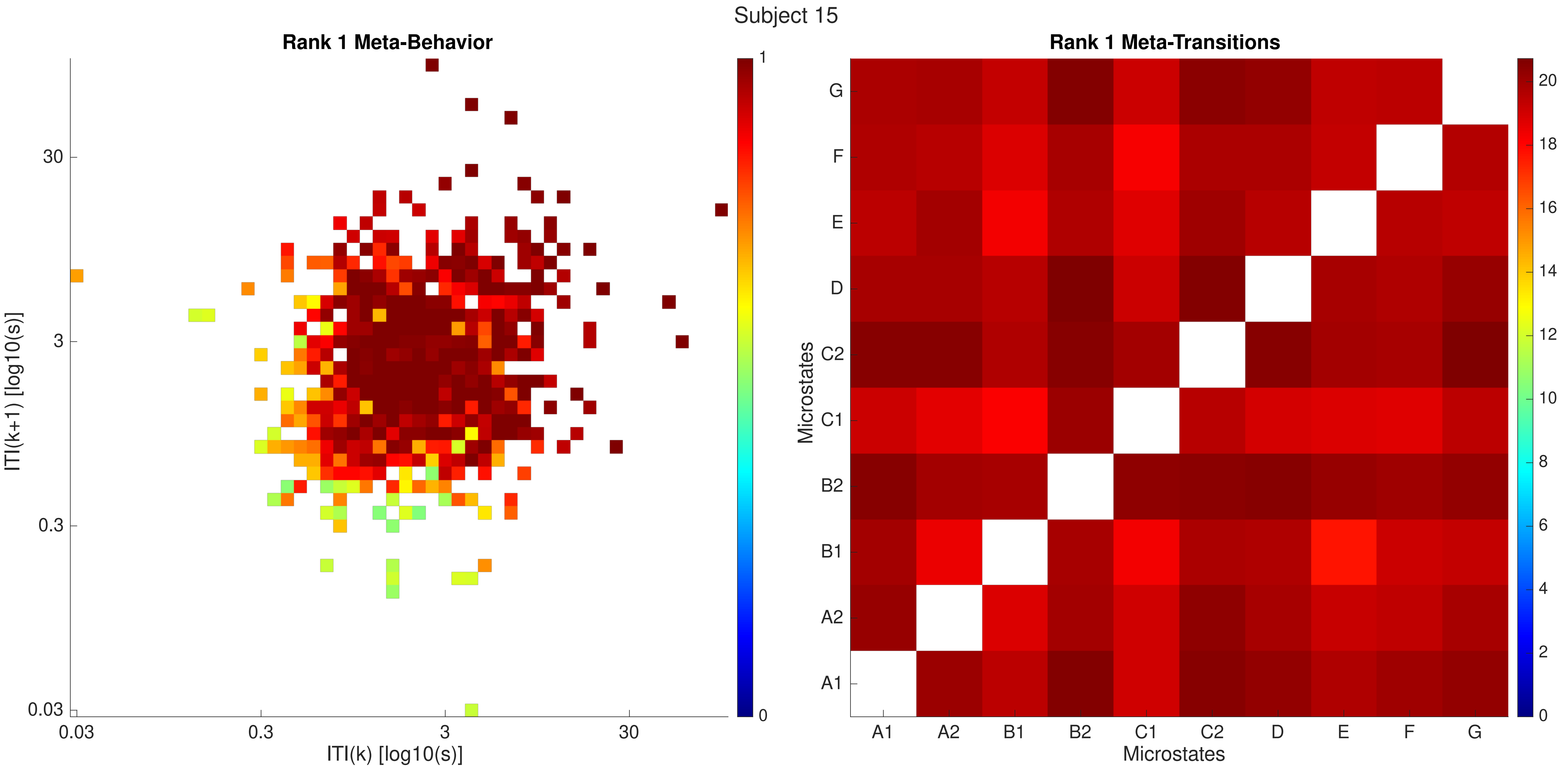
Subject 15
Rank 1 Meta-Behavior
ITI(k+1) [log10(s)]
ITI(k) [log10(s)]
0.03
0.3
3
30
0
1
Rank 1 Meta-Transitions
Microstates
A1
A2
B1
B2
C1
C2
D
E
F
G
0
2
4
6
8
10
12
14
16
18
20

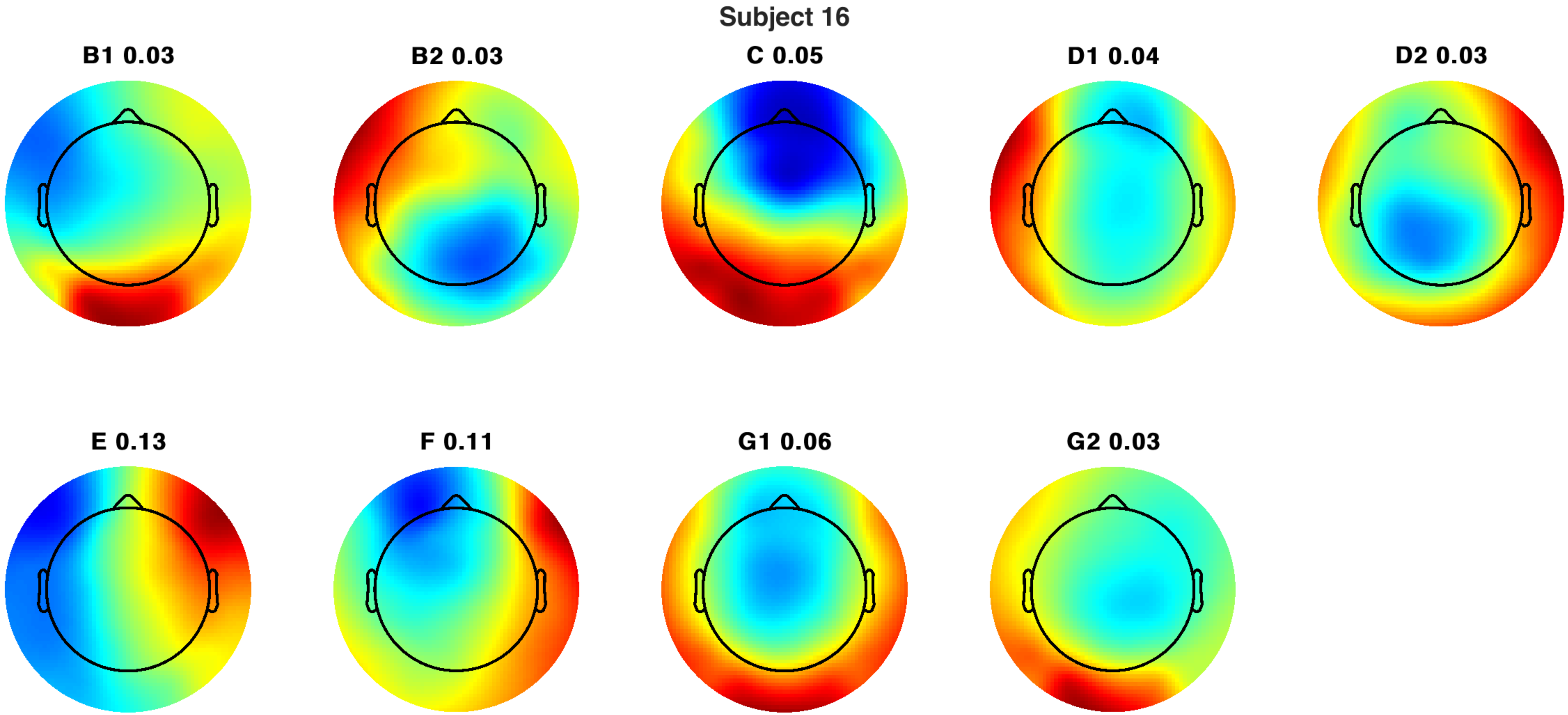
Subject 16
B1 0.03
B2 0.03
C 0.05
D1 0.04
D2 0.03
E 0.13
F 0.11
G1 0.06
G2 0.03

Subject 16

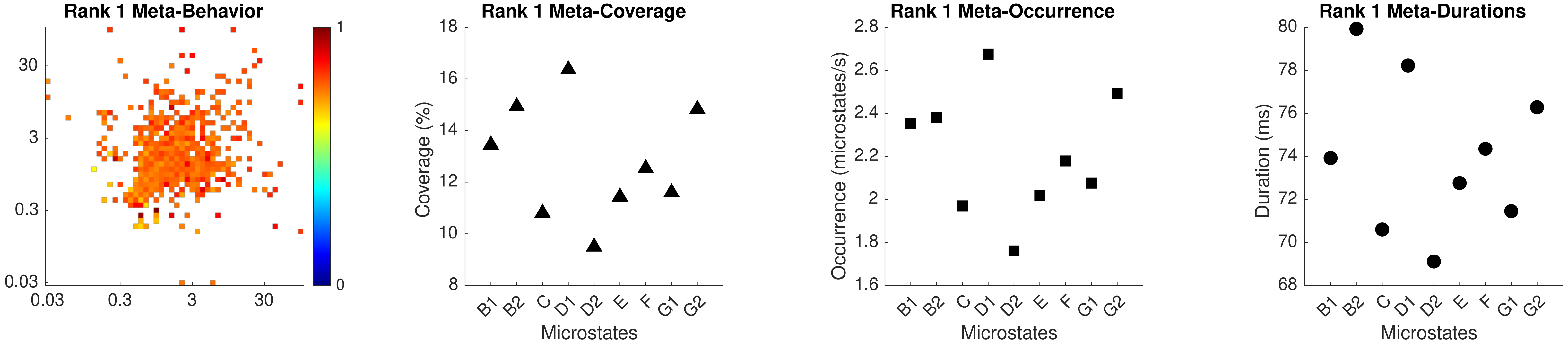

Rank 1 Meta-Behavior
30
3
0.3
0.03
0.03
0.3
3
30
1
0
Rank 1 Meta-Coverage
Coverage (%)
Microstates
B1
B2
C
D1
D2
E
F
G1
G2
Rank 1 Meta-Occurrence
Occurrence (microstates/s)
Microstates
Rank 1 Meta-Durations
Duration (ms)
Microstates

Subject 16

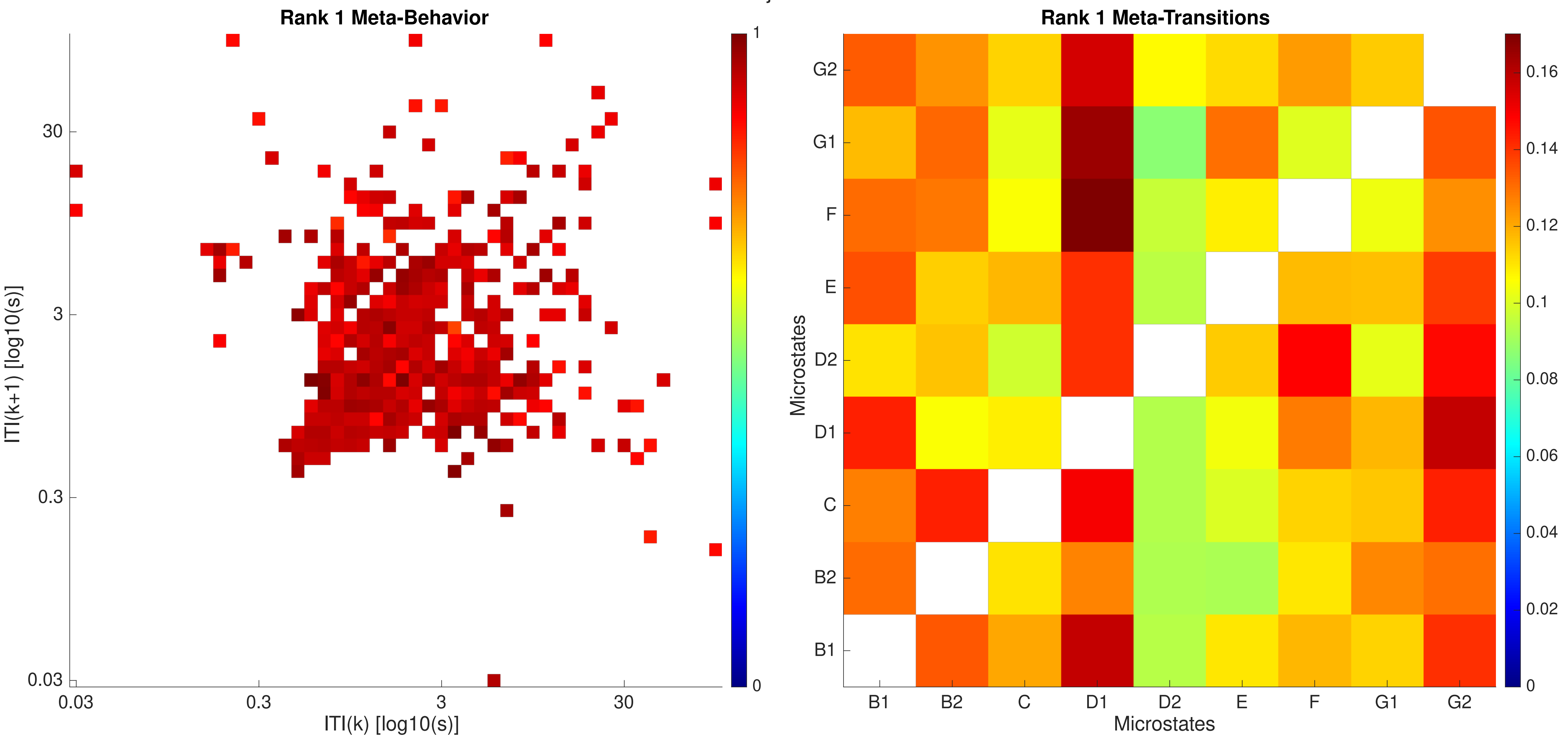

Rank 1 Meta-Behavior
ITI(k+1) [log10(s)]
30
3
0.3
0.03
0.03
0.3
3
30
ITI(k) [log10(s)]
1
0
Rank 1 Meta-Transitions
Microstates
G2
G1
F
E
D2
D1
C
B2
B1
B1
B2
C
D1
D2
E
F
G1
G2
Microstates
0.16
0.14
0.12
0.1
0.08
0.06
0.04
0.02
0

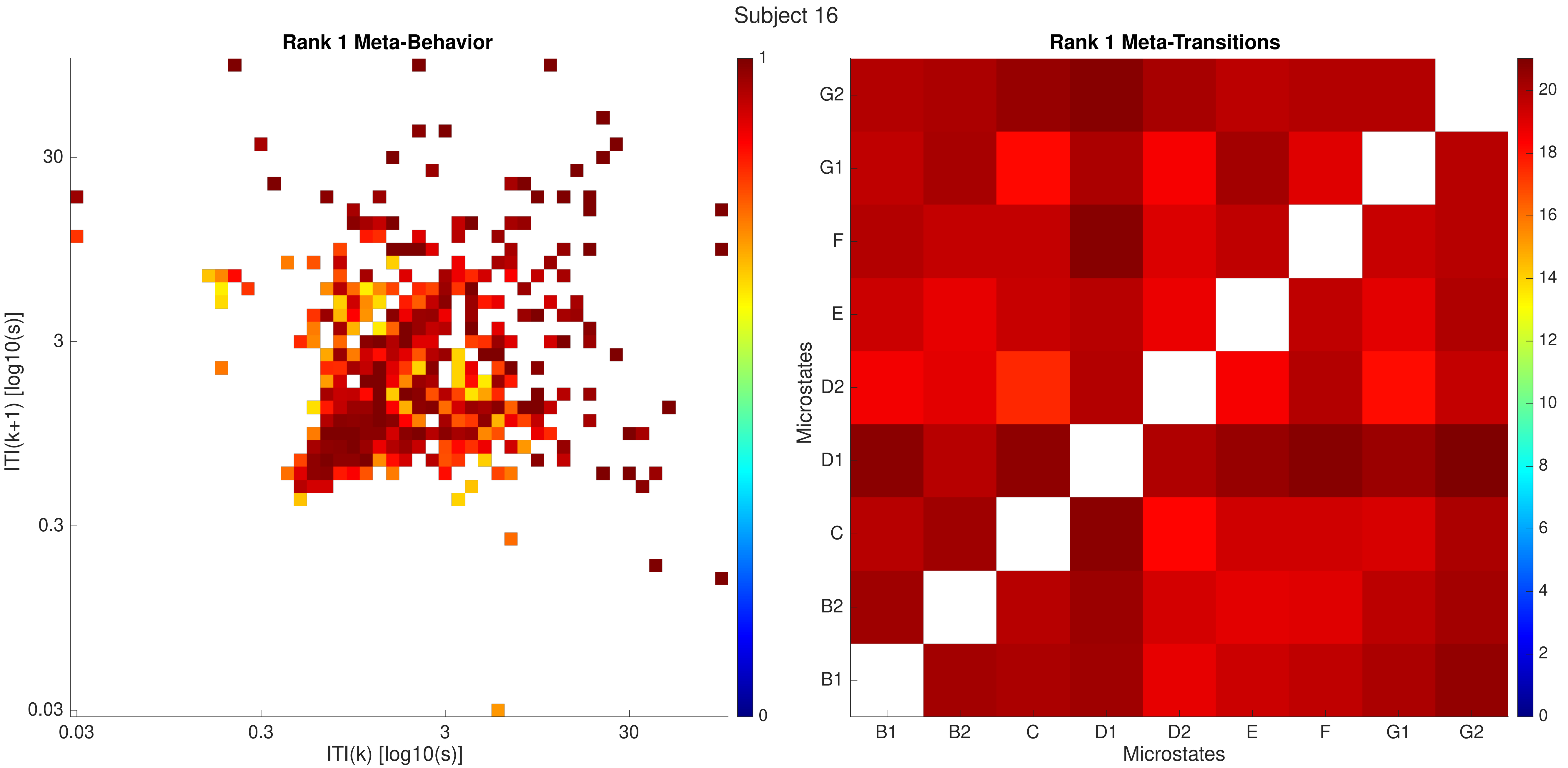
Subject 16
Rank 1 Meta-Behavior
ITI(k+1) [log10(s)]
ITI(k) [log10(s)]
0.03
0.3
3
30
0
1
Rank 1 Meta-Transitions
Microstates
B1
B2
C
D1
D2
E
F
G1
G2
0
2
4
6
8
10
12
14
16
18
20

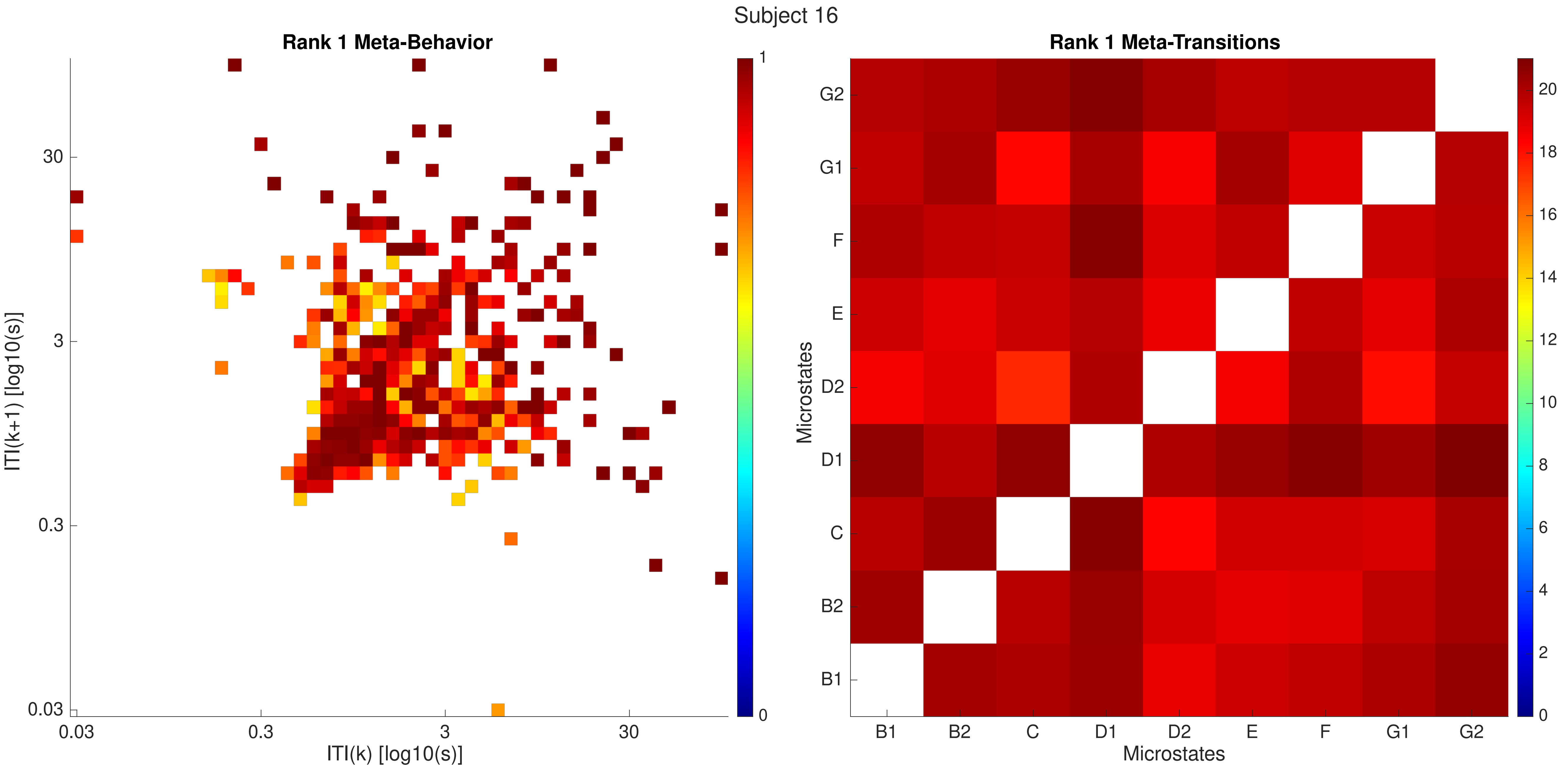
Subject 16
Rank 1 Meta-Behavior
ITI(k+1) [log10(s)]
30
3
0.3
0.03
0.03
0.3
3
30
ITI(k) [log10(s)]
1
0
Rank 1 Meta-Transitions
Microstates
G2
G1
F
E
D2
D1
C
B2
B1
B1
B2
C
D1
D2
E
F
G1
G2
Microstates
20
18
16
14
12
10
8
6
4
2
0

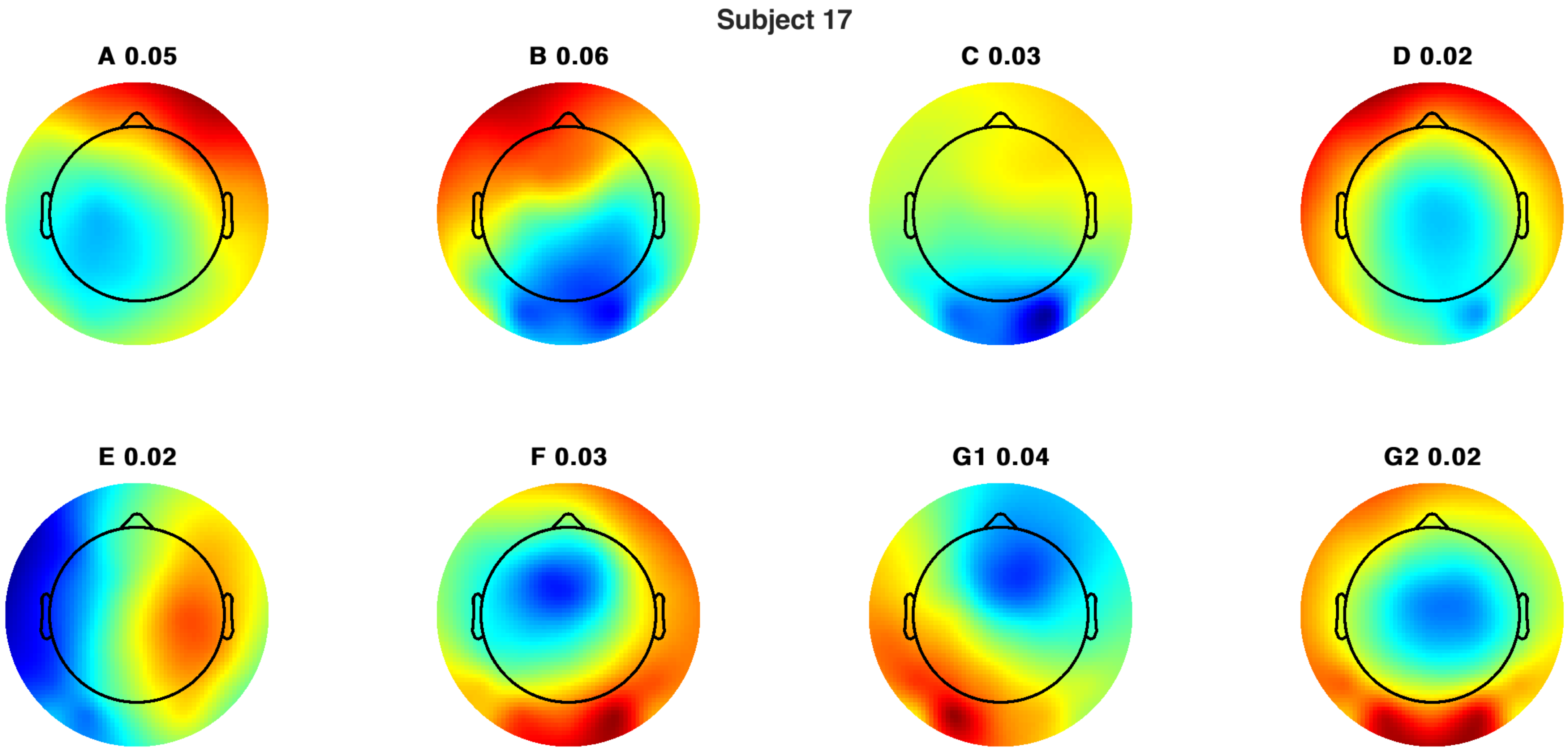
Subject 17
A 0.05
B 0.06
C 0.03
D 0.02
E 0.02
F 0.03
G1 0.04
G2 0.02

Subject 17

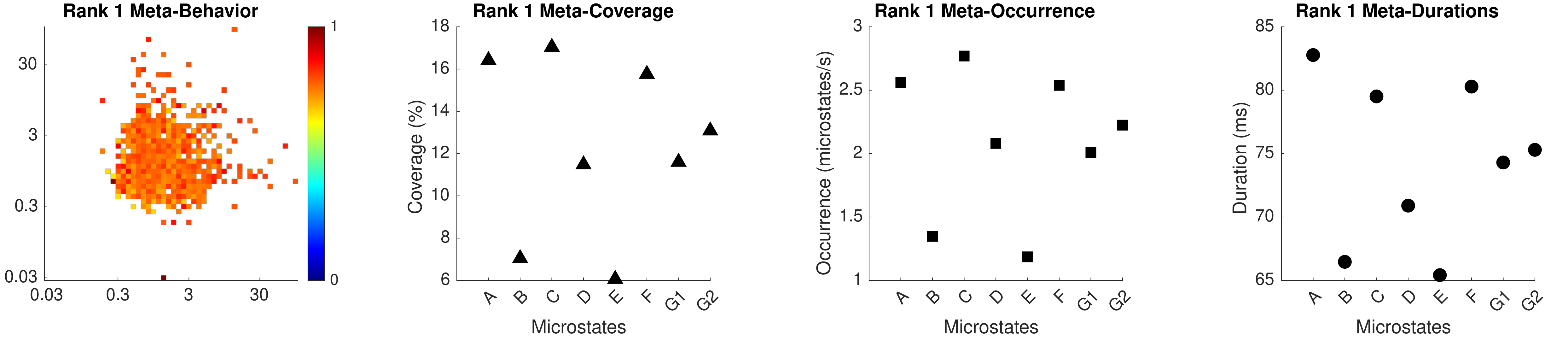

Rank 1 Meta-Behavior
30
3
0.3
0.03
0.03
0.3
3
30
1
0
Rank 1 Meta-Coverage
Coverage (%)
18
16
14
12
10
8
6
A
B
C
D
E
F
G1
G2
Microstates
Rank 1 Meta-Occurrence
Occurrence (microstates/s)
3
2.5
2
1.5
1
A
B
C
D
E
F
G1
G2
Microstates
Rank 1 Meta-Durations
Duration (ms)
85
80
75
70
65
A
B
C
D
E
F
G1
G2
Microstates

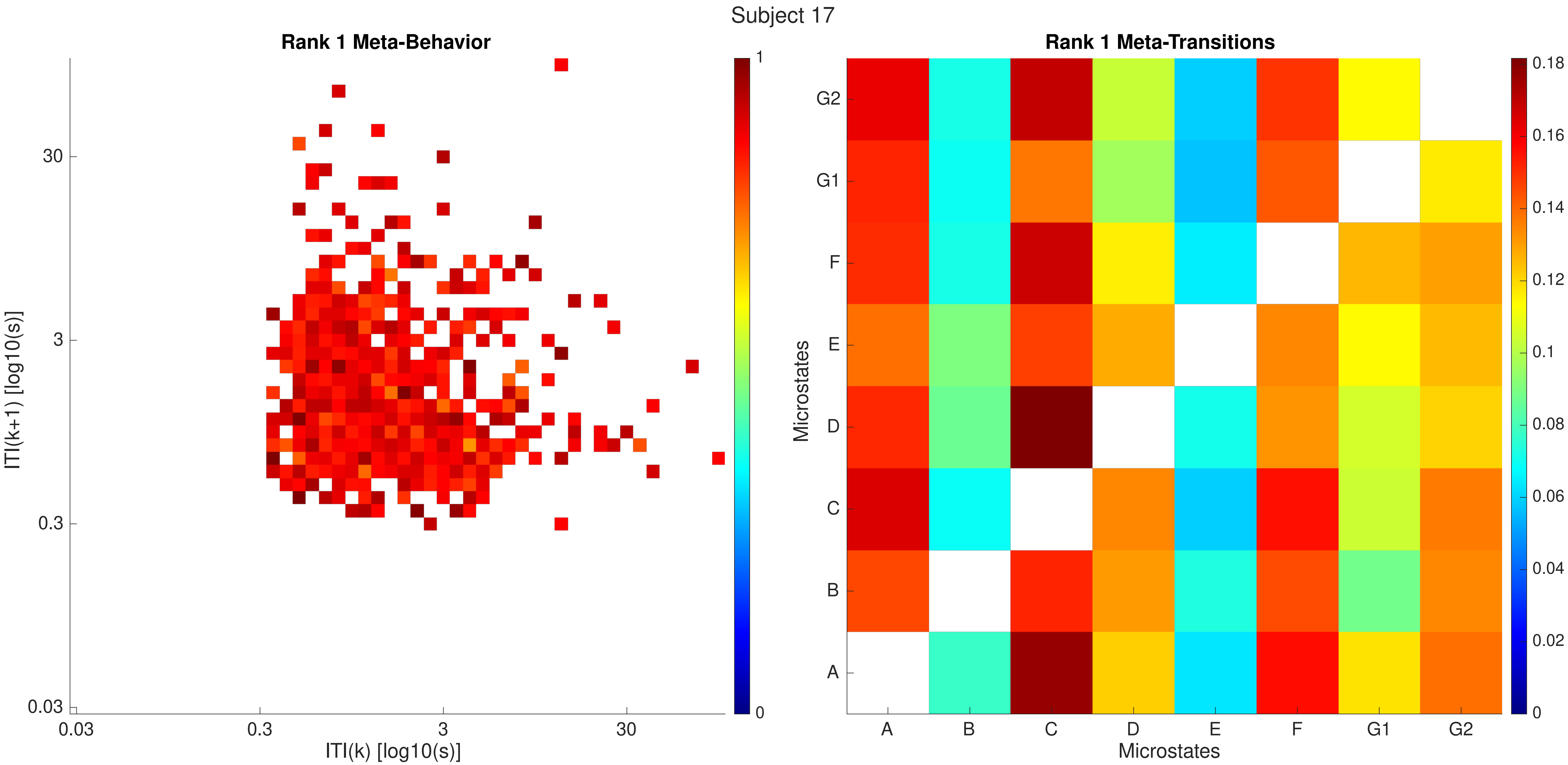
Subject 17
Rank 1 Meta-Behavior
ITI(k+1) [log10(s)]
30
3
0.3
0.03
ITI(k) [log10(s)]
0.03
0.3
3
30
1
0
Rank 1 Meta-Transitions
Microstates
G2
G1
F
E
D
C
B
A
Microstates
A
B
C
D
E
F
G1
G2
0.18
0.16
0.14
0.12
0.1
0.08
0.06
0.04
0.02
0

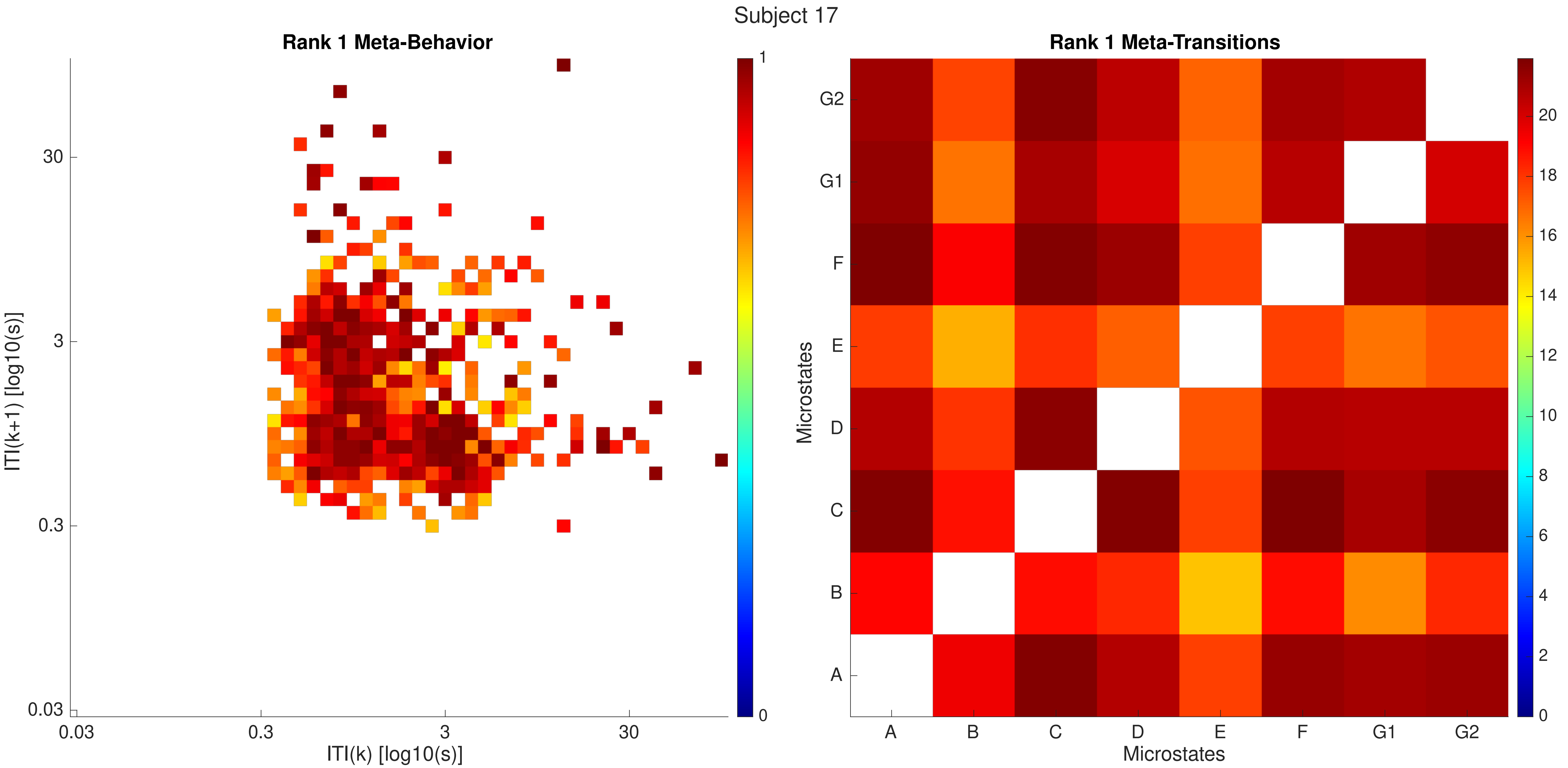

Subject 17
Rank 1 Meta-Behavior
ITI(k+1) [log10(s)]
ITI(k) [log10(s)]
0.03
0.3
3
30
0
1
Rank 1 Meta-Transitions
Microstates
A
B
C
D
E
F
G1
G2
0
2
4
6
8
10
12
14
16
18
20

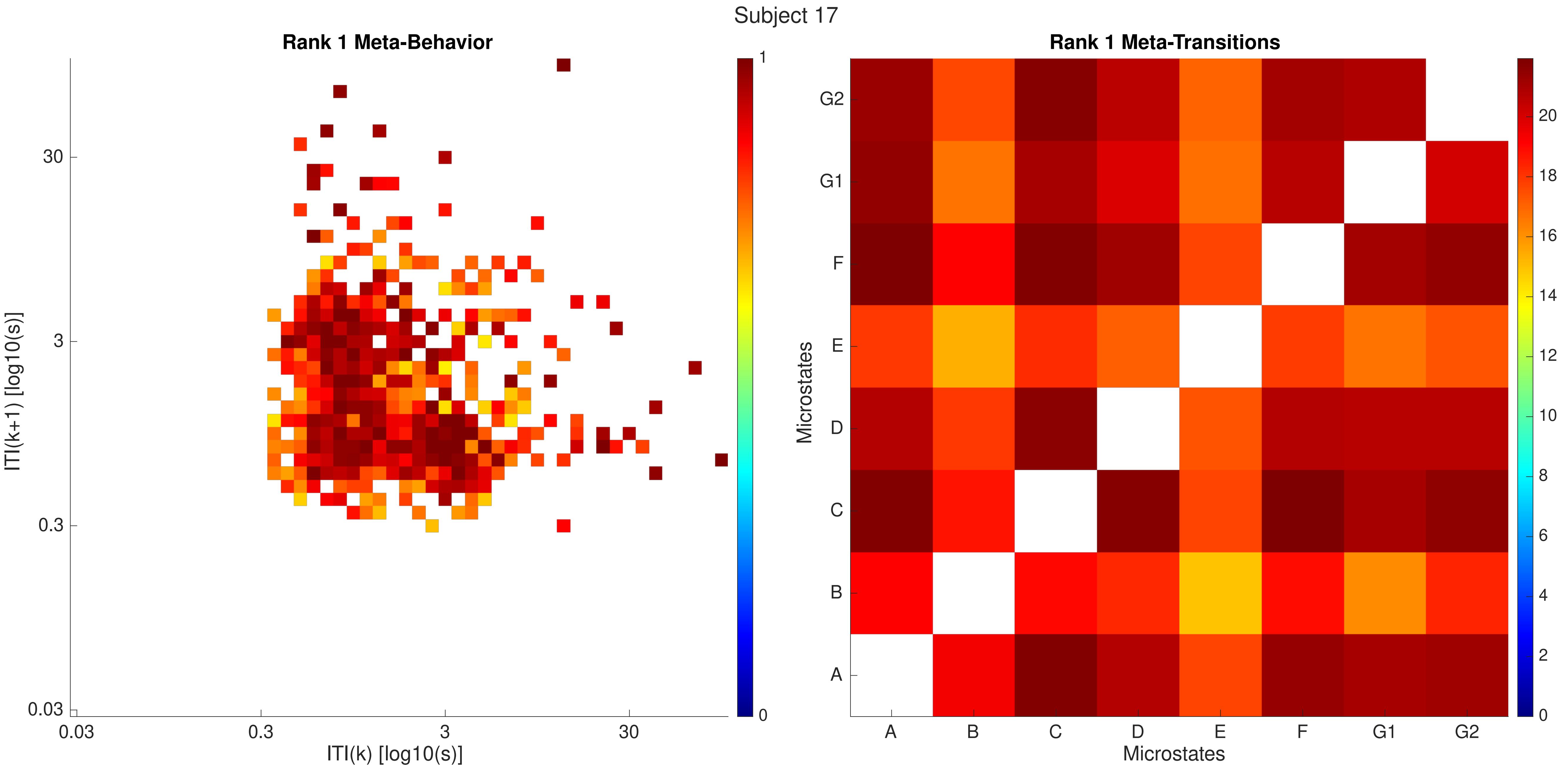

Subject 17
Rank 1 Meta-Behavior
ITI(k+1) [log10(s)]
ITI(k) [log10(s)]
0.03
0.3
3
30
1
0
Rank 1 Meta-Transitions
Microstates
A
B
C
D
E
F
G1
G2
0
2
4
6
8
10
12
14
16
18
20

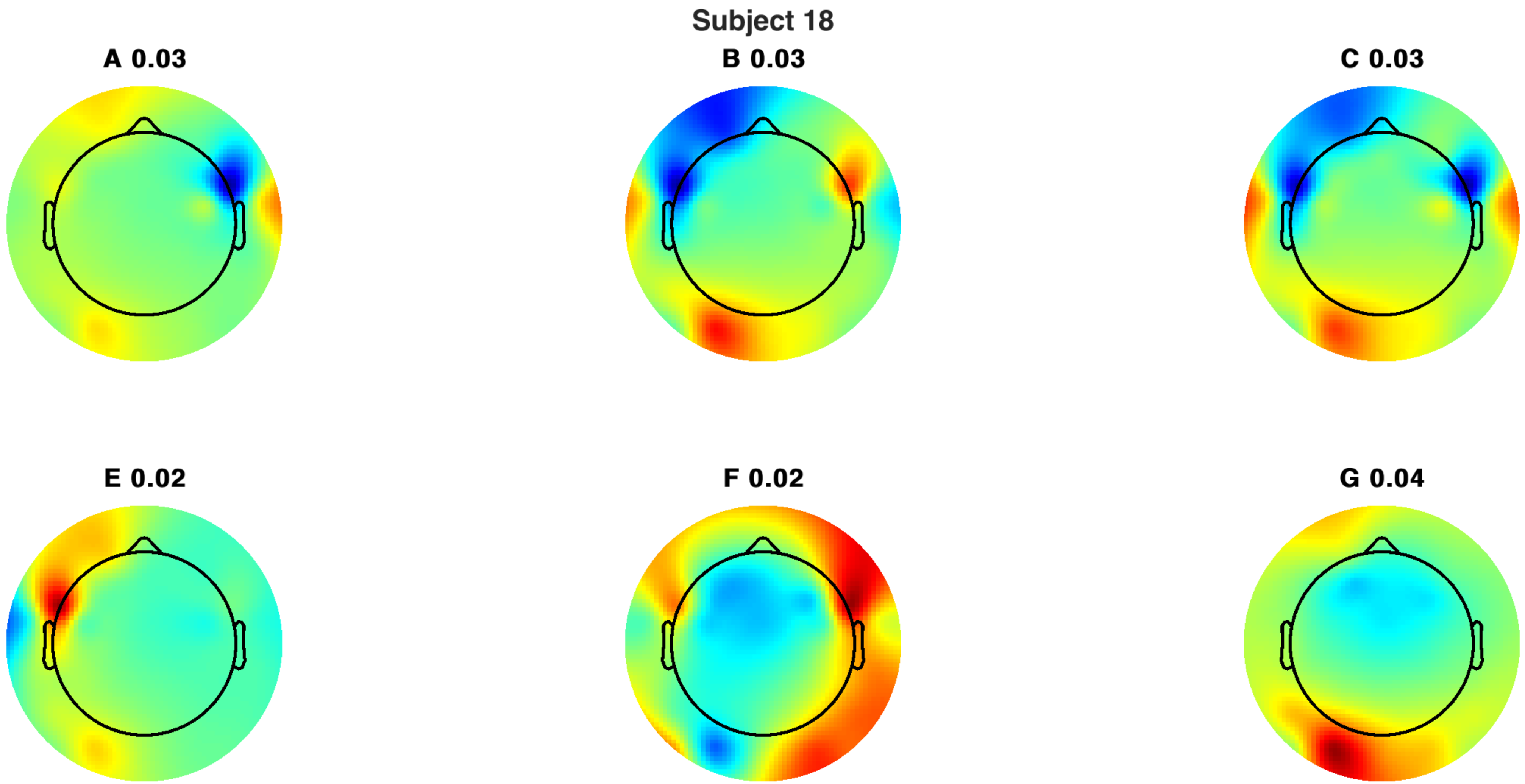
Subject 18
A 0.03
B 0.03
C 0.03
E 0.02
F 0.02
G 0.04

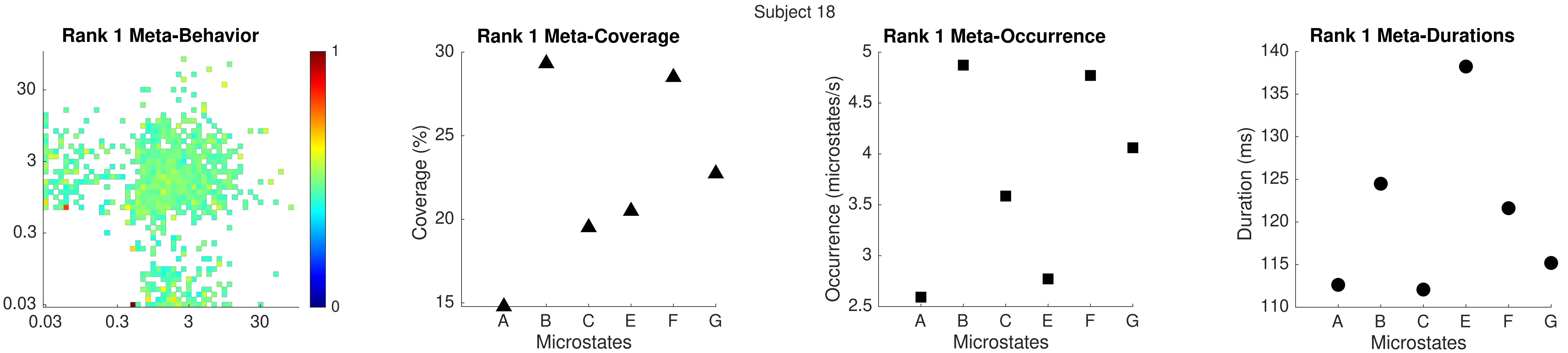

Subject 18
Rank 1 Meta-Behavior
30
3
0.3
0.03
0.03
0.3
3
30
1
0
Rank 1 Meta-Coverage
Coverage (%)
30
25
20
15
A
B
C
E
F
G
Microstates
Rank 1 Meta-Occurrence
Occurrence (microstates/s)
5
4.5
4
3.5
3
2.5
A
B
C
E
F
G
Microstates
Rank 1 Meta-Durations
Duration (ms)
140
135
130
125
120
115
110
A
B
C
E
F
G
Microstates

Subject 18

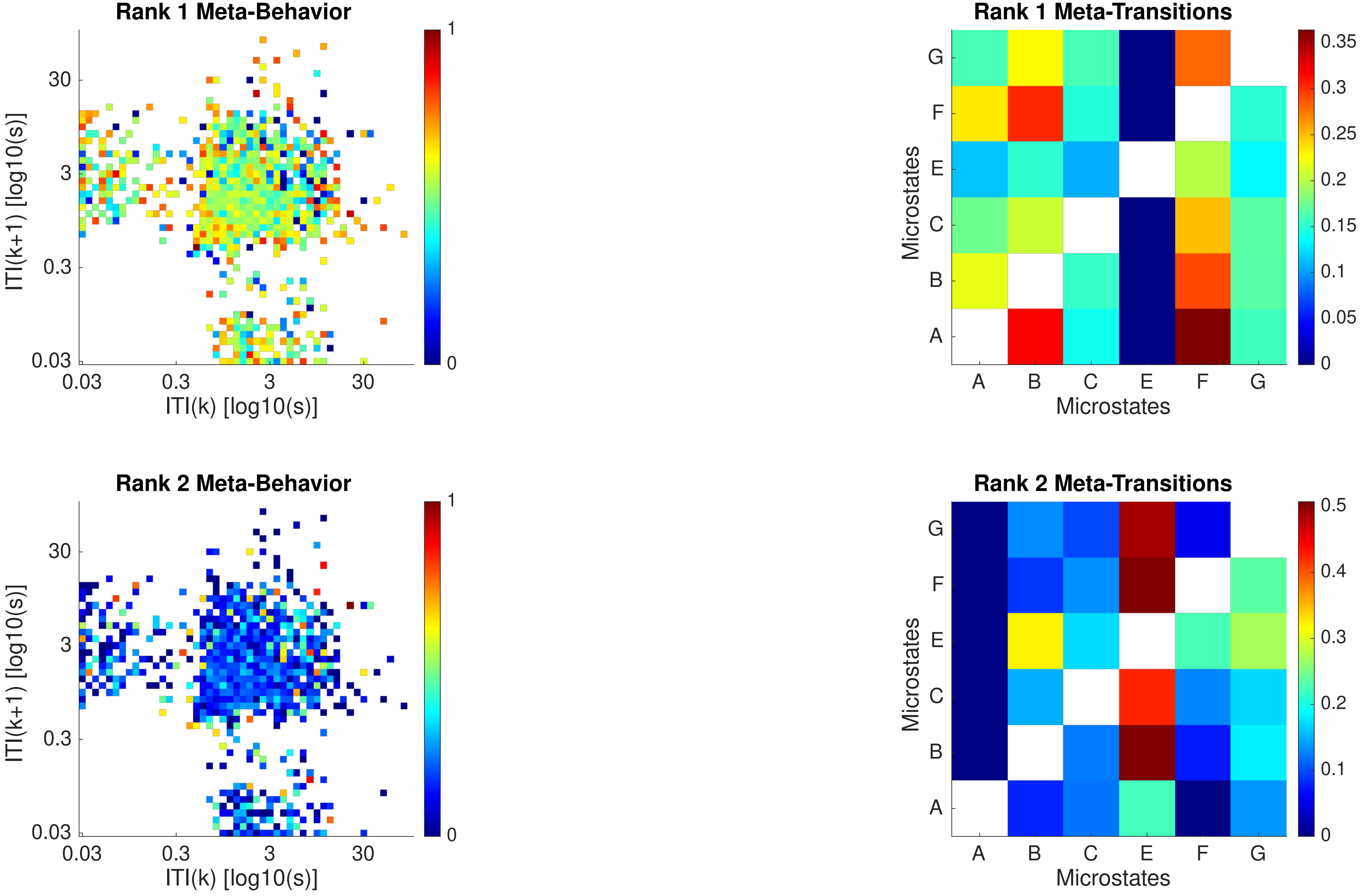
Rank 1 Meta-Behavior
ITI(k+1) [log10(s)]
ITI(k) [log10(s)]
0.03
0.3
3
30
1
0
Rank 1 Meta-Transitions
Microstates
A
B
C
E
F
G
0.35
0.3
0.25
0.2
0.15
0.1
0.05
0
Rank 2 Meta-Behavior
ITI(k+1) [log10(s)]
ITI(k) [log10(s)]
Rank 2 Meta-Transitions
Microstates
0.5
0.4
0.3
0.2
0.1
0

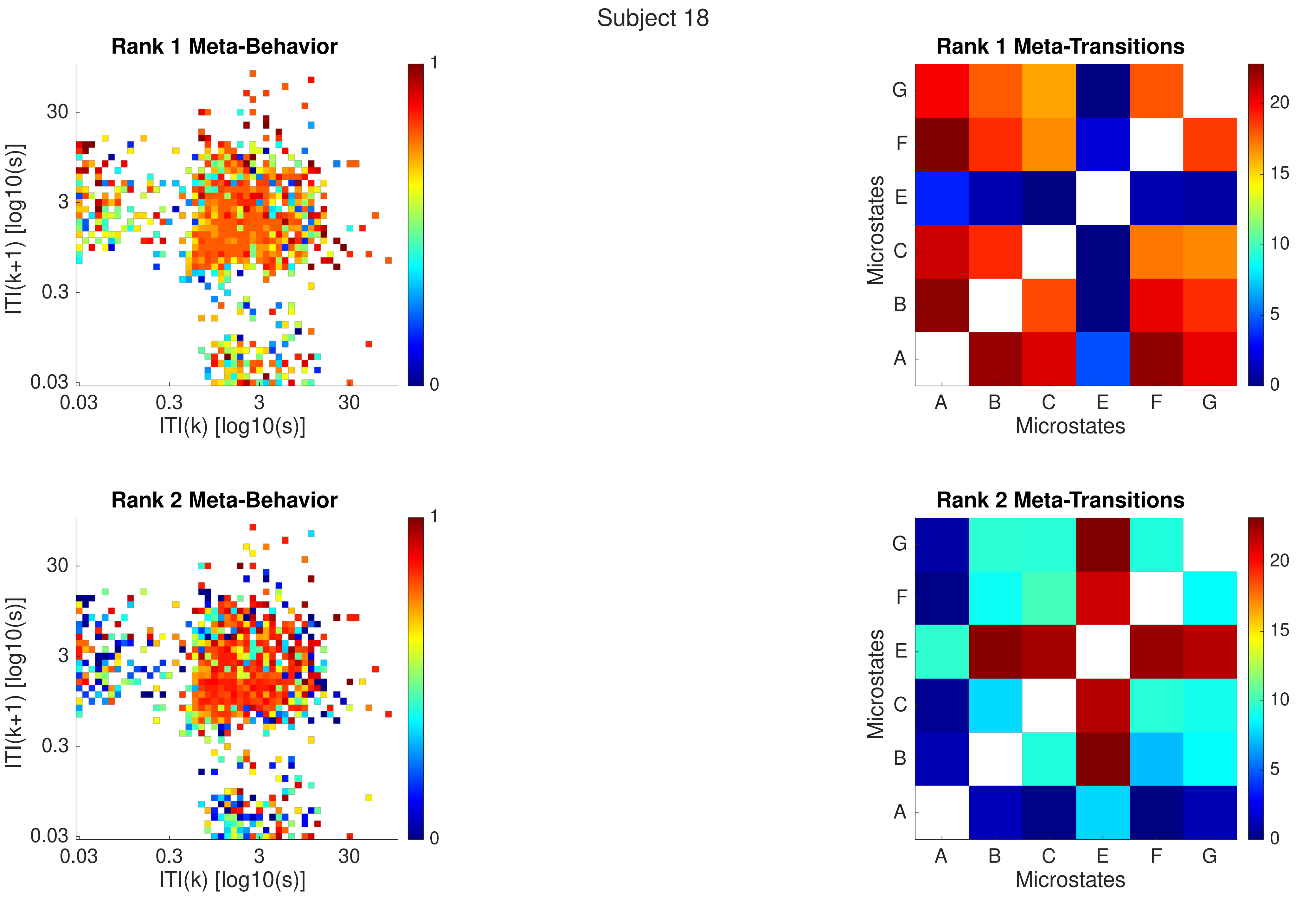

Subject 18
Rank 1 Meta-Behavior
ITI(k+1) [log10(s)]
ITI(k) [log10(s)]
30
3
0.3
0.03
0.03
0.3
3
30
1
0
Rank 1 Meta-Transitions
Microstates
G
F
E
C
B
A
A
B
C
E
F
G
Microstates
20
15
10
5
0
Rank 2 Meta-Behavior
ITI(k+1) [log10(s)]
ITI(k) [log10(s)]
Rank 2 Meta-Transitions
Microstates
Microstates

Subject 18

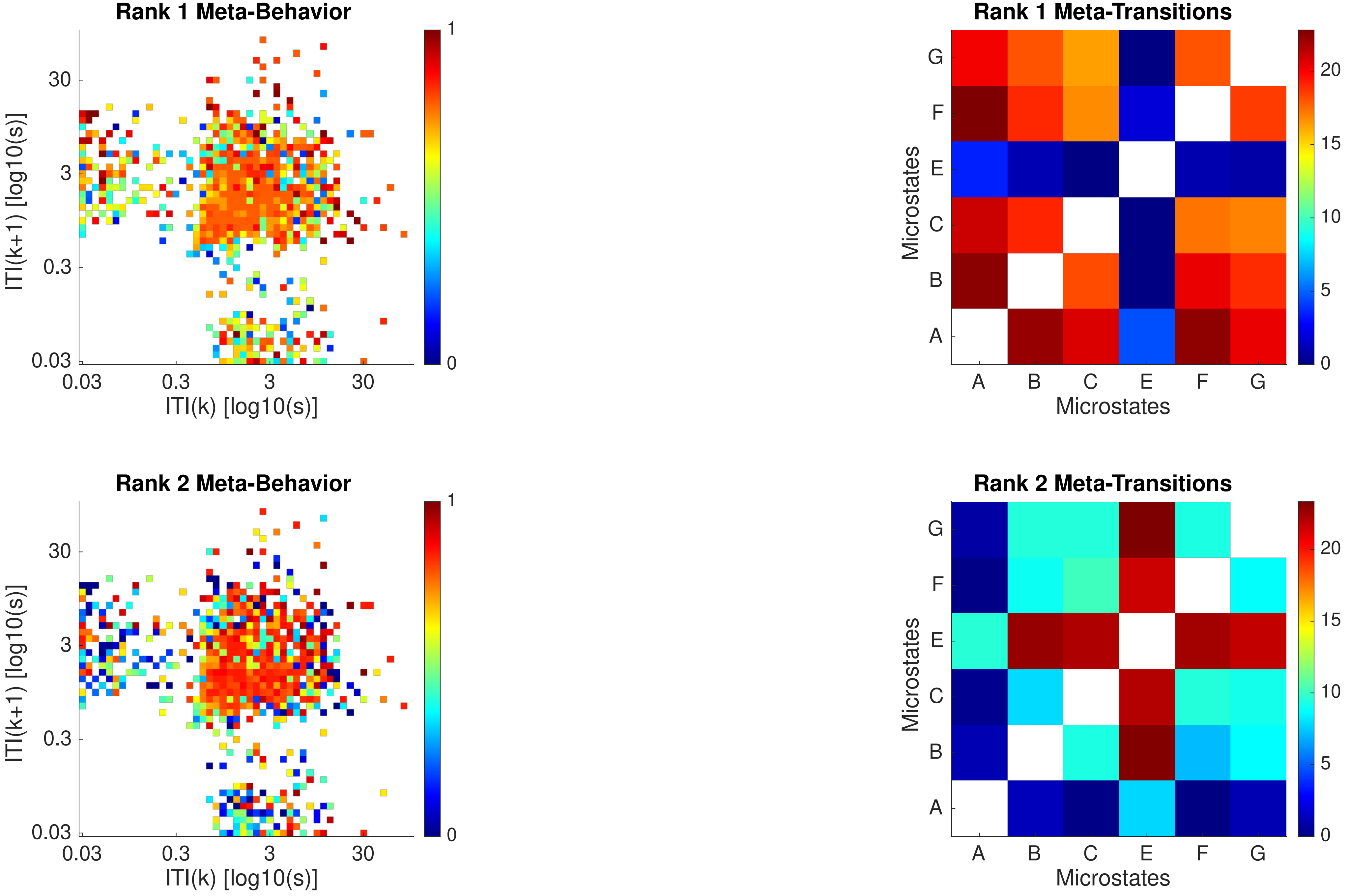
Rank 1 Meta-Behavior
ITI(k+1) [log10(s)]
ITI(k) [log10(s)]
0.03
0.3
3
30
0
1
Rank 1 Meta-Transitions
Microstates
Microstates
A
B
C
E
F
G
0
5
10
15
20
Rank 2 Meta-Behavior
ITI(k+1) [log10(s)]
ITI(k) [log10(s)]
0.03
0.3
3
30
0
1
Rank 2 Meta-Transitions
Microstates
Microstates
A
B
C
E
F
G
0
5
10
15
20

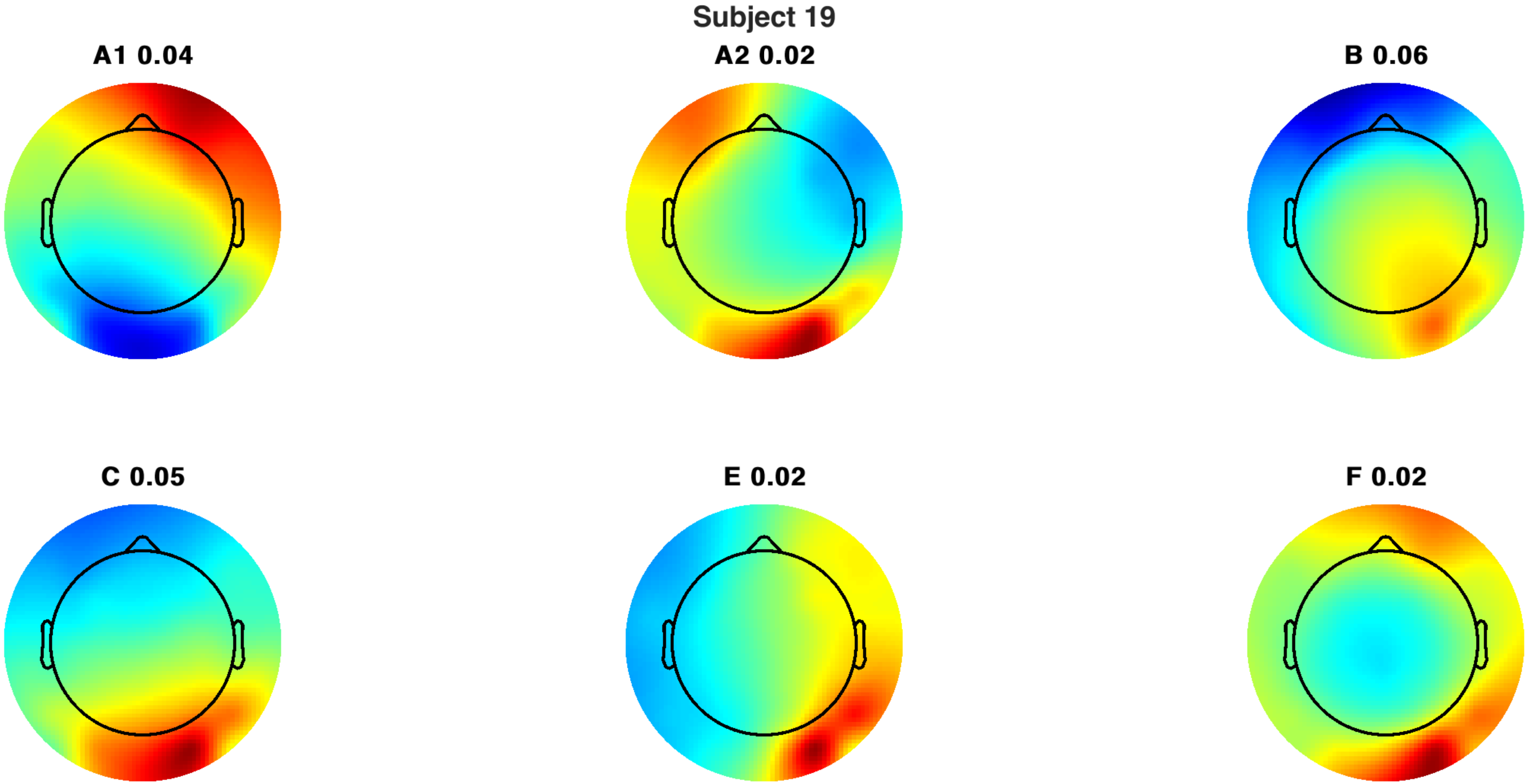
Subject 19
A1 0.04
A2 0.02
B 0.06
C 0.05
E 0.02
F 0.02

Subject 19

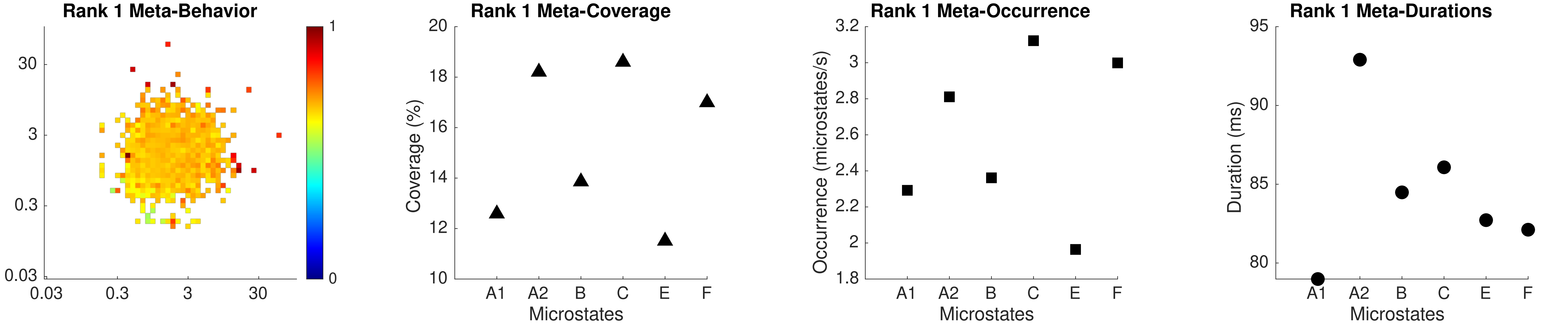

Rank 1 Meta-Behavior
30
3
0.3
0.03
0.03
0.3
3
30
1
0
Rank 1 Meta-Coverage
Coverage (%)
20
18
16
14
12
10
A1
A2
B
C
E
F
Microstates
Rank 1 Meta-Occurrence
Occurrence (microstates/s)
3.2
3
2.8
2.6
2.4
2.2
2
1.8
A1
A2
B
C
E
F
Microstates
Rank 1 Meta-Durations
Duration (ms)
95
90
85
80
A1
A2
B
C
E
F
Microstates

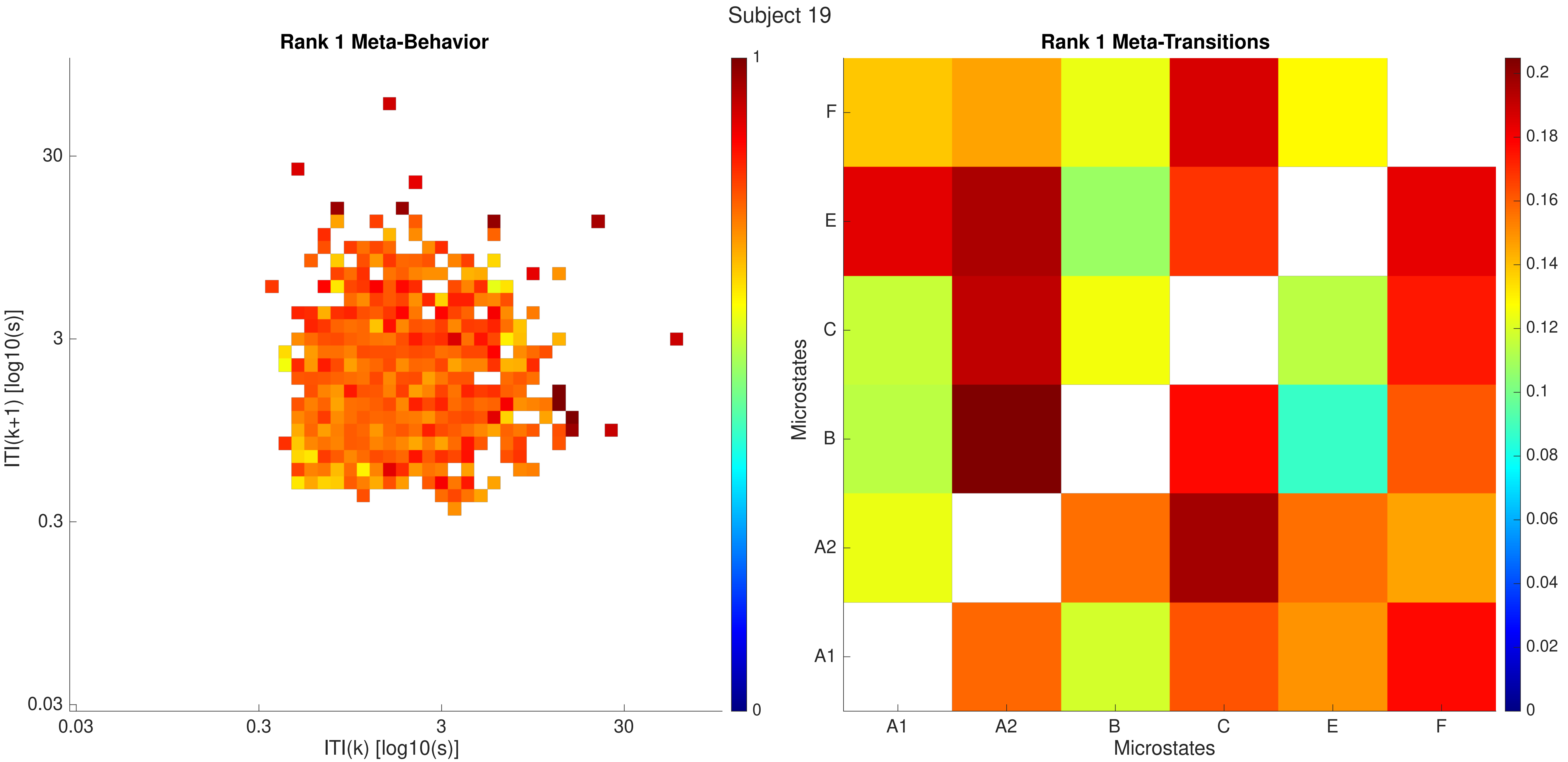
Subject 19
Rank 1 Meta-Behavior
ITI(k+1) [log10(s)]
30
3
0.3
0.03
0.03
0.3
3
30
ITI(k) [log10(s)]
1
0
Rank 1 Meta-Transitions
Microstates
F
E
C
B
A2
A1
A1
A2
B
C
E
F
Microstates
0.2
0.18
0.16
0.14
0.12
0.1
0.08
0.06
0.04
0.02
0

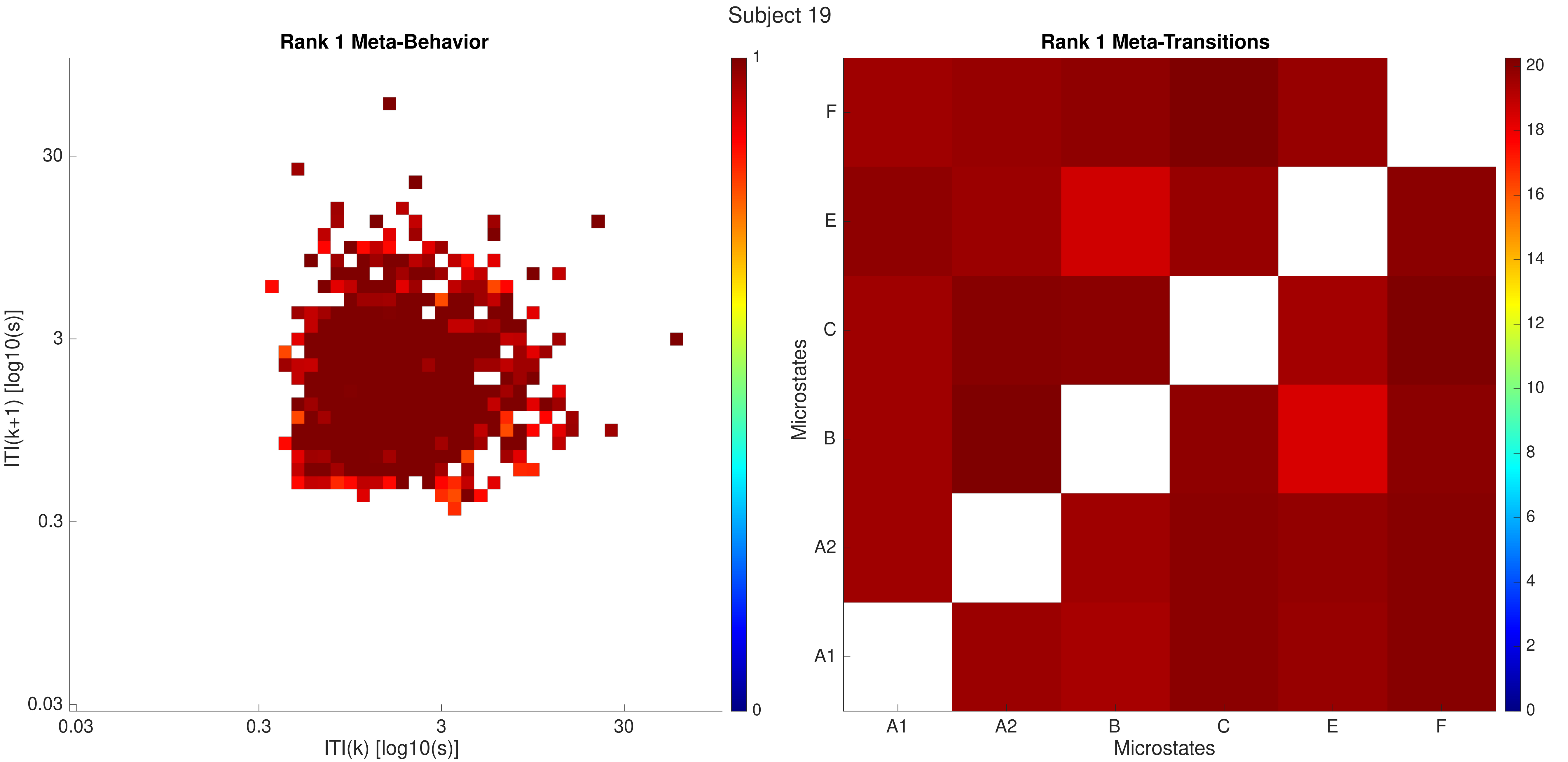

Subject 19
Rank 1 Meta-Behavior
ITI(k+1) [log10(s)]
30
3
0.3
0.03
ITI(k) [log10(s)]
0.03
0.3
3
30
1
0
Rank 1 Meta-Transitions
Microstates
F
E
C
B
A2
A1
Microstates
A1
A2
B
C
E
F
20
18
16
14
12
10
8
6
4
2
0

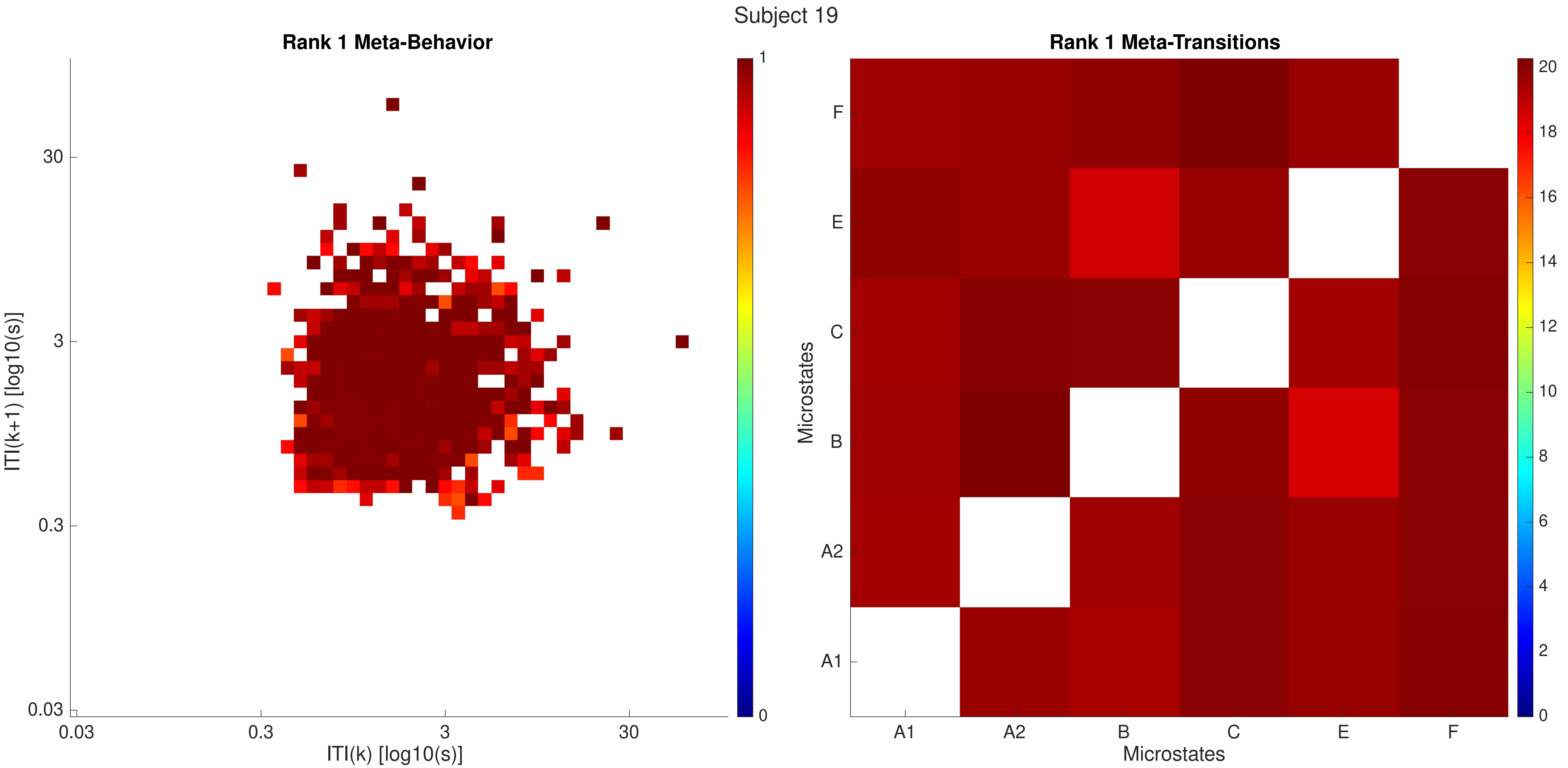
Subject 19
Rank 1 Meta-Behavior
30
3
0.3
0.03
ITI(k+1) [log10(s)]
0.03
0.3
3
30
ITI(k) [log10(s)]
1
0
Rank 1 Meta-Transitions
F
E
C
B
A2
A1
Microstates
A1
A2
B
C
E
F
Microstates
20
18
16
14
12
10
8
6
4
2
0

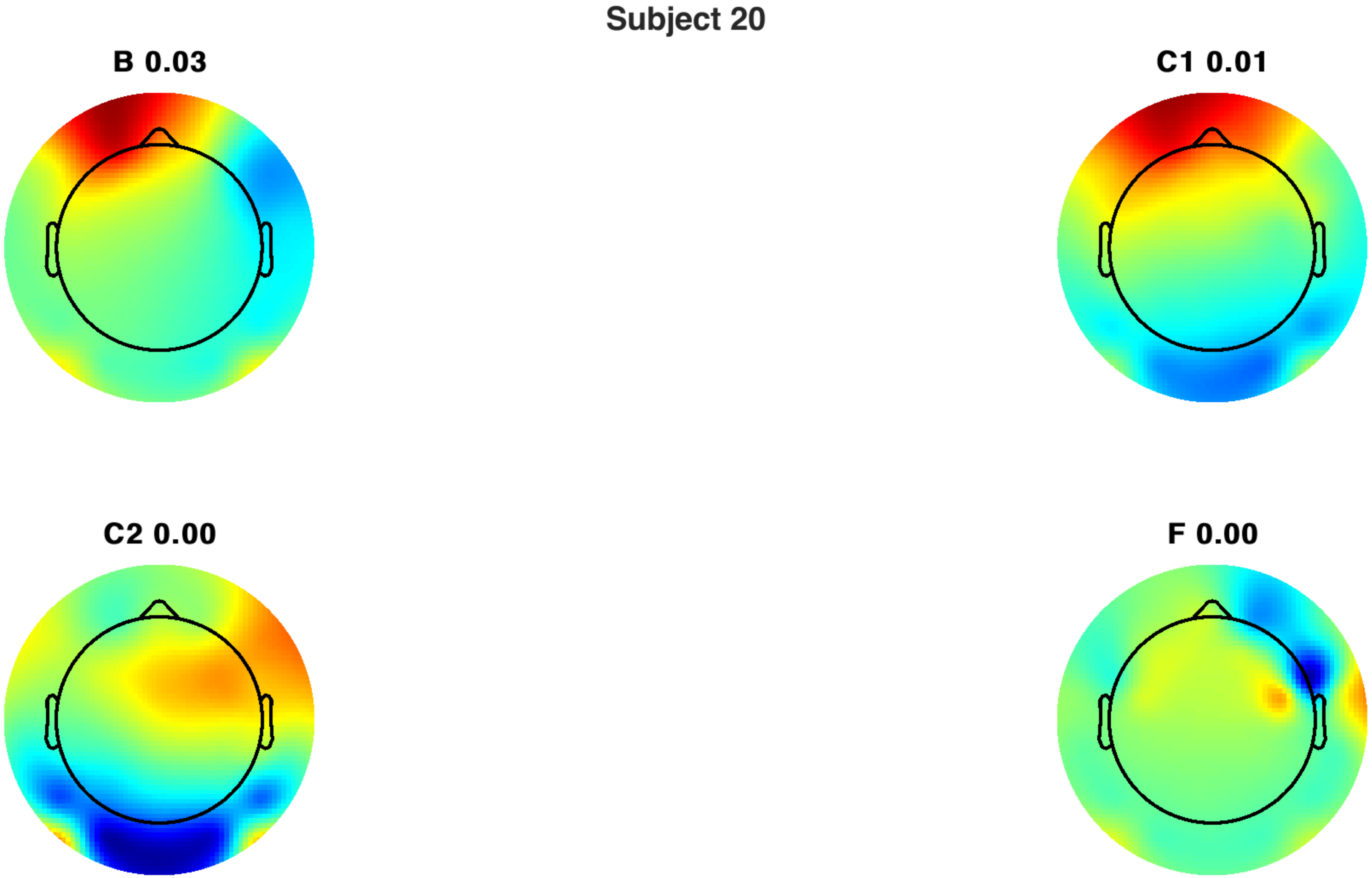
Subject 20
B 0.03
C1 0.01
C2 0.00
F 0.00

Subject 20

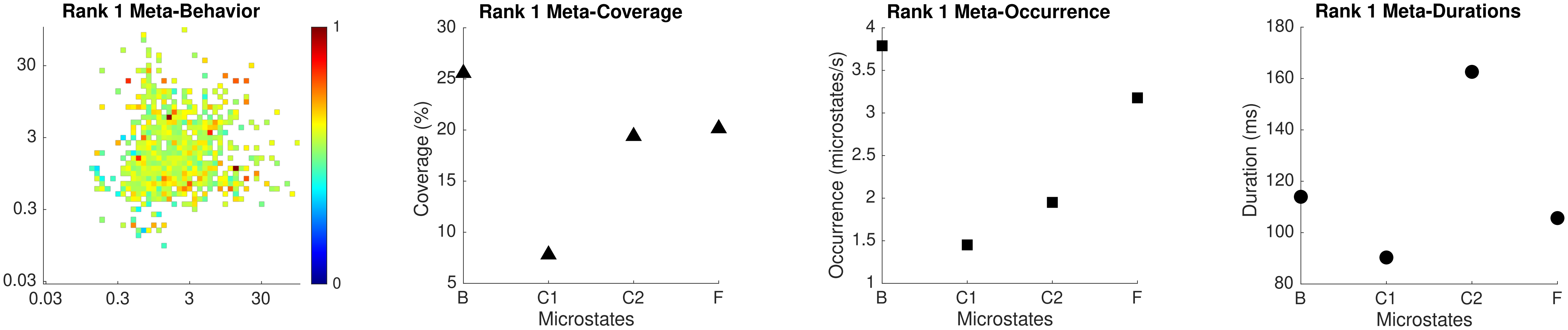

Rank 1 Meta-Behavior
30
3
0.3
0.03
0.03
0.3
3
30
1
0
Rank 1 Meta-Coverage
Coverage (%)
Microstates
B
C1
C2
F
Rank 1 Meta-Occurrence
Occurrence (microstates/s)
Microstates
Rank 1 Meta-Durations
Duration (ms)
Microstates

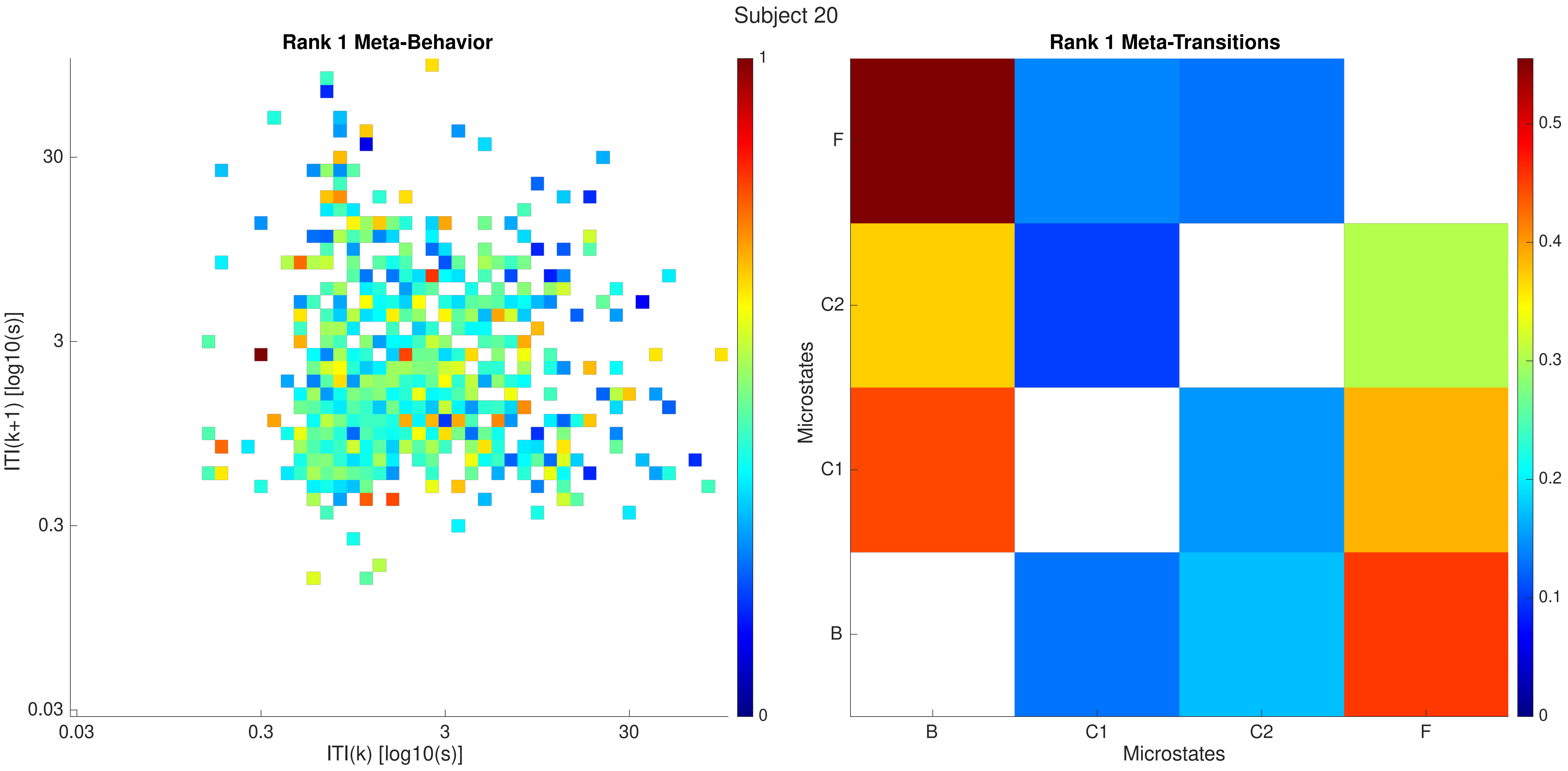

Subject 20
Rank 1 Meta-Behavior
ITI(k+1) [log10(s)]
ITI(k) [log10(s)]
0.03
0.3
3
30
0
1
Rank 1 Meta-Transitions
Microstates
B
C1
C2
F
0
0.1
0.2
0.3
0.4
0.5

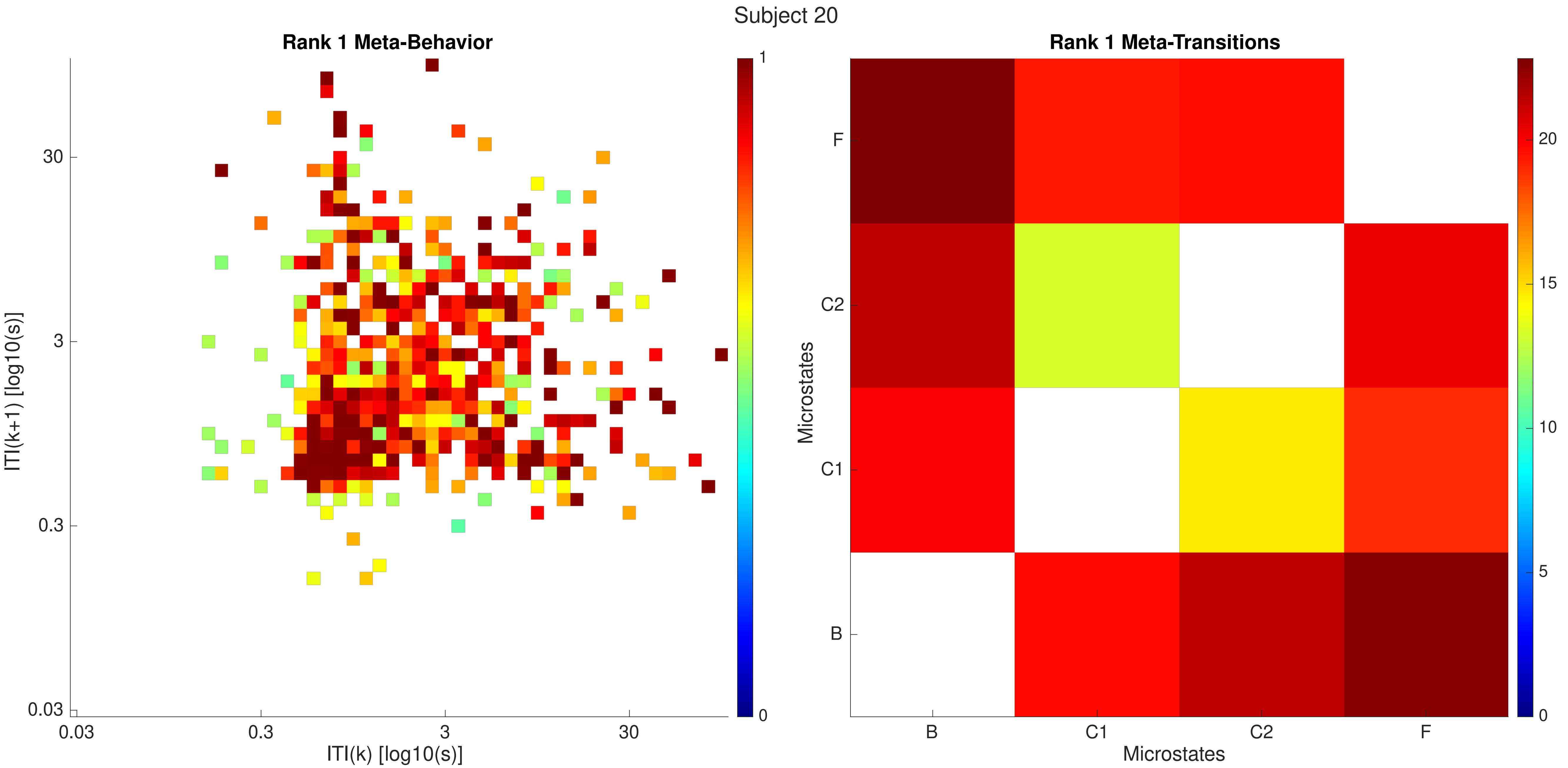
Subject 20
Rank 1 Meta-Behavior
ITI(k+1) [log10(s)]
ITI(k) [log10(s)]
0.03
0.3
3
30
0
1
Rank 1 Meta-Transitions
Microstates
B
C1
C2
F
Microstates
0
5
10
15
20

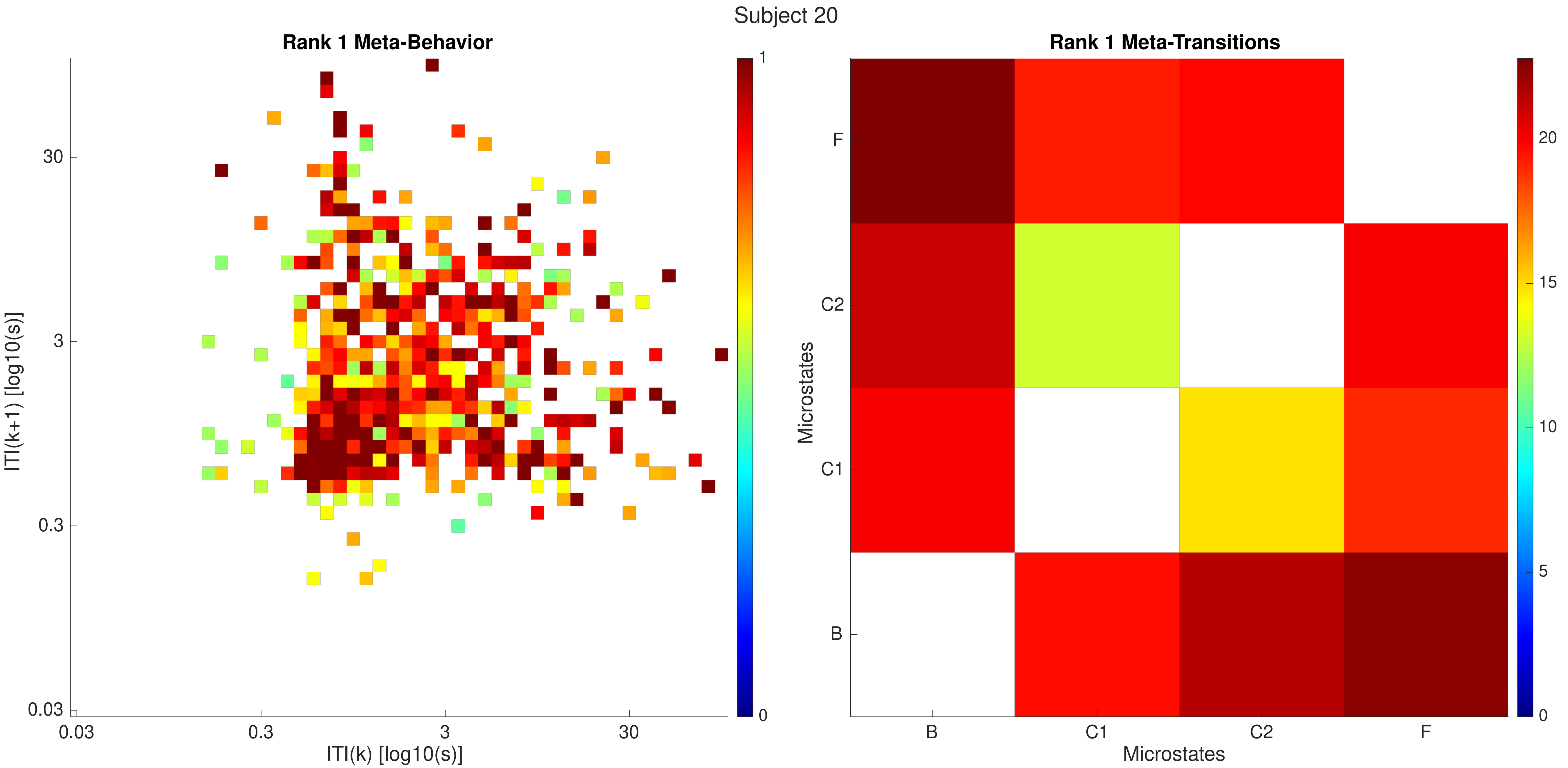

Subject 20
Rank 1 Meta-Behavior
ITI(k+1) [log10(s)]
ITI(k) [log10(s)]
0.03
0.3
3
30
0
1
Rank 1 Meta-Transitions
Microstates
B
C1
C2
F
0
5
10
15
20

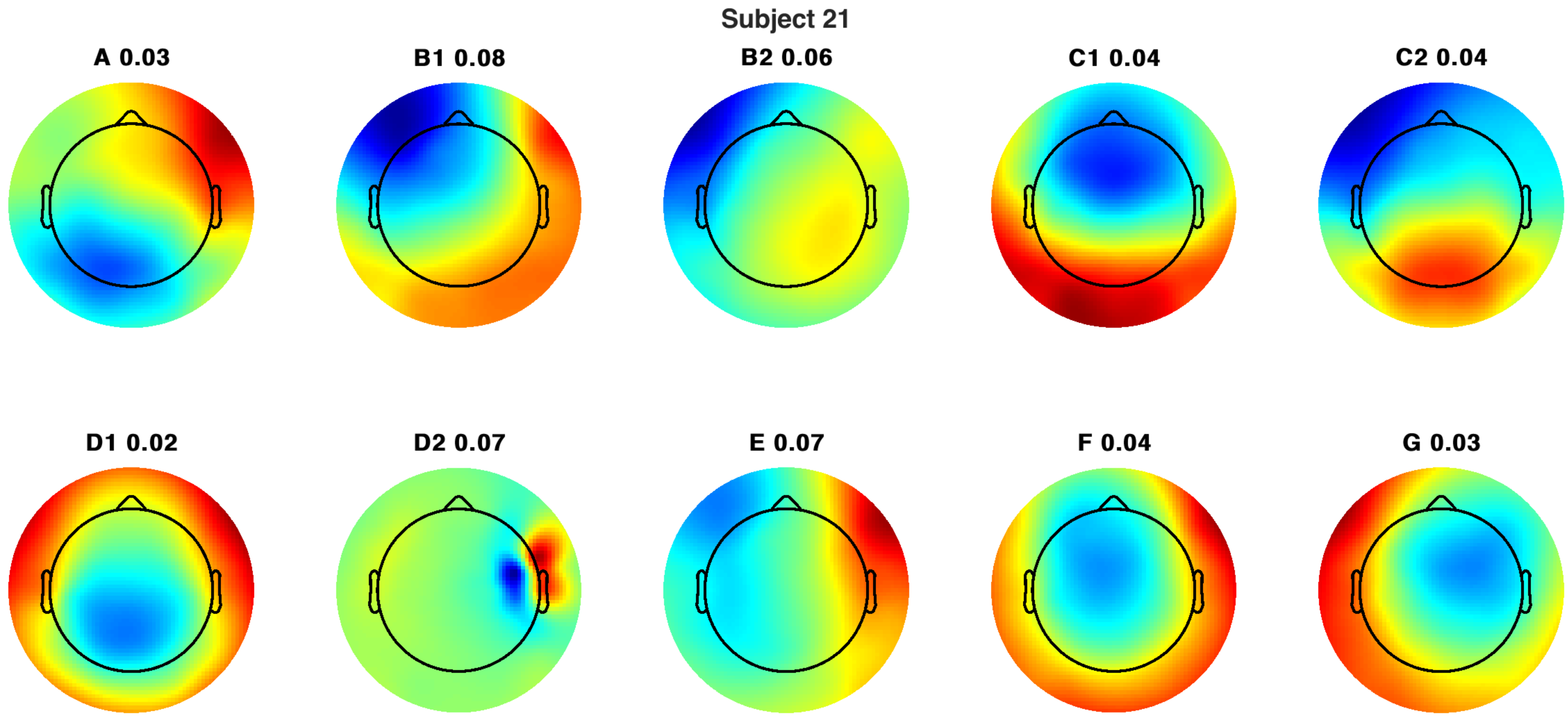
Subject 21
A 0.03
B1 0.08
B2 0.06
C1 0.04
C2 0.04
D1 0.02
D2 0.07
E 0.07
F 0.04
G 0.03

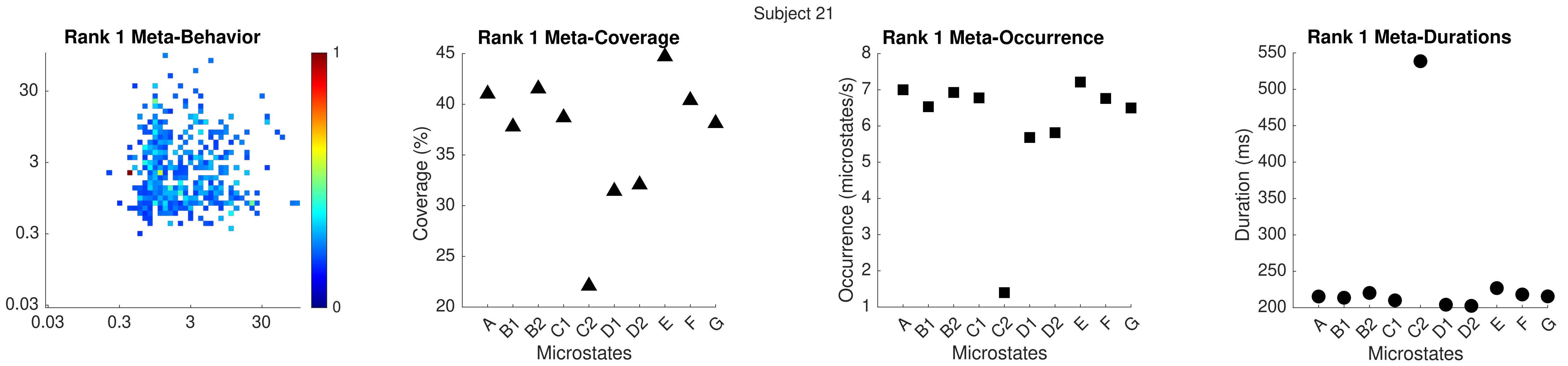
Subject 21
Rank 1 Meta-Behavior
Rank 1 Meta-Coverage
Coverage (%)
Microstates
Rank 1 Meta-Occurrence
Occurrence (microstates/s)
Microstates
Rank 1 Meta-Durations
Duration (ms)
Microstates
A
B1
B2
C1
C2
D1
D2
E
F
G

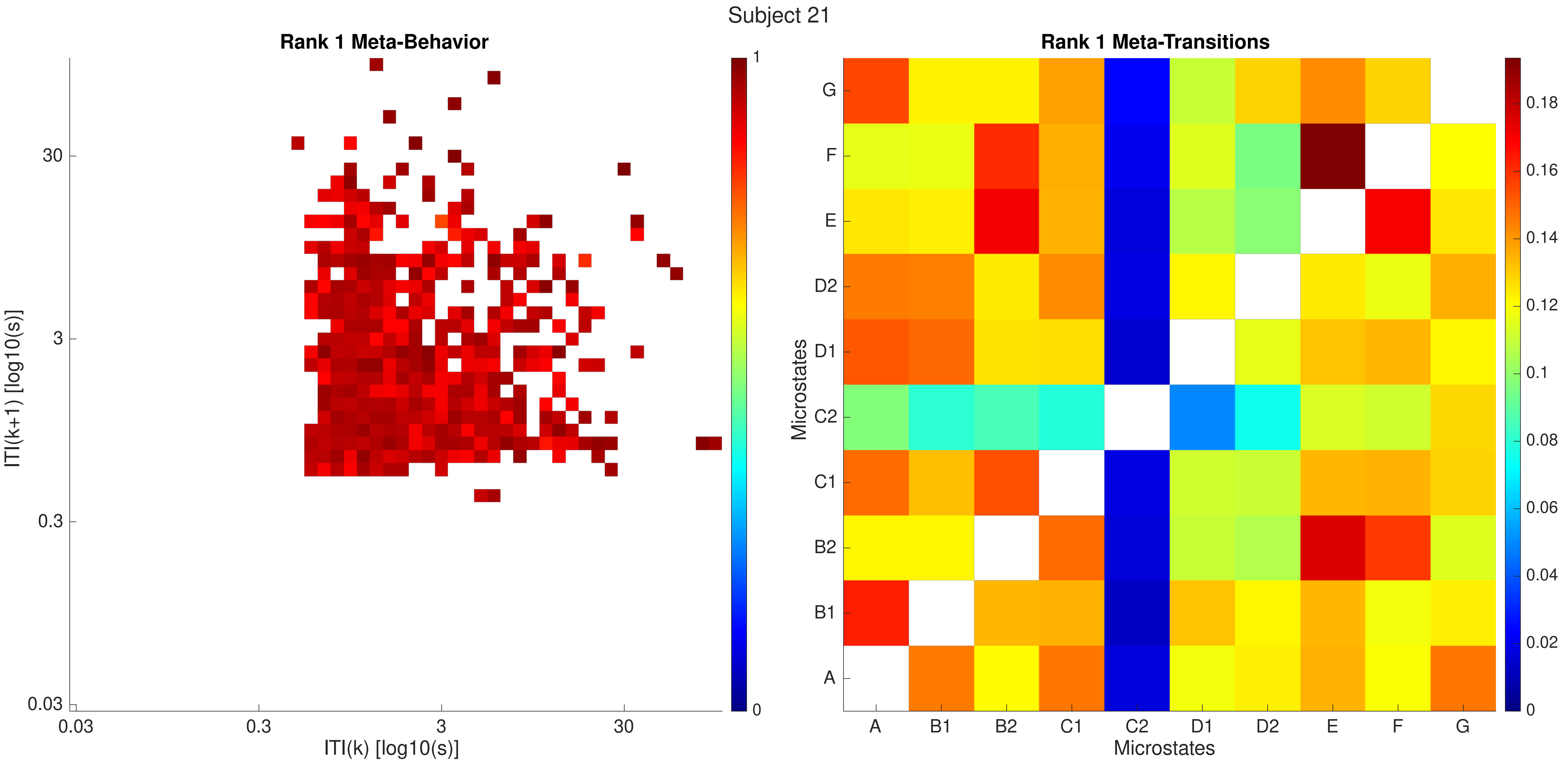

Subject 21
Rank 1 Meta-Behavior
ITI(k+1) [log10(s)]
ITI(k) [log10(s)]
0.03
0.3
3
30
0
1
Rank 1 Meta-Transitions
Microstates
A
B1
B2
C1
C2
D1
D2
E
F
G
0
0.02
0.04
0.06
0.08
0.1
0.12
0.14
0.16
0.18

Subject 21

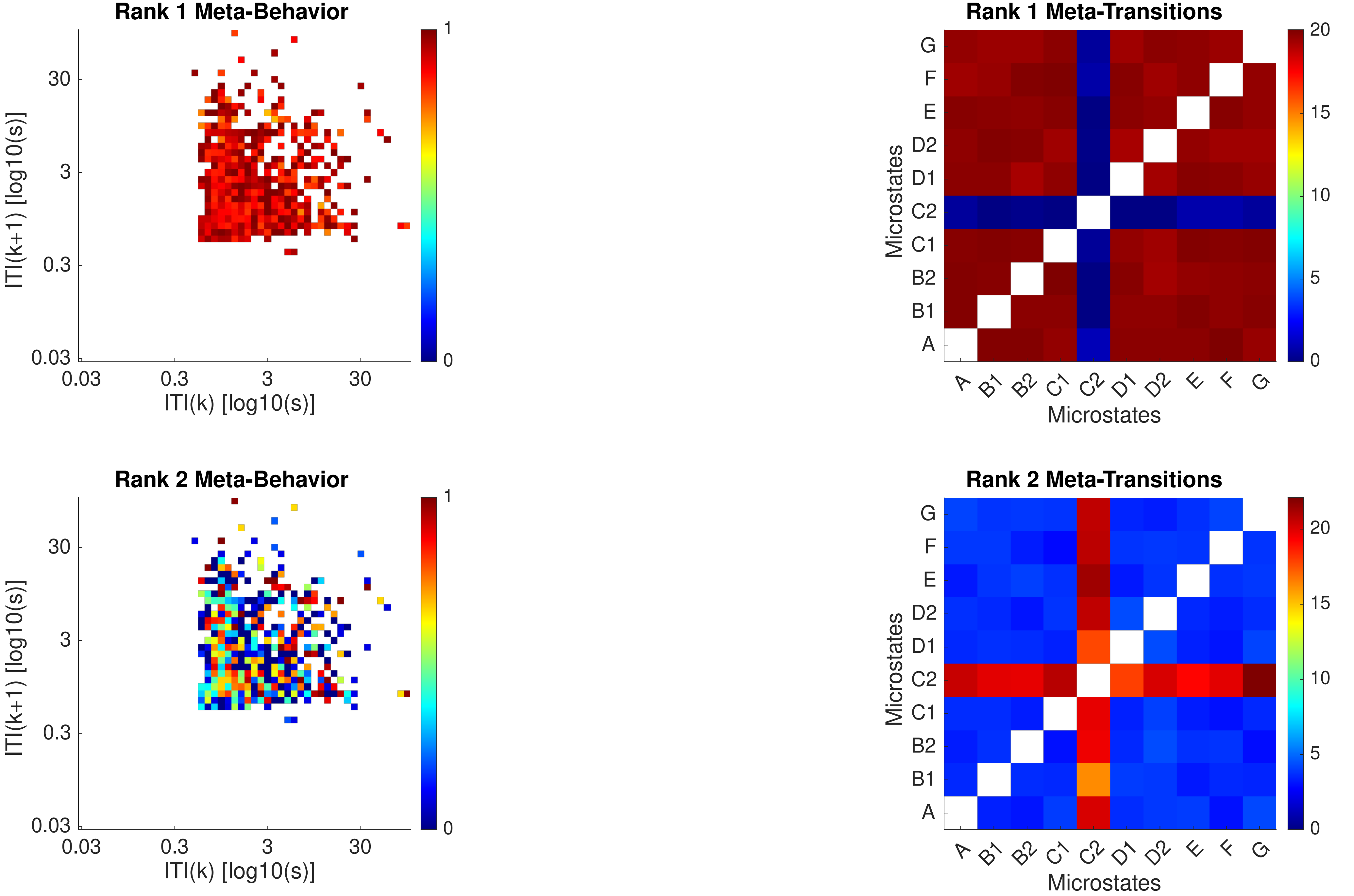

Rank 1 Meta-Behavior
ITI(k+1) [log10(s)]
ITI(k) [log10(s)]
0.03
0.3
3
30
0
1
Rank 1 Meta-Transitions
Microstates
A
B1
B2
C1
C2
D1
D2
E
F
G
0
5
10
15
20
Rank 2 Meta-Behavior
ITI(k+1) [log10(s)]
ITI(k) [log10(s)]
Rank 2 Meta-Transitions
Microstates

Subject 21

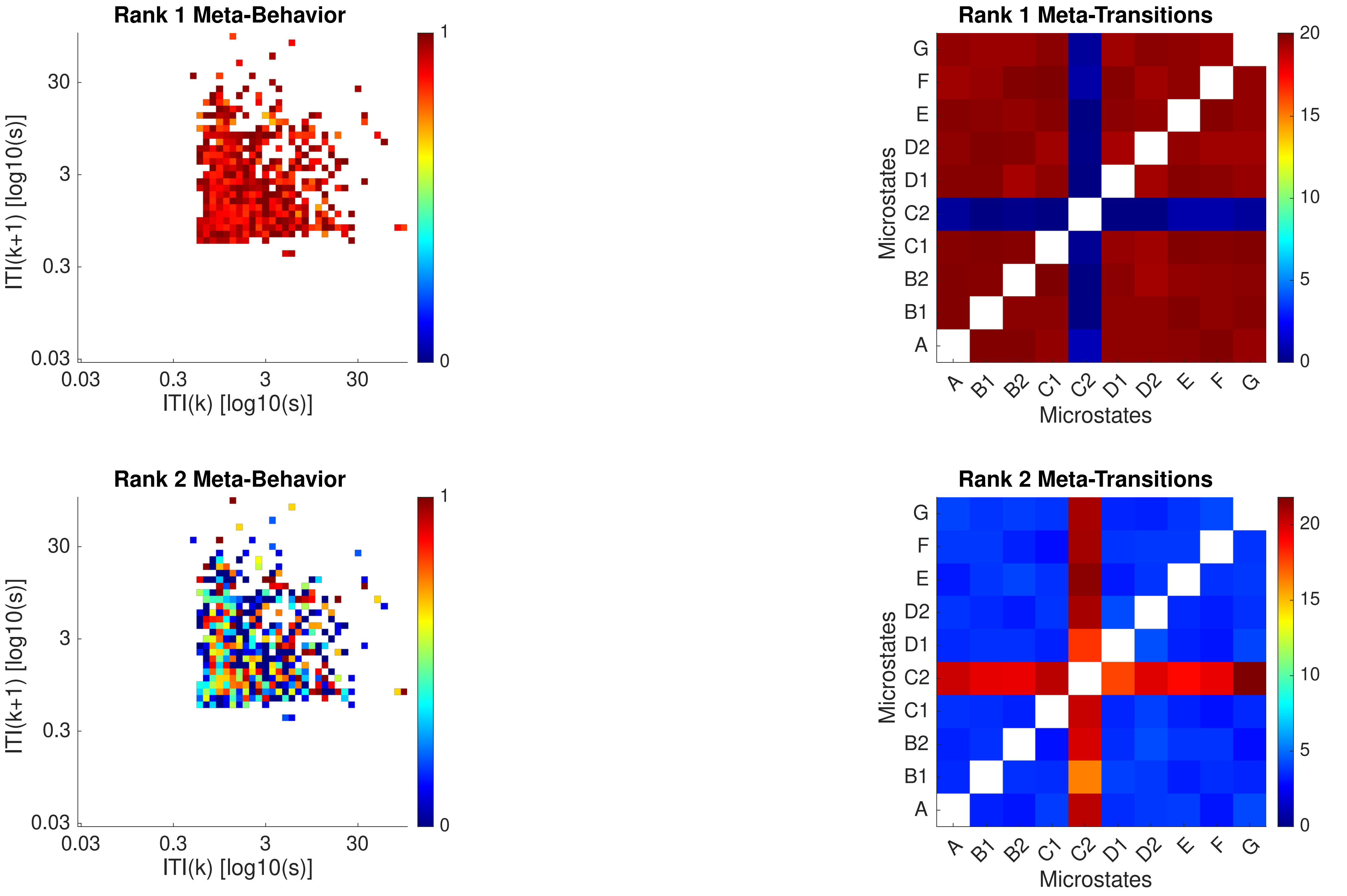

Rank 1 Meta-Behavior
ITI(k+1) [log10(s)]
ITI(k) [log10(s)]
0.03
0.3
3
30
0
1
Rank 1 Meta-Transitions
Microstates
A
B1
B2
C1
C2
D1
D2
E
F
G
0
5
10
15
20
Rank 2 Meta-Behavior
Rank 2 Meta-Transitions

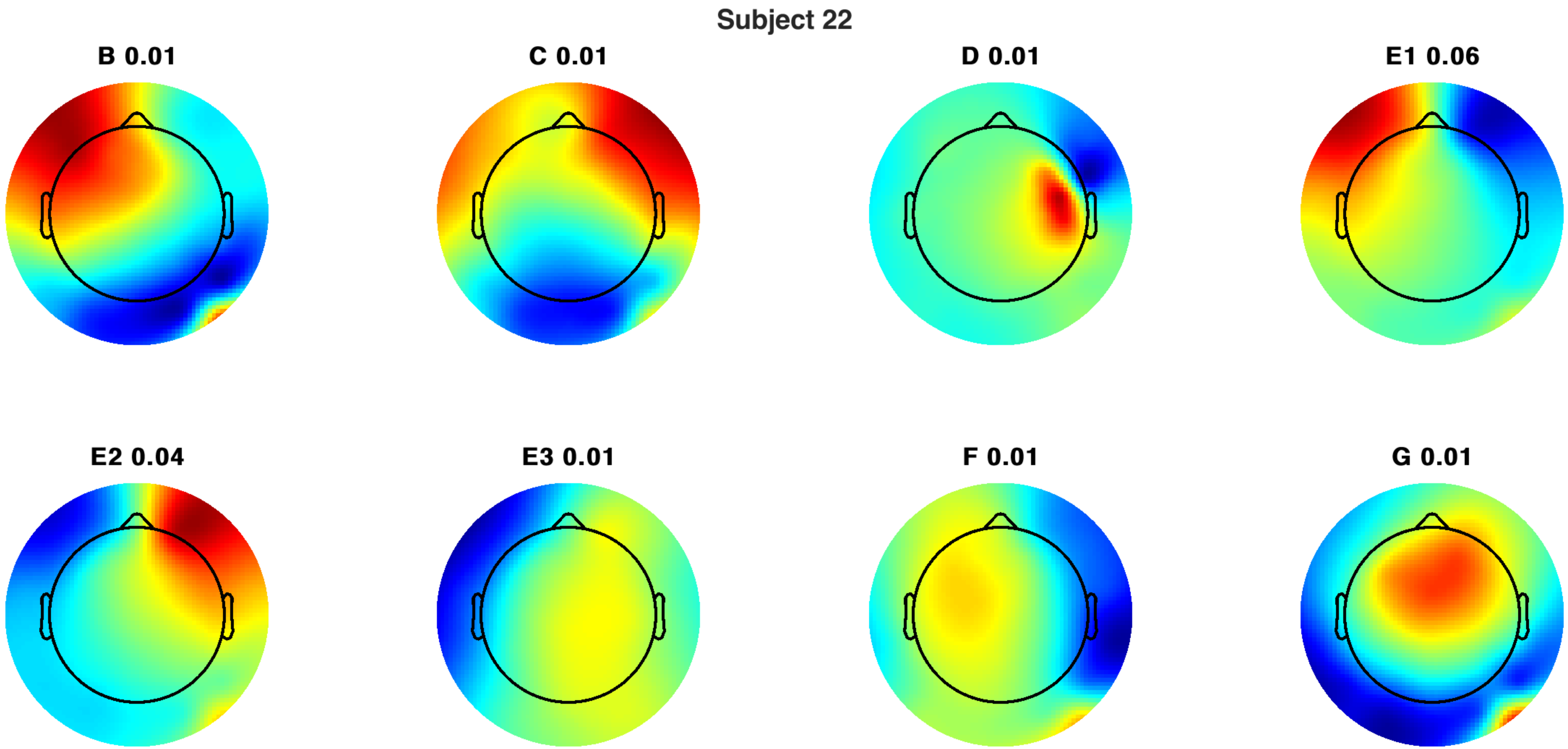
Subject 22
B 0.01
C 0.01
D 0.01
E1 0.06
E2 0.04
E3 0.01
F 0.01
G 0.01

Subject 22

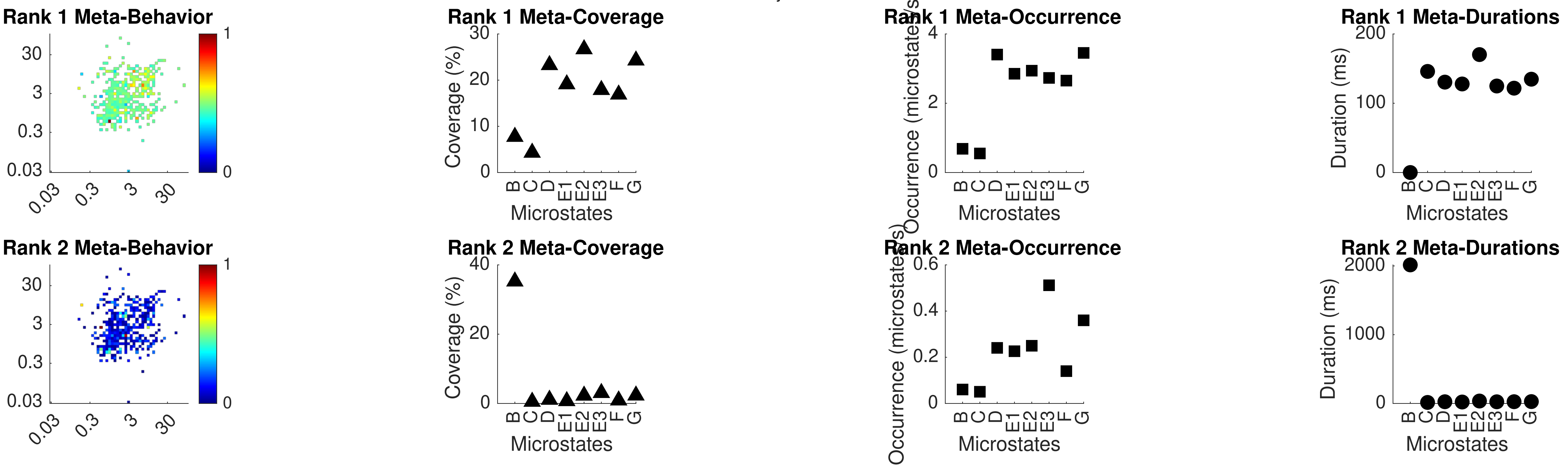
Rank 1 Meta-Behavior
Rank 1 Meta-Coverage
Coverage (%)
Microstates
Rank 1 Meta-Occurrence
Occurrence (microstates/s)
Rank 1 Meta-Durations
Duration (ms)
Rank 2 Meta-Behavior
Rank 2 Meta-Coverage
Rank 2 Meta-Occurrence
Rank 2 Meta-Durations
B C D E1 E2 E3 F G
0.03 0.3 3 30

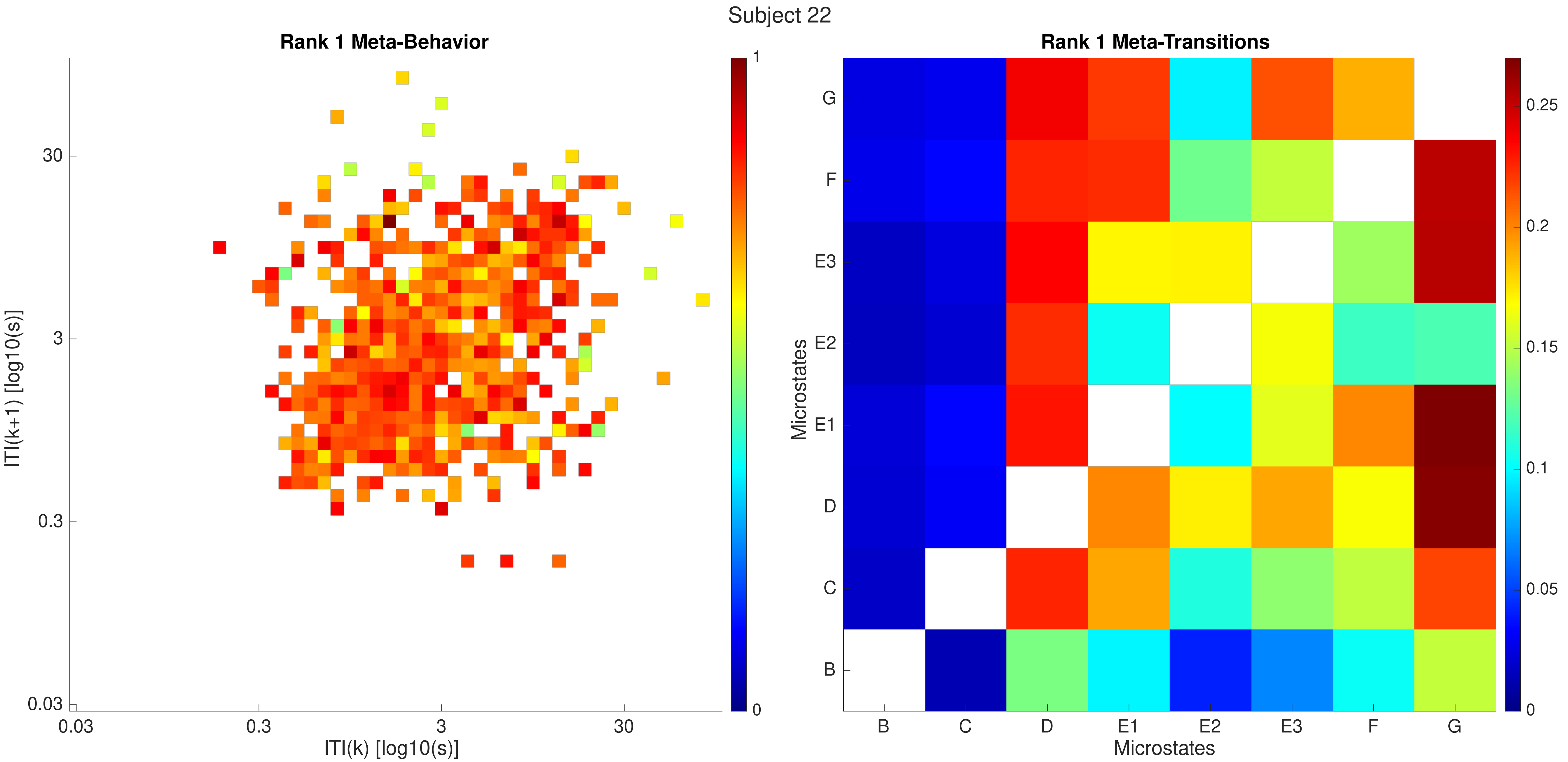
Subject 22
Rank 1 Meta-Behavior
ITI(k+1) [log10(s)]
ITI(k) [log10(s)]
0.03
0.3
3
30
0
1
Rank 1 Meta-Transitions
Microstates
B
C
D
E1
E2
E3
F
G
0
0.05
0.1
0.15
0.2
0.25

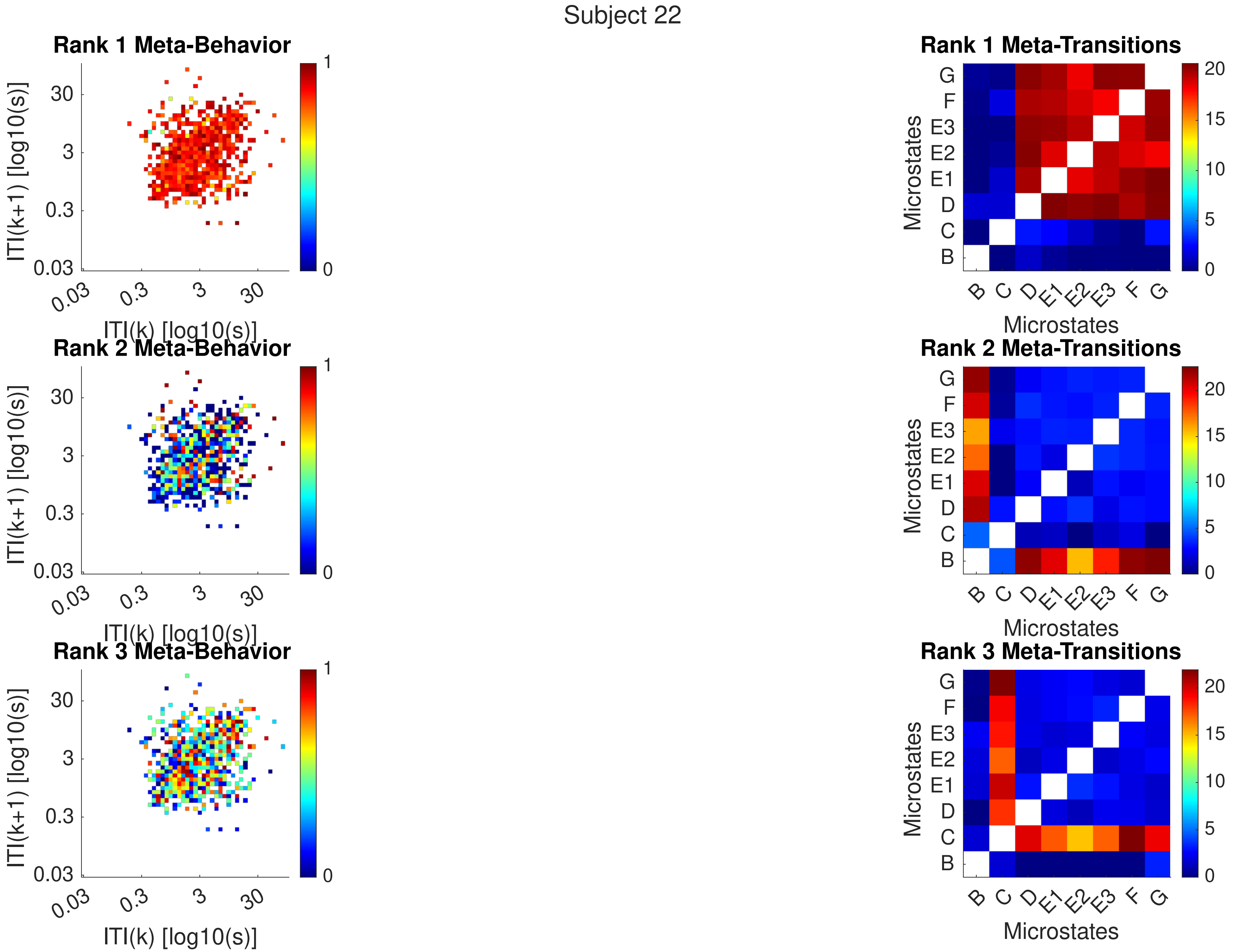
Subject 22
Rank 1 Meta-Behavior
Rank 2 Meta-Behavior
Rank 3 Meta-Behavior
ITI(k+1) [log10(s)]
ITI(k) [log10(s)]
0.03
0.3
3
30
1
0
Rank 1 Meta-Transitions
Rank 2 Meta-Transitions
Rank 3 Meta-Transitions
Microstates
B
C
D
E1
E2
E3
F
G
0
5
10
15
20

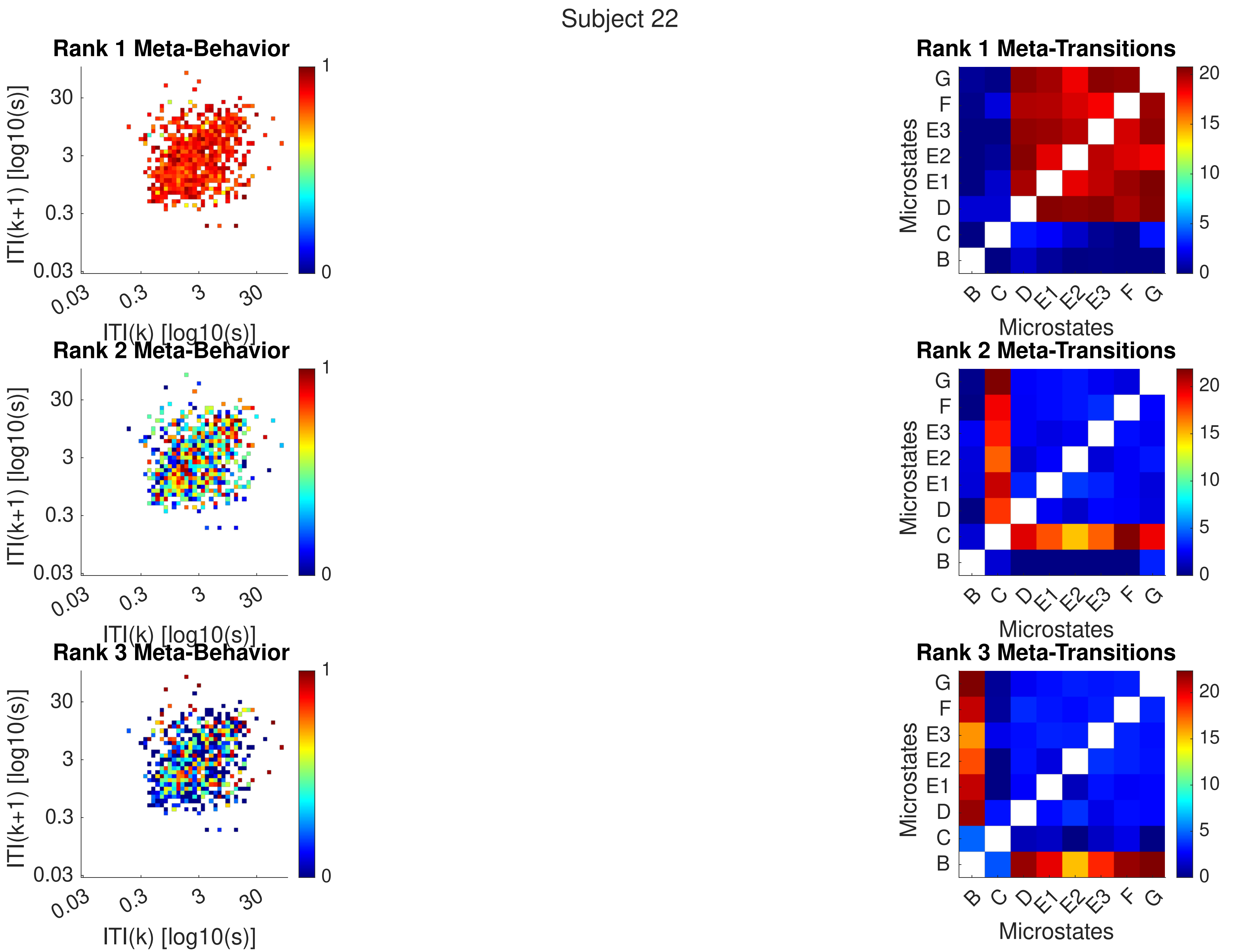
Subject 22
Rank 1 Meta-Behavior
Rank 2 Meta-Behavior
Rank 3 Meta-Behavior
ITI(k+1) [log10(s)]
ITI(k) [log10(s)]
0.03
0.3
3
30
0
1
Rank 1 Meta-Transitions
Rank 2 Meta-Transitions
Rank 3 Meta-Transitions
Microstates
B
C
D
E1
E2
E3
F
G
0
5
10
15
20

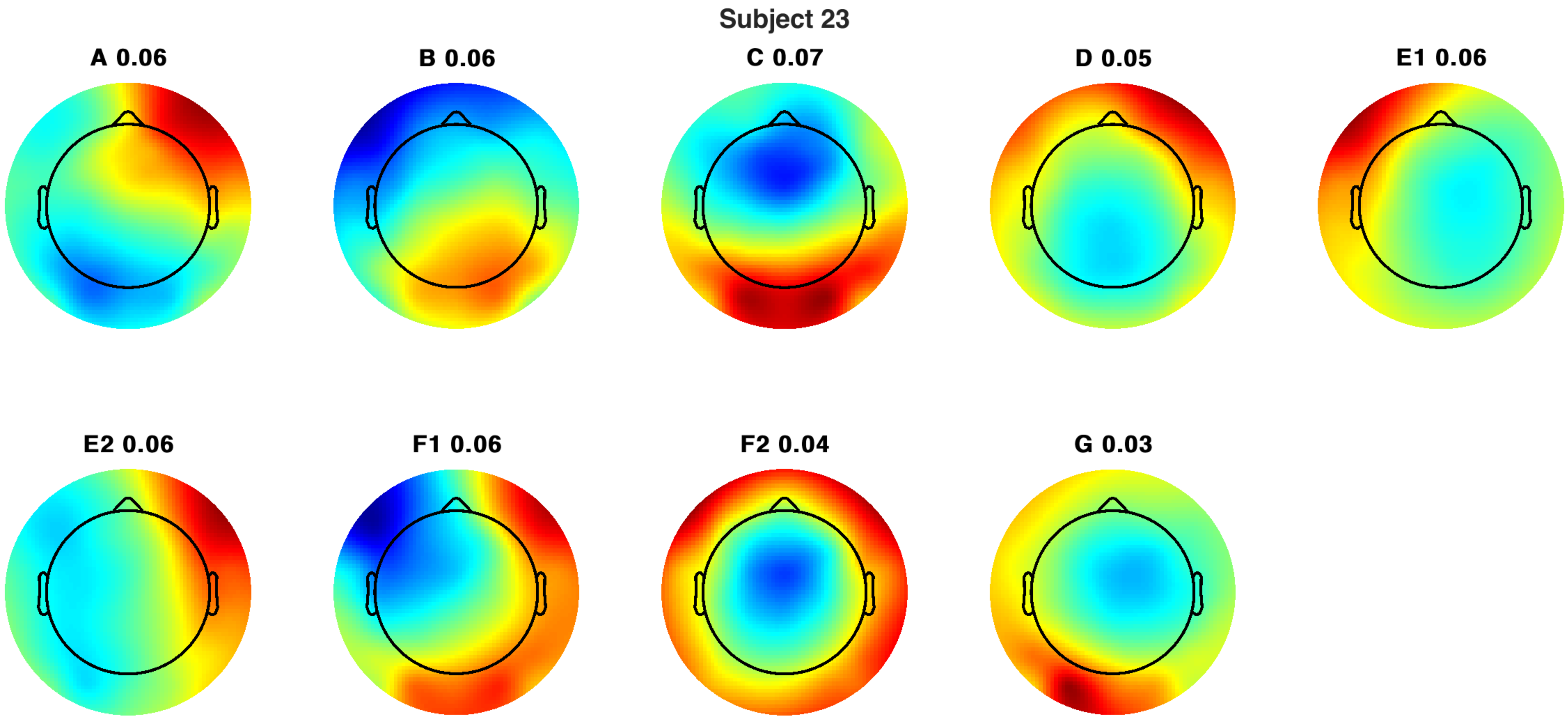
Subject 23
A 0.06
B 0.06
C 0.07
D 0.05
E1 0.06
E2 0.06
F1 0.06
F2 0.04
G 0.03

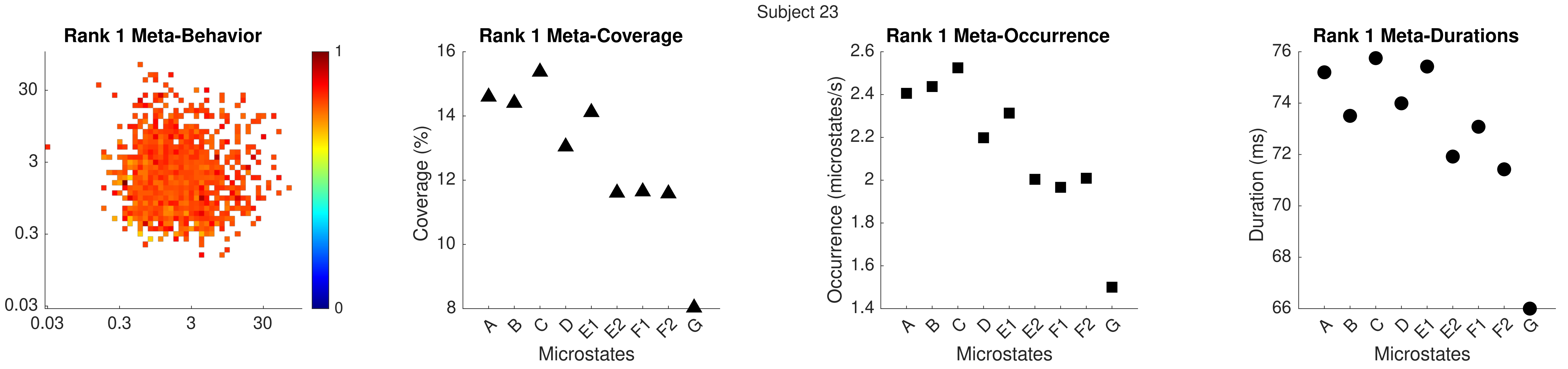

Subject 23
Rank 1 Meta-Behavior
0.03
0.3
3
30
0
1
Rank 1 Meta-Coverage
Coverage (%)
Microstates
A B C D E1 E2 F1 F2 G
Rank 1 Meta-Occurrence
Occurrence (microstates/s)
Microstates
Rank 1 Meta-Durations
Duration (ms)
Microstates

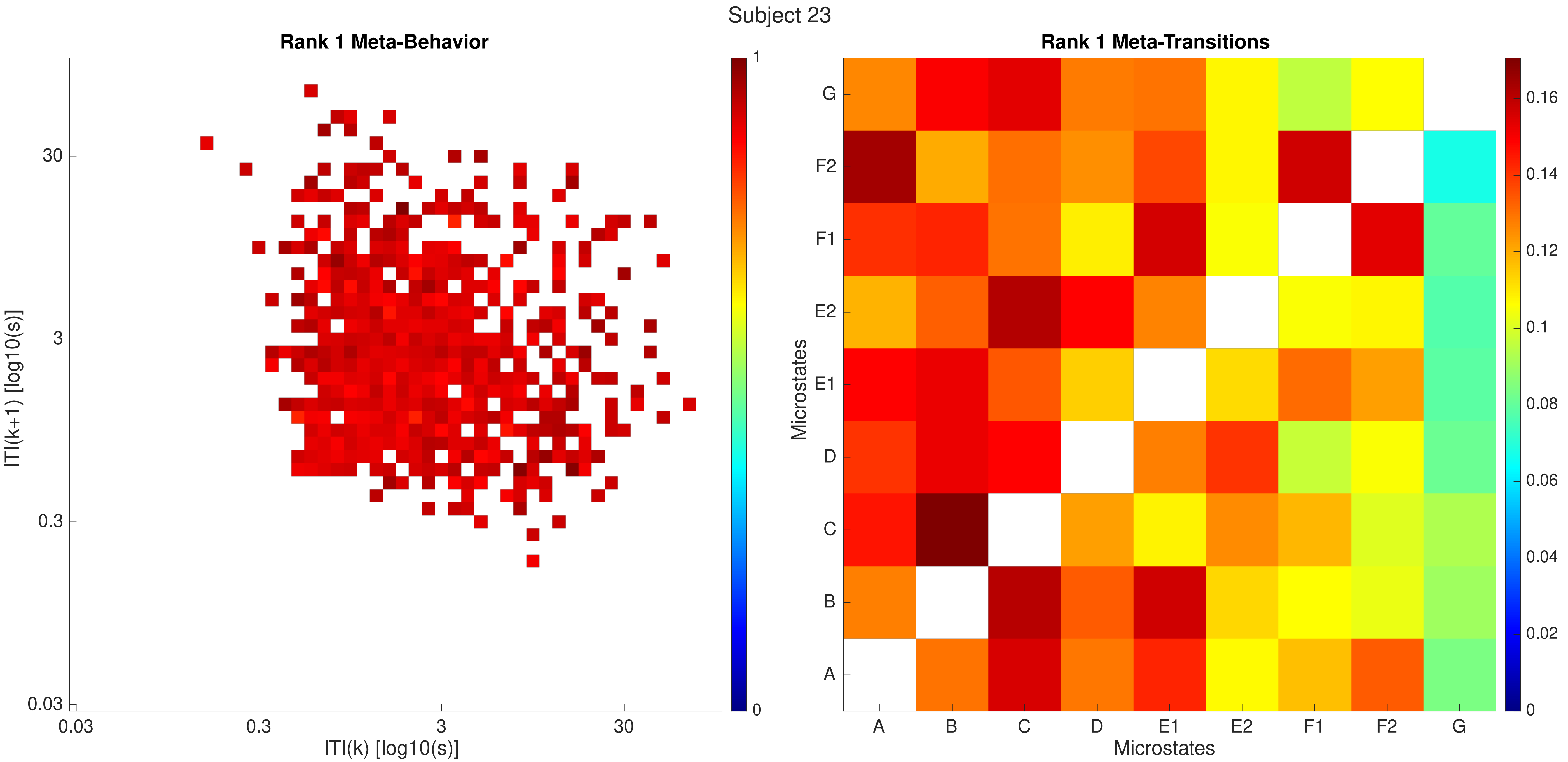

Subject 23
Rank 1 Meta-Behavior
ITI(k+1) [log10(s)]
ITI(k) [log10(s)]
0.03
0.3
3
30
0
1
Rank 1 Meta-Transitions
Microstates
A
B
C
D
E1
E2
F1
F2
G
0
0.02
0.04
0.06
0.08
0.1
0.12
0.14
0.16

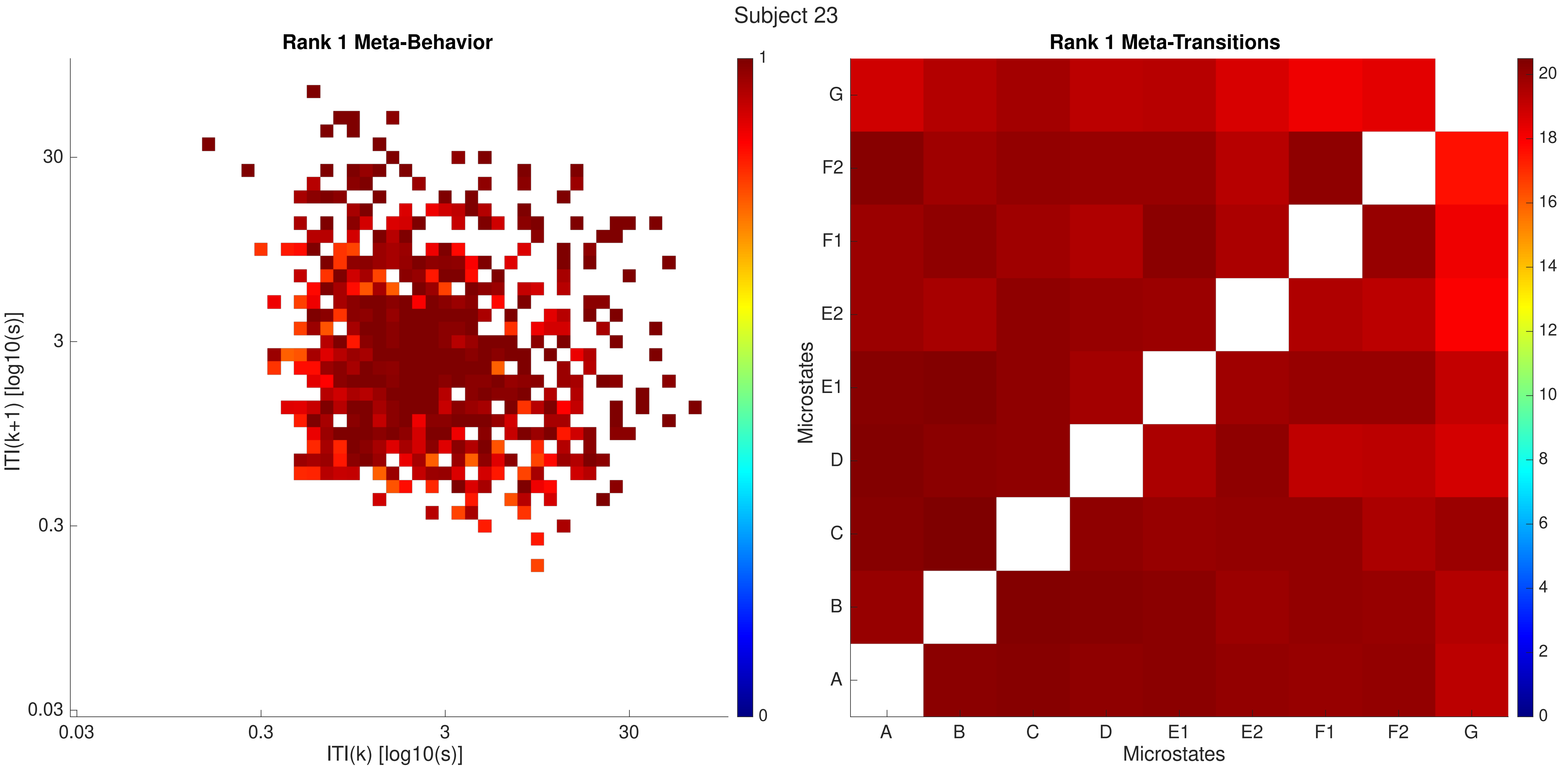
Subject 23
Rank 1 Meta-Behavior
ITI(k+1) [log10(s)]
30
3
0.3
0.03
ITI(k) [log10(s)]
0.03
0.3
3
30
1
0
Rank 1 Meta-Transitions
Microstates
G
F2
F1
E2
E1
D
C
B
A
Microstates
A
B
C
D
E1
E2
F1
F2
G
20
18
16
14
12
10
8
6
4
2
0

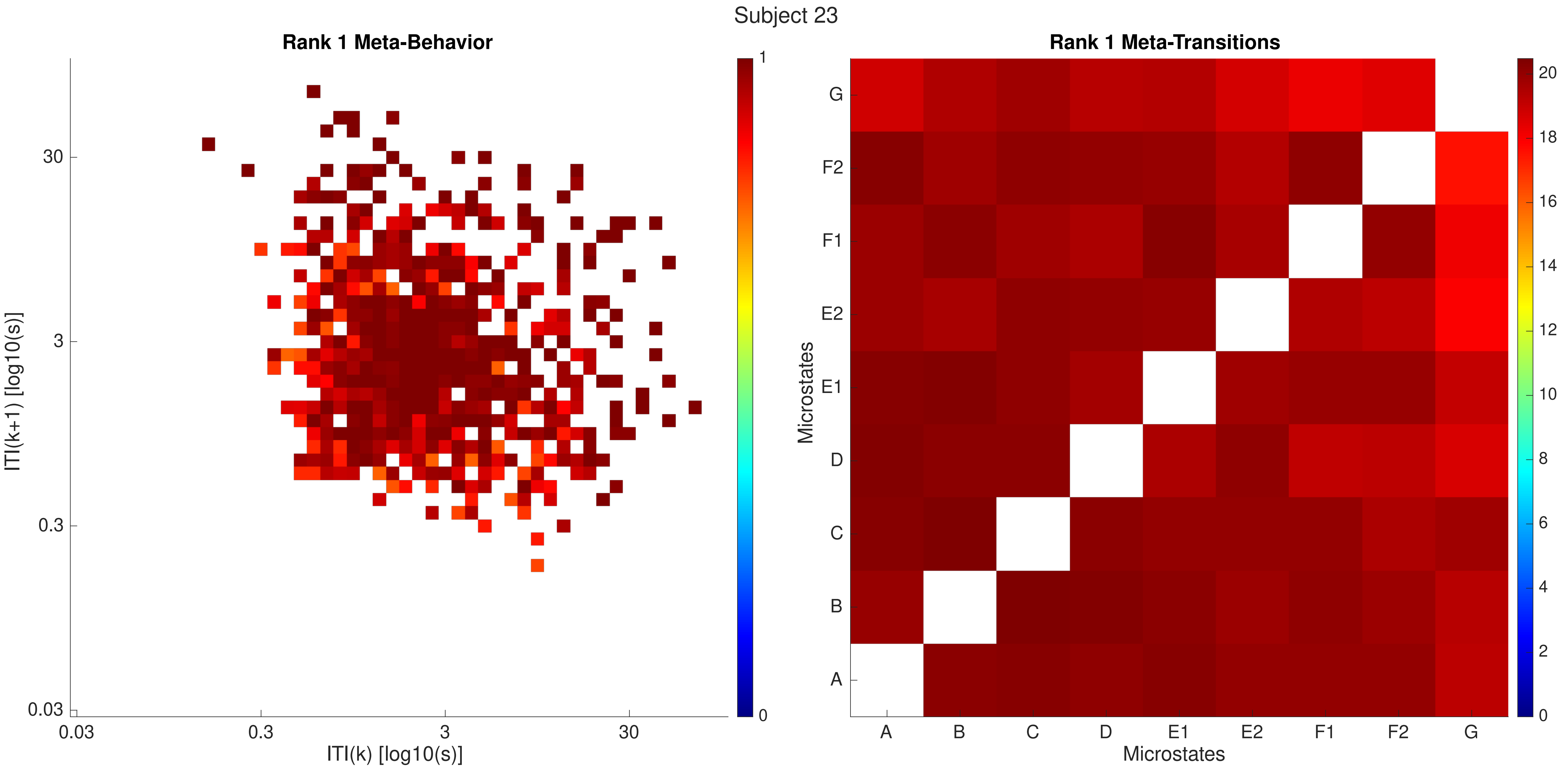

Subject 23
Rank 1 Meta-Behavior
ITI(k+1) [log10(s)]
30
3
0.3
0.03
0.03
0.3
3
30
ITI(k) [log10(s)]
1
0
Rank 1 Meta-Transitions
Microstates
G
F2
F1
E2
E1
D
C
B
A
A
B
C
D
E1
E2
F1
F2
G
Microstates
20
18
16
14
12
10
8
6
4
2
0

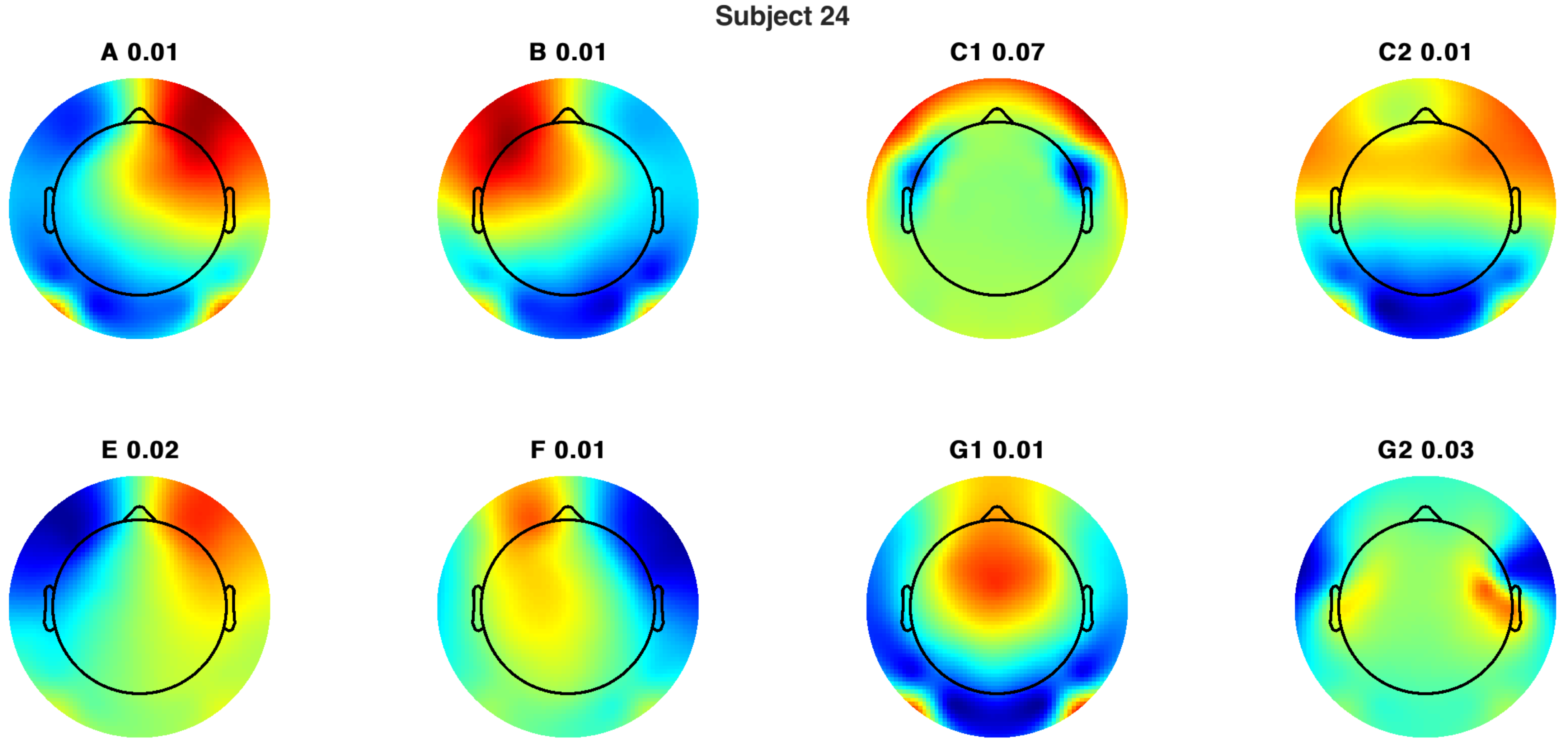
Subject 24
A 0.01
B 0.01
C1 0.07
C2 0.01
E 0.02
F 0.01
G1 0.01
G2 0.03

Subject 24

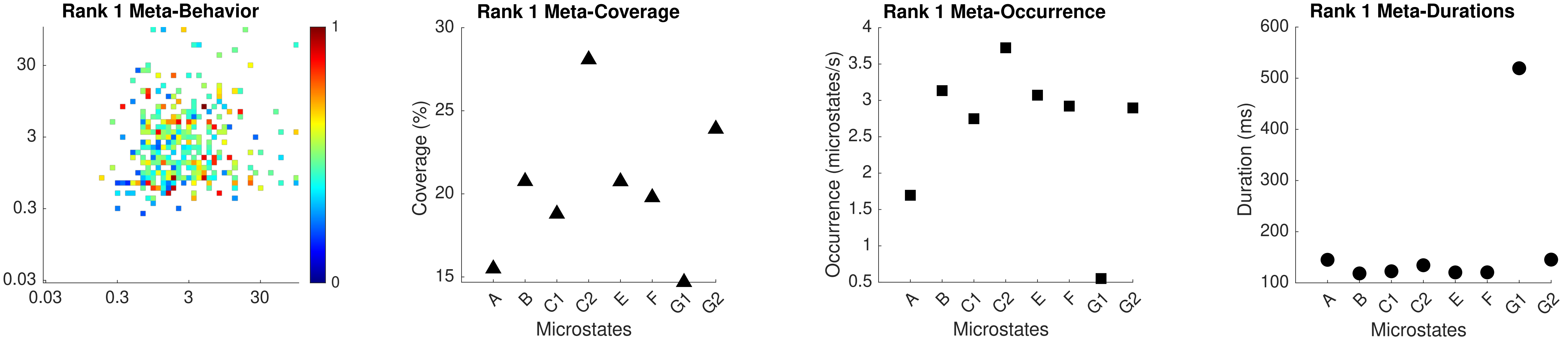

Rank 1 Meta-Behavior
Rank 1 Meta-Coverage
Rank 1 Meta-Occurrence
Rank 1 Meta-Durations
Coverage (%)
Occurrence (microstates/s)
Duration (ms)
Microstates
A
B
C1
C2
E
F
G1
G2

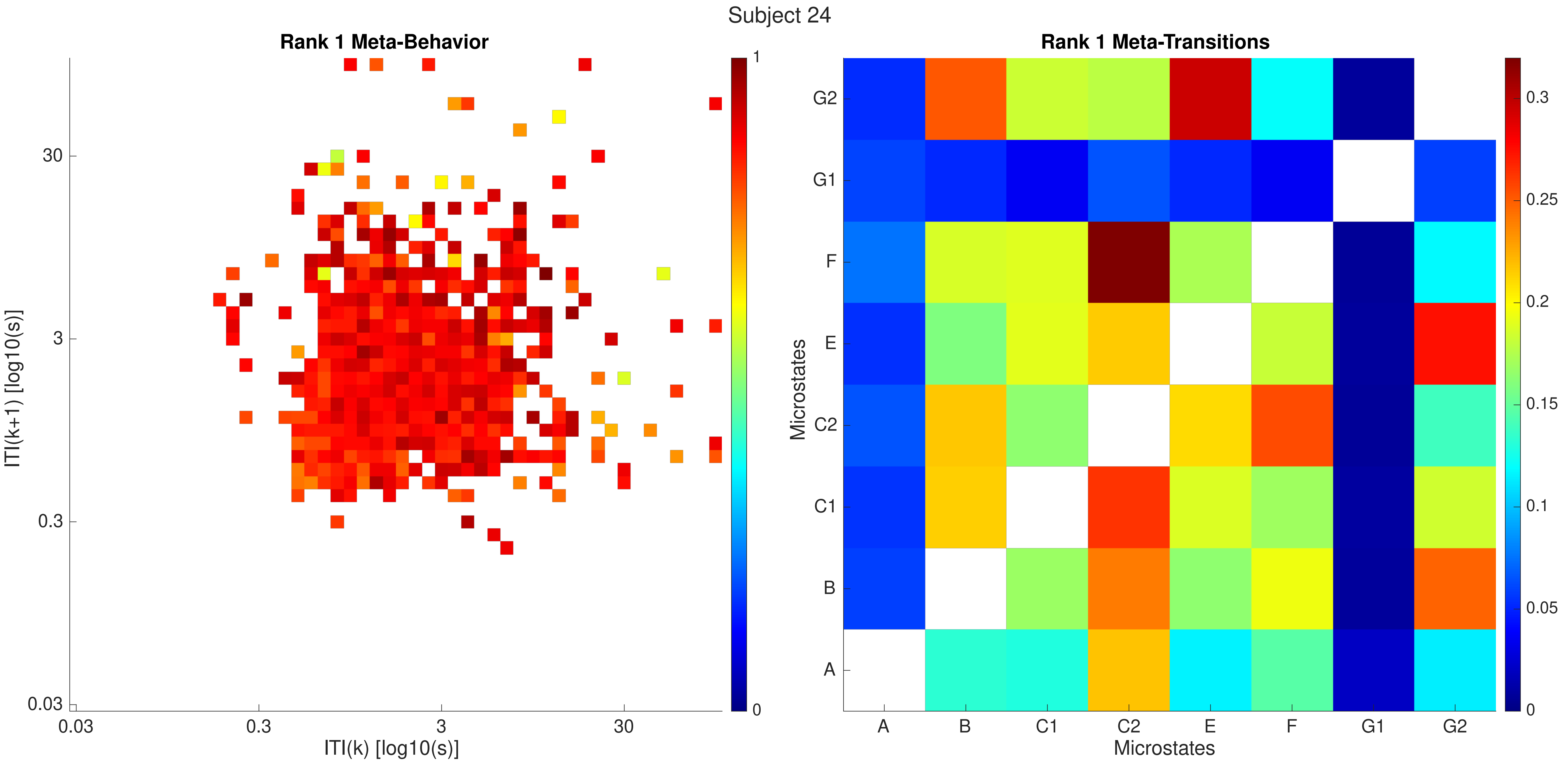

Subject 24
Rank 1 Meta-Behavior
ITI(k+1) [log10(s)]
ITI(k) [log10(s)]
0.03
0.3
3
30
1
0
Rank 1 Meta-Transitions
Microstates
A
B
C1
C2
E
F
G1
G2
0
0.05
0.1
0.15
0.2
0.25
0.3

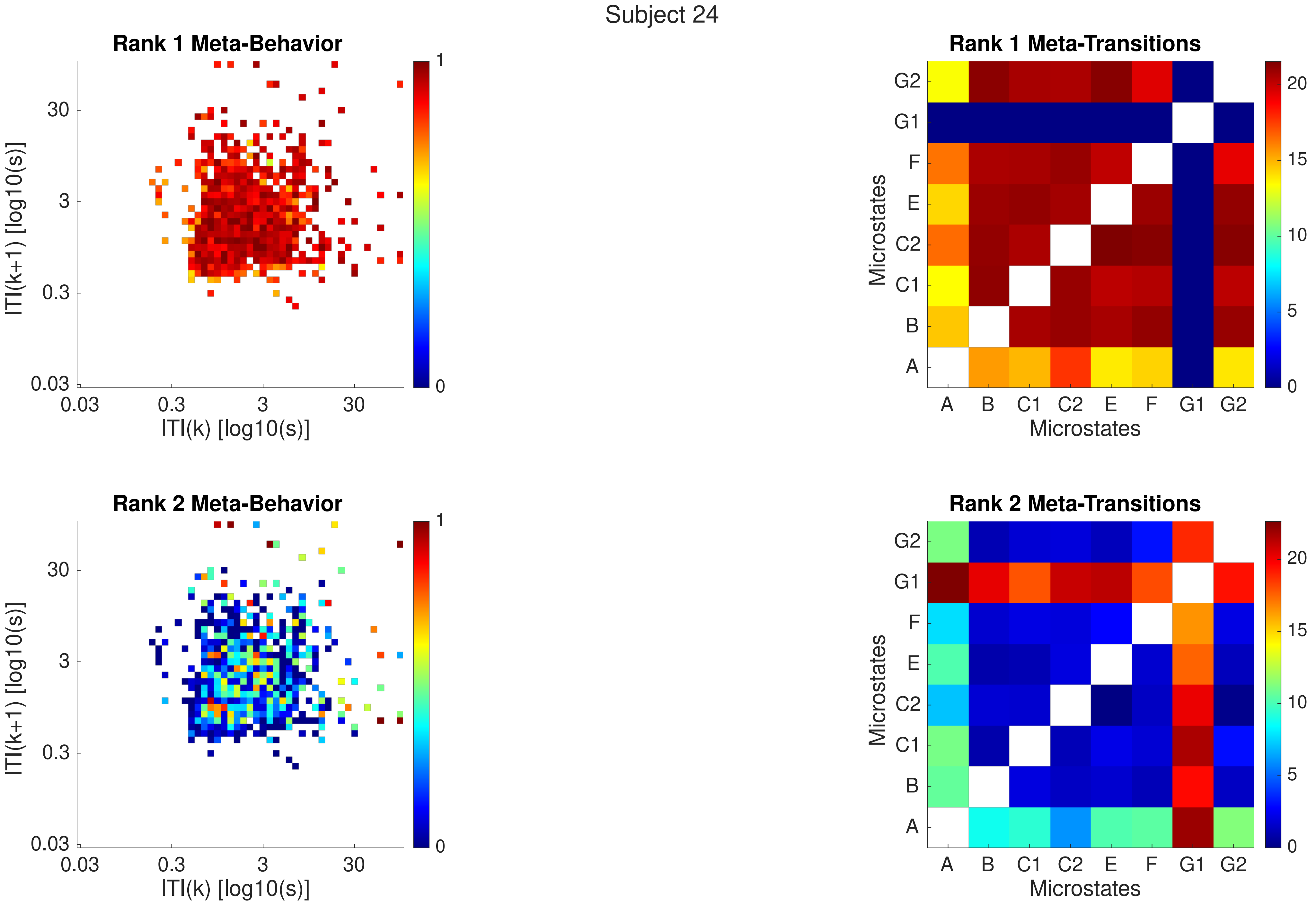
Subject 24
Rank 1 Meta-Behavior
ITI(k+1) [log10(s)]
ITI(k) [log10(s)]
0.03
0.3
3
30
0
1
Rank 1 Meta-Transitions
Microstates
A
B
C1
C2
E
F
G1
G2
0
5
10
15
20
Rank 2 Meta-Behavior
Rank 2 Meta-Transitions

Subject 24

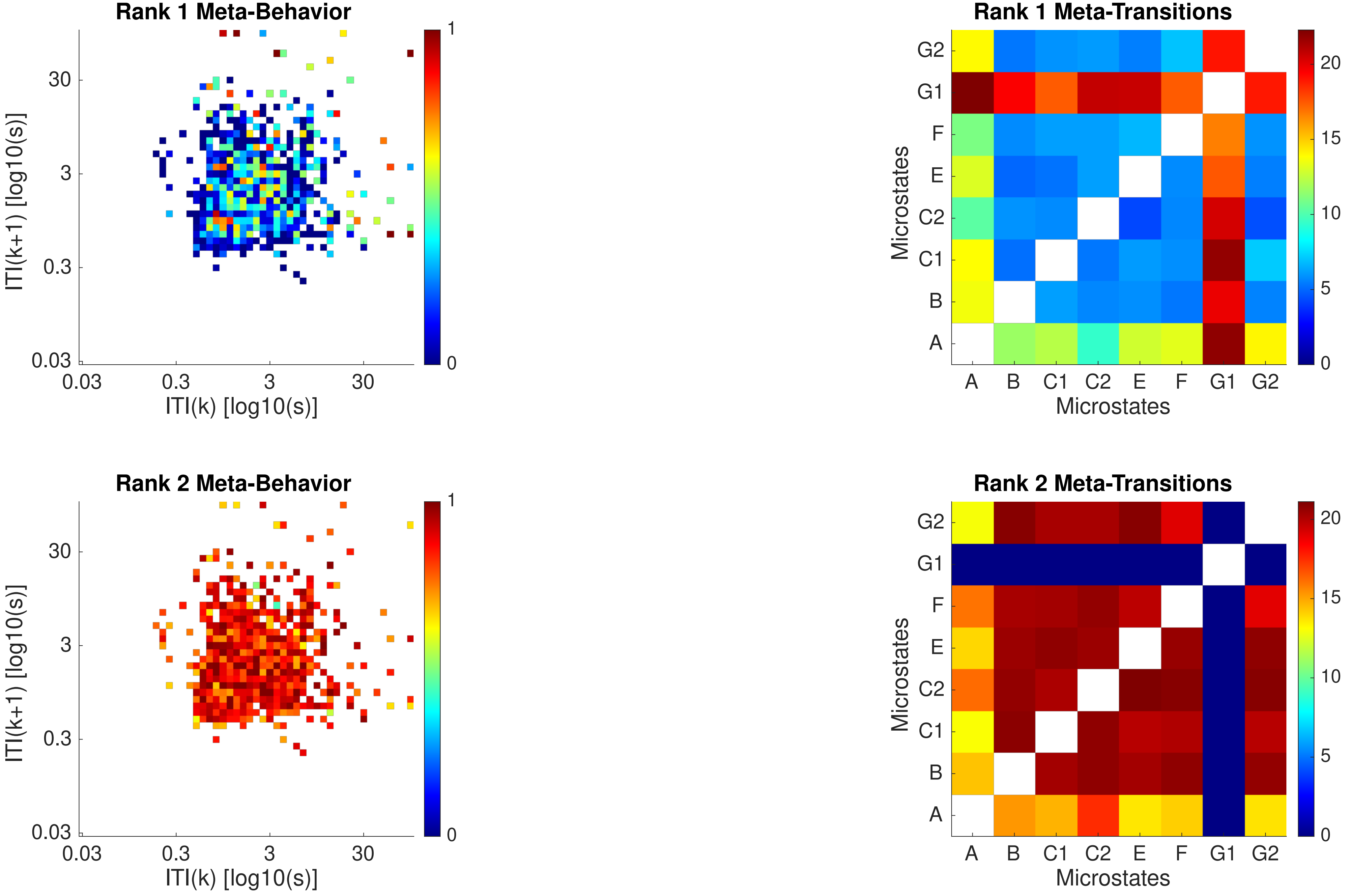

Rank 1 Meta-Behavior
ITI(k+1) [log10(s)]
ITI(k) [log10(s)]
0.03
0.3
3
30
0
1
Rank 1 Meta-Transitions
Microstates
A
B
C1
C2
E
F
G1
G2
0
5
10
15
20
Rank 2 Meta-Behavior
ITI(k+1) [log10(s)]
ITI(k) [log10(s)]
Rank 2 Meta-Transitions
Microstates

Subject 25

B 0.00

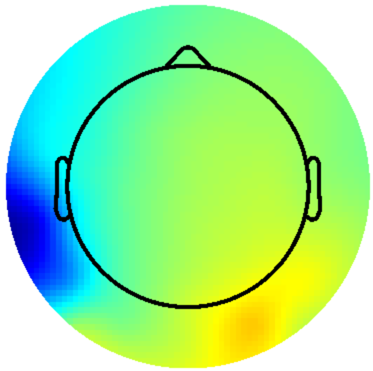

E 0.00

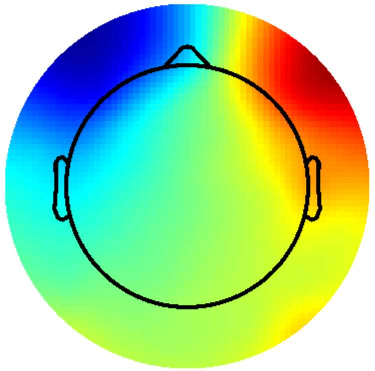

Subject 25

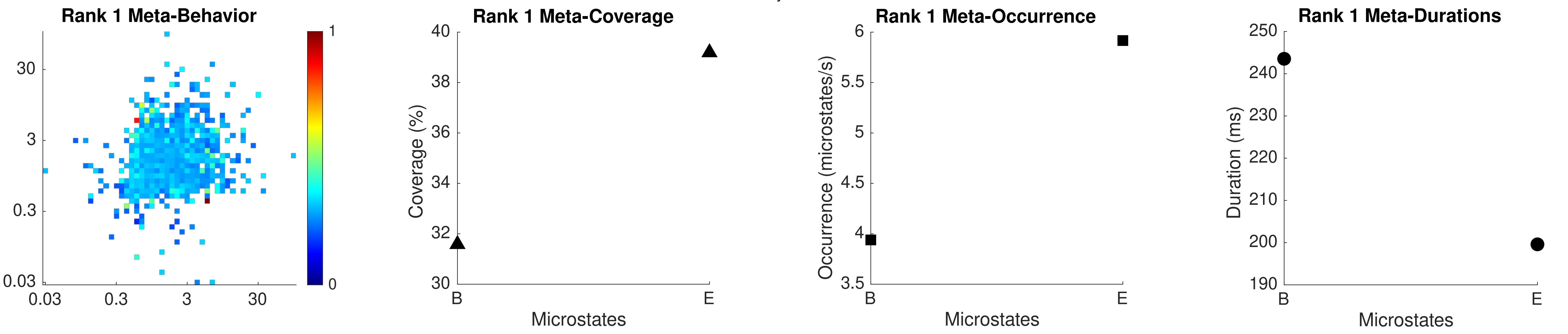
Rank 1 Meta-Behavior
30
3
0.3
0.03
0.03
0.3
3
30
1
0
Rank 1 Meta-Coverage
Coverage (%)
40
38
36
34
32
30
B
E
Microstates
Rank 1 Meta-Occurrence
Occurrence (microstates/s)
6
5.5
5
4.5
4
3.5
B
E
Microstates
Rank 1 Meta-Durations
Duration (ms)
250
240
230
220
210
200
190
B
E
Microstates

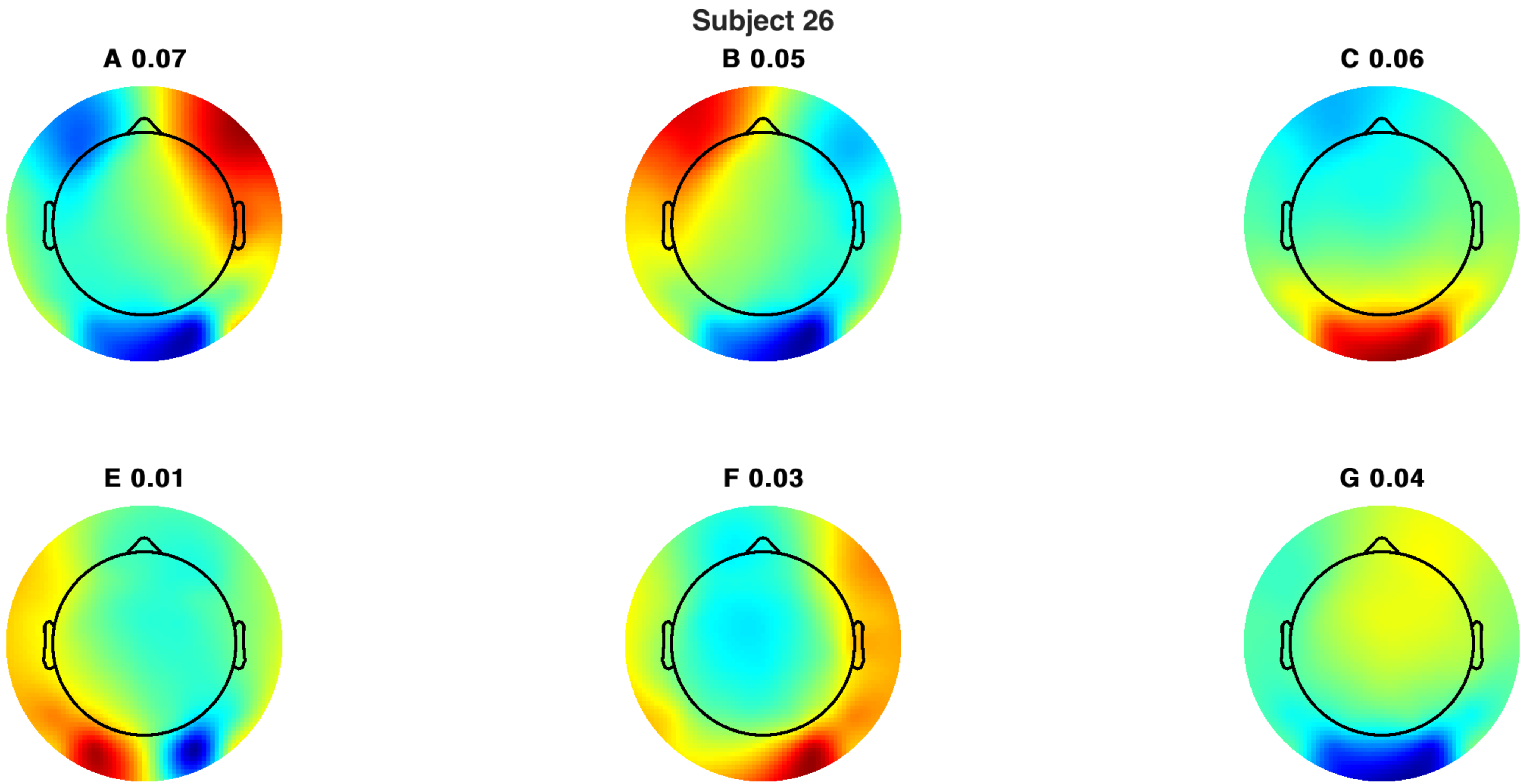
Subject 26
A 0.07
B 0.05
C 0.06
E 0.01
F 0.03
G 0.04

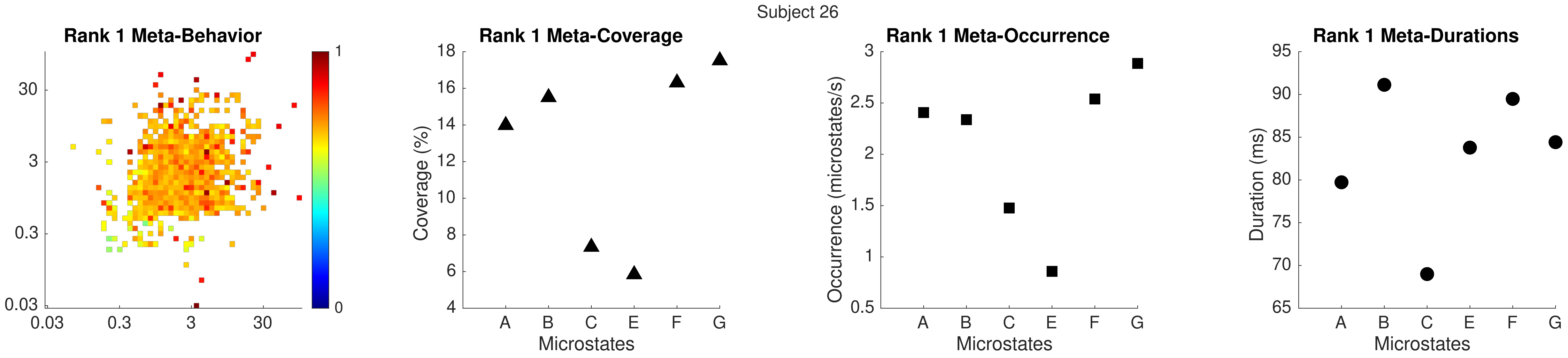
Subject 26
Rank 1 Meta-Behavior
30
3
0.3
0.03
0.03
0.3
3
30
1
0
Rank 1 Meta-Coverage
Coverage (%)
18
16
14
12
10
8
6
4
A
B
C
E
F
G
Microstates
Rank 1 Meta-Occurrence
Occurrence (microstates/s)
3
2.5
2
1.5
1
0.5
A
B
C
E
F
G
Microstates
Rank 1 Meta-Durations
Duration (ms)
95
90
85
80
75
70
65
A
B
C
E
F
G
Microstates

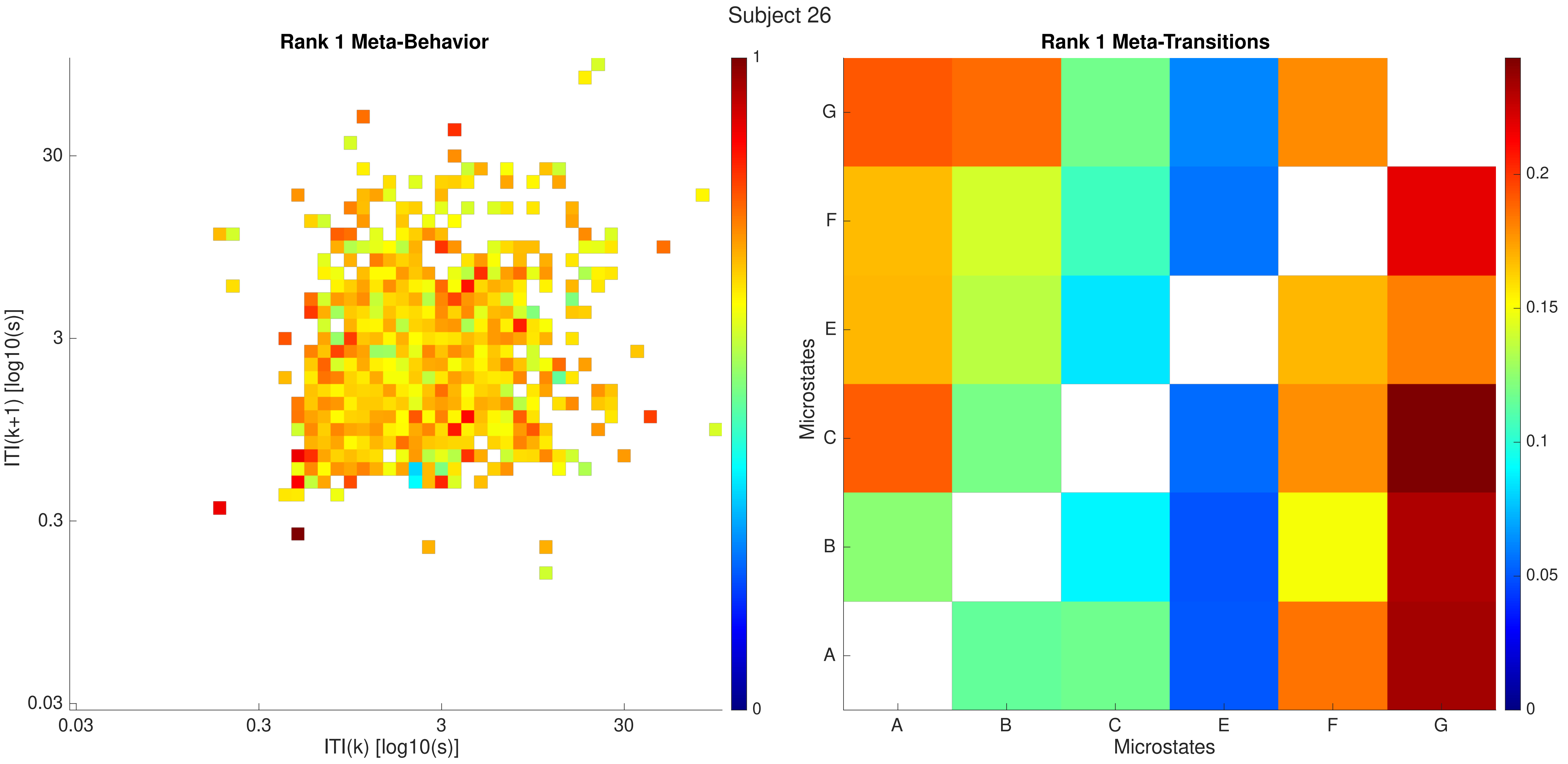
Subject 26
Rank 1 Meta-Behavior
ITI(k+1) [log10(s)]
ITI(k) [log10(s)]
0.03
0.3
3
30
0
1
Rank 1 Meta-Transitions
Microstates
A
B
C
E
F
G
0
0.05
0.1
0.15
0.2

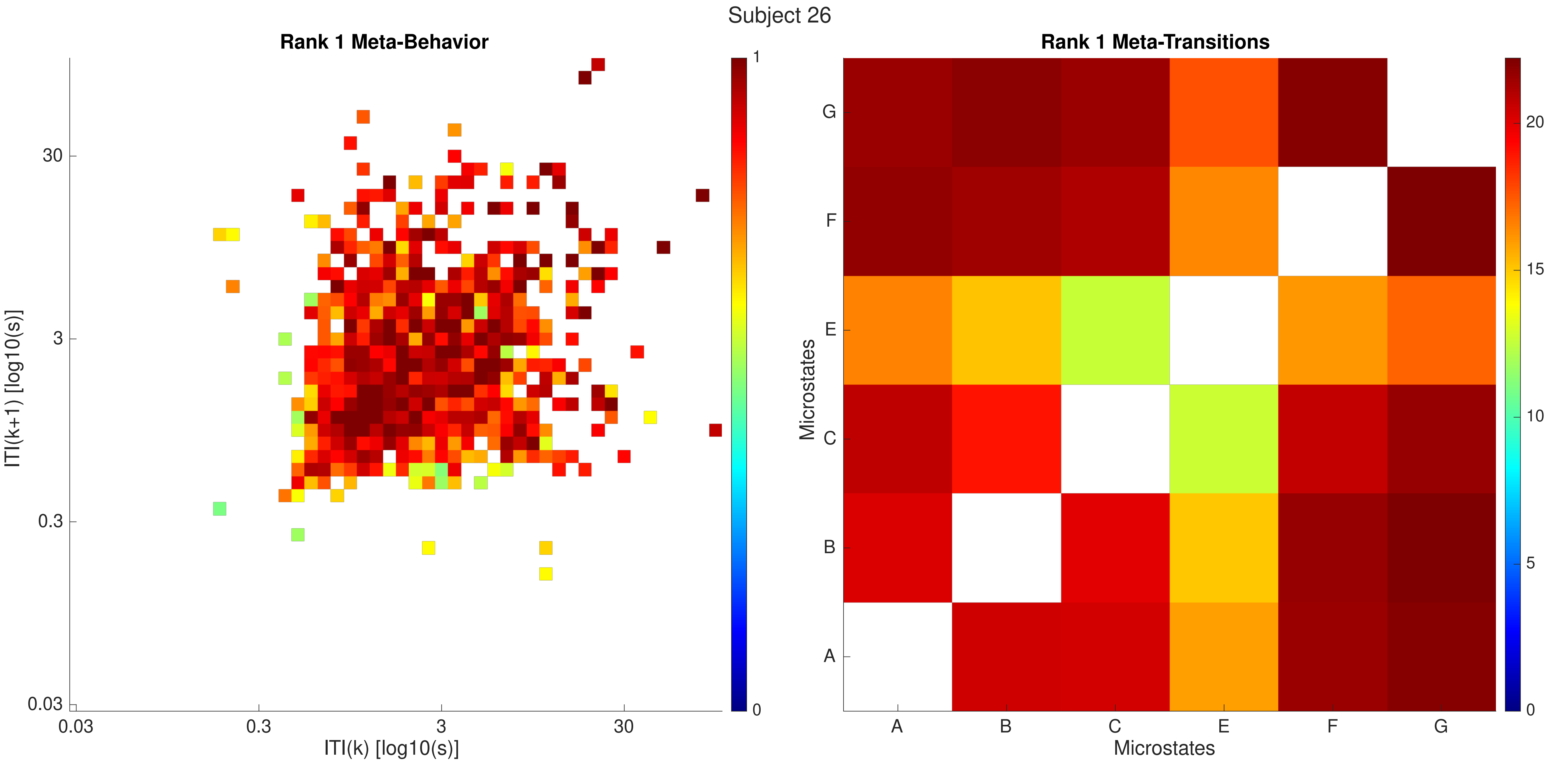
Subject 26
Rank 1 Meta-Behavior
ITI(k+1) [log10(s)]
30
3
0.3
0.03
0.03
0.3
3
30
ITI(k) [log10(s)]
1
0
Rank 1 Meta-Transitions
Microstates
G
F
E
C
B
A
A
B
C
E
F
G
Microstates
20
15
10
5
0

Subject 26

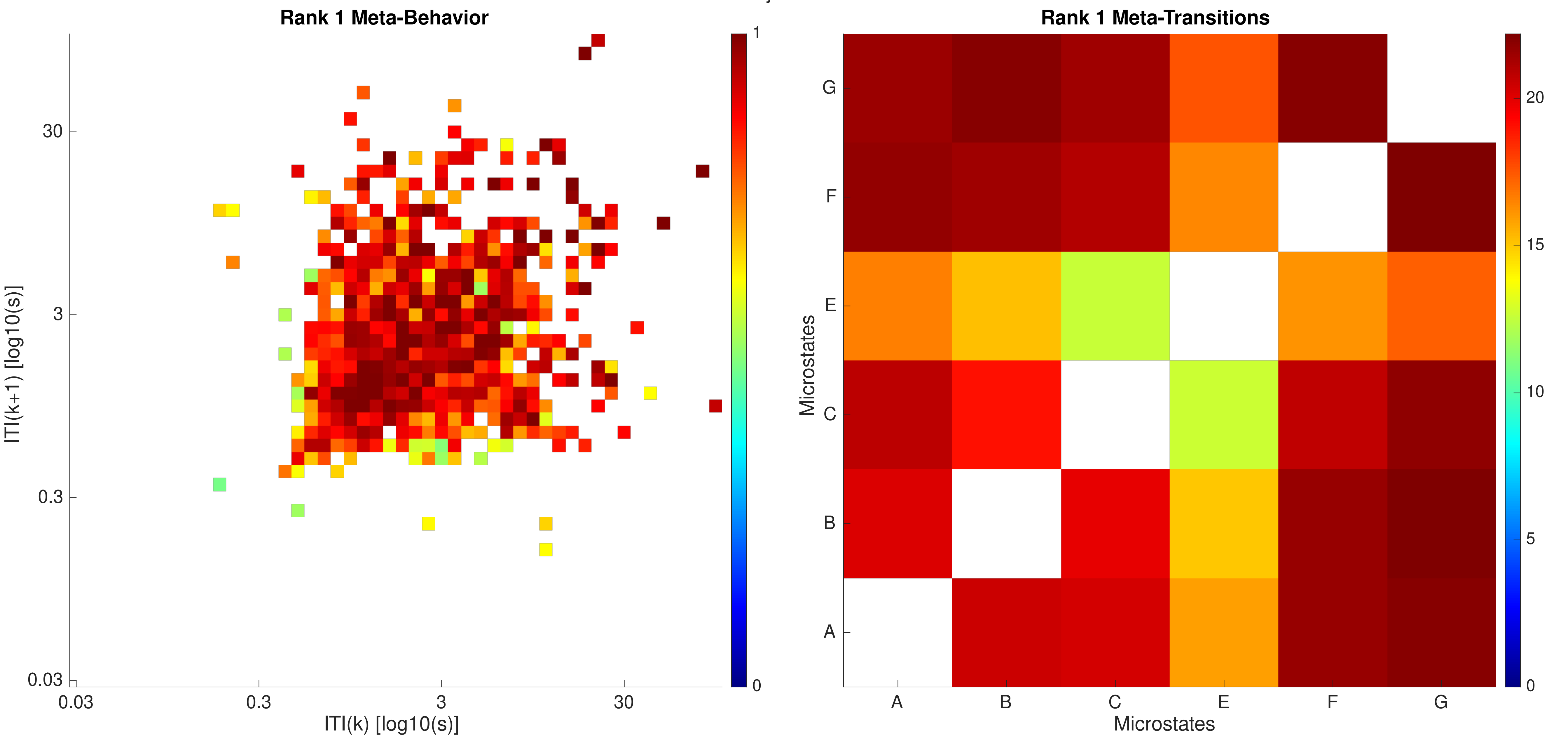

Rank 1 Meta-Behavior
ITI(k+1) [log10(s)]
ITI(k) [log10(s)]
0.03
0.3
3
30
0
1
Rank 1 Meta-Transitions
Microstates
A
B
C
E
F
G
0
5
10
15
20

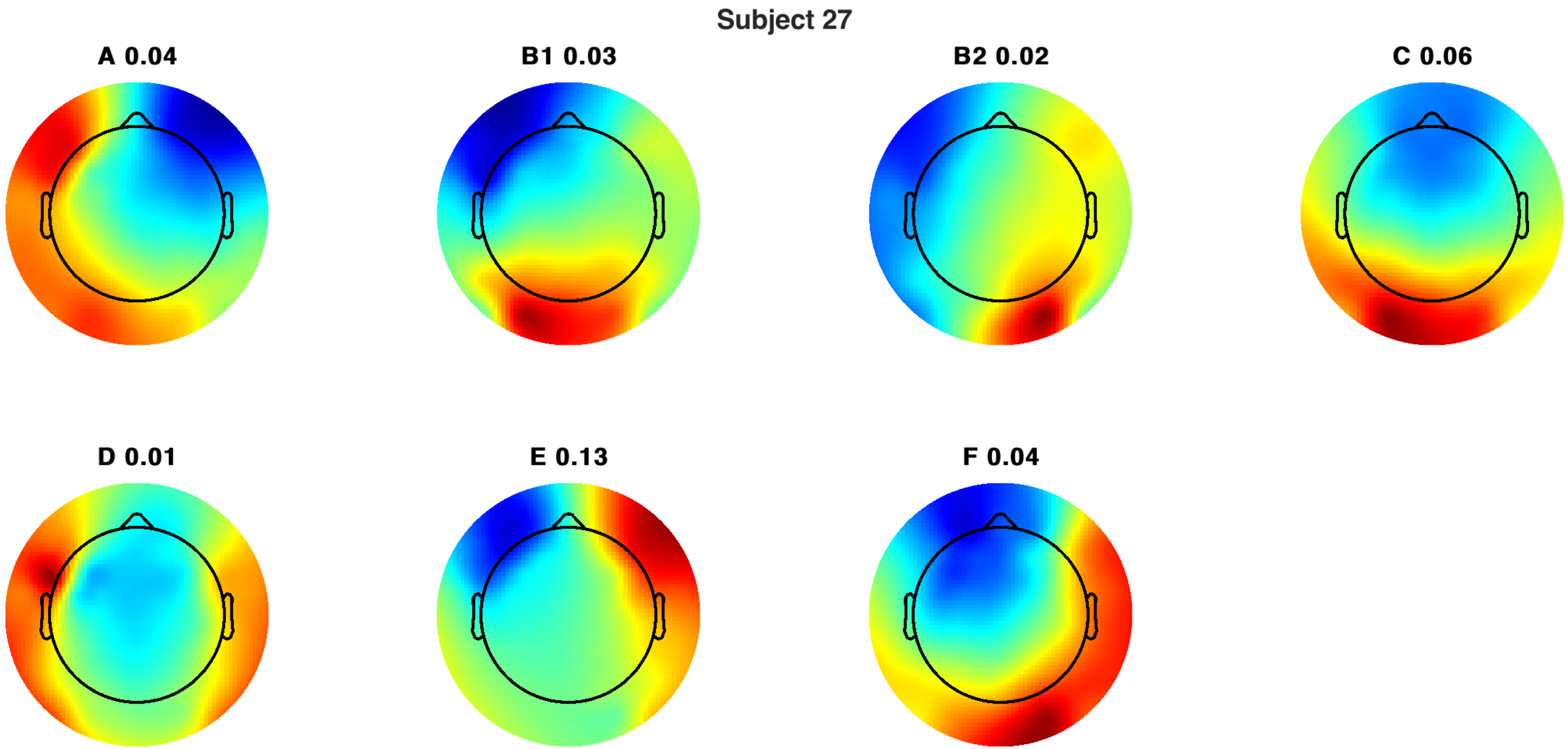
Subject 27
A 0.04
B1 0.03
B2 0.02
C 0.06
D 0.01
E 0.13
F 0.04

Subject 27

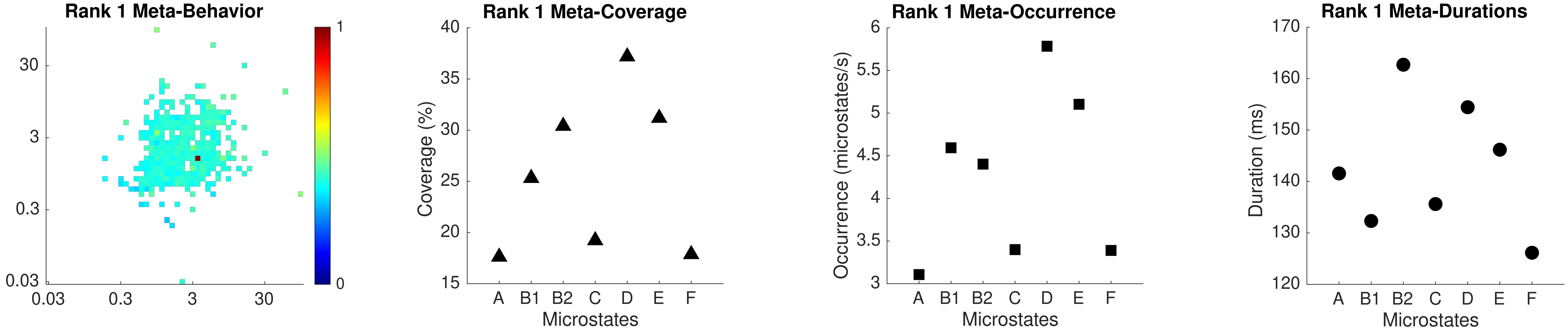

Rank 1 Meta-Behavior
Rank 1 Meta-Coverage
Coverage (%)
Microstates
Rank 1 Meta-Occurrence
Occurrence (microstates/s)
Microstates
Rank 1 Meta-Durations
Duration (ms)
Microstates
A B1 B2 C D E F

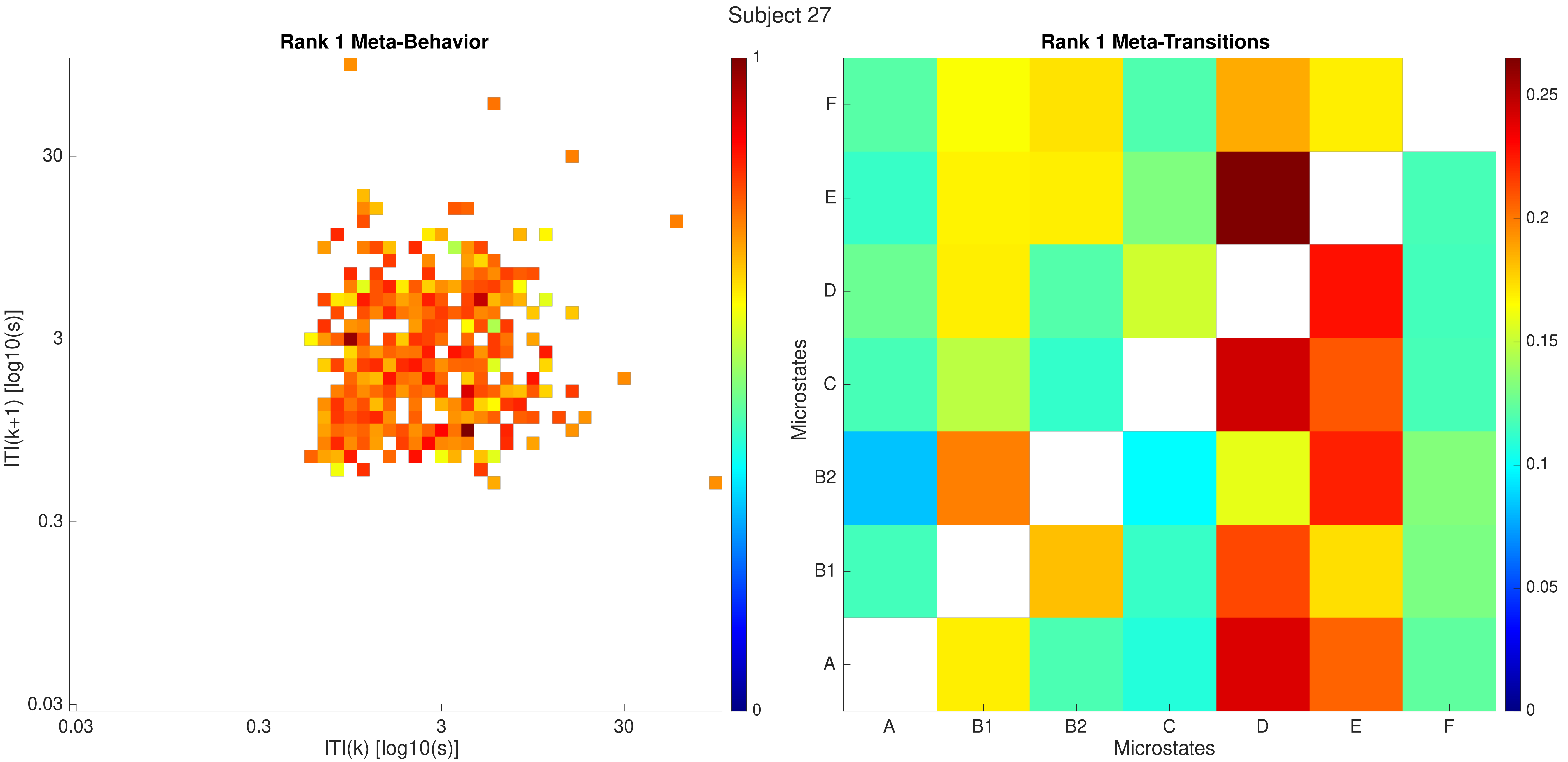

Subject 27
Rank 1 Meta-Behavior
ITI(k+1) [log10(s)]
ITI(k) [log10(s)]
0.03
0.3
3
30
1
0
Rank 1 Meta-Transitions
Microstates
A
B1
B2
C
D
E
F
0.25
0.2
0.15
0.1
0.05
0

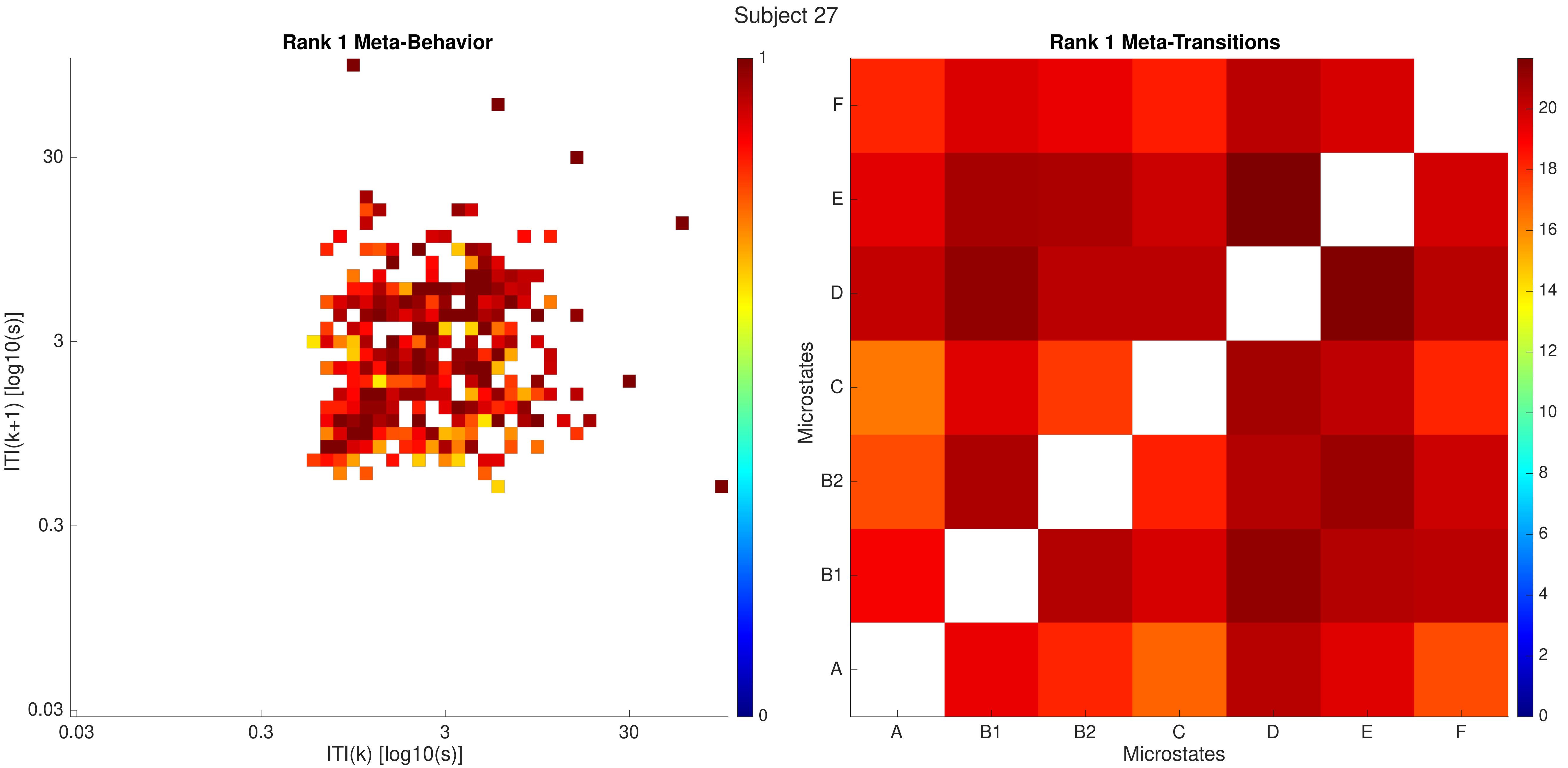
Subject 27
Rank 1 Meta-Behavior
ITI(k+1) [log10(s)]
ITI(k) [log10(s)]
0.03
0.3
3
30
0
1
Rank 1 Meta-Transitions
Microstates
A
B1
B2
C
D
E
F
0
2
4
6
8
10
12
14
16
18
20

Subject 27

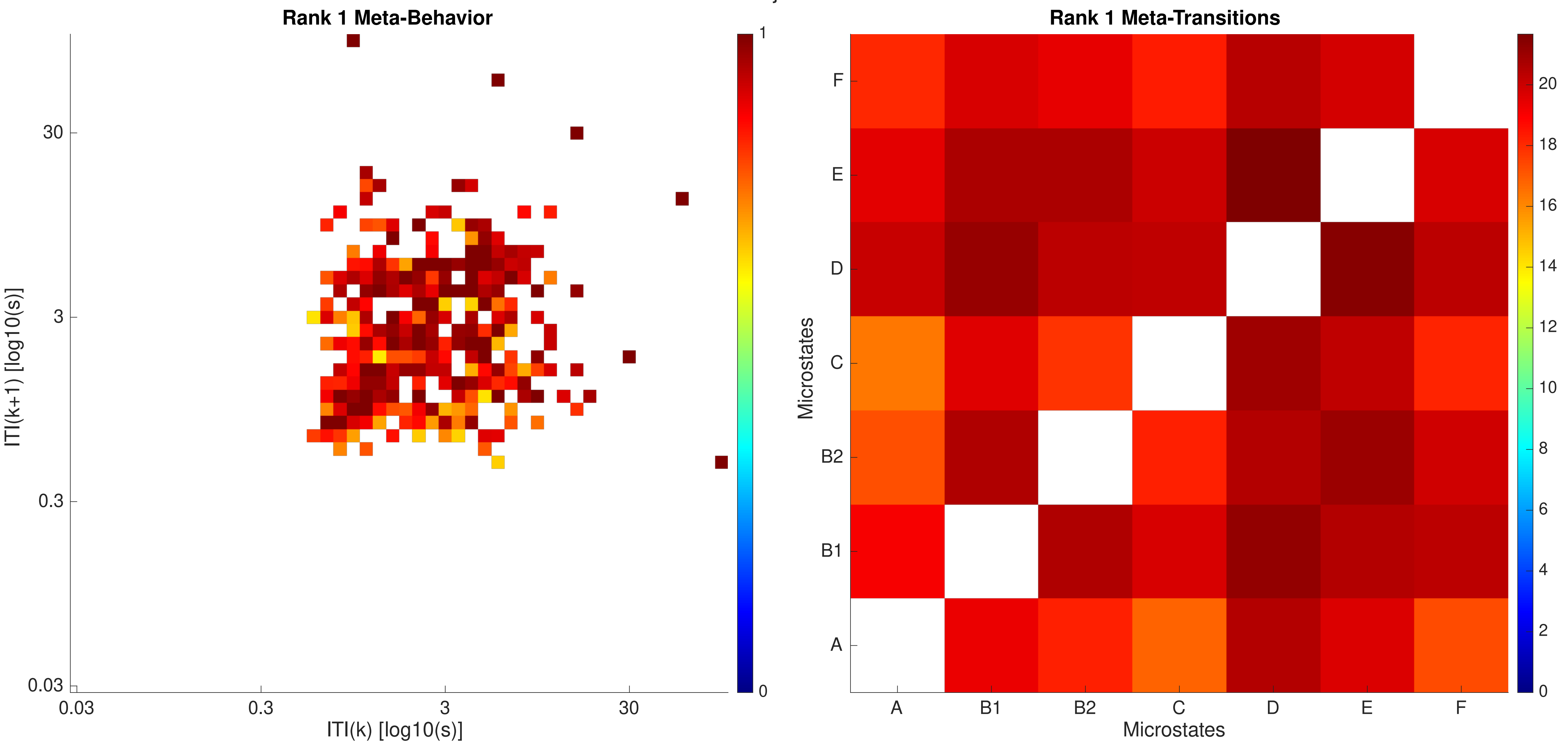

Rank 1 Meta-Behavior
ITI(k+1) [log10(s)]
ITI(k) [log10(s)]
0.03
0.3
3
30
1
0
Rank 1 Meta-Transitions
Microstates
A
B1
B2
C
D
E
F
0
2
4
6
8
10
12
14
16
18
20

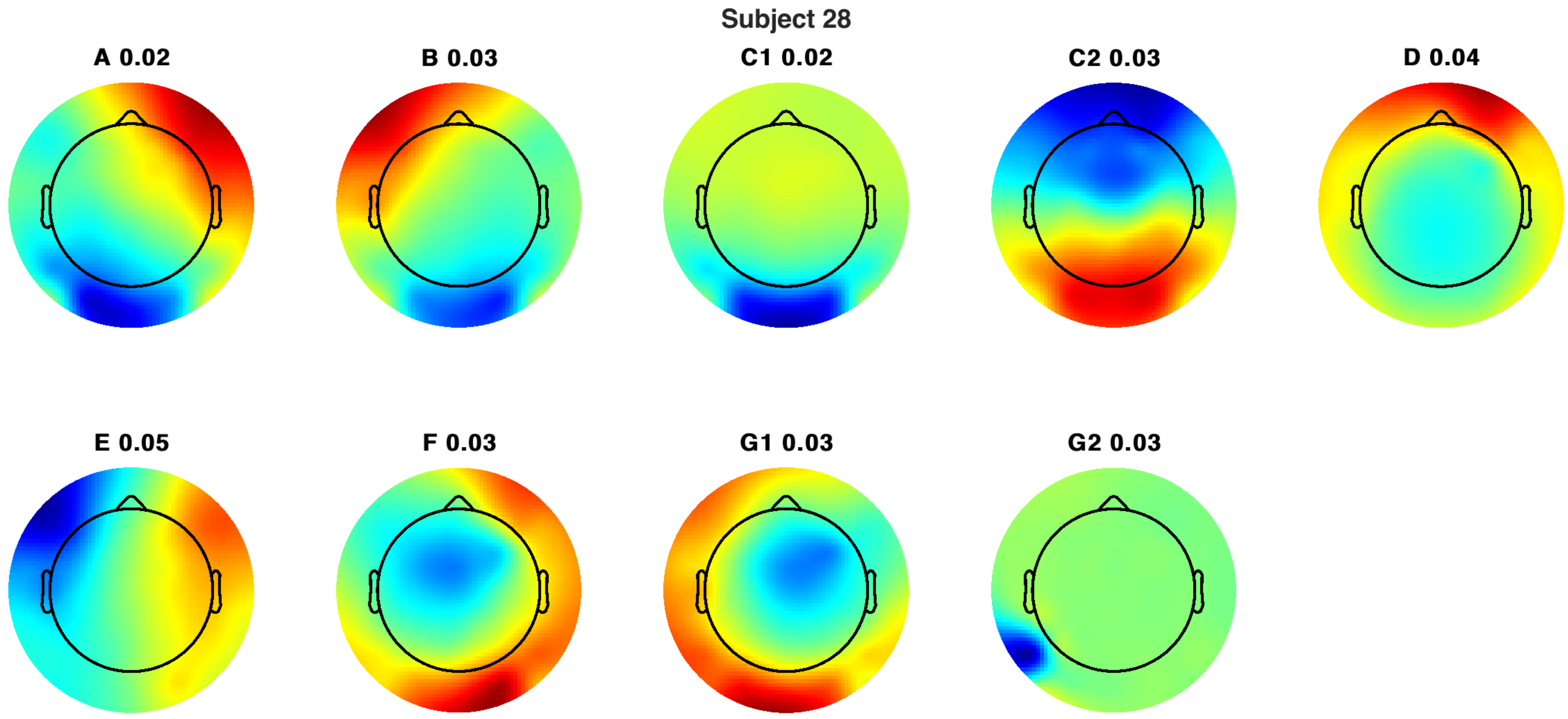
Subject 28
A 0.02
B 0.03
C1 0.02
C2 0.03
D 0.04
E 0.05
F 0.03
G1 0.03
G2 0.03

Subject 28

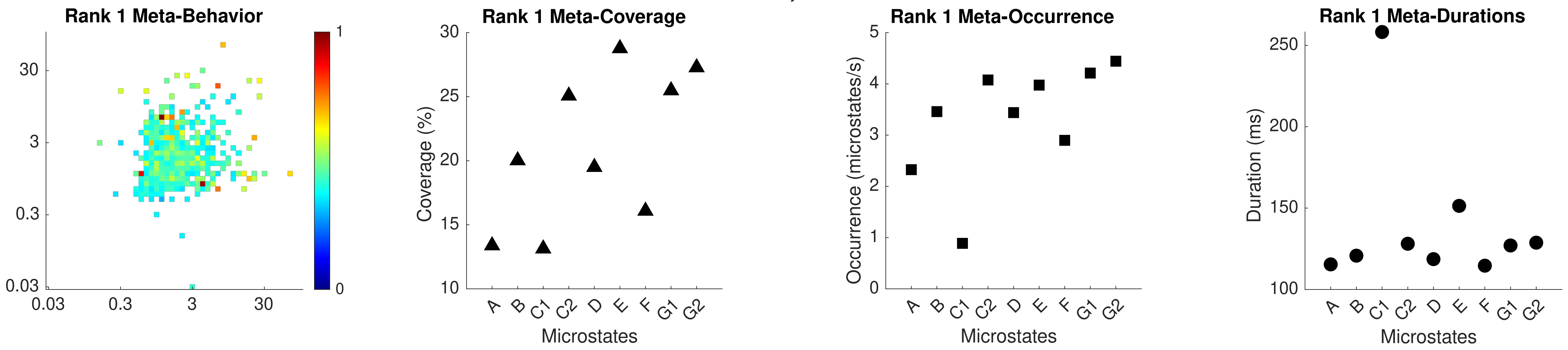

Rank 1 Meta-Behavior
Rank 1 Meta-Coverage
Rank 1 Meta-Occurrence
Rank 1 Meta-Durations
Coverage (%)
Occurrence (microstates/s)
Duration (ms)
Microstates
A B C1 C2 D E F G1 G2

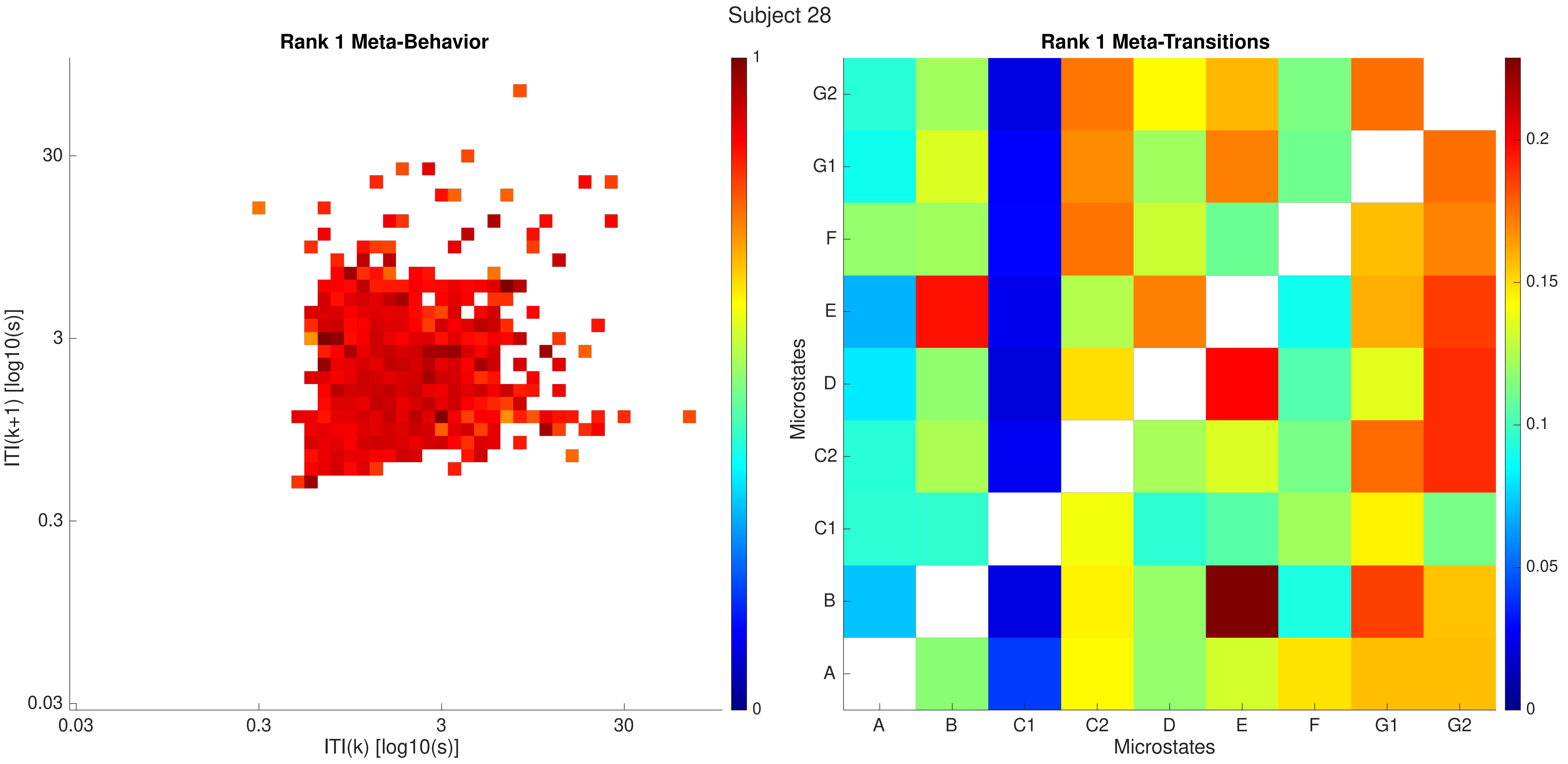

Subject 28
Rank 1 Meta-Behavior
ITI(k+1) [log10(s)]
ITI(k) [log10(s)]
0.03
0.3
3
30
1
0
Rank 1 Meta-Transitions
Microstates
A
B
C1
C2
D
E
F
G1
G2
0
0.05
0.1
0.15
0.2

Subject 28

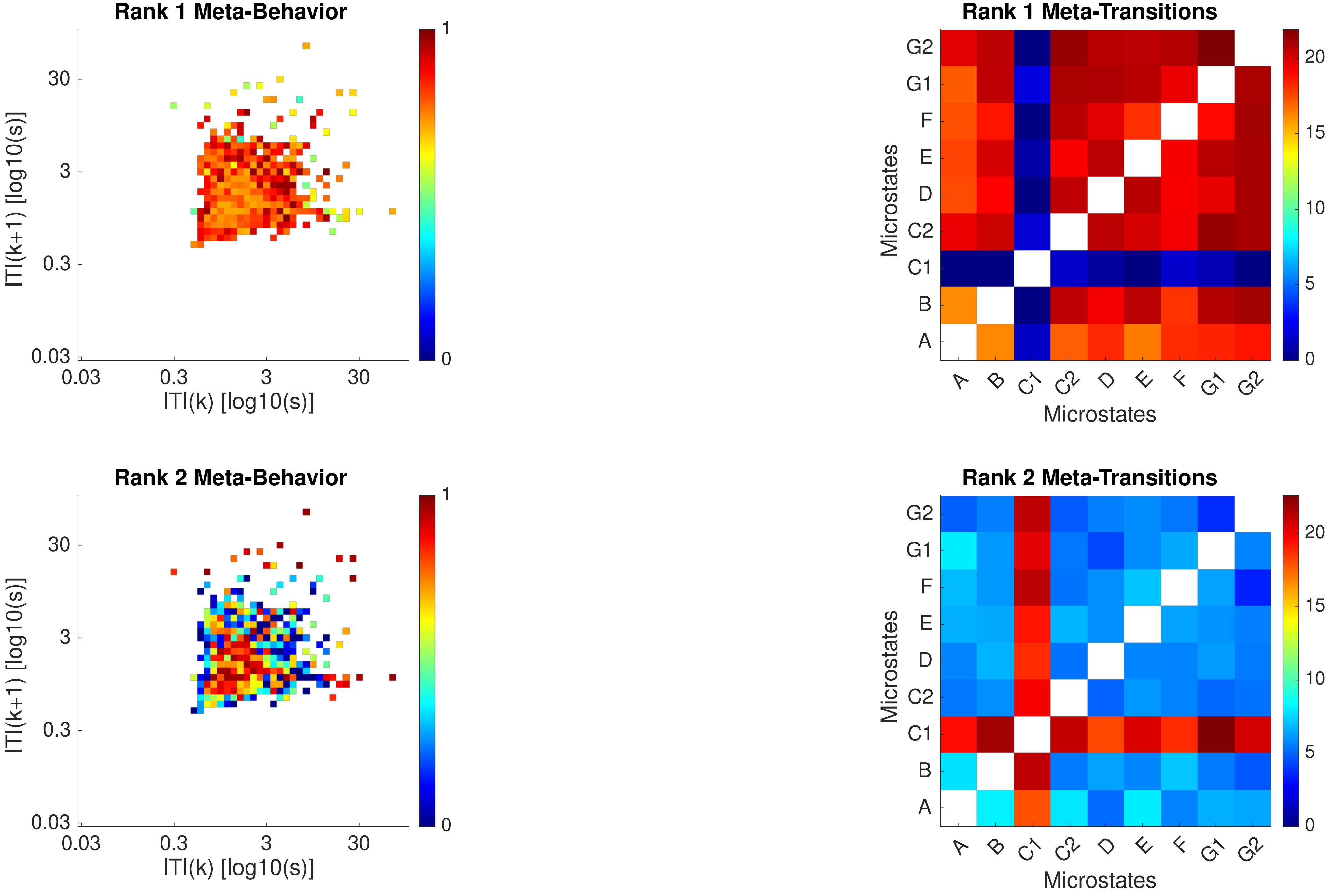
Rank 1 Meta-Behavior
Rank 1 Meta-Transitions
Rank 2 Meta-Behavior
Rank 2 Meta-Transitions
ITI(k+1) [log10(s)]
ITI(k) [log10(s)]
0.03
0.3
3
30
0
1
Microstates
A
B
C1
C2
D
E
F
G1
G2
0
5
10
15
20

Subject 28

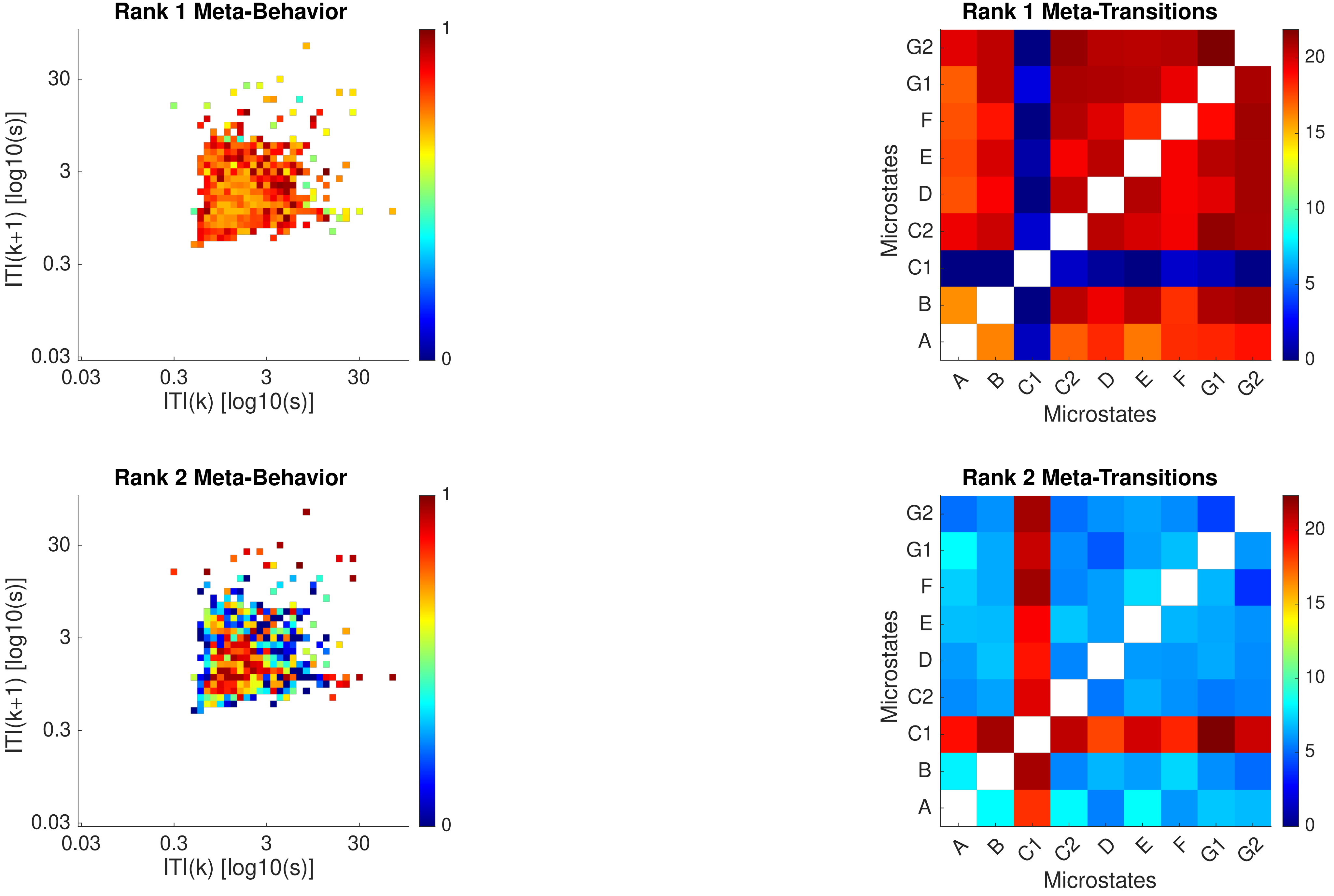

Rank 1 Meta-Behavior
ITI(k+1) [log10(s)]
ITI(k) [log10(s)]
0.03
0.3
3
30
0
1
Rank 1 Meta-Transitions
Microstates
A
B
C1
C2
D
E
F
G1
G2
0
5
10
15
20
Rank 2 Meta-Behavior
Rank 2 Meta-Transitions

Subject 29

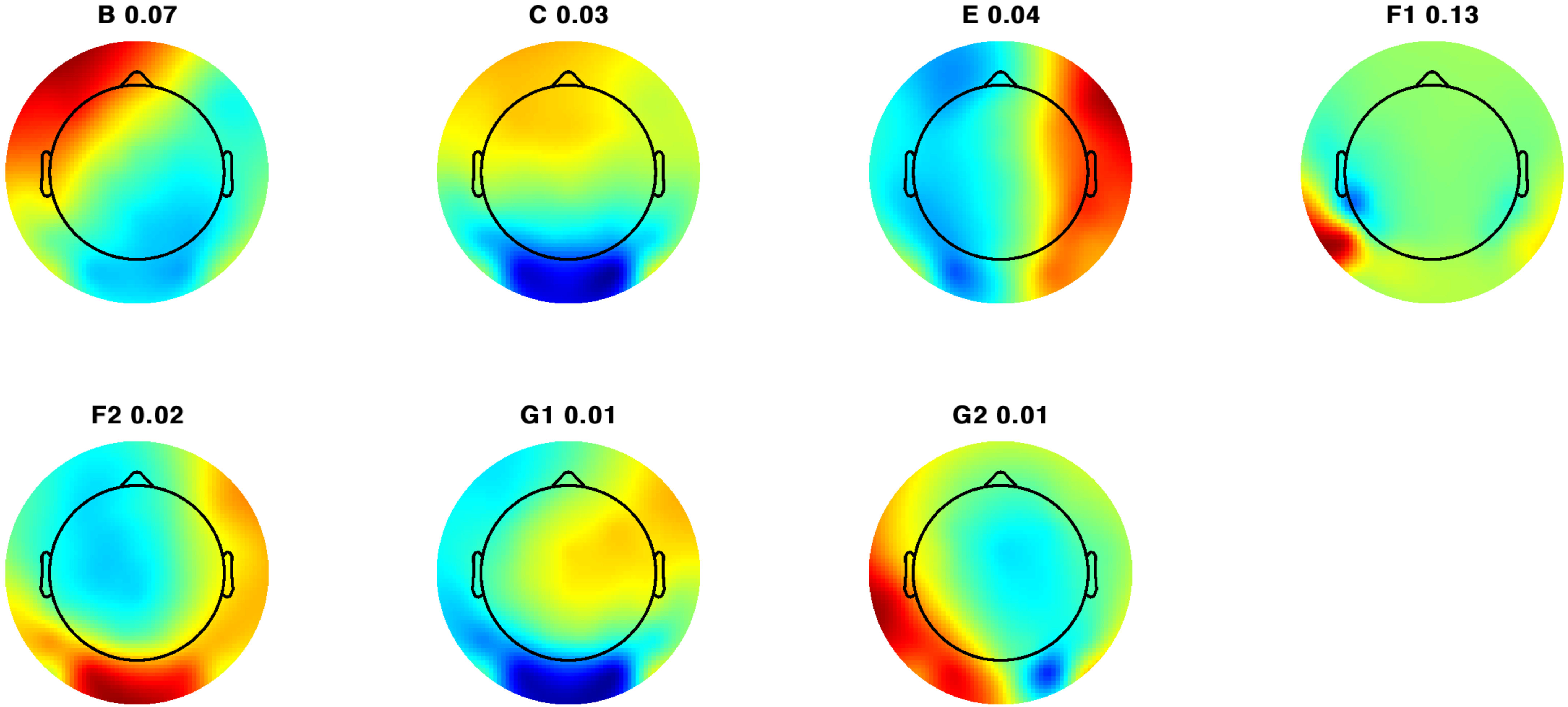

Subject 29

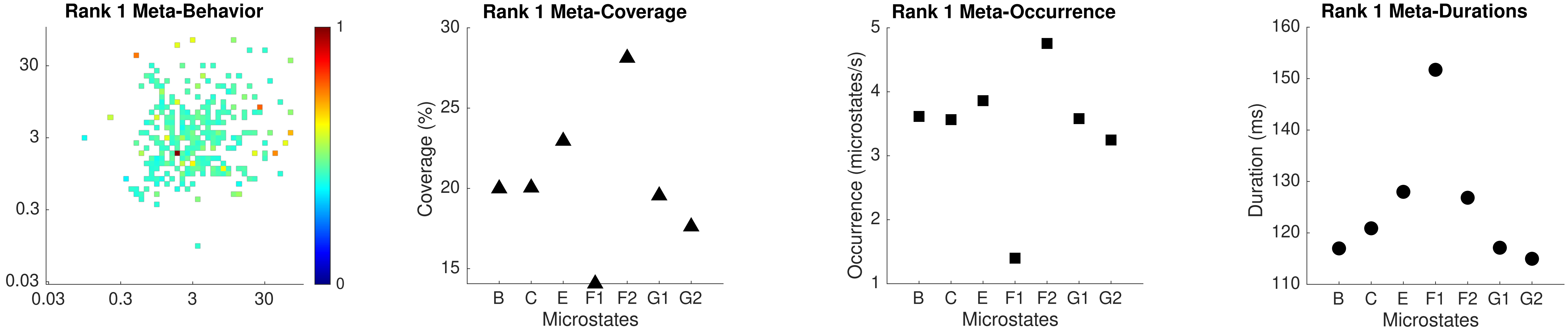
Rank 1 Meta-Behavior
30
3
0.3
0.03
0.03
0.3
3
30
1
0
Rank 1 Meta-Coverage
Coverage (%)
30
25
20
15
B
C
E
F1
F2
G1
G2
Microstates
Rank 1 Meta-Occurrence
Occurrence (microstates/s)
5
4
3
2
1
B
C
E
F1
F2
G1
G2
Microstates
Rank 1 Meta-Durations
Duration (ms)
160
150
140
130
120
110
B
C
E
F1
F2
G1
G2
Microstates

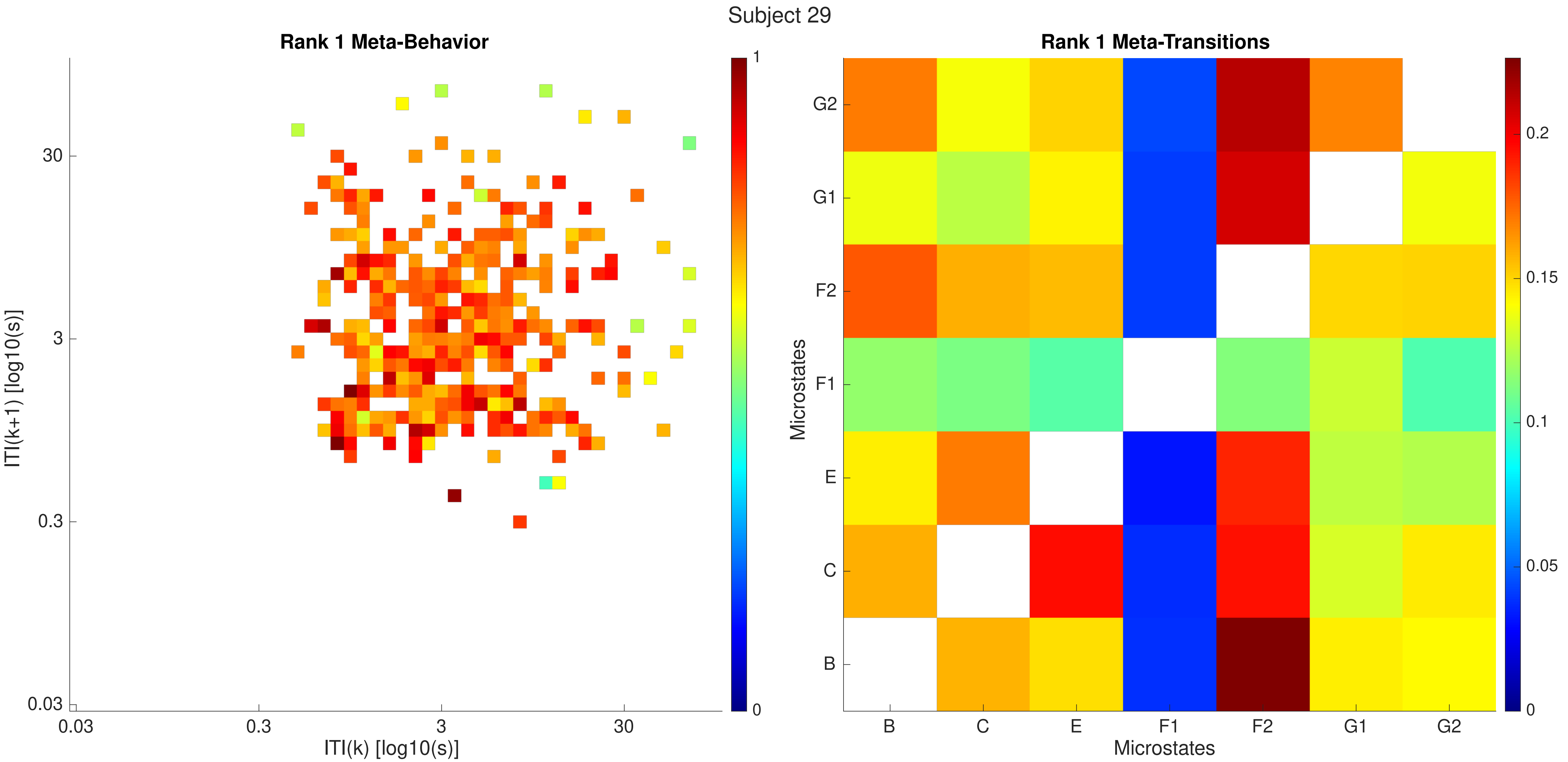

Subject 29
Rank 1 Meta-Behavior
ITI(k+1) [log10(s)]
ITI(k) [log10(s)]
0.03
0.3
3
30
1
0
Rank 1 Meta-Transitions
Microstates
G2
G1
F2
F1
E
C
B
0.2
0.15
0.1
0.05
0

Subject 29

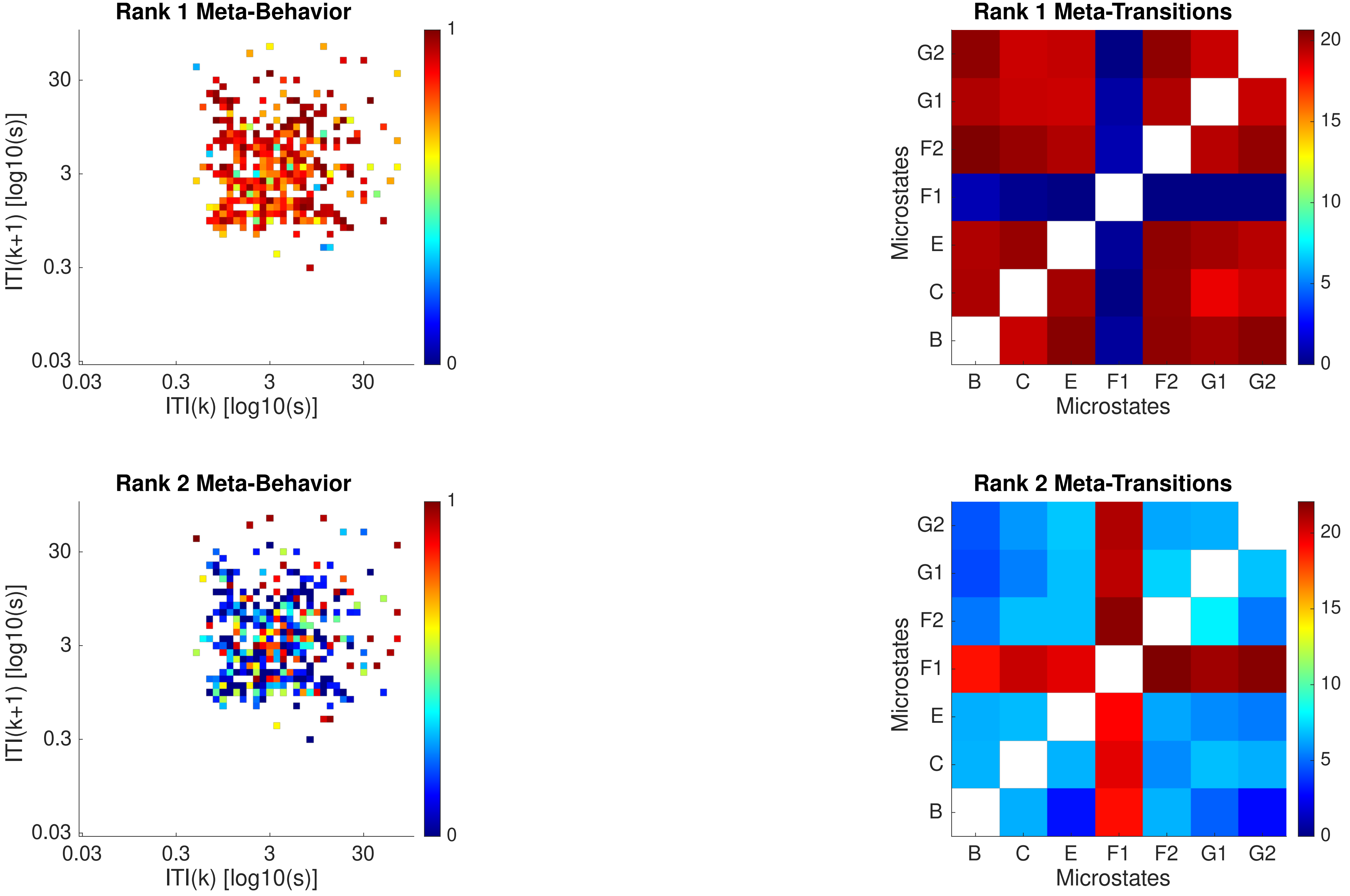
Rank 1 Meta-Behavior
ITI(k+1) [log10(s)]
ITI(k) [log10(s)]
Rank 1 Meta-Transitions
Microstates
B C E F1 F2 G1 G2
Rank 2 Meta-Behavior
ITI(k+1) [log10(s)]
ITI(k) [log10(s)]
Rank 2 Meta-Transitions
Microstates
B C E F1 F2 G1 G2

Subject 29

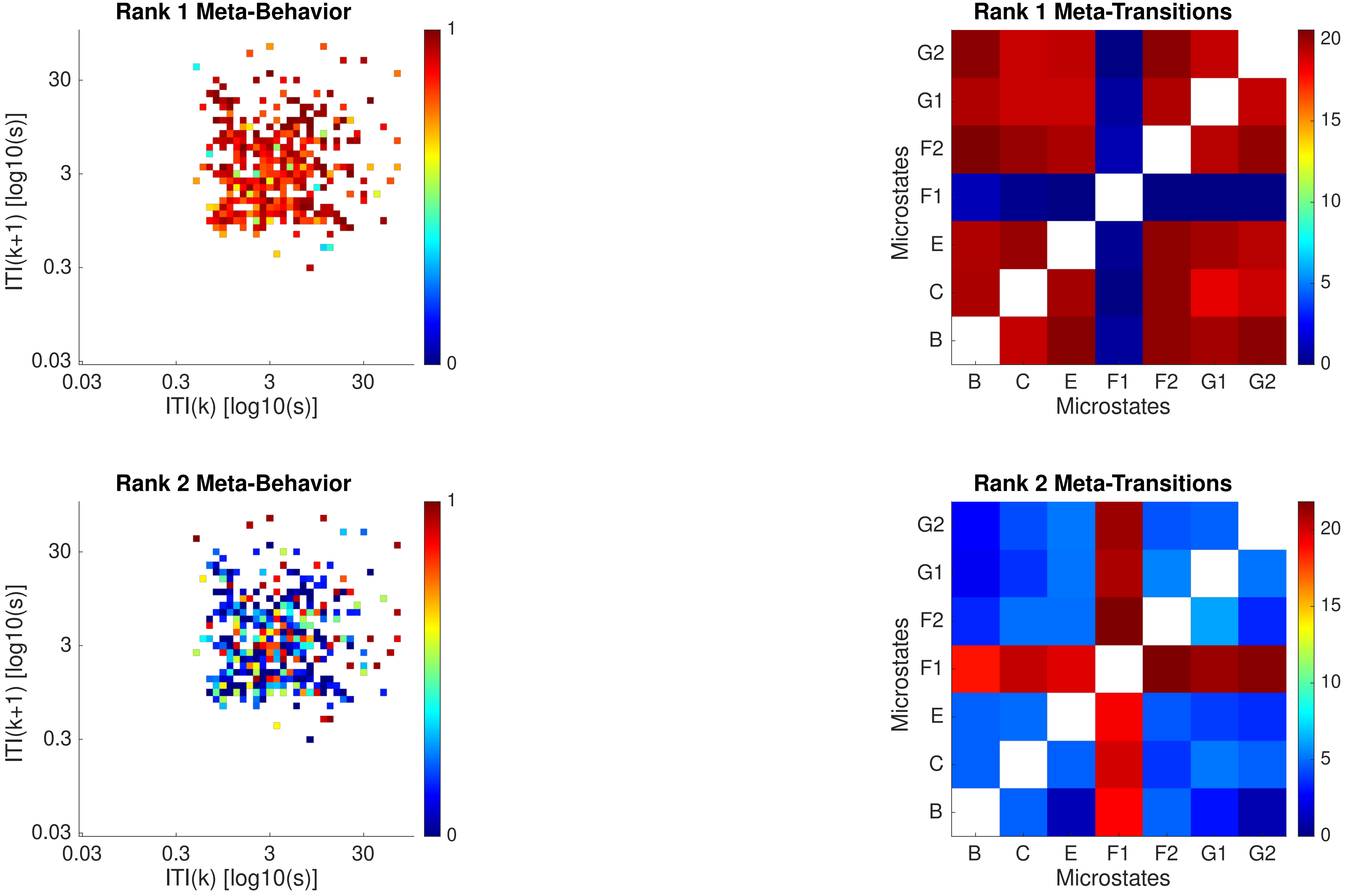

Rank 1 Meta-Behavior
ITI(k+1) [log10(s)]
ITI(k) [log10(s)]
0.03
0.3
3
30
0
1
Rank 1 Meta-Transitions
Microstates
B
C
E
F1
F2
G1
G2
0
5
10
15
20
Rank 2 Meta-Behavior
Rank 2 Meta-Transitions

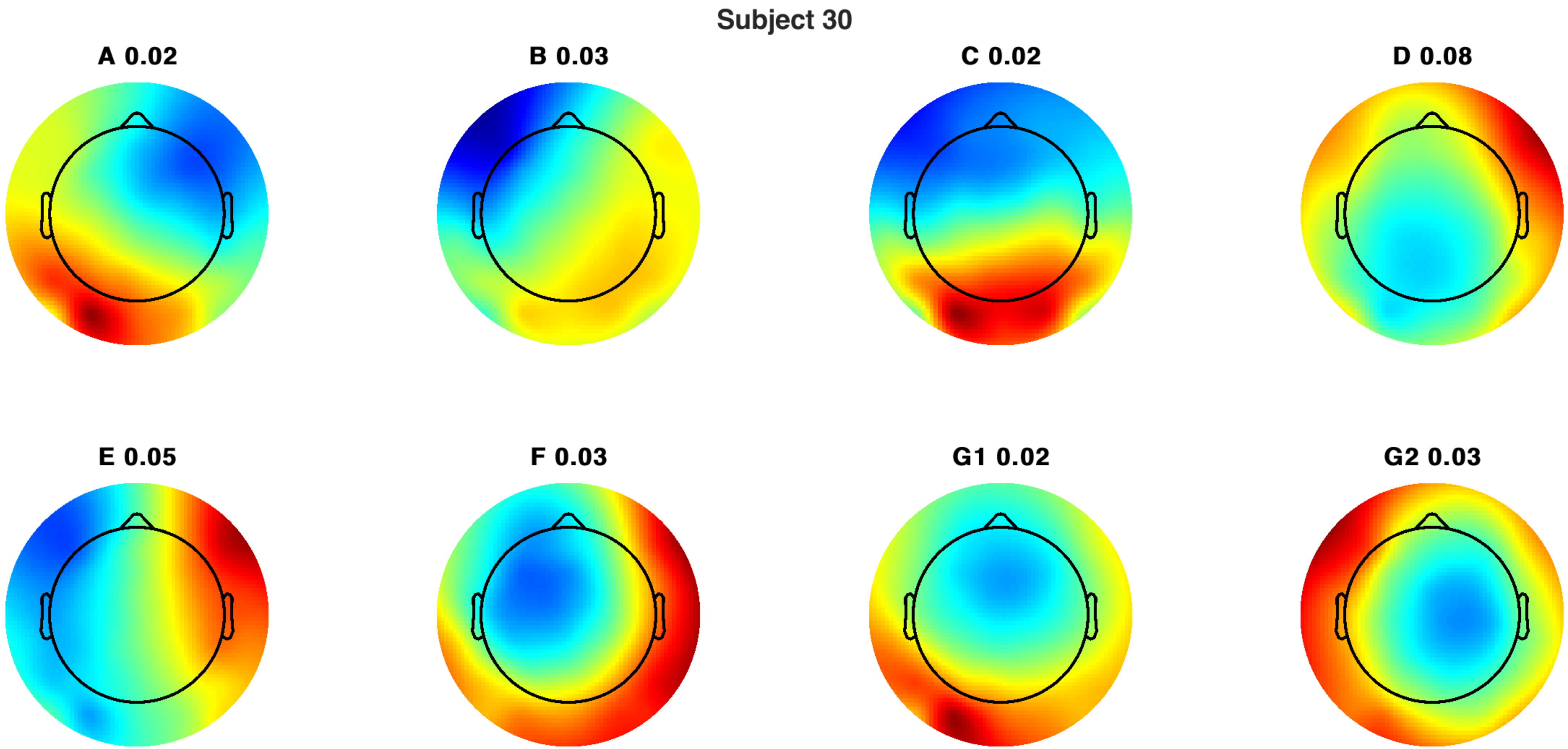
Subject 30
A 0.02
B 0.03
C 0.02
D 0.08
E 0.05
F 0.03
G1 0.02
G2 0.03

Subject 30

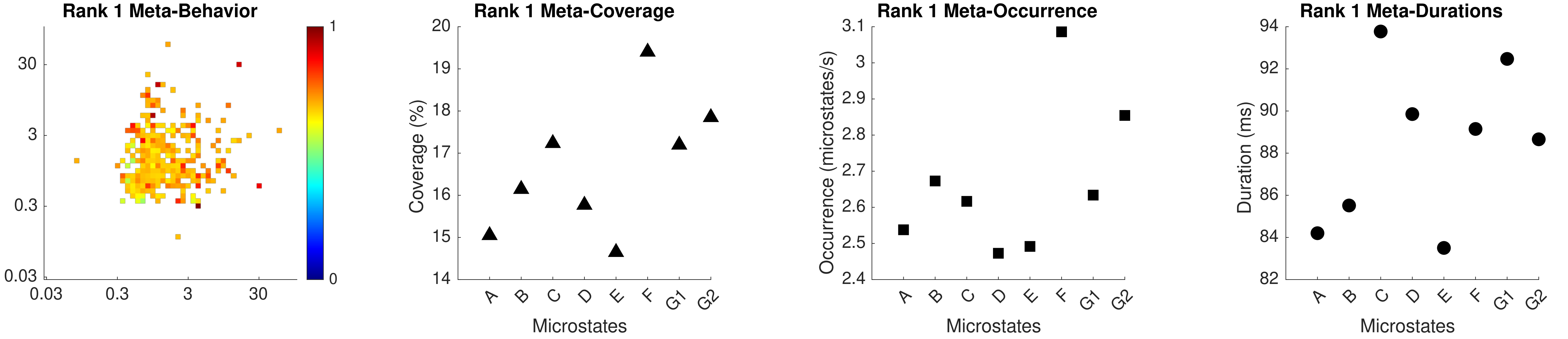
Rank 1 Meta-Behavior
Rank 1 Meta-Coverage
Rank 1 Meta-Occurrence
Rank 1 Meta-Durations
Coverage (%)
Occurrence (microstates/s)
Duration (ms)
Microstates
A B C D E F G1 G2

Subject 30

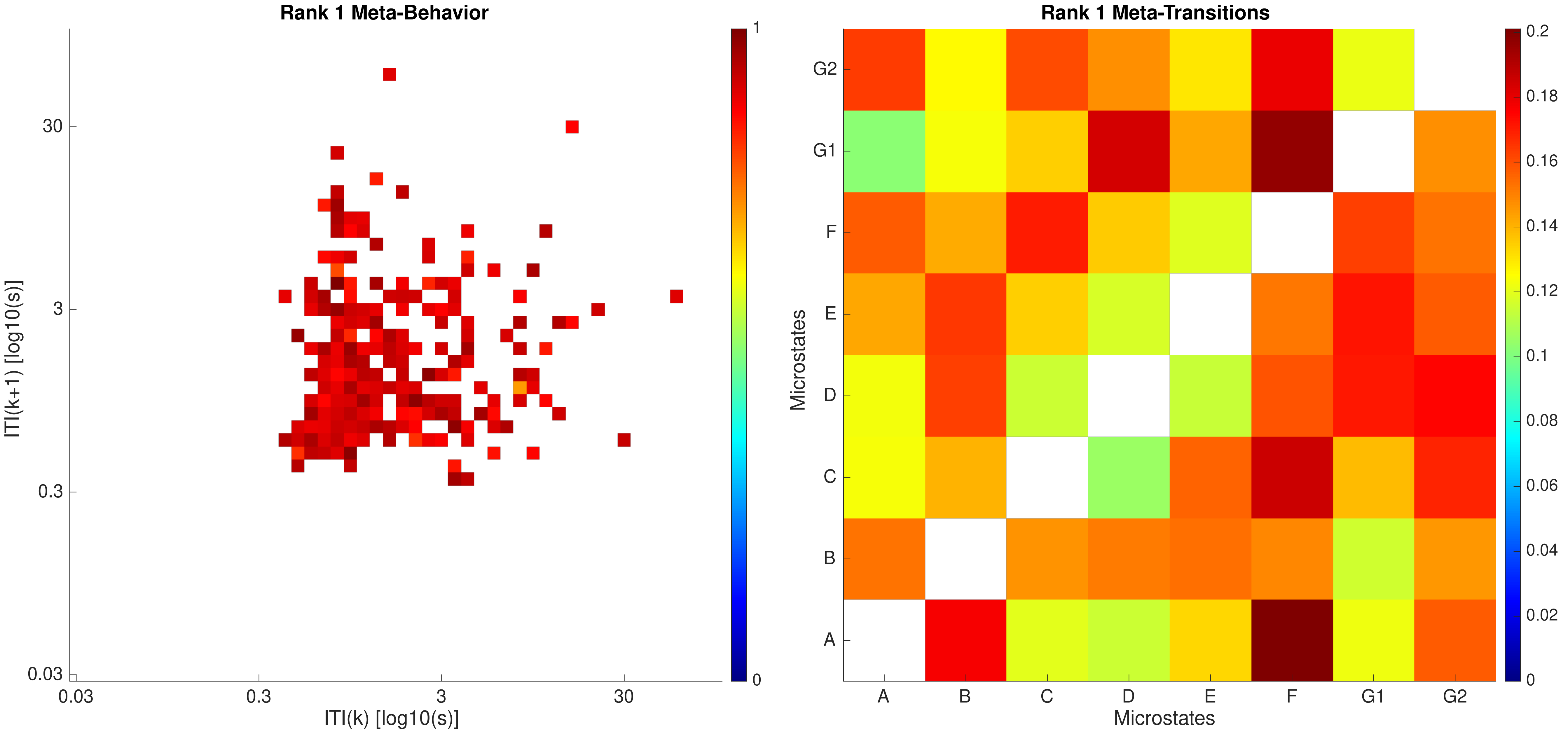
Rank 1 Meta-Behavior
ITI(k+1) [log10(s)]
ITI(k) [log10(s)]
0.03
0.3
3
30
1
0
Rank 1 Meta-Transitions
Microstates
A
B
C
D
E
F
G1
G2
0.2
0.18
0.16
0.14
0.12
0.1
0.08
0.06
0.04
0.02
0

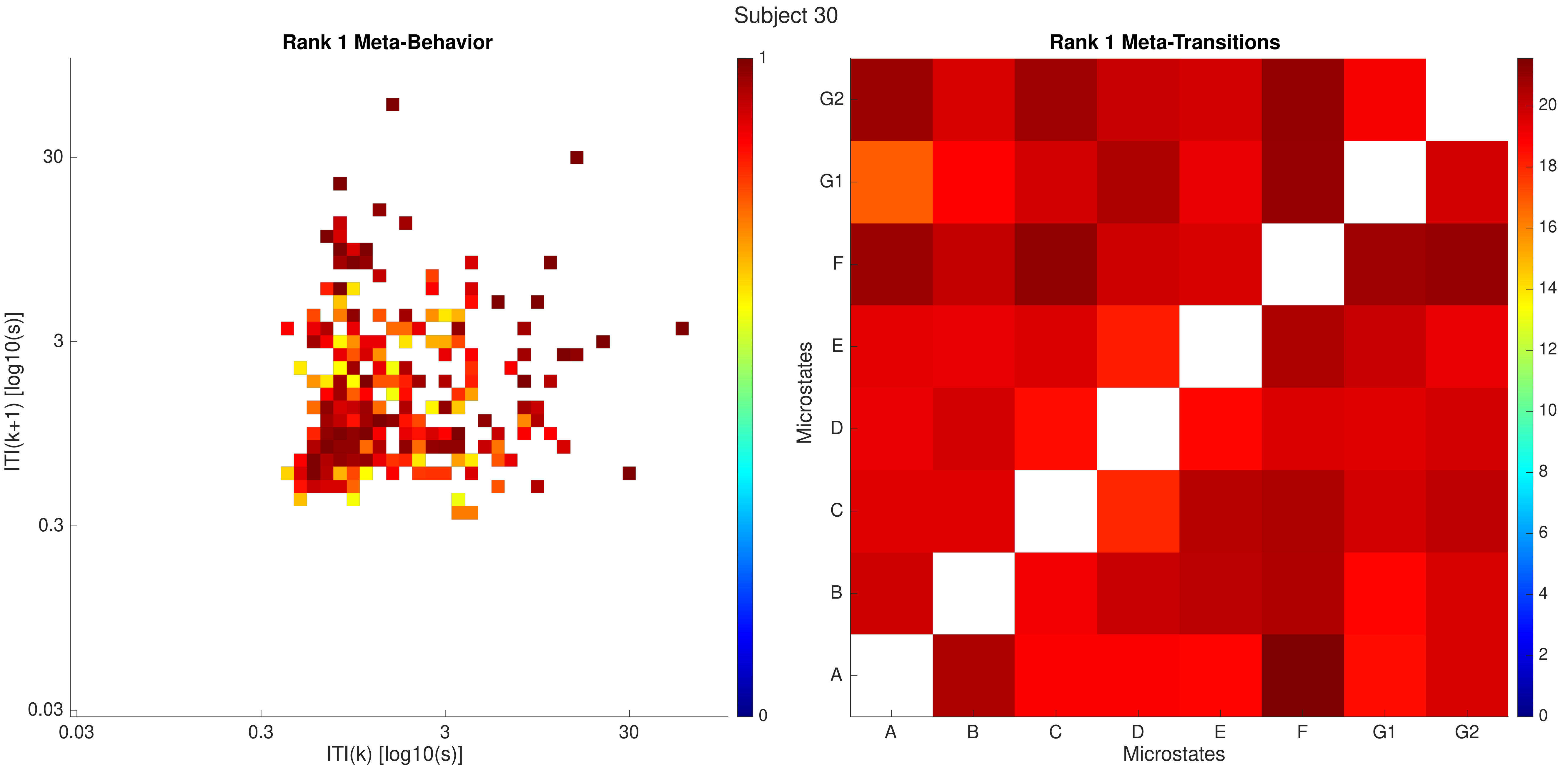

Subject 30
Rank 1 Meta-Behavior
ITI(k+1) [log10(s)]
30
3
0.3
0.03
0.03
0.3
3
30
ITI(k) [log10(s)]
1
0
Rank 1 Meta-Transitions
Microstates
G2
G1
F
E
D
C
B
A
A
B
C
D
E
F
G1
G2
Microstates
20
18
16
14
12
10
8
6
4
2
0

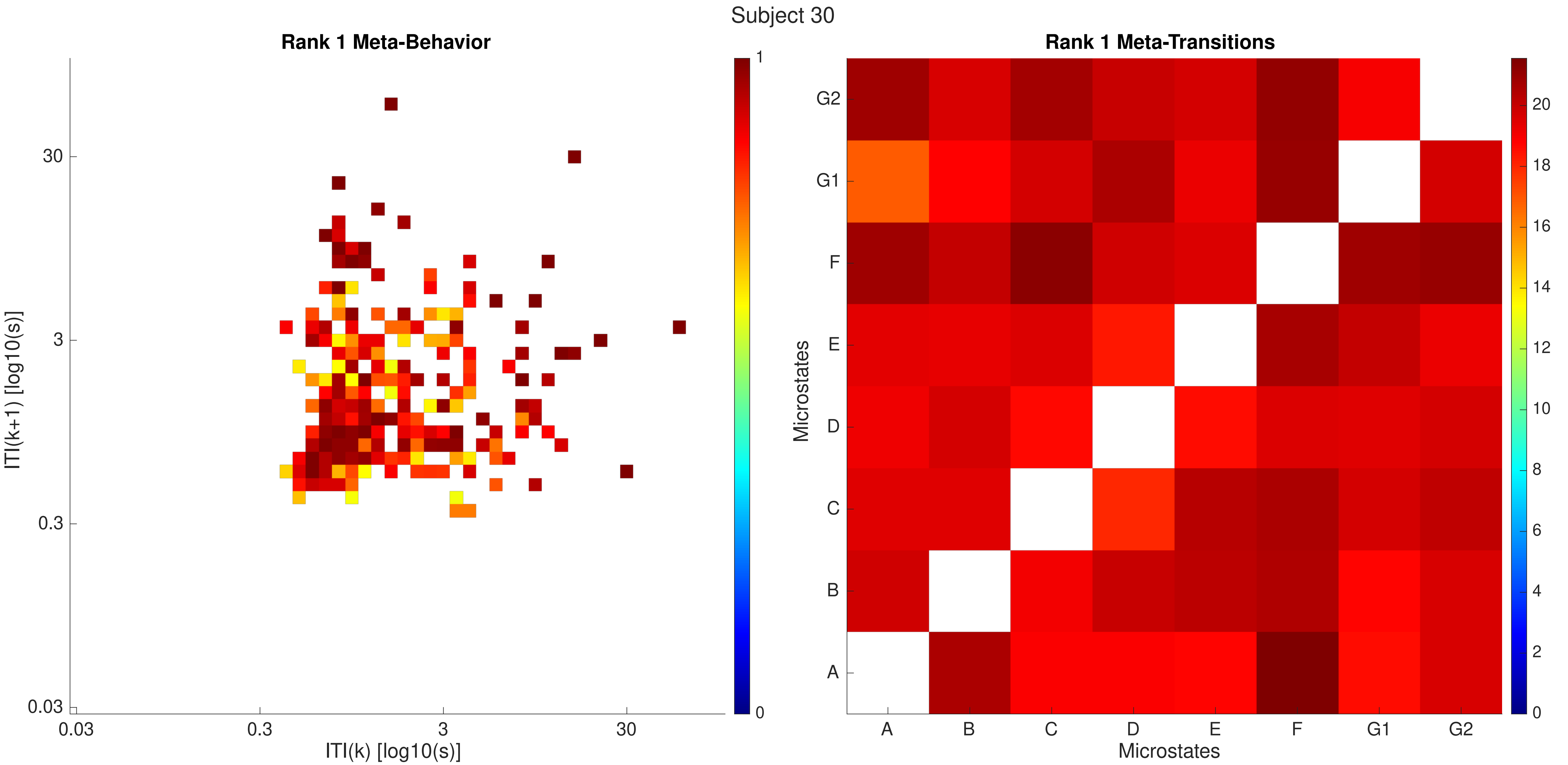

Subject 30
Rank 1 Meta-Behavior
ITI(k+1) [log10(s)]
ITI(k) [log10(s)]
0.03
0.3
3
30
0
1
Rank 1 Meta-Transitions
Microstates
A
B
C
D
E
F
G1
G2
0
2
4
6
8
10
12
14
16
18
20

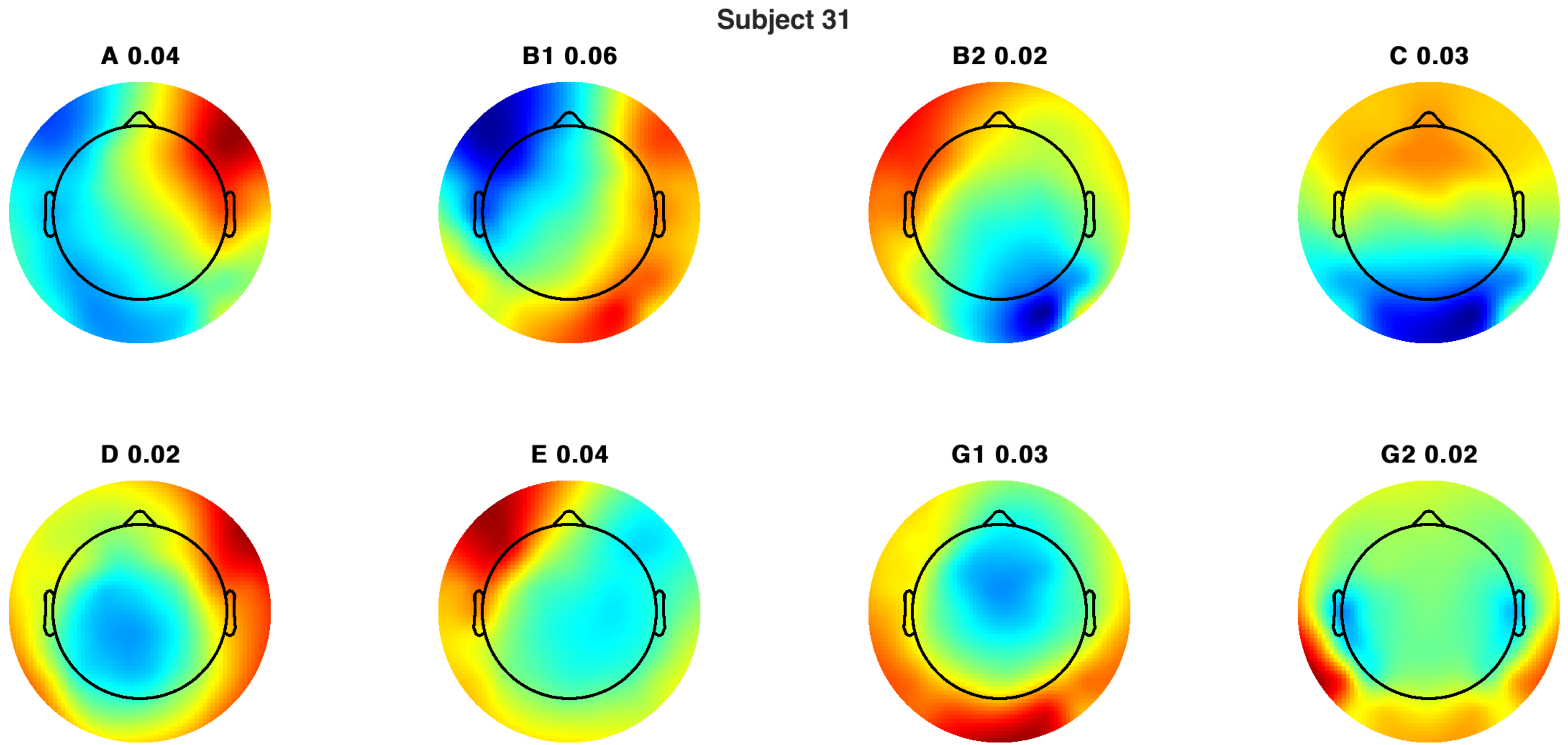
Subject 31
A 0.04
B1 0.06
B2 0.02
C 0.03
D 0.02
E 0.04
G1 0.03
G2 0.02

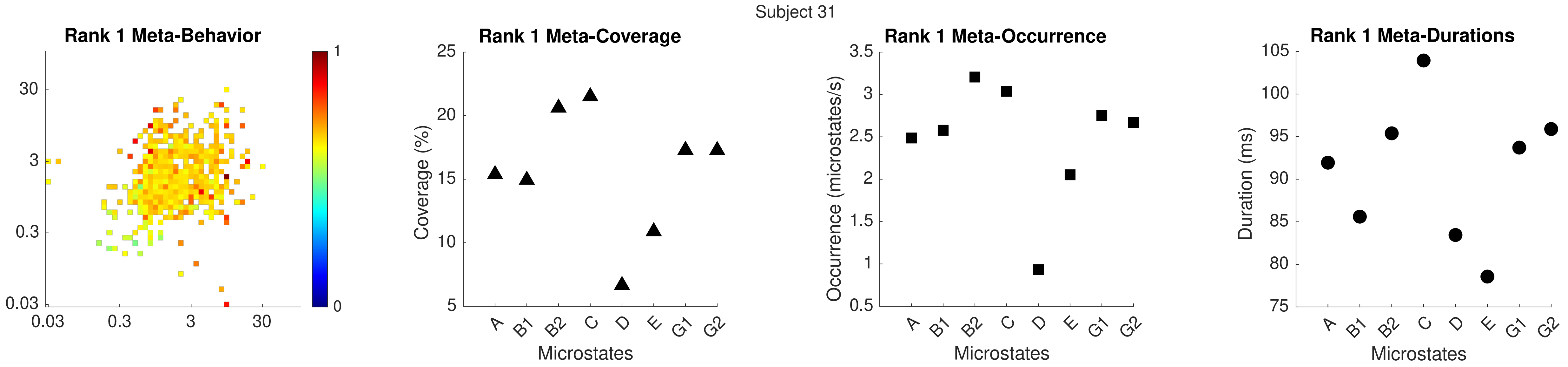

Subject 31
Rank 1 Meta-Behavior
Rank 1 Meta-Coverage
Coverage (%)
Microstates
Rank 1 Meta-Occurrence
Occurrence (microstates/s)
Rank 1 Meta-Durations
Duration (ms)
A B1 B2 C D E G1 G2

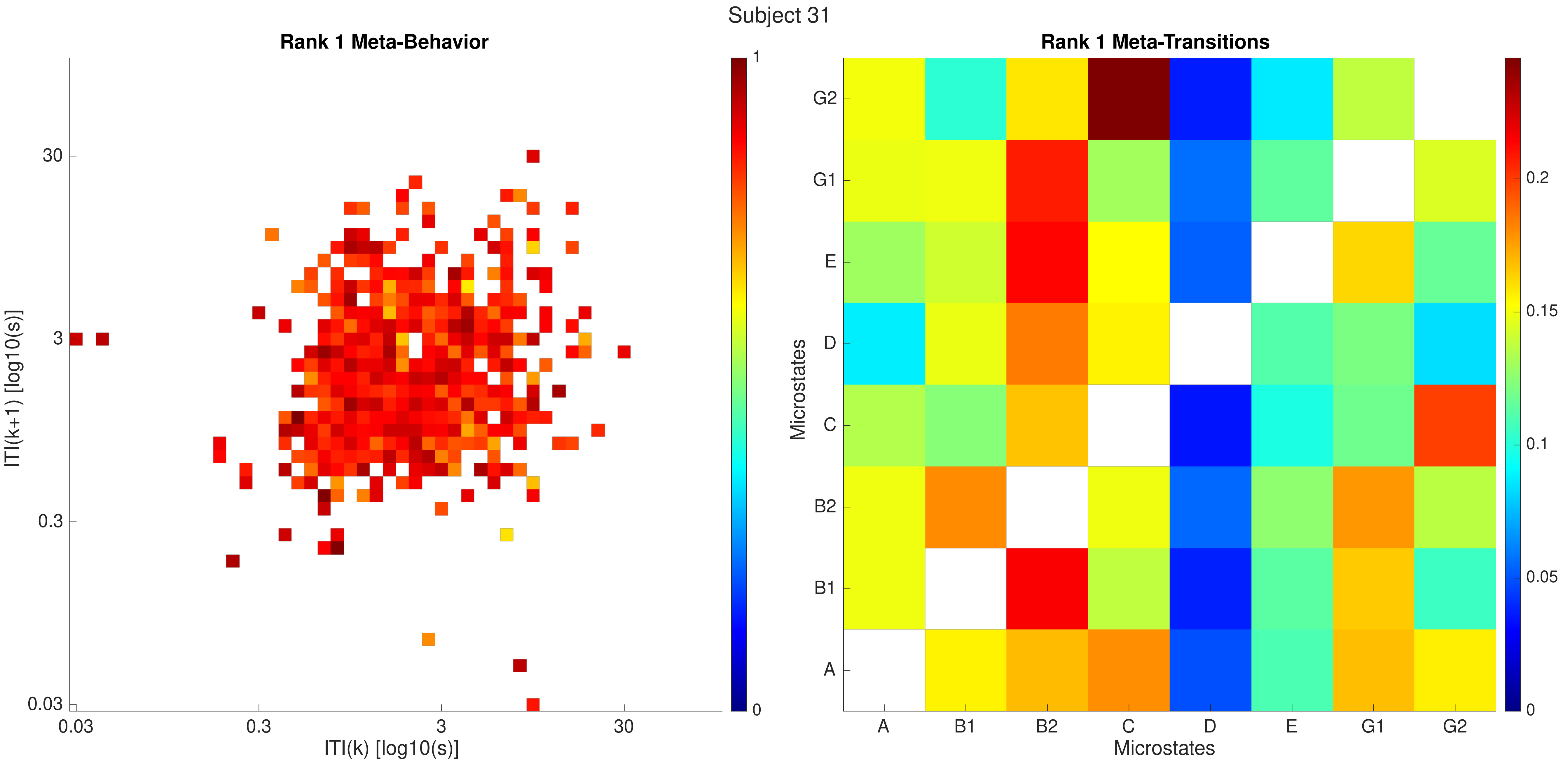

Subject 31
Rank 1 Meta-Behavior
ITI(k+1) [log10(s)]
ITI(k) [log10(s)]
0.03
0.3
3
30
1
0
Rank 1 Meta-Transitions
Microstates
A
B1
B2
C
D
E
G1
G2
0.2
0.15
0.1
0.05

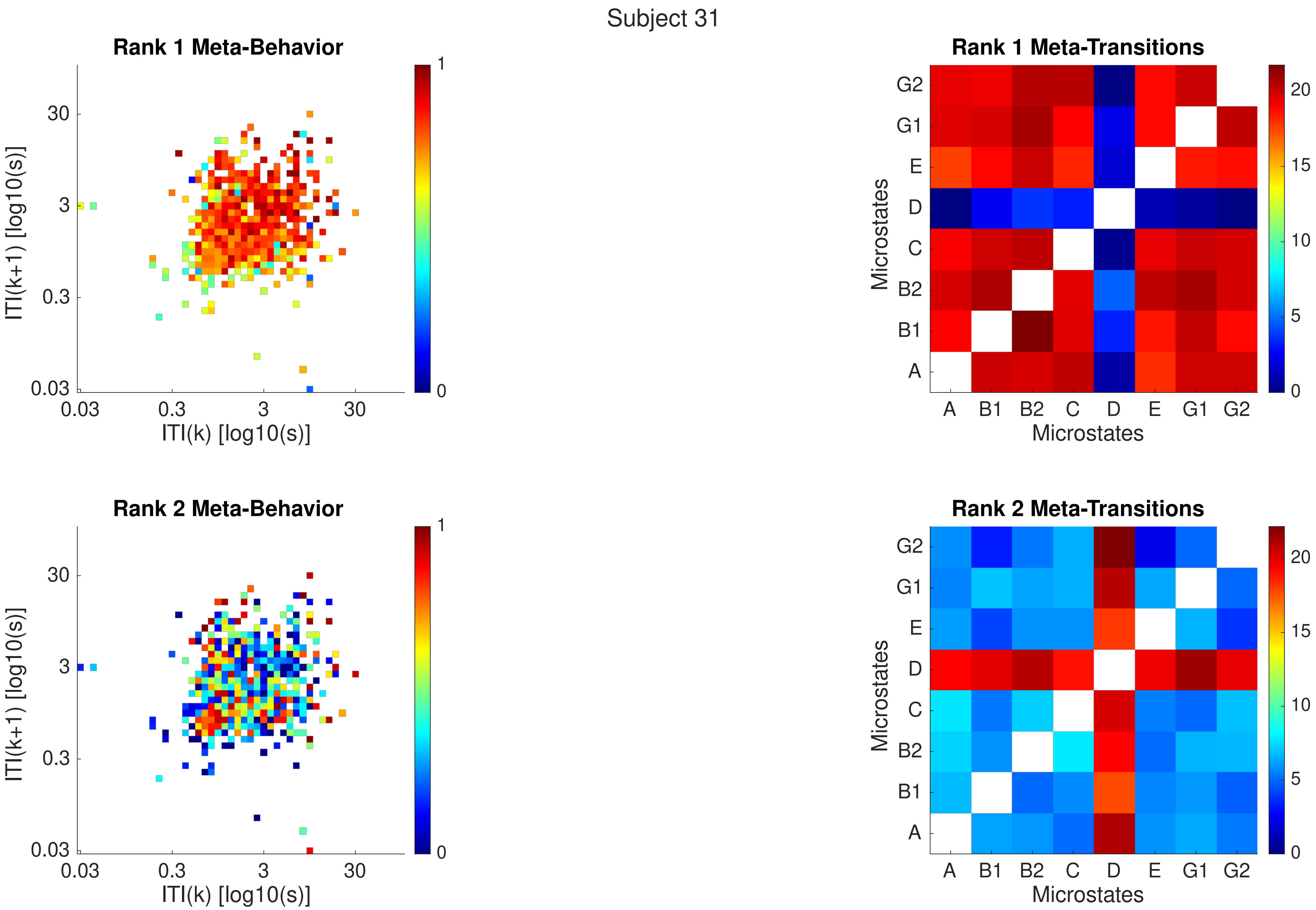
Subject 31
Rank 1 Meta-Behavior
ITI(k+1) [log10(s)]
ITI(k) [log10(s)]
30
3
0.3
0.03
0.03
0.3
3
30
1
0
Rank 1 Meta-Transitions
Microstates
Microstates
G2
G1
E
D
C
B2
B1
A
A
B1
B2
C
D
E
G1
G2
20
15
10
5
0
Rank 2 Meta-Behavior
ITI(k+1) [log10(s)]
ITI(k) [log10(s)]
30
3
0.3
0.03
0.03
0.3
3
30
1
0
Rank 2 Meta-Transitions
Microstates
Microstates
G2
G1
E
D
C
B2
B1
A
A
B1
B2
C
D
E
G1
G2
20
15
10
5
0

Subject 31

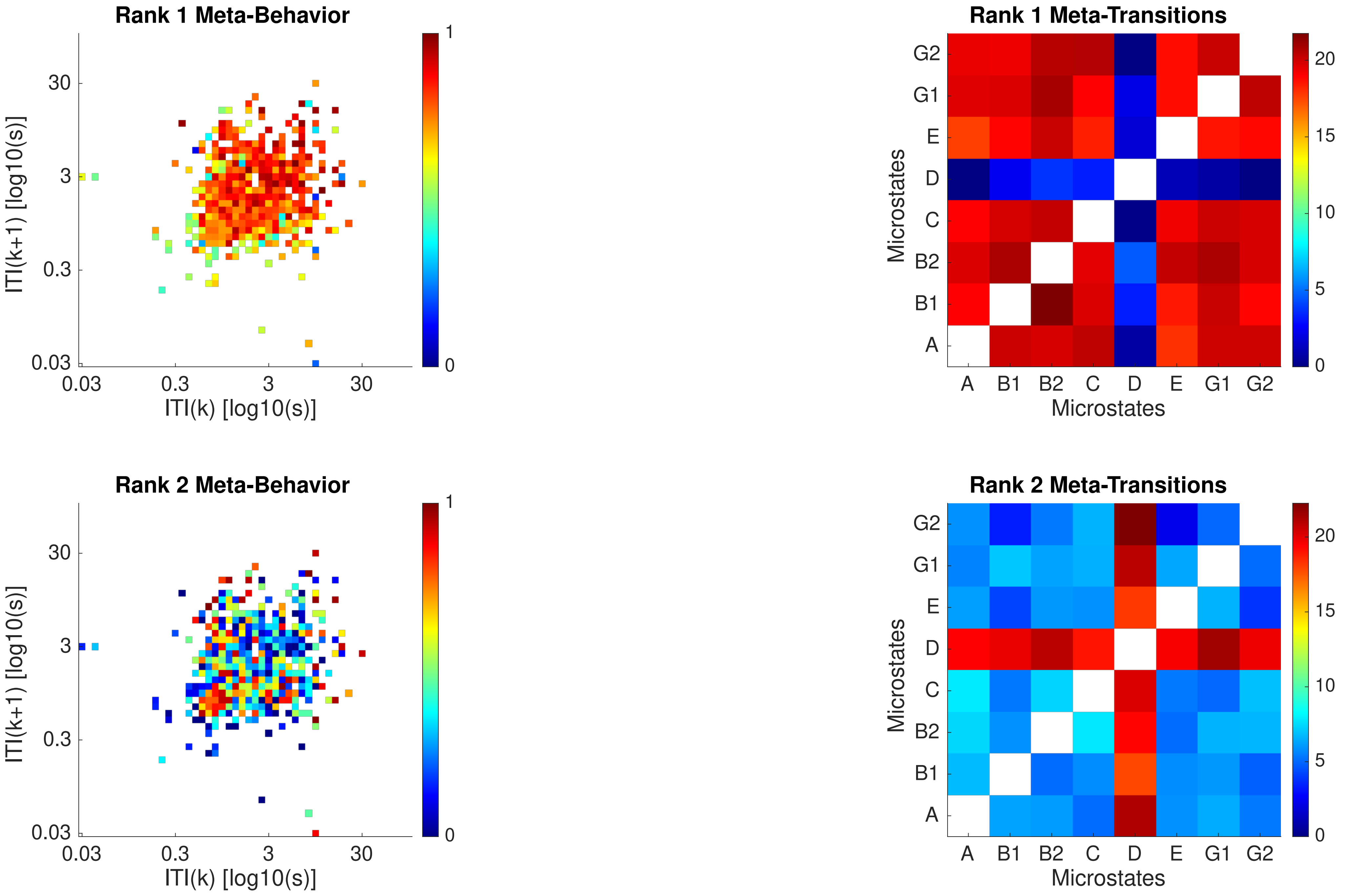

Rank 1 Meta-Behavior
ITI(k+1) [log10(s)]
ITI(k) [log10(s)]
0.03
0.3
3
30
0
1
Rank 1 Meta-Transitions
Microstates
A
B1
B2
C
D
E
G1
G2
0
5
10
15
20
Rank 2 Meta-Behavior
ITI(k+1) [log10(s)]
ITI(k) [log10(s)]
Rank 2 Meta-Transitions
Microstates

Subject 32

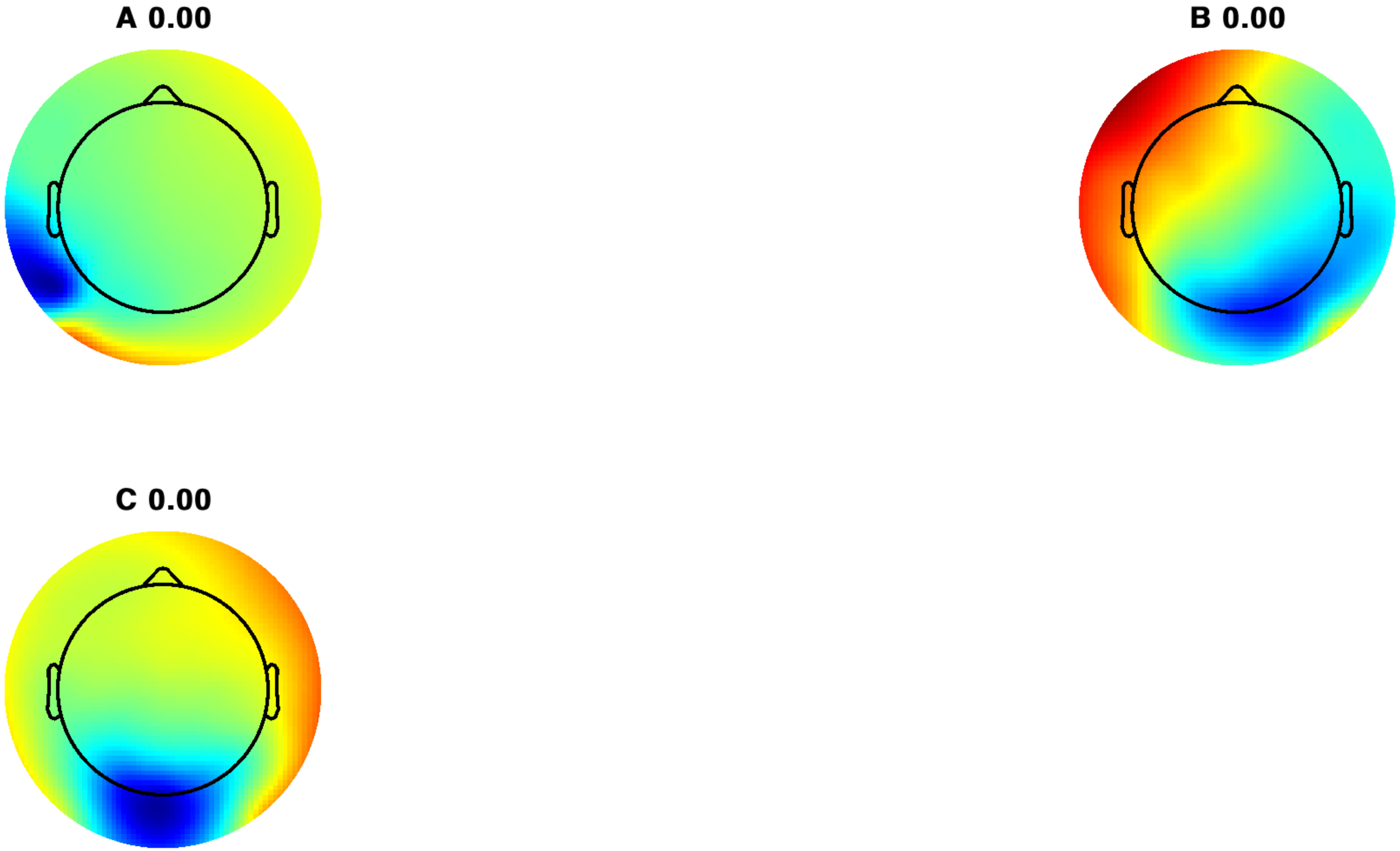

Subject 32

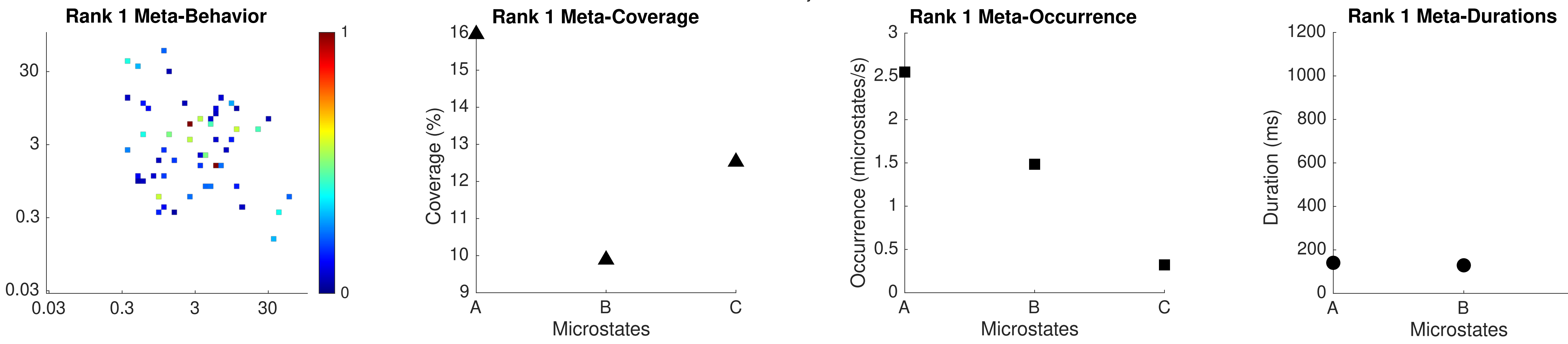

Rank 1 Meta-Behavior
30
3
0.3
0.03
0.03
0.3
3
30
1
0
Rank 1 Meta-Coverage
Coverage (%)
16
15
14
13
12
11
10
9
A
B
C
Microstates
Rank 1 Meta-Occurrence
Occurrence (microstates/s)
3
2.5
2
1.5
1
0.5
0
A
B
C
Microstates
Rank 1 Meta-Durations
Duration (ms)
1200
1000
800
600
400
200
0
A
B
C
Microstates

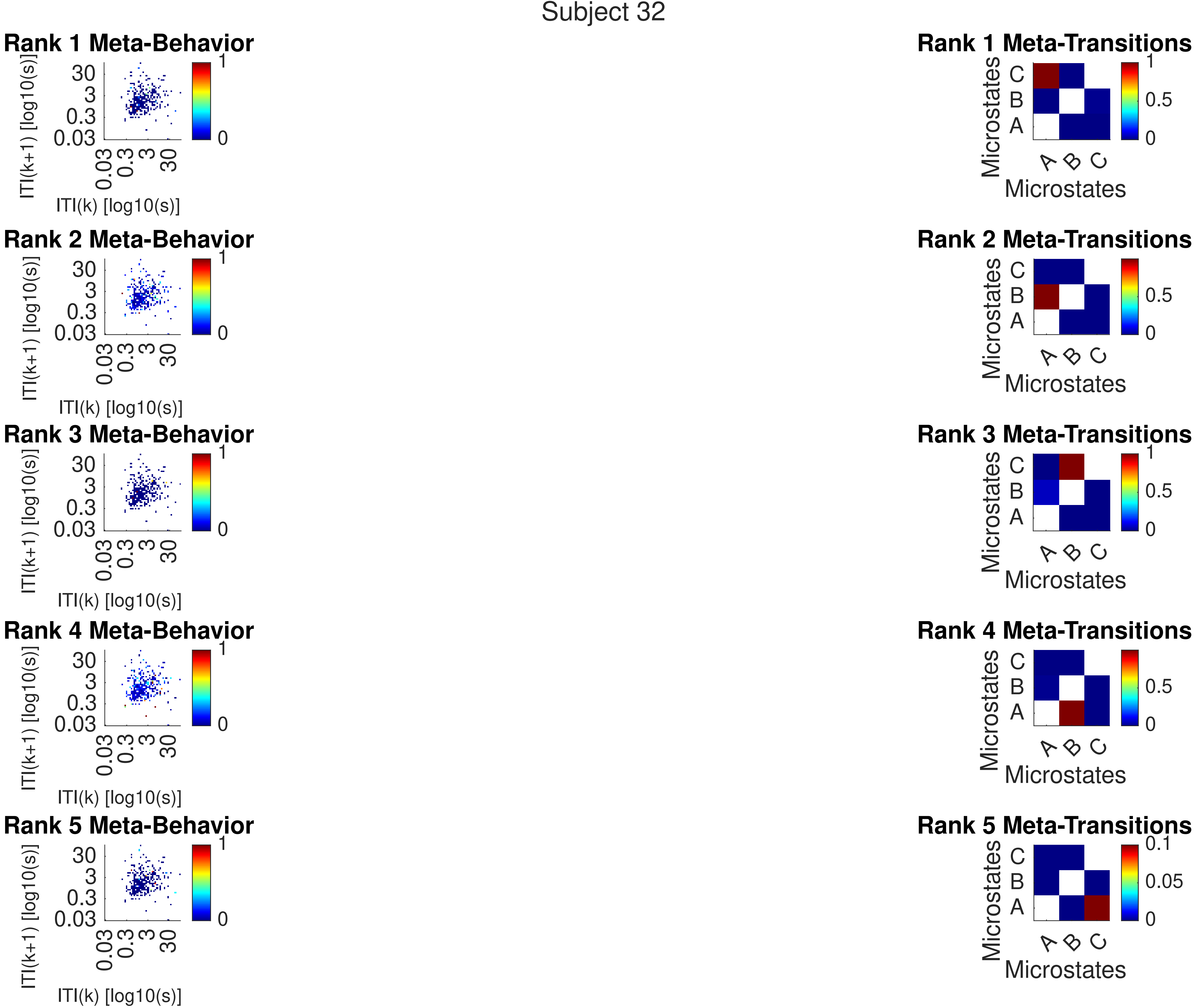
Subject 32
Rank 1 Meta-Behavior
Rank 2 Meta-Behavior
Rank 3 Meta-Behavior
Rank 4 Meta-Behavior
Rank 5 Meta-Behavior
ITI(k+1) [log10(s)]
ITI(k) [log10(s)]
30
3
0.3
0.03
1
0
Rank 1 Meta-Transitions
Rank 2 Meta-Transitions
Rank 3 Meta-Transitions
Rank 4 Meta-Transitions
Rank 5 Meta-Transitions
Microstates
A
B
C
0.5
0.1
0.05

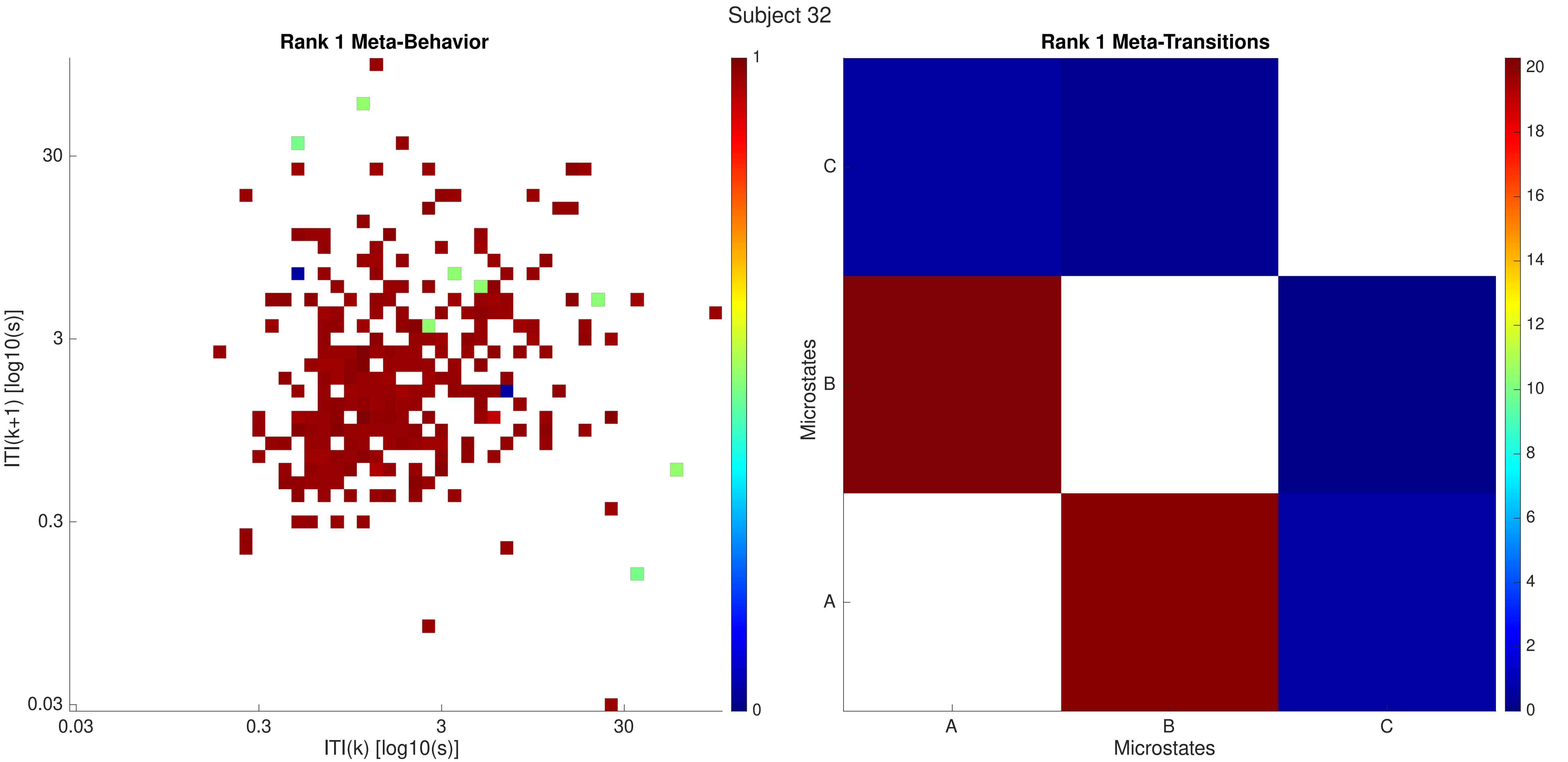

Subject 32
Rank 1 Meta-Behavior
ITI(k+1) [log10(s)]
ITI(k) [log10(s)]
0.03
0.3
3
30
1
0
Rank 1 Meta-Transitions
Microstates
A
B
C
20
18
16
14
12
10
8
6
4
2
0

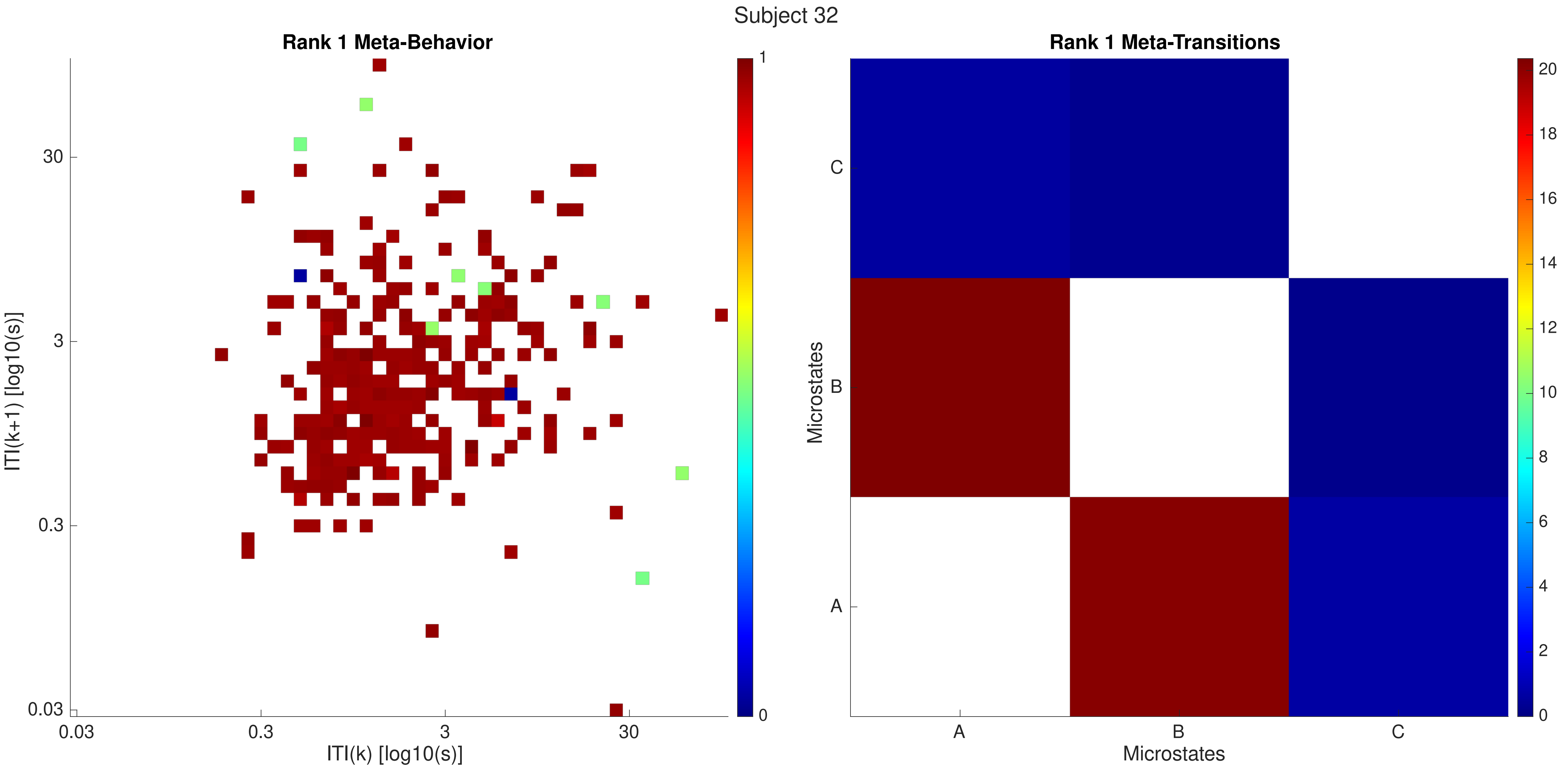
Subject 32
Rank 1 Meta-Behavior
ITI(k+1) [log10(s)]
ITI(k) [log10(s)]
30
3
0.3
0.03
1
0
Rank 1 Meta-Transitions
Microstates
A
B
C
20
18
16
14
12
10
8
6
4
2
0

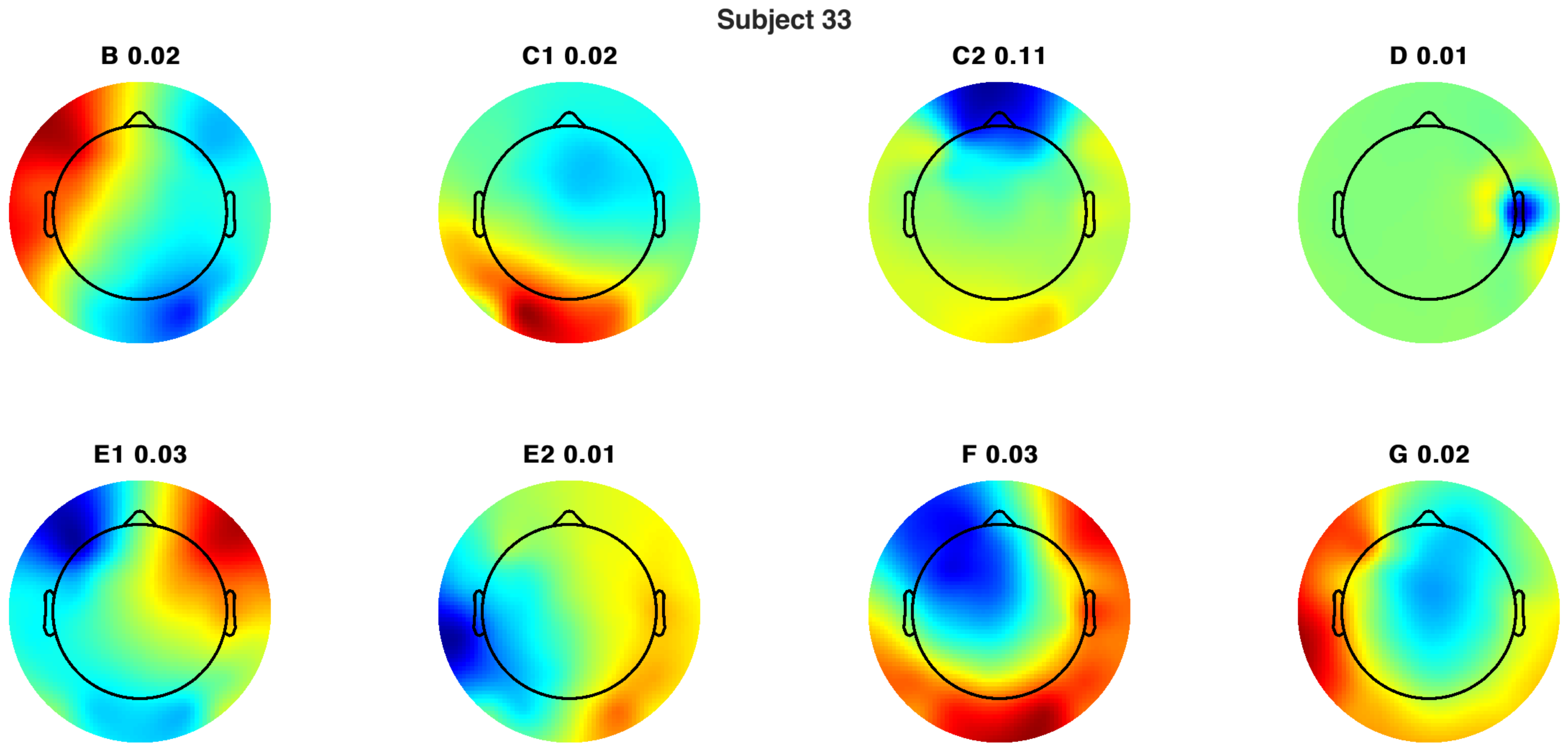
Subject 33
B 0.02
C1 0.02
C2 0.11
D 0.01
E1 0.03
E2 0.01
F 0.03
G 0.02

Subject 33

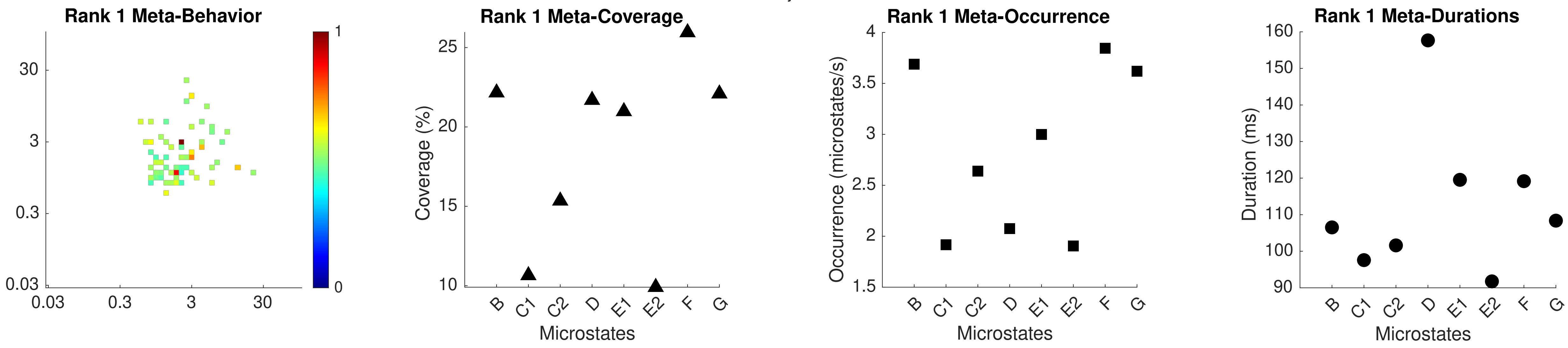

Rank 1 Meta-Behavior
30
3
0.3
0.03
0.03
0.3
3
30
1
0
Rank 1 Meta-Coverage
Coverage (%)
25
20
15
10
B
C1
C2
D
E1
E2
F
G
Microstates
Rank 1 Meta-Occurrence
Occurrence (microstates/s)
4
3.5
3
2.5
2
1.5
B
C1
C2
D
E1
E2
F
G
Microstates
Rank 1 Meta-Durations
Duration (ms)
160
150
140
130
120
110
100
90
B
C1
C2
D
E1
E2
F
G
Microstates

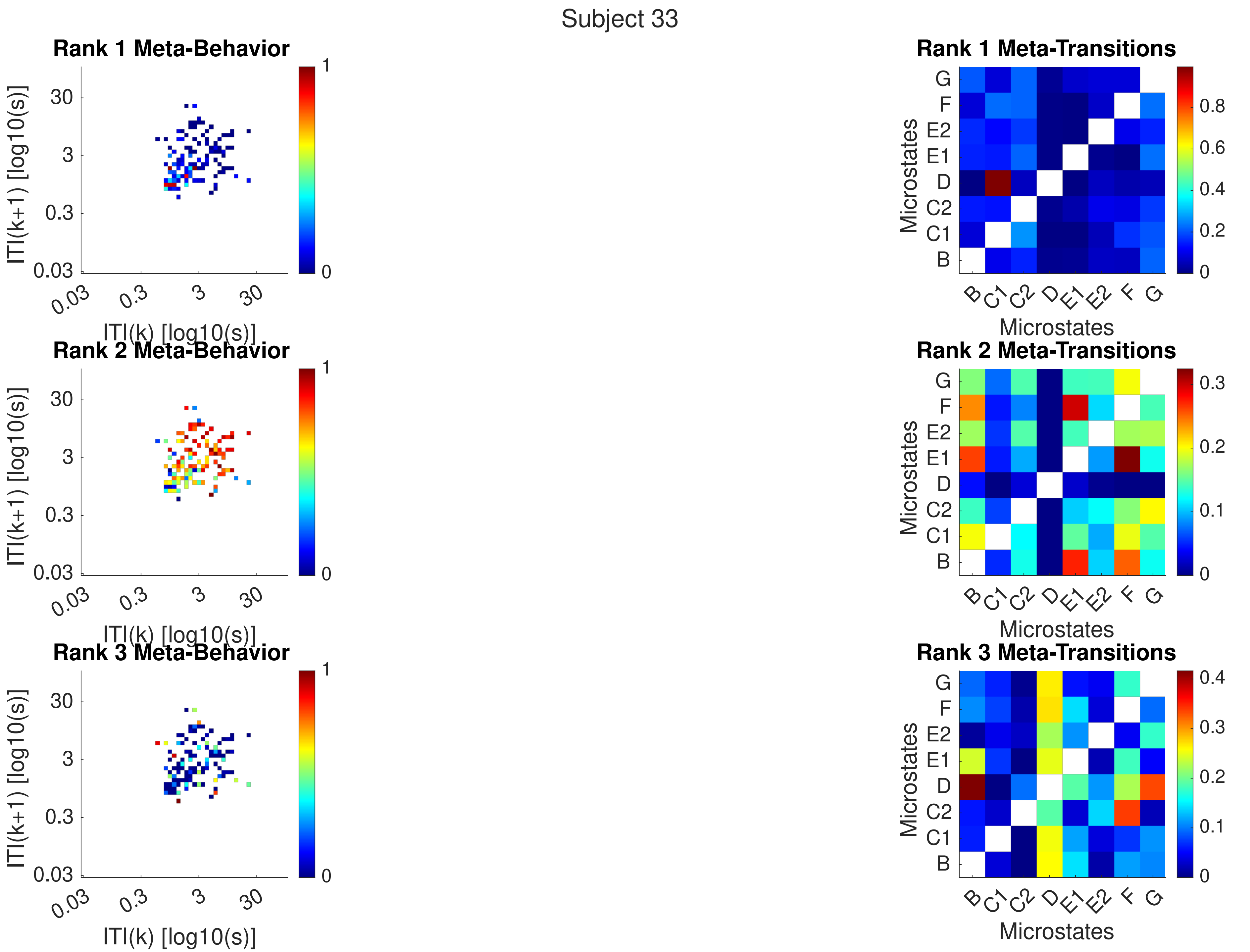
Subject 33
Rank 1 Meta-Behavior
Rank 2 Meta-Behavior
Rank 3 Meta-Behavior
ITI(k+1) [log10(s)]
ITI(k) [log10(s)]
0.03
0.3
3
30
0
1
Rank 1 Meta-Transitions
Rank 2 Meta-Transitions
Rank 3 Meta-Transitions
Microstates
B
C1
C2
D
E1
E2
F
G
0.2
0.4
0.6
0.8
0.1
0.3

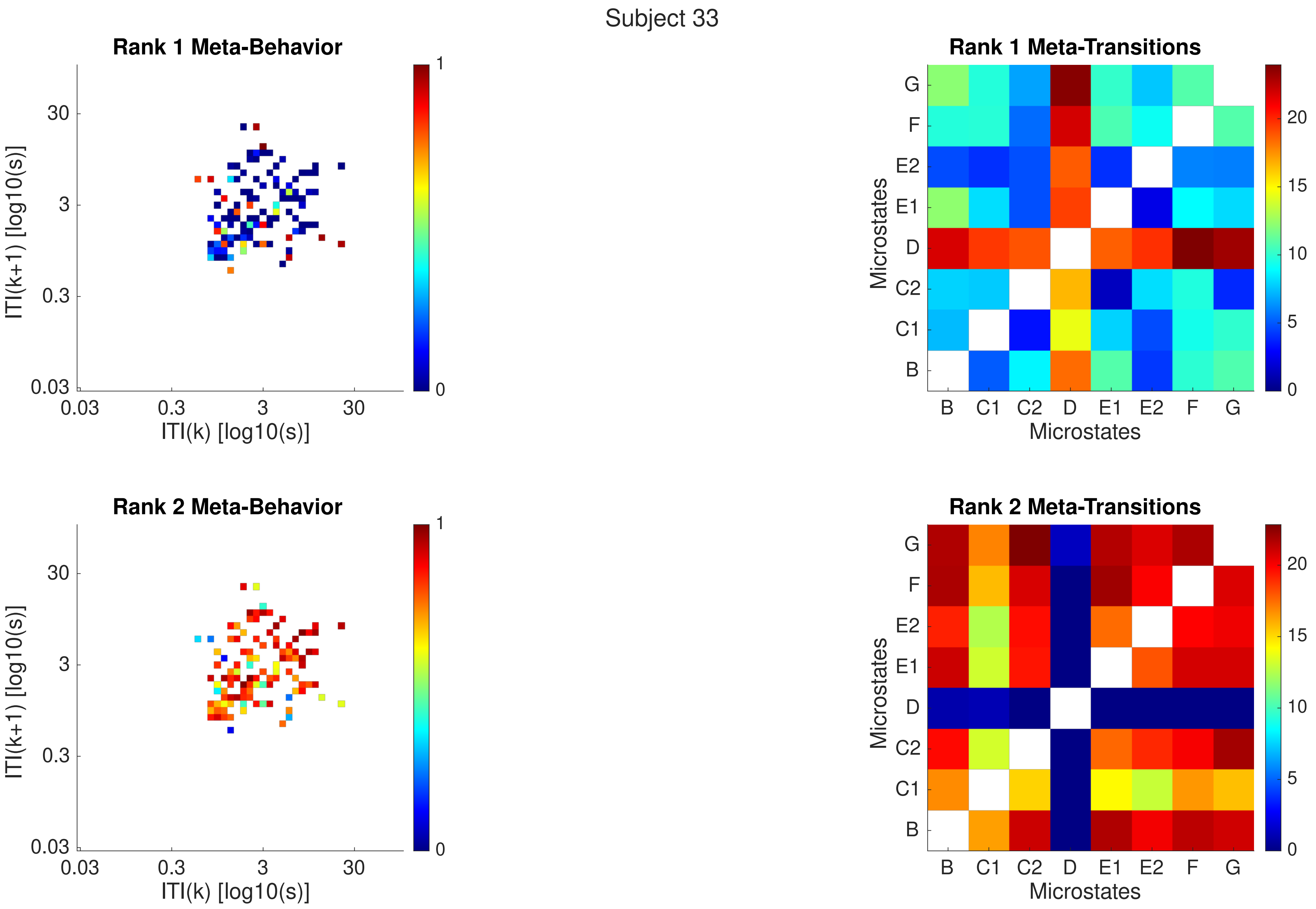
Subject 33
Rank 1 Meta-Behavior
ITI(k+1) [log10(s)]
ITI(k) [log10(s)]
0.03
0.3
3
30
0
1
Rank 1 Meta-Transitions
Microstates
B
C1
C2
D
E1
E2
F
G
0
5
10
15
20
Rank 2 Meta-Behavior
ITI(k+1) [log10(s)]
ITI(k) [log10(s)]
Rank 2 Meta-Transitions
Microstates

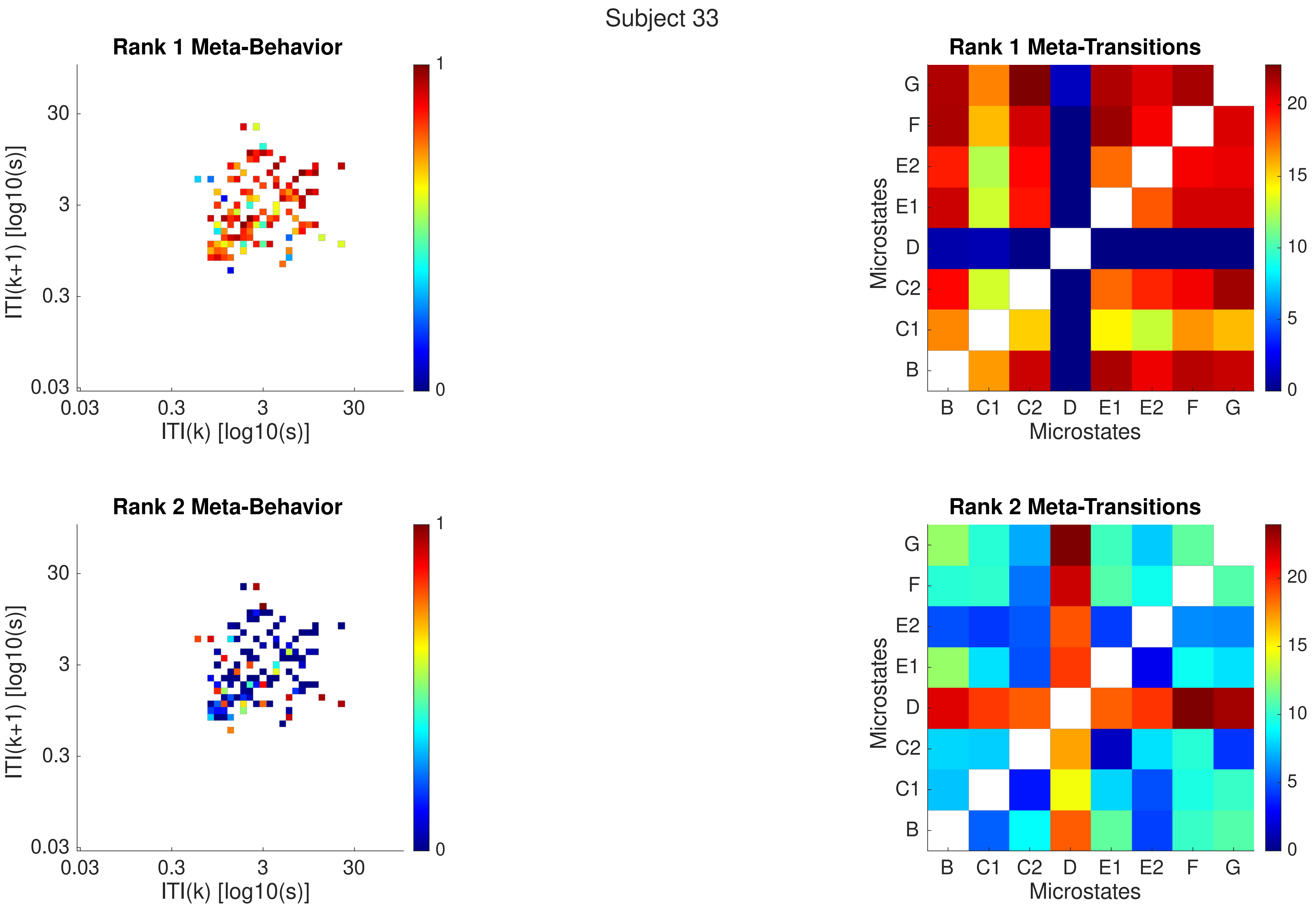
Subject 33
Rank 1 Meta-Behavior
ITI(k+1) [log10(s)]
ITI(k) [log10(s)]
0.03
0.3
3
30
0
1
Rank 1 Meta-Transitions
Microstates
B
C1
C2
D
E1
E2
F
G
0
5
10
15
20
Rank 2 Meta-Behavior
ITI(k+1) [log10(s)]
ITI(k) [log10(s)]
Rank 2 Meta-Transitions
Microstates

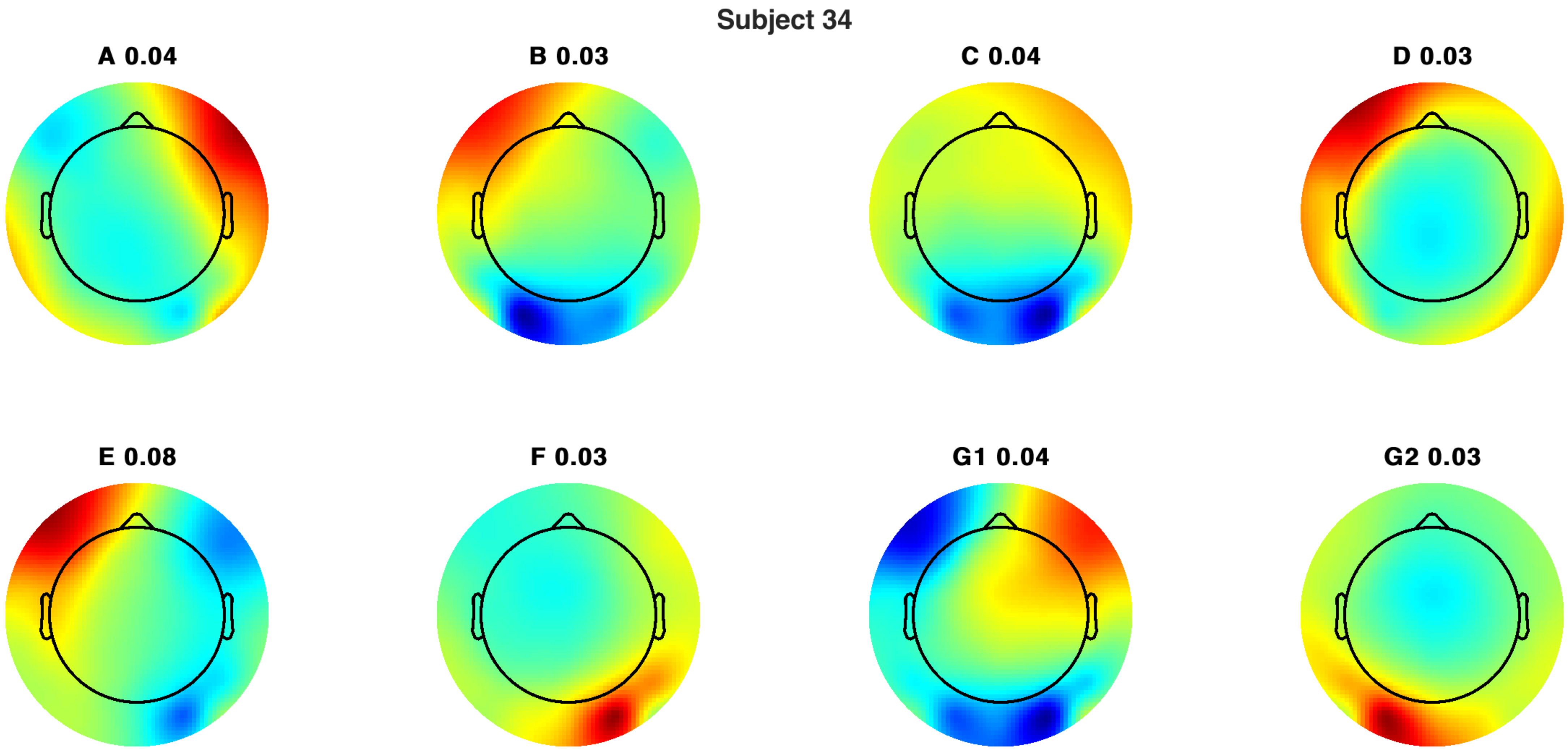
Subject 34
A 0.04
B 0.03
C 0.04
D 0.03
E 0.08
F 0.03
G1 0.04
G2 0.03

Subject 34

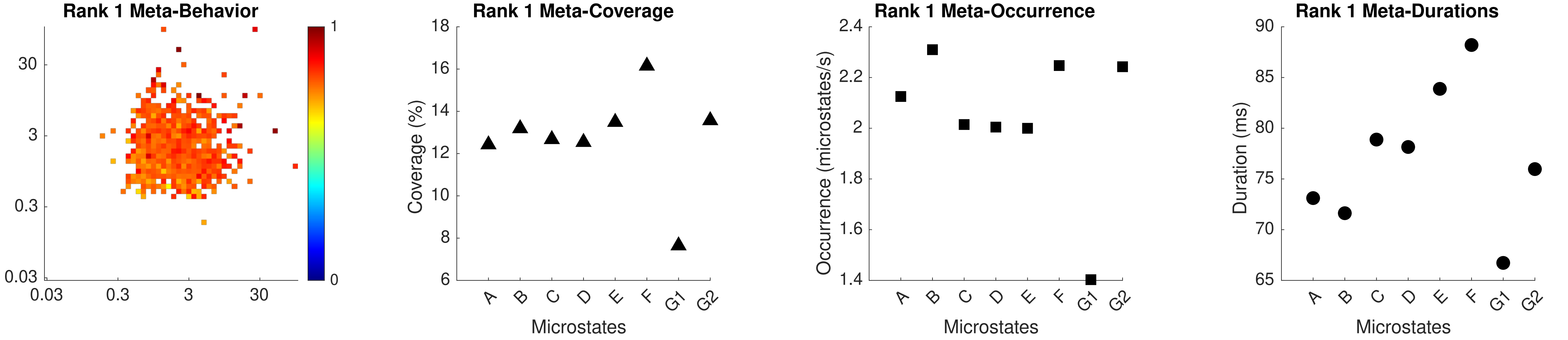
Rank 1 Meta-Behavior
30
3
0.3
0.03
0.03
0.3
3
30
1
0
Rank 1 Meta-Coverage
Coverage (%)
18
16
14
12
10
8
6
A
B
C
D
E
F
G1
G2
Microstates
Rank 1 Meta-Occurrence
Occurrence (microstates/s)
2.4
2.2
2
1.8
1.6
1.4
A
B
C
D
E
F
G1
G2
Microstates
Rank 1 Meta-Durations
Duration (ms)
90
85
80
75
70
65
A
B
C
D
E
F
G1
G2
Microstates

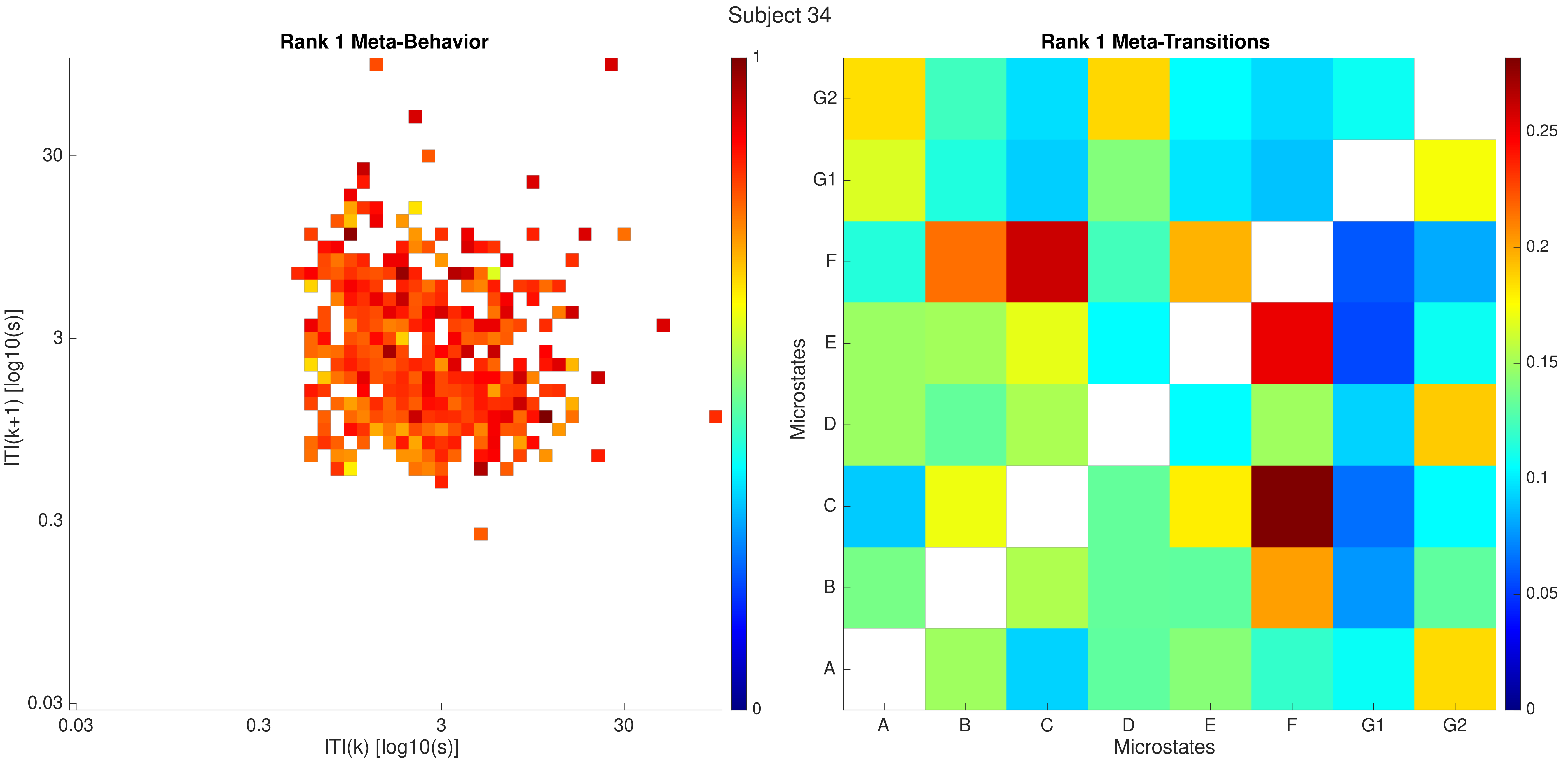

Subject 34
Rank 1 Meta-Behavior
ITI(k+1) [log10(s)]
30
3
0.3
0.03
ITI(k) [log10(s)]
1
0
Rank 1 Meta-Transitions
Microstates
G2
G1
F
E
D
C
B
A
0.25
0.2
0.15
0.1
0.05

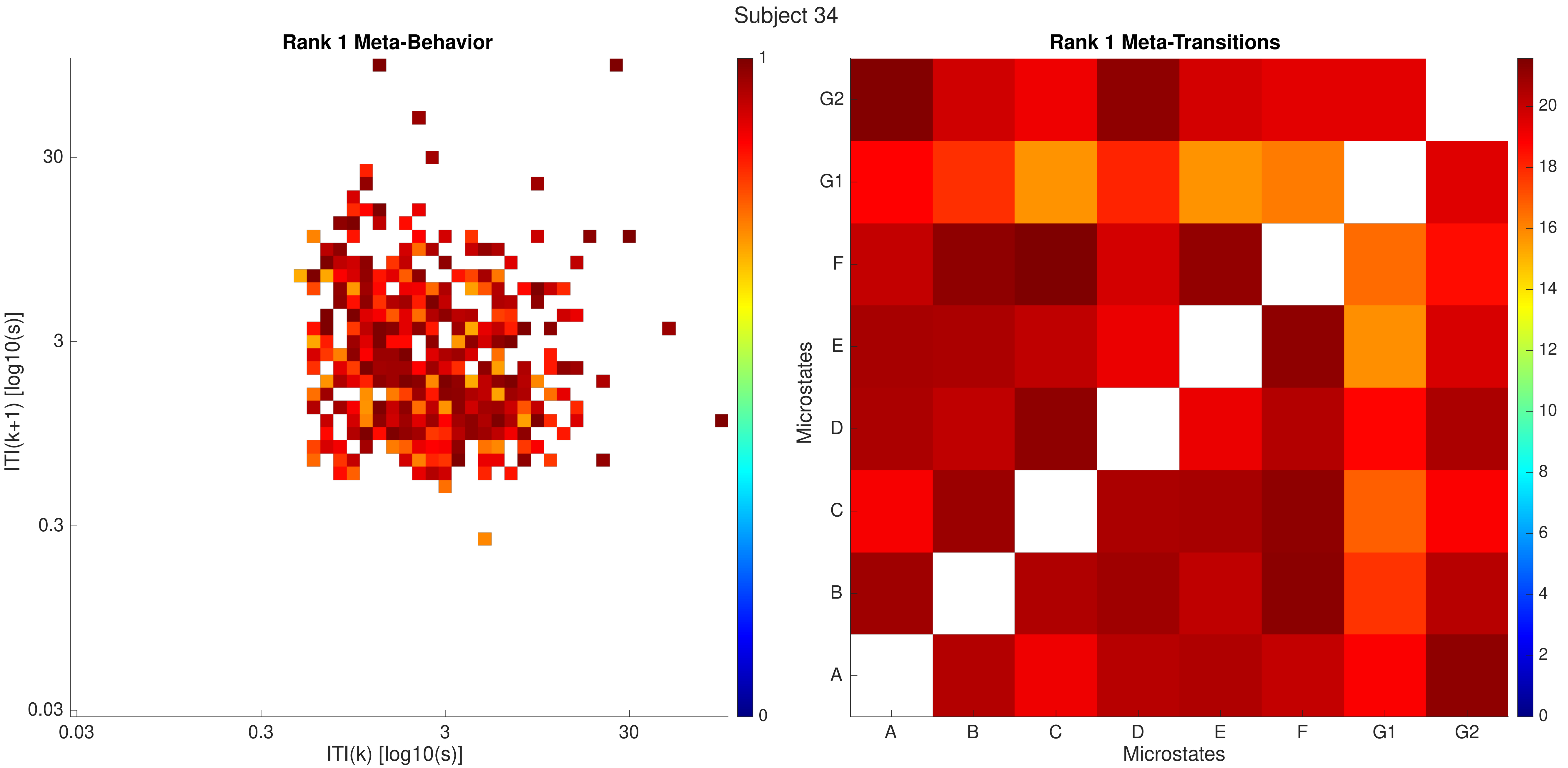
Subject 34
Rank 1 Meta-Behavior
ITI(k+1) [log10(s)]
30
3
0.3
0.03
0.03
0.3
3
30
ITI(k) [log10(s)]
1
0
Rank 1 Meta-Transitions
Microstates
G2
G1
F
E
D
C
B
A
A
B
C
D
E
F
G1
G2
Microstates
20
18
16
14
12
10
8
6
4
2
0

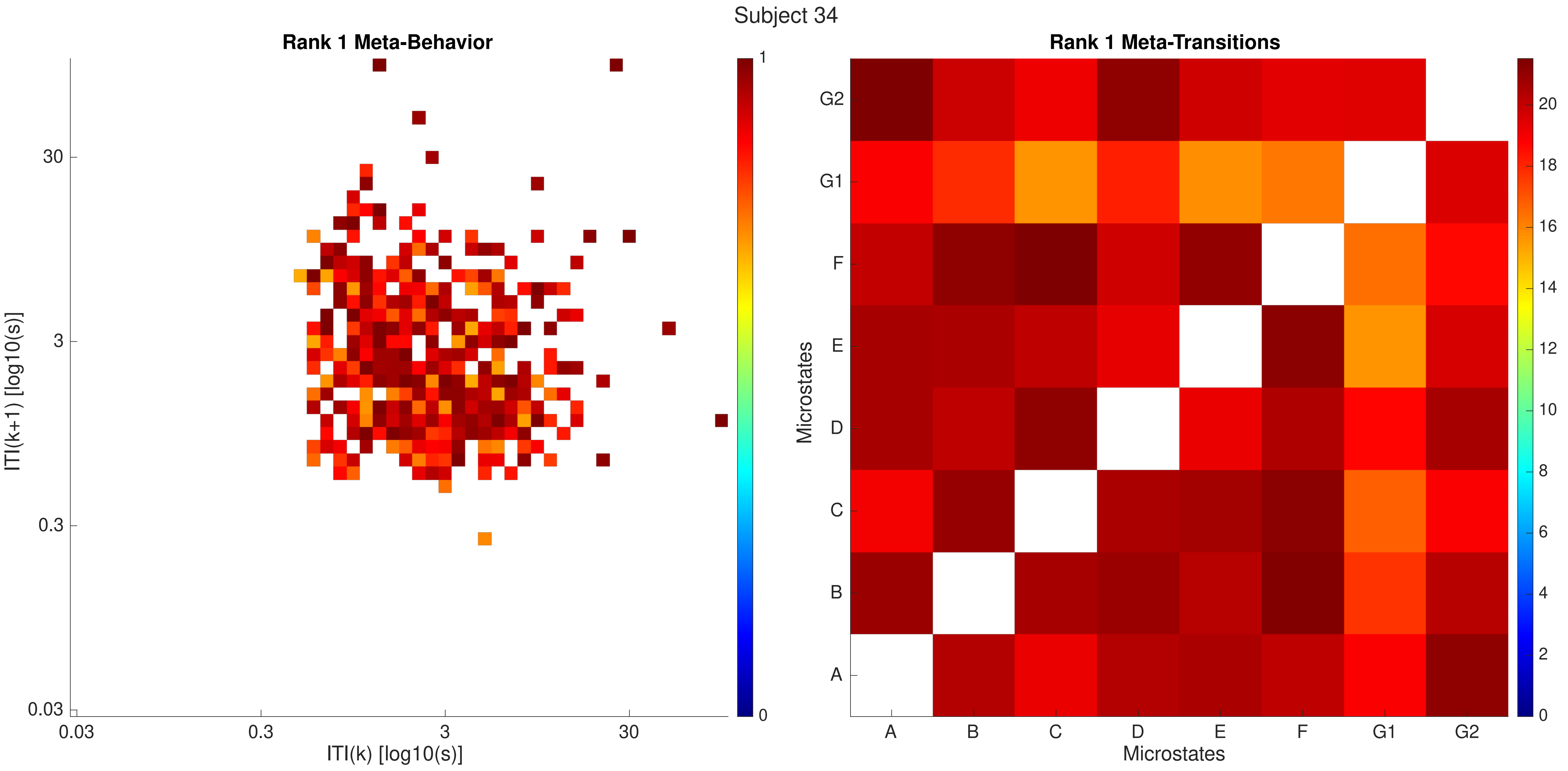

Subject 34
Rank 1 Meta-Behavior
ITI(k+1) [log10(s)]
30
3
0.3
0.03
0.03
0.3
3
30
ITI(k) [log10(s)]
1
0
Rank 1 Meta-Transitions
Microstates
G2
G1
F
E
D
C
B
A
A
B
C
D
E
F
G1
G2
Microstates
20
18
16
14
12
10
8
6
4
2
0

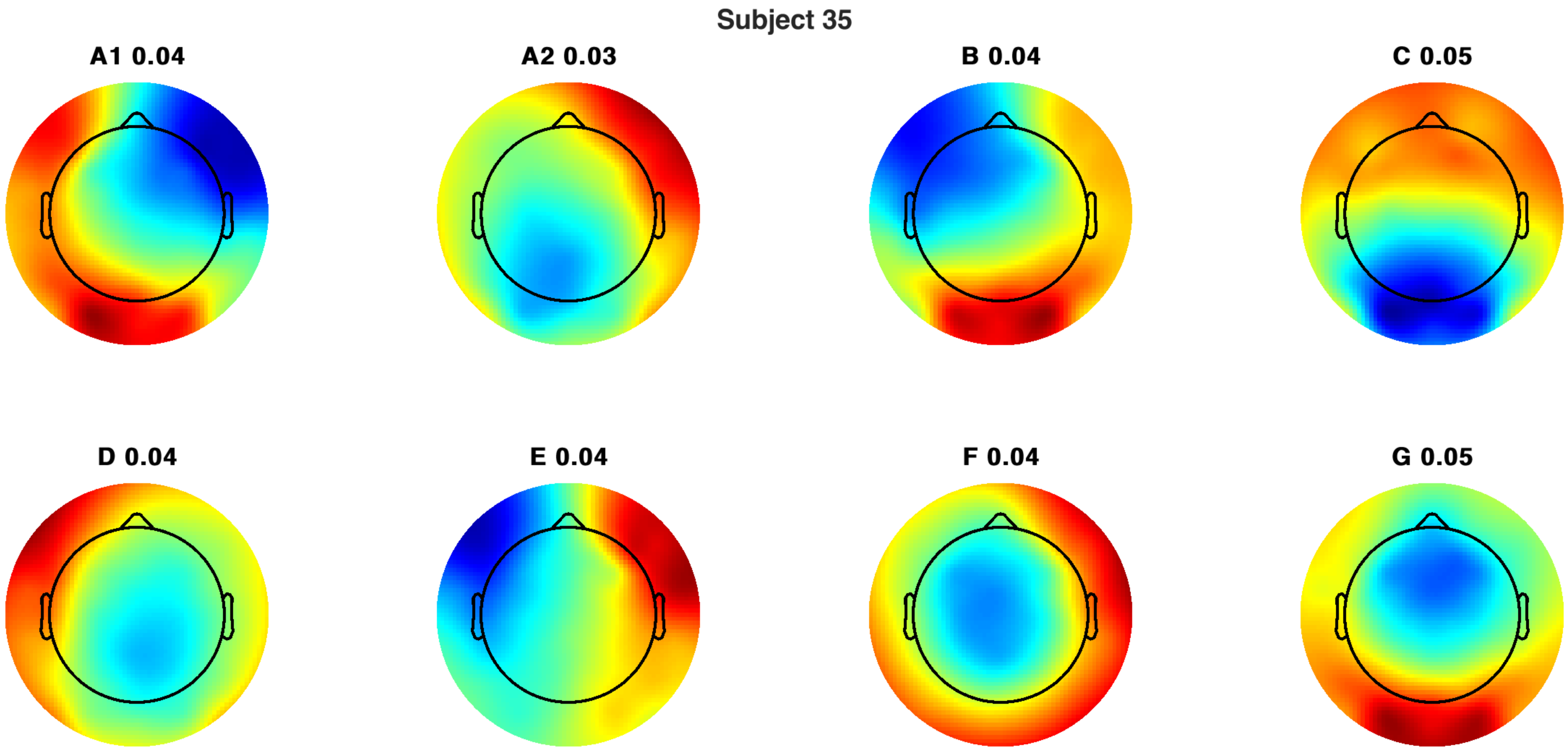
Subject 35
A1 0.04
A2 0.03
B 0.04
C 0.05
D 0.04
E 0.04
F 0.04
G 0.05

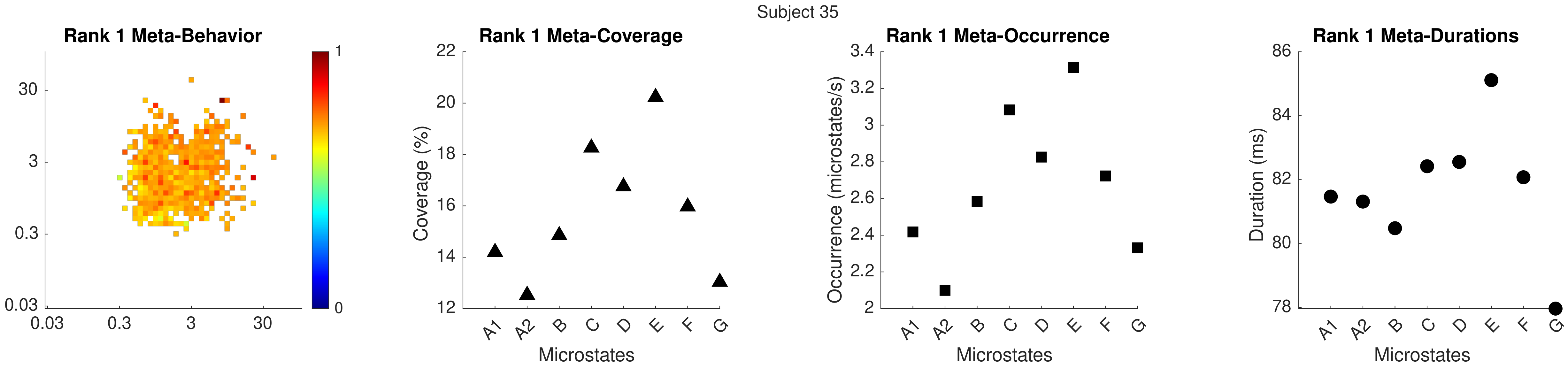
Subject 35
Rank 1 Meta-Behavior
Rank 1 Meta-Coverage
Coverage (%)
Microstates
Rank 1 Meta-Occurrence
Occurrence (microstates/s)
Microstates
Rank 1 Meta-Durations
Duration (ms)
Microstates

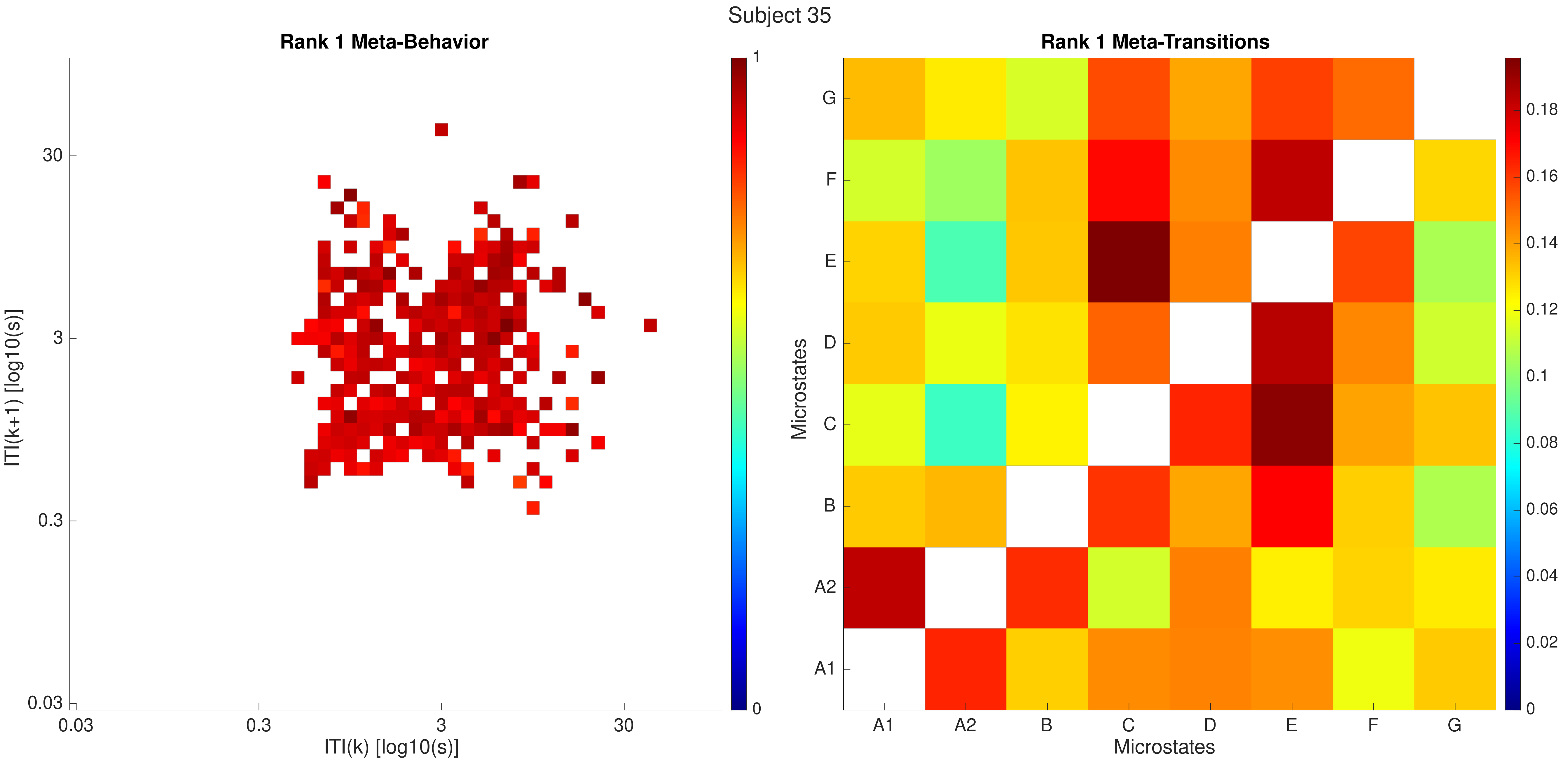

Subject 35
Rank 1 Meta-Behavior
ITI(k+1) [log10(s)]
ITI(k) [log10(s)]
0.03
0.3
3
30
1
0
Rank 1 Meta-Transitions
Microstates
A1
A2
B
C
D
E
F
G
0.18
0.16
0.14
0.12
0.1
0.08
0.06
0.04
0.02
0

Subject 35

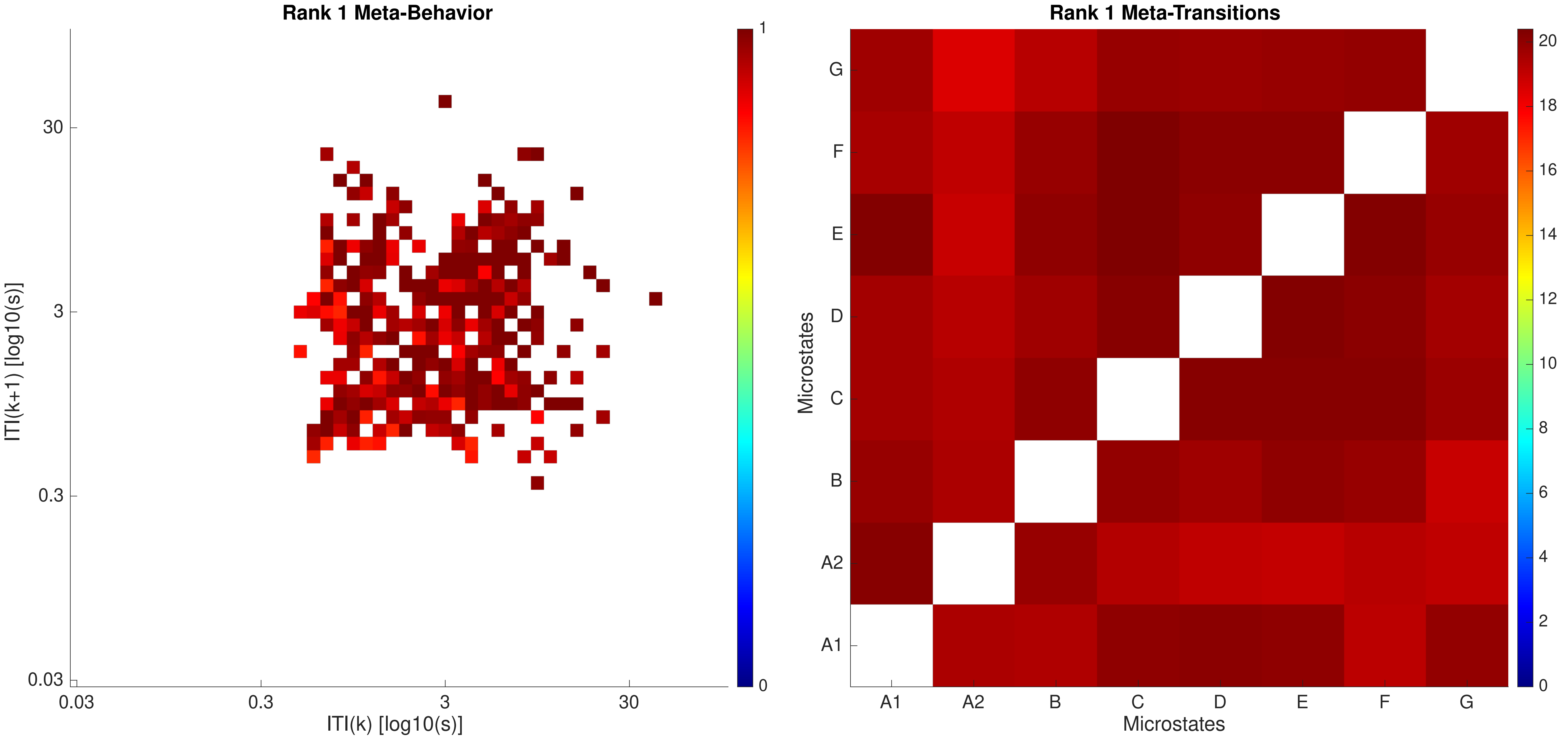

Rank 1 Meta-Behavior
ITI(k+1) [log10(s)]
ITI(k) [log10(s)]
0.03
0.3
3
30
0
1
Rank 1 Meta-Transitions
Microstates
A1
A2
B
C
D
E
F
G
0
2
4
6
8
10
12
14
16
18
20

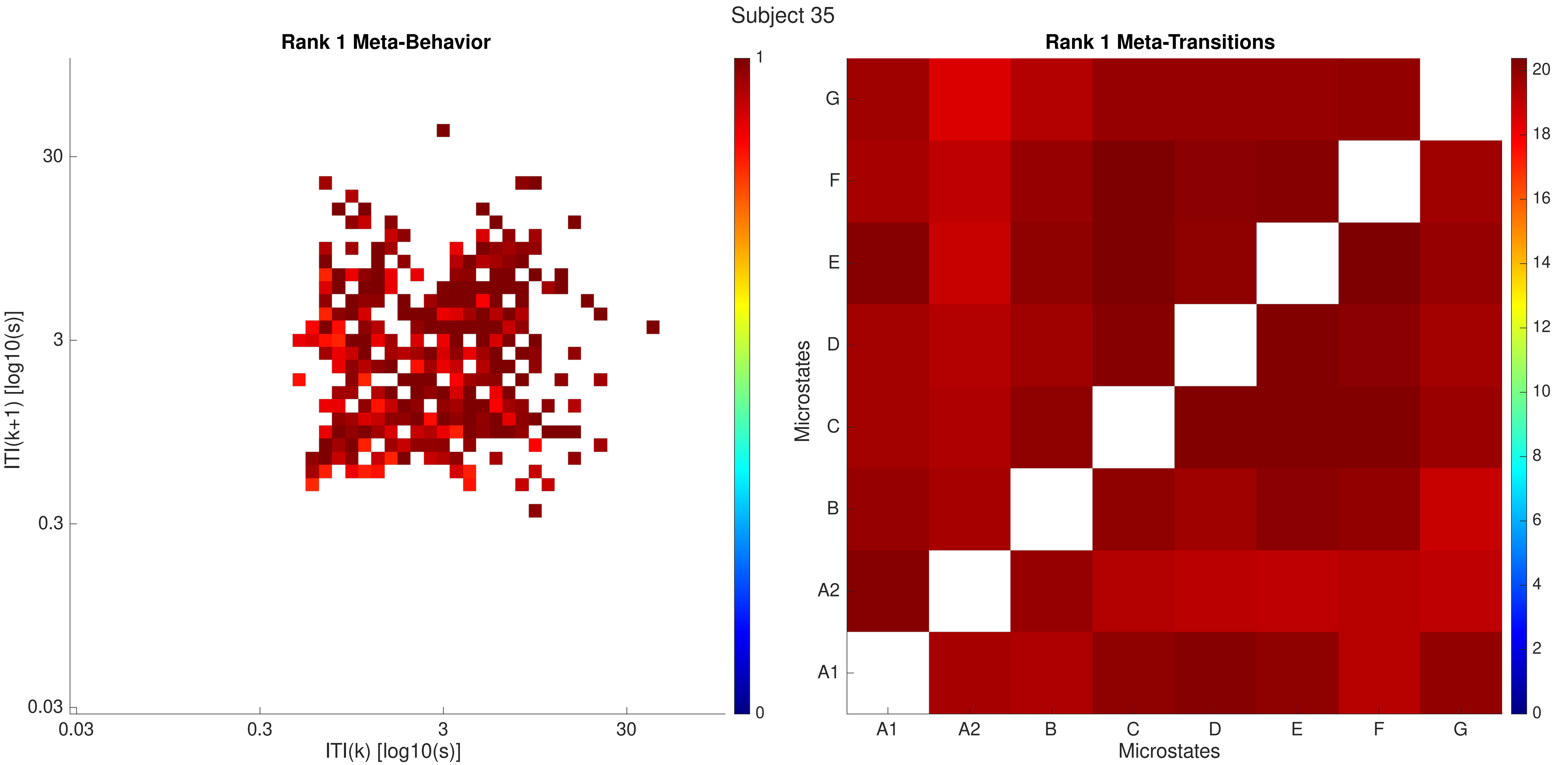

Subject 35
Rank 1 Meta-Behavior
ITI(k+1) [log10(s)]
ITI(k) [log10(s)]
0.03
0.3
3
30
0
1
Rank 1 Meta-Transitions
Microstates
A1
A2
B
C
D
E
F
G
0
2
4
6
8
10
12
14
16
18
20

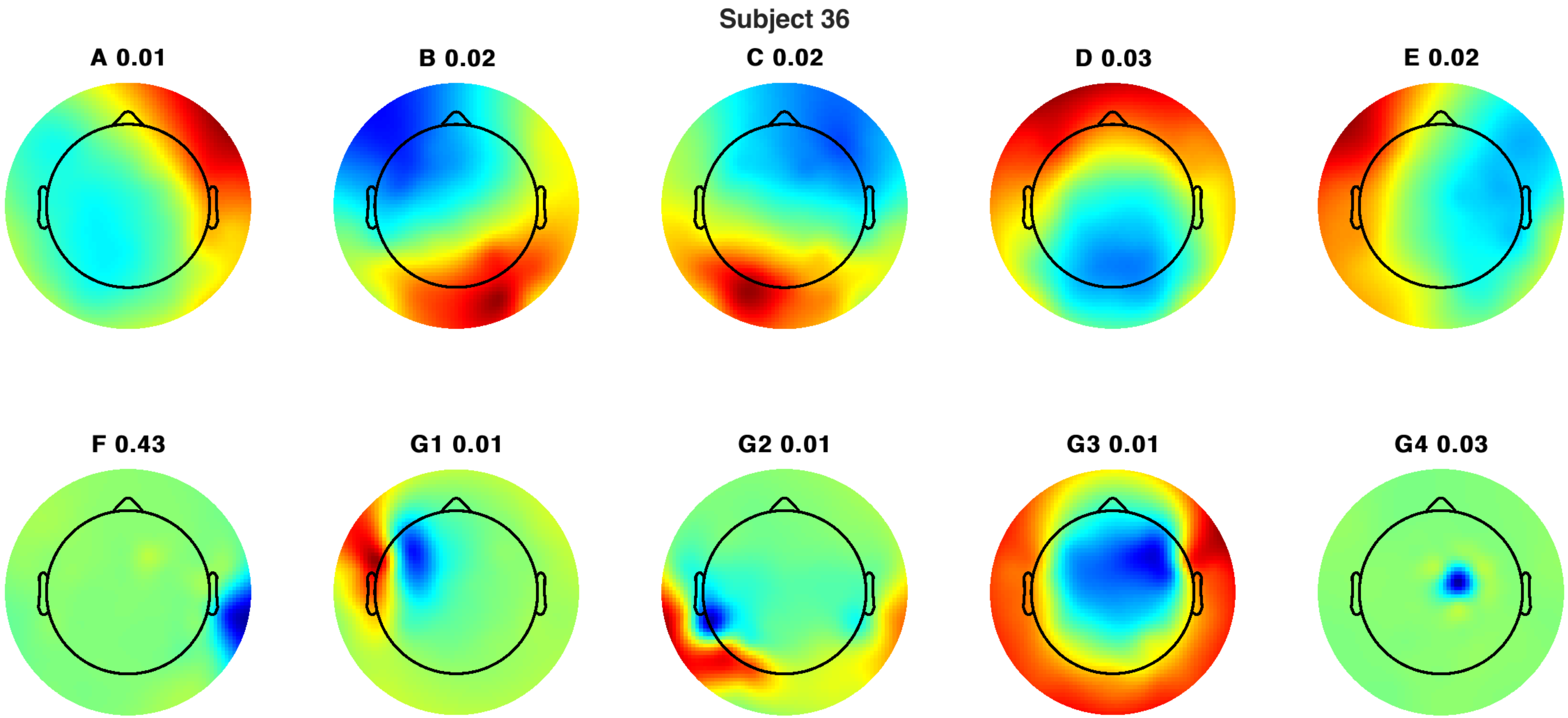
Subject 36
A 0.01
B 0.02
C 0.02
D 0.03
E 0.02
F 0.43
G1 0.01
G2 0.01
G3 0.01
G4 0.03

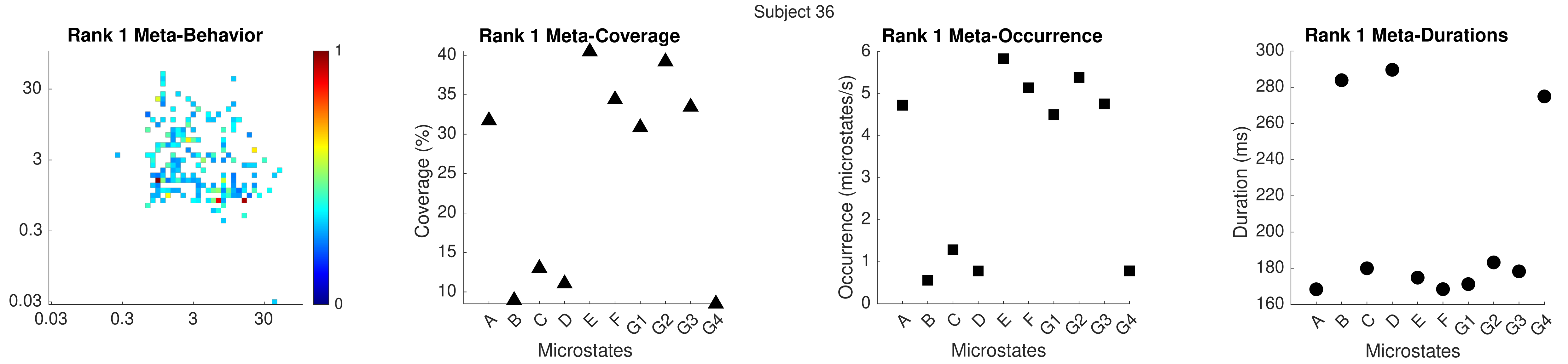
Subject 36
Rank 1 Meta-Behavior
Rank 1 Meta-Coverage
Coverage (%)
Microstates
Rank 1 Meta-Occurrence
Occurrence (microstates/s)
Microstates
Rank 1 Meta-Durations
Duration (ms)
Microstates

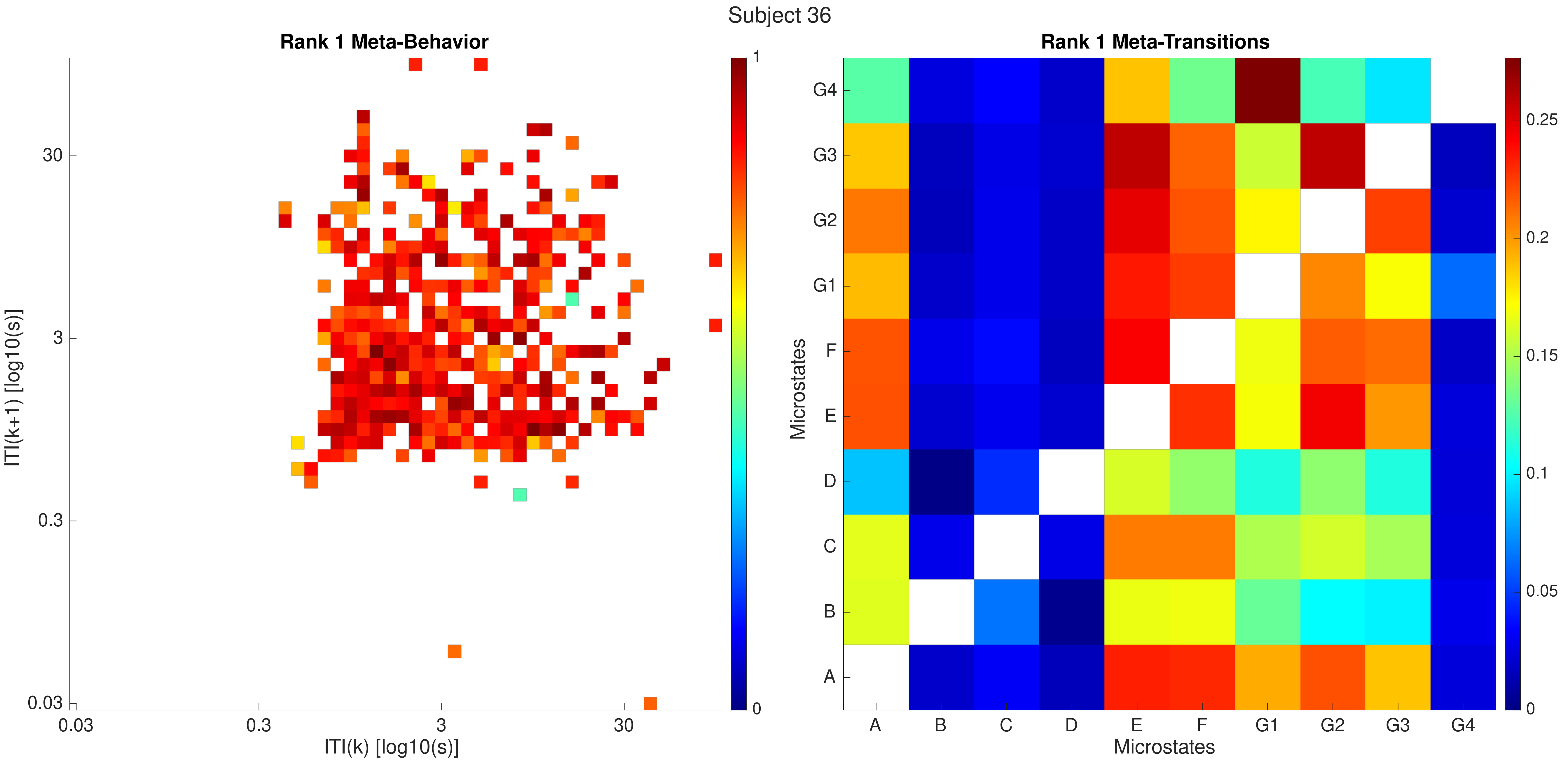
Subject 36
Rank 1 Meta-Behavior
ITI(k+1) [log10(s)]
ITI(k) [log10(s)]
0.03
0.3
3
30
1
0
Rank 1 Meta-Transitions
Microstates
A
B
C
D
E
F
G1
G2
G3
G4
0
0.05
0.1
0.15
0.2
0.25

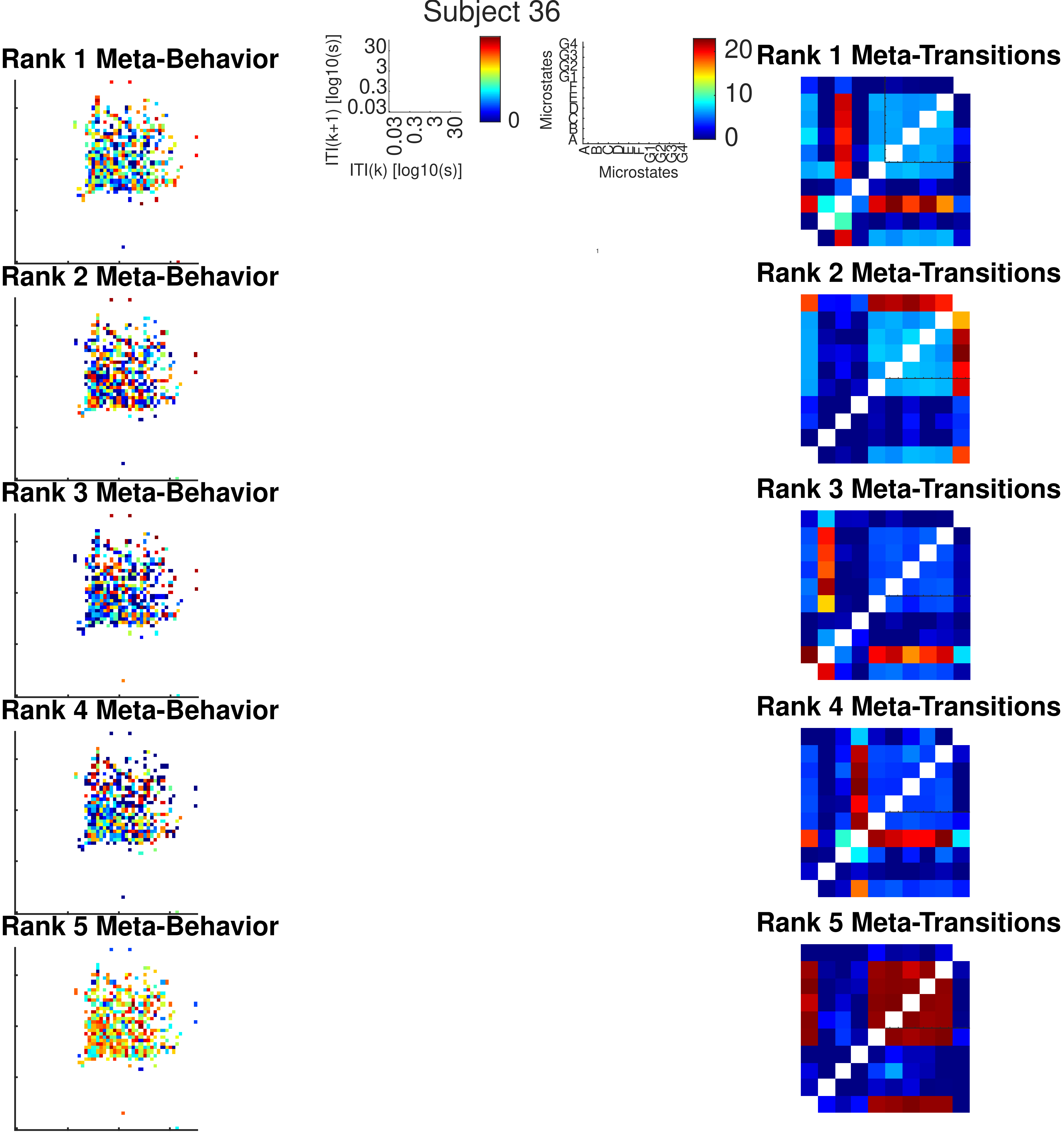

Subject 36
Rank 1 Meta-Behavior
Rank 2 Meta-Behavior
Rank 3 Meta-Behavior
Rank 4 Meta-Behavior
Rank 5 Meta-Behavior
ITI(k+1) [log10(s)]
30
3
0.3
0.03
ITI(k) [log10(s)]
0
Microstates
A B C D E F G1 G2 G3 G4
20
10
0
Rank 1 Meta-Transitions
Rank 2 Meta-Transitions
Rank 3 Meta-Transitions
Rank 4 Meta-Transitions
Rank 5 Meta-Transitions

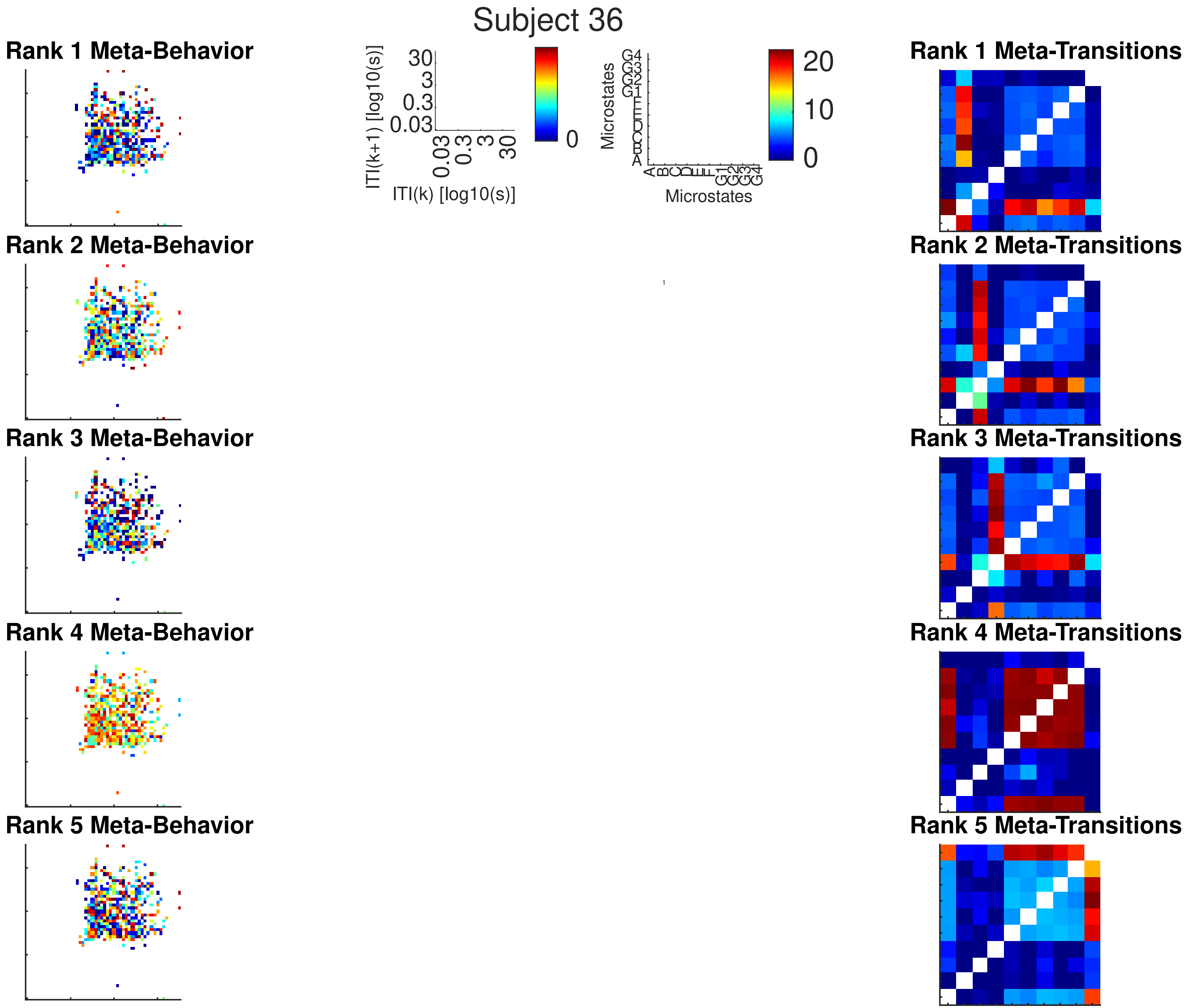
Subject 36
Rank 1 Meta-Behavior
Rank 2 Meta-Behavior
Rank 3 Meta-Behavior
Rank 4 Meta-Behavior
Rank 5 Meta-Behavior
ITI(k+1) [log10(s)]
30
3
0.3
0.03
ITI(k) [log10(s)]
0
Microstates
A
B
C
D
E
F
G1
G2
G3
G4
20
10
0
Rank 1 Meta-Transitions
Rank 2 Meta-Transitions
Rank 3 Meta-Transitions
Rank 4 Meta-Transitions
Rank 5 Meta-Transitions

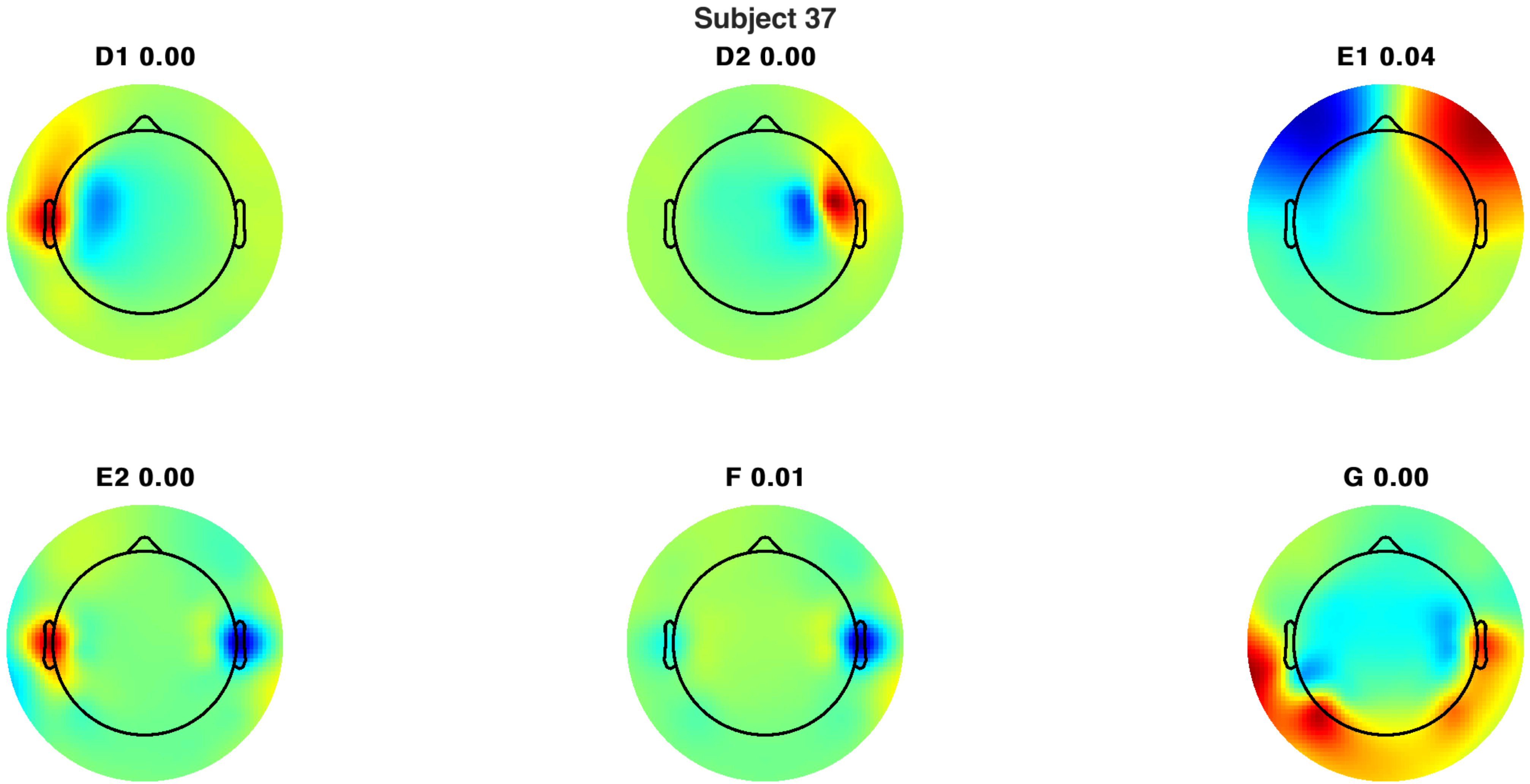
Subject 37
D1 0.00
D2 0.00
E1 0.04
E2 0.00
F 0.01
G 0.00

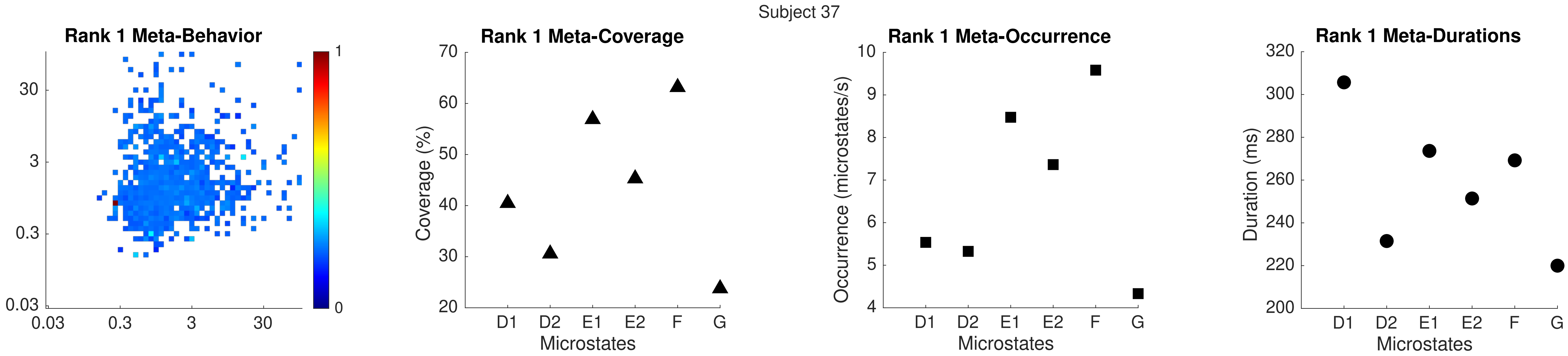
Subject 37
Rank 1 Meta-Behavior
30
3
0.3
0.03
0.03
0.3
3
30
1
0
Rank 1 Meta-Coverage
Coverage (%)
70
60
50
40
30
20
D1
D2
E1
E2
F
G
Microstates
Rank 1 Meta-Occurrence
Occurrence (microstates/s)
10
9
8
7
6
5
4
D1
D2
E1
E2
F
G
Microstates
Rank 1 Meta-Durations
Duration (ms)
320
300
280
260
240
220
200
D1
D2
E1
E2
F
G
Microstates

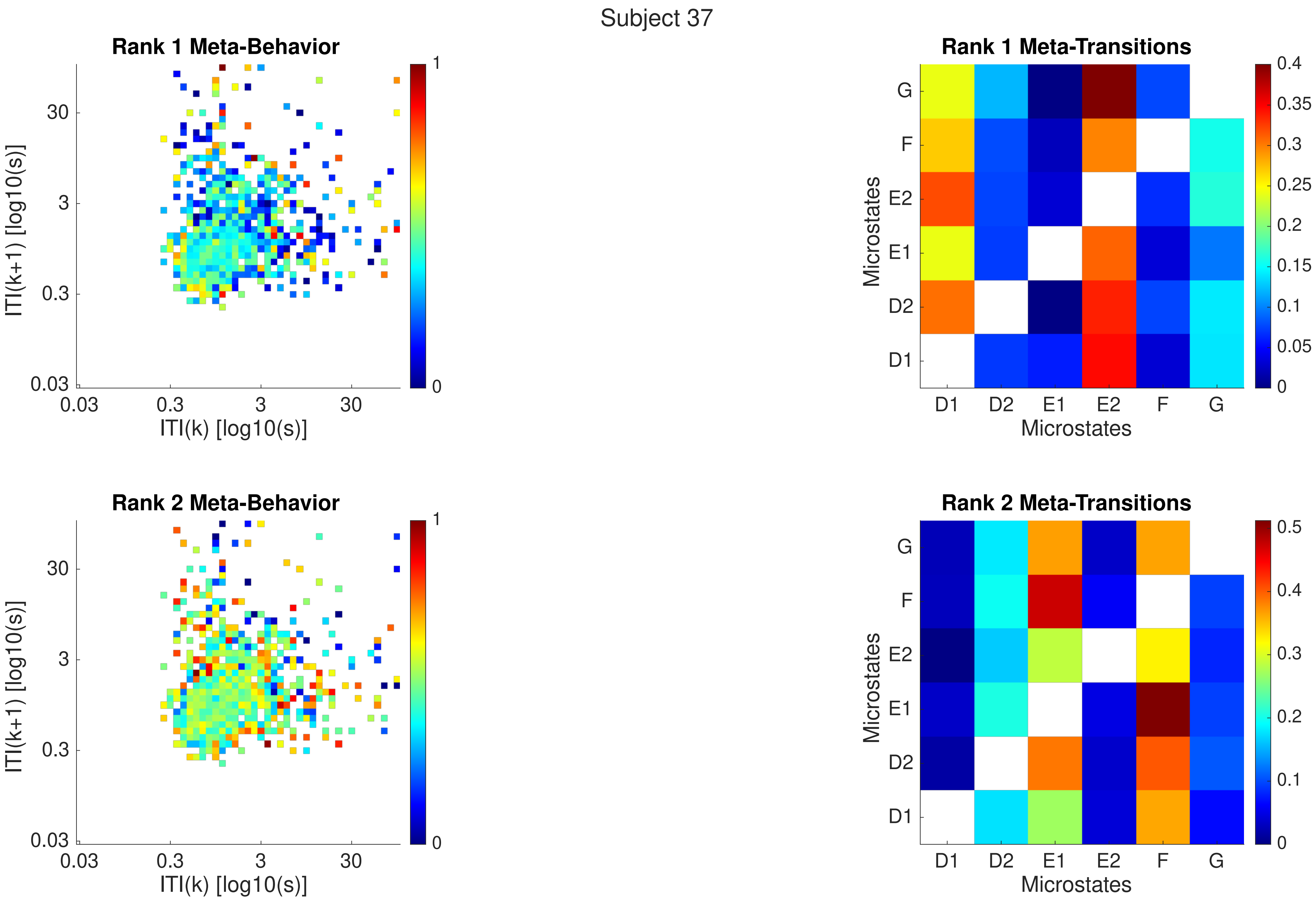
Subject 37
Rank 1 Meta-Behavior
Rank 1 Meta-Transitions
Rank 2 Meta-Behavior
Rank 2 Meta-Transitions
ITI(k+1) [log10(s)]
ITI(k) [log10(s)]
0.03
0.3
3
30
0
1
Microstates
D1
D2
E1
E2
F
G
0
0.05
0.1
0.15
0.2
0.25
0.3
0.35
0.4
0.5

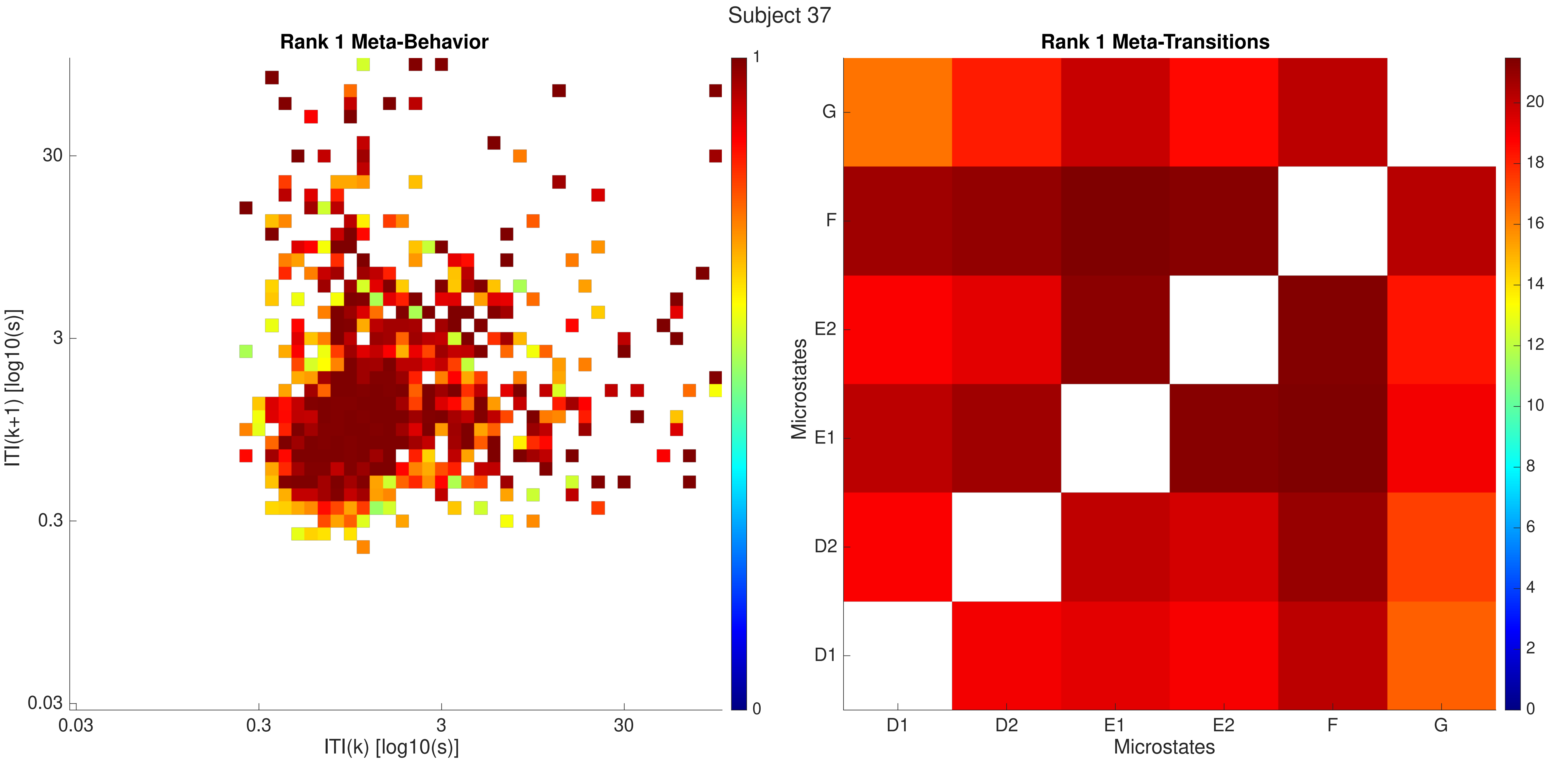

Subject 37
Rank 1 Meta-Behavior
Rank 1 Meta-Transitions
ITI(k+1) [log10(s)]
ITI(k) [log10(s)]
0.03
0.3
3
30
0
1
Microstates
D1
D2
E1
E2
F
G
0
2
4
6
8
10
12
14
16
18
20

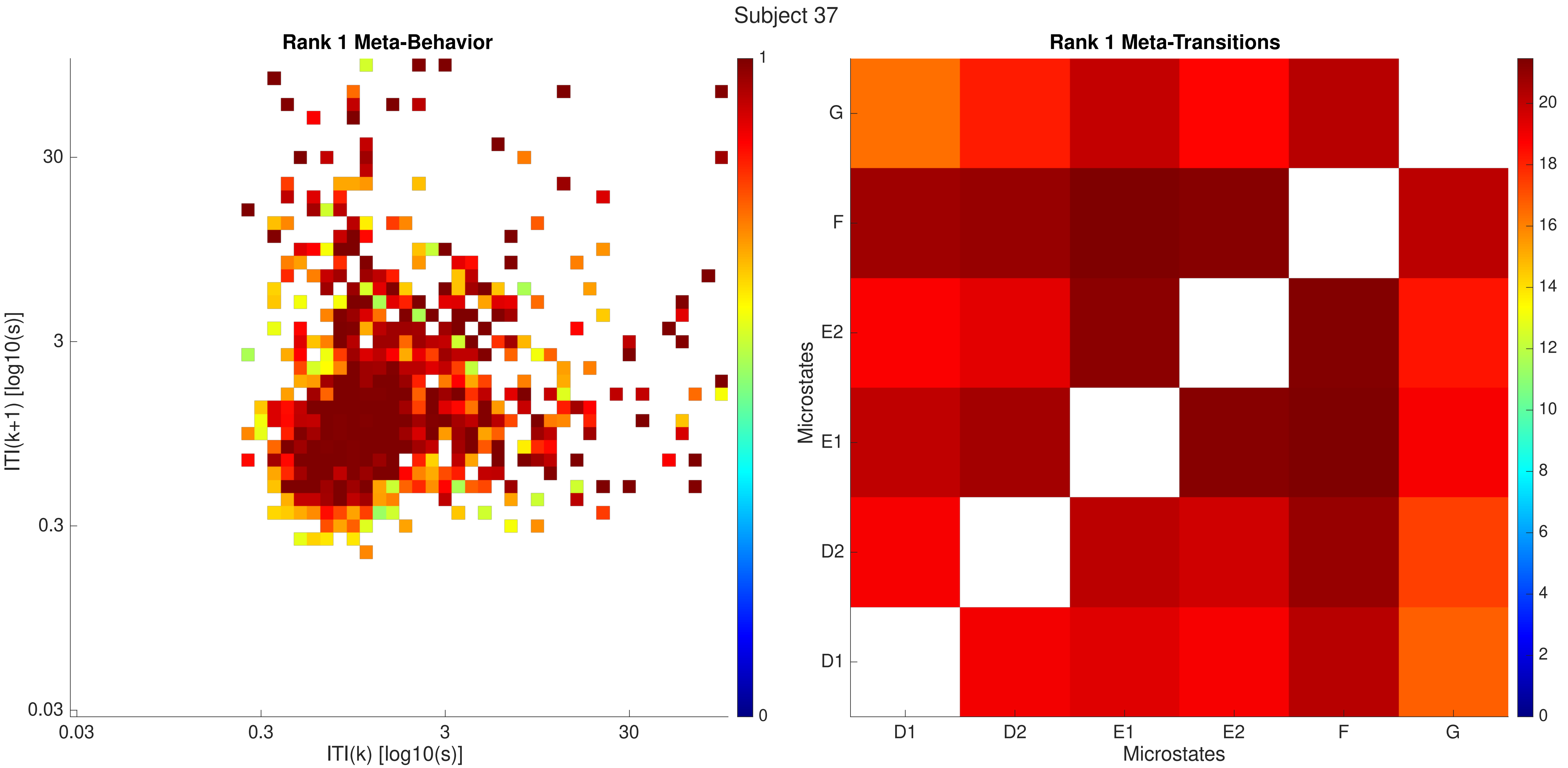

Subject 37
Rank 1 Meta-Behavior
ITI(k+1) [log10(s)]
ITI(k) [log10(s)]
0.03
0.3
3
30
0
1
Rank 1 Meta-Transitions
Microstates
D1
D2
E1
E2
F
G
0
2
4
6
8
10
12
14
16
18
20

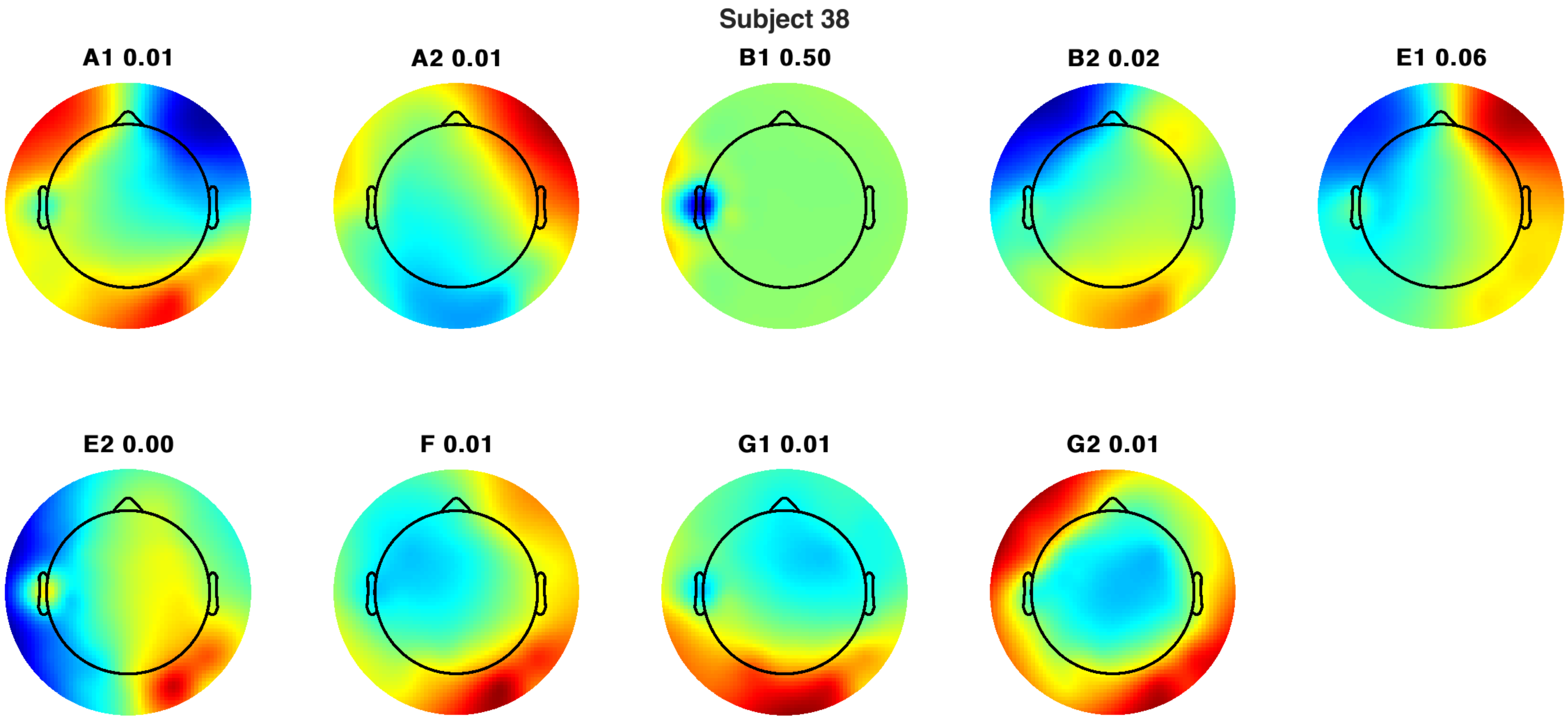
Subject 38
A1 0.01
A2 0.01
B1 0.50
B2 0.02
E1 0.06
E2 0.00
F 0.01
G1 0.01
G2 0.01

Subject 38

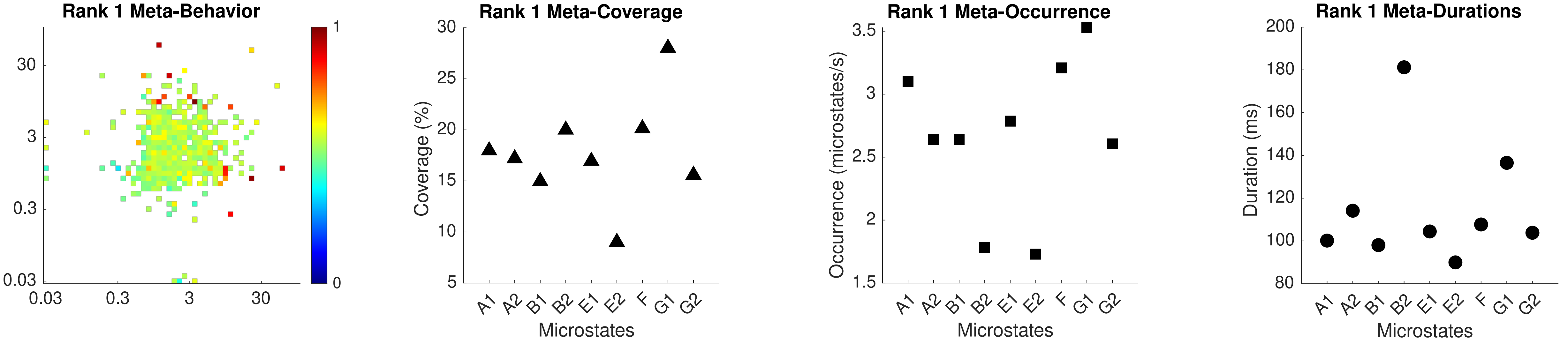

Rank 1 Meta-Behavior
30
3
0.3
0.03
0.03
0.3
3
30
1
0
Rank 1 Meta-Coverage
Coverage (%)
30
25
20
15
10
5
A1
A2
B1
B2
E1
E2
F
G1
G2
Microstates
Rank 1 Meta-Occurrence
Occurrence (microstates/s)
3.5
3
2.5
2
1.5
A1
A2
B1
B2
E1
E2
F
G1
G2
Microstates
Rank 1 Meta-Durations
Duration (ms)
200
180
160
140
120
100
80
A1
A2
B1
B2
E1
E2
F
G1
G2
Microstates

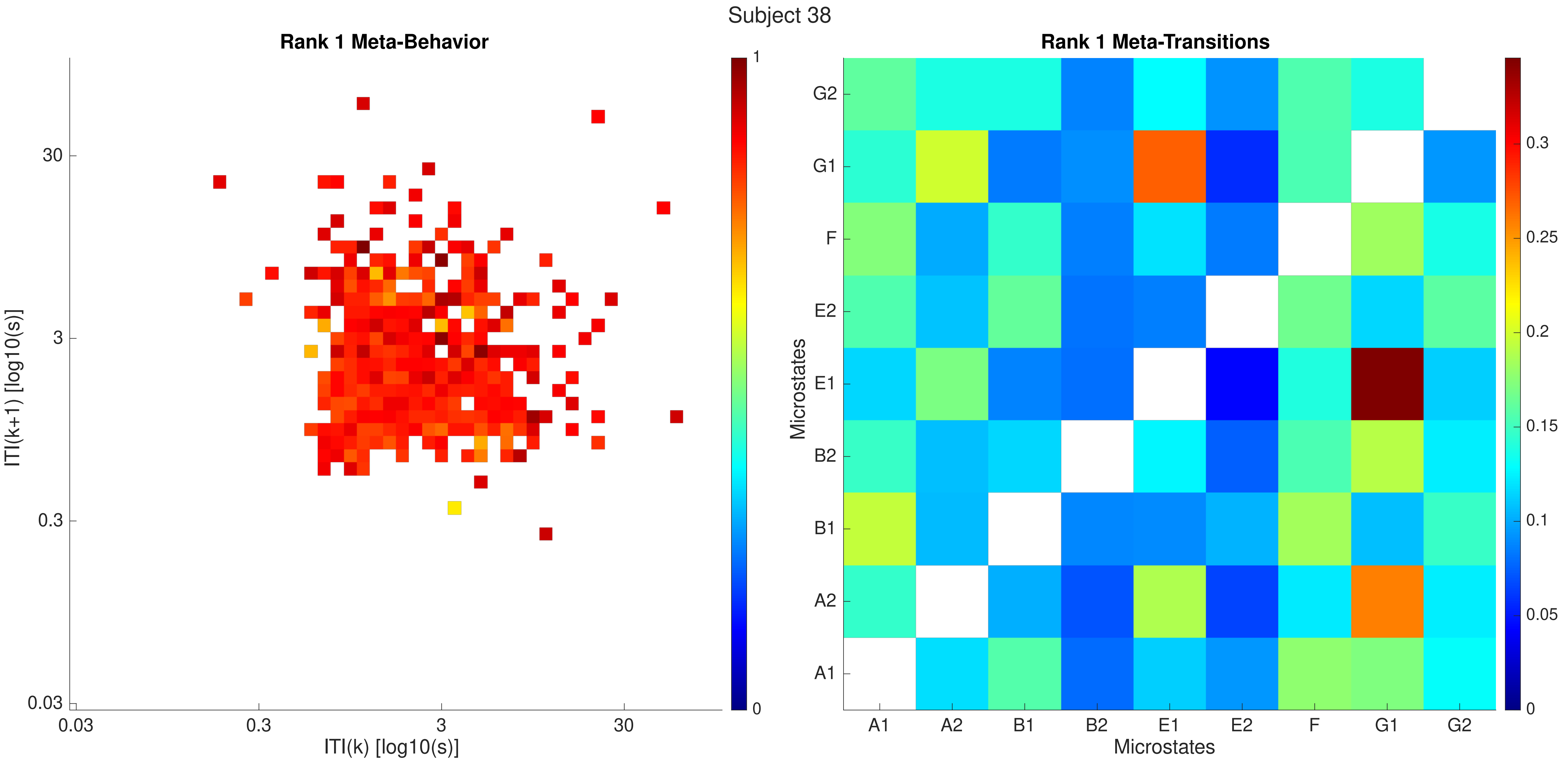

Subject 38
Rank 1 Meta-Behavior
ITI(k+1) [log10(s)]
ITI(k) [log10(s)]
0.03
0.3
3
30
0
1
Rank 1 Meta-Transitions
Microstates
A1
A2
B1
B2
E1
E2
F
G1
G2
0
0.05
0.1
0.15
0.2
0.25
0.3

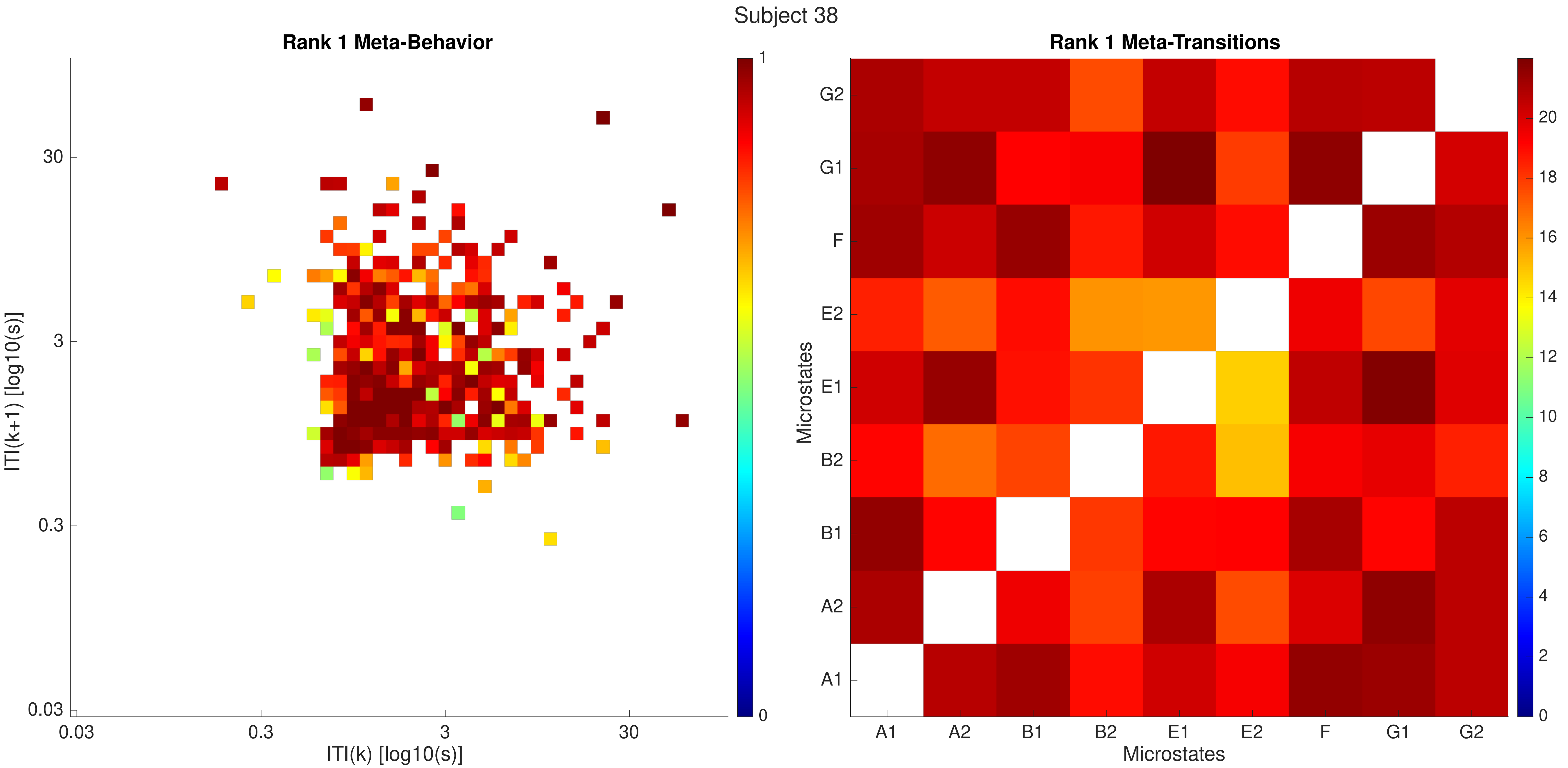
Subject 38
Rank 1 Meta-Behavior
ITI(k+1) [log10(s)]
ITI(k) [log10(s)]
0.03
0.3
3
30
0
1
Rank 1 Meta-Transitions
Microstates
A1
A2
B1
B2
E1
E2
F
G1
G2
0
2
4
6
8
10
12
14
16
18
20

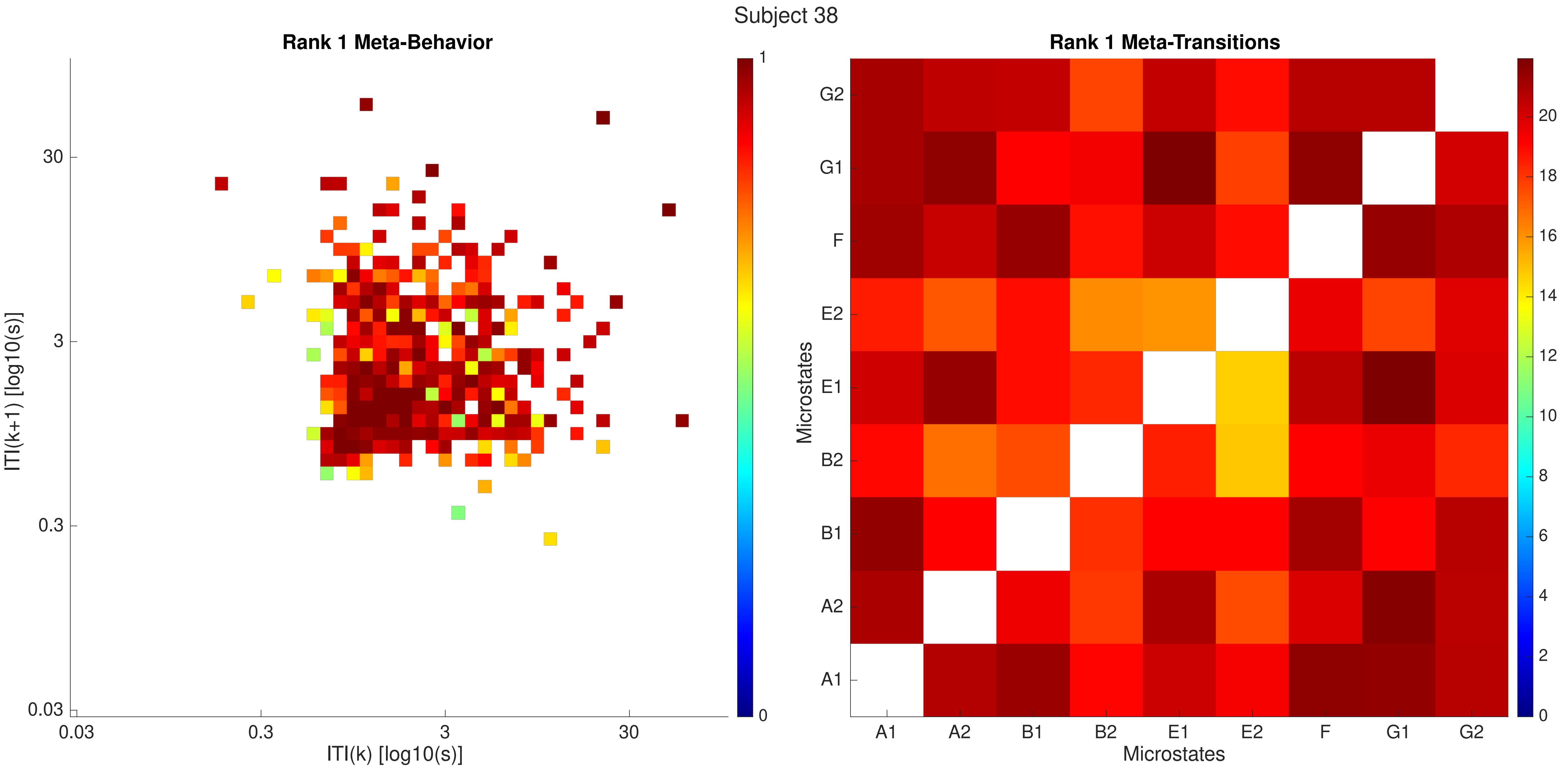
Subject 38
Rank 1 Meta-Behavior
ITI(k+1) [log10(s)]
ITI(k) [log10(s)]
0.03
0.3
3
30
0
1
Rank 1 Meta-Transitions
Microstates
A1
A2
B1
B2
E1
E2
F
G1
G2
0
2
4
6
8
10
12
14
16
18
20

Subject 39

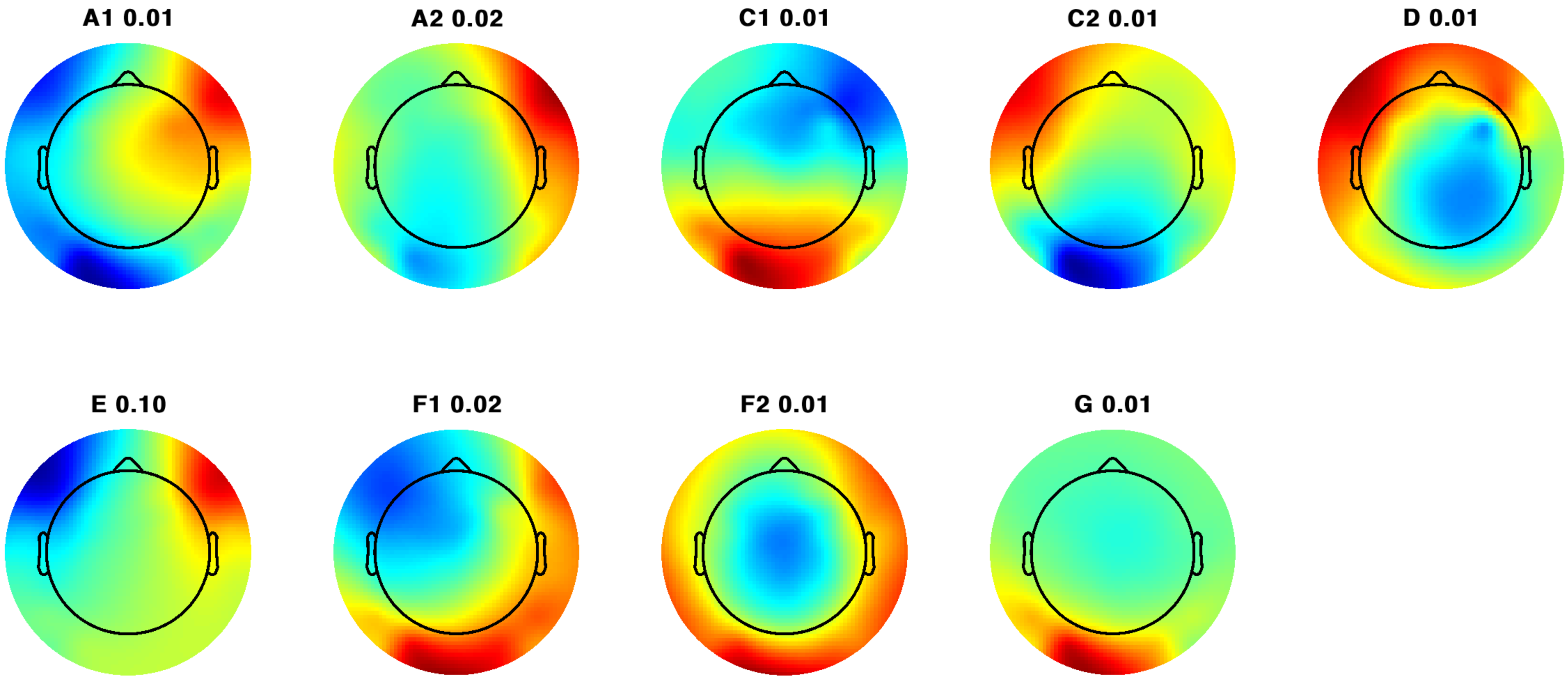

Subject 39

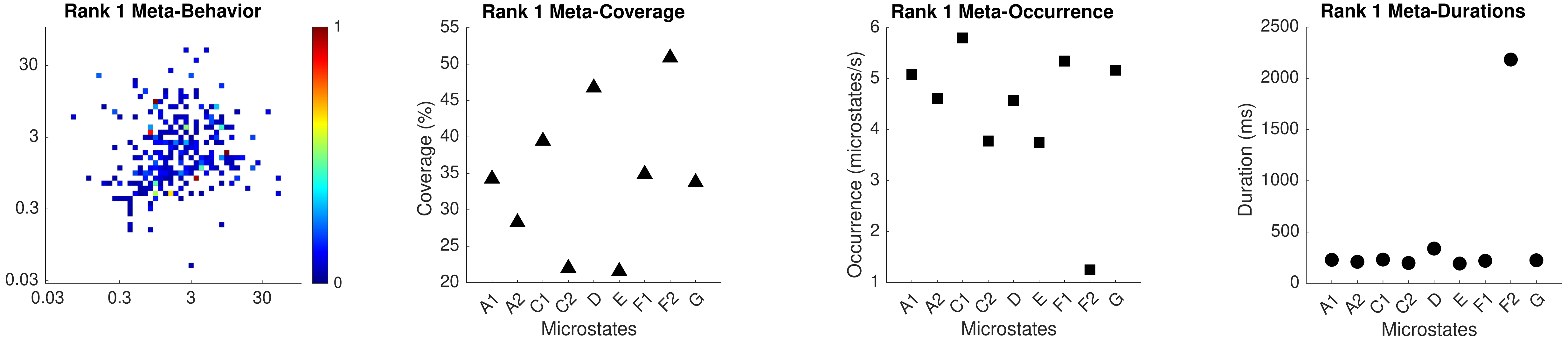

Rank 1 Meta-Behavior
30
3
0.3
0.03
0.03
0.3
3
30
1
0
Rank 1 Meta-Coverage
Coverage (%)
55
50
45
40
35
30
25
20
A1
A2
C1
C2
D
E
F1
F2
G
Microstates
Rank 1 Meta-Occurrence
Occurrence (microstates/s)
6
5
4
3
2
1
A1
A2
C1
C2
D
E
F1
F2
G
Microstates
Rank 1 Meta-Durations
Duration (ms)
2500
2000
1500
1000
500
0
A1
A2
C1
C2
D
E
F1
F2
G
Microstates

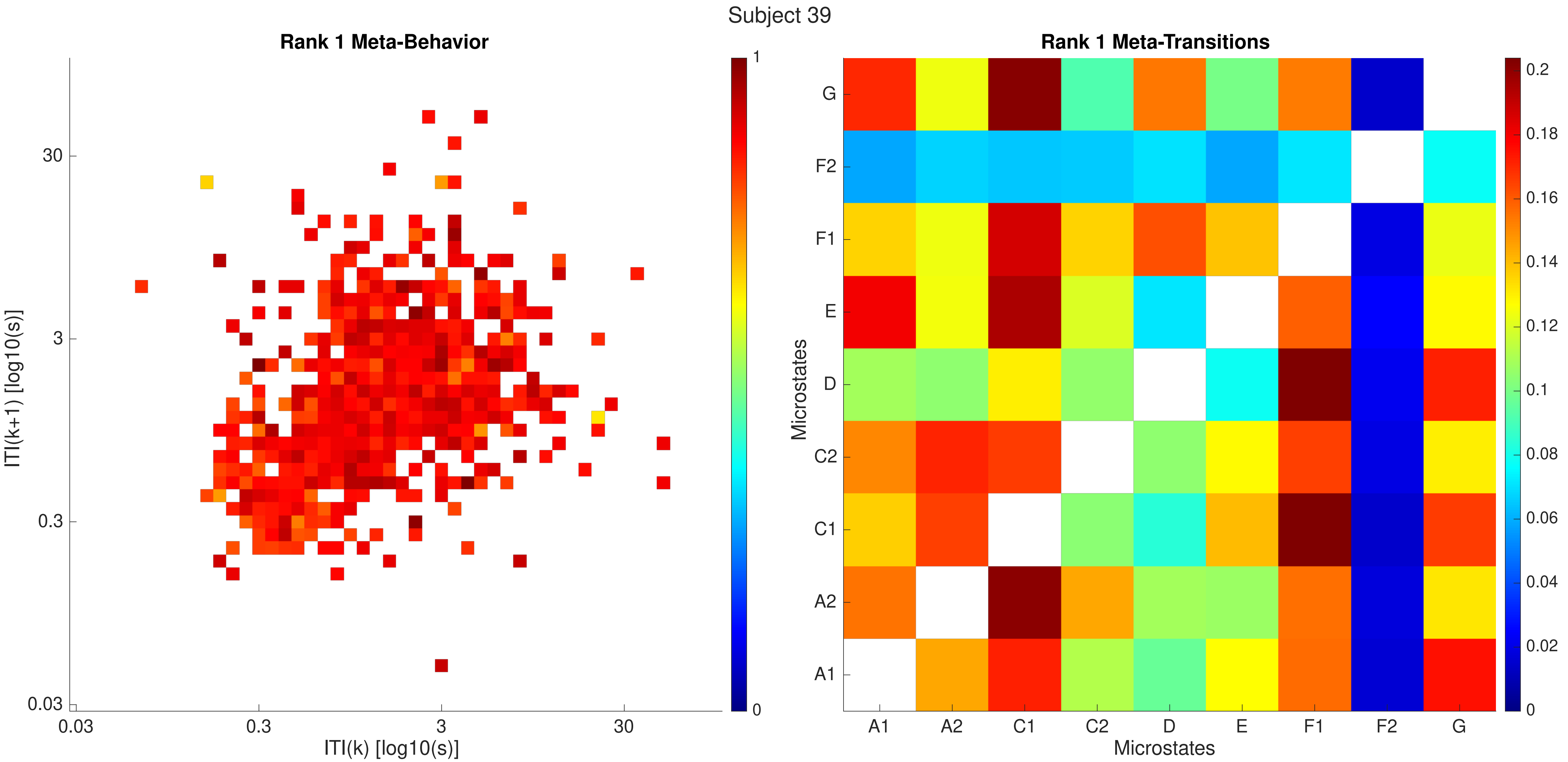

Subject 39
Rank 1 Meta-Behavior
ITI(k+1) [log10(s)]
ITI(k) [log10(s)]
0.03
0.3
3
30
0
1
Rank 1 Meta-Transitions
Microstates
A1
A2
C1
C2
D
E
F1
F2
G
0
0.02
0.04
0.06
0.08
0.1
0.12
0.14
0.16
0.18
0.2

Subject 39

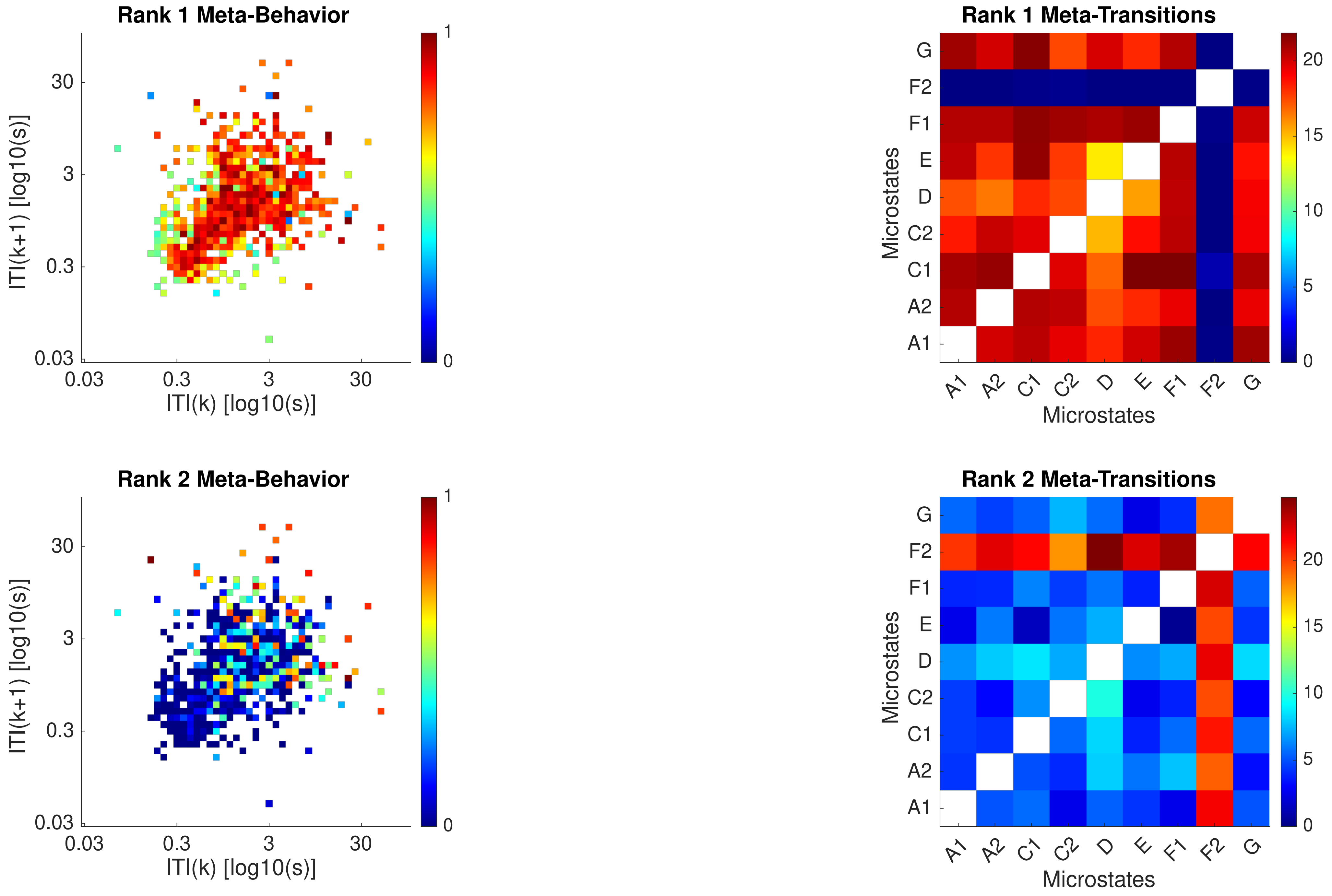
Rank 1 Meta-Behavior
ITI(k+1) [log10(s)]
ITI(k) [log10(s)]
0.03
0.3
3
30
0
1
Rank 1 Meta-Transitions
Microstates
A1
A2
C1
C2
D
E
F1
F2
G
0
5
10
15
20
Rank 2 Meta-Behavior
Rank 2 Meta-Transitions

Subject 39

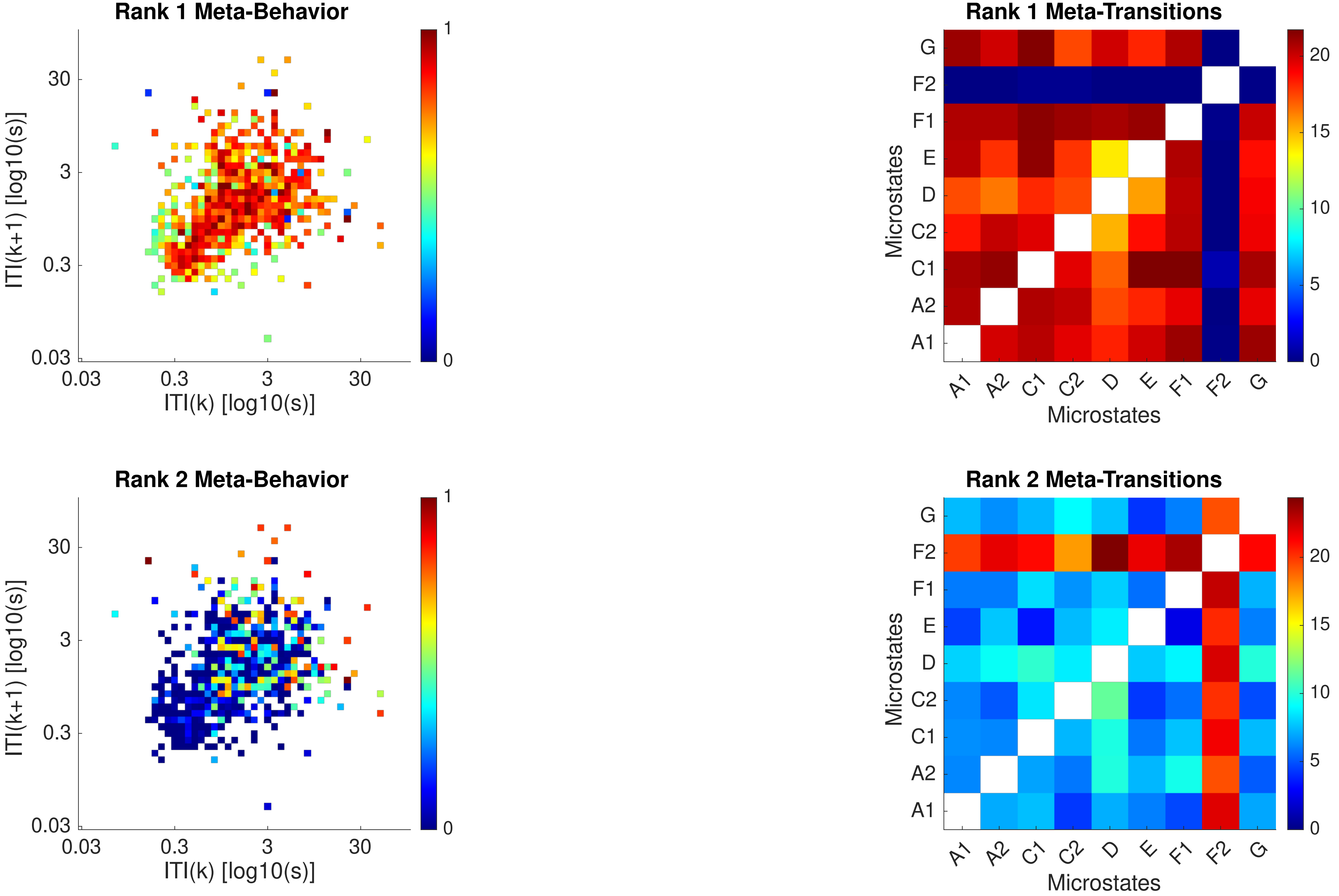

Rank 1 Meta-Behavior
Rank 1 Meta-Transitions
Rank 2 Meta-Behavior
Rank 2 Meta-Transitions
ITI(k+1) [log10(s)]
ITI(k) [log10(s)]
0.03
0.3
3
30
0
1
Microstates
A1
A2
C1
C2
D
E
F1
F2
G
0
5
10
15
20

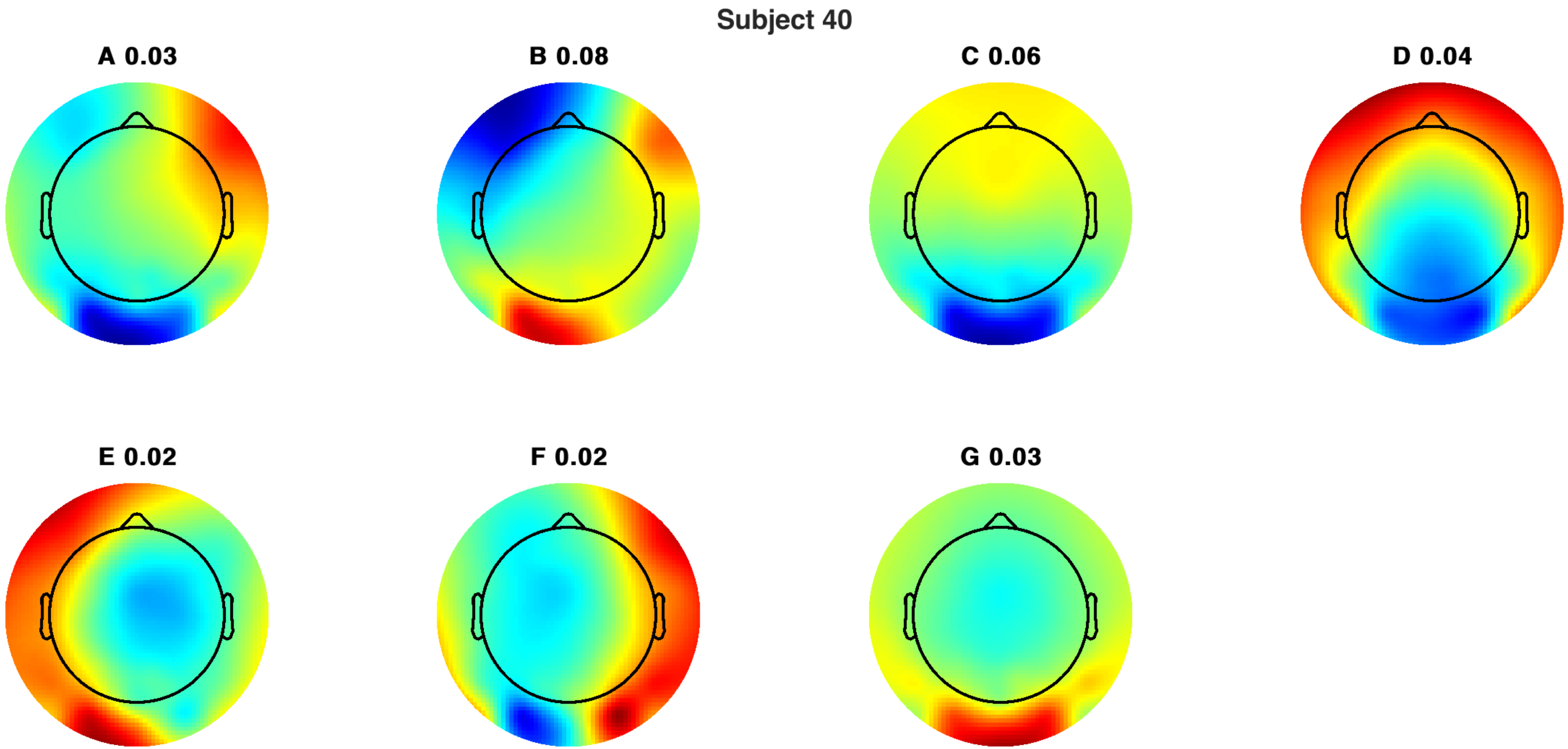
Subject 40
A 0.03
B 0.08
C 0.06
D 0.04
E 0.02
F 0.02
G 0.03

Subject 40

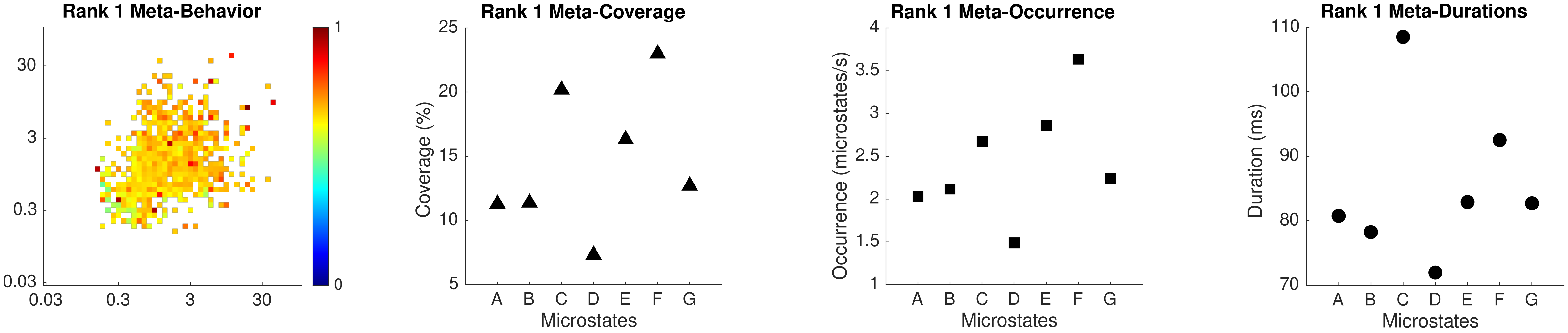
Rank 1 Meta-Behavior
30
3
0.3
0.03
0.03
0.3
3
30
1
0
Rank 1 Meta-Coverage
Coverage (%)
25
20
15
10
5
A B C D E F G
Microstates
Rank 1 Meta-Occurrence
Occurrence (microstates/s)
4
3.5
3
2.5
2
1.5
1
A B C D E F G
Microstates
Rank 1 Meta-Durations
Duration (ms)
110
100
90
80
70
A B C D E F G
Microstates

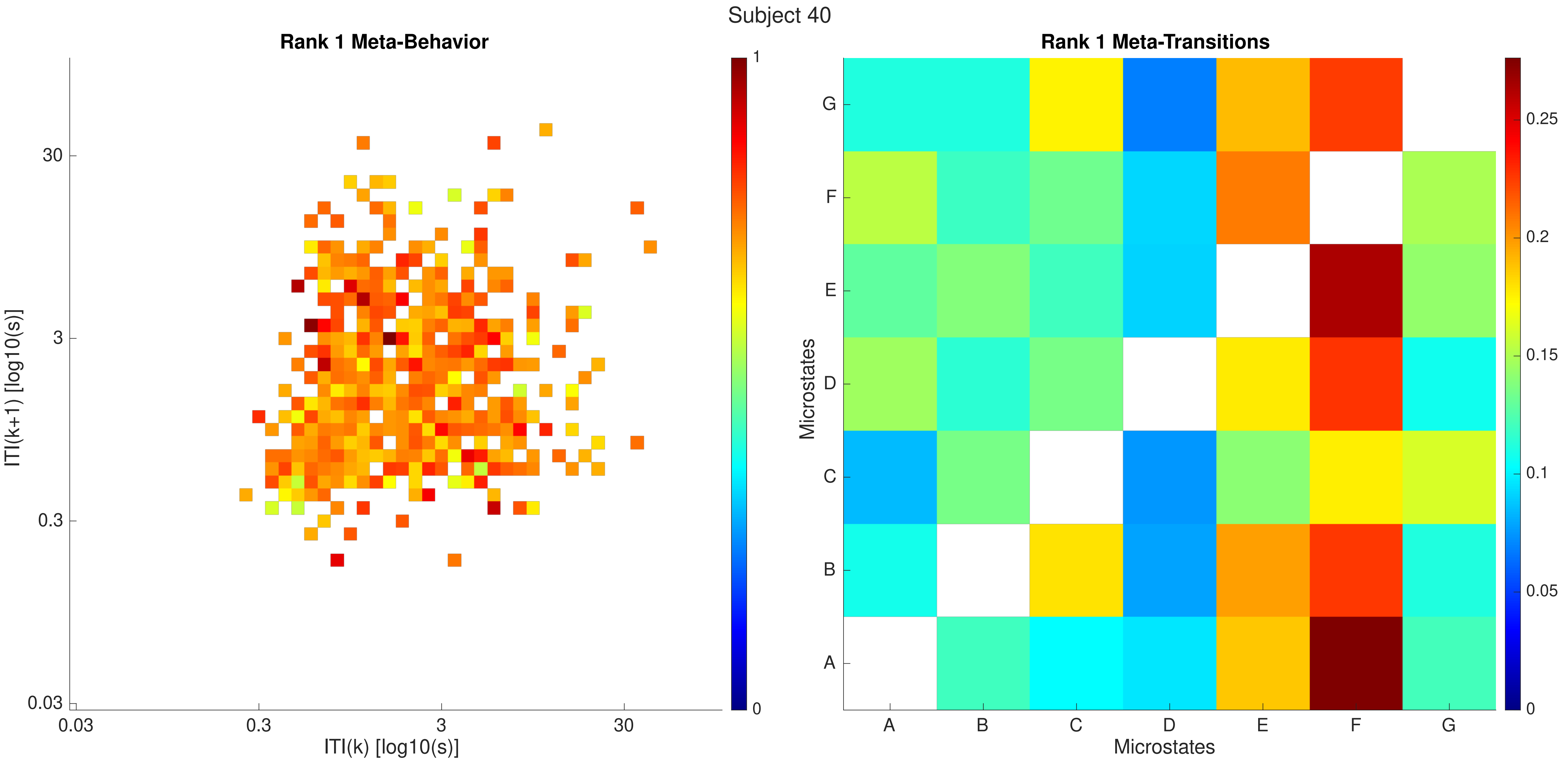
Subject 40
Rank 1 Meta-Behavior
ITI(k+1) [log10(s)]
ITI(k) [log10(s)]
0.03
0.3
3
30
0
1
Rank 1 Meta-Transitions
Microstates
A
B
C
D
E
F
G
0
0.05
0.1
0.15
0.2
0.25

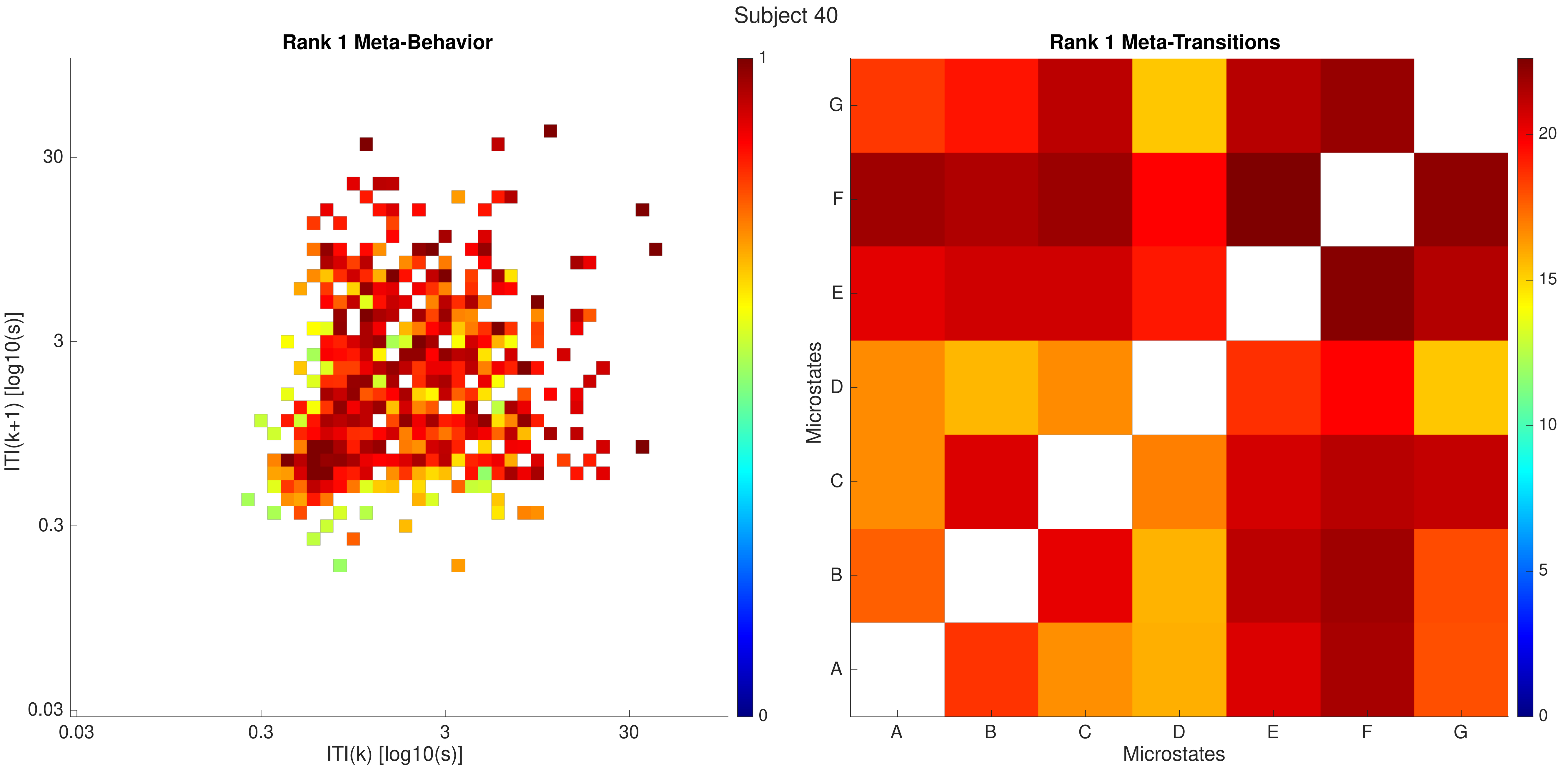

Subject 40
Rank 1 Meta-Behavior
ITI(k+1) [log10(s)]
30
3
0.3
0.03
0.03
0.3
3
30
ITI(k) [log10(s)]
1
0
Rank 1 Meta-Transitions
Microstates
G
F
E
D
C
B
A
A
B
C
D
E
F
G
Microstates
20
15
10
5
0

Subject 40

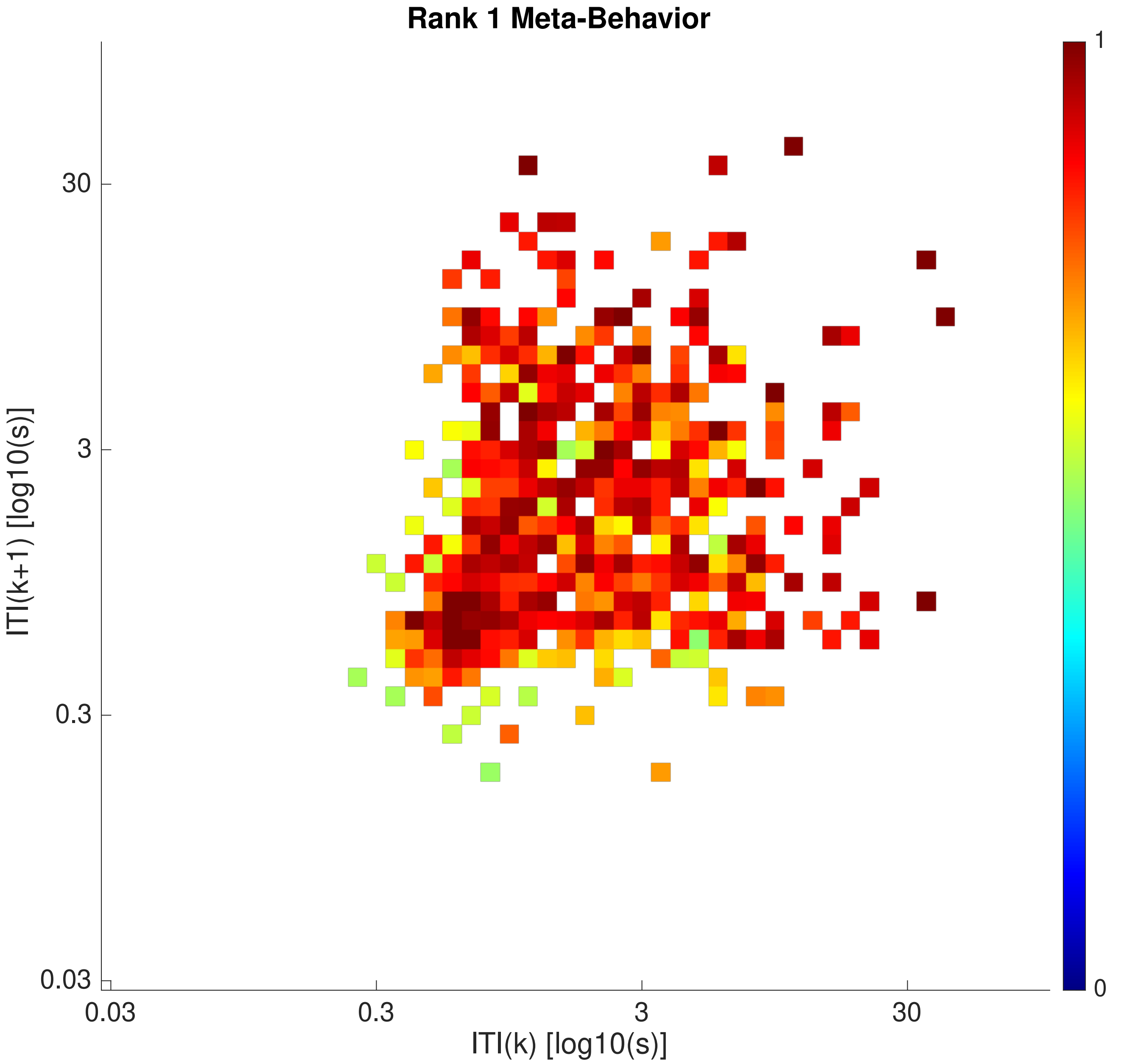

Rank 1 Meta-Behavior
30
3
0.3
0.03
ITI(k+1) [log10(s)]
0.03
0.3
3
30
ITI(k) [log10(s)]
1
0


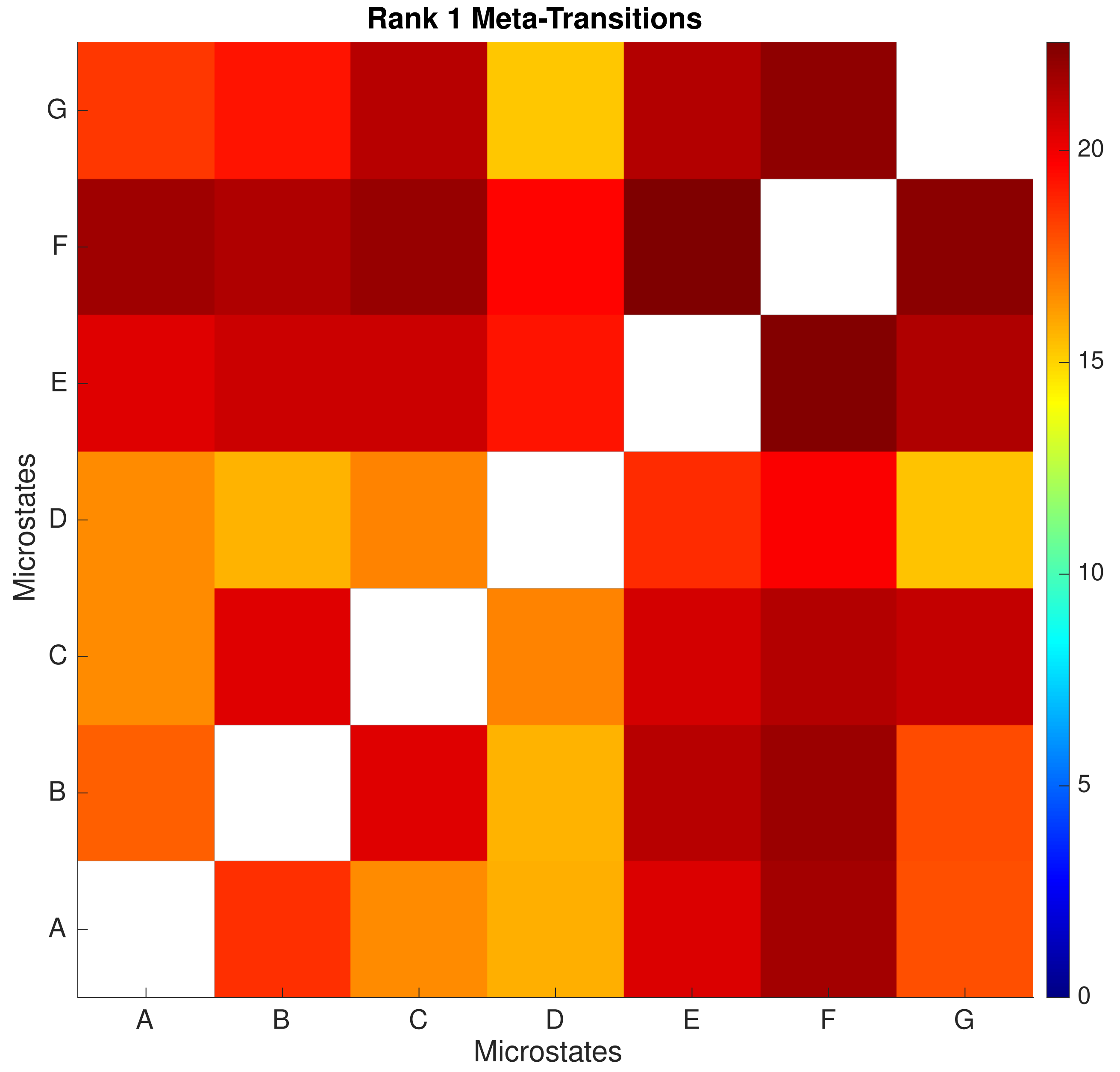

Rank 1 Meta-Transitions
G
F
E
D
C
B
A
Microstates
A
B
C
D
E
F
G
Microstates
20
15
10
5
0

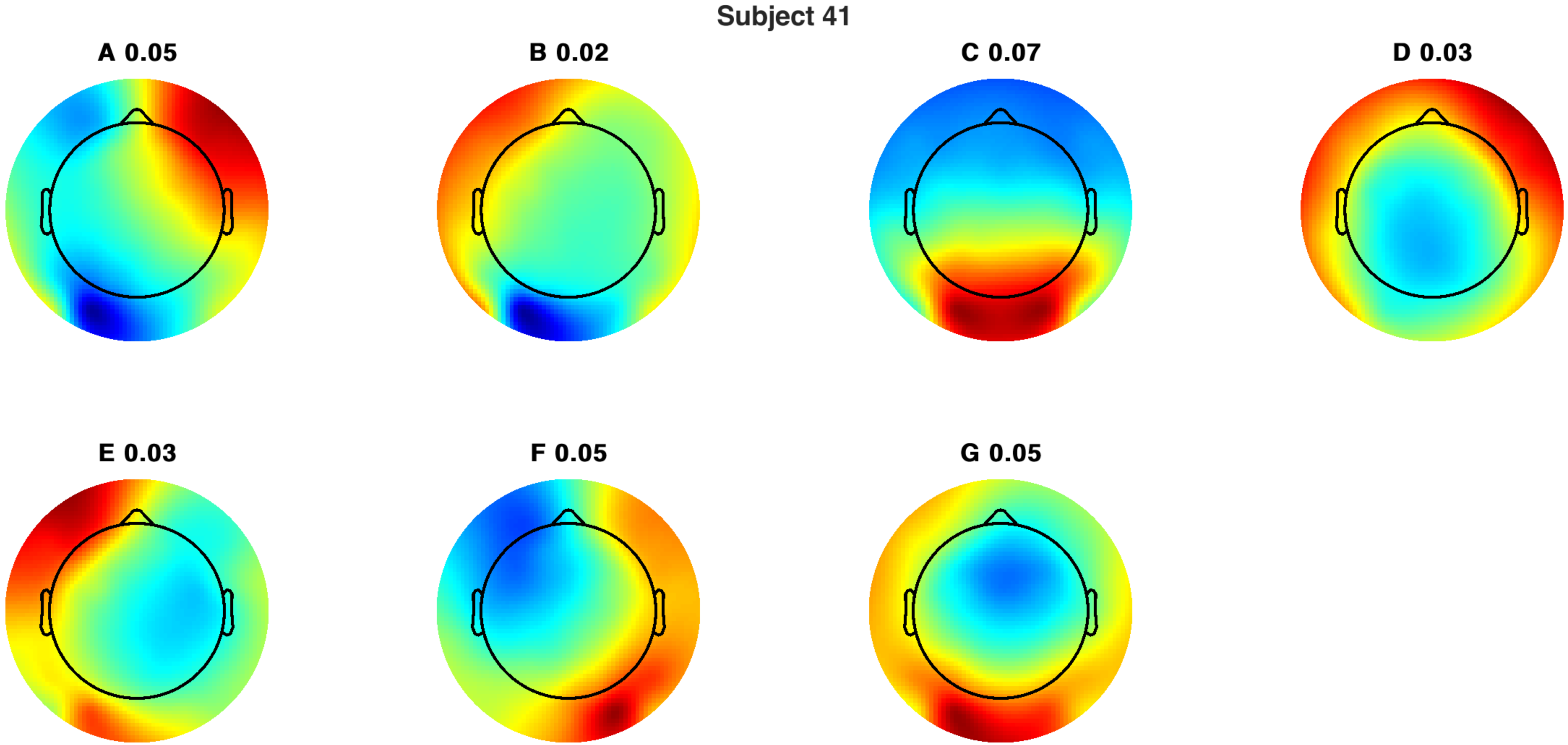
Subject 41
A 0.05
B 0.02
C 0.07
D 0.03
E 0.03
F 0.05
G 0.05

Subject 41

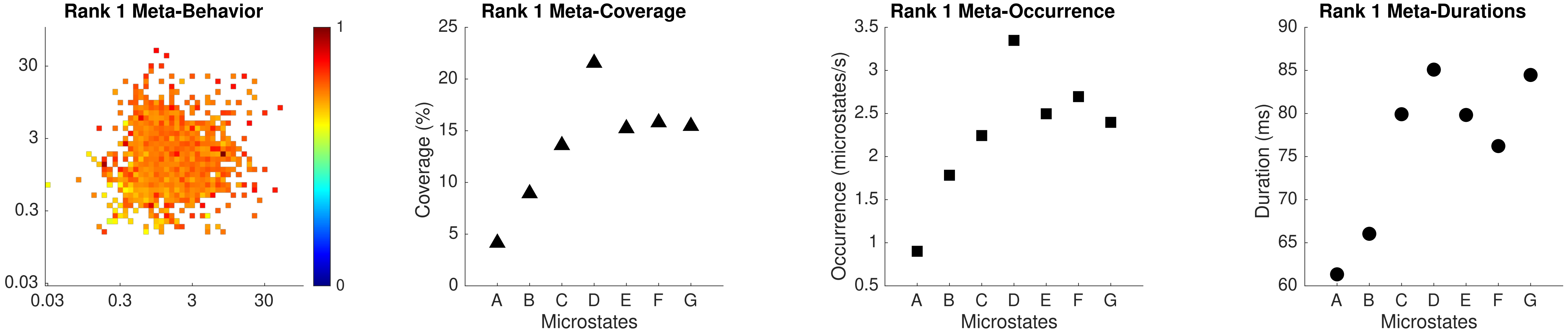

Rank 1 Meta-Behavior
Rank 1 Meta-Coverage
Coverage (%)
Microstates
Rank 1 Meta-Occurrence
Occurrence (microstates/s)
Microstates
Rank 1 Meta-Durations
Duration (ms)
Microstates

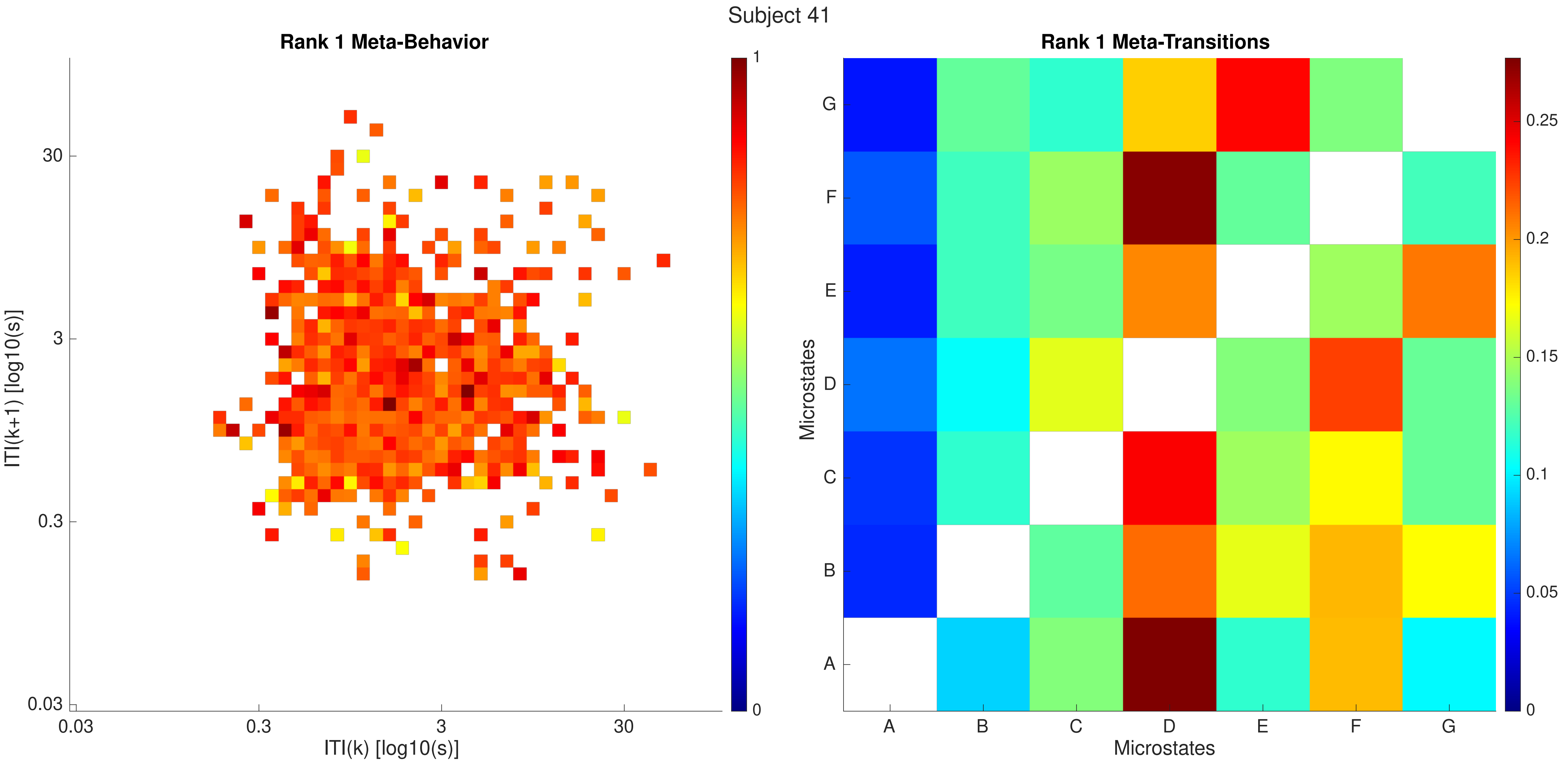
Subject 41
Rank 1 Meta-Behavior
ITI(k+1) [log10(s)]
ITI(k) [log10(s)]
0.03
0.3
3
30
0
1
Rank 1 Meta-Transitions
Microstates
A
B
C
D
E
F
G
0
0.05
0.1
0.15
0.2
0.25

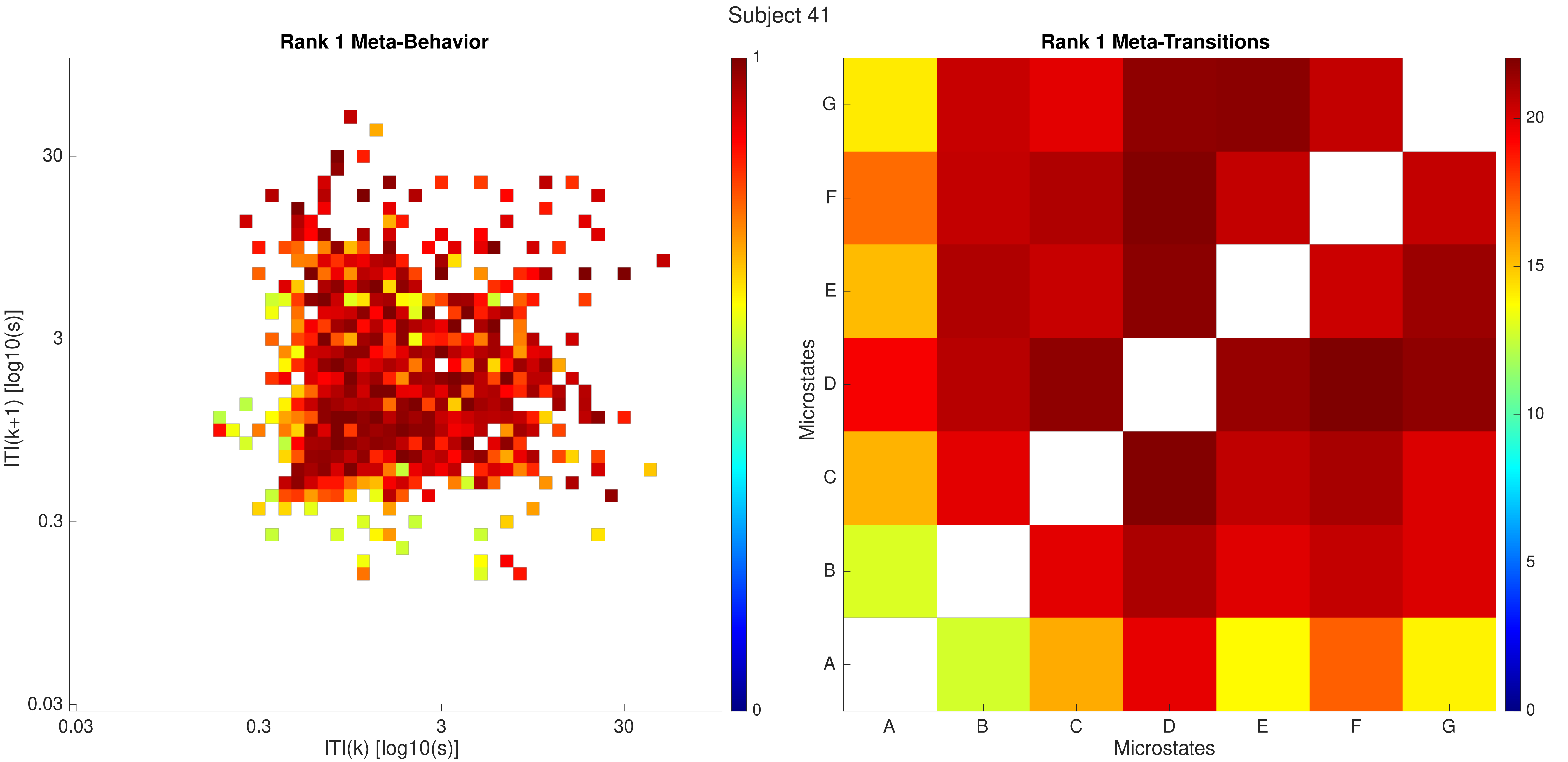

Subject 41
Rank 1 Meta-Behavior
ITI(k+1) [log10(s)]
ITI(k) [log10(s)]
0.03
0.3
3
30
0
1
Rank 1 Meta-Transitions
Microstates
A
B
C
D
E
F
G
0
5
10
15
20

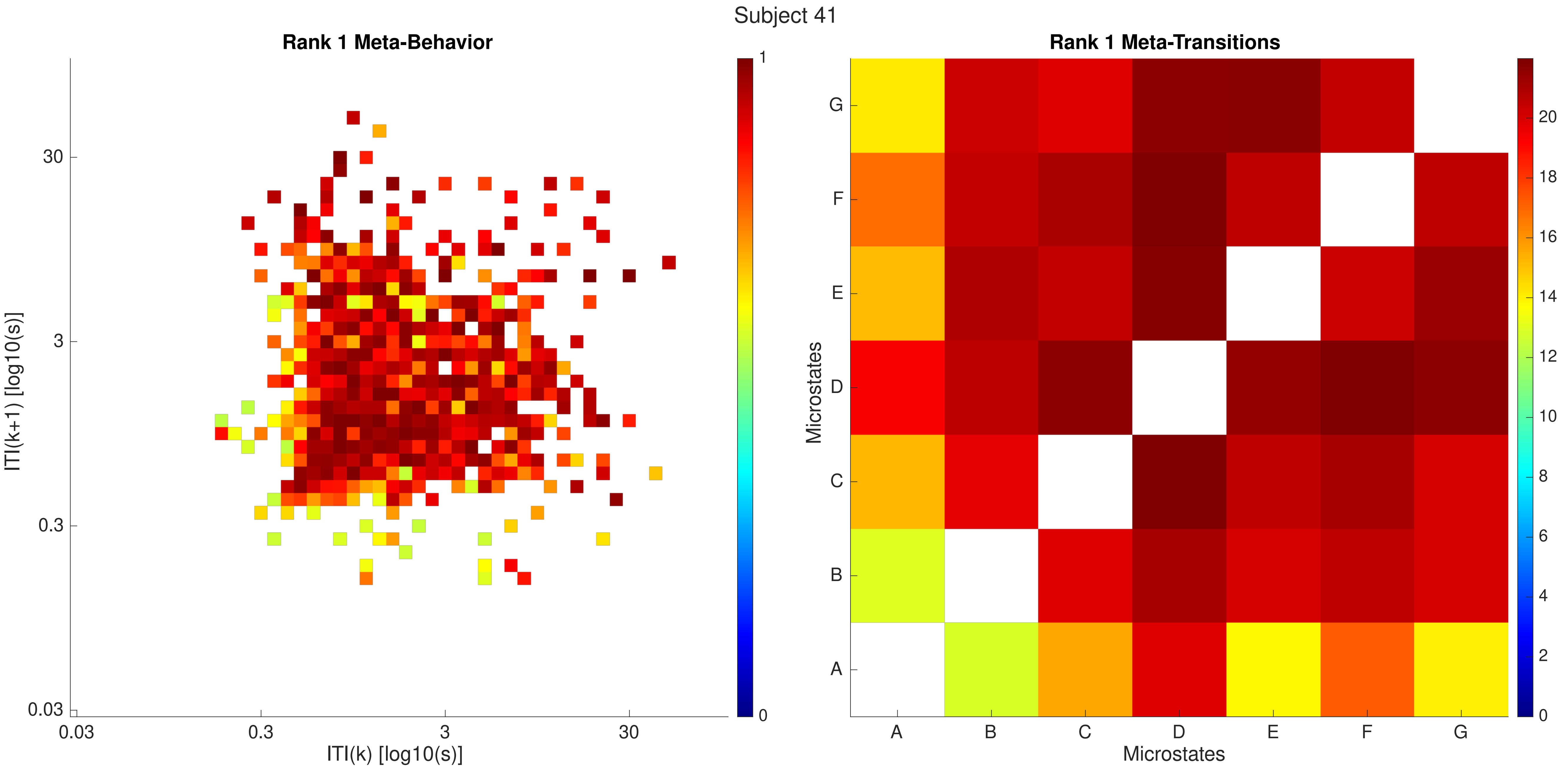
Subject 41
Rank 1 Meta-Behavior
ITI(k+1) [log10(s)]
ITI(k) [log10(s)]
0.03
0.3
3
30
0
1
Rank 1 Meta-Transitions
Microstates
A
B
C
D
E
F
G
0
2
4
6
8
10
12
14
16
18
20

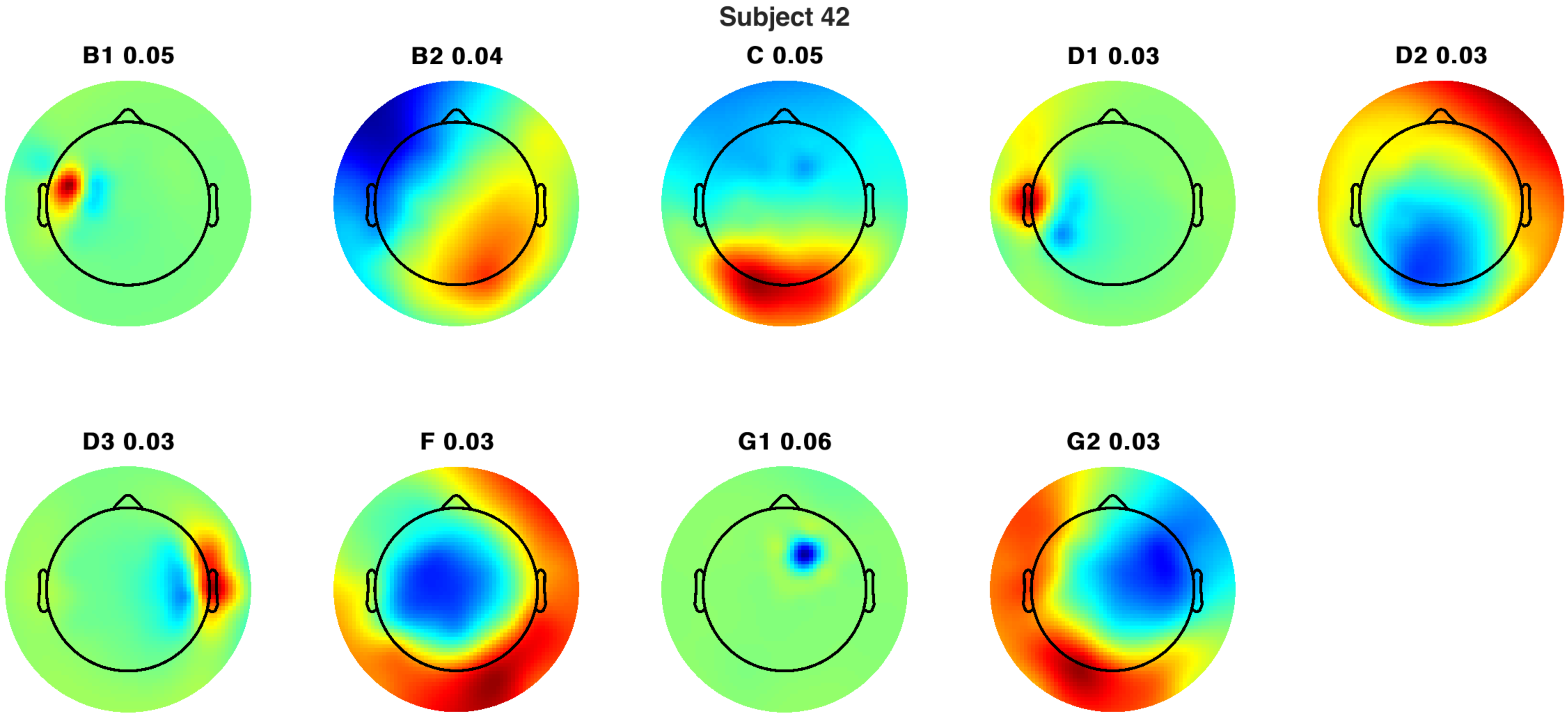
Subject 42
B1 0.05
B2 0.04
C 0.05
D1 0.03
D2 0.03
D3 0.03
F 0.03
G1 0.06
G2 0.03

Subject 42

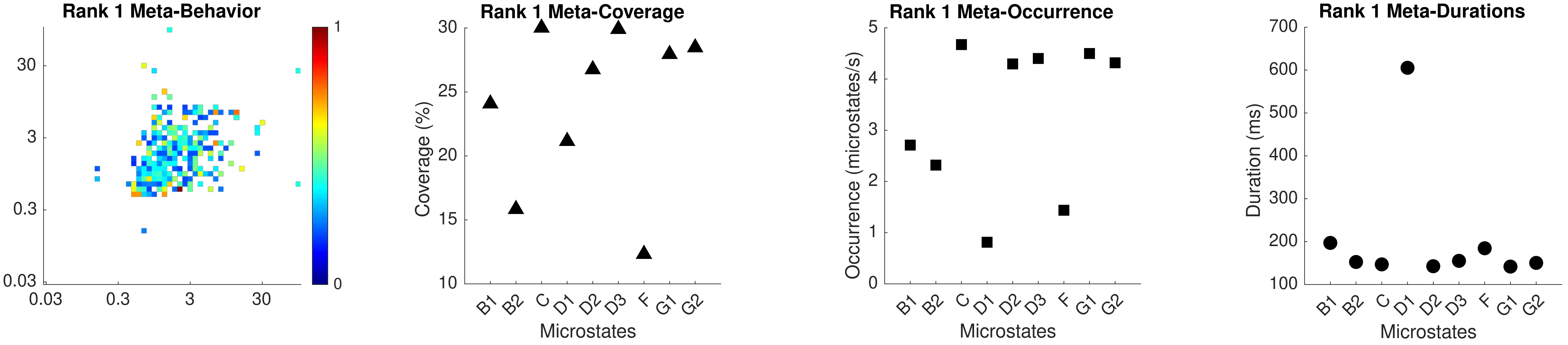

Rank 1 Meta-Behavior
Rank 1 Meta-Coverage
Coverage (%)
Microstates
Rank 1 Meta-Occurrence
Occurrence (microstates/s)
Microstates
Rank 1 Meta-Durations
Duration (ms)
Microstates
B1 B2 C D1 D2 D3 F G1 G2

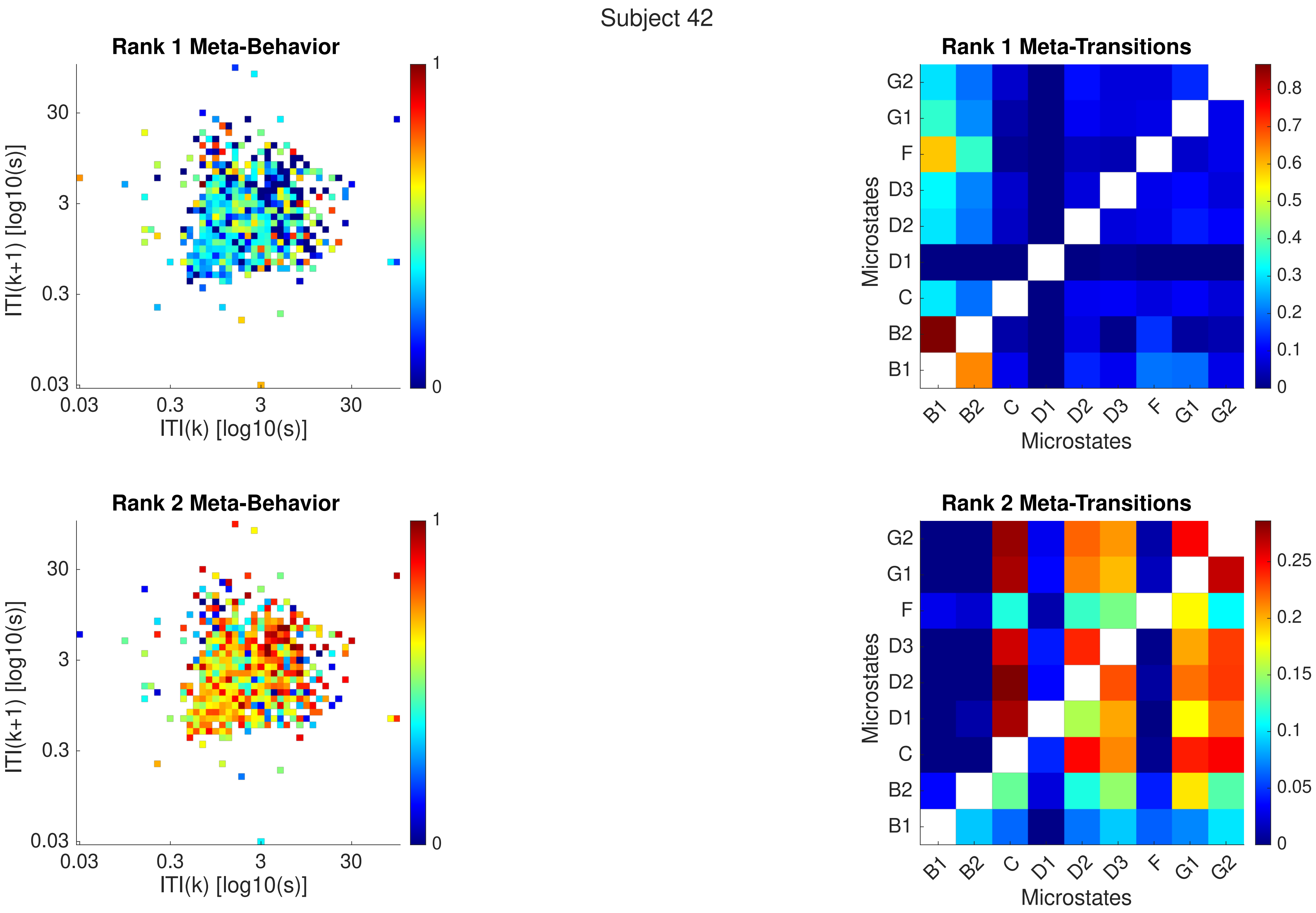
Subject 42
Rank 1 Meta-Behavior
ITI(k+1) [log10(s)]
ITI(k) [log10(s)]
0.03
0.3
3
30
1
0
Rank 1 Meta-Transitions
Microstates
G2
G1
F
D3
D2
D1
C
B2
B1
0.8
0.7
0.6
0.5
0.4
0.3
0.2
0.1
0
Rank 2 Meta-Behavior
ITI(k+1) [log10(s)]
ITI(k) [log10(s)]
Rank 2 Meta-Transitions
Microstates
0.25
0.2
0.15
0.1
0.05

Subject 42

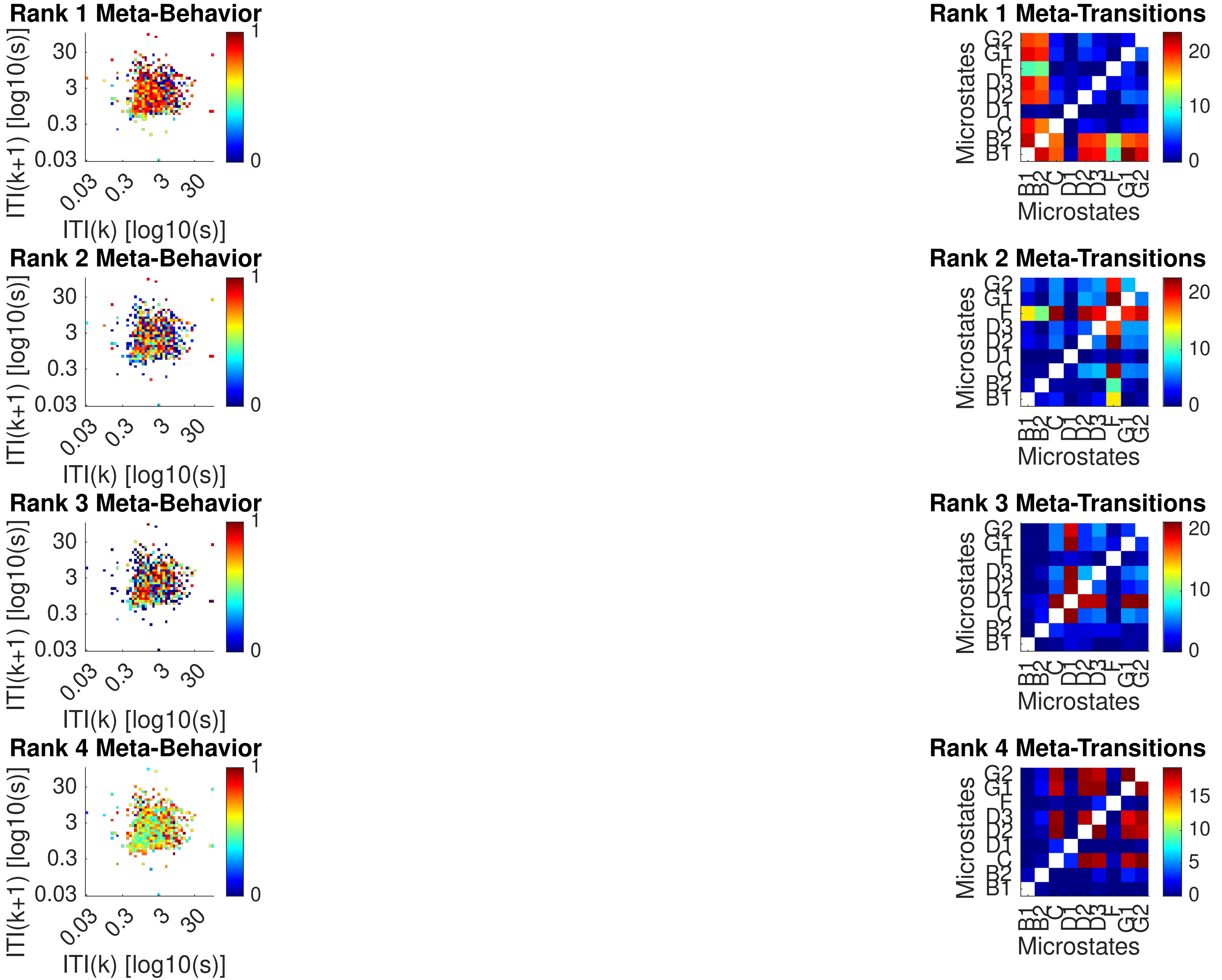

Rank 1 Meta-Behavior
Rank 2 Meta-Behavior
Rank 3 Meta-Behavior
Rank 4 Meta-Behavior
ITI(k+1) [log10(s)]
ITI(k) [log10(s)]
0.03
0.3
3
30
0
1
Rank 1 Meta-Transitions
Rank 2 Meta-Transitions
Rank 3 Meta-Transitions
Rank 4 Meta-Transitions
Microstates
B1
B2
C
D1
D2
D3
F
G1
G2
0
5
10
15
20

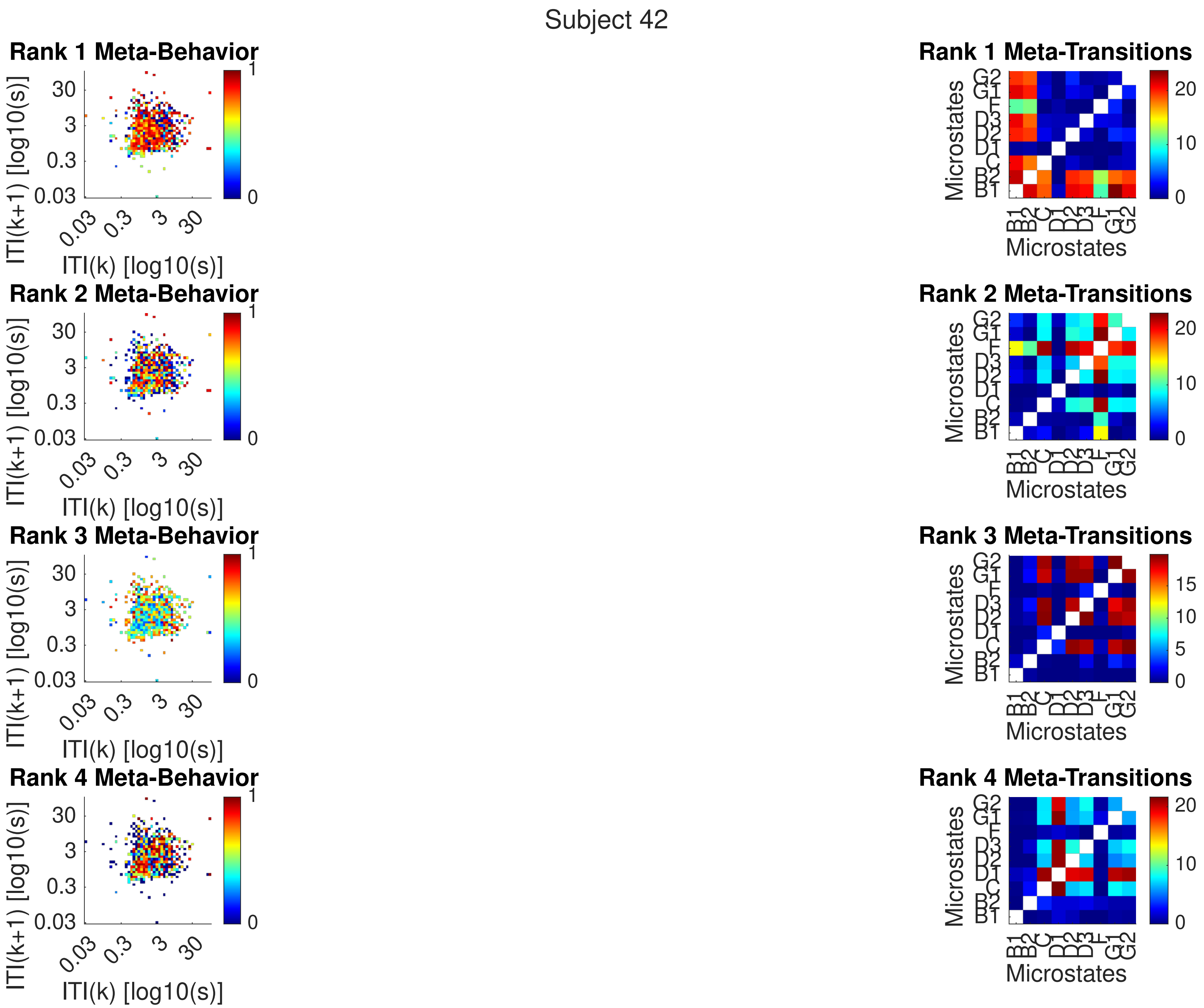

Subject 42
Rank 1 Meta-Behavior
Rank 2 Meta-Behavior
Rank 3 Meta-Behavior
Rank 4 Meta-Behavior
ITI(k+1) [log10(s)]
ITI(k) [log10(s)]
0.03
0.3
3
30
0
1
Rank 1 Meta-Transitions
Rank 2 Meta-Transitions
Rank 3 Meta-Transitions
Rank 4 Meta-Transitions
Microstates
B1
B2
C
D1
D2
D3
F
G1
G2
0
5
10
15
20

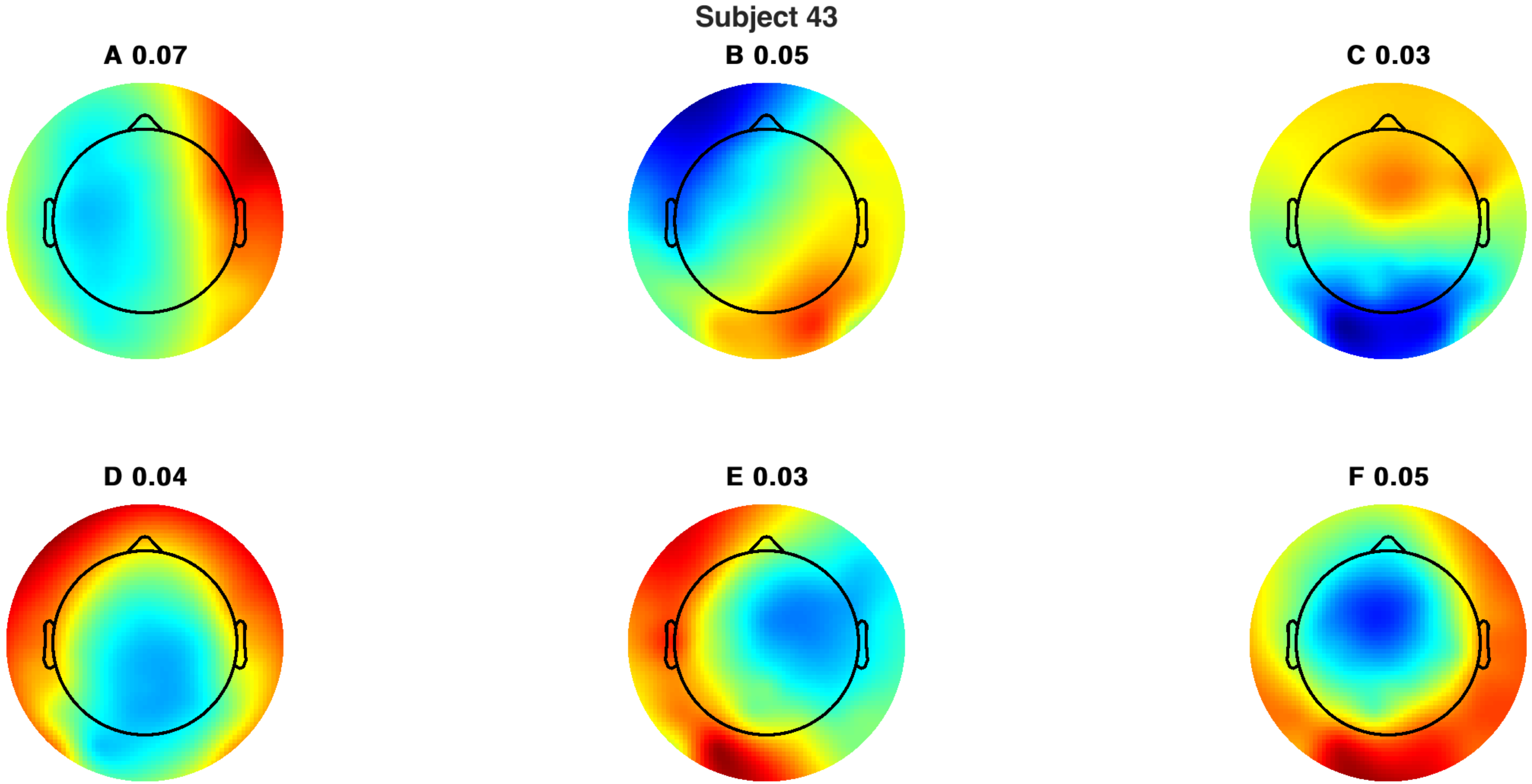
Subject 43
A 0.07
B 0.05
C 0.03
D 0.04
E 0.03
F 0.05

Subject 43

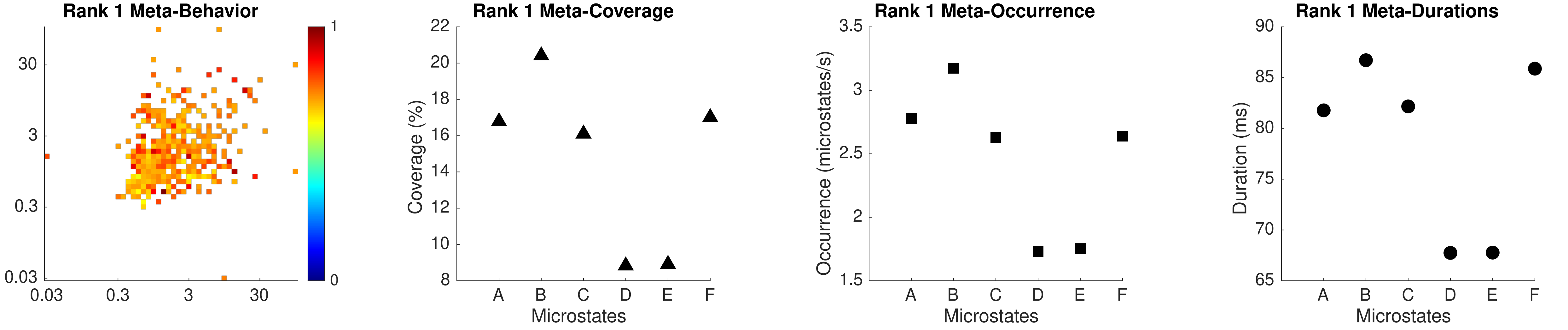

Rank 1 Meta-Behavior
30
3
0.3
0.03
0.03
0.3
3
30
1
0
Rank 1 Meta-Coverage
Coverage (%)
22
20
18
16
14
12
10
8
A
B
C
D
E
F
Microstates
Rank 1 Meta-Occurrence
Occurrence (microstates/s)
3.5
3
2.5
2
1.5
A
B
C
D
E
F
Microstates
Rank 1 Meta-Durations
Duration (ms)
90
85
80
75
70
65
A
B
C
D
E
F
Microstates

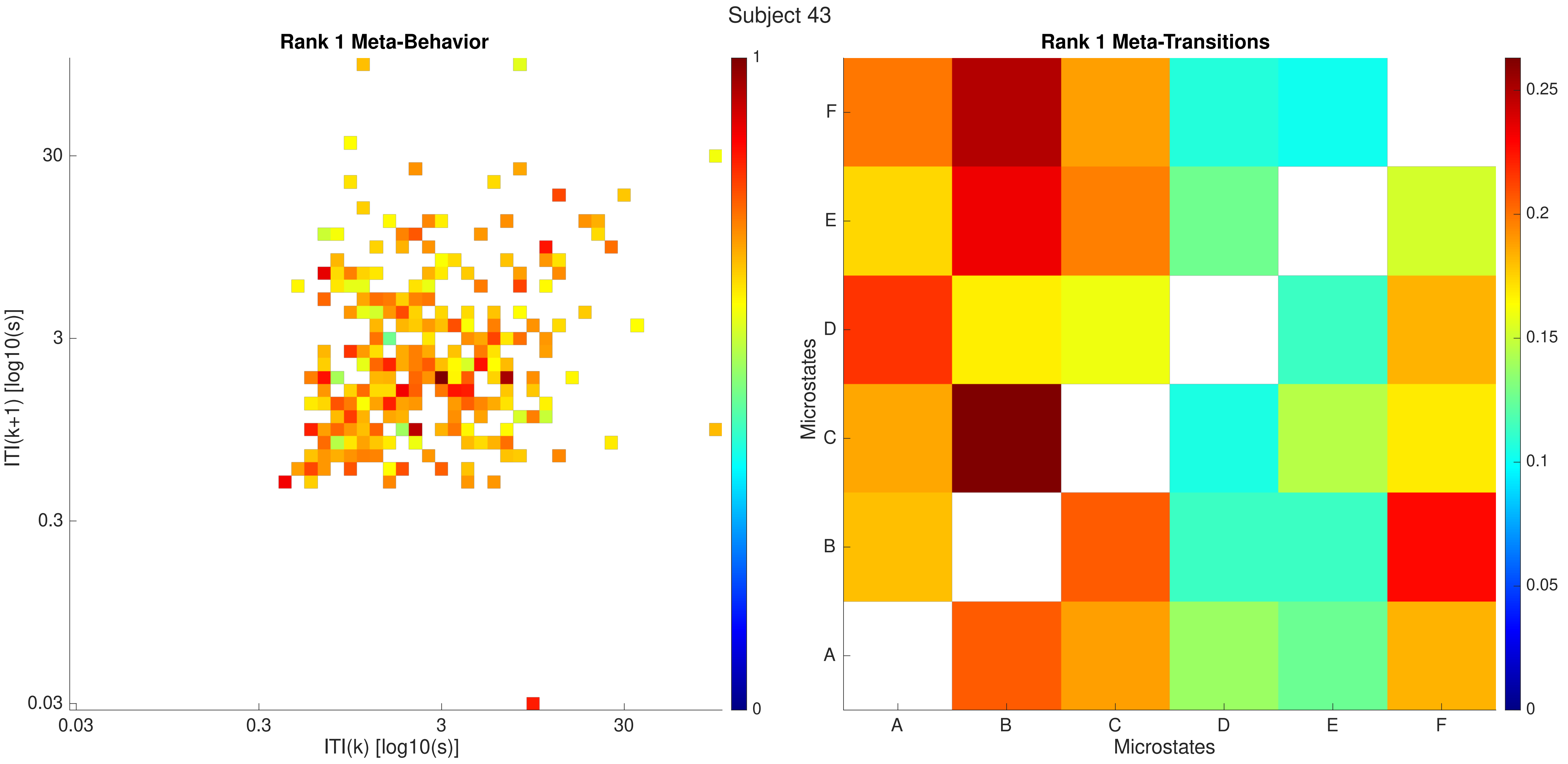

Subject 43
Rank 1 Meta-Behavior
ITI(k+1) [log10(s)]
30
3
0.3
0.03
0.03
0.3
3
30
ITI(k) [log10(s)]
1
0
Rank 1 Meta-Transitions
Microstates
F
E
D
C
B
A
A
B
C
D
E
F
Microstates
0.25
0.2
0.15
0.1
0.05
0

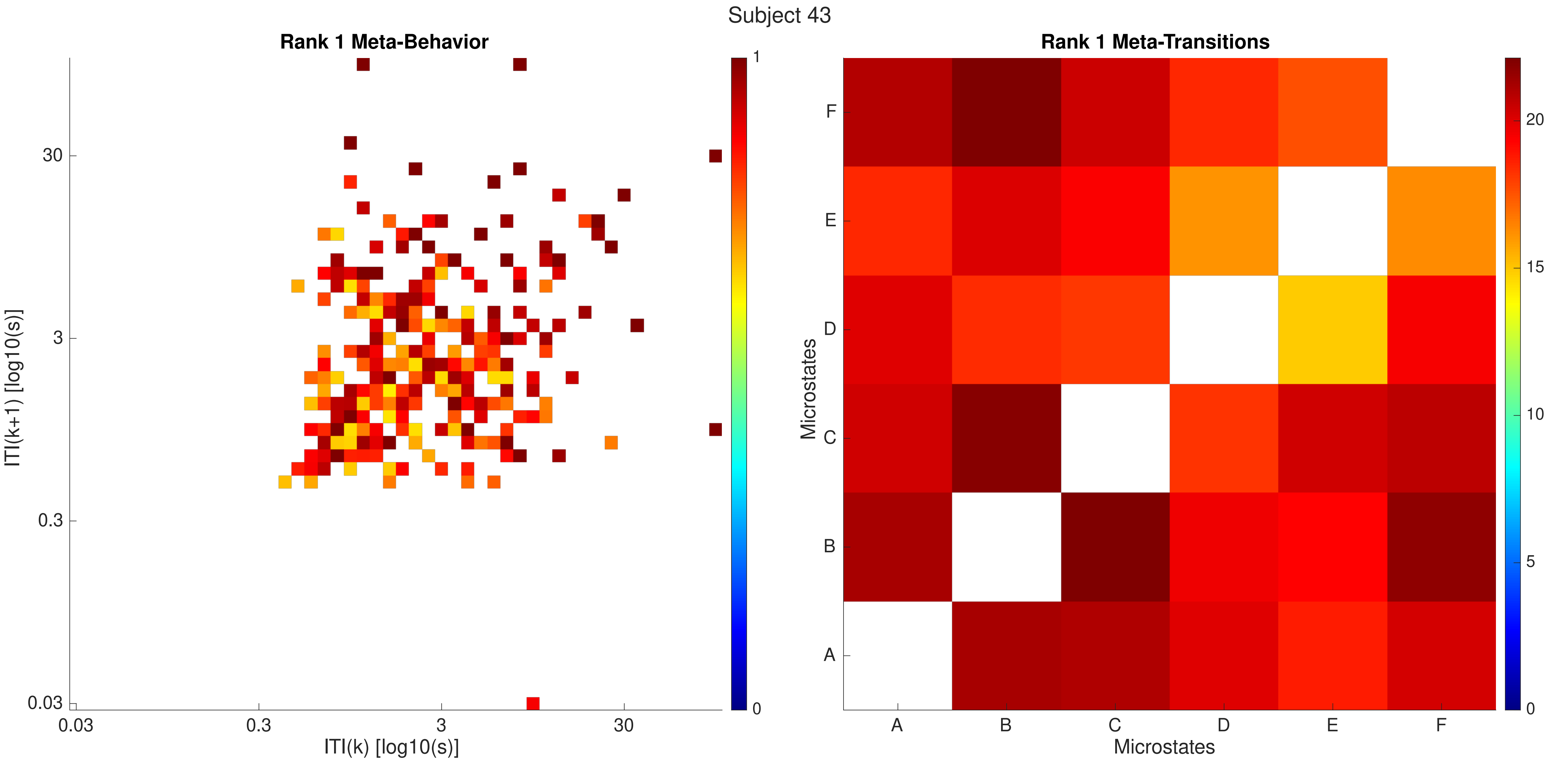
Subject 43
Rank 1 Meta-Behavior
ITI(k+1) [log10(s)]
30
3
0.3
0.03
0.03
0.3
3
30
ITI(k) [log10(s)]
1
0
Rank 1 Meta-Transitions
Microstates
F
E
D
C
B
A
A
B
C
D
E
F
Microstates
20
15
10
5
0

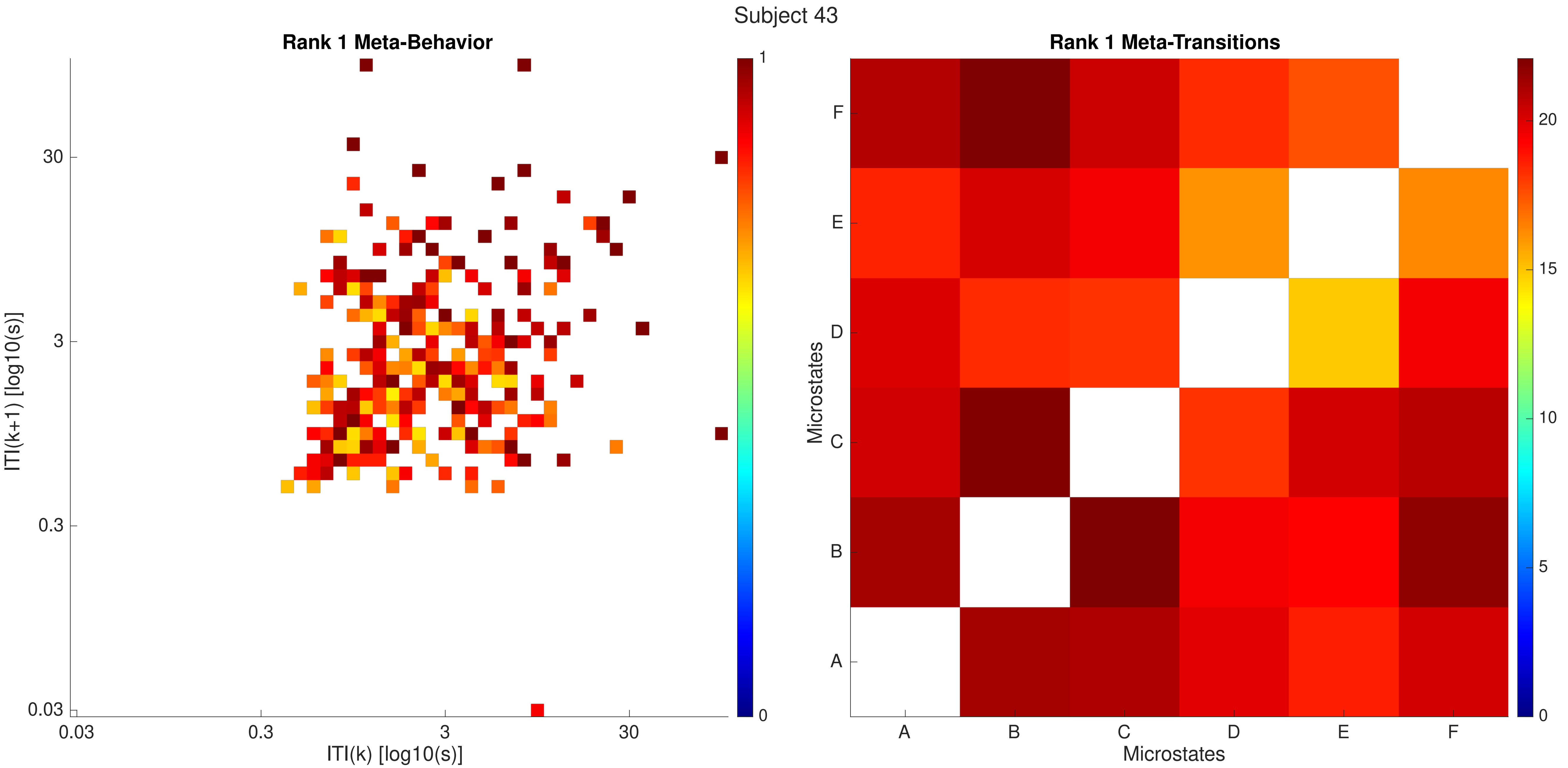

Subject 43
Rank 1 Meta-Behavior
ITI(k+1) [log10(s)]
ITI(k) [log10(s)]
0.03
0.3
3
30
1
0
Rank 1 Meta-Transitions
Microstates
Microstates
A
B
C
D
E
F
0
5
10
15
20

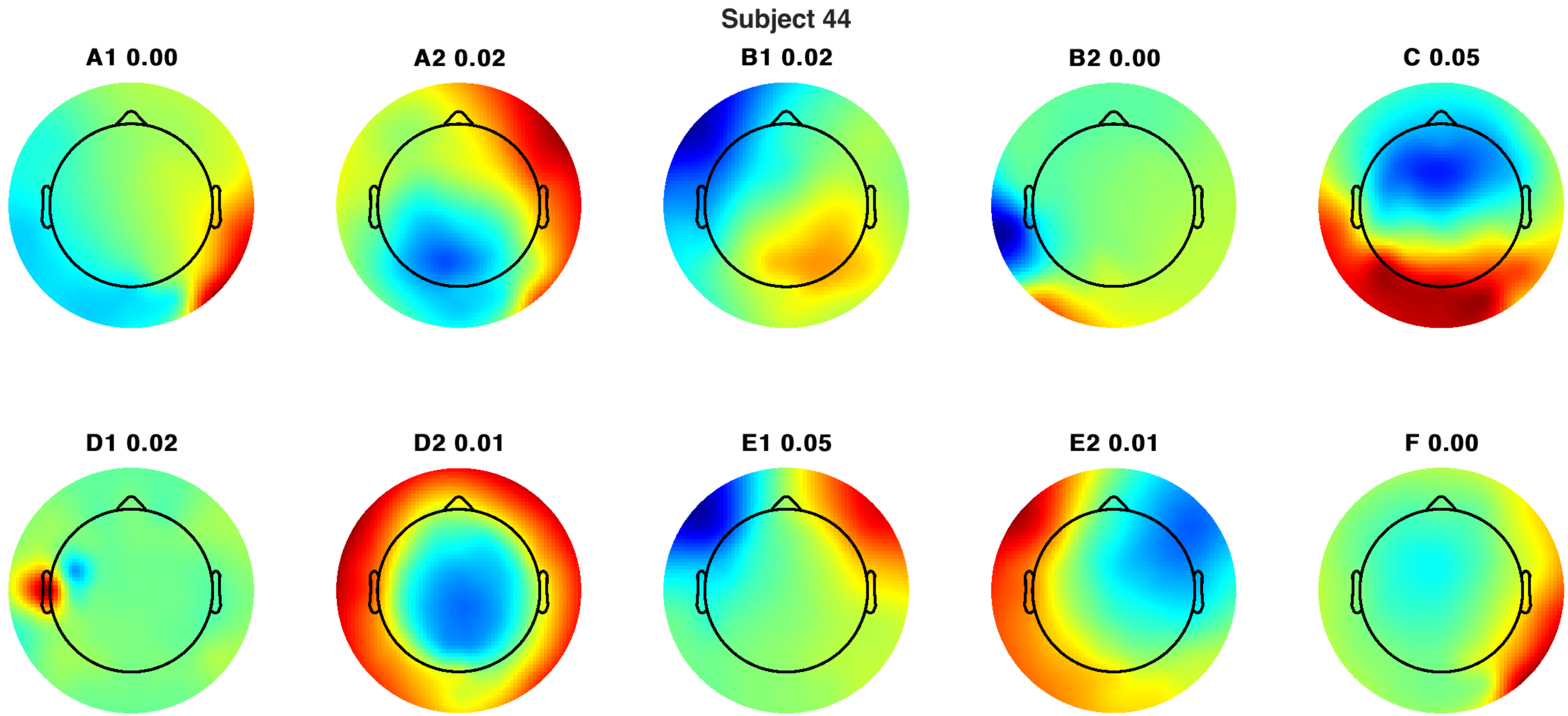
Subject 44
A1 0.00
A2 0.02
B1 0.02
B2 0.00
C 0.05
D1 0.02
D2 0.01
E1 0.05
E2 0.01
F 0.00

Subject 44

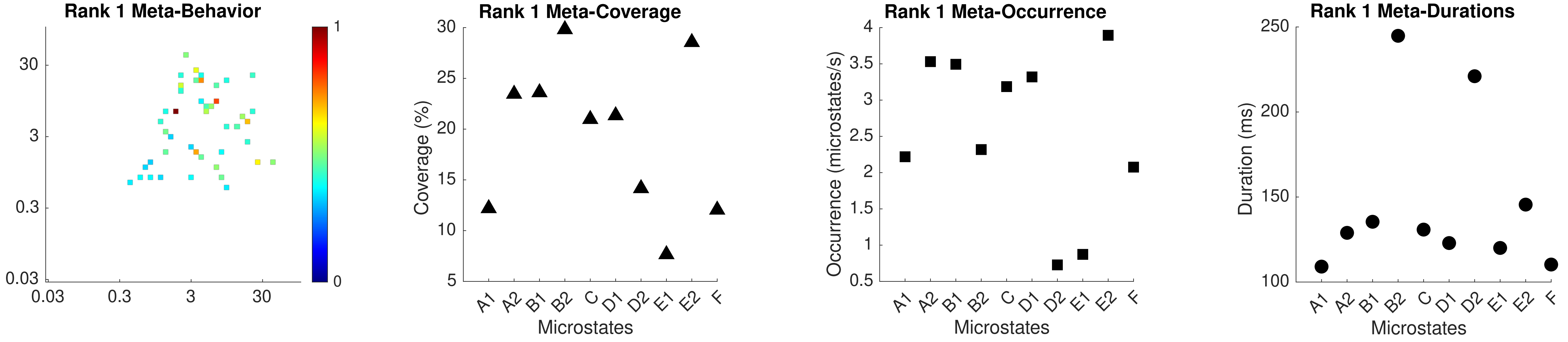

Rank 1 Meta-Behavior
30
3
0.3
0.03
0.03
0.3
3
30
1
0
Rank 1 Meta-Coverage
Coverage (%)
30
25
20
15
10
5
A1
A2
B1
B2
C
D1
D2
E1
E2
F
Microstates
Rank 1 Meta-Occurrence
Occurrence (microstates/s)
4
3.5
3
2.5
2
1.5
1
0.5
A1
A2
B1
B2
C
D1
D2
E1
E2
F
Microstates
Rank 1 Meta-Durations
Duration (ms)
250
200
150
100
A1
A2
B1
B2
C
D1
D2
E1
E2
F
Microstates

Subject 44

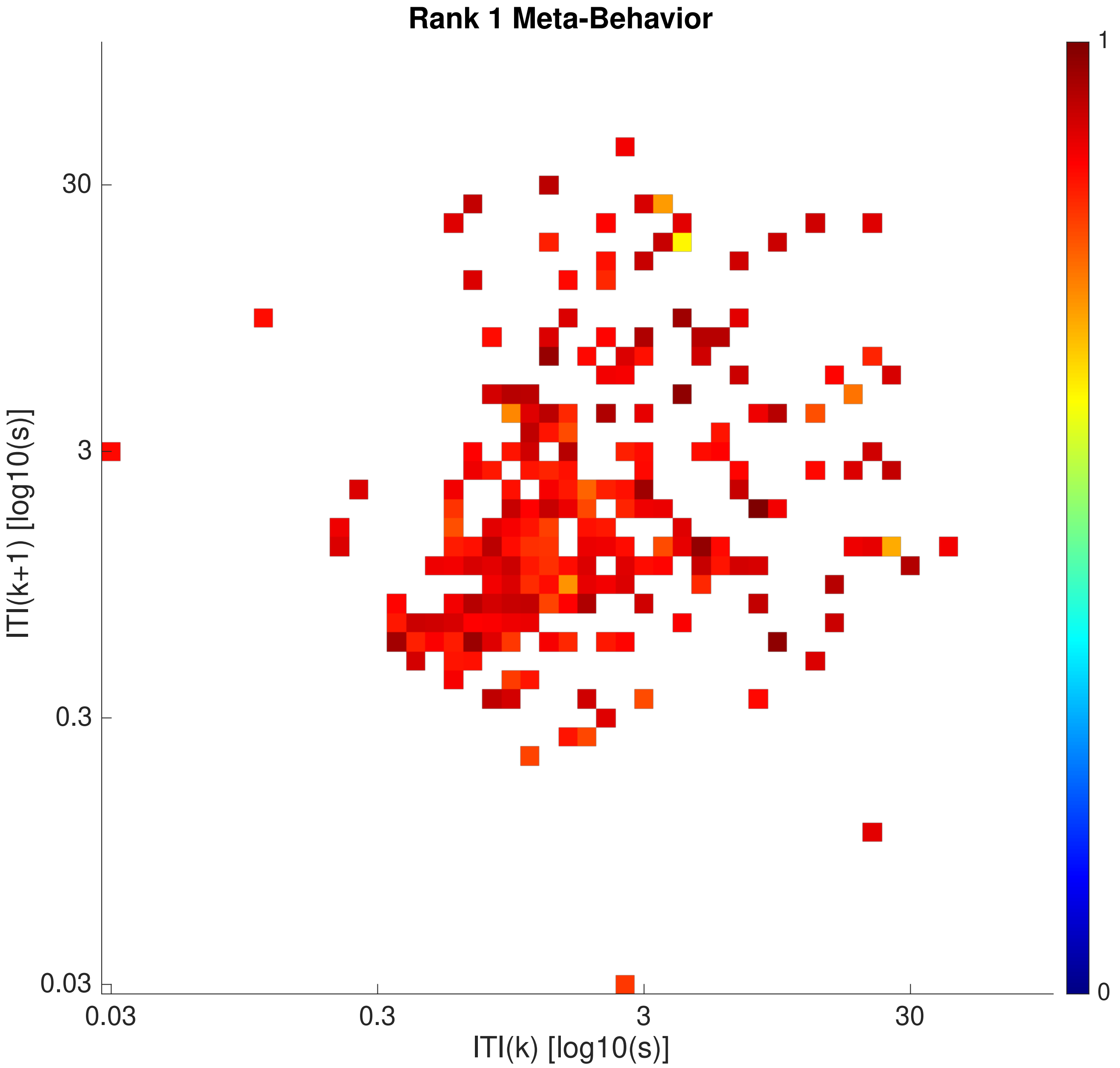

Rank 1 Meta-Behavior
ITI(k+1) [log10(s)]
ITI(k) [log10(s)]
0.03
0.3
3
30
0
1


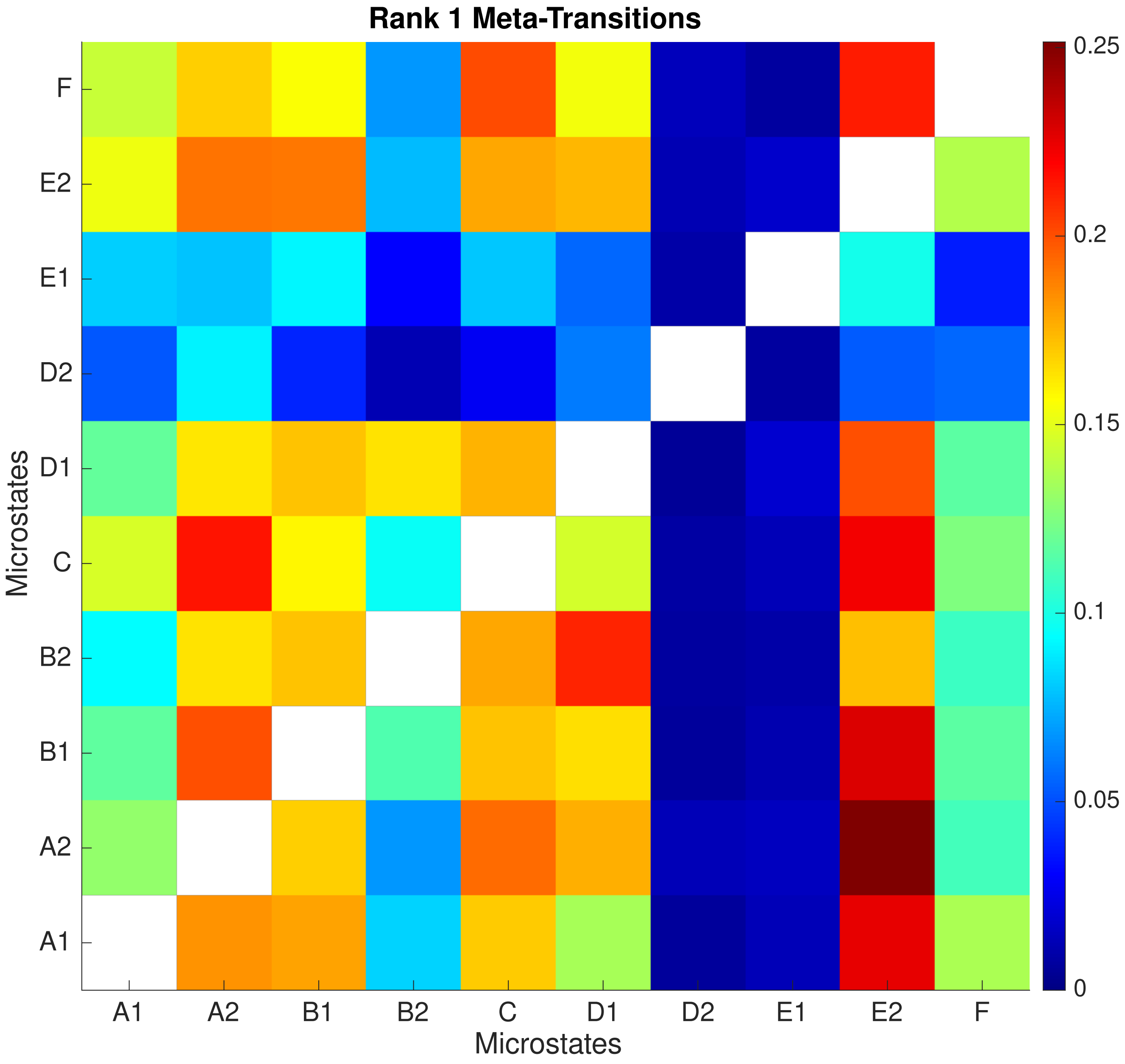

Rank 1 Meta-Transitions
Microstates
A1
A2
B1
B2
C
D1
D2
E1
E2
F
0
0.05
0.1
0.15
0.2
0.25

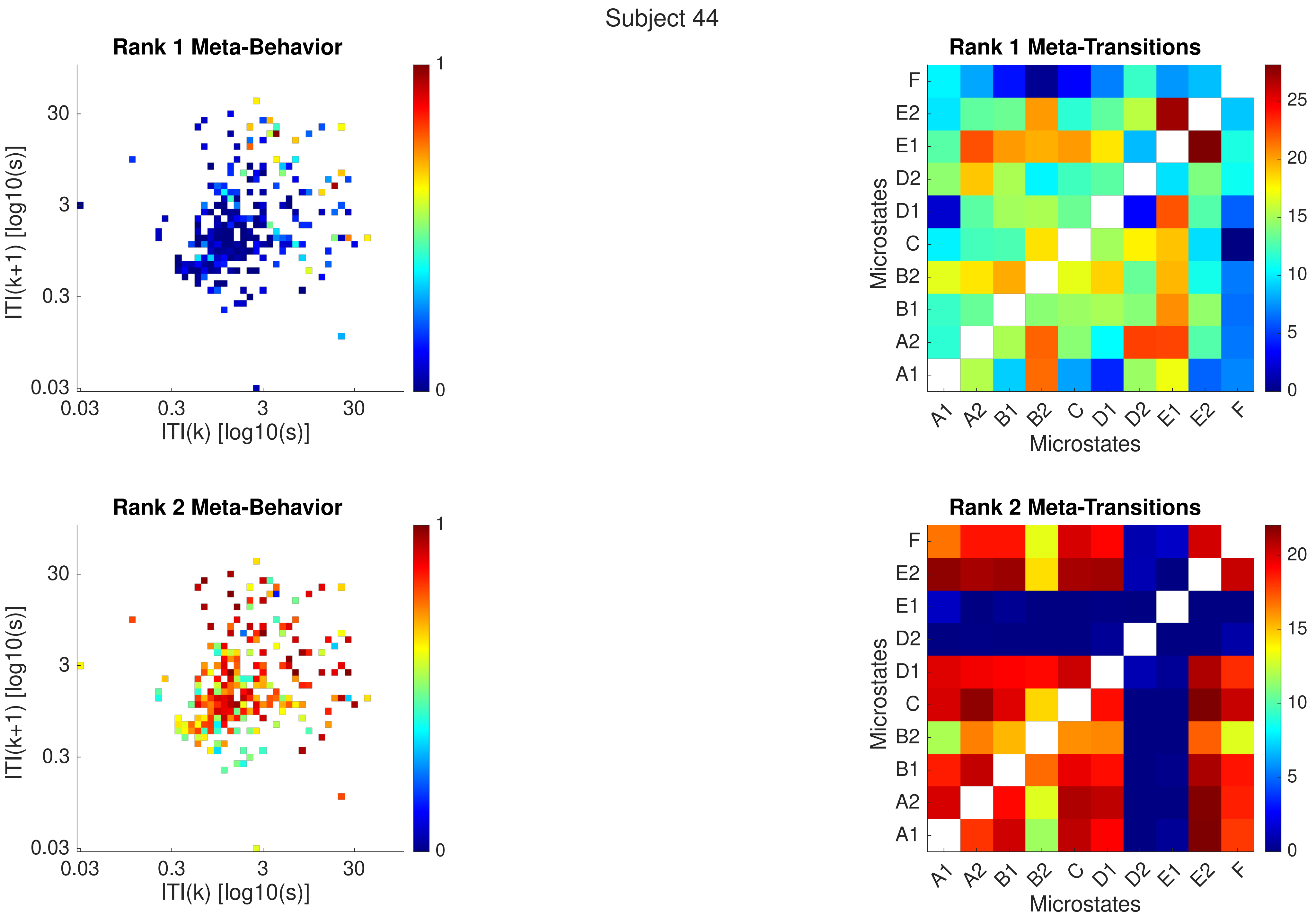
Subject 44
Rank 1 Meta-Behavior
ITI(k+1) [log10(s)]
ITI(k) [log10(s)]
0.03
0.3
3
30
0
1
Rank 1 Meta-Transitions
Microstates
F
E2
E1
D2
D1
C
B2
B1
A2
A1
0
5
10
15
20
25
Rank 2 Meta-Behavior
Rank 2 Meta-Transitions

Subject 44

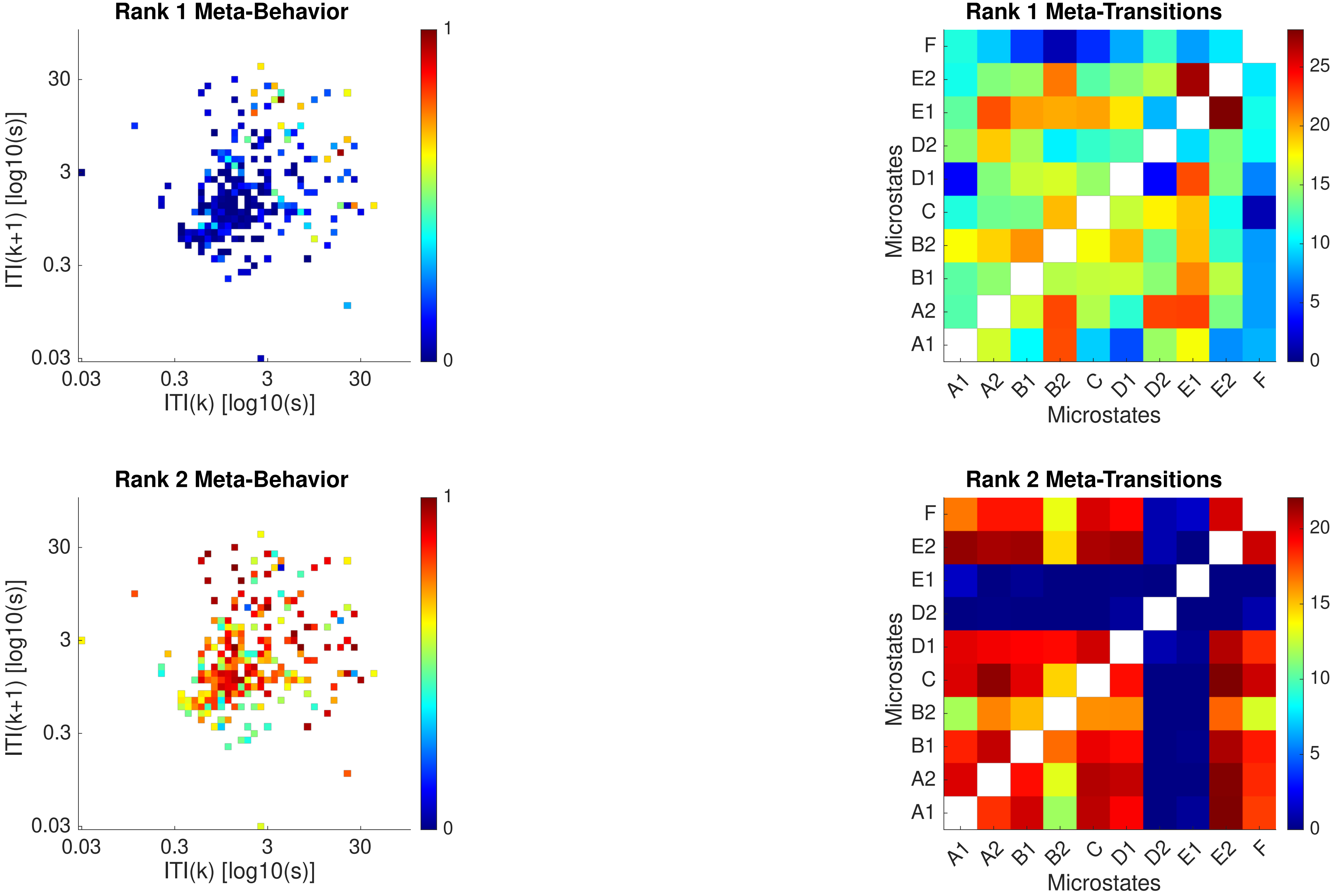

Rank 1 Meta-Behavior
ITI(k+1) [log10(s)]
ITI(k) [log10(s)]
0.03
0.3
3
30
Rank 1 Meta-Transitions
Microstates
A1
A2
B1
B2
C
D1
D2
E1
E2
F
Rank 2 Meta-Behavior
Rank 2 Meta-Transitions

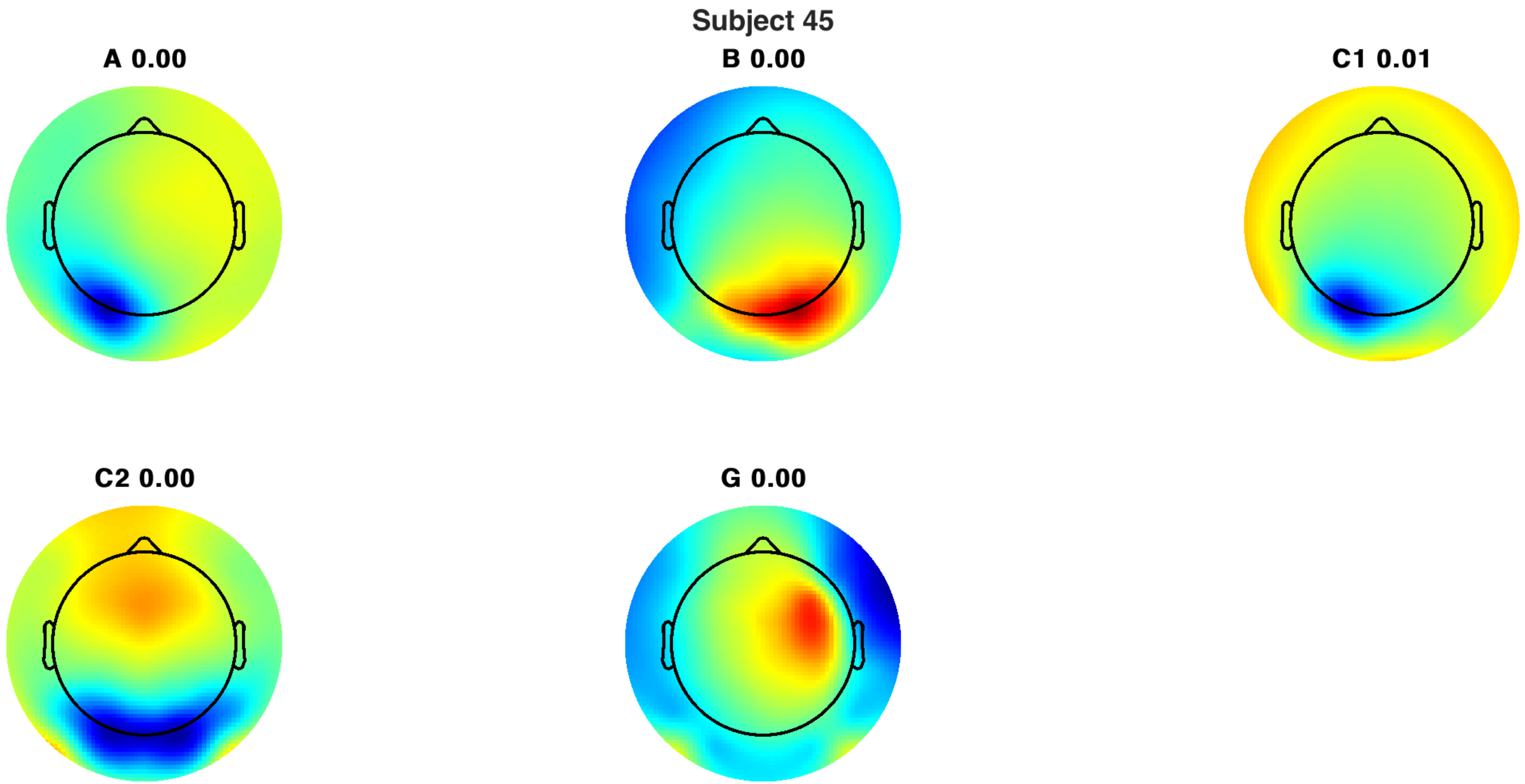
Subject 45
A 0.00
B 0.00
C1 0.01
C2 0.00
G 0.00

Subject 45

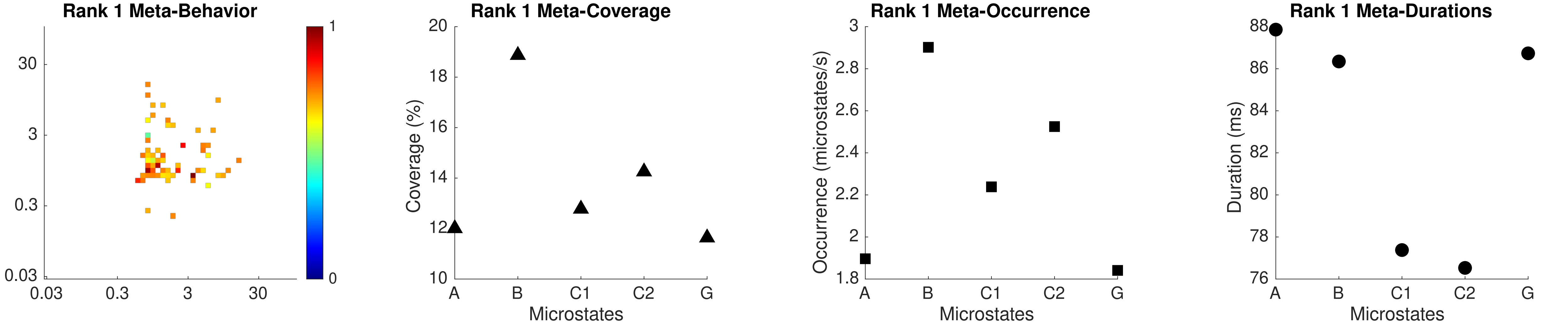

Rank 1 Meta-Behavior
30
3
0.3
0.03
0.03
0.3
3
30
1
0
Rank 1 Meta-Coverage
Coverage (%)
20
18
16
14
12
10
A
B
C1
C2
G
Microstates
Rank 1 Meta-Occurrence
Occurrence (microstates/s)
3
2.8
2.6
2.4
2.2
2
1.8
A
B
C1
C2
G
Microstates
Rank 1 Meta-Durations
Duration (ms)
88
86
84
82
80
78
76
A
B
C1
C2
G
Microstates

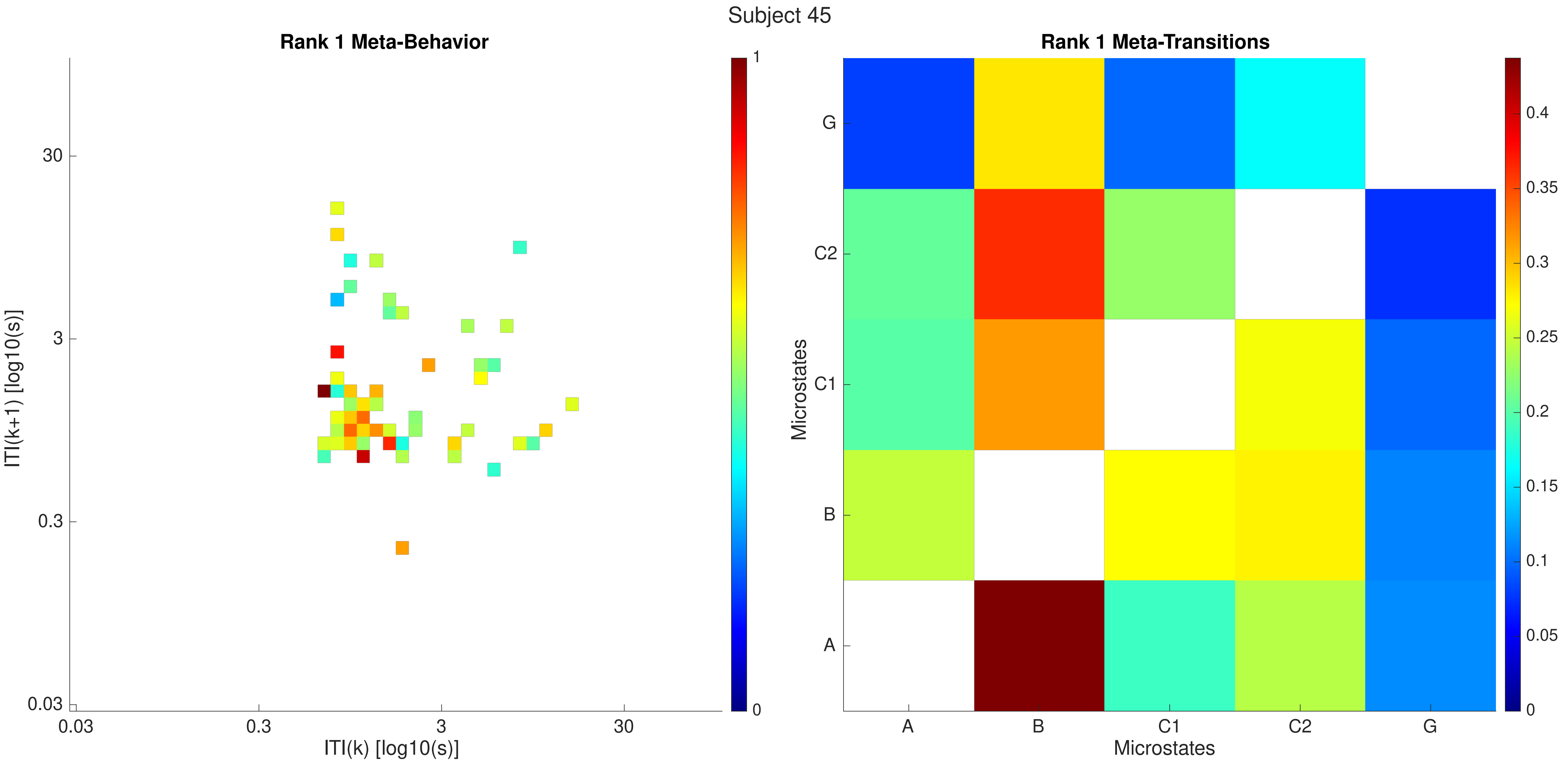

Subject 45
Rank 1 Meta-Behavior
ITI(k+1) [log10(s)]
ITI(k) [log10(s)]
30
3
0.3
0.03
0.03
0.3
3
30
1
0
Rank 1 Meta-Transitions
Microstates
Microstates
G
C2
C1
B
A
A
B
C1
C2
G
0.4
0.35
0.3
0.25
0.2
0.15
0.1
0.05
0

Subject 45

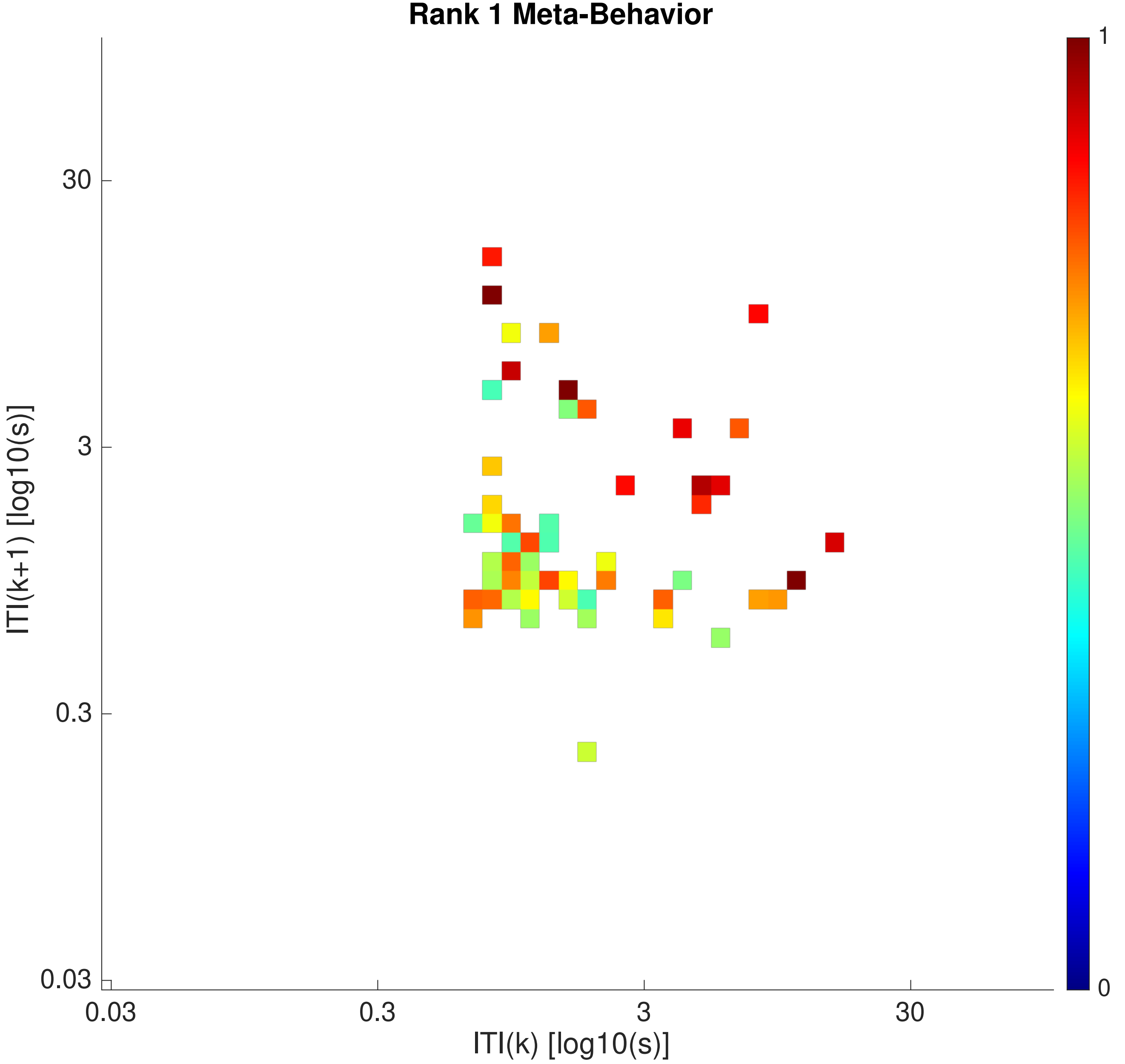

Rank 1 Meta-Behavior
ITI(k+1) [log10(s)]
30
3
0.3
0.03
ITI(k) [log10(s)]
0.03
0.3
3
30
1
0


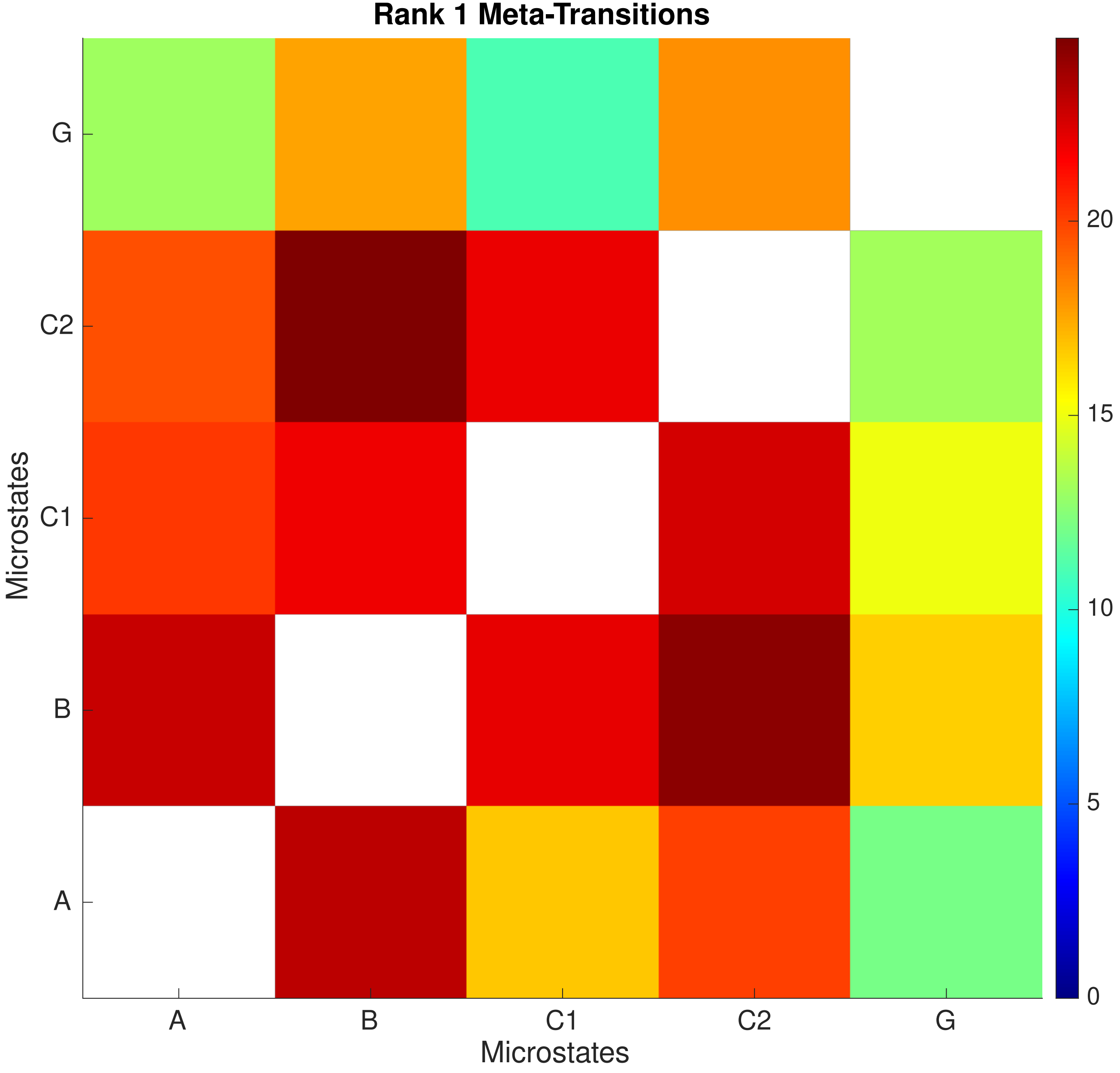

Rank 1 Meta-Transitions
Microstates
G
C2
C1
B
A
Microstates
A
B
C1
C2
G
20
15
10
5
0

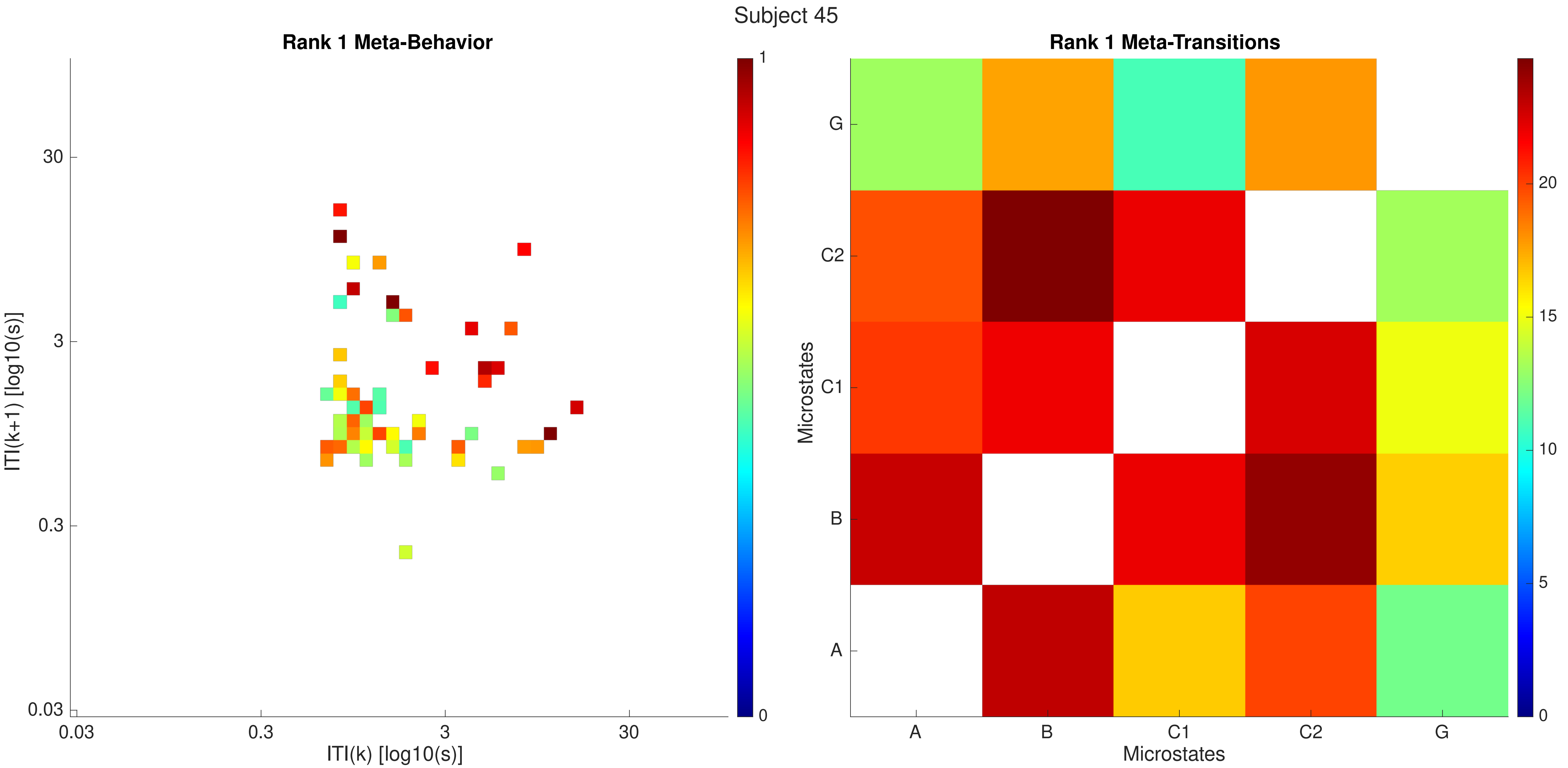
Subject 45
Rank 1 Meta-Behavior
ITI(k+1) [log10(s)]
30
3
0.3
0.03
0.03
0.3
3
30
ITI(k) [log10(s)]
1
0
Rank 1 Meta-Transitions
Microstates
G
C2
C1
B
A
A
B
C1
C2
G
Microstates
20
15
10
5
0

Subject 46

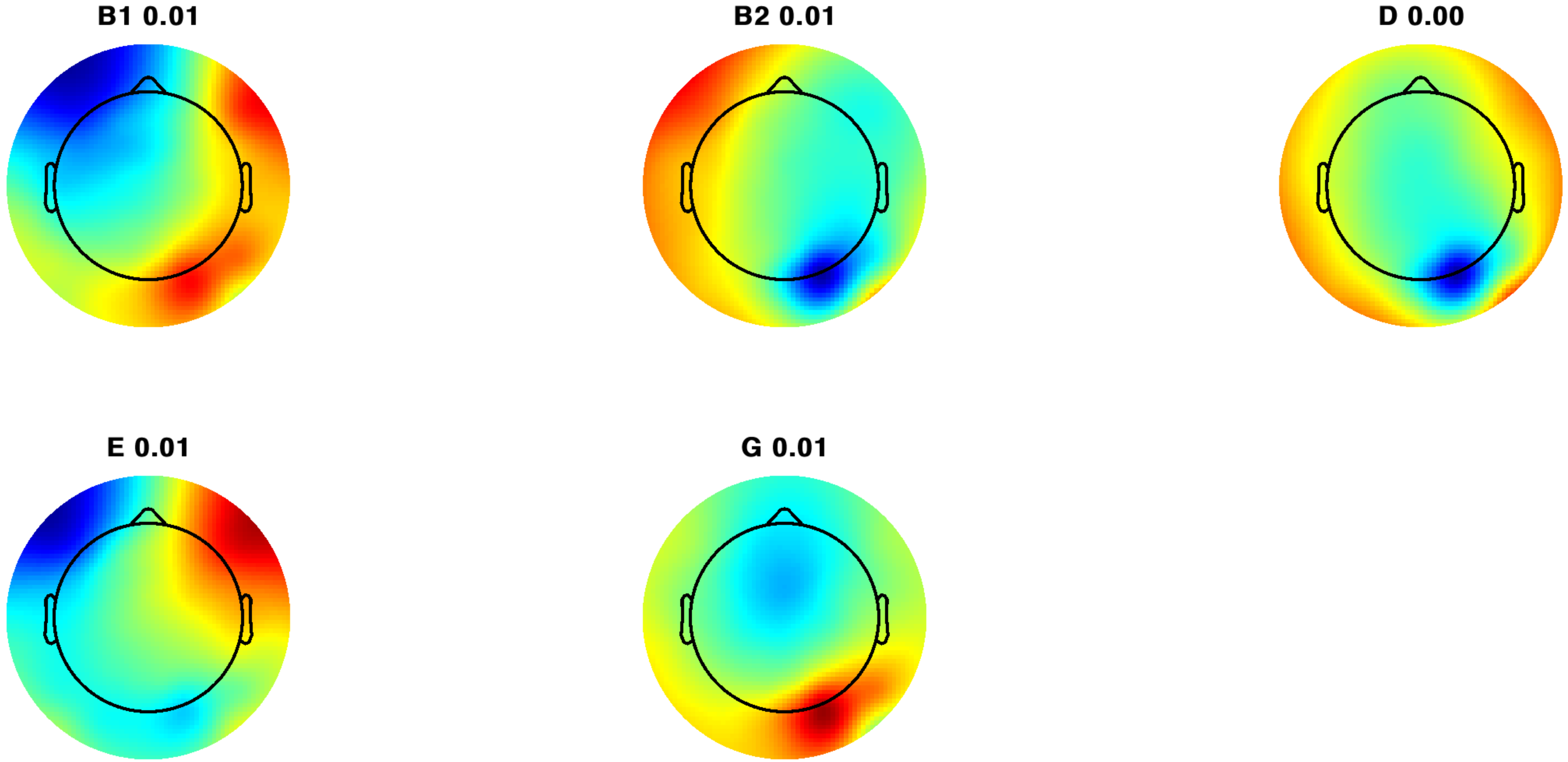

Subject 46

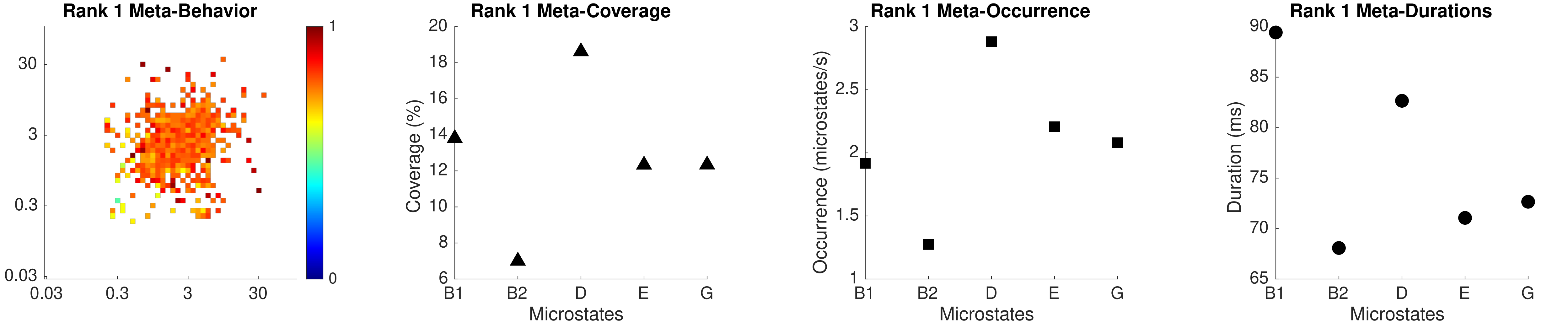

Rank 1 Meta-Behavior
30
3
0.3
0.03
0.03
0.3
3
30
1
0
Rank 1 Meta-Coverage
Coverage (%)
20
18
16
14
12
10
8
6
B1
B2
D
E
G
Microstates
Rank 1 Meta-Occurrence
Occurrence (microstates/s)
3
2.5
2
1.5
1
B1
B2
D
E
G
Microstates
Rank 1 Meta-Durations
Duration (ms)
90
85
80
75
70
65
B1
B2
D
E
G
Microstates

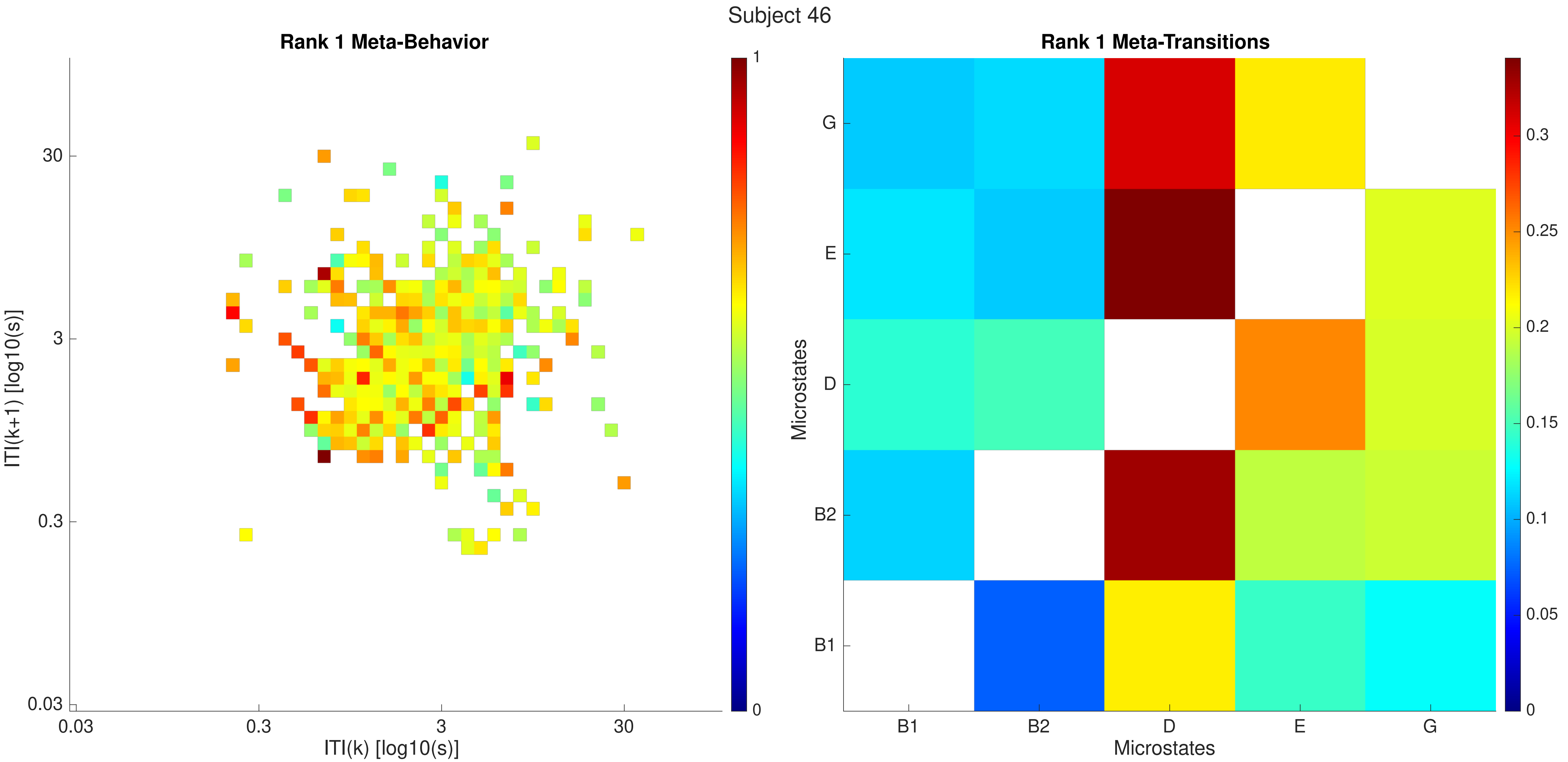

Subject 46
Rank 1 Meta-Behavior
ITI(k+1) [log10(s)]
ITI(k) [log10(s)]
0.03
0.3
3
30
1
0
Rank 1 Meta-Transitions
Microstates
B1
B2
D
E
G
0
0.05
0.1
0.15
0.2
0.25
0.3

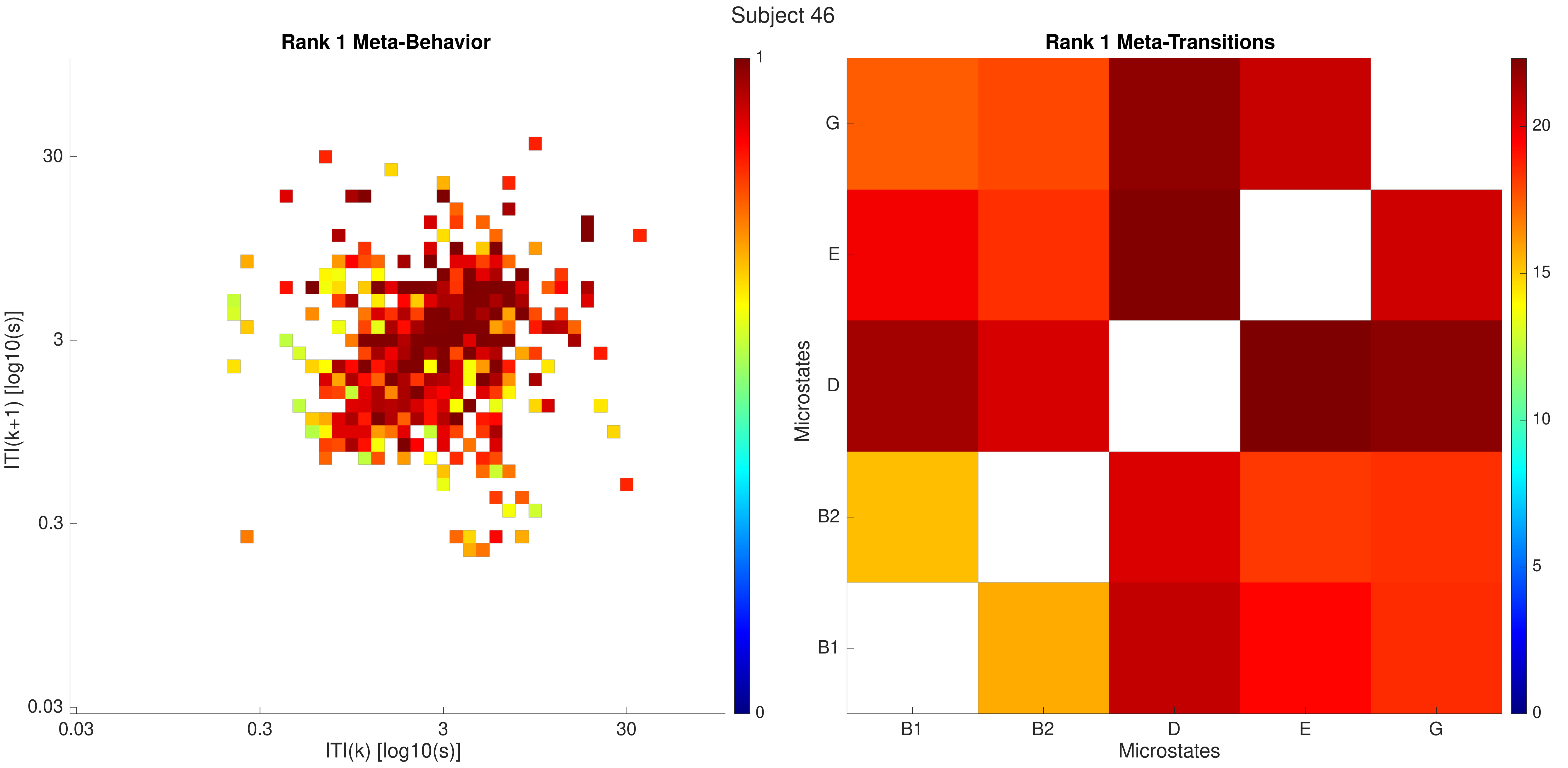

Subject 46
Rank 1 Meta-Behavior
ITI(k+1) [log10(s)]
ITI(k) [log10(s)]
0.03
0.3
3
30
0
1
Rank 1 Meta-Transitions
Microstates
B1
B2
D
E
G
0
5
10
15
20

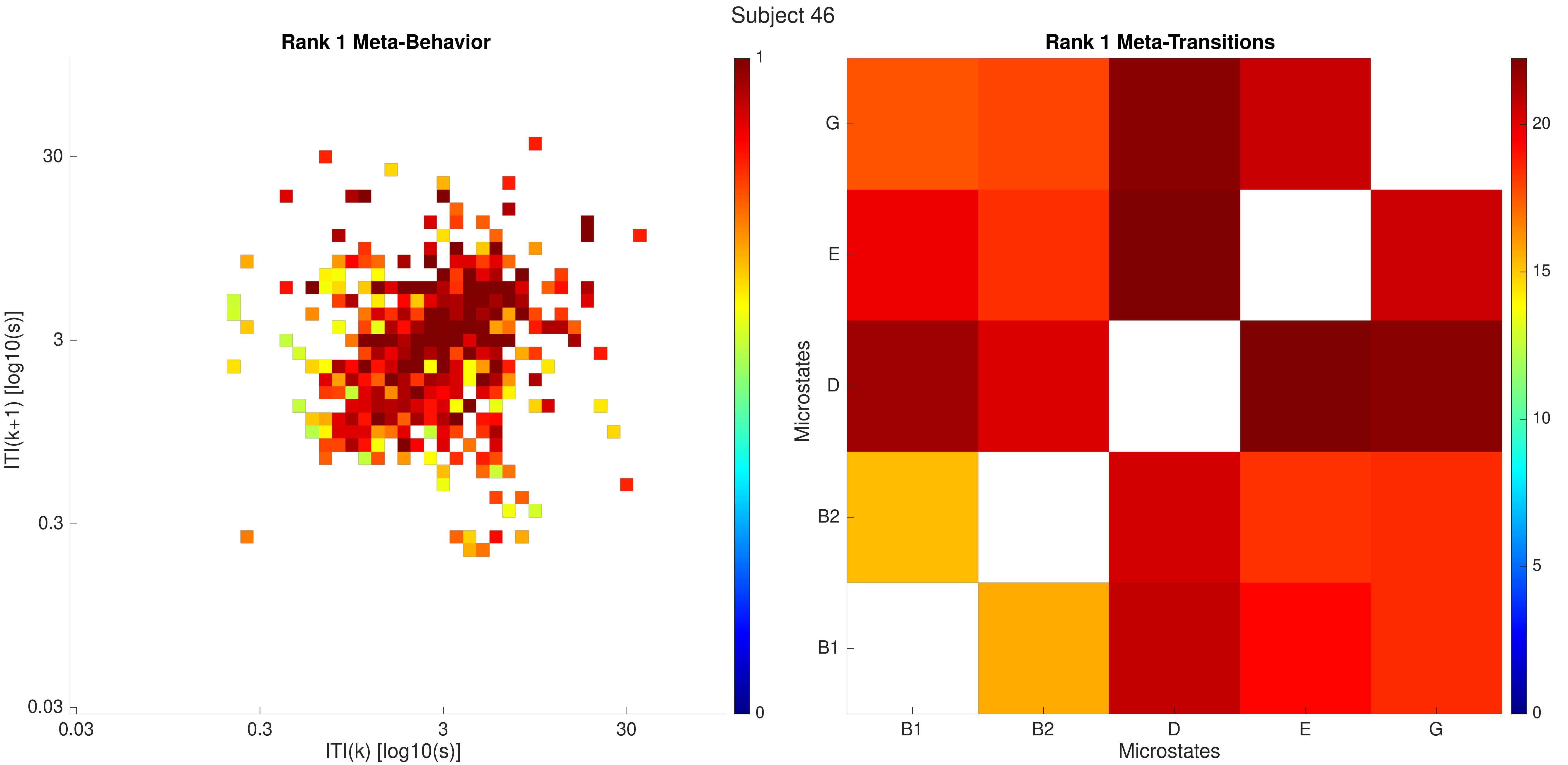
Subject 46
Rank 1 Meta-Behavior
ITI(k+1) [log10(s)]
30
3
0.3
0.03
0.03
0.3
3
30
ITI(k) [log10(s)]
1
0
Rank 1 Meta-Transitions
Microstates
G
E
D
B2
B1
B1
B2
D
E
G
Microstates
20
15
10
5
0

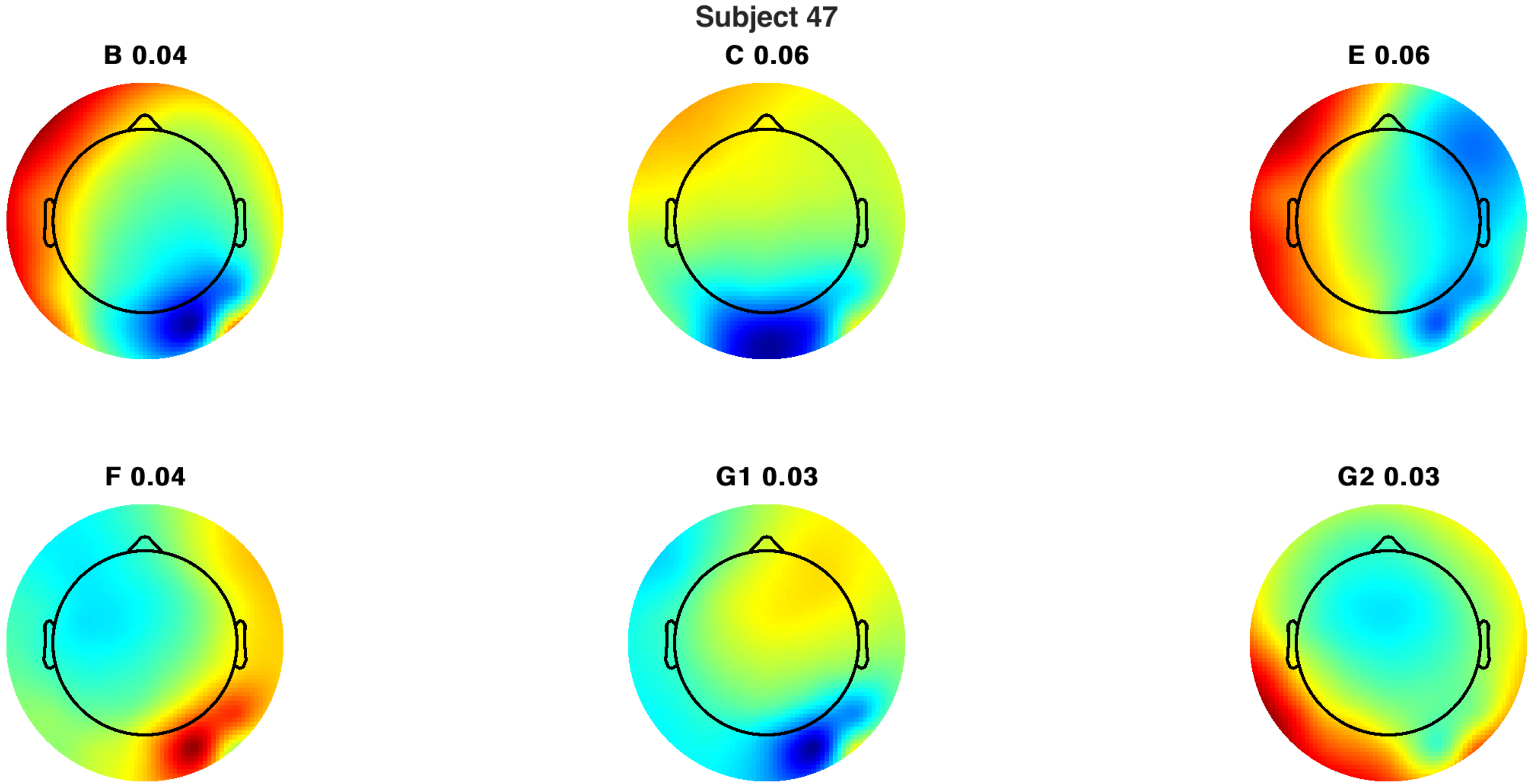
Subject 47
B 0.04
C 0.06
E 0.06
F 0.04
G1 0.03
G2 0.03

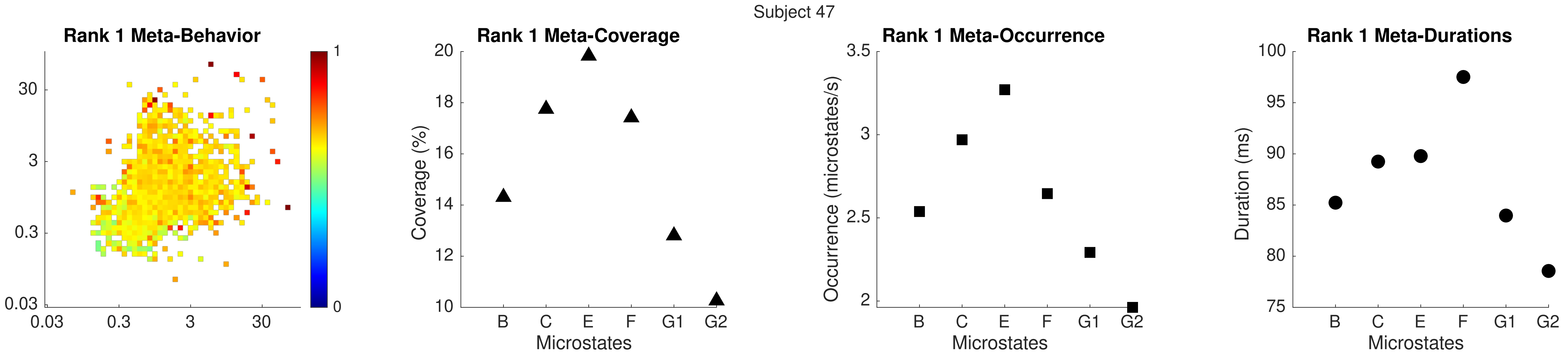

Subject 47
Rank 1 Meta-Behavior
30
3
0.3
0.03
0.03
0.3
3
30
1
0
Rank 1 Meta-Coverage
Coverage (%)
20
18
16
14
12
10
B
C
E
F
G1
G2
Microstates
Rank 1 Meta-Occurrence
Occurrence (microstates/s)
3.5
3
2.5
2
B
C
E
F
G1
G2
Microstates
Rank 1 Meta-Durations
Duration (ms)
100
95
90
85
80
75
B
C
E
F
G1
G2
Microstates

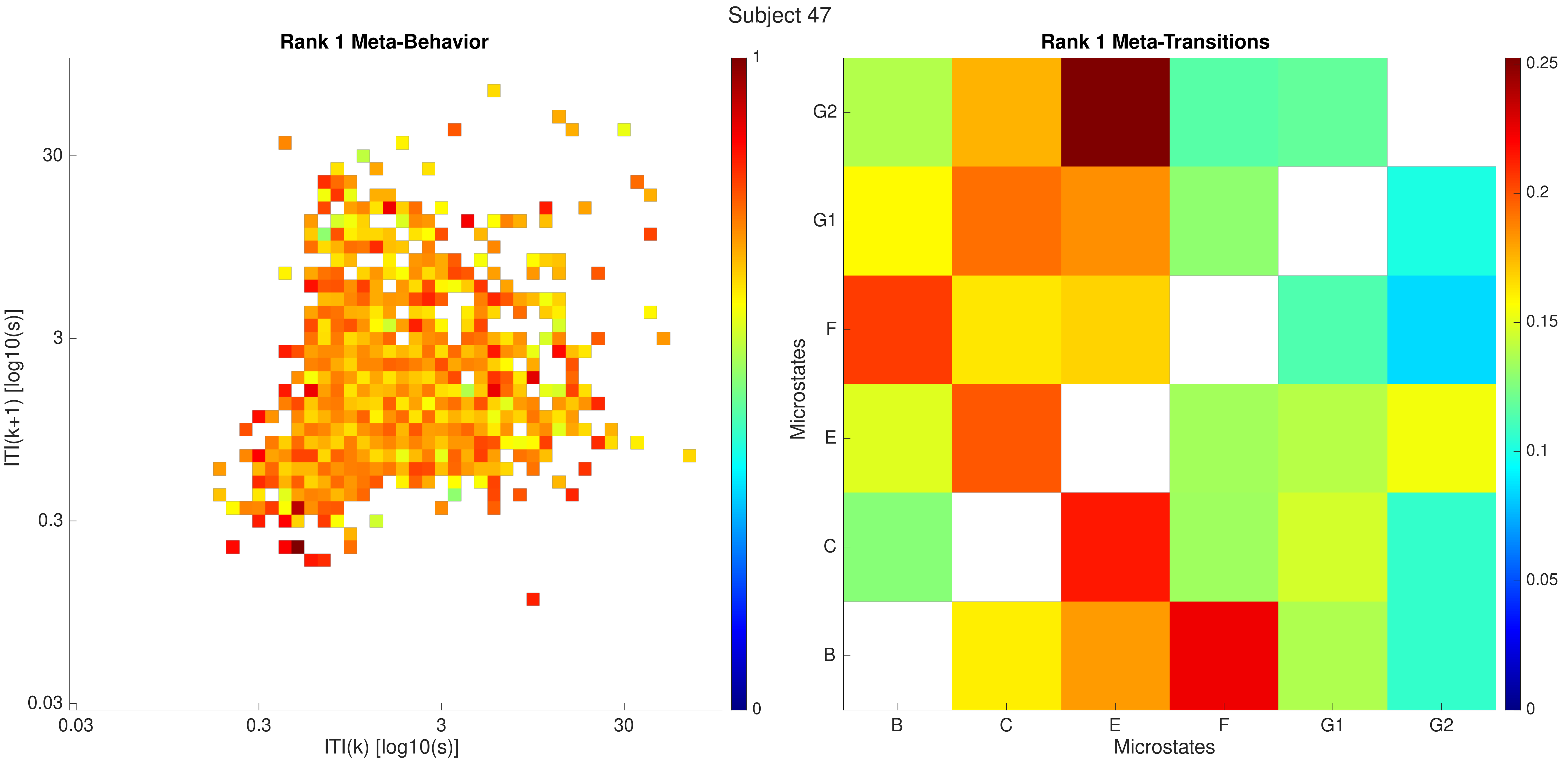
Subject 47
Rank 1 Meta-Behavior
ITI(k+1) [log10(s)]
ITI(k) [log10(s)]
0.03
0.3
3
30
0
1
Rank 1 Meta-Transitions
Microstates
Microstates
B
C
E
F
G1
G2
0
0.05
0.1
0.15
0.2
0.25

Subject 47

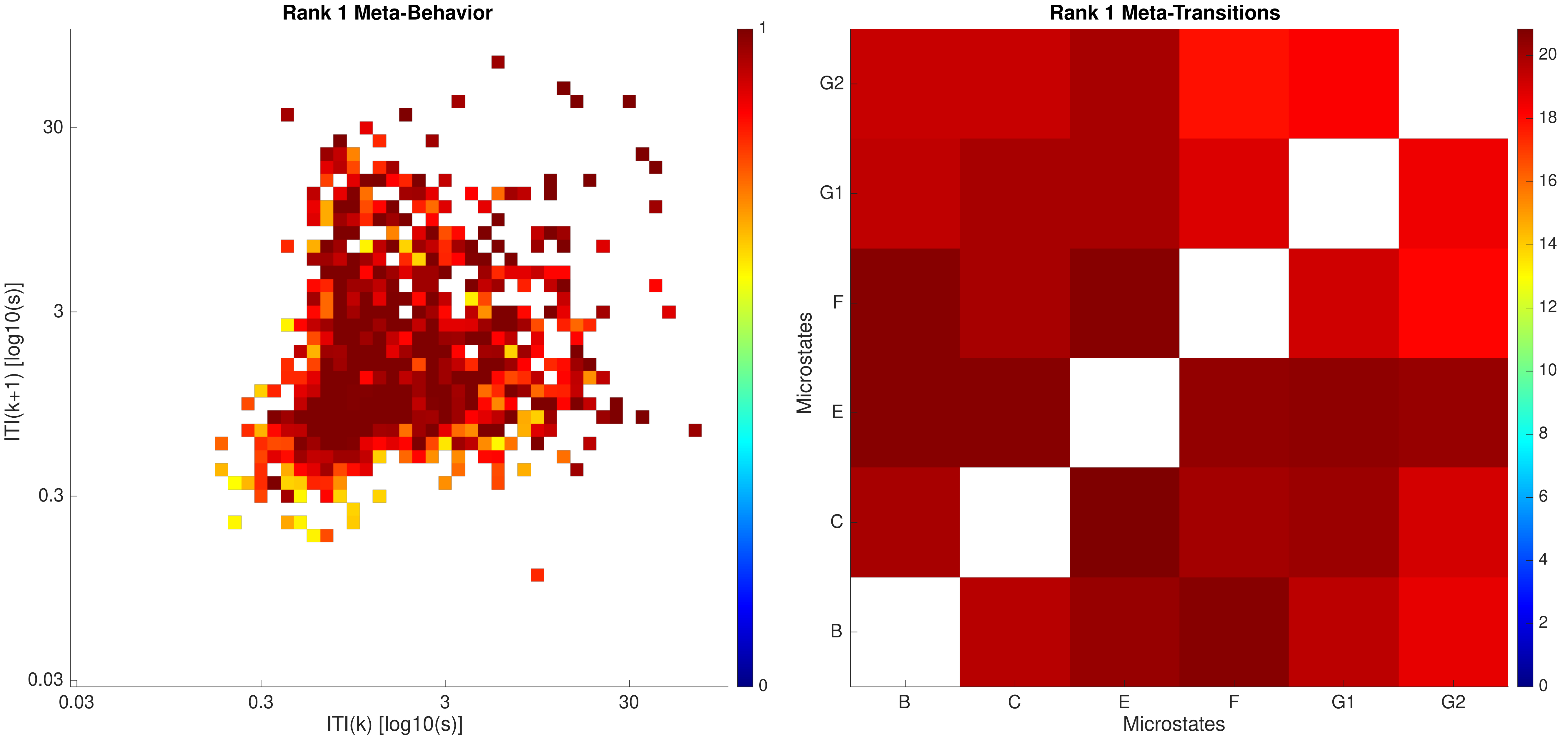

Rank 1 Meta-Behavior
ITI(k+1) [log10(s)]
ITI(k) [log10(s)]
0.03
0.3
3
30
1
0
Rank 1 Meta-Transitions
Microstates
B
C
E
F
G1
G2
0
2
4
6
8
10
12
14
16
18
20

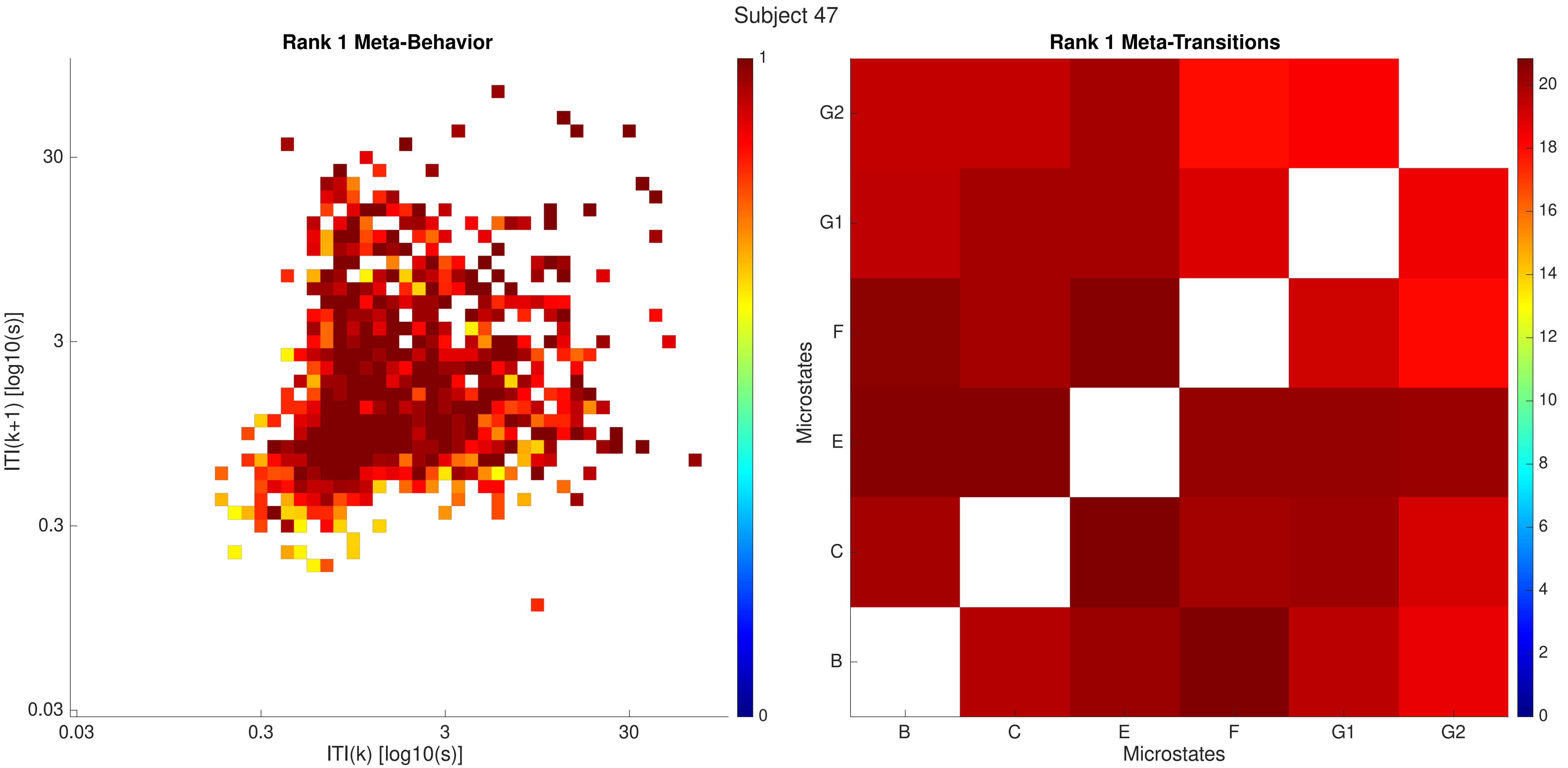
Subject 47
Rank 1 Meta-Behavior
ITI(k+1) [log10(s)]
ITI(k) [log10(s)]
0.03
0.3
3
30
0
1
Rank 1 Meta-Transitions
Microstates
Microstates
B
C
E
F
G1
G2
0
2
4
6
8
10
12
14
16
18
20

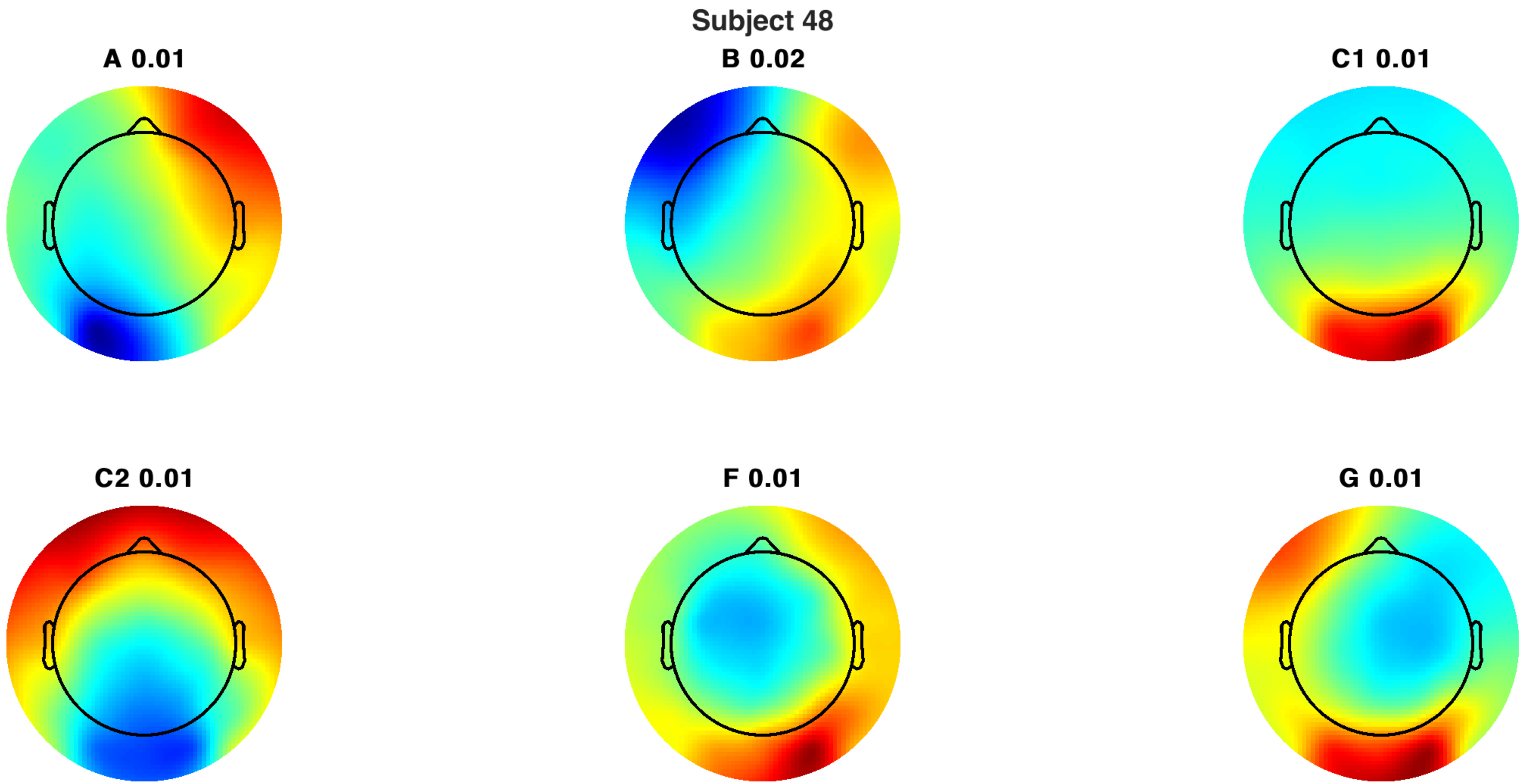
Subject 48
A 0.01
B 0.02
C1 0.01
C2 0.01
F 0.01
G 0.01

Subject 48

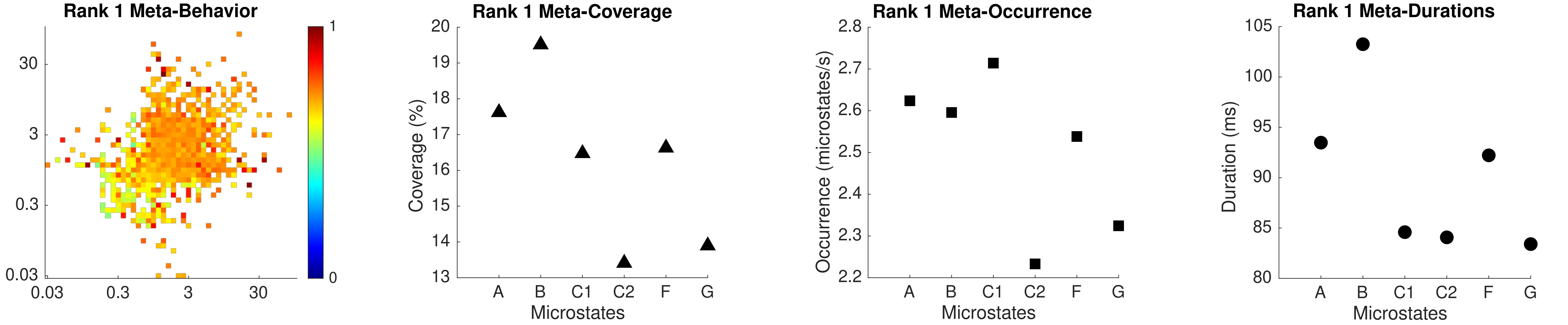

Rank 1 Meta-Behavior
30
3
0.3
0.03
0.03
0.3
3
30
1
0
Rank 1 Meta-Coverage
Coverage (%)
20
19
18
17
16
15
14
13
A
B
C1
C2
F
G
Microstates
Rank 1 Meta-Occurrence
Occurrence (microstates/s)
2.8
2.7
2.6
2.5
2.4
2.3
2.2
A
B
C1
C2
F
G
Microstates
Rank 1 Meta-Durations
Duration (ms)
105
100
95
90
85
80
A
B
C1
C2
F
G
Microstates

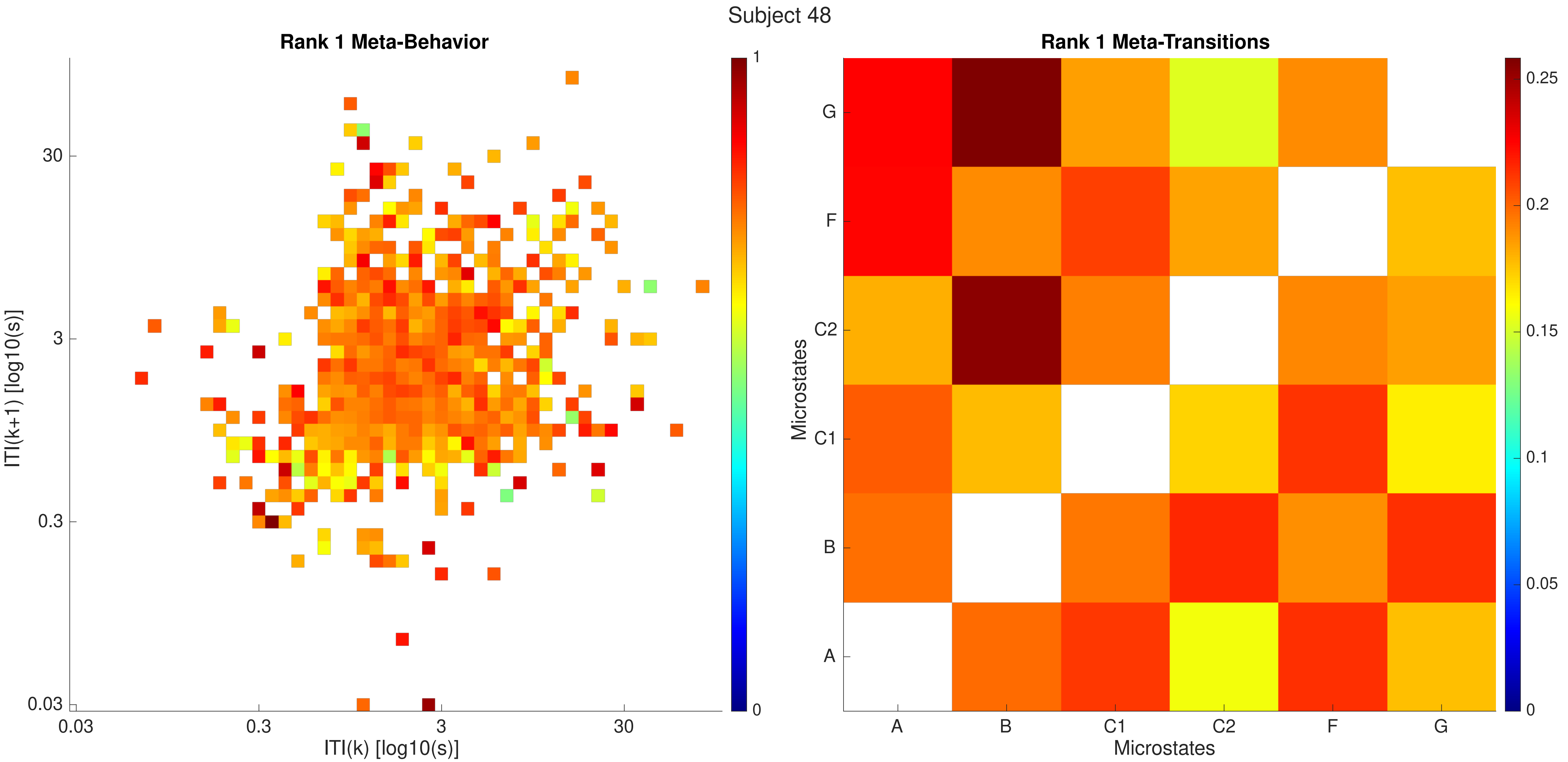

Subject 48
Rank 1 Meta-Behavior
ITI(k+1) [log10(s)]
ITI(k) [log10(s)]
30
3
0.3
0.03
0.03
0.3
3
30
1
0
Rank 1 Meta-Transitions
Microstates
Microstates
G
F
C2
C1
B
A
A
B
C1
C2
F
G
0.25
0.2
0.15
0.1
0.05
0

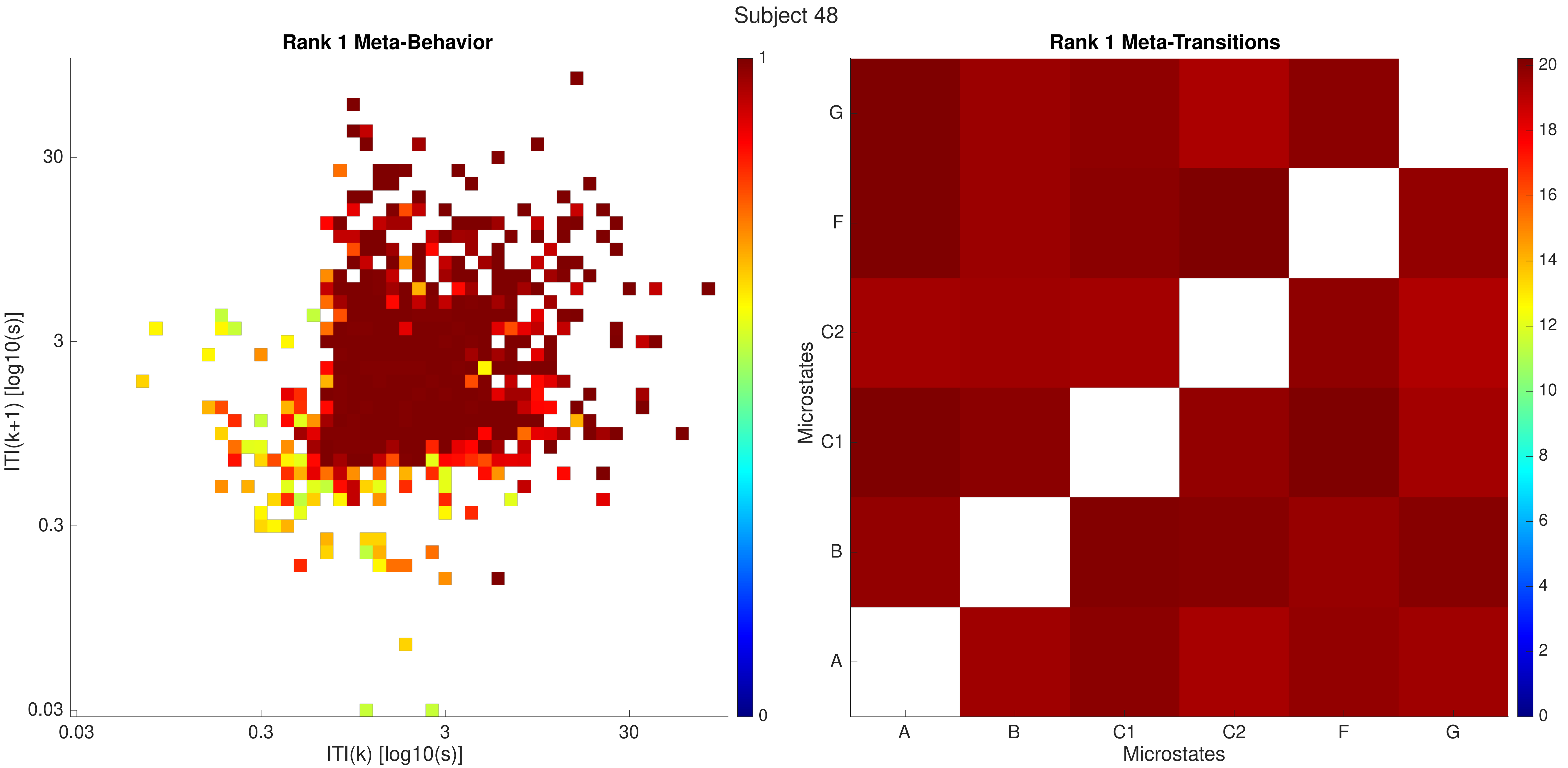

Subject 48
Rank 1 Meta-Behavior
ITI(k+1) [log10(s)]
ITI(k) [log10(s)]
0.03
0.3
3
30
0
1
Rank 1 Meta-Transitions
Microstates
Microstates
A
B
C1
C2
F
G
0
2
4
6
8
10
12
14
16
18
20

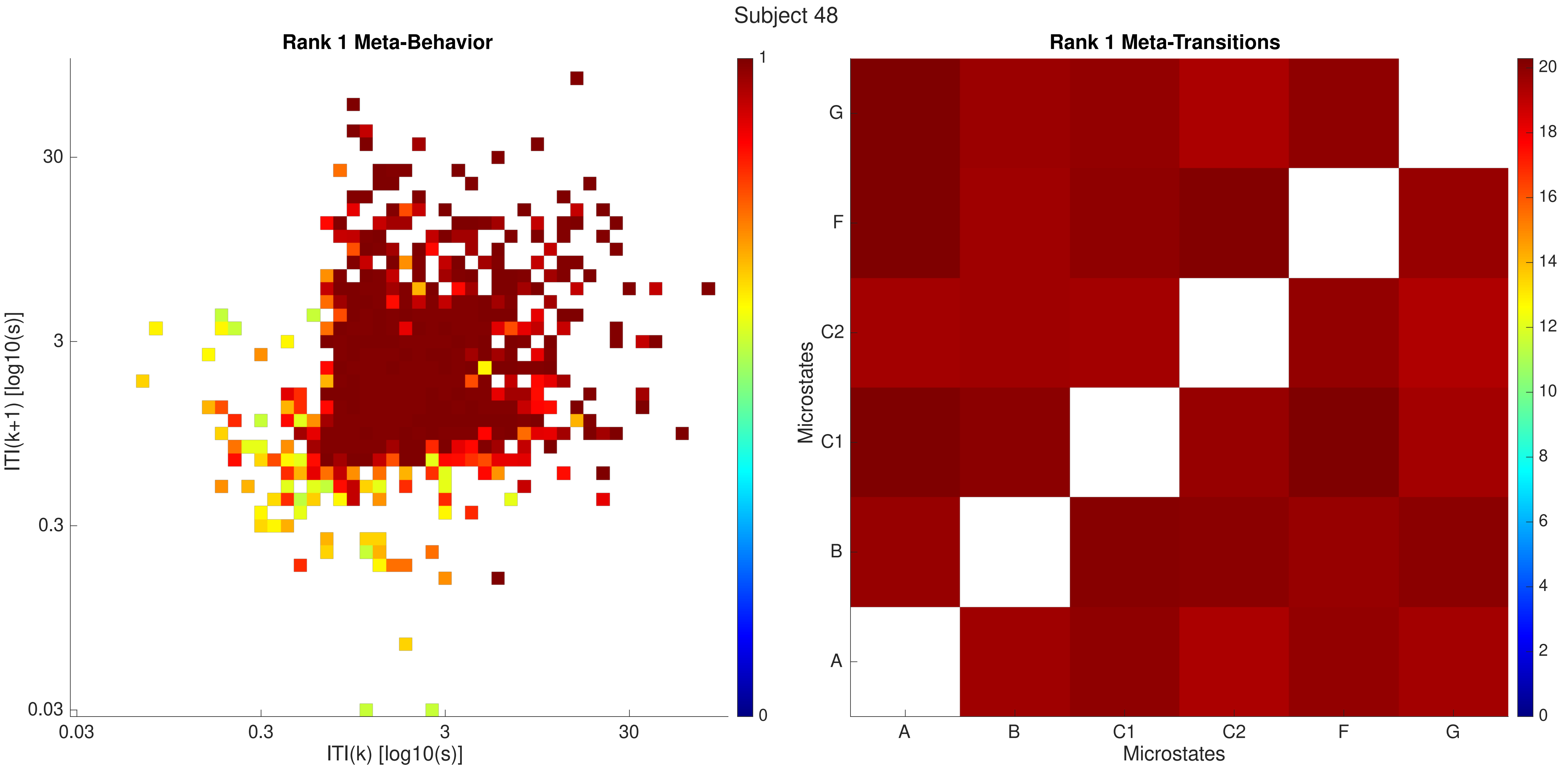
Subject 48
Rank 1 Meta-Behavior
ITI(k+1) [log10(s)]
ITI(k) [log10(s)]
0.03
0.3
3
30
0
1
Rank 1 Meta-Transitions
Microstates
Microstates
A
B
C1
C2
F
G
0
2
4
6
8
10
12
14
16
18
20

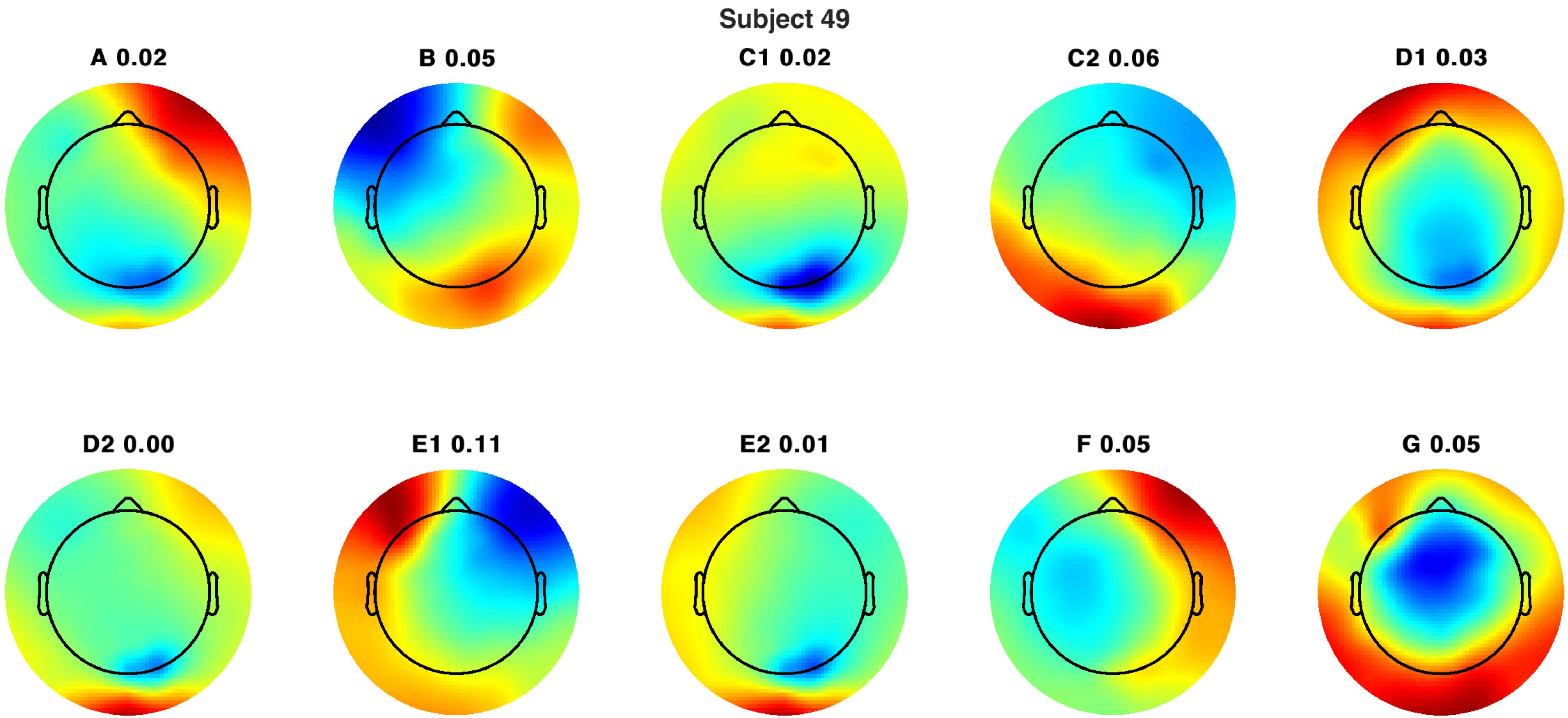
Subject 49
A 0.02
B 0.05
C1 0.02
C2 0.06
D1 0.03
D2 0.00
E1 0.11
E2 0.01
F 0.05
G 0.05

Subject 49

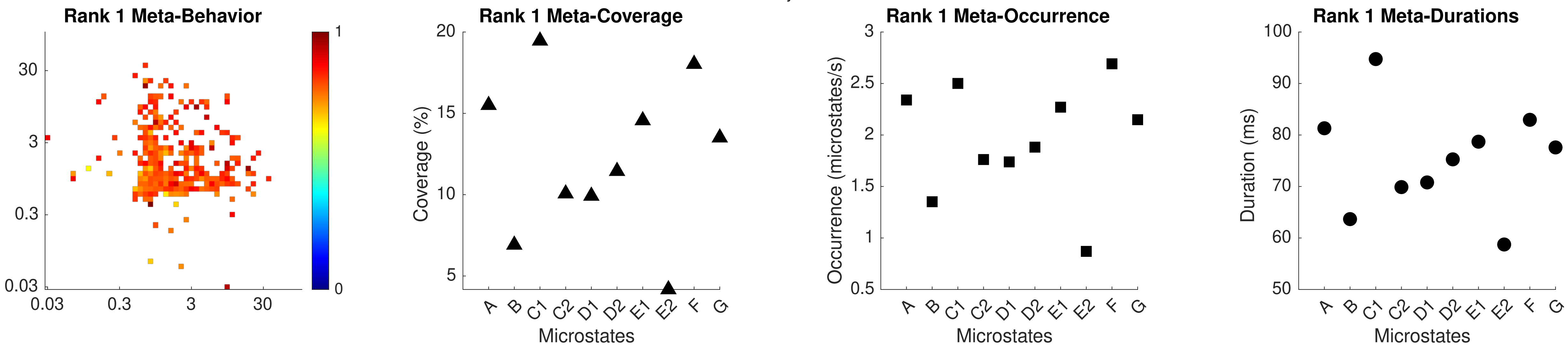

Rank 1 Meta-Behavior
30
3
0.3
0.03
0.03
0.3
3
30
1
0
Rank 1 Meta-Coverage
Coverage (%)
20
15
10
5
A B C1 C2 D1 D2 E1 E2 F G
Microstates
Rank 1 Meta-Occurrence
Occurrence (microstates/s)
3
2.5
2
1.5
1
0.5
A B C1 C2 D1 D2 E1 E2 F G
Microstates
Rank 1 Meta-Durations
Duration (ms)
100
90
80
70
60
50
A B C1 C2 D1 D2 E1 E2 F G
Microstates

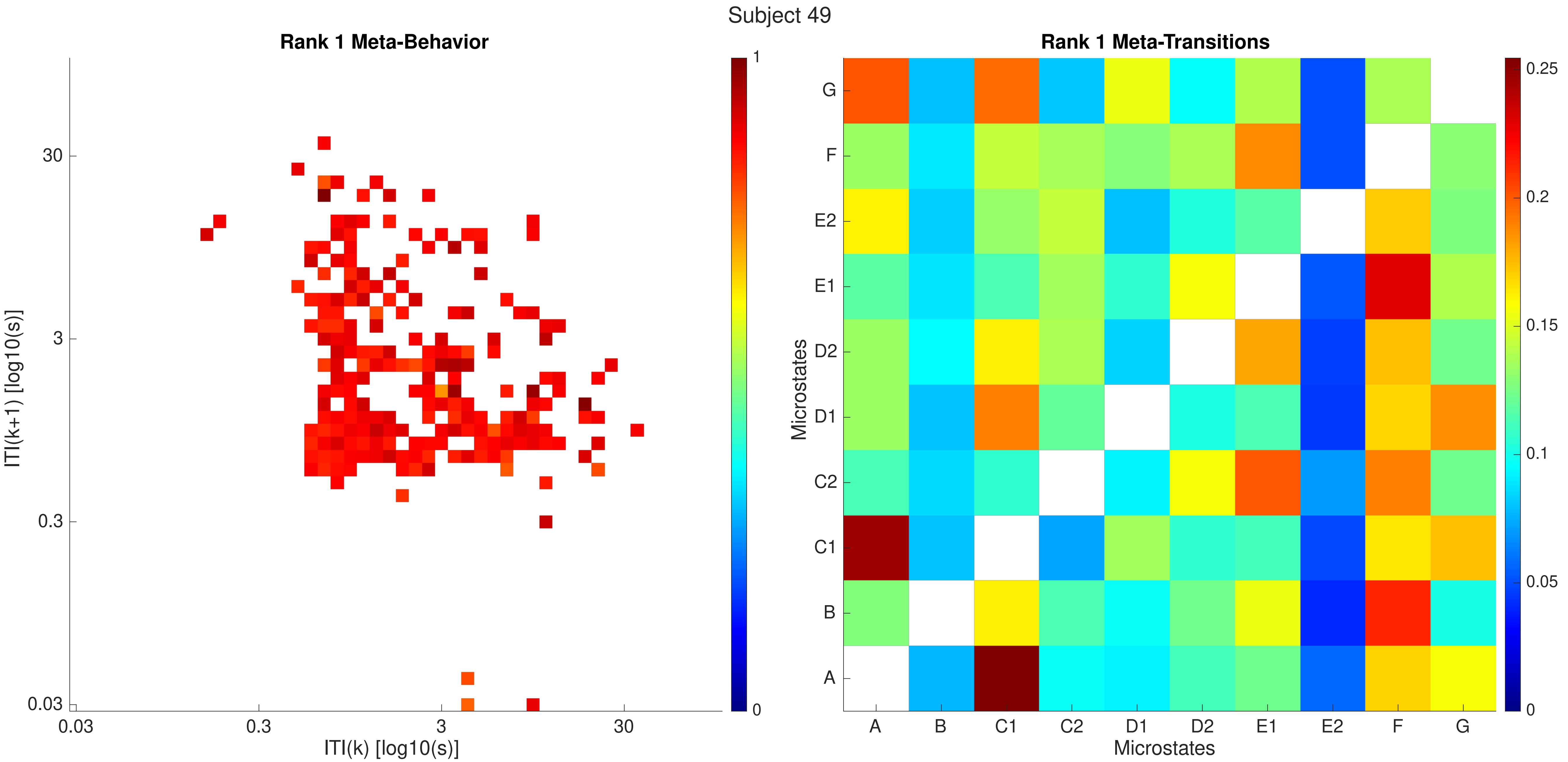
Subject 49
Rank 1 Meta-Behavior
ITI(k+1) [log10(s)]
ITI(k) [log10(s)]
0.03
0.3
3
30
1
0
Rank 1 Meta-Transitions
Microstates
A
B
C1
C2
D1
D2
E1
E2
F
G
0.25
0.2
0.15
0.1
0.05
0

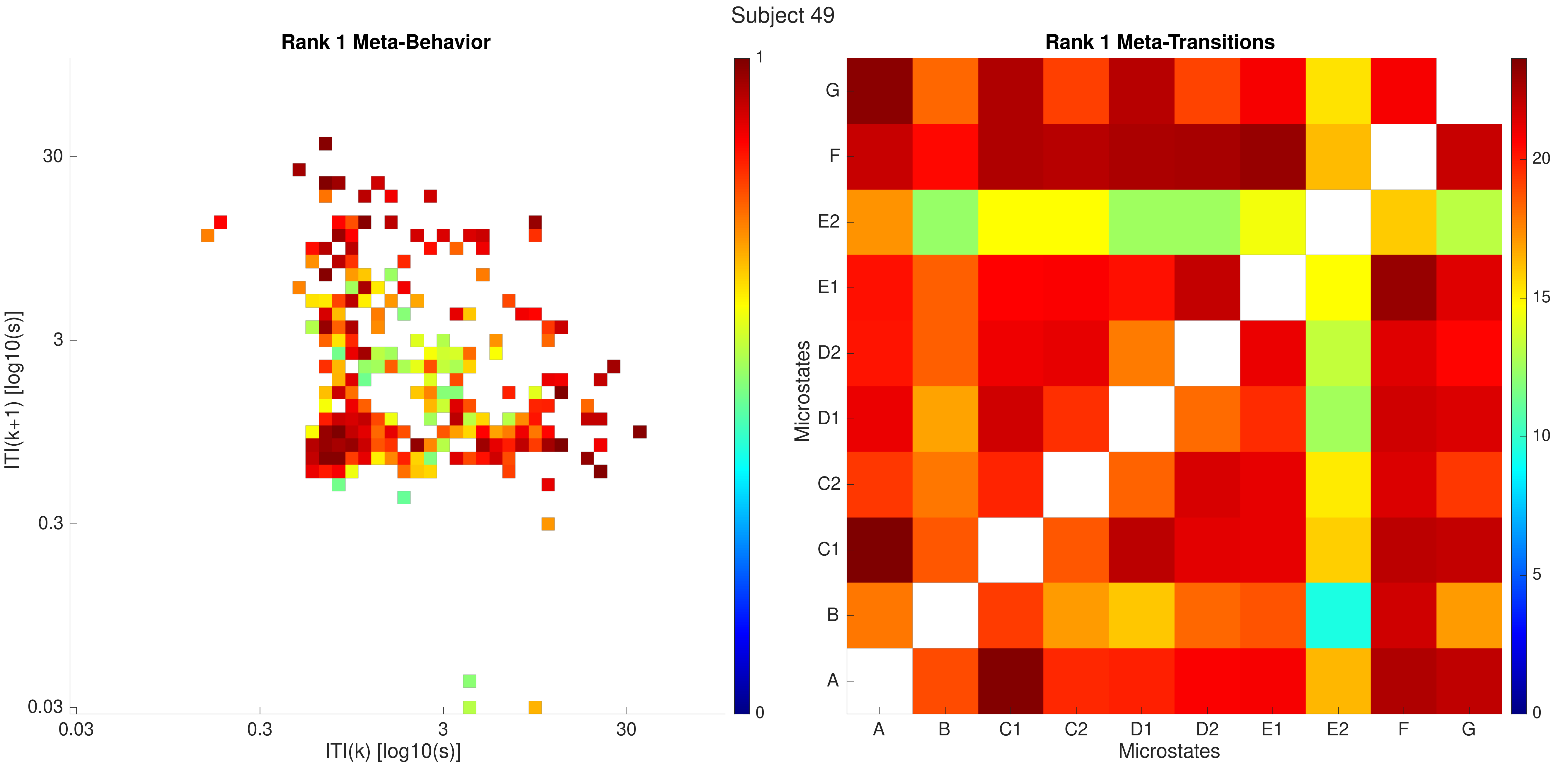

Subject 49
Rank 1 Meta-Behavior
ITI(k+1) [log10(s)]
ITI(k) [log10(s)]
0.03
0.3
3
30
0
1
Rank 1 Meta-Transitions
Microstates
A
B
C1
C2
D1
D2
E1
E2
F
G
0
5
10
15
20

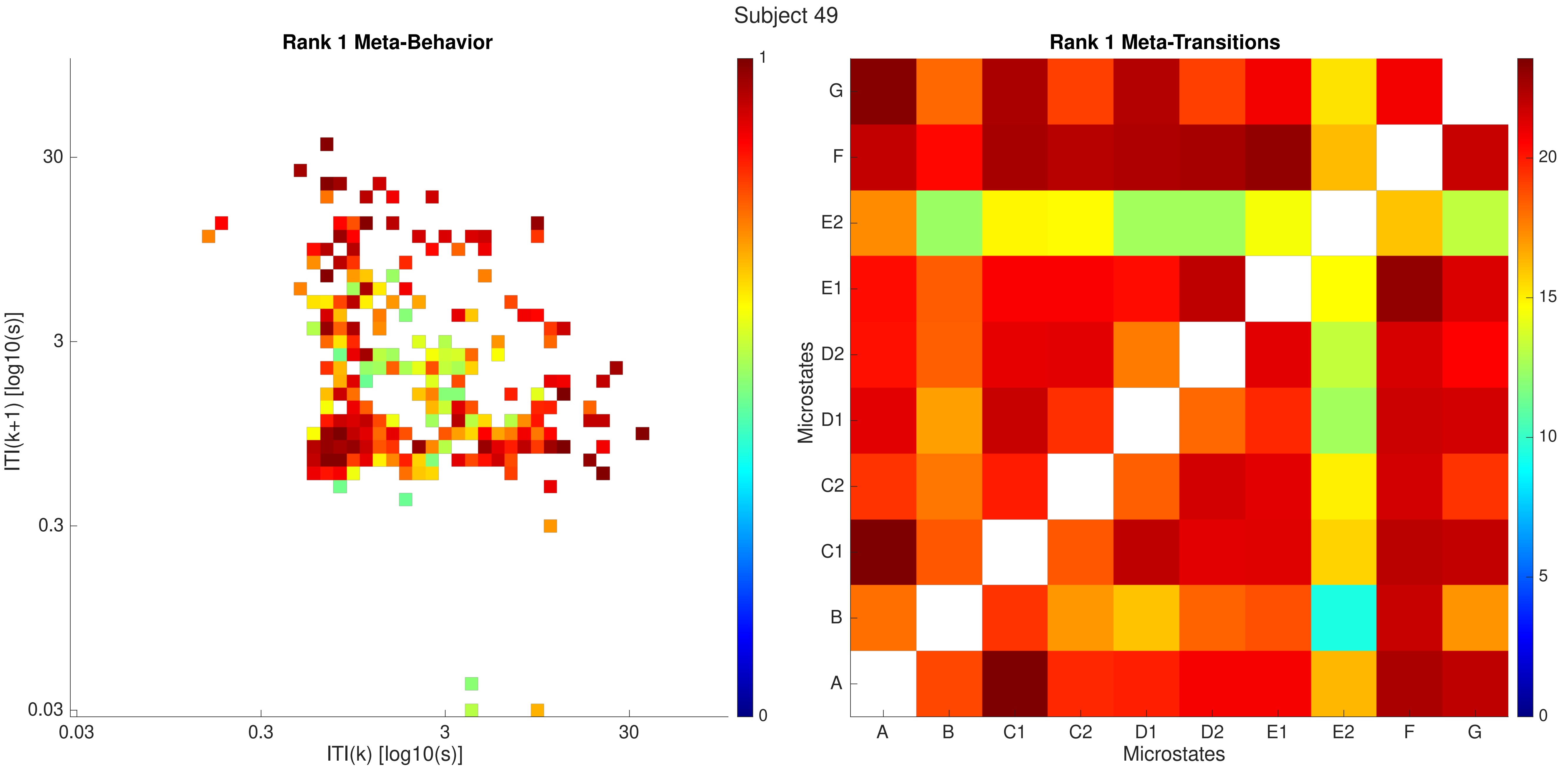

Subject 49
Rank 1 Meta-Behavior
ITI(k+1) [log10(s)]
ITI(k) [log10(s)]
30
3
0.3
0.03
1
0
Rank 1 Meta-Transitions
Microstates
A
B
C1
C2
D1
D2
E1
E2
F
G
20
15
10
5
0

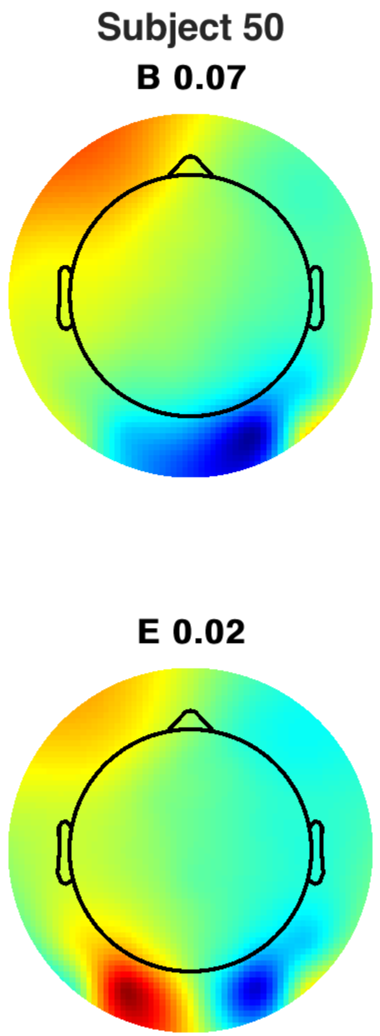
Subject 50
B 0.07
E 0.02

Subject 50

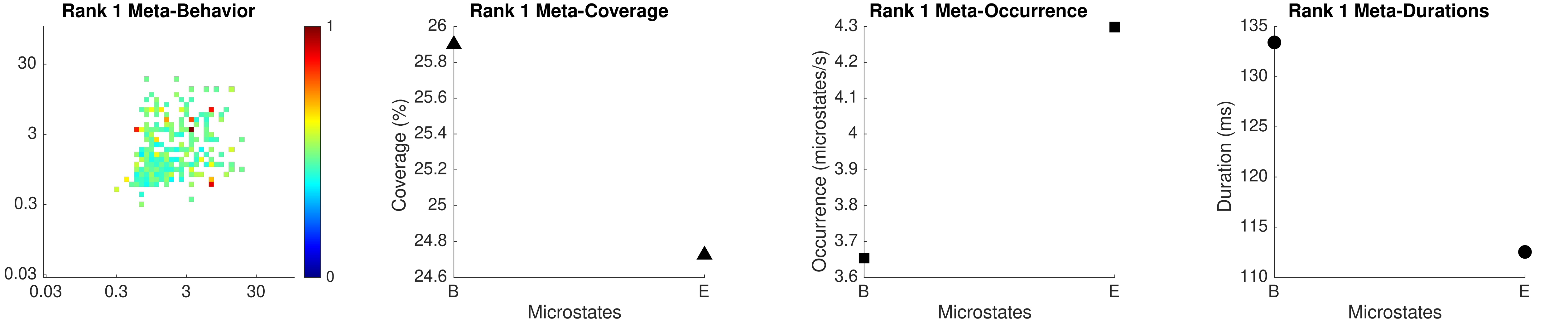

Rank 1 Meta-Behavior
30
3
0.3
0.03
0.03
0.3
3
30
1
0
Rank 1 Meta-Coverage
Coverage (%)
26
25.8
25.6
25.4
25.2
25
24.8
24.6
B
E
Microstates
Rank 1 Meta-Occurrence
Occurrence (microstates/s)
4.3
4.2
4.1
4
3.9
3.8
3.7
3.6
B
E
Microstates
Rank 1 Meta-Durations
Duration (ms)
135
130
125
120
115
110
B
E
Microstates

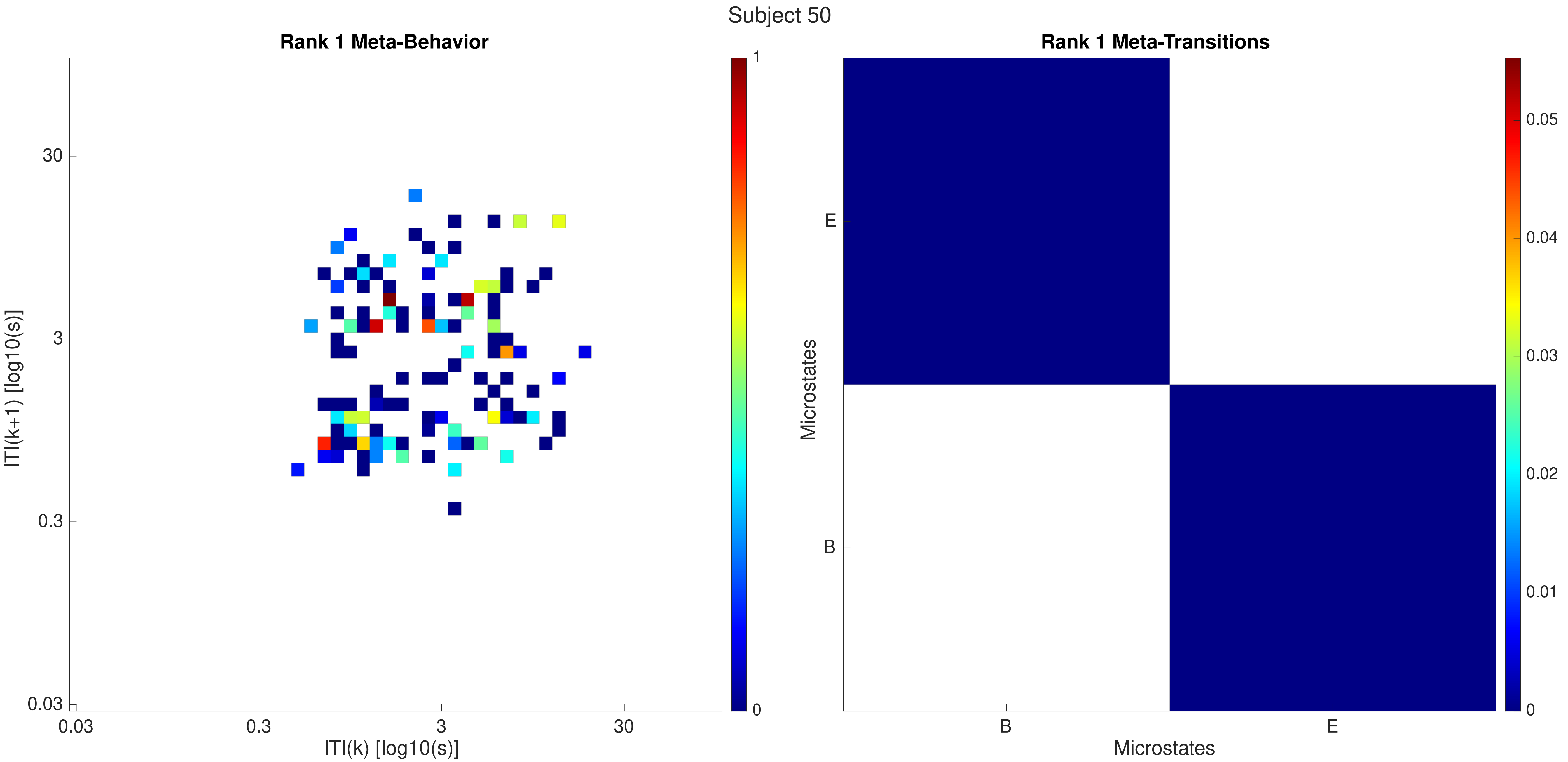

Subject 50
Rank 1 Meta-Behavior
ITI(k+1) [log10(s)]
30
3
0.3
0.03
ITI(k) [log10(s)]
1
0
Rank 1 Meta-Transitions
Microstates
E
B
0.05
0.04
0.03
0.02
0.01

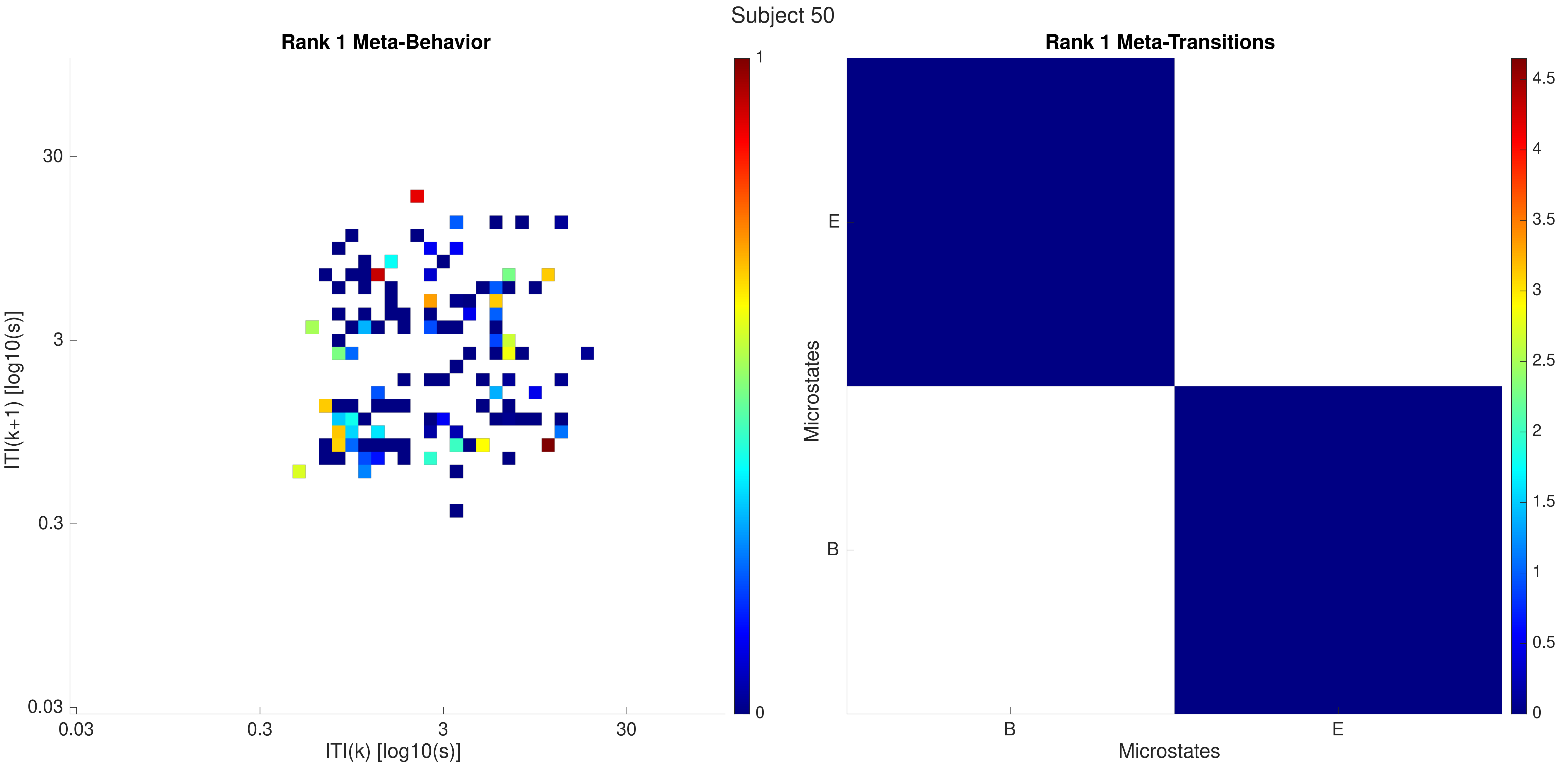

Subject 50
Rank 1 Meta-Behavior
ITI(k+1) [log10(s)]
30
3
0.3
0.03
0.03
0.3
3
30
ITI(k) [log10(s)]
1
0
Rank 1 Meta-Transitions
Microstates
E
B
B
E
Microstates
4.5
4
3.5
3
2.5
2
1.5
1
0.5
0

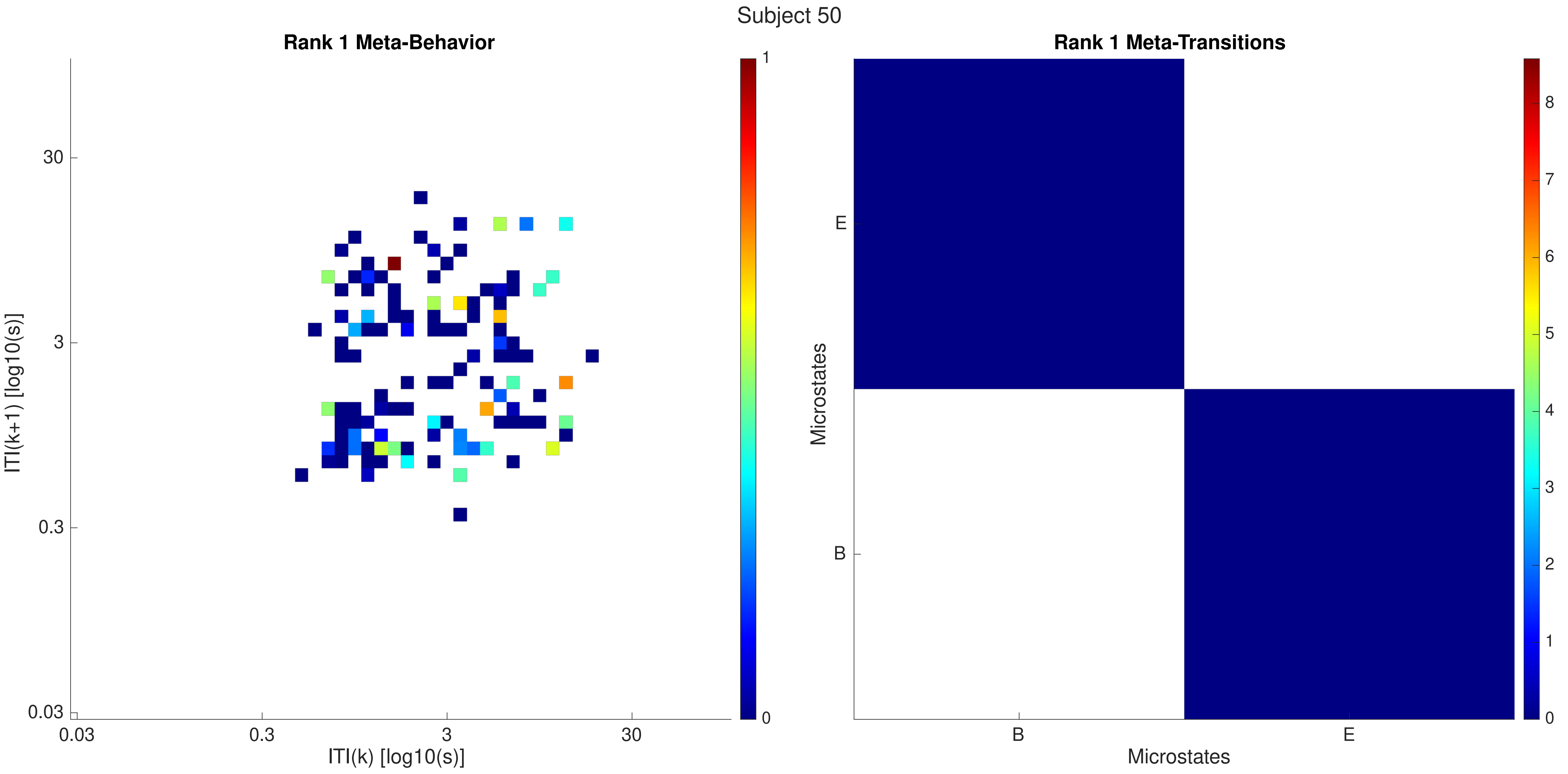

Subject 50
Rank 1 Meta-Behavior
ITI(k+1) [log10(s)]
ITI(k) [log10(s)]
0.03
0.3
3
30
1
0
Rank 1 Meta-Transitions
Microstates
Microstates
E
B
0
1
2
3
4
5
6
7
8

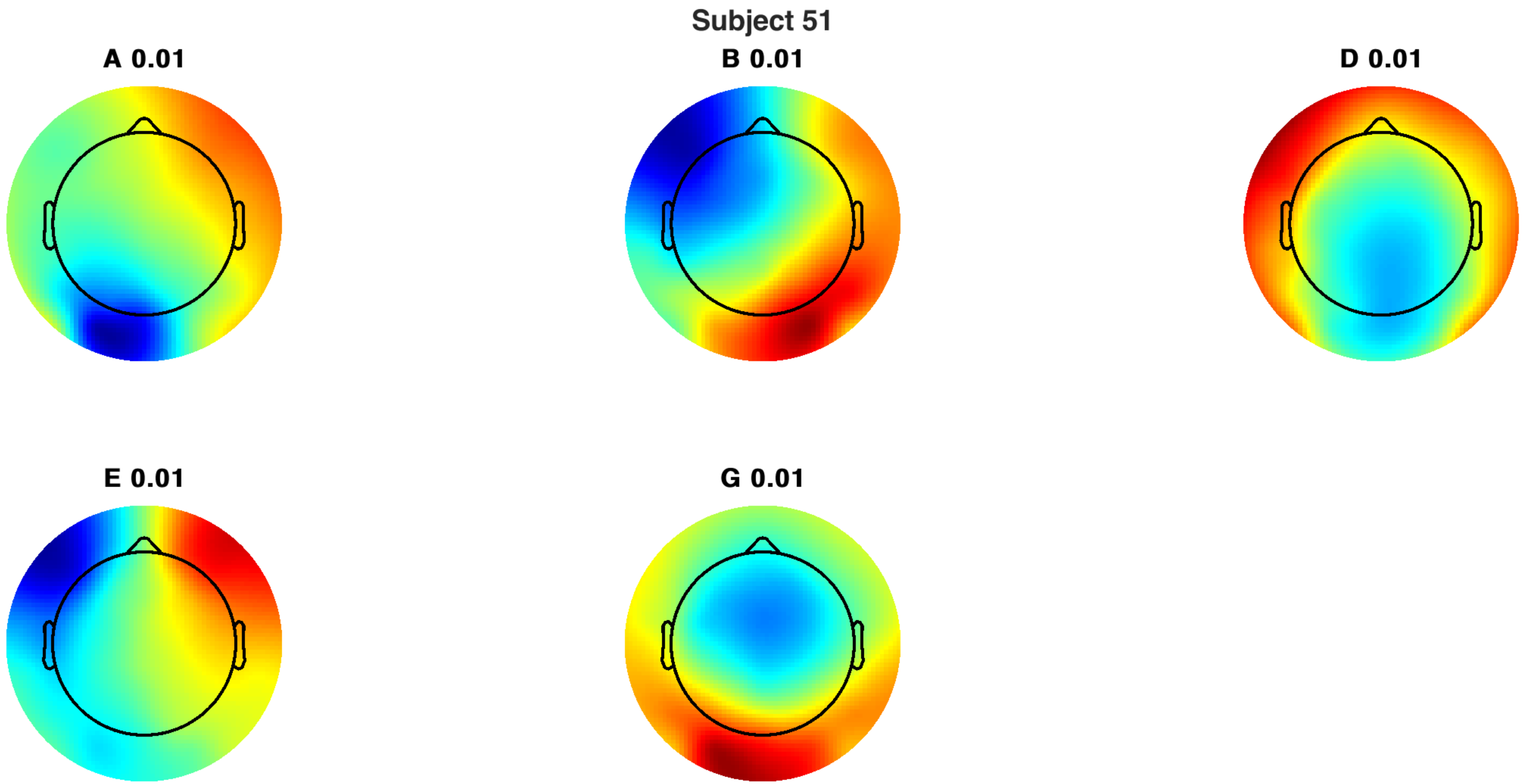
Subject 51
A 0.01
B 0.01
D 0.01
E 0.01
G 0.01

Subject 51

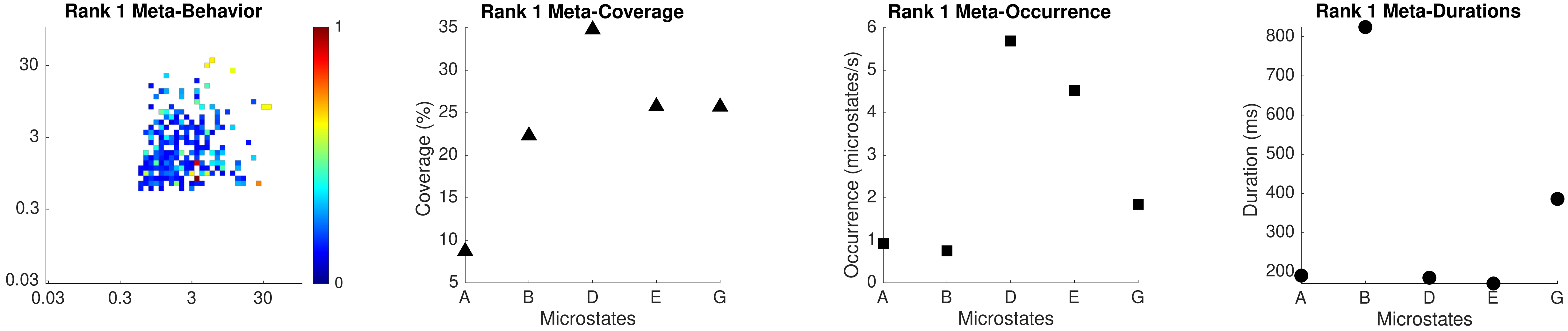

Rank 1 Meta-Behavior
30
3
0.3
0.03
0.03
0.3
3
30
1
0
Rank 1 Meta-Coverage
Coverage (%)
35
30
25
20
15
10
5
A
B
D
E
G
Microstates
Rank 1 Meta-Occurrence
Occurrence (microstates/s)
6
5
4
3
2
1
0
A
B
D
E
G
Microstates
Rank 1 Meta-Durations
Duration (ms)
800
700
600
500
400
300
200
A
B
D
E
G
Microstates

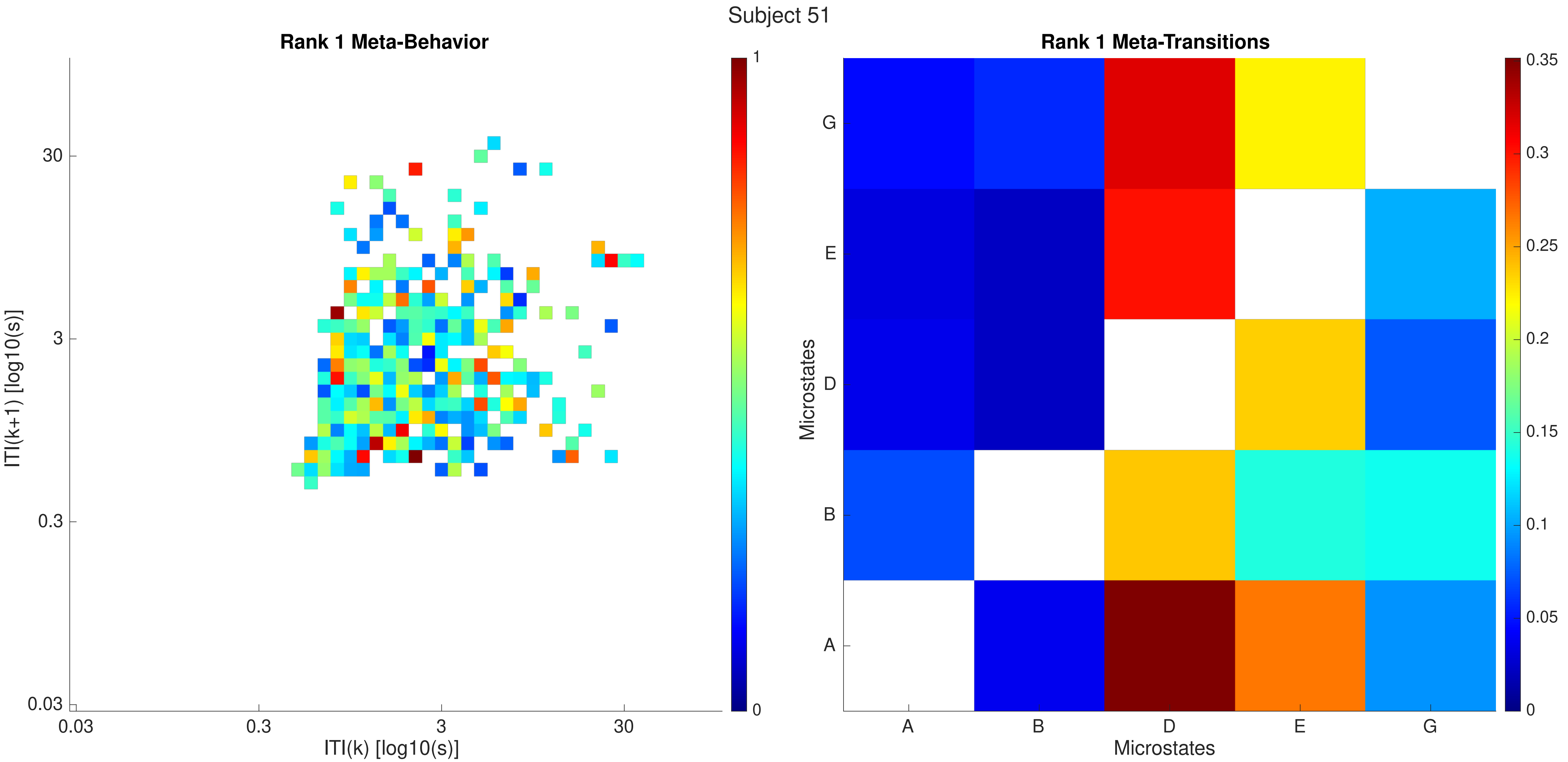

Subject 51
Rank 1 Meta-Behavior
ITI(k+1) [log10(s)]
30
3
0.3
0.03
0.03
0.3
3
30
ITI(k) [log10(s)]
1
0
Rank 1 Meta-Transitions
Microstates
G
E
D
B
A
A
B
D
E
G
Microstates
0.35
0.3
0.25
0.2
0.15
0.1
0.05
0

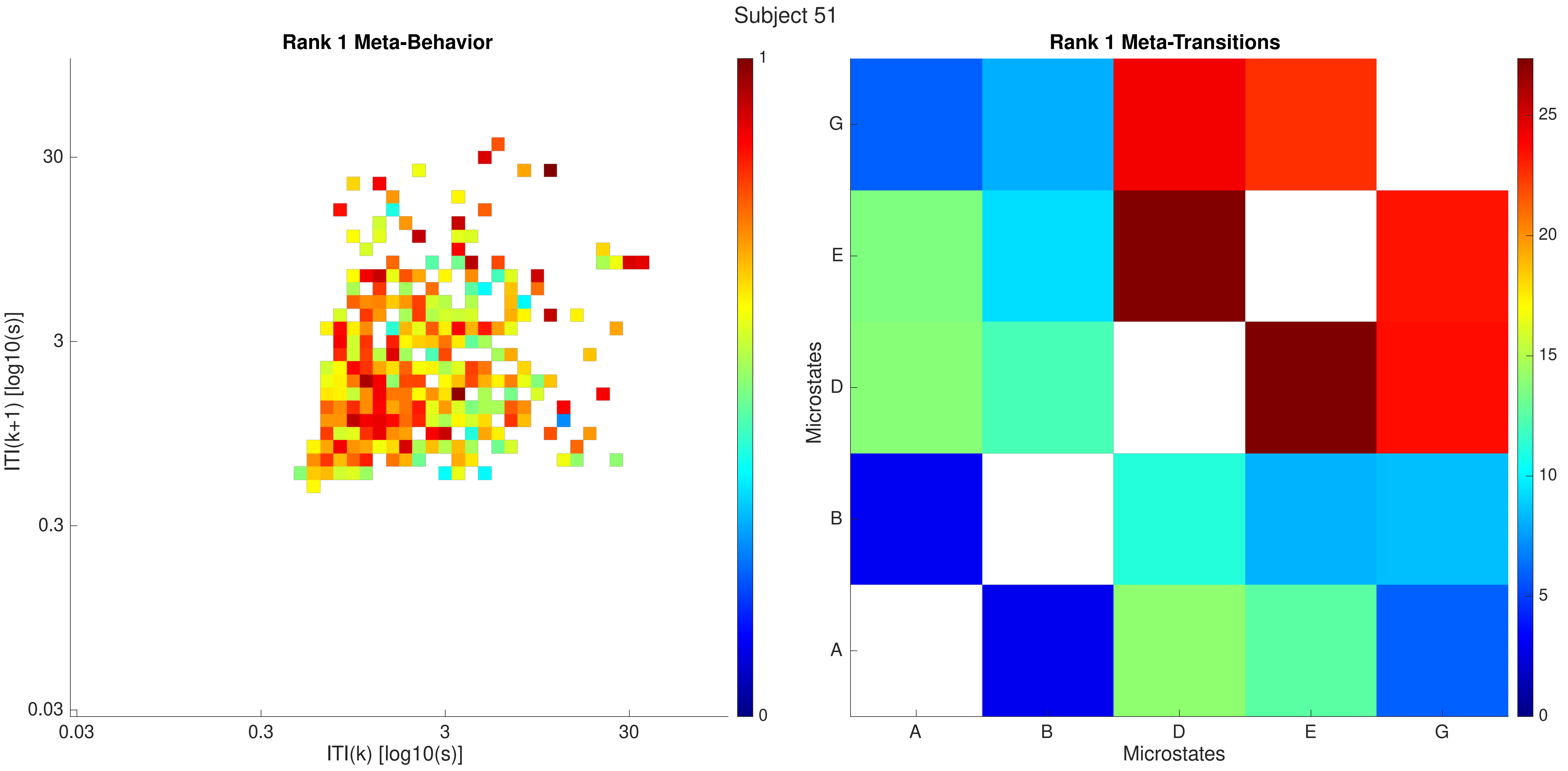

Subject 51
Rank 1 Meta-Behavior
ITI(k+1) [log10(s)]
30
3
0.3
0.03
0.03
0.3
3
30
ITI(k) [log10(s)]
1
0
Rank 1 Meta-Transitions
Microstates
G
E
D
B
A
A
B
D
E
G
Microstates
25
20
15
10
5
0

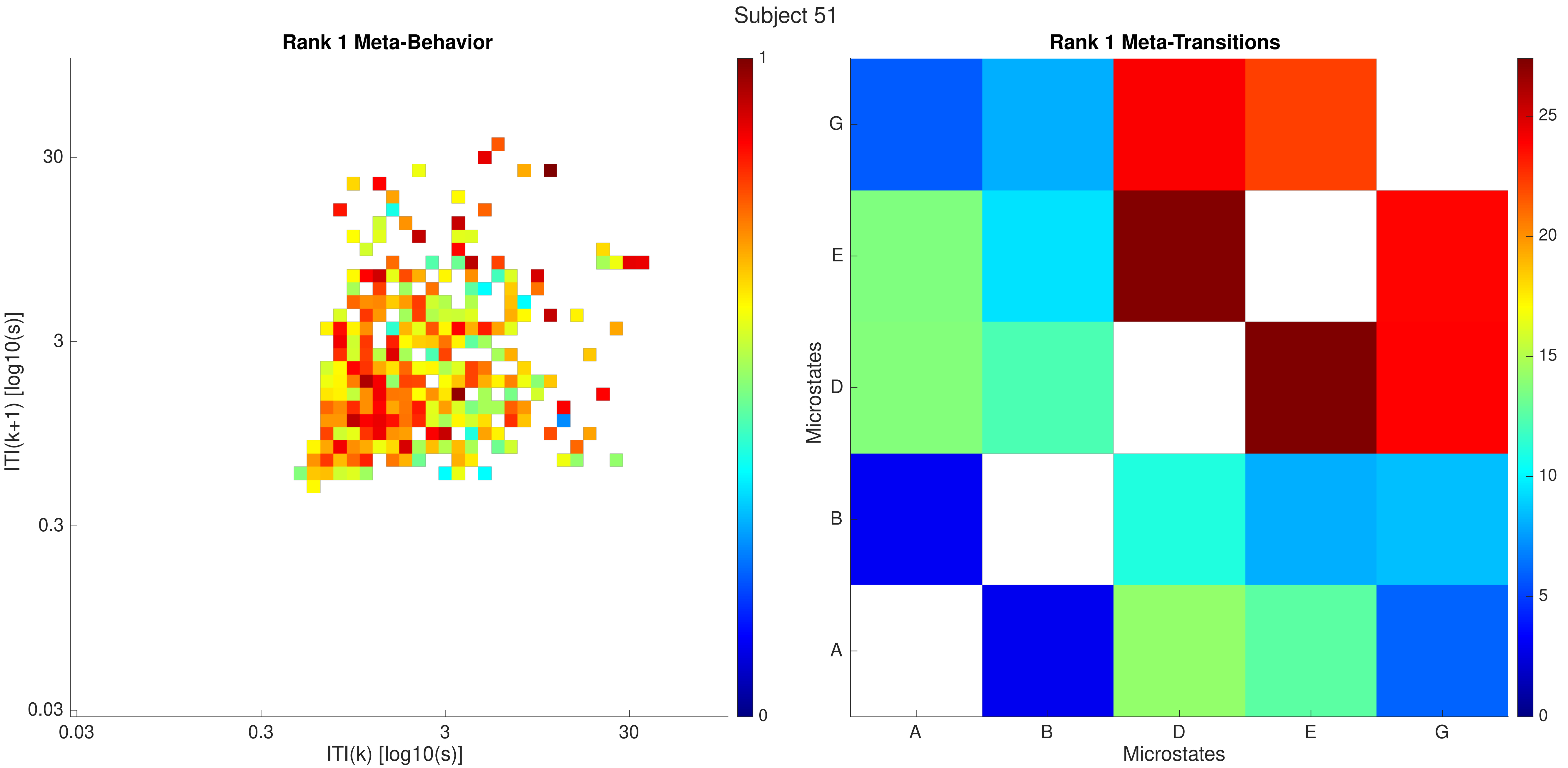
Subject 51
Rank 1 Meta-Behavior
ITI(k+1) [log10(s)]
ITI(k) [log10(s)]
0.03
0.3
3
30
0
1
Rank 1 Meta-Transitions
Microstates
Microstates
A
B
D
E
G
0
5
10
15
20
25

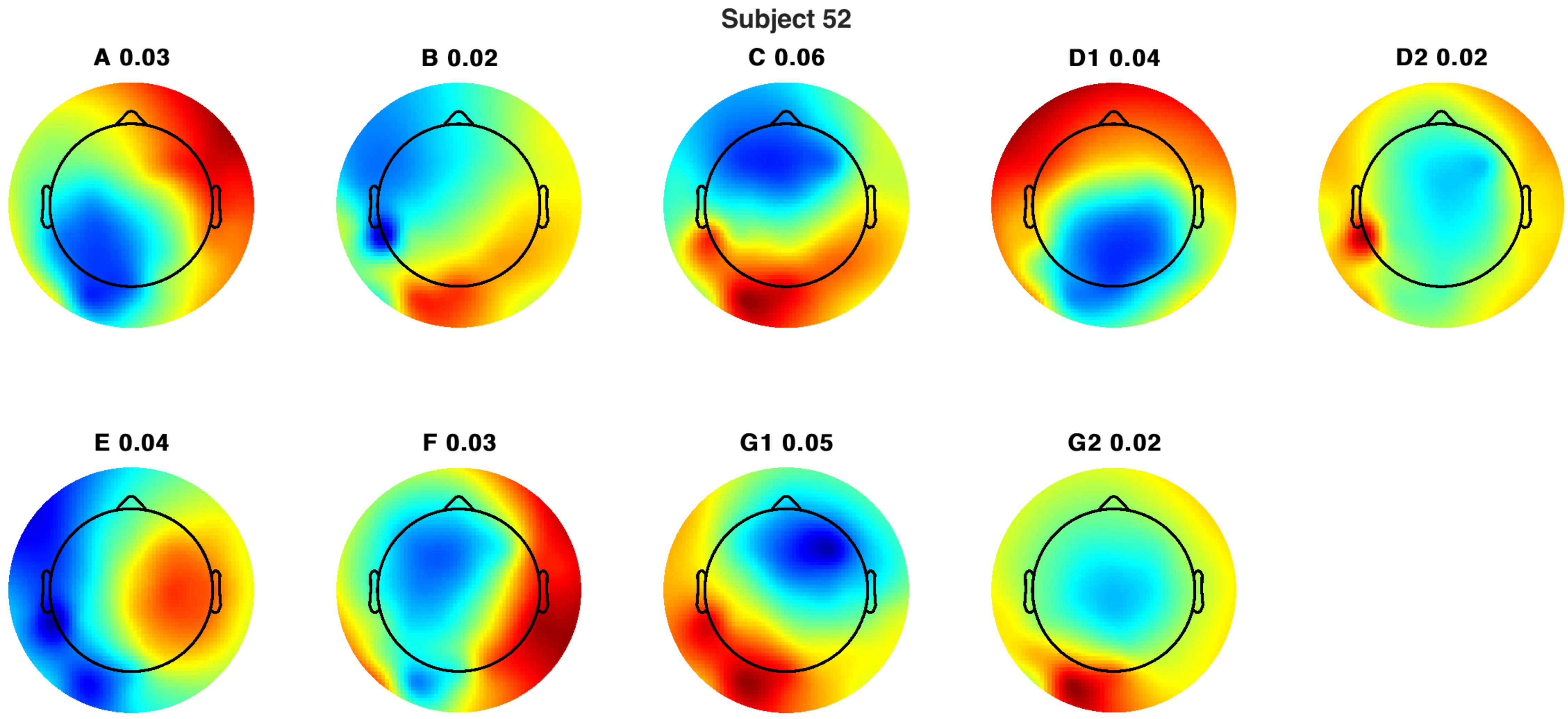

Subject 52
A 0.03
B 0.02
C 0.06
D1 0.04
D2 0.02
E 0.04
F 0.03
G1 0.05
G2 0.02

Subject 52

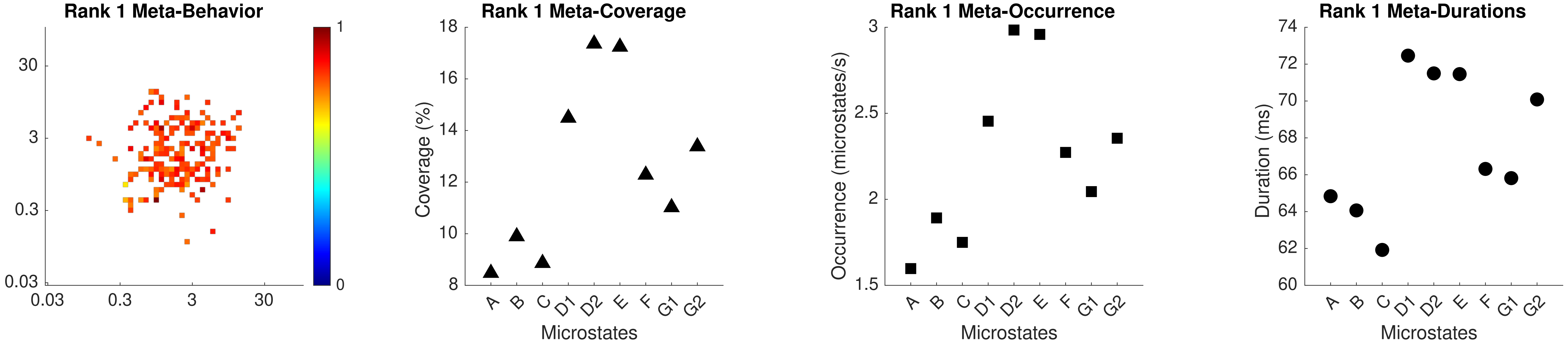

Rank 1 Meta-Behavior
30
3
0.3
0.03
0.03
0.3
3
30
1
0
Rank 1 Meta-Coverage
Coverage (%)
18
16
14
12
10
8
A
B
C
D1
D2
E
F
G1
G2
Microstates
Rank 1 Meta-Occurrence
Occurrence (microstates/s)
3
2.5
2
1.5
A
B
C
D1
D2
E
F
G1
G2
Microstates
Rank 1 Meta-Durations
Duration (ms)
74
72
70
68
66
64
62
60
A
B
C
D1
D2
E
F
G1
G2
Microstates

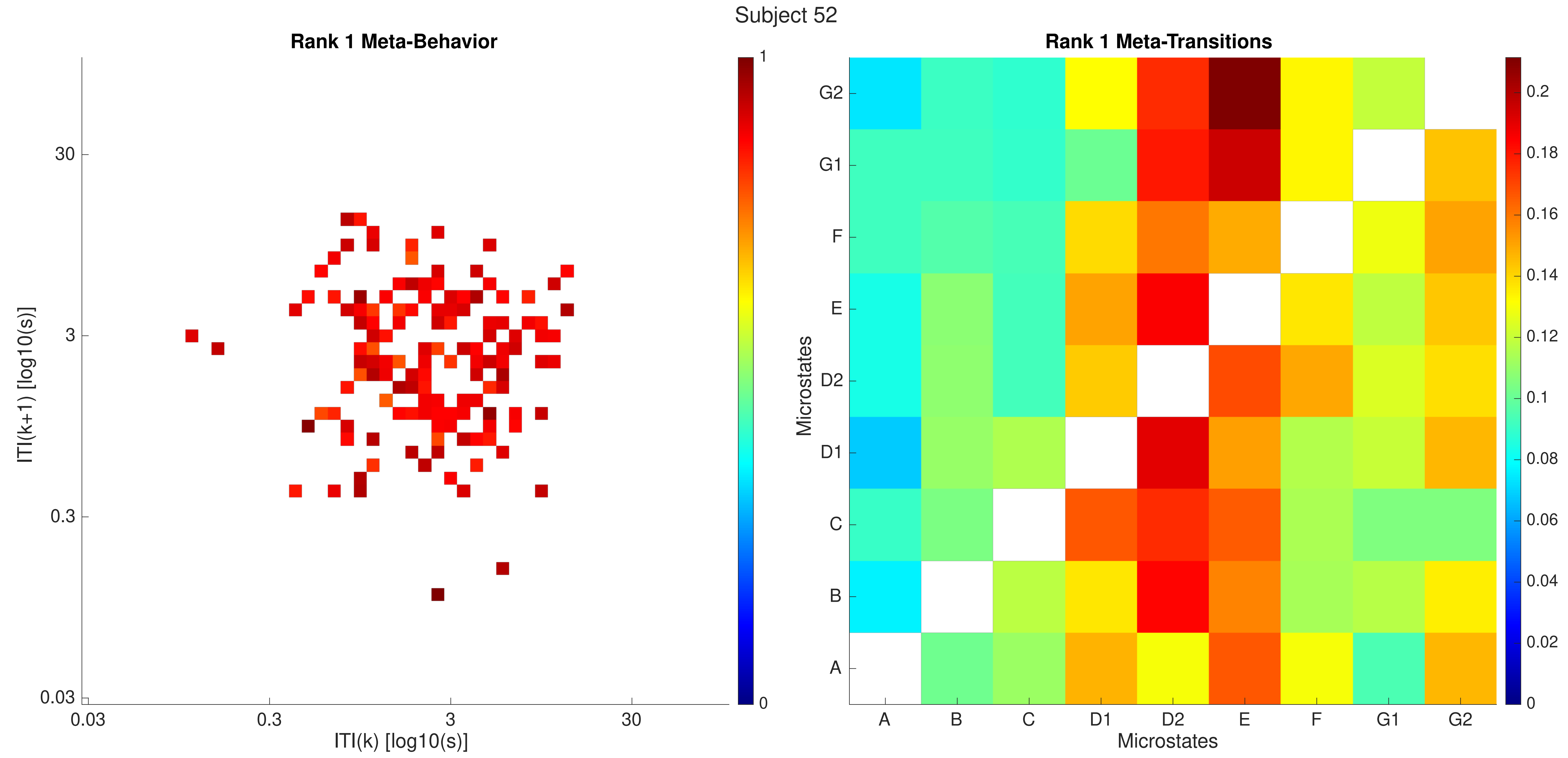
Subject 52
Rank 1 Meta-Behavior
ITI(k+1) [log10(s)]
ITI(k) [log10(s)]
30
3
0.3
0.03
1
0
Rank 1 Meta-Transitions
Microstates
A
B
C
D1
D2
E
F
G1
G2
0.2
0.18
0.16
0.14
0.12
0.1
0.08
0.06
0.04
0.02
0

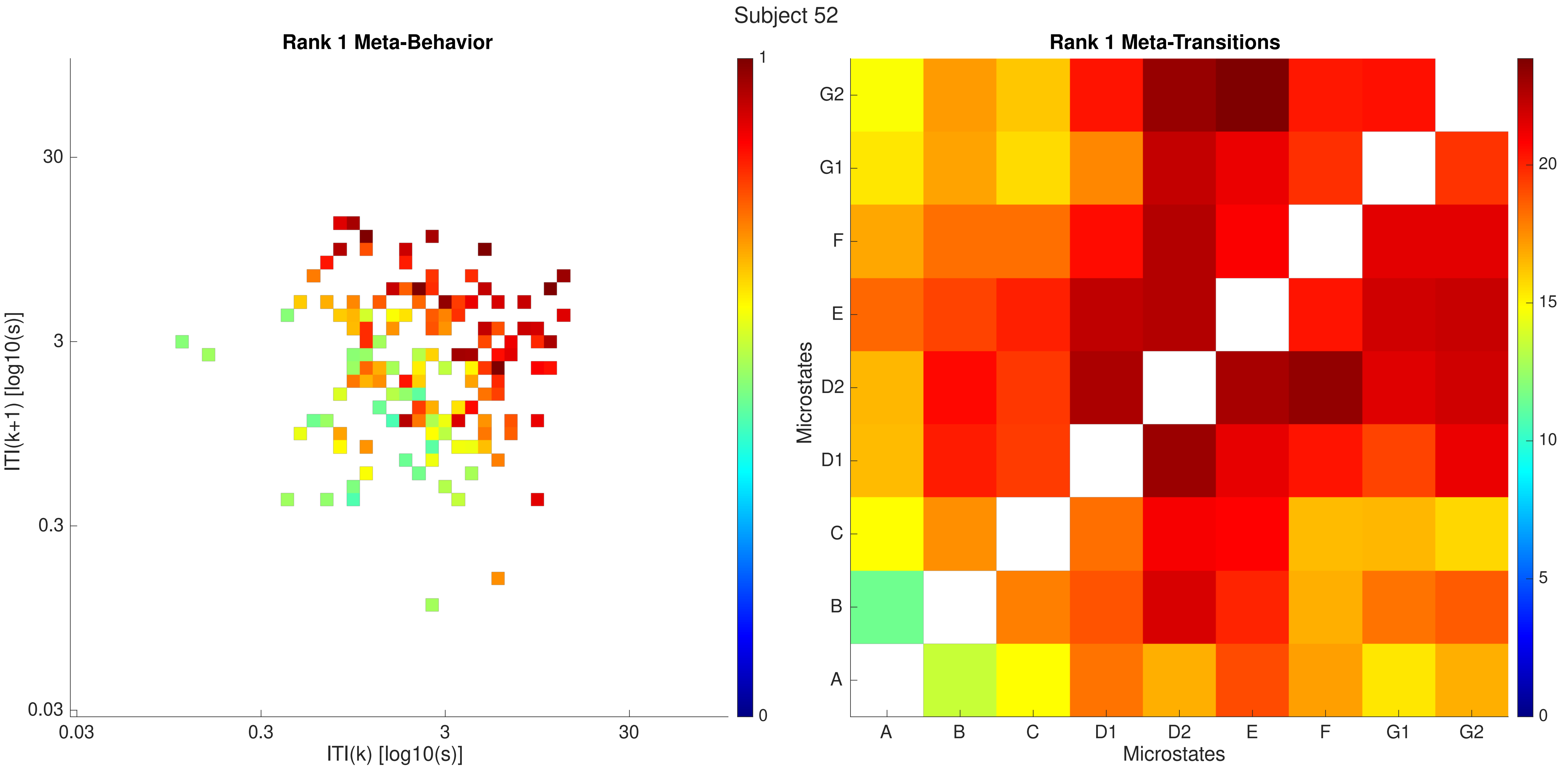
Subject 52
Rank 1 Meta-Behavior
ITI(k+1) [log10(s)]
ITI(k) [log10(s)]
0.03
0.3
3
30
1
0
Rank 1 Meta-Transitions
Microstates
A
B
C
D1
D2
E
F
G1
G2
0
5
10
15
20

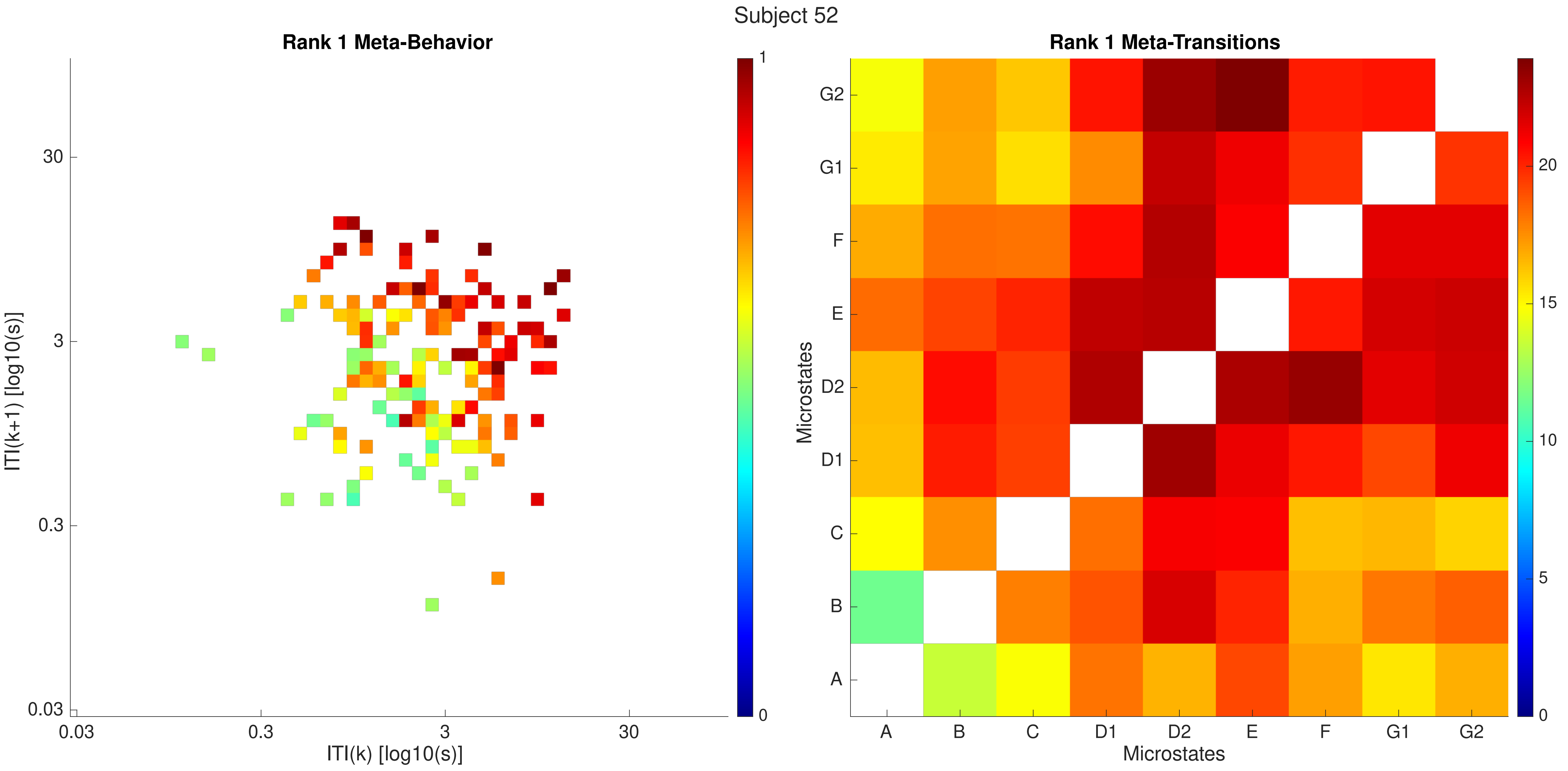
Subject 52
Rank 1 Meta-Behavior
ITI(k+1) [log10(s)]
ITI(k) [log10(s)]
0.03
0.3
3
30
0
1
Rank 1 Meta-Transitions
Microstates
A
B
C
D1
D2
E
F
G1
G2
0
5
10
15
20

Subject 53

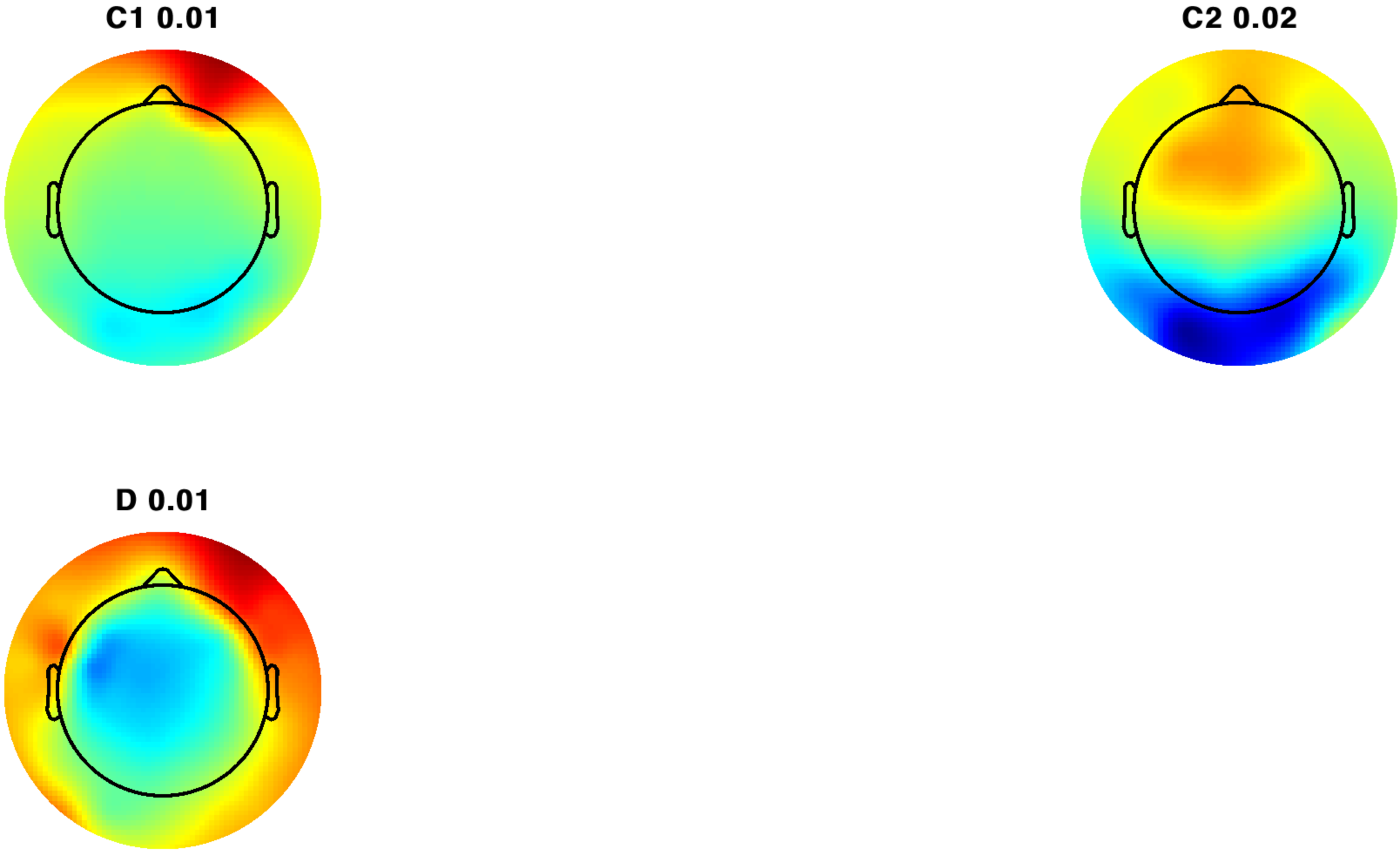

Subject 53

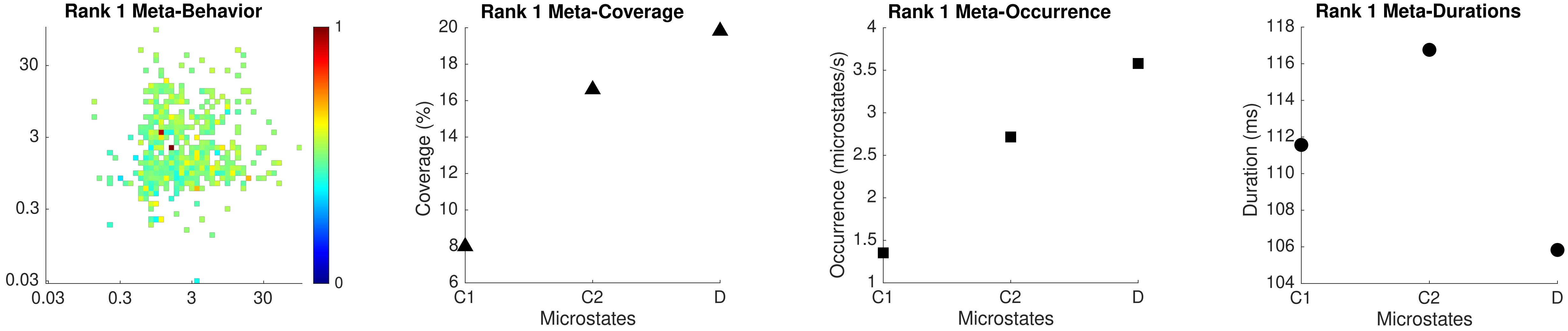
Rank 1 Meta-Behavior
30
3
0.3
0.03
0.03
0.3
3
30
1
0
Rank 1 Meta-Coverage
Coverage (%)
20
18
16
14
12
10
8
6
C1
C2
D
Microstates
Rank 1 Meta-Occurrence
Occurrence (microstates/s)
4
3.5
3
2.5
2
1.5
1
C1
C2
D
Microstates
Rank 1 Meta-Durations
Duration (ms)
118
116
114
112
110
108
106
104
C1
C2
D
Microstates

Subject 53

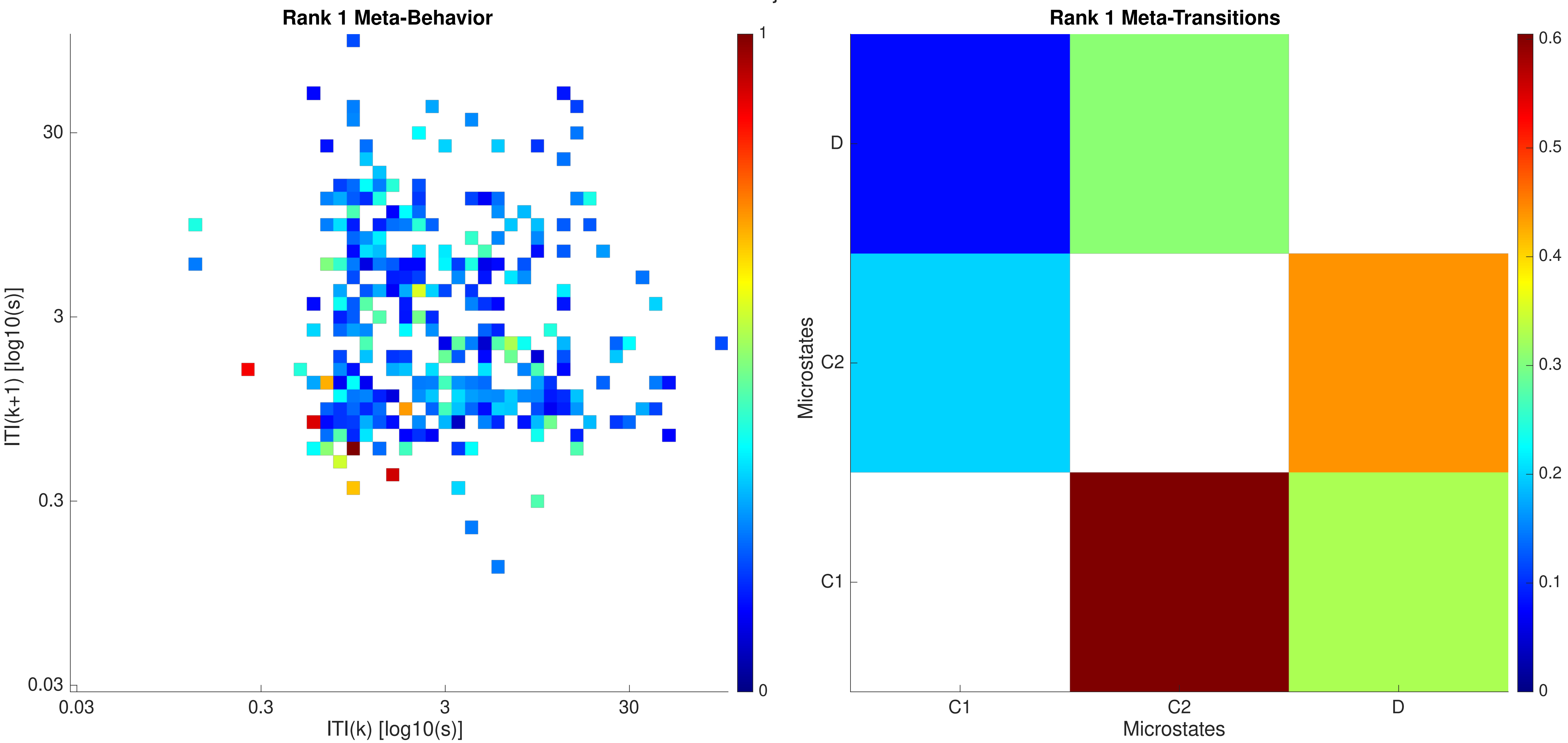
Rank 1 Meta-Behavior
ITI(k+1) [log10(s)]
ITI(k) [log10(s)]
30
3
0.3
0.03
0.03
0.3
3
30
1
0
Rank 1 Meta-Transitions
Microstates
D
C2
C1
C1
C2
D
Microstates
0.6
0.5
0.4
0.3
0.2
0.1
0

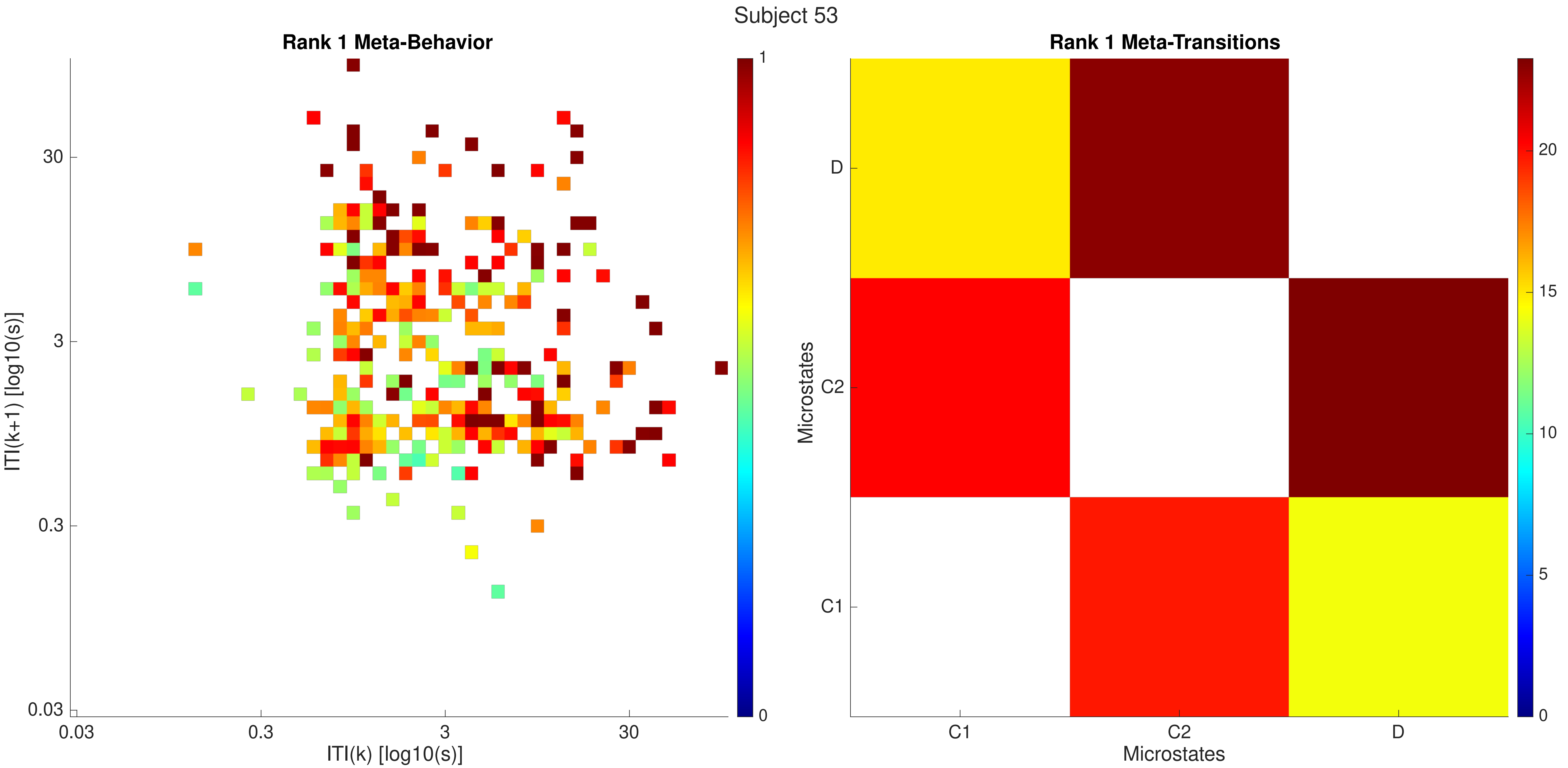

Subject 53
Rank 1 Meta-Behavior
ITI(k+1) [log10(s)]
ITI(k) [log10(s)]
0.03
0.3
3
30
0
1
Rank 1 Meta-Transitions
Microstates
C1
C2
D
0
5
10
15
20

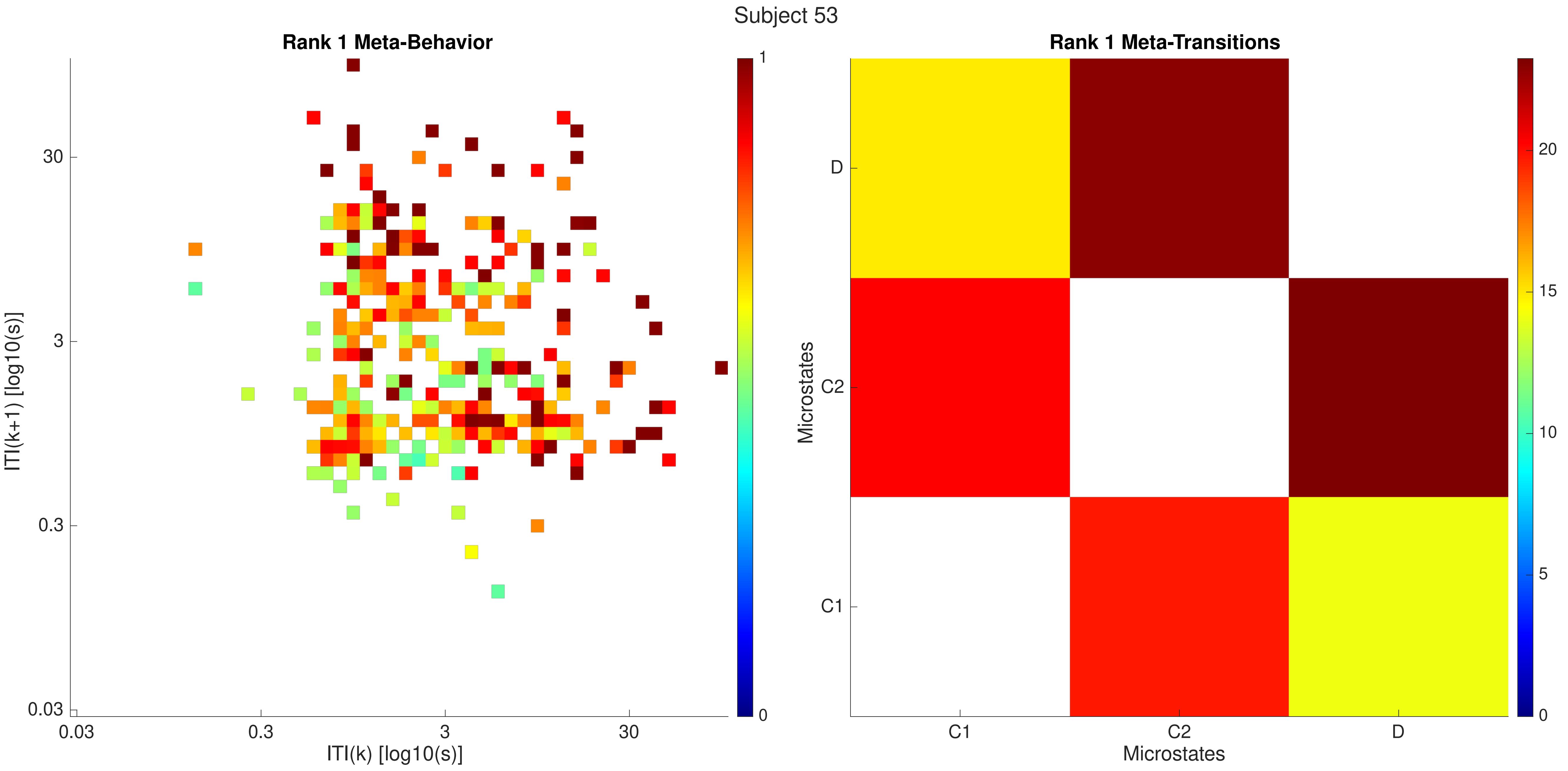
Subject 53
Rank 1 Meta-Behavior
ITI(k+1) [log10(s)]
30
3
0.3
0.03
0.03
0.3
3
30
ITI(k) [log10(s)]
1
0
Rank 1 Meta-Transitions
Microstates
D
C2
C1
C1
C2
D
Microstates
20
15
10
5
0

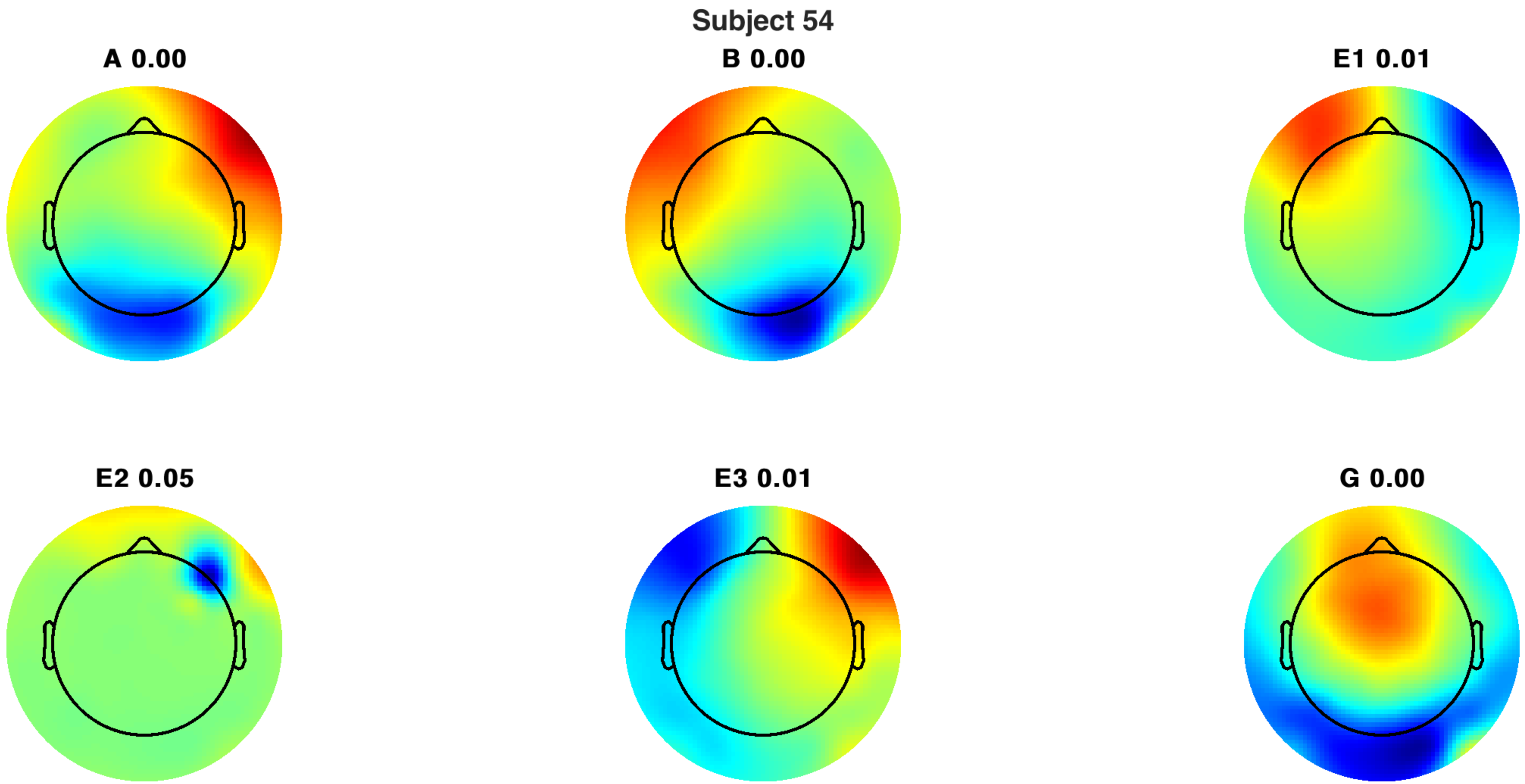
Subject 54
A 0.00
B 0.00
E1 0.01
E2 0.05
E3 0.01
G 0.00

Subject 54

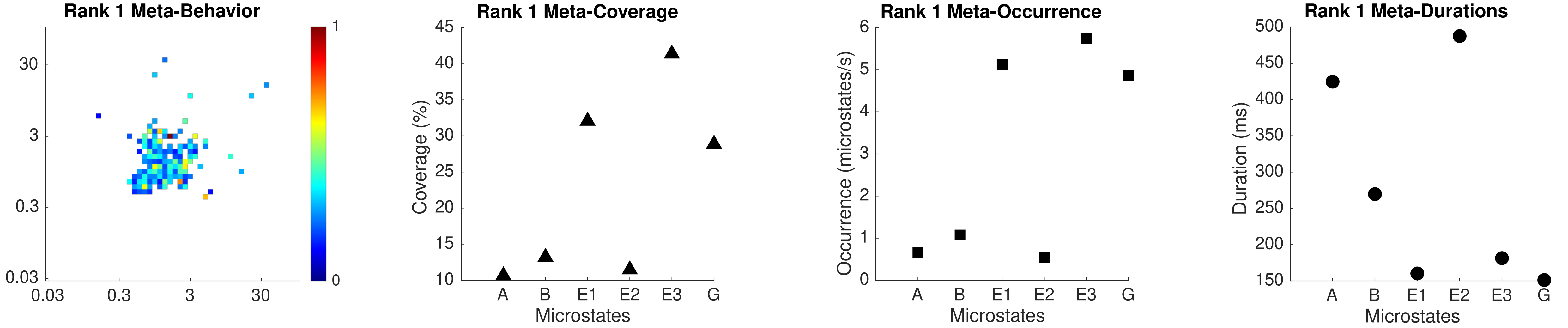

Rank 1 Meta-Behavior
30
3
0.3
0.03
0.03
0.3
3
30
1
0
Rank 1 Meta-Coverage
Coverage (%)
45
40
35
30
25
20
15
10
A
B
E1
E2
E3
G
Microstates
Rank 1 Meta-Occurrence
Occurrence (microstates/s)
6
5
4
3
2
1
0
A
B
E1
E2
E3
G
Microstates
Rank 1 Meta-Durations
Duration (ms)
500
450
400
350
300
250
200
150
A
B
E1
E2
E3
G
Microstates

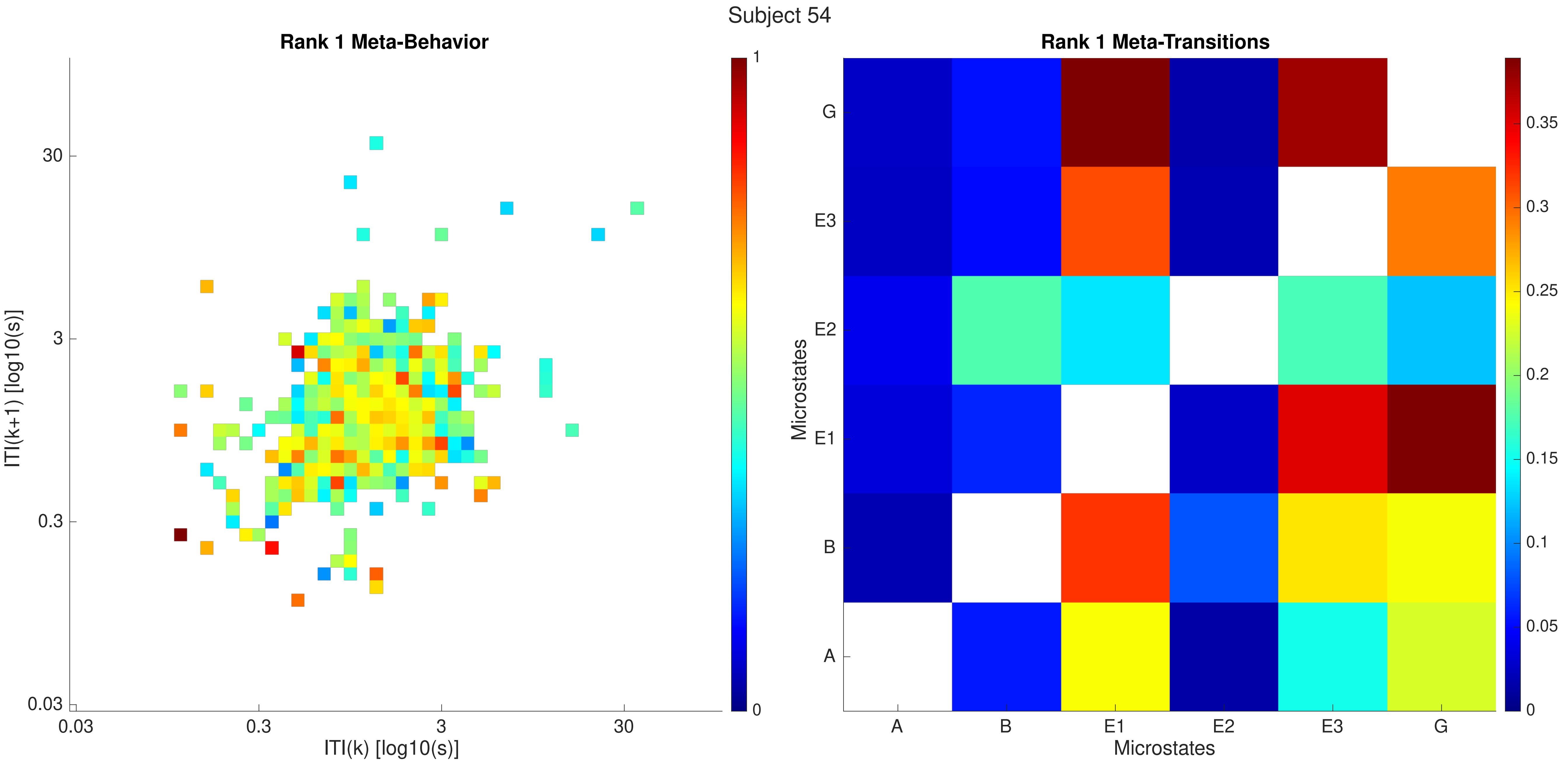
Subject 54
Rank 1 Meta-Behavior
ITI(k+1) [log10(s)]
ITI(k) [log10(s)]
0.03
0.3
3
30
1
0
Rank 1 Meta-Transitions
Microstates
A
B
E1
E2
E3
G
0
0.05
0.1
0.15
0.2
0.25
0.3
0.35

Subject 54

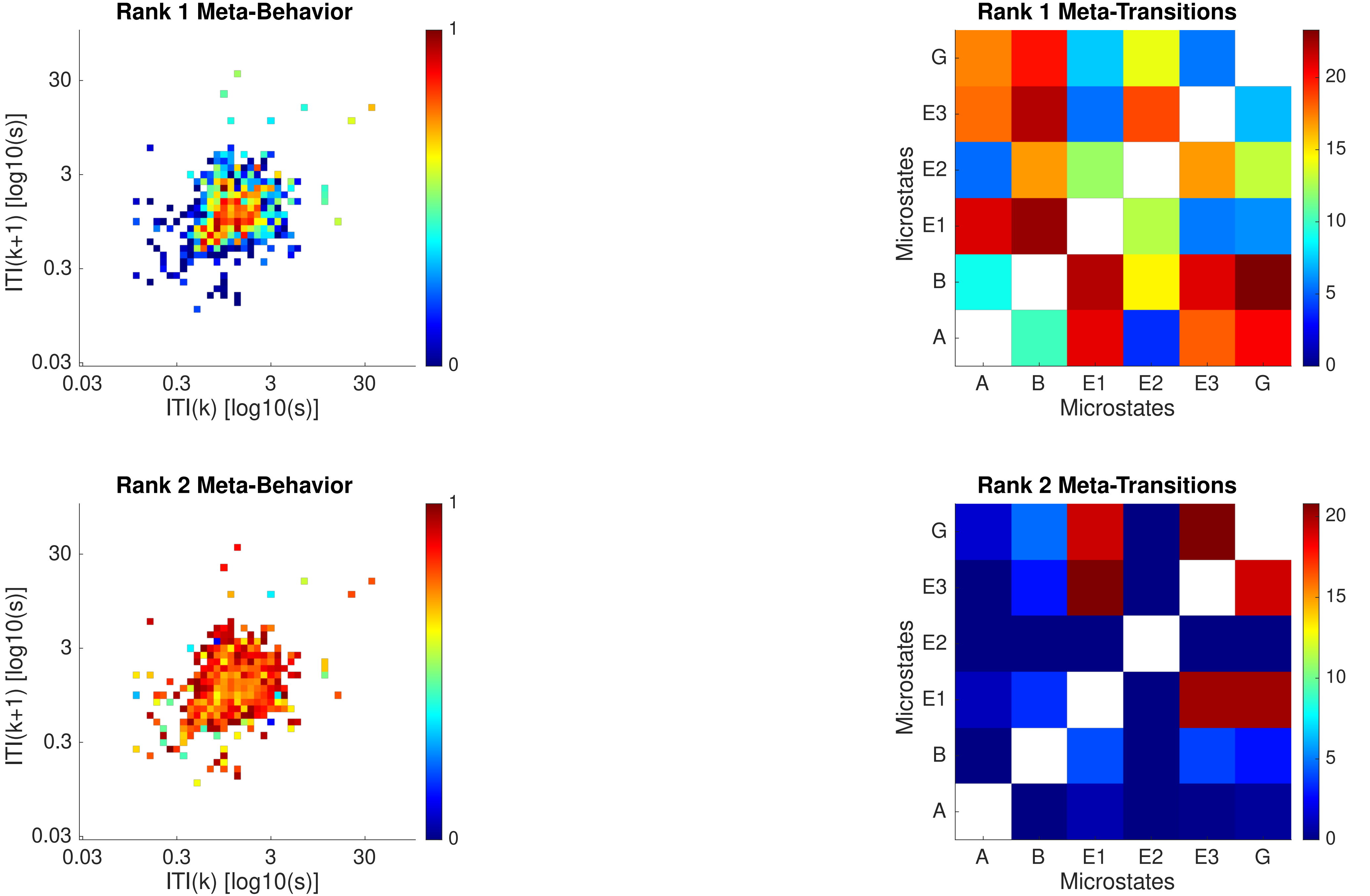

Rank 1 Meta-Behavior
ITI(k+1) [log10(s)]
ITI(k) [log10(s)]
0.03
0.3
3
30
0
1
Rank 1 Meta-Transitions
Microstates
A
B
E1
E2
E3
G
0
5
10
15
20
Rank 2 Meta-Behavior
Rank 2 Meta-Transitions

Subject 54

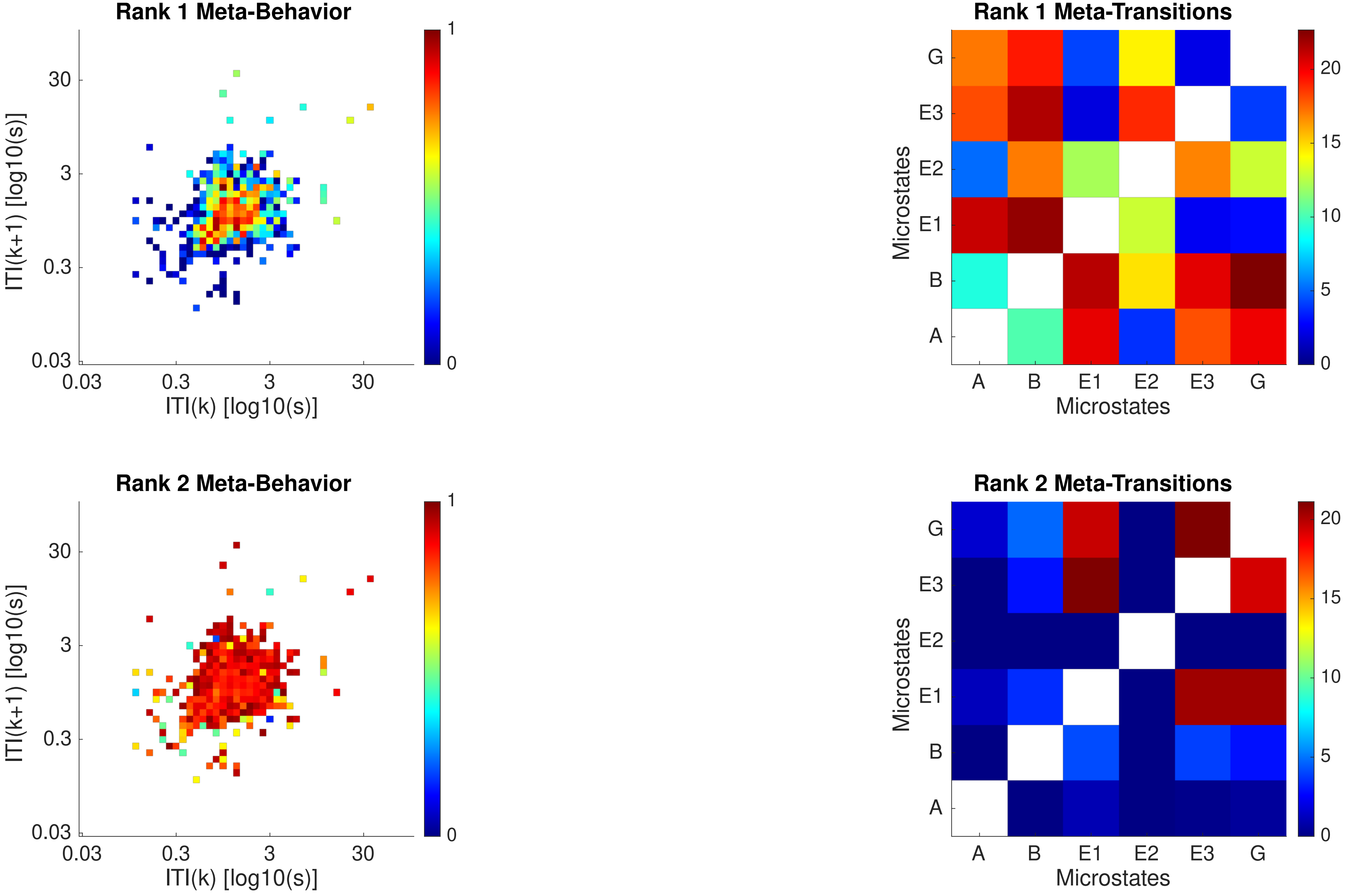
Rank 1 Meta-Behavior
ITI(k+1) [log10(s)]
ITI(k) [log10(s)]
0.03
0.3
3
30
0
1
Rank 1 Meta-Transitions
Microstates
G
E3
E2
E1
B
A
0
5
10
15
20
Rank 2 Meta-Behavior
ITI(k+1) [log10(s)]
ITI(k) [log10(s)]
Rank 2 Meta-Transitions
Microstates

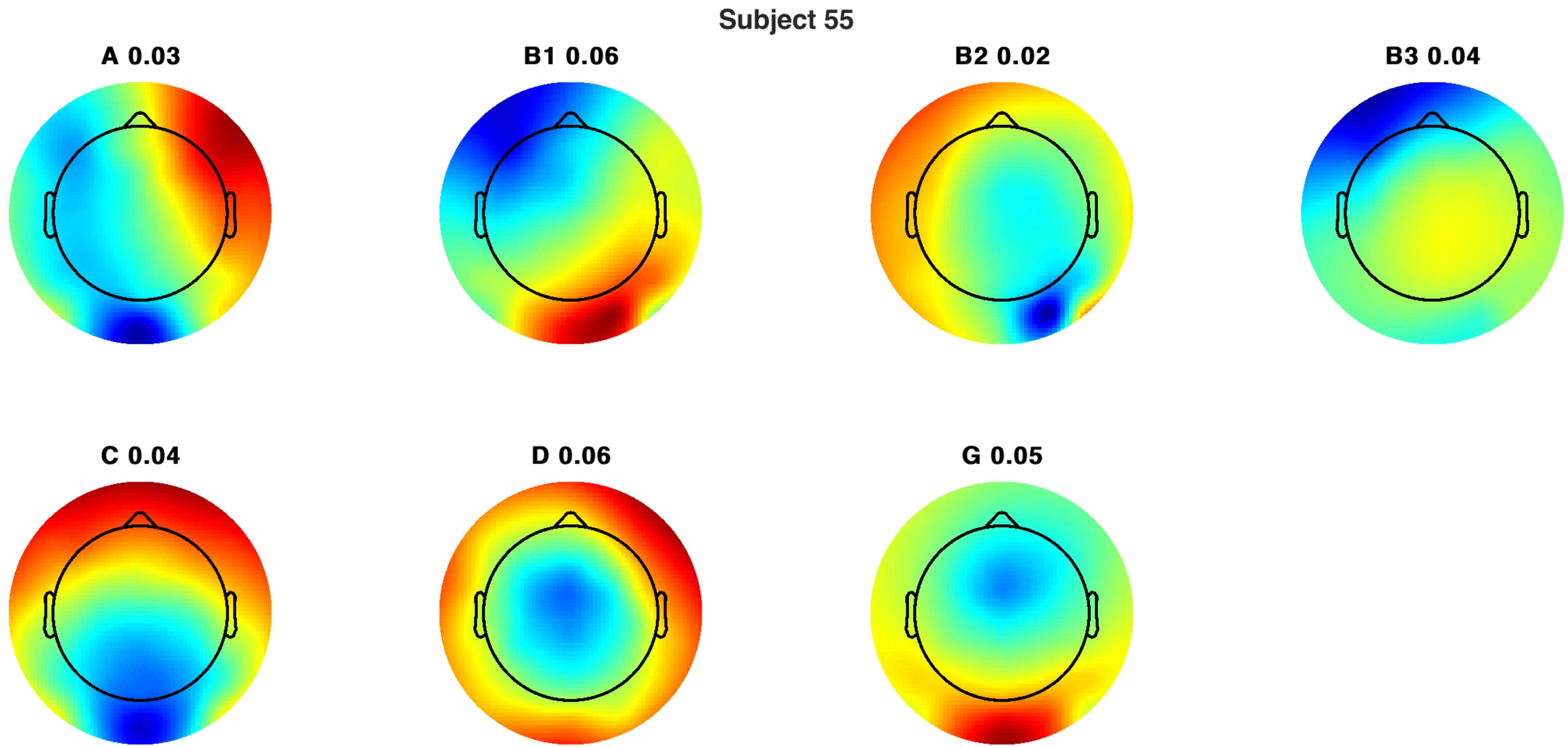
Subject 55
A 0.03
B1 0.06
B2 0.02
B3 0.04
C 0.04
D 0.06
G 0.05

Subject 55

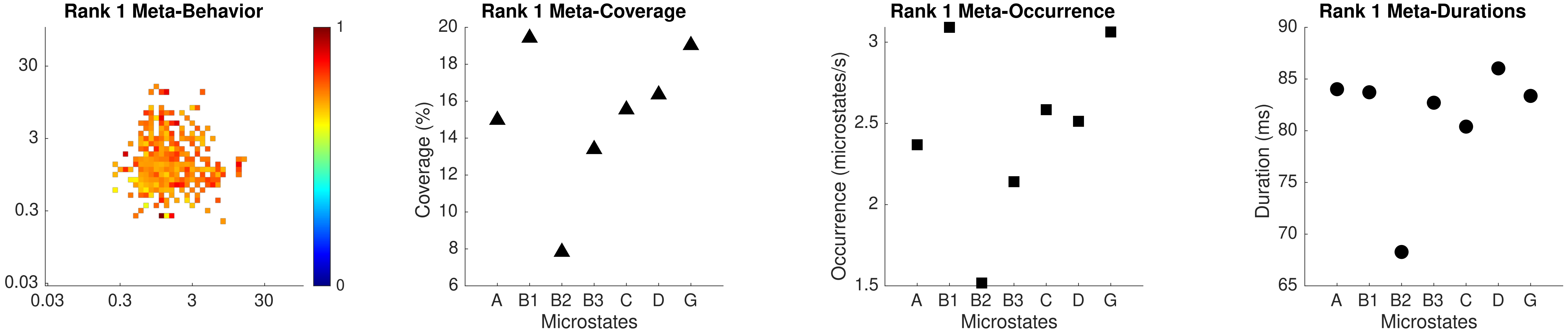

Rank 1 Meta-Behavior
30
3
0.3
0.03
0.03
0.3
3
30
1
0
Rank 1 Meta-Coverage
Coverage (%)
20
18
16
14
12
10
8
6
A
B1
B2
B3
C
D
G
Microstates
Rank 1 Meta-Occurrence
Occurrence (microstates/s)
3
2.5
2
1.5
A
B1
B2
B3
C
D
G
Microstates
Rank 1 Meta-Durations
Duration (ms)
90
85
80
75
70
65
A
B1
B2
B3
C
D
G
Microstates

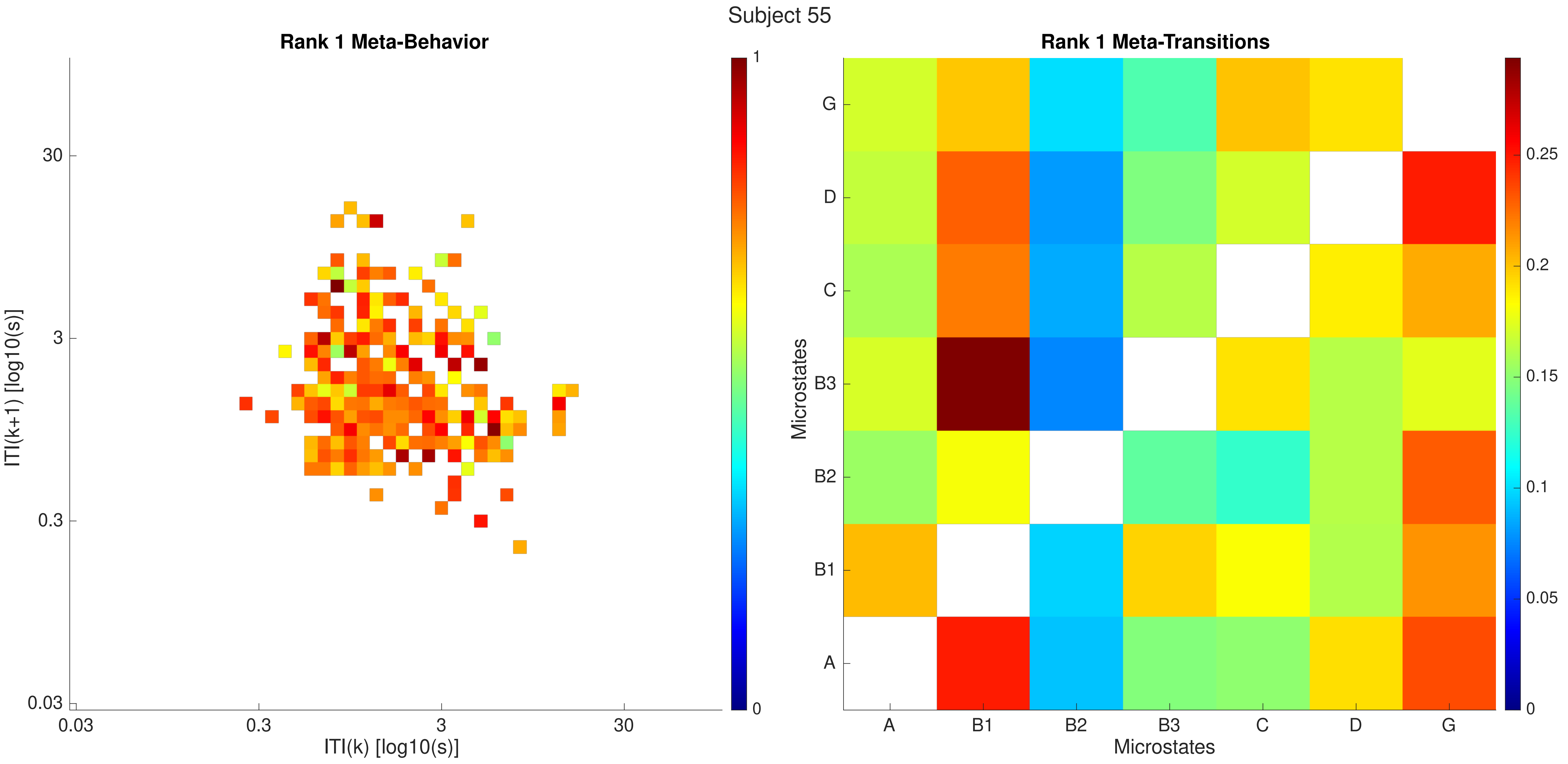

Subject 55
Rank 1 Meta-Behavior
ITI(k+1) [log10(s)]
ITI(k) [log10(s)]
0.03
0.3
3
30
1
0
Rank 1 Meta-Transitions
Microstates
Microstates
A
B1
B2
B3
C
D
G
0
0.05
0.1
0.15
0.2
0.25

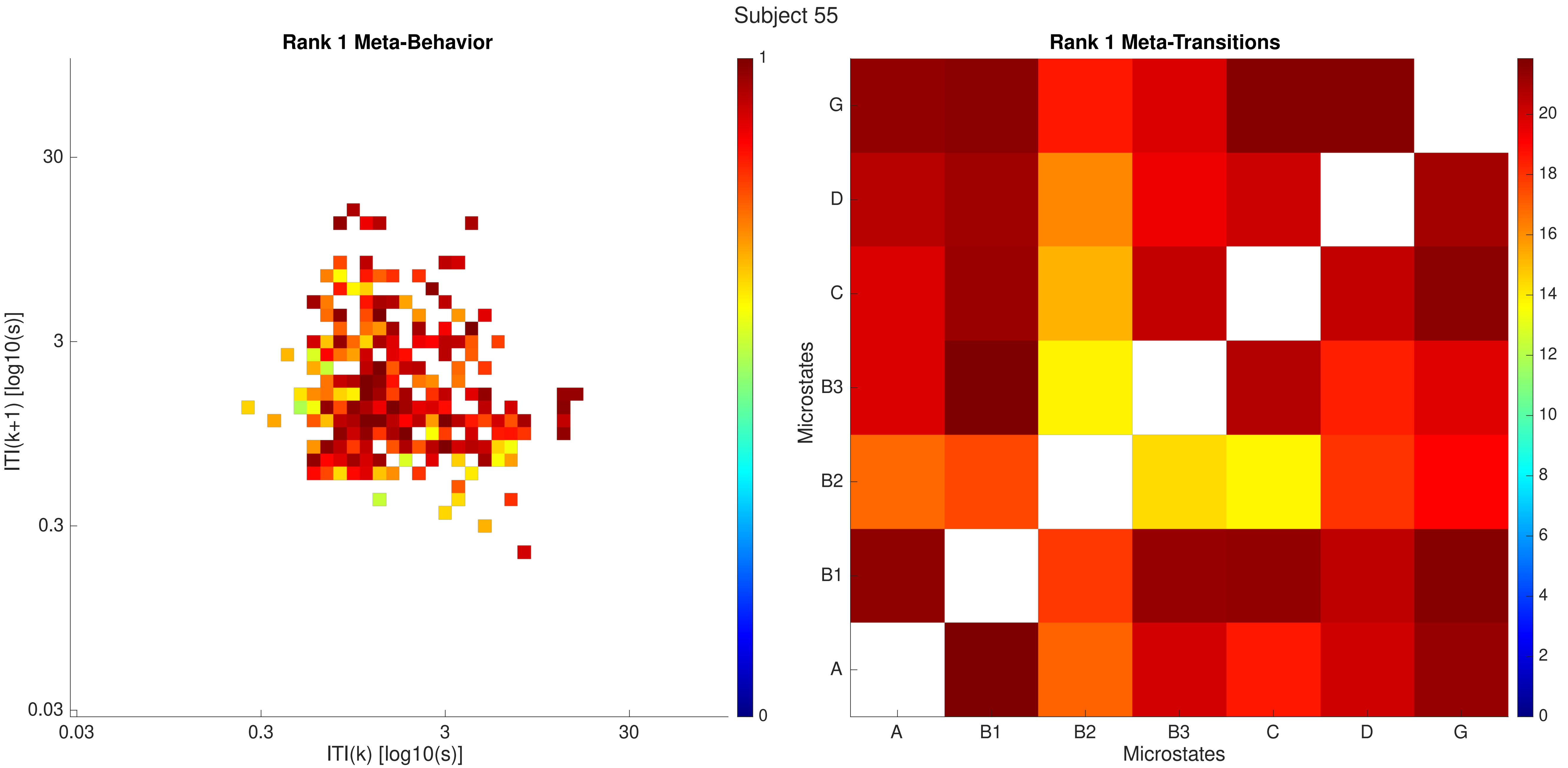
Subject 55
Rank 1 Meta-Behavior
ITI(k+1) [log10(s)]
30
3
0.3
0.03
ITI(k) [log10(s)]
0.03
0.3
3
30
1
0
Rank 1 Meta-Transitions
Microstates
G
D
C
B3
B2
B1
A
Microstates
A
B1
B2
B3
C
D
G
20
18
16
14
12
10
8
6
4
2
0

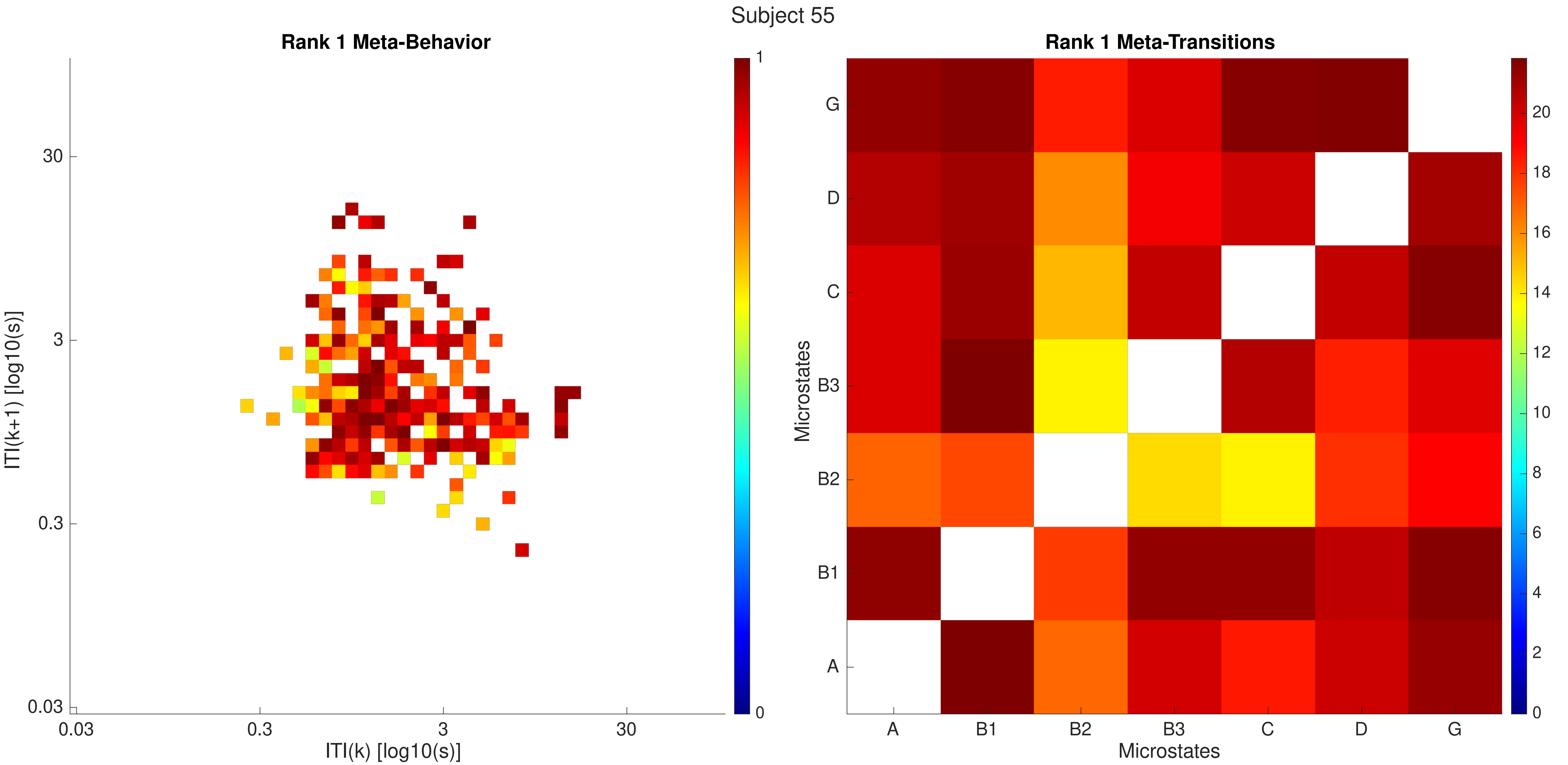

Subject 55
Rank 1 Meta-Behavior
ITI(k+1) [log10(s)]
30
3
0.3
0.03
0.03
0.3
3
30
ITI(k) [log10(s)]
1
0
Rank 1 Meta-Transitions
Microstates
G
D
C
B3
B2
B1
A
A
B1
B2
B3
C
D
G
Microstates
20
18
16
14
12
10
8
6
4
2
0

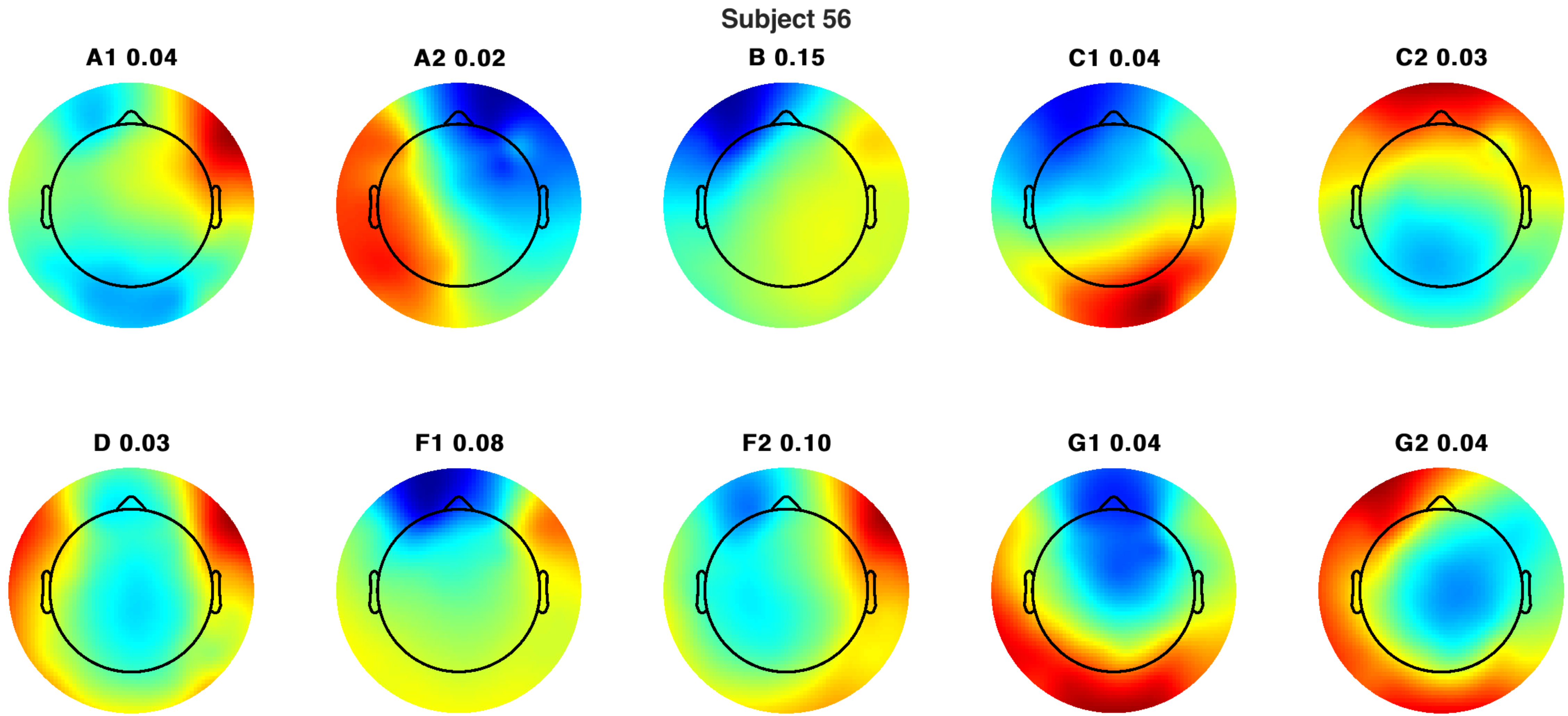
Subject 56
A1 0.04
A2 0.02
B 0.15
C1 0.04
C2 0.03
D 0.03
F1 0.08
F2 0.10
G1 0.04
G2 0.04

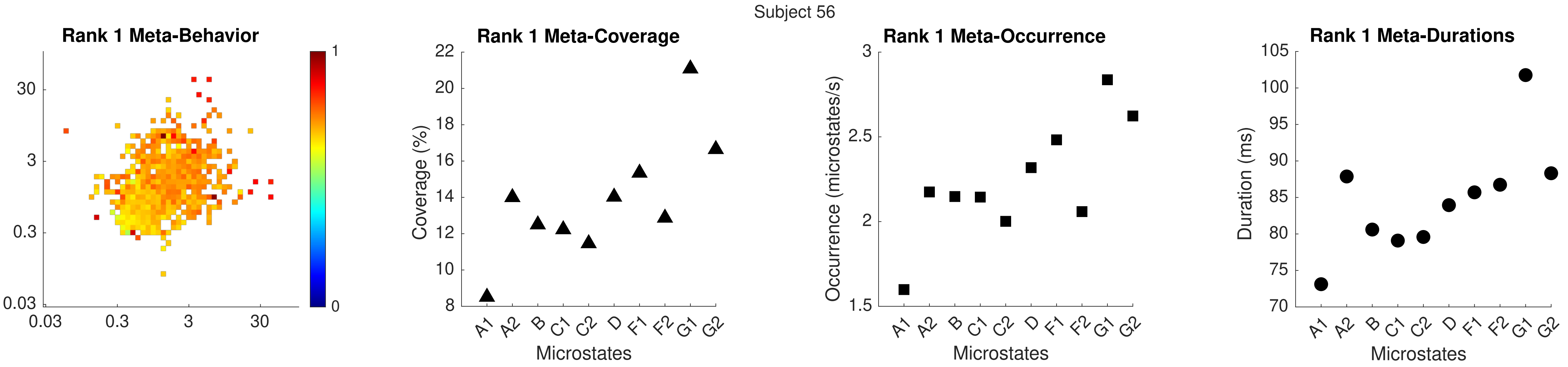
Subject 56
Rank 1 Meta-Behavior
Rank 1 Meta-Coverage
Rank 1 Meta-Occurrence
Rank 1 Meta-Durations
Coverage (%)
Occurrence (microstates/s)
Duration (ms)
Microstates
A1 A2 B C1 C2 D F1 F2 G1 G2

Subject 56

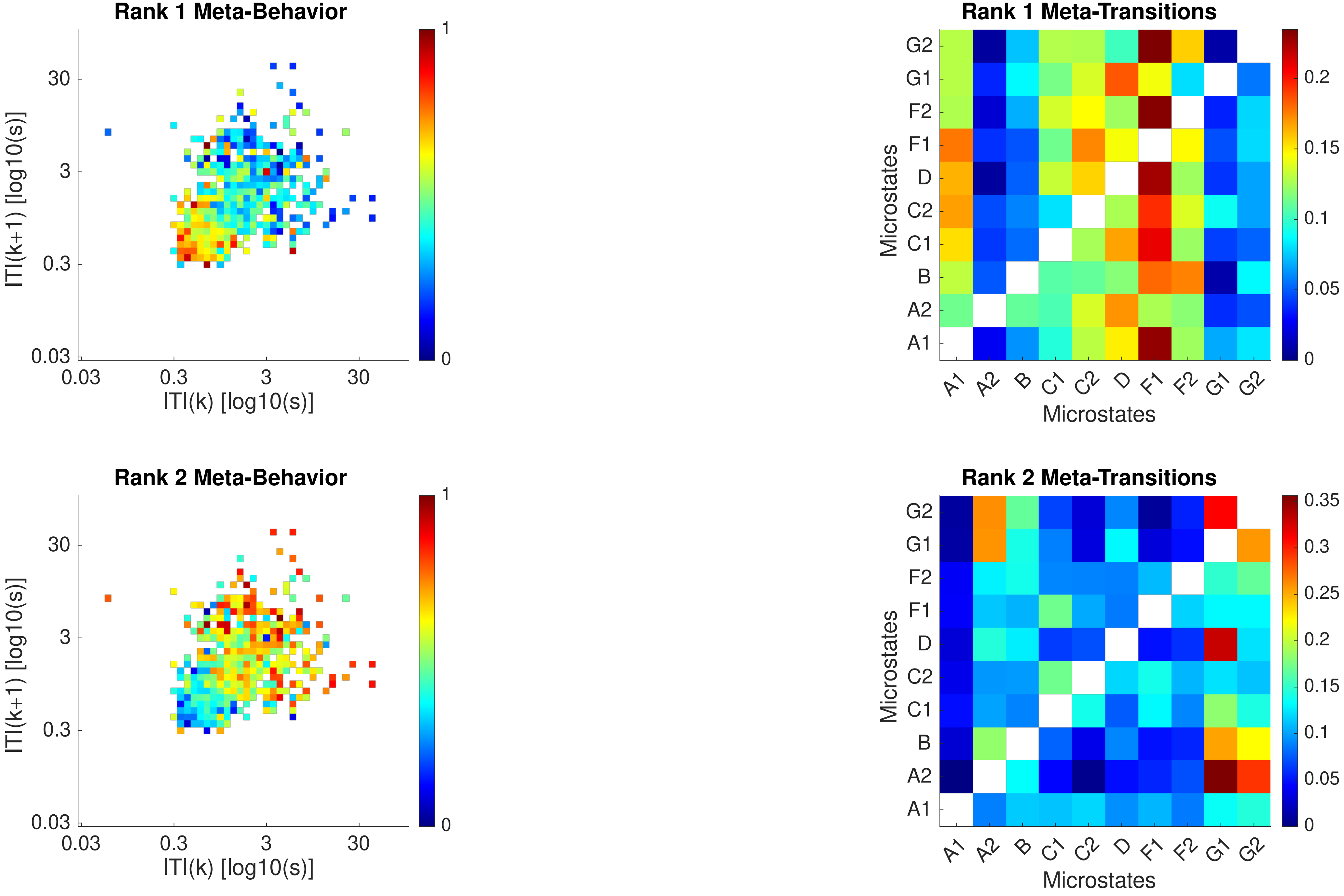

Rank 1 Meta-Behavior
ITI(k+1) [log10(s)]
ITI(k) [log10(s)]
0.03
0.3
3
30
0
1
Rank 1 Meta-Transitions
Microstates
A1
A2
B
C1
C2
D
F1
F2
G1
G2
0
0.05
0.1
0.15
0.2
Rank 2 Meta-Behavior
ITI(k+1) [log10(s)]
ITI(k) [log10(s)]
Rank 2 Meta-Transitions
Microstates
0.25
0.3
0.35

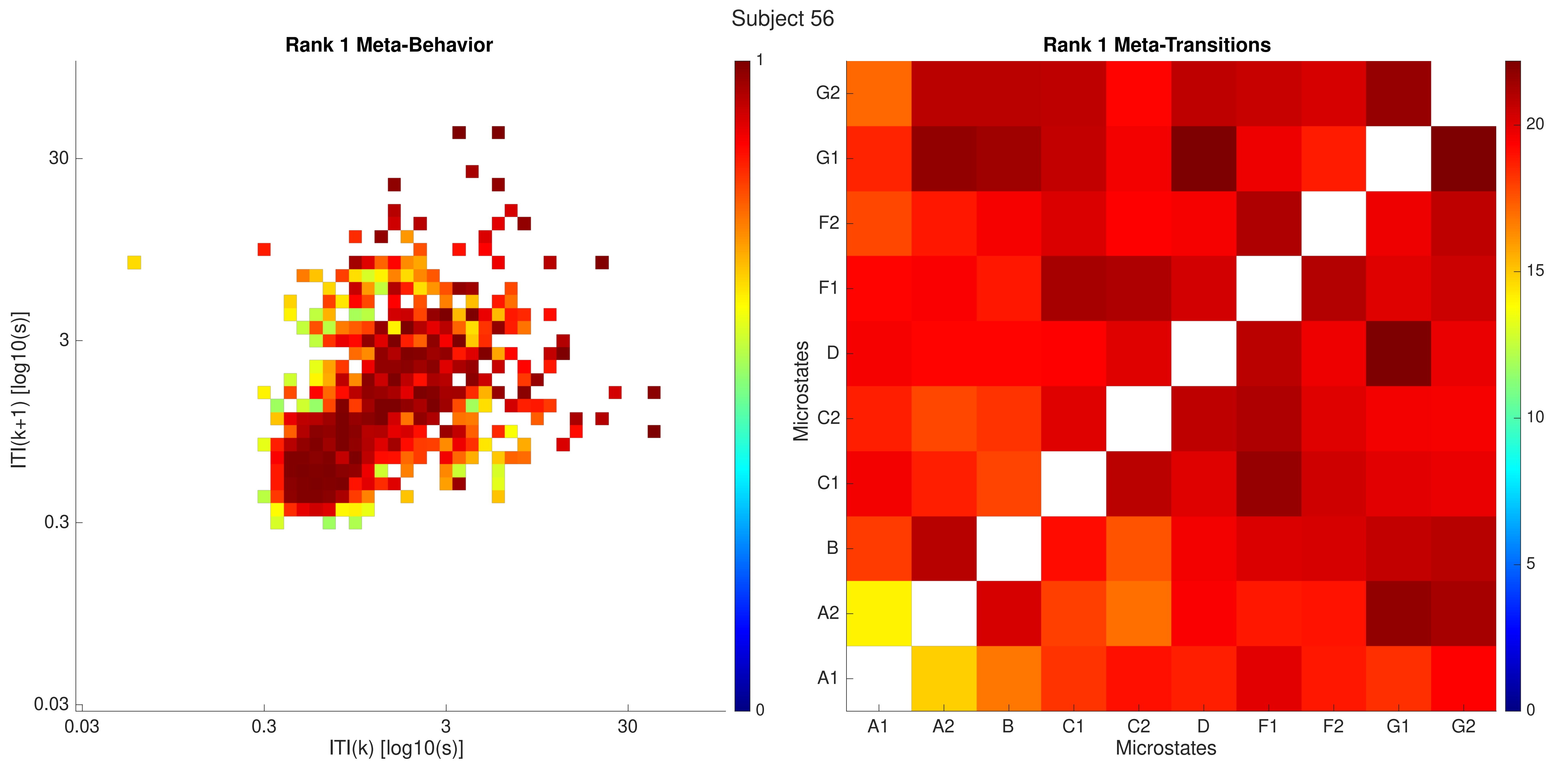
Subject 56
Rank 1 Meta-Behavior
ITI(k+1) [log10(s)]
ITI(k) [log10(s)]
0.03
0.3
3
30
0
1
Rank 1 Meta-Transitions
Microstates
A1
A2
B
C1
C2
D
F1
F2
G1
G2
0
5
10
15
20

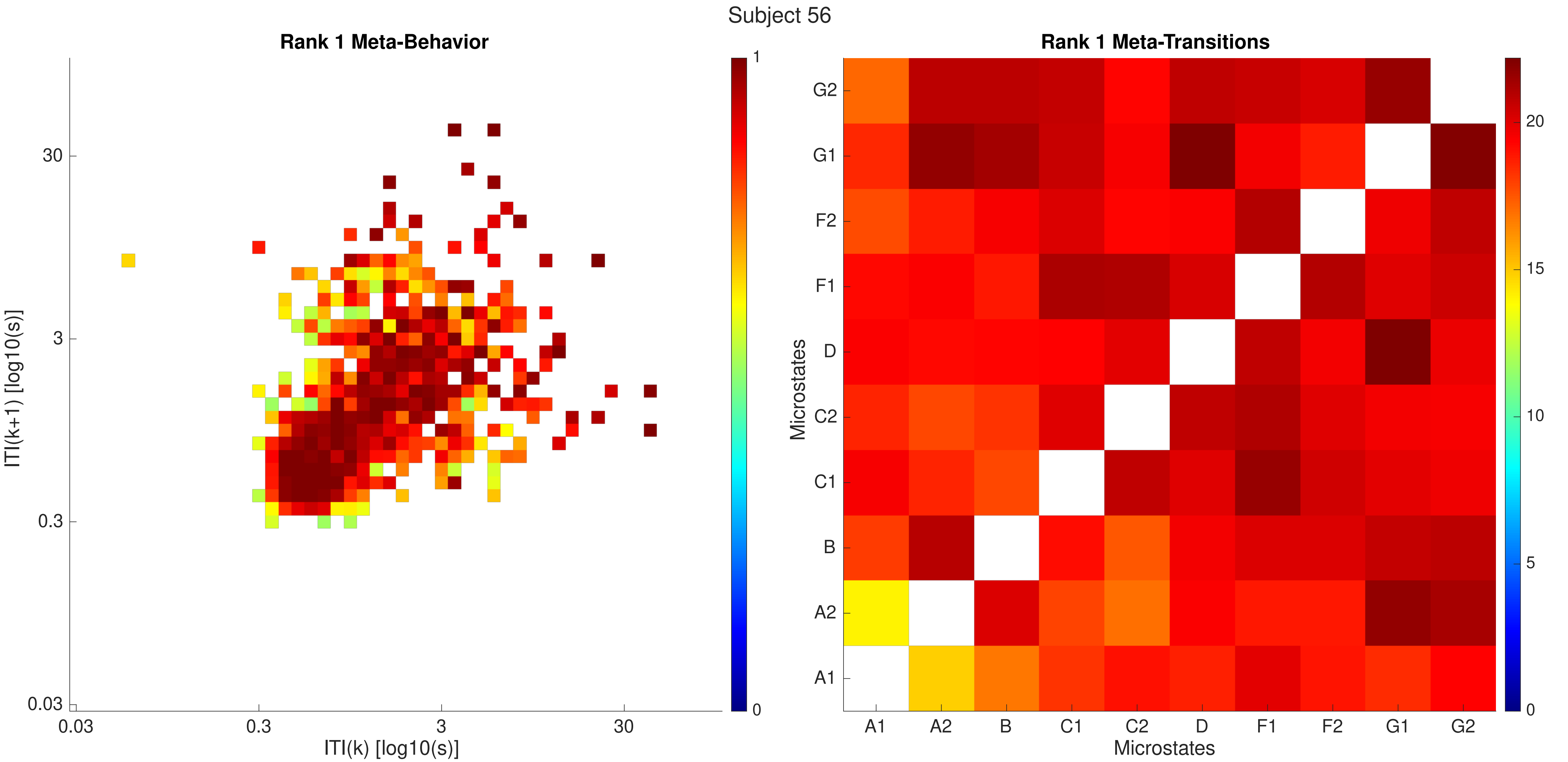

Subject 56
Rank 1 Meta-Behavior
ITI(k+1) [log10(s)]
ITI(k) [log10(s)]
0.03
0.3
3
30
1
0
Rank 1 Meta-Transitions
Microstates
A1
A2
B
C1
C2
D
F1
F2
G1
G2
0
5
10
15
20

Subject 57

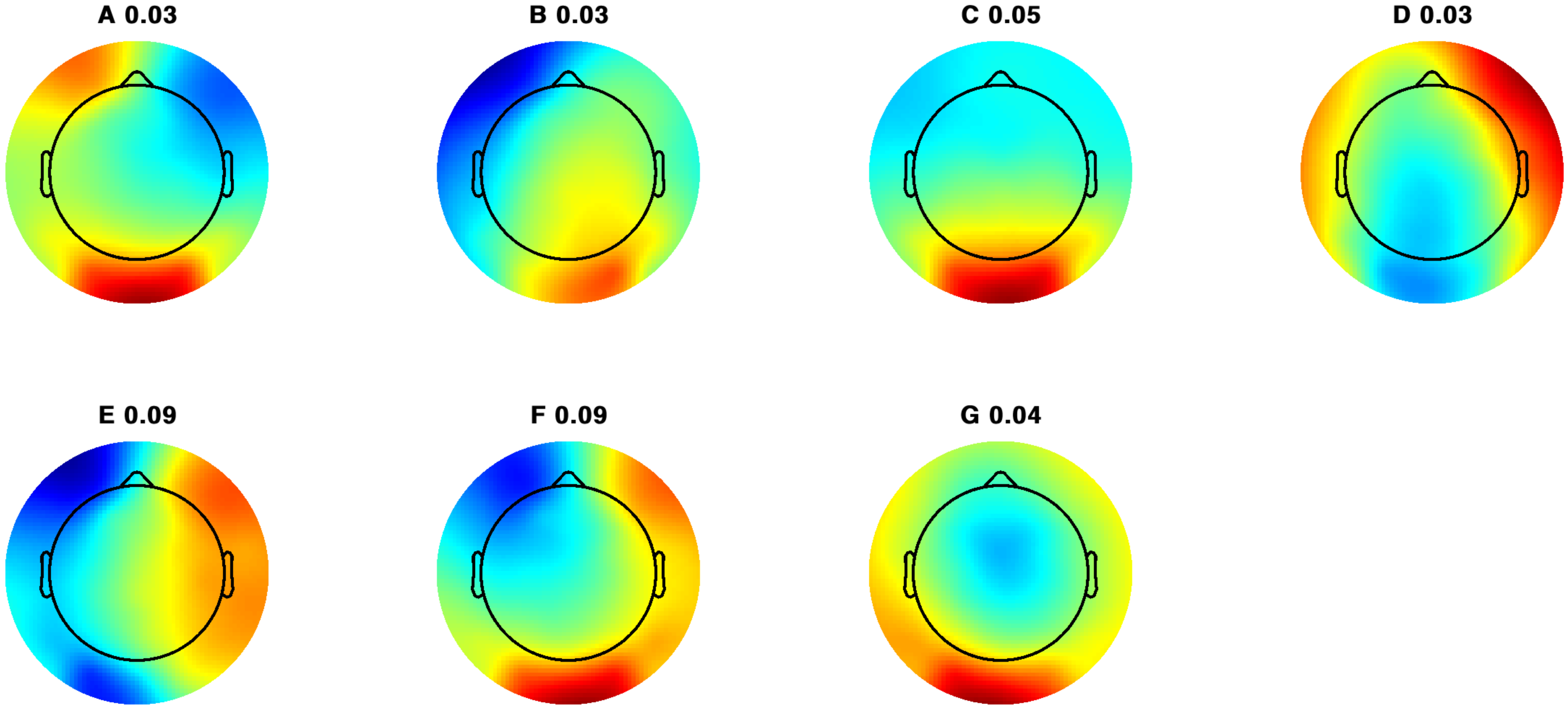

Subject 57

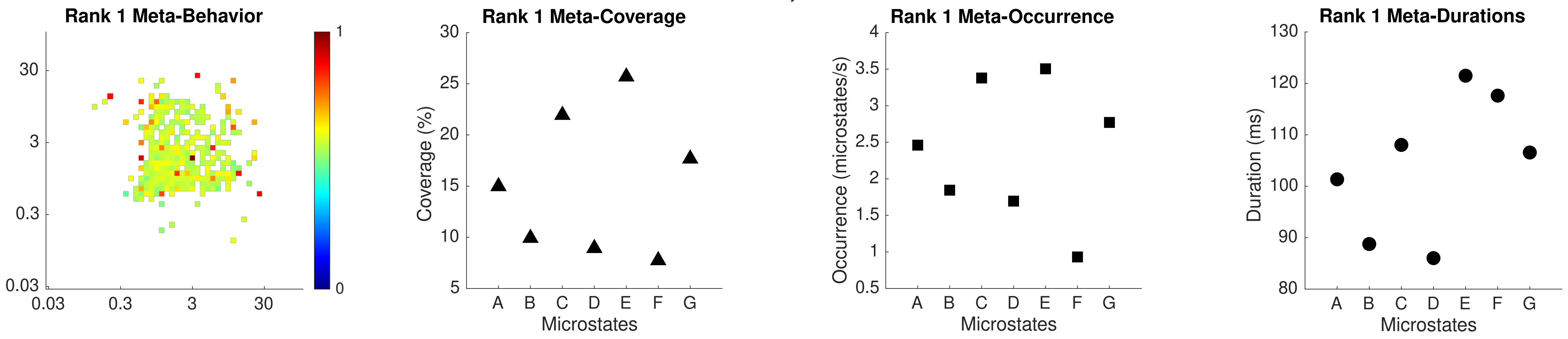

Rank 1 Meta-Behavior
30
3
0.3
0.03
0.03
0.3
3
30
1
0
Rank 1 Meta-Coverage
Coverage (%)
Microstates
A B C D E F G
Rank 1 Meta-Occurrence
Occurrence (microstates/s)
Microstates
Rank 1 Meta-Durations
Duration (ms)
Microstates

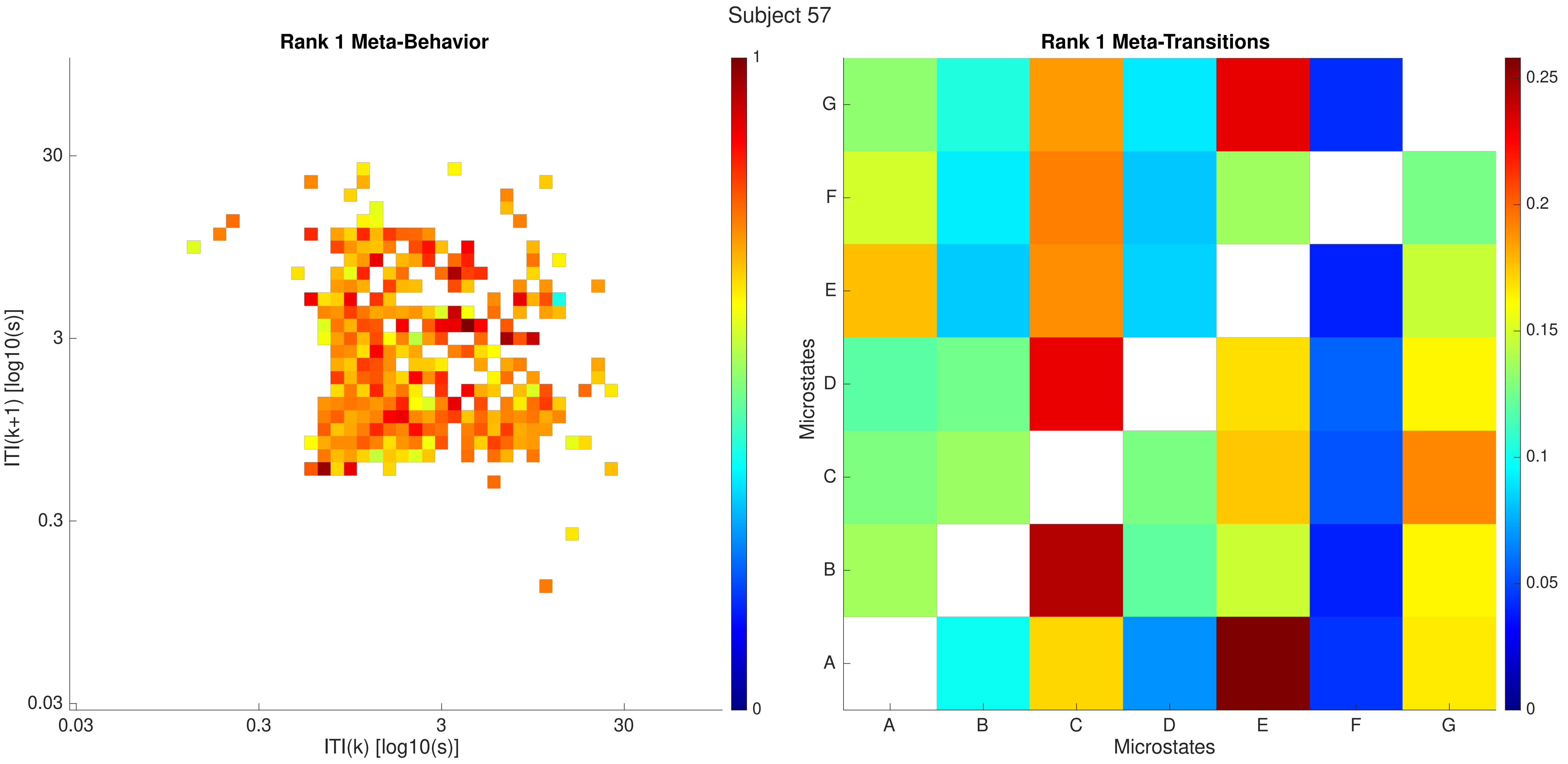
Subject 57
Rank 1 Meta-Behavior
ITI(k+1) [log10(s)]
30
3
0.3
0.03
ITI(k) [log10(s)]
1
0
Rank 1 Meta-Transitions
Microstates
G
F
E
D
C
B
A
0.25
0.2
0.15
0.1
0.05

Subject 57

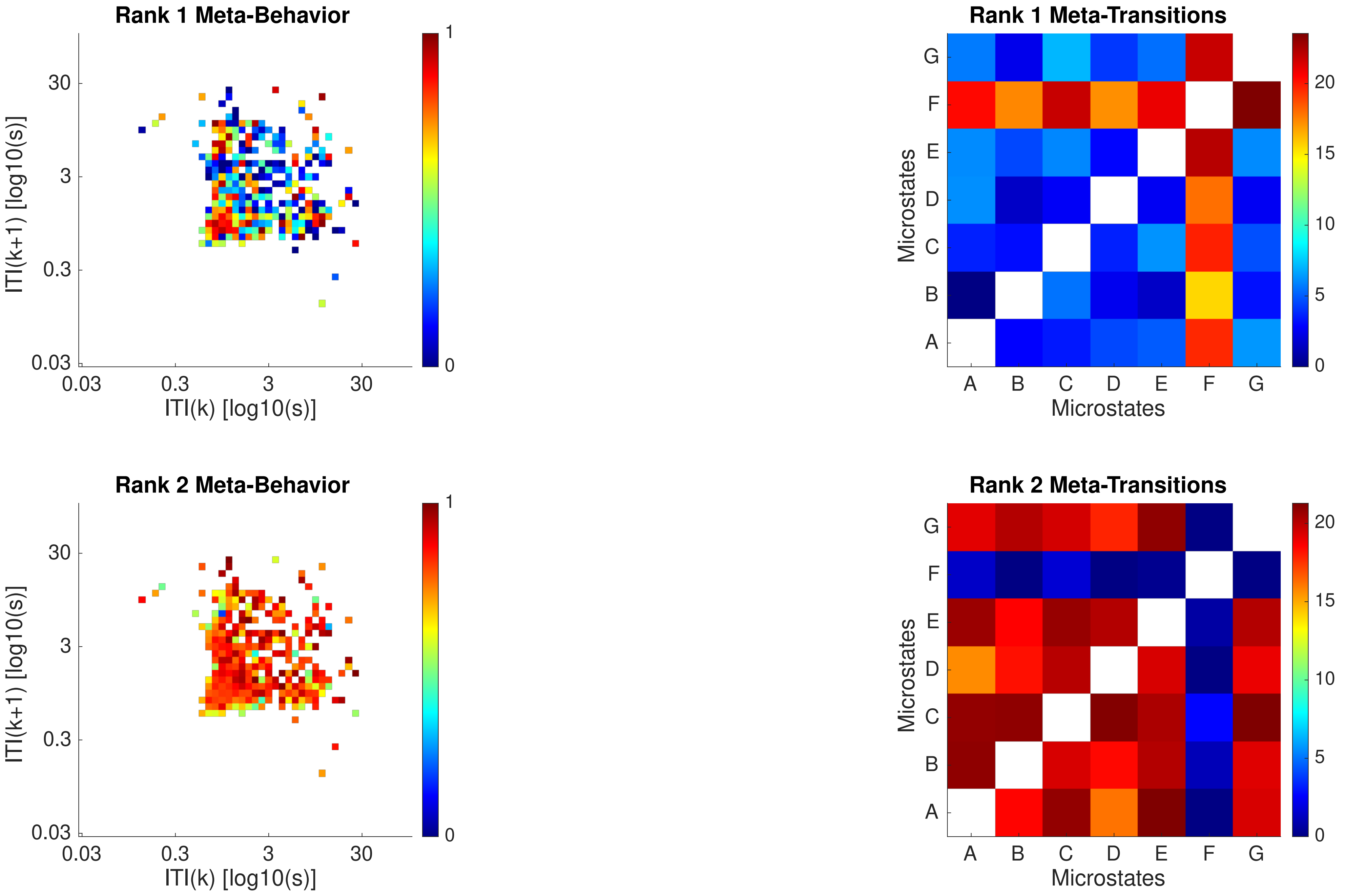
Rank 1 Meta-Behavior
ITI(k+1) [log10(s)]
ITI(k) [log10(s)]
0.03
0.3
3
30
Rank 1 Meta-Transitions
Microstates
A
B
C
D
E
F
G
Rank 2 Meta-Behavior
Rank 2 Meta-Transitions
0
5
10
15
20
1

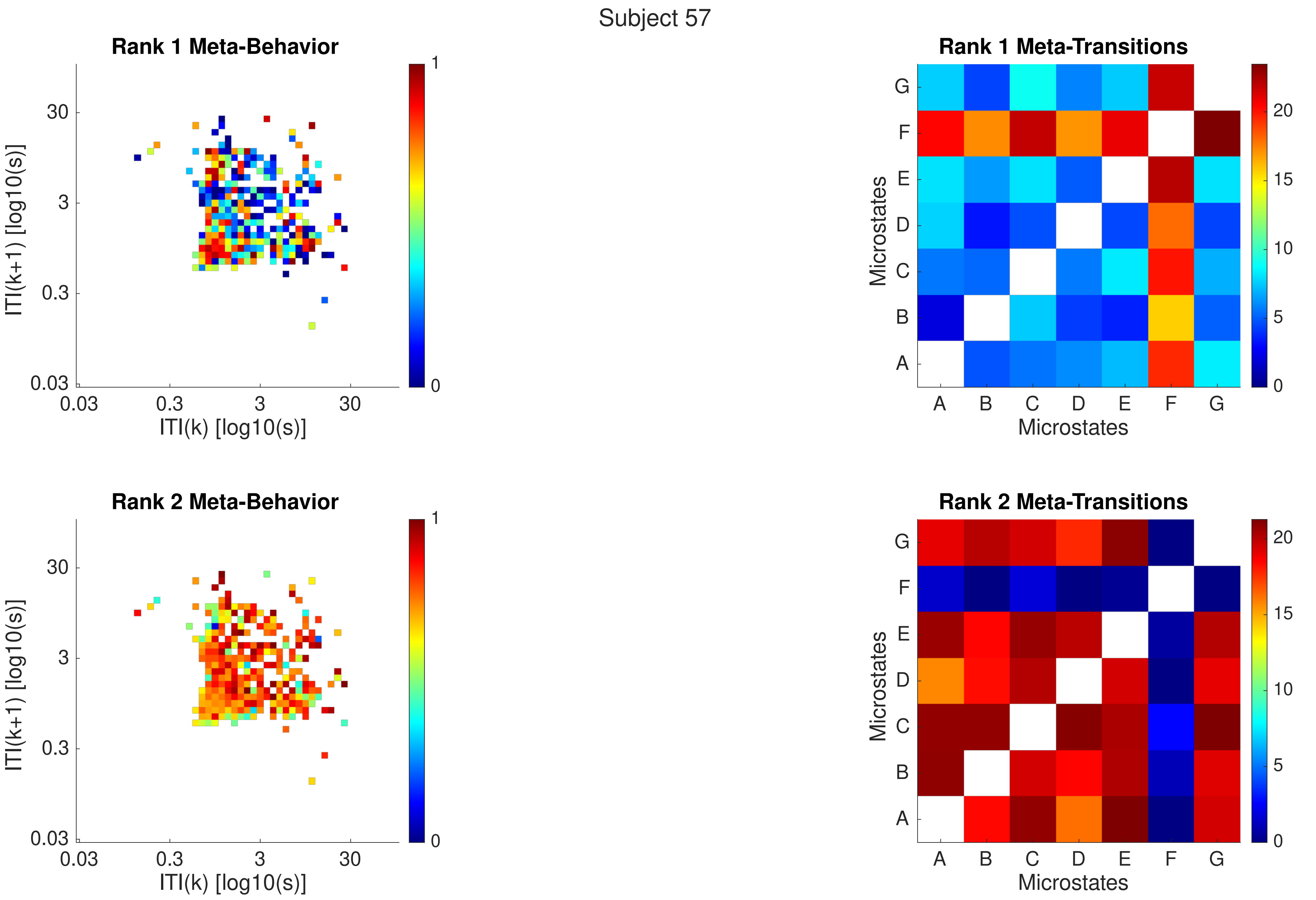

Subject 57
Rank 1 Meta-Behavior
ITI(k+1) [log10(s)]
ITI(k) [log10(s)]
0.03
0.3
3
30
0
1
Rank 1 Meta-Transitions
Microstates
A
B
C
D
E
F
G
0
5
10
15
20
Rank 2 Meta-Behavior
ITI(k+1) [log10(s)]
ITI(k) [log10(s)]
Rank 2 Meta-Transitions
Microstates

Subject 58

C 0.08

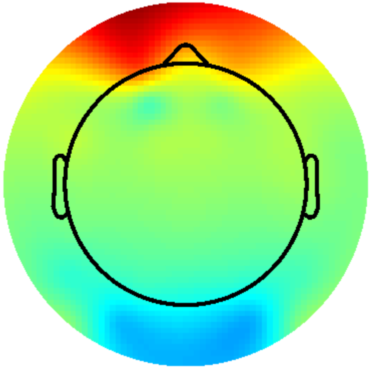

G 0.03

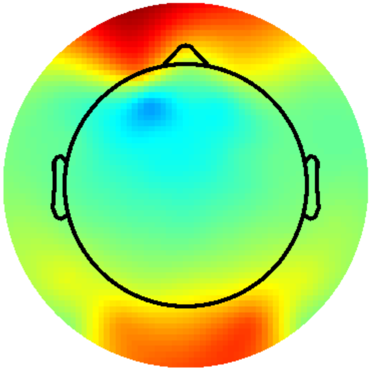

Subject 58

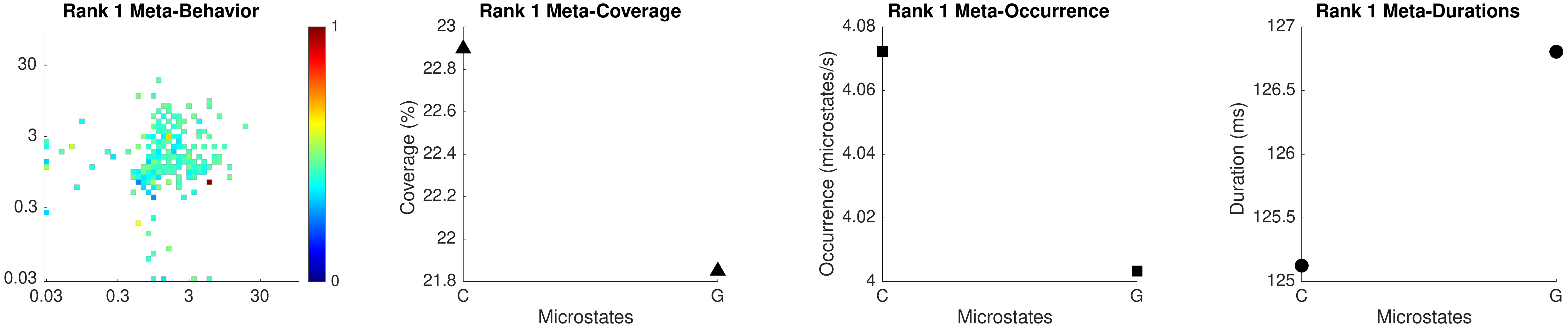
Rank 1 Meta-Behavior
30
3
0.3
0.03
0.03
0.3
3
30
1
0
Rank 1 Meta-Coverage
Coverage (%)
23
22.8
22.6
22.4
22.2
22
21.8
C
G
Microstates
Rank 1 Meta-Occurrence
Occurrence (microstates/s)
4.08
4.06
4.04
4.02
4
C
G
Microstates
Rank 1 Meta-Durations
Duration (ms)
127
126.5
126
125.5
125
C
G
Microstates

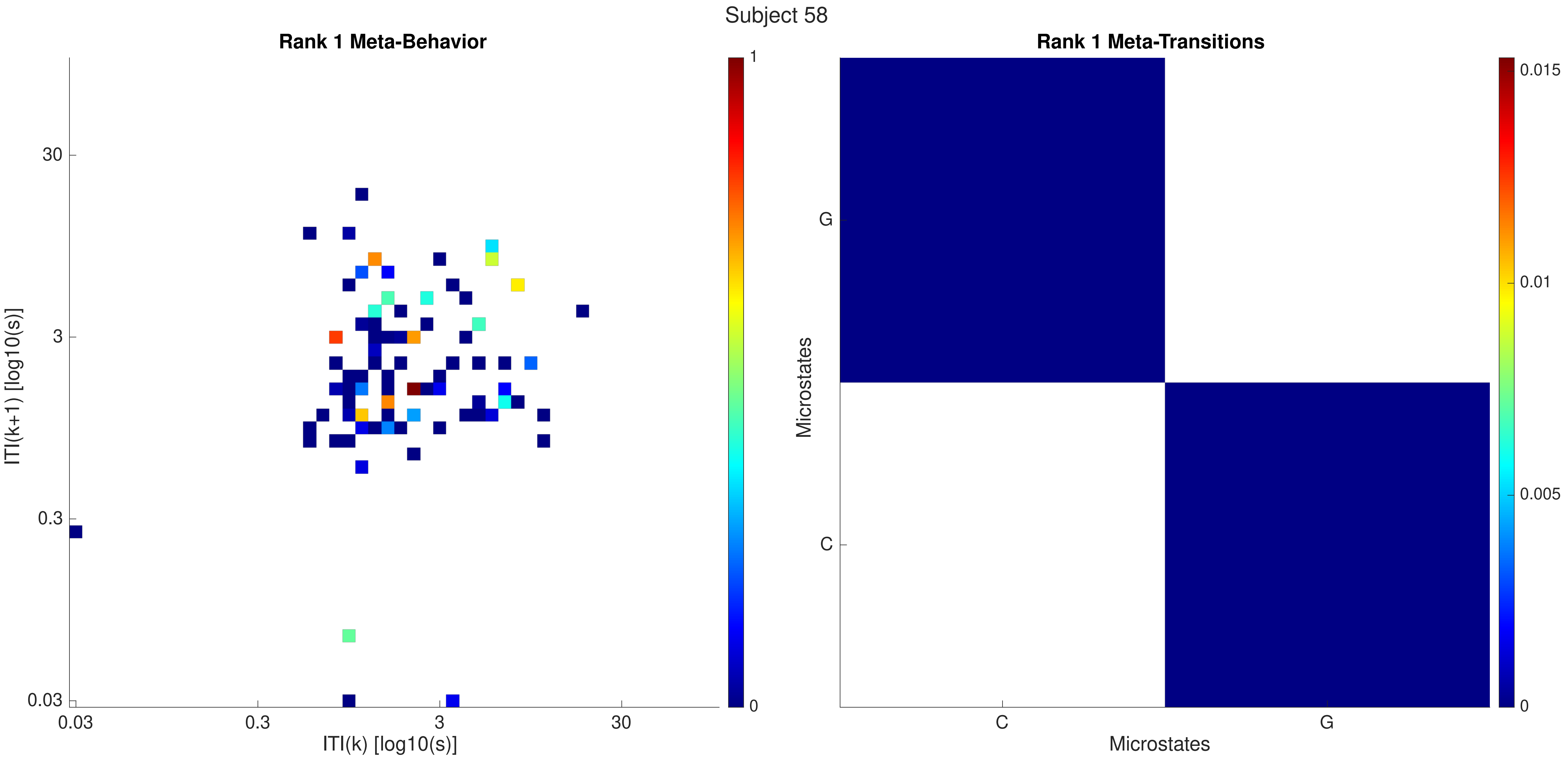

Subject 58
Rank 1 Meta-Behavior
Rank 1 Meta-Transitions
ITI(k+1) [log10(s)]
ITI(k) [log10(s)]
0.03
0.3
3
30
1
0
Microstates
C
G
0.015
0.01
0.005

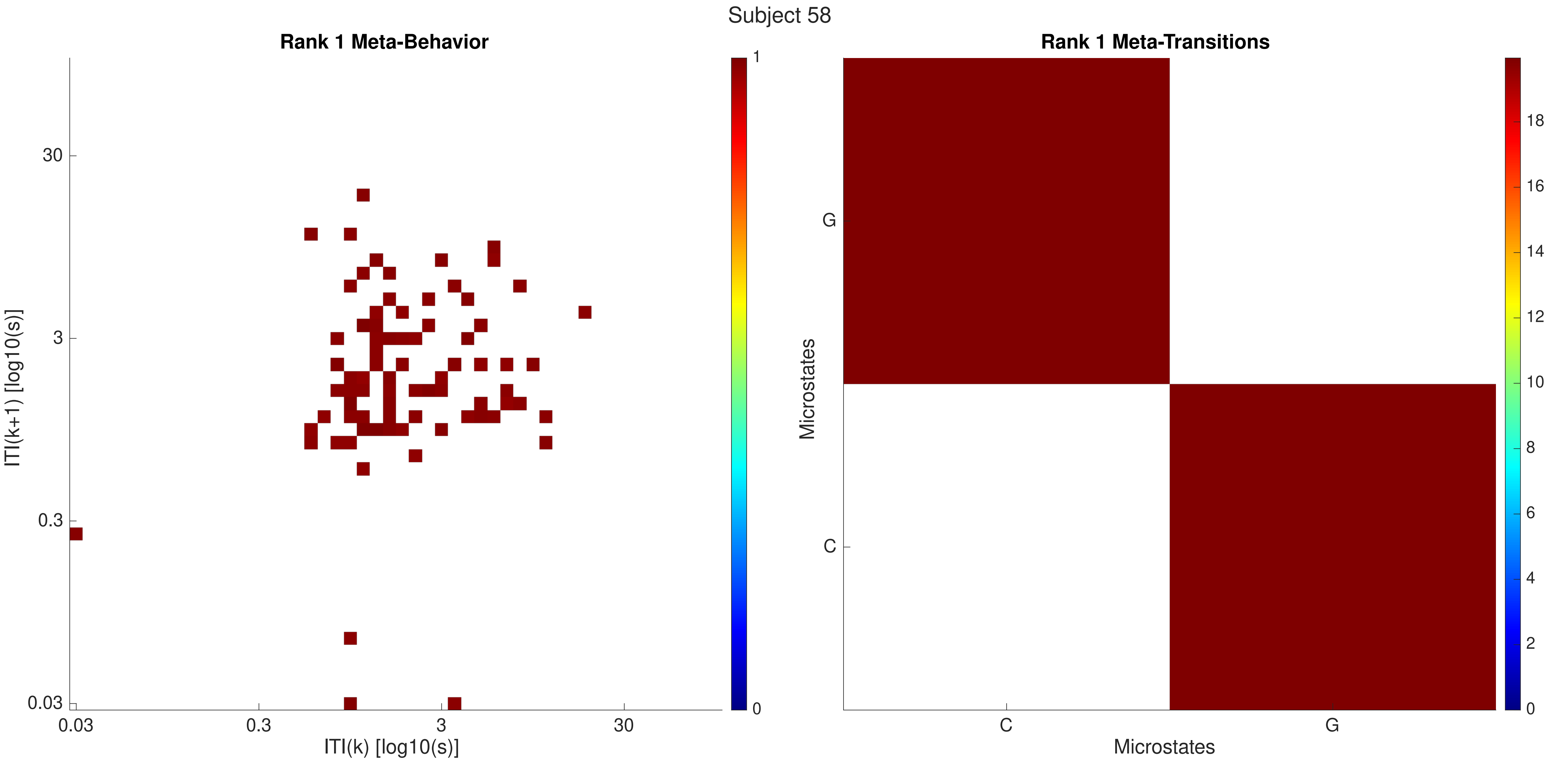

Subject 58
Rank 1 Meta-Behavior
ITI(k+1) [log10(s)]
30
3
0.3
0.03
ITI(k) [log10(s)]
0.03
0.3
3
30
1
0
Rank 1 Meta-Transitions
Microstates
G
C
Microstates
C
G
18
16
14
12
10
8
6
4
2
0

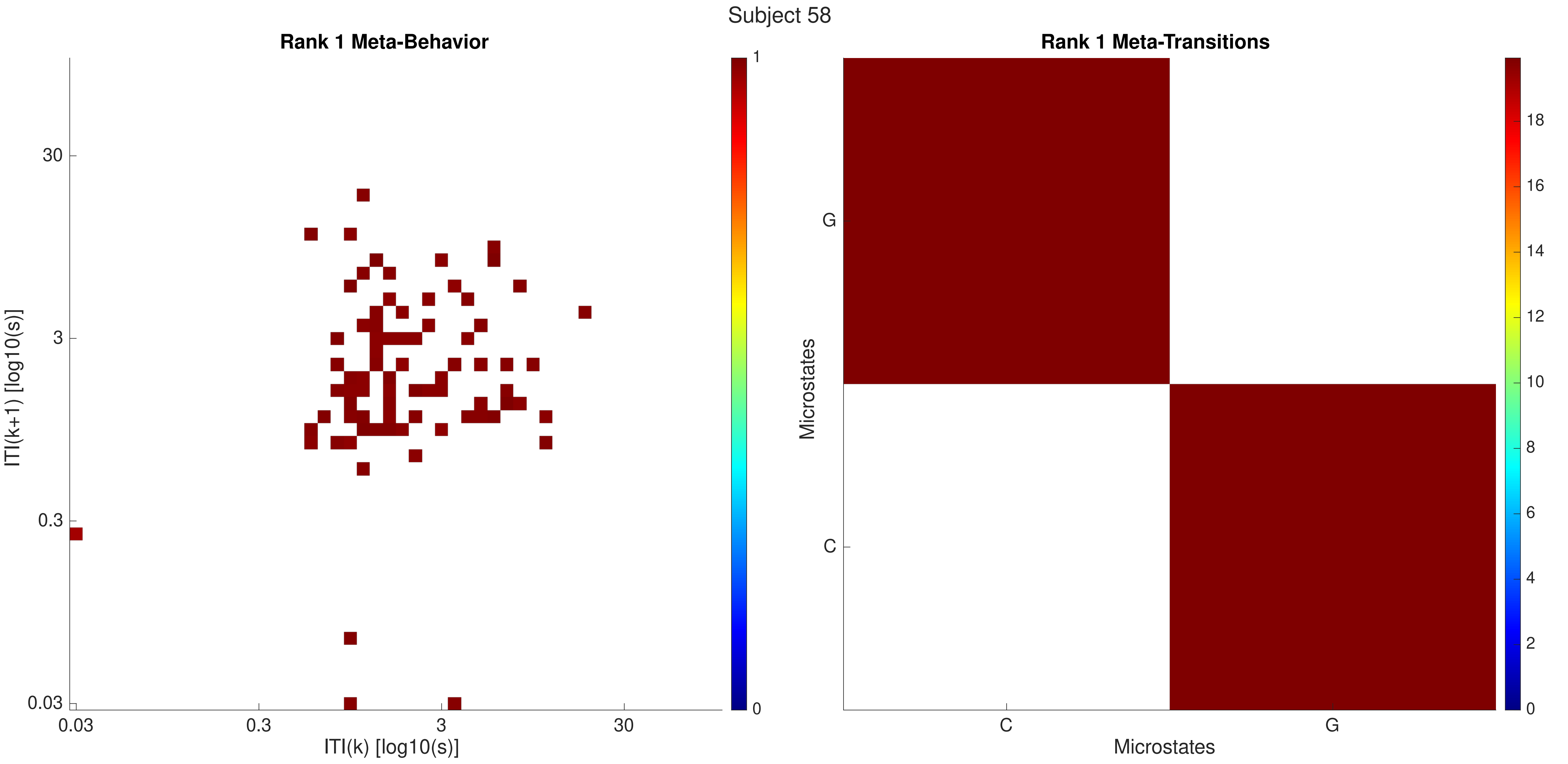
Subject 58
Rank 1 Meta-Behavior
ITI(k+1) [log10(s)]
30
3
0.3
0.03
0.03
0.3
3
30
ITI(k) [log10(s)]
1
0
Rank 1 Meta-Transitions
Microstates
G
C
C
G
Microstates
18
16
14
12
10
8
6
4
2
0

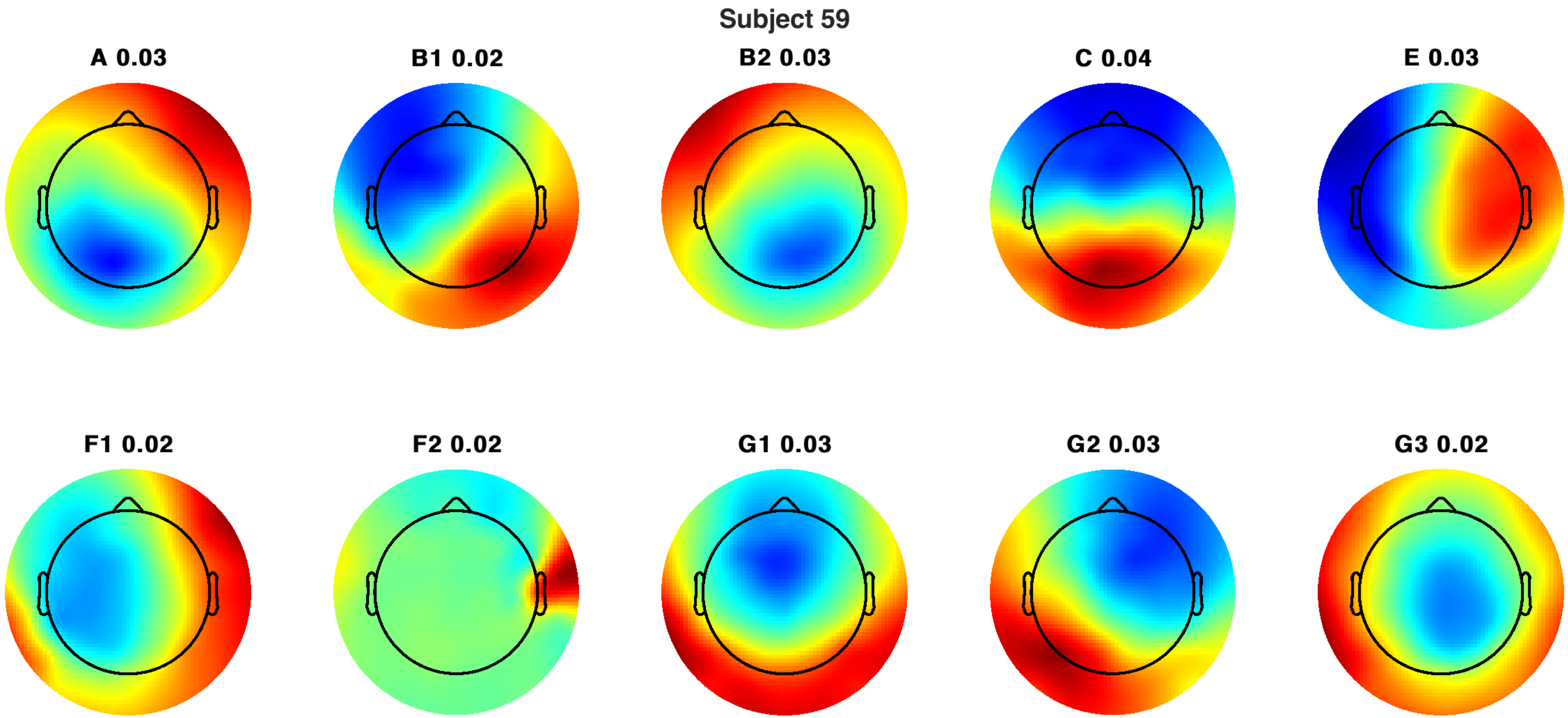
Subject 59
A 0.03
B1 0.02
B2 0.03
C 0.04
E 0.03
F1 0.02
F2 0.02
G1 0.03
G2 0.03
G3 0.02

Subject 59

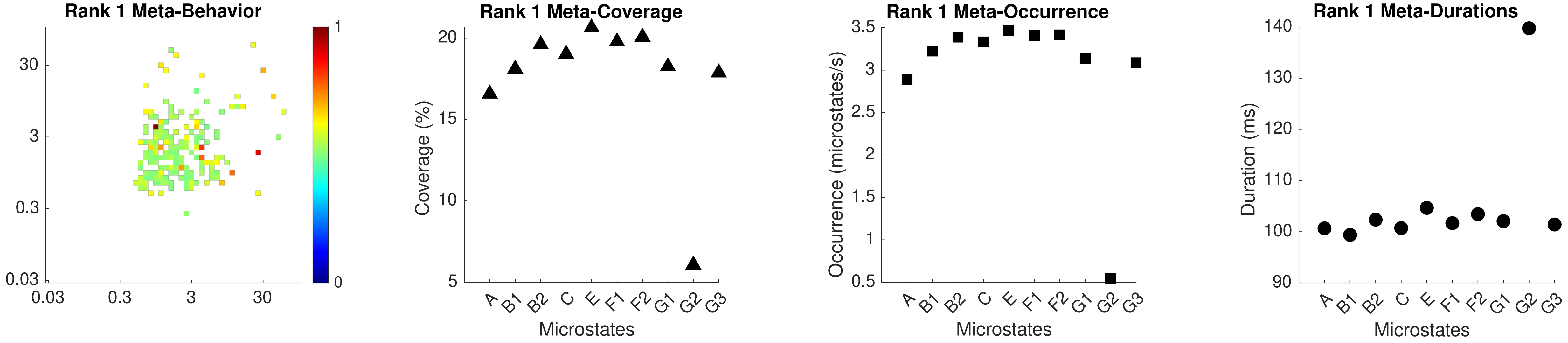

Rank 1 Meta-Behavior
30
3
0.3
0.03
0.03
0.3
3
30
1
0
Rank 1 Meta-Coverage
Coverage (%)
20
15
10
5
A
B1
B2
C
E
F1
F2
G1
G2
G3
Microstates
Rank 1 Meta-Occurrence
Occurrence (microstates/s)
3.5
3
2.5
2
1.5
1
0.5
A
B1
B2
C
E
F1
F2
G1
G2
G3
Microstates
Rank 1 Meta-Durations
Duration (ms)
140
130
120
110
100
90
A
B1
B2
C
E
F1
F2
G1
G2
G3
Microstates

Subject 59

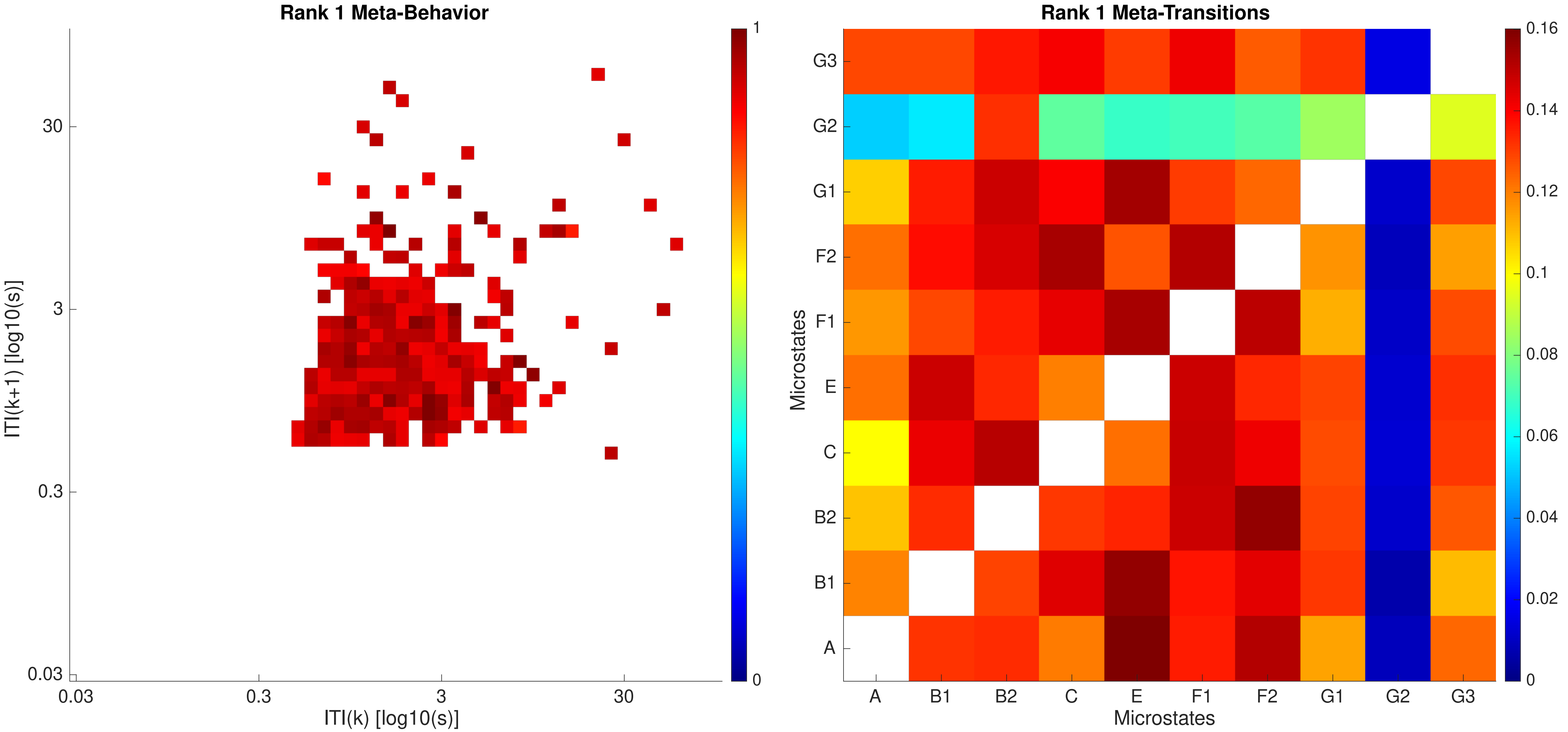

Rank 1 Meta-Behavior
ITI(k+1) [log10(s)]
ITI(k) [log10(s)]
0.03
0.3
3
30
0
1
Rank 1 Meta-Transitions
Microstates
A
B1
B2
C
E
F1
F2
G1
G2
G3
0
0.02
0.04
0.06
0.08
0.1
0.12
0.14
0.16

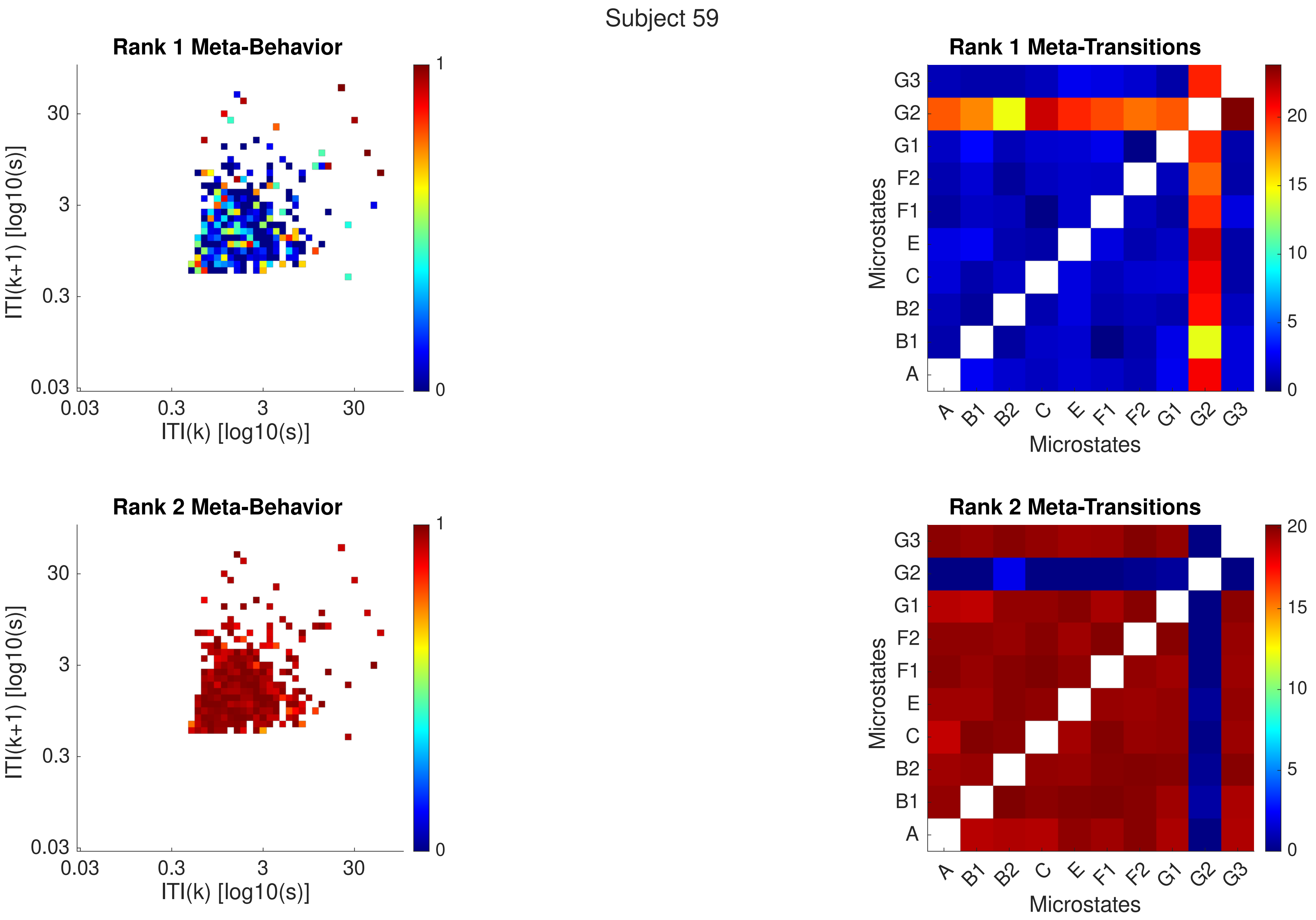
Subject 59
Rank 1 Meta-Behavior
Rank 1 Meta-Transitions
Rank 2 Meta-Behavior
Rank 2 Meta-Transitions
ITI(k+1) [log10(s)]
ITI(k) [log10(s)]
0.03
0.3
3
30
0
1
Microstates
A
B1
B2
C
E
F1
F2
G1
G2
G3
0
5
10
15
20

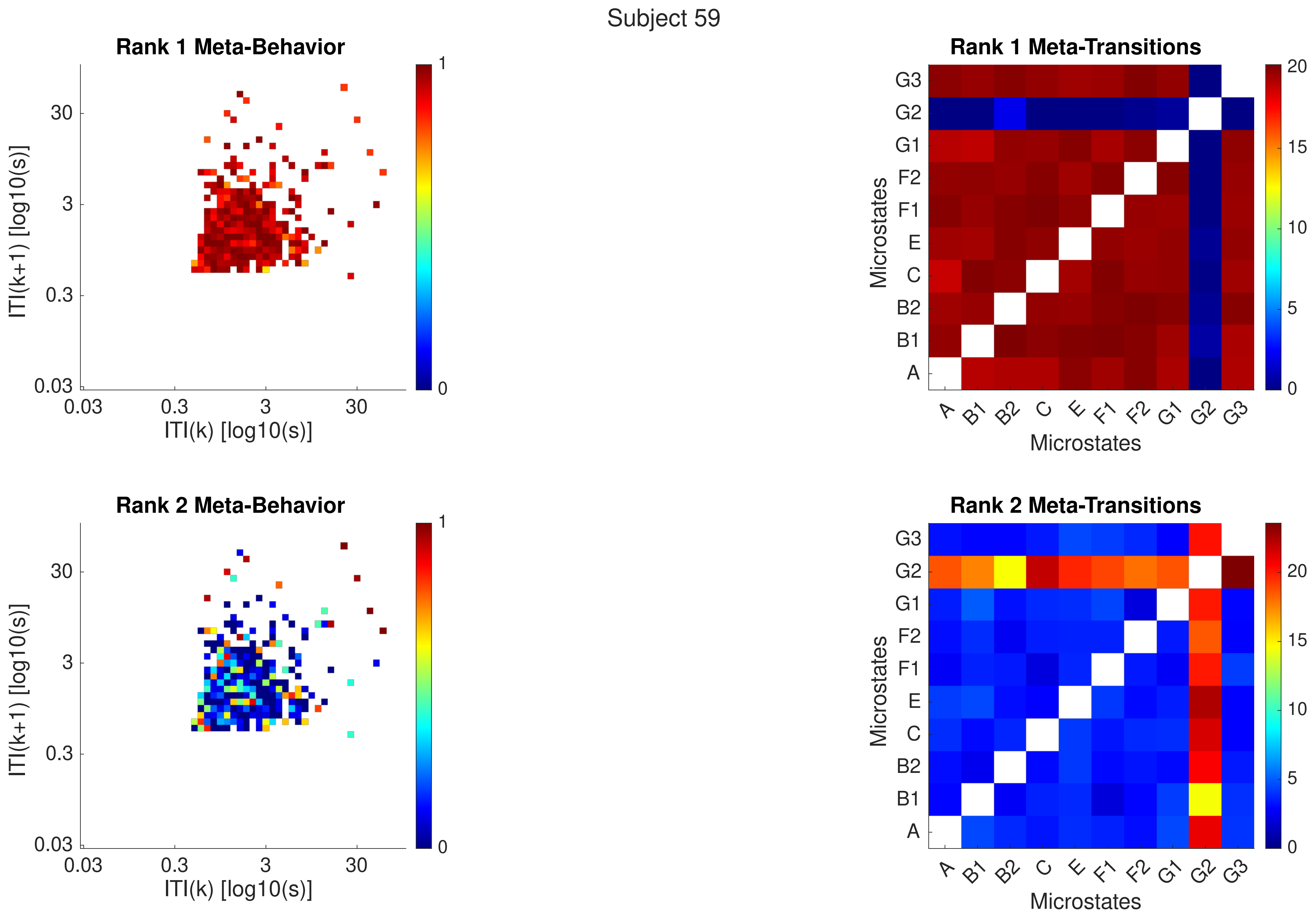
Subject 59
Rank 1 Meta-Behavior
ITI(k+1) [log10(s)]
ITI(k) [log10(s)]
0.03
0.3
3
30
0
1
Rank 1 Meta-Transitions
Microstates
A
B1
B2
C
E
F1
F2
G1
G2
G3
0
5
10
15
20
Rank 2 Meta-Behavior
Rank 2 Meta-Transitions

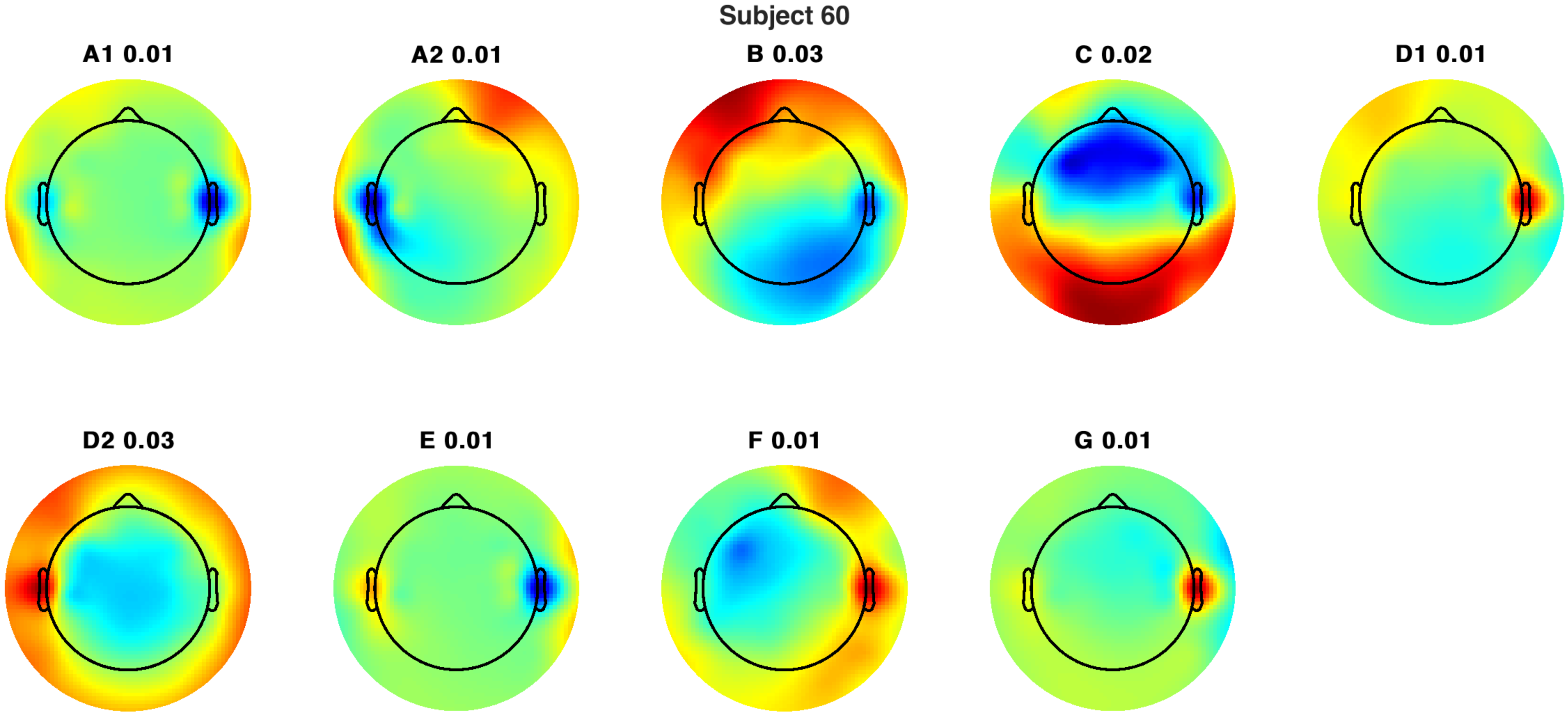

Subject 60
A1 0.01
A2 0.01
B 0.03
C 0.02
D1 0.01
D2 0.03
E 0.01
F 0.01
G 0.01

Subject 60

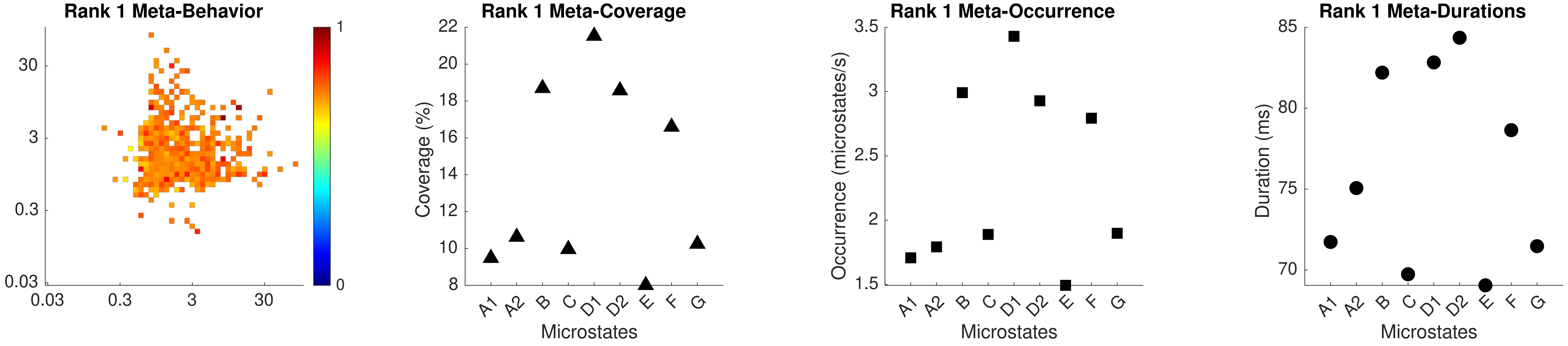

Rank 1 Meta-Behavior
30
3
0.3
0.03
0.03
0.3
3
30
1
0
Rank 1 Meta-Coverage
Coverage (%)
22
20
18
16
14
12
10
8
A1 A2 B C D1 D2 E F G
Microstates
Rank 1 Meta-Occurrence
Occurrence (microstates/s)
3.5
3
2.5
2
1.5
A1 A2 B C D1 D2 E F G
Microstates
Rank 1 Meta-Durations
Duration (ms)
85
80
75
70
A1 A2 B C D1 D2 E F G
Microstates

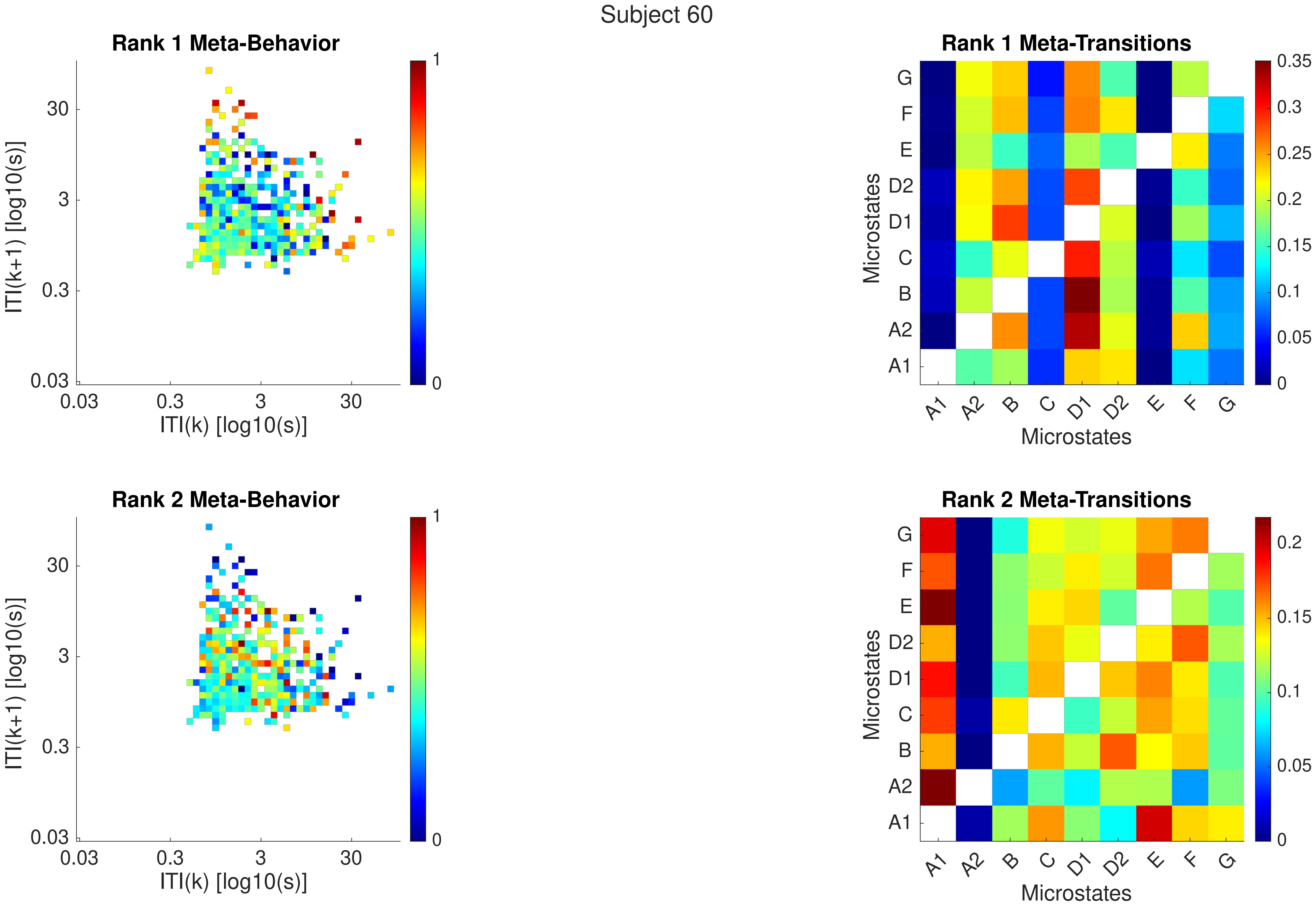
Subject 60
Rank 1 Meta-Behavior
Rank 1 Meta-Transitions
Rank 2 Meta-Behavior
Rank 2 Meta-Transitions
ITI(k+1) [log10(s)]
ITI(k) [log10(s)]
0.03
0.3
3
30
0
1
Microstates
A1
A2
B
C
D1
D2
E
F
G
0
0.05
0.1
0.15
0.2
0.25
0.3
0.35

Subject 60

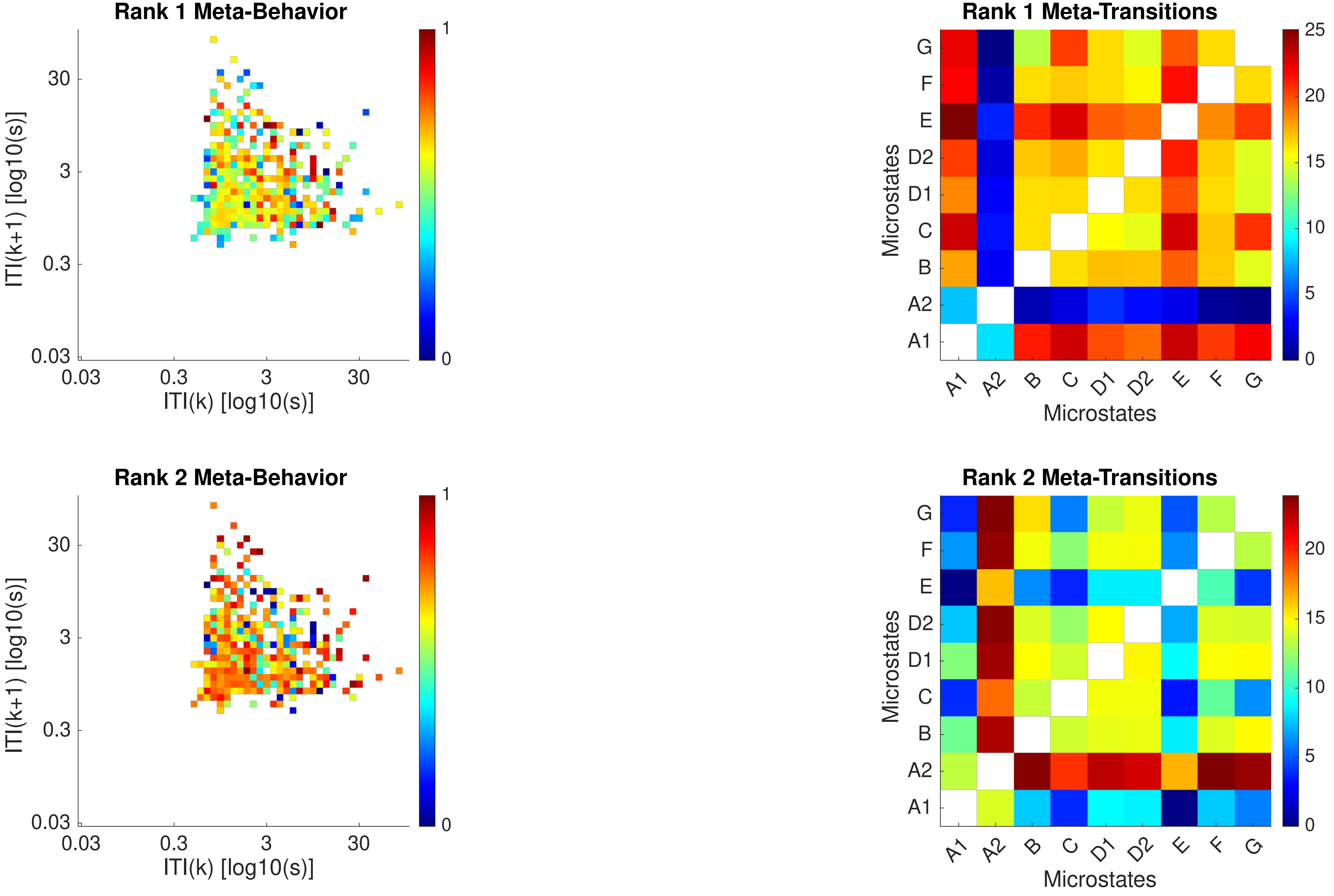

Rank 1 Meta-Behavior
ITI(k+1) [log10(s)]
ITI(k) [log10(s)]
0.03
0.3
3
30
0
1
Rank 1 Meta-Transitions
Microstates
A1
A2
B
C
D1
D2
E
F
G
0
5
10
15
20
25
Rank 2 Meta-Behavior
Rank 2 Meta-Transitions

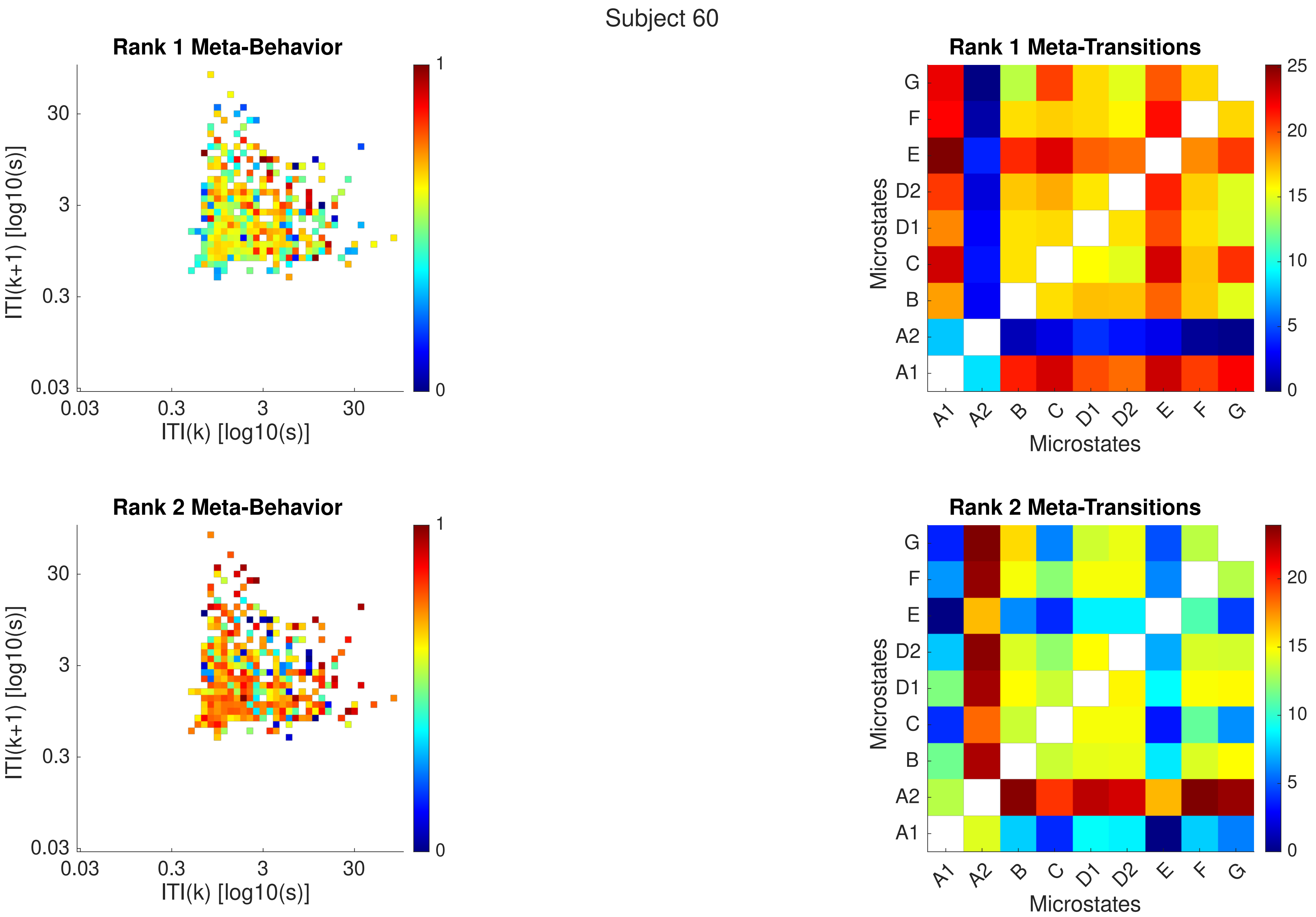

Subject 60
Rank 1 Meta-Behavior
ITI(k+1) [log10(s)]
ITI(k) [log10(s)]
30
3
0.3
0.03
1
0
Rank 1 Meta-Transitions
Microstates
A1
A2
B
C
D1
D2
E
F
G
25
20
15
10
5
0
Rank 2 Meta-Behavior
Rank 2 Meta-Transitions

Subject 61

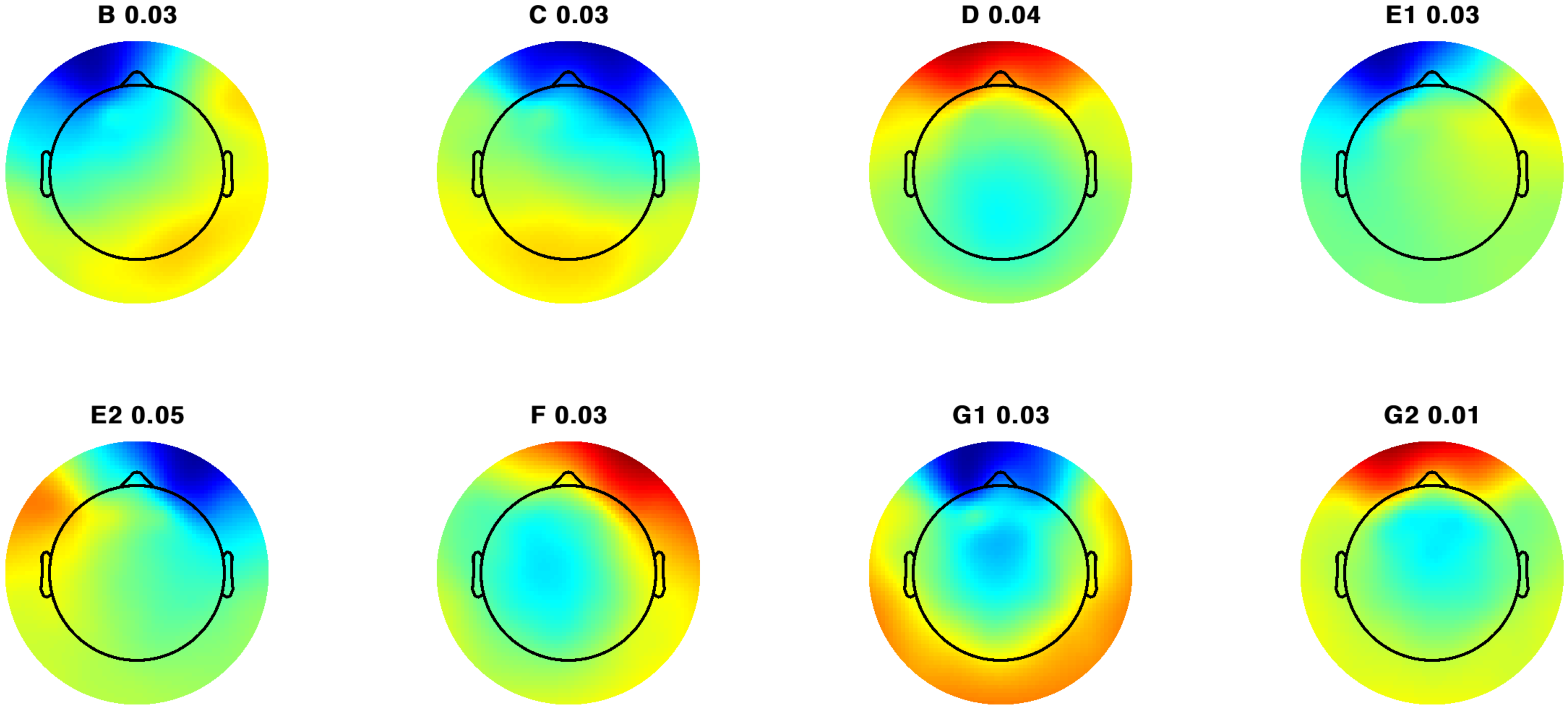

Subject 61

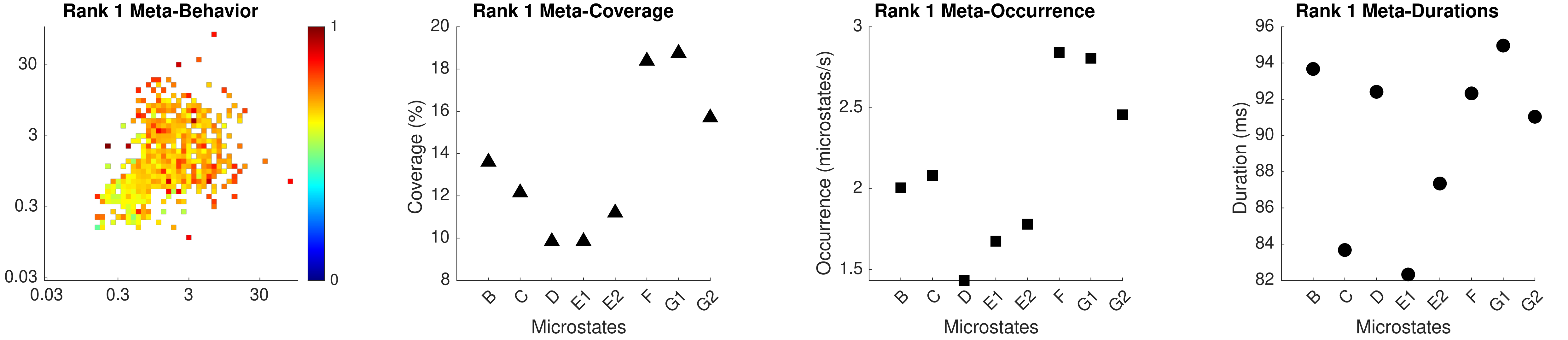

Rank 1 Meta-Behavior
30
3
0.3
0.03
0.03
0.3
3
30
1
0
Rank 1 Meta-Coverage
Coverage (%)
20
18
16
14
12
10
8
B
C
D
E1
E2
F
G1
G2
Microstates
Rank 1 Meta-Occurrence
Occurrence (microstates/s)
3
2.5
2
1.5
Microstates
Rank 1 Meta-Durations
Duration (ms)
96
94
92
90
88
86
84
82
Microstates

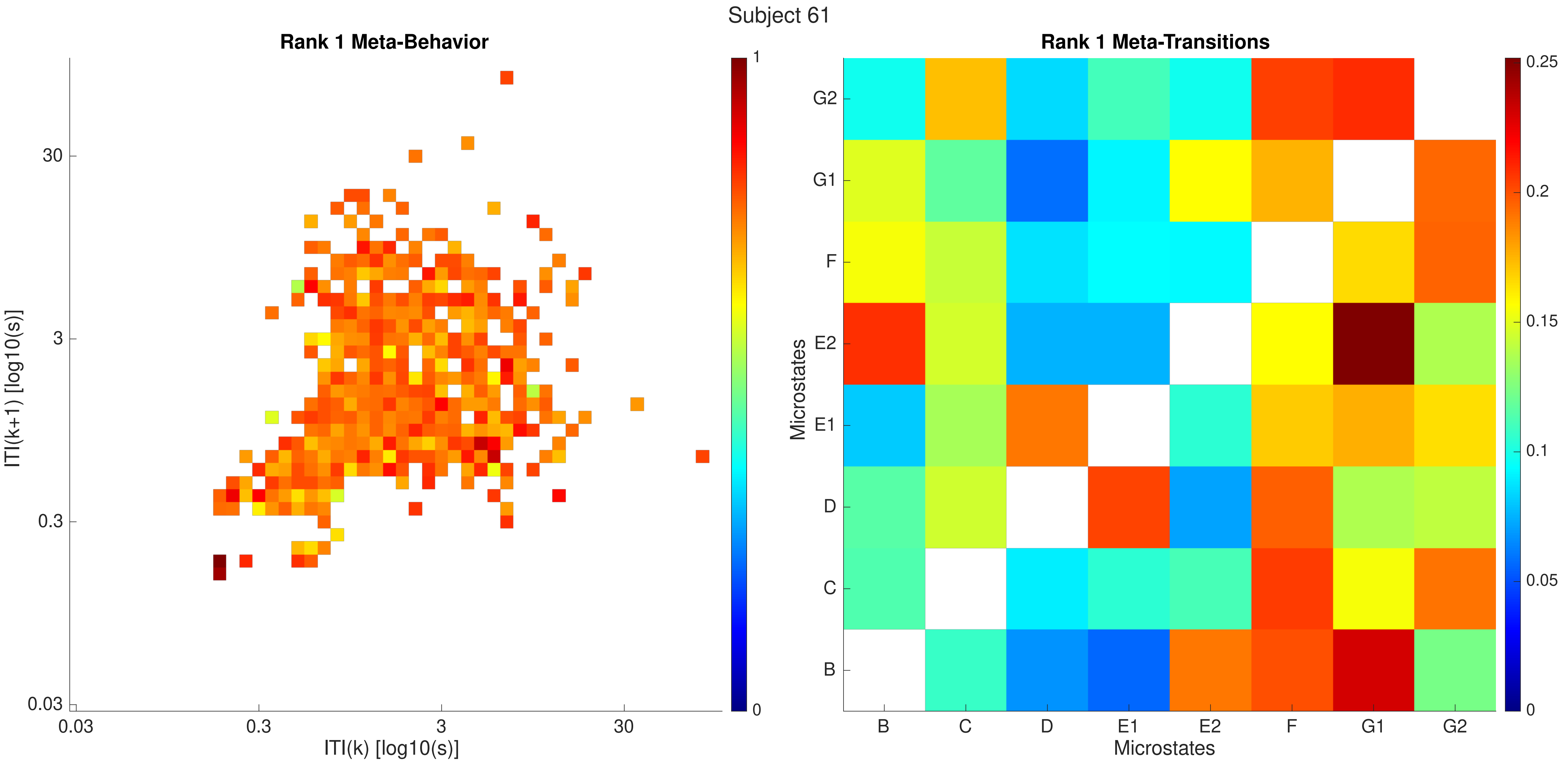

Subject 61
Rank 1 Meta-Behavior
ITI(k+1) [log10(s)]
30
3
0.3
0.03
ITI(k) [log10(s)]
1
0
Rank 1 Meta-Transitions
Microstates
G2
G1
F
E2
E1
D
C
B
0.25
0.2
0.15
0.1
0.05

Subject 61

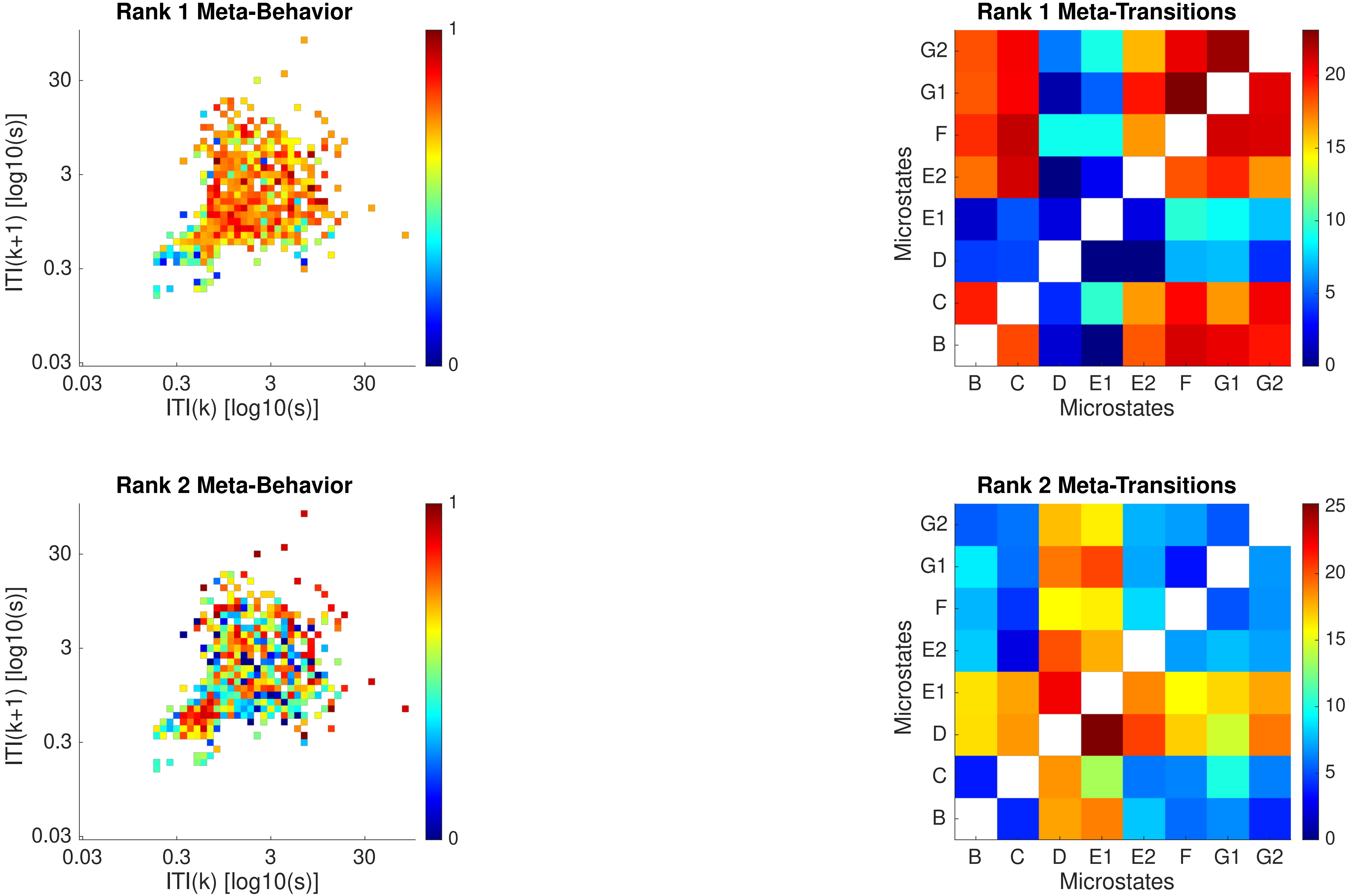

Rank 1 Meta-Behavior
ITI(k+1) [log10(s)]
ITI(k) [log10(s)]
0.03
0.3
3
30
0
1
Rank 1 Meta-Transitions
Microstates
B
C
D
E1
E2
F
G1
G2
0
5
10
15
20
Rank 2 Meta-Behavior
ITI(k+1) [log10(s)]
ITI(k) [log10(s)]
Rank 2 Meta-Transitions
Microstates
25

Subject 61

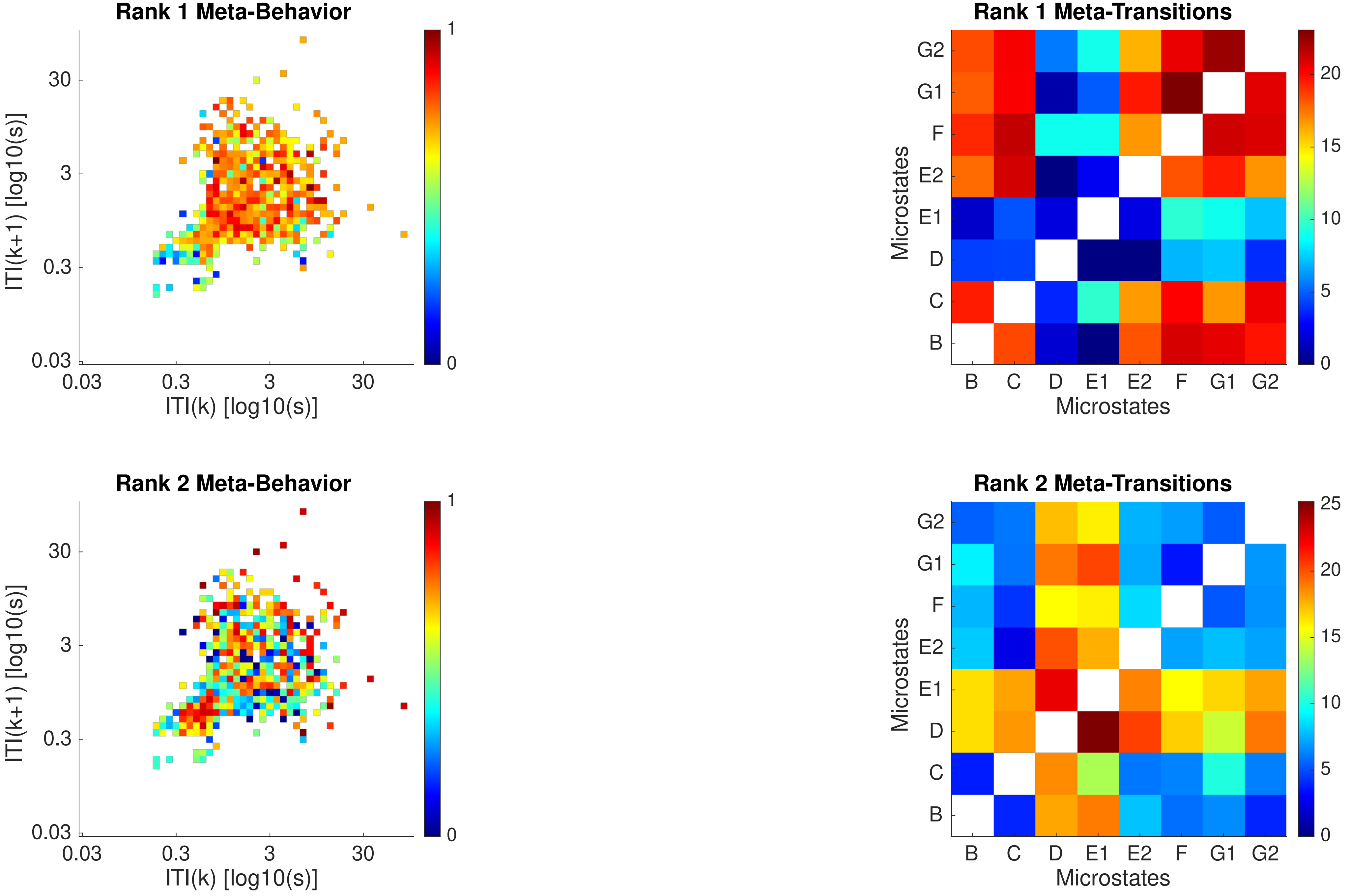
Rank 1 Meta-Behavior
Rank 1 Meta-Transitions
Rank 2 Meta-Behavior
Rank 2 Meta-Transitions
ITI(k+1) [log10(s)]
ITI(k) [log10(s)]
0.03
0.3
3
30
0
1
Microstates
B
C
D
E1
E2
F
G1
G2
0
5
10
15
20
25
