## Supplementary Notes for "Neural microstates in real-world behaviour captured on the smartphone"

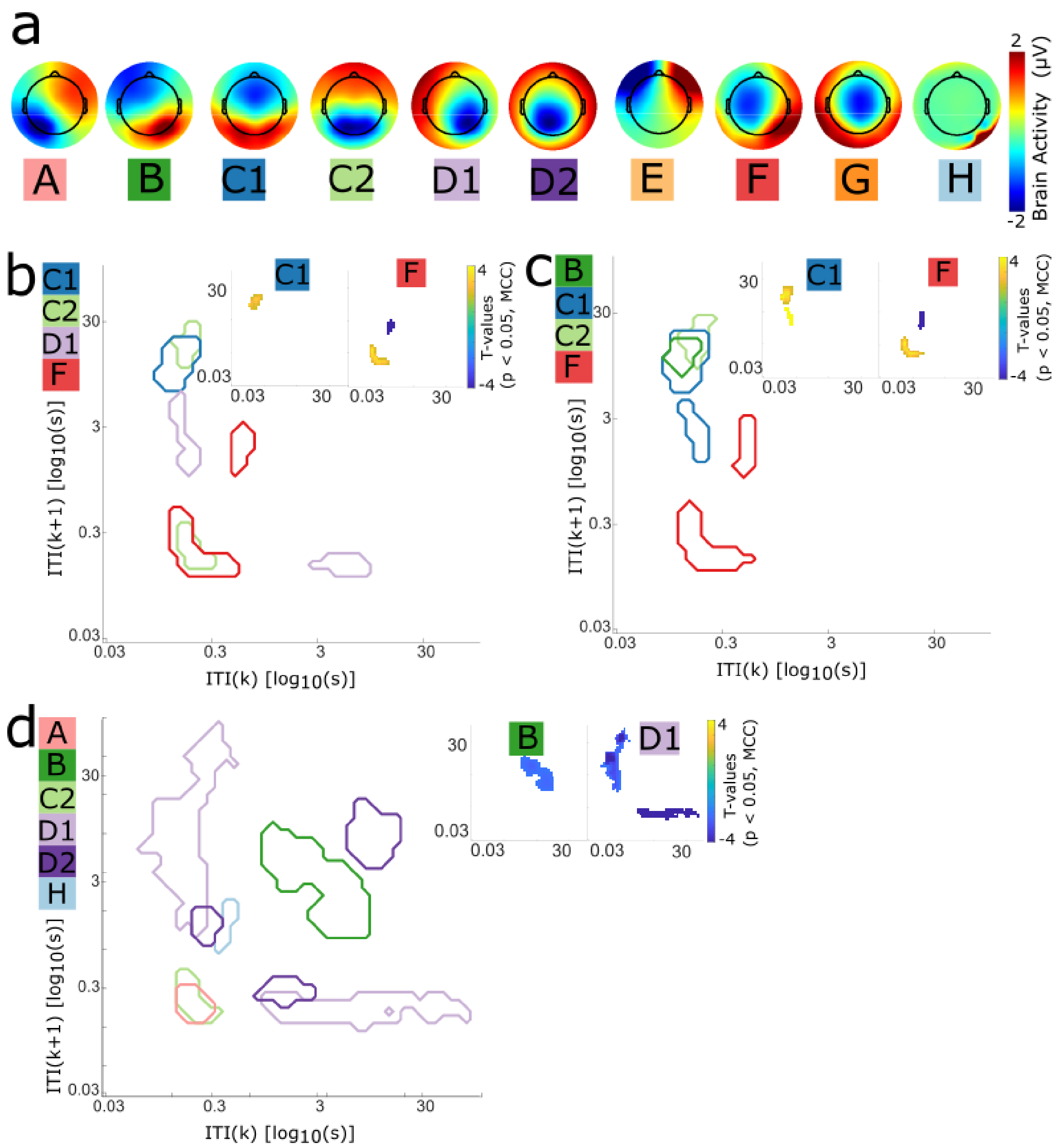


**Supplementary Notes Figure 1.** Microstates identified at rest without restricting the number of microstates to four resulted in 10 microstates. **a)** The microstates captured at rest were similar to microstates during behavior. We followed the same naming convention we used for the behavioral states. Microstate C and D appear to be split in two states. Overall, there may be more than four microstates in eyes closed rest. **b)** Univariate linear regressions reveal a relationship between JIDs and microstate parameters across the population. Contours of the JID show significant relationships to the different microstate coverage parameters. T-values of the coverage parameter and historical smartphone behaviours show significant relationships between microstate C, C', D and F. Microstate C increases in fast followed by slow intervals, similar to our previous findings in four states. Microstate F and C’ increase during faster behaviors. Microstate D decreases during fast to slow and slow to fast behaviors. The same pattern observed in the four states for microstate D is not found here. One possible explanation for these differences is that likely microstate F and G were previously clustered as microstate D and now these are merely separated. The t-values were corrected for multiple comparisons (1000 bootstraps, $\alpha = 0.05$). **c)** Same as b for the occurrence parameter. **d)** Same as b for the duration parameter.

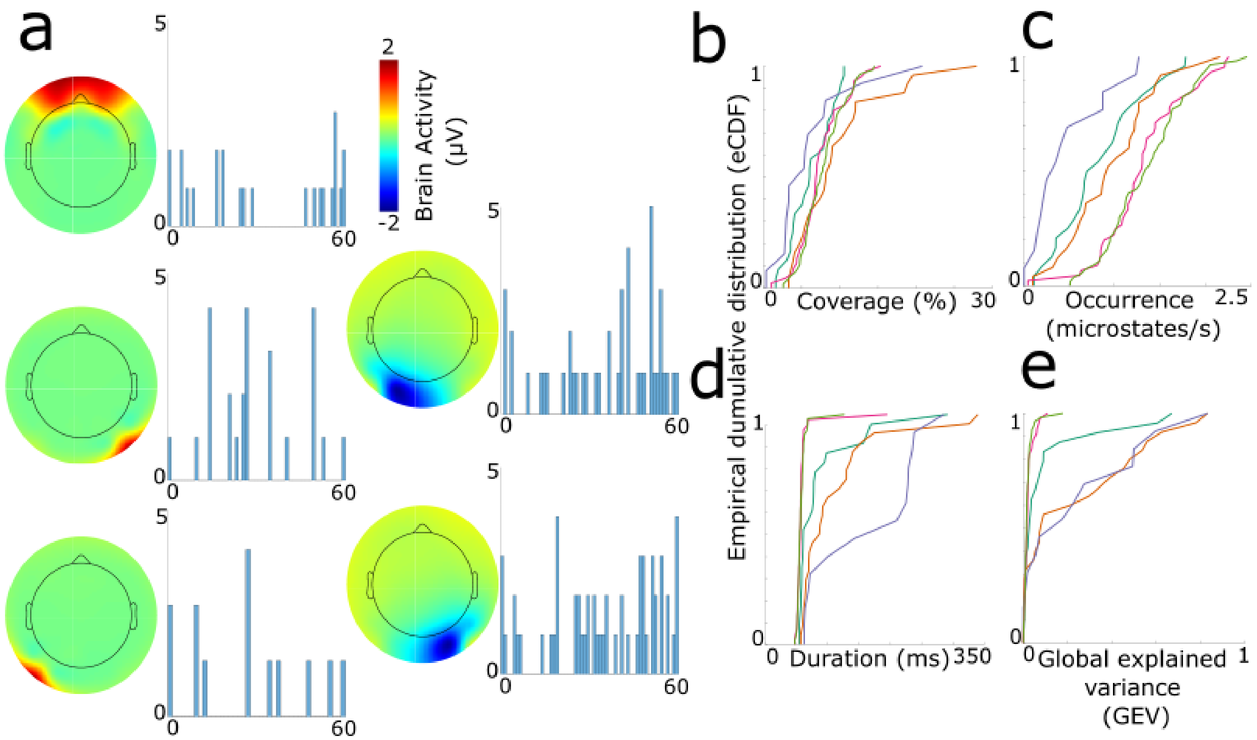


**Supplementary Notes Figure 2.** Microstates during smartphone behavior **(a)** We identified twelve population-level microstates during smartphone behavior five of which shown here displayed large amplitudes over the frontopolar and posterior electrodes. Histograms show the participants clustered in the population microstates. **(b to e)** Empirical cumulative distribution function (ECDF) of the microstate parameters across the population. The ECDF of **(b)** displays coverage, **(c)** median occurrence, **(d)** median duration and **(e)** the global explained variance (GEV). There was substantial variance across the microstate parameters accompanying them. All these five microstates explained ~32% global variance across the population. Despite the higher GEV values, the spatial patterns of these microstates indicate that they may be dominated by muscle-related artifacts. Therefore, the relationship between these five *artefactual* microstates and behaviours was not further considered.

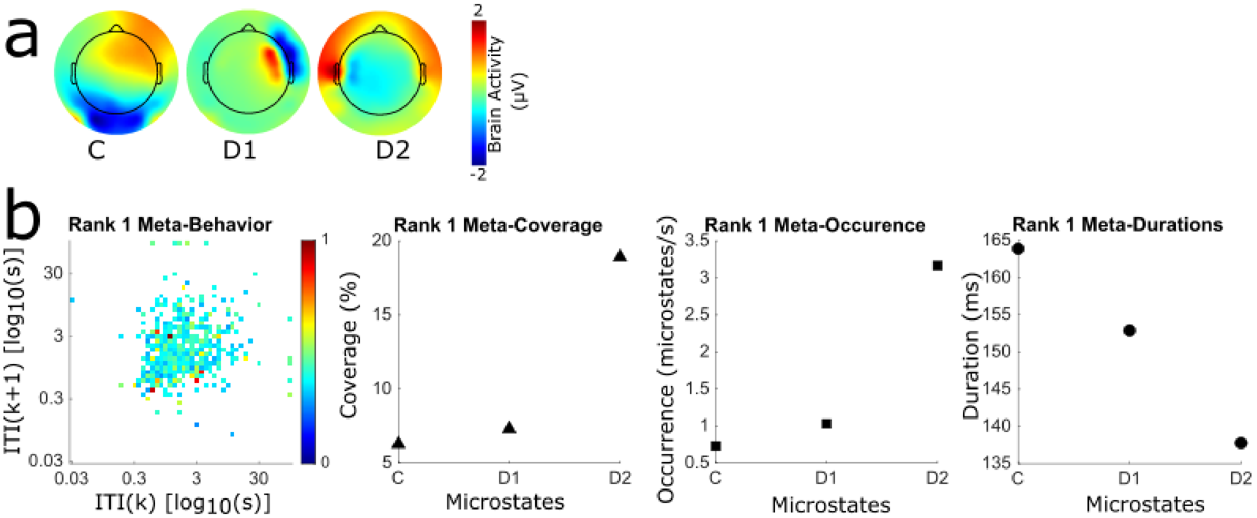


**Supplementary Notes Figure 3.** The NNMF found no consistent configuration of microstate parameters across the behaviors for a subset of the participants (N = 22). **a)** Topographical plots of microstates for one example individual. The microstate maps do not match typical microstates across the population. **b)** NNMF decompositions of microstate parameters (meta-parameters) expressed across smartphone behavior (meta-behavior) for one example subject. The meta-parameters commonly occur in only a few two-dimensional bins. Microstates that appear to be dominated by artifacts have considerable variance in the parameters. NNMF may have picked up the large variance and was unable to find a common pattern. Still,

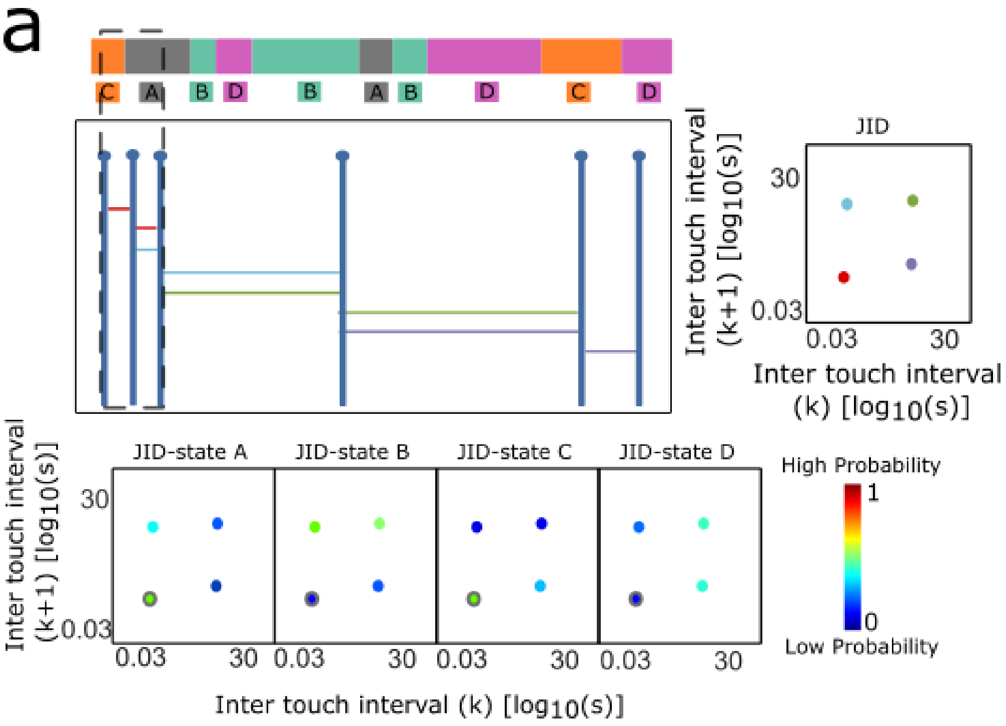


**Supplementary notes Figure 4. Method diagram**. In contrast to the NNMF which captured the configuration of microstate parameters during diverse behaviors, we addressed each microstate parameter separately. We used a block-bootstrapping approach to address whether each microstate parameter exhibited a statistically significantly relationship to the JIDs (1000 bootstraps, $\alpha = 0.05$). First, the method to compute JIDs was adjusted to calculate the microstate parameters across the varying behavioural dynamics. During two inter-touch intervals (K and K+1), multiple microstates were active. The probability of observing a microstate with a specific occurrence and duration across the inter-touch intervals was calculated. For the occurrence, the average number of times a microstate appeared during the inter-touch intervals was divided by the total number of active microstates during the intervals (K and K+1). For the duration, the average duration of a microstate was divided by the total duration of the inter-touch intervals. The coverage was not considered as it is a percentage across the recording while we are considering intervals. The probabilities were aggregated based on the median in each two-dimensional bin (Supplementary Figure X). For each bin, we created 1000 bootstraps by randomizing the inter-touch intervals and then calculating the parameters. This created a random distribution for each microstate parameter. If the original microstate parameter value was larger than the 95th percentile of the distribution, then, it was considered statistically significant different from a random distribution. Then we used spatiotemporal clustering with LIMO EEG for multiple comparison correction (Pernet et al., 2011).

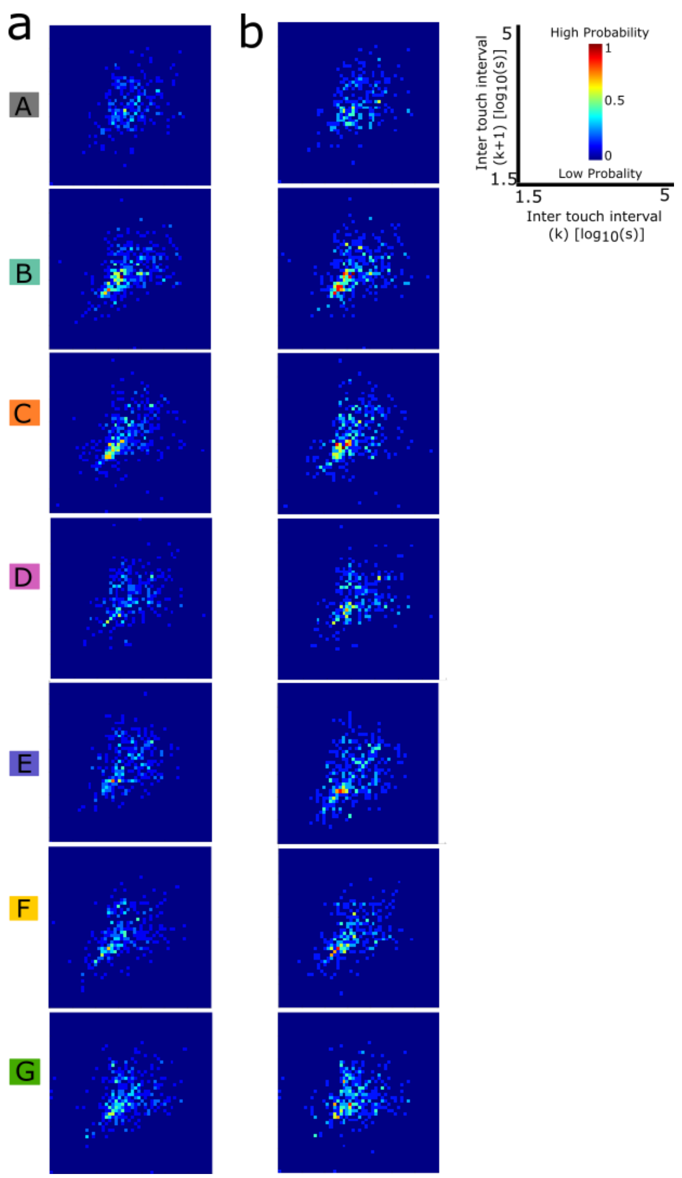


**Supplementary notes Figure 5. Relationship between microstate parameters and smartphone behaviors.** Statistically significant JID bins were aggregated across all the participants to reveal consistent patterns in the population. Within individuals, the number of statistically significant bins were small (less than 10 %) and spread around the whole behavioral space, indicating no clusters of behaviors were significant. This result was observed for the duration, occurrence, and coverage parameters. **(a)** To observe population-wide patterns, we summed the individual results for microstate occurrence, corrected by min-max normalization using the smallest and largest number of significant participants (min =0, max=7). **(b)** Same as a for the microstates with a specific duration. Overall, the largest number of consistently significant behavioral bin occurred in 11% of the participants. The parameters were observed across different types of behaviors, be it fast or slow. There were some marginal differences across the different microstates. These results suggest that separately each microstate parameter does not strongly contribute to differences in smartphone behaviors within or between individuals.

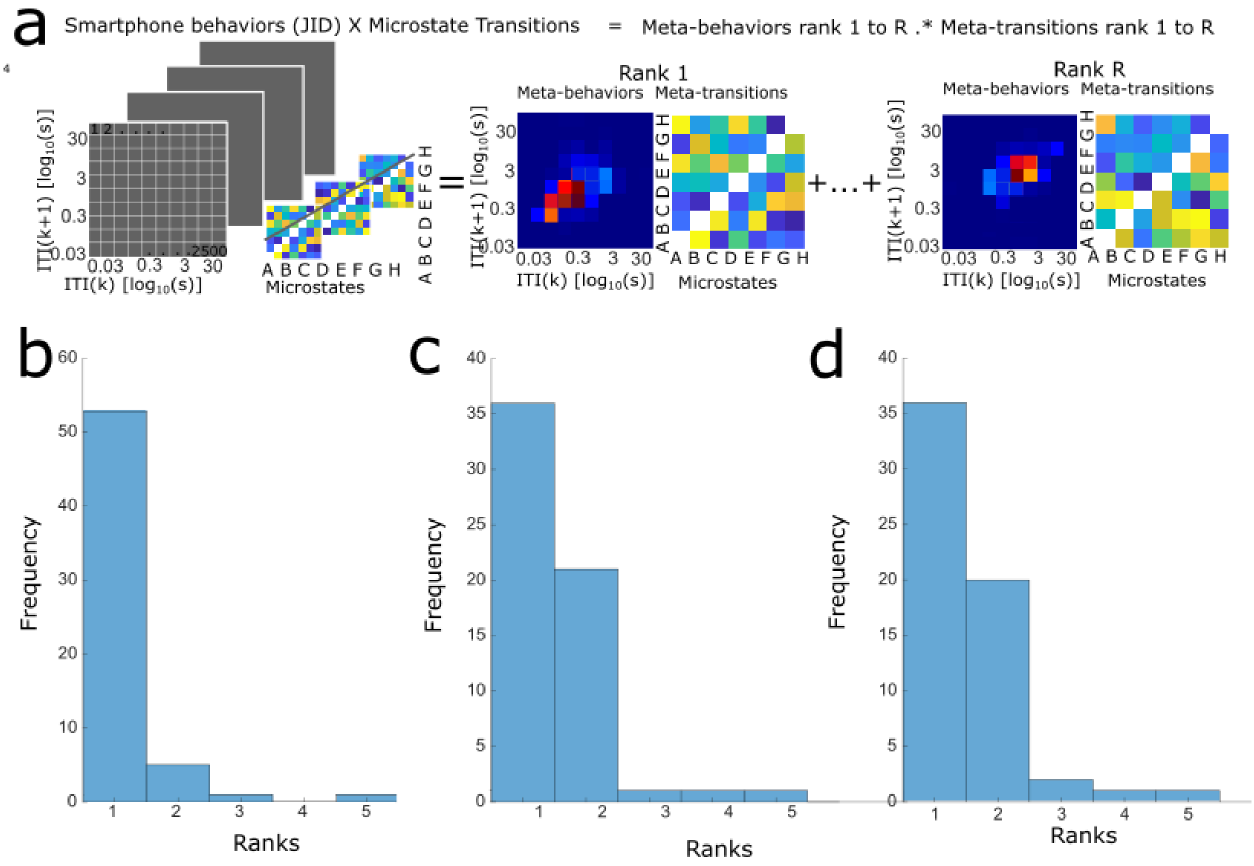


**Supplementary Notes Figure 6.** Transition probabilities and smartphone behaviors. **a)** Sketch showing how non-negative matrix factorization (NNMF) decomposes the microstate transitions during each two-dimensional bin of smartphone behaviours. **(b to d)** Histograms for number of ranks from non-negative matrix decompositions of **b)** transition probabilities, **c)** transition durations before and **d)** transition durations after.

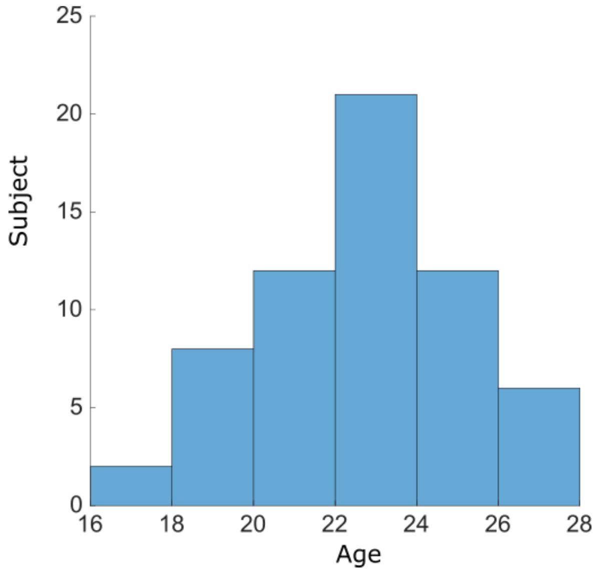


**Supplementary Notes Figure 7.** Age distribution of participants.
